# Patient-derived xenografts and organoids model therapy response in prostate cancer

**DOI:** 10.1101/2020.03.17.994350

**Authors:** Sofia Karkampouna, Federico La Manna, Maria R. De Filippo, Mirjam Kiener, Marta De Menna, Eugenio Zoni, Joël Grosjean, Irena Klima, Andrea Garofoli, Marco Bolis, Jean-Philippe Theurillat, Vera Genitsch, David Keller, Tijmen H. Booij, Christian U. Stirnimann, Kenneth Eng, Andrea Sboner, Charlotte K. Y. Ng, Salvatore Piscuoglio, Gray PC, Martin Spahn, Mark A. Rubin, George N. Thalmann, Marianna Kruithof-de Julio

## Abstract

Therapy resistance and metastatic processes in prostate cancer (PCa) remain undefined, due to lack of experimental models that mimic different disease stages. We describe a novel androgen-dependent PCa patient-derived xenograft (PDX) model from treatment-naïve, soft tissue metastasis (PNPCa). RNA and whole-exome sequencing of the PDX tissue and organoids confirmed transcriptomic and genomic similarity to primary tumor. PNPCa harbours *BRCA2 and CHD1* somatic mutations, shows an *SPOP/FOXA1*-like transcriptomic signature and microsatellite instability, which occurs in 3% of advanced PCa and has never been modelled *in vivo*. Comparison of the treatment-naïve PNPCa with additional metastatic PDXs (BM18, LAPC9), in a medium-throughput organoid screen of FDA-approved compounds, revealed differential drug sensitivities. Multikinase inhibitors (ponatinib, sunitinib, sorafenib) were broadly effective on all PDX- and patient-derived organoids from advanced cases with acquired resistance to standard-of-care compounds. This proof-of-principle study may provide a preclinical tool to screen drug responses to standard-of-care and newly identified, repurposed compounds.

## Introduction

Prostate cancer (PCa) is the second most commonly diagnosed cancer type and the fifth leading cause of cancer death in men worldwide [1]. Androgen deprivation therapy has been used to hamper tumor growth due to the hormone sensitivity of the prostate. However, a subset of tumors will acquire resistance and reoccur as castration-resistant prostate cancer (CRPC). Novel classes of androgen inhibitors, such as enzalutamide [2] and abiraterone [3], are used for CRPC cases, however acquisition of resistance and intratumor heterogeneity limits their efficiency, thus compelling the use of drugs with different mechanisms of action. The lack of available experimental models of early stage, treatment-naive PCa is a major restriction in preclinical PCa research.

Patient-derived xenografts (PDX) are used to address intra-tumor characteristics and drug response since they model the original tumor in a more representative manner than other models such as two-dimensional cell culture [4]. Various PDX study programs have evaluated the take of various PDX models of primary and metastatic PCa [5-7], with the use of different immunocompromised strains, sites of implantations (subrenal [8], subcutaneous [9], orthotopic [5], intrafemoral [10]), grafting of biopsies, cells, circulatory tumor cells [11] and patient-derived organoids [9, 12]. Nonetheless, the few PCa PDXs which are available and suitable for characterizing androgen dependency in PCa are predominantly from metastasis [13] (e.g. BM18 [14] and LAPC9 [15] PDX, androgen-dependent and -independent, respectively). Overall, tumor take is higher from metastatic cases, than primary PCa, grafted in severely immunocompromised mice strains [5, 16] thus underrepresenting early stage cases.

Tumor-derived organoids recapitulate features of naturally occurring tumors such as cellular phenotype, heterogeneity, drug response and overall complexity more efficiently than 2D cell lines, while providing an alternative to animal models [17]. The most widely used methodology is isolation of tumor epithelial cells cultured in Matrigel in presence of a tumor type-specific panel of growth factors [18]. Several studies have demonstrated that drug response in organoids correlates with concomitant genomic profile and may predict clinical outcome [9, 19, 20]. Large scale drug screening on primary PCa organoids have not been performed, with the exception of CRPC neuroendocrine PCa [12], mainly due to the low proliferation rate of PCa organoids and limited availability of material (e.g. needle biopsies). The extent to which PCa PDX and organoids can model key features of therapy resistance and drug response remains unclear.

In this study, we describe the development of a PDX model derived from a treatment-naïve soft tissue metastasis (PNPCa), with androgen sensitive characteristics. Molecular characterization revealed unique genomic features including *CHD1* and *BRCA2* mutations as well as high microsatellite instability (MSI-H). To assess whether therapy resistance preexists in this treatment-naïve PCa case, we developed a method for organoid derivation that facilitates *in vitro* immunological assays and drug screening. Using PDX organoids from three different models, we established a pipeline for medium-throughput organoid drug screen. Lastly, we implemented the outcome of this analysis into a clinically-relevant, near-patient *in vitro* tool.

## Results

### Establishment of clinically relevant models for human PCa

We have generated a novel PDX model from soft tissue metastasis subcutaneously implanted in male, NOD-*scid* IL2Rgammanull (NSG) immunodeficient mice (PN met needle biopsies) (**Fig.1A)**. Histological evaluation of tissue morphology (**Fig.1B, Sup.Fig.1A**), presence of luminal markers PSA, NKX3.1, AR, CK8 expression and absence of CK5+ basal cells (**Fig.1B, Sup.Fig.1B-D)** indicate stable luminal epithelial morphology among the primary TUR-P tumor, the PNmet and PDX1-6 passages. The PDX was established in two different immunocompromised strains, with and without testosterone supplementation (**Sup.Fig.1E**). Flow cytometric analysis showed that cells from PNPCa PDX are positive for prostate-specific membrane antigen (PSMA), E-cadherin (E-Cad) and Integrin α-6 (CD49f), supporting the epithelial and prostatic origin of the tissue. In addition, about 37% of the cells were stem cell marker CD44+, while a minor fraction of cells stained positively for endothelial markers CD36 (6%) and CD146 (1%) (**Fig.1C)**.

**Figure 1.**
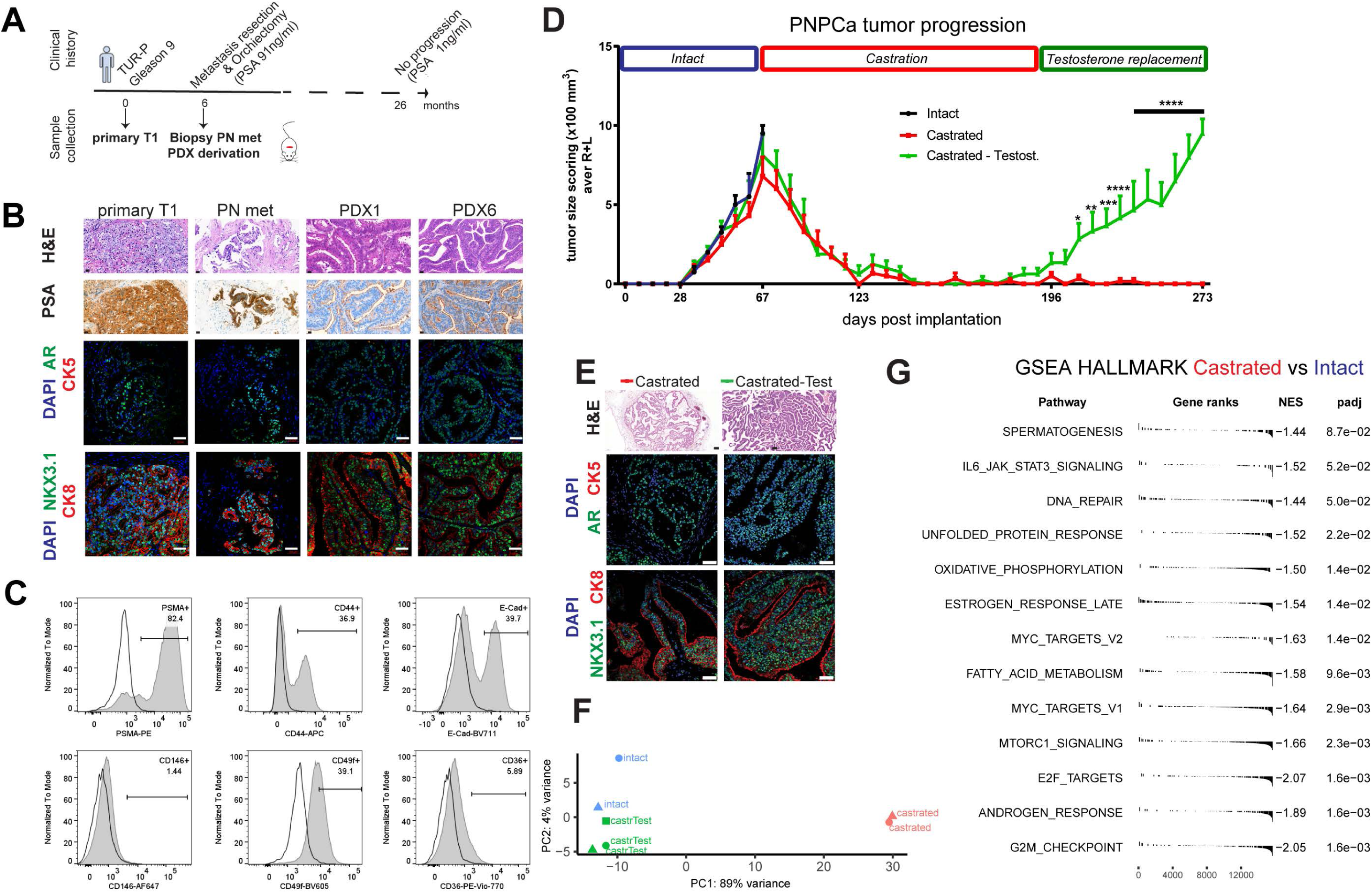
Establishment of a novel androgen dependent, patient derived xenograft from an early, treatment-naïve prostate cancer metastasis. **A**. Scheme of clinical history and respective obtained samples (primary tumor TUR-P (“T1”) and penile metastasis needle biopsies used to establish the PDX model (PNPCa) and subsequent passages. **B**. Histological morphology of primary TURP tumor, penile metastasis (PN met) from PCa and the PDX passages 1 and 6 (PDX1, PDX6) derived from the metastasis needle biopsy implantation (PNPCa), as assessed by Hematoxylin and Eosin staining (H&E). Scale bars 20um. *Top to bottom panels*; PSA protein expression. Scale bars 20um. Expression of AR (green), CK5 (red) assessed by immunofluorescence, DAPI (blue) marks the nuclei. Scale bars 50um. Expression of NKX3.1 (green), CK8 (red) assessed by immunofluorescence. Scale bars 50um. **C**. Flowcytometry analysis of epithelial and prostate-specific marker expression in PNPCa PDX tissue. FcR-blocked PNPCa cells were stained with antibodies against CD44, E-Cadherin, PSMA, CD49f, CD36 and CD146. **D**. PDX tumor growth progression in time. Groups; 1.Intact tumors (collected at max size, N=2), 2.Castrated (N=5), 3.Castrated followed by Testosterone re-administration (Castrated-Testosterone) (N=4, N=3 from day 203-252, N=2 from day 252 to 273). Tumor scoring was performed weekly by routine palpation; values represent average calculation of the tumors of all animals per group (considering 2 tumors, left L and right R of each animal). Error bars represent SEM, is calculated considering No of animals for each time point. Ordinary two-way ANOVA with Tukey`s multiple comparison correction was performed, p < 0.05 (*), p < 0.01 (**). **E**. *Top to bottom panels*; Histological H&E staining of representative tumors from Castrated and Castrated-Testosterone hosts. Immunofluorescence staining for AR and CK5, CK8, NKX3.1 and CK8, CD44 and Ki67. DAPI marks the nuclei. Scale bars 50um. **F**. Principal component analysis plot of the gene expression of the 500 most variable genes on all samples. **G**. Gene set enrichment analysis plot of statistically significant (adjusted p-value < 0.05) enrichment of HALLMARK pathways based on the differential expression analysis of the Castrated versus the Intact group.

Tumor growth properties in response to androgen levels, were addressed *in vivo*. The PDX (passage 6) was implanted subcutaneously in NSG mice (N=11) receiving weekly testosterone injections until tumors reached adequate volume (approx.1000mm3) (**Fig.1D**). PDX tumors were collected as intact control (day 67, N=2) and castration was performed on the remaining animals (N=9). Weekly tumor growth assessment indicated progressive tumor regression (**Fig.1D**), reaching non-palpable tumors by day 151. After ten days of consistently non-palpable tumors (day 151-161) animals were categorized in **1**. castrated group (monitoring whether spontaneous regrowth occurs without re-administration of testosterone (total N=5; N=2 collected as control at day 108, or day 41 post-castration, and N=3 remained in experiment as prolonged castration group)) and **2**.) castrated-testosterone group to assess the extent of tumor regrowth upon androgen re-administration (N=4). All tumors reformed upon testosterone with statistically significant tumor burden (p < 0.05 (*) on day 231, p < 0.0001 (****) from day238 (**Fig.1D;** Castrated (red line), Castrated-Testosterone (green line), **SI Table 1**). At endpoint, tumors were collected for histological and transcriptomic analysis. Histologically, castration induced reduction of epithelial NKX3.1+, CK8+, AR+ glands, (**Fig.1E**; Castrated group), which is reversible upon testosterone re-administration (**Fig.1E**; Castrated-Testosterone groups). Spontaneous tumor regrowth was not observed within the time period checked (206 days post-castration), therefore no androgen-independent regrowth was detected (**Fig.1D**). However, tumor regrowth was observed after testosterone-re-administration in animals maintained under prolonged castration (40 weeks). Analysis of different organs (liver, lung, prostate, lymph node, femur and tibia) at end point collection, indicated macroscopic foci on all lung tissues (**Sup.Fig.2A)**, human panCK cell infiltration of lymph node in 1 case (**Sup.Fig.2B)**, loss of normal epithelial architecture in anterior prostate (**Sup.Fig.2B)** and potential micrometastases areas of scattered panCK+ (human specific) cells residing in the bone (**Sup.Fig.2C)**, although no apparent lesion was detectable by X-ray (**Sup.Fig.2D)**. Principal component analysis (PCA) of RNASeq (**Fig.1F**) and hierarchical clustering (**Sup.Fig.3**) indicated high transcriptomic correlation among tumors from intact hosts and those that received testosterone after surgical castration, while tumors from castrated hosts further diverged from the aforementioned groups (**Fig.1F**). Differential expression analysis revealed lower expression of genes in metabolic pathways, mTOR, MYC and AR response pathways in the Castrated group compared to the Intact tumor groups (**Fig.1G**), which is indicative of the biological processes taking place after androgen deprivation treatment. Androgen ablation by castration causes tumor regression whereas testosterone re-administration induces tumor regrowth, rendering the PNPCa an androgen-dependent model from a treatment naïve case.

### Molecular analysis revealed genomic and transcriptomic stability among the PCa xenograft and organoid models

To further explore the *in vitro* characteristics of the PDX tumor cells, we developed an organoid culture method from bulk tumor tissue that allows organoids to grow in suspension conditions, with no requirement for extracellular matrix support (e.g. Matrigel). PNPCa-PDX tissue-derived organoids exhibit a luminal phenotype evident by PSA, AR and CK8 expression (**Fig.2A, Sup.Fig.4)**. Previously established advanced PCa PDX models (BM18 [14], LAPC9 [15]) were used for comparison with the PNPCa. While both PDXs were derived from bone metastasis, the LAPC9 PDX represents advanced, androgen-independent PCa, whereas the BM18 PDX retains androgen sensitivity **(Sup.Fig.5A-B)**. Organoids derived from both PDXs display morphological features of budding acinar and adenocarcinoma-like organoids and express CK8, PSA and AR **(Fig.2A)**. Tumorigenic potential of BM18 and LAPC9 organoids was confirmed by performing subrenal grafting **(Sup.Fig.5C-H)** or intraprostatic inoculation **(Sup.Fig.5-N)**. The growth kinetic of organoids shows that their viability is enhanced when they are grown in organoid media containing androgens (dihydrotestosterone, DHT) (**Fig.2B**).

**Figure 2.**
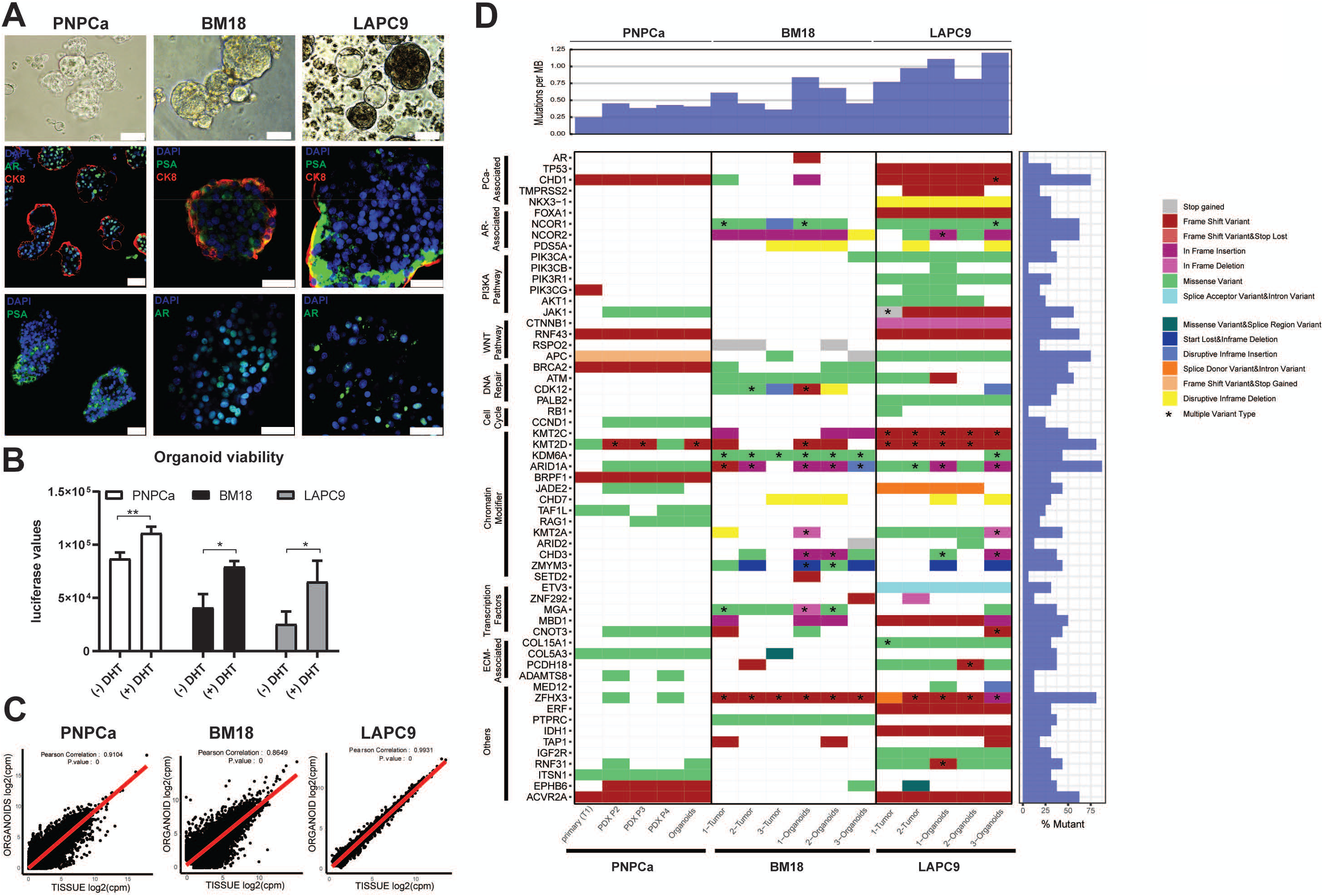
Mutational landscape of PDX and PDX-derived organoids from PNPCa, and advanced androgen (in) dependent BM18 and LAPC9 models. **A**. Morphology of PNPCa, BM18 and LAPC9 PDX-derived organoids; brightfield images, whole mount immunofluorescence staining and 3D projection of z-stack of organoids stained for PSA, AR, and CK8. DAPI marks the nuclei. Scale bars 50um. **B**. Viability assay of organoids derived from PNPCa, BM18 and LAPC9 tumor tissues and exposed to dihydrotestosterone (+/- DHT) for 48 hours. Luciferase values (ATP release, CellTiter Glo 3D) are proportional to cell viability. Two-tailed t-test, *p<0.05, **p<0.01. **C**. Correlation plots of gene expression between PNPCa PDX tissue (N=3) and organoids (N=2) (Pearson correlation coefficient r=0.91), BM18 PDX tissue (N=2) and organoids (N=2), (Pearson correlation coefficient r=0.86), LAPC9 PDX tissue (N=2) and organoids (N=2) (Pearson correlation coefficient r=0.99). **D**. Somatic mutation analysis of WES sequencing of tissue and organoids of PNPCa, BM18 and LAPC9 PDX. Columns represent different samples, while rows represent genes categorised per pathway. Types of genetic aberration detected are indicated in different colors. Multiple types of mutations per gene are indicated with an asterisk. Mutation frequency per MB is indicated on the top histogram. Frequency of mutation in the tested samples is shown in the right histogram.

Next we determined whether the established PDX and PDX-derived organoids comprehensively recapitulate the originating tumor in terms of transcriptomic and genomic profile. RNA sequencing was performed on three distinct PDX passages of the PNPCa (P2, P3 and P4) and two PDX-derived organoid samples (Org1, Org2) (**Sup.Fig.4**), as well as BM18 and LAPC9 (tumor tissue and organoids). Gene expression levels (**Fig.2C**) showed high correlation, among the PDX tissues and the PDX-derived organoids, for all tested models, supporting the use of our established organoid culture system to preserve the transcriptional profile. We assessed the genomic landscape of the PNPCa by subjecting the originating primary tumor TUR-P (T1), its matched germline control (N1), 3 PDX passages (P2, P3, P4) and the PDX-derived organoids (Org2) to whole exome sequencing (WES) (**Sup.Fig.4, Fig.2D**). Copy number analysis showed that the derivative organoid models retained overall patterns of copy number alterations (CNAs) (**Sup.Fig.6A**).

Of the total somatic non-synonymous mutations (**Sup.Fig.7A-B, SI Table 2**) found in the primary tumors, 54%, 53%, 51% and 53% were observed in the corresponding organoids and PDX models at passage 2, 3 and 4 respectively (**Sup.Fig.6B**). Nearly all non-synonymous somatic mutations in *bona fide* cancer genes were preserved in the PNPCa models, including truncating (loss of function) mutations in *CHD1* (p.Asn310fs), *ACVR2A* (p.Lys437fs), *RNF43* (p.Gly659fs), *APC* (p.Asp802fs) and *BRCA2* (p.Asn863fs) (**Fig.2D, Sup.Fig.7A**). These mutations show stable or increasing cancer cell fraction (CCF) in the PDXs compared to the primary T1, potentially due to clonal selection. Only two frameshift mutations in cancer genes were lost in the PDX tumors; *SPEN* and *PIK3CG* (**Fig.2D**). No mutations in the *AR* gene were identified; mutations in other AR pathway gene members are the cancer related *UBE3A* gene, while mutations in *ARID1A, NCOA1, KDM3A* were observed only in the primary tumor (**Sup.Fig.7B**). Mutations with homogeneously high prevalence on all samples (≥80% CCF) are found in the gene loci of *CHD1, APC, RNF43* and *KMT2D* (**Sup.Fig.8**). *BRCA2* and *ACVR2A* mutations were found in 60-90% CCF in P2, P3, P4 and Org2 and in ≤20% CCF of T1 (**Sup.Fig.8**).

We compared gene expression of PNPCa to those of genetically defined subgroups obtained from TCGA. Principal components analysis (PCA) plot depicts the variance among gene signatures of PCa cases with CHD1 homozygous deletion (**Sup.Fig.9A**) and from cases with mutant *FOXA1, SPOP, CHD1*, ETS rearrangements (ERG, ETV1, ETV4) (**Sup.Fig.9B**). We evaluated the activity of gene-signatures that are characteristic of specific PCa subtypes and quantified the activity of gene-sets using a single-sample gene-set enrichment analysis approach (ssGSEA). Signatures of ETS and SPOP/FOXA1 subgroups are divergent with respect to each other, while the PNPCa tumors and organoids, clustered between the two categories (**Sup.Fig.9C**) and closely to CHD1 homozygous-deletion group (**Sup.Fig.9A**), in agreement with presence of a truncating mutation in *CHD1* (**Fig.2D**).

WES was additionally performed on the LAPC9 and BM18 PDX tumors and PDX-derived organoids. Mutational load was increased in the LAPC9 compared to the BM18 and PNPCa (**Fig.2D**, upper plot), in line with the its aggressive, androgen-independent phenotype *in vivo*. At the genomic level, frameshift mutations of high impact were detected in the BM18 were *ATM, KDM6A* and *ZHFX3* mutations (**Fig.2D**). LAPC9 PDX had a *NKX3*.*1* deletion, along with frameshift mutations in *TP53, CHD1, FOXA1, ERG* and *PI3K* genes and in genes of the Wnt pathway (*APC, CTNNB1, RNF43*) (**Fig.2D**). In both models, the mutational profile was conserved between the PDX tumor and the PDX-derived organoids, similarly to PNPCa (**Fig.2D**).

### Functional testing of targeted treatments according to the genomic profile of the PNPCa PDX and organoids

We next characterized the functional implications of specific mutational alterations of the PNPCa tumor. PNPCa PDX harbors a somatic truncating mutation in the *BRCA2* (p.Asn863fs) gene. Patients with germline *BRCA2* defects have earlier disease onset, a higher rate of 5-year metastatic progression and poor survival compared to non-carriers [21-23]. Tumors with *BRCA1/2* defects and defective DNA repair mechanism are particularly sensitive to DNA damaging agents such as radiotherapy [24] and to PARP inhibitors such as olaparib, which is used for prostate, breast and ovarian cancer [25]. To functionally assess the therapy response of the PNPCa model, we have focused on its actionable genomic characteristics and available treatment options related to this specific genomic profile. *In vitro* viability assays of irradiated organoids confirmed the sensitivity of the *BRCA2*-mutant PNPCa PDX-derived cells to irradiation, compared to organoids derived from other PDX models (BM18, LAPC9) (**Fig.3A-C**) confirming the sensitivity of this tumor to irradiation in this experimental setup.

**Figure 3.**
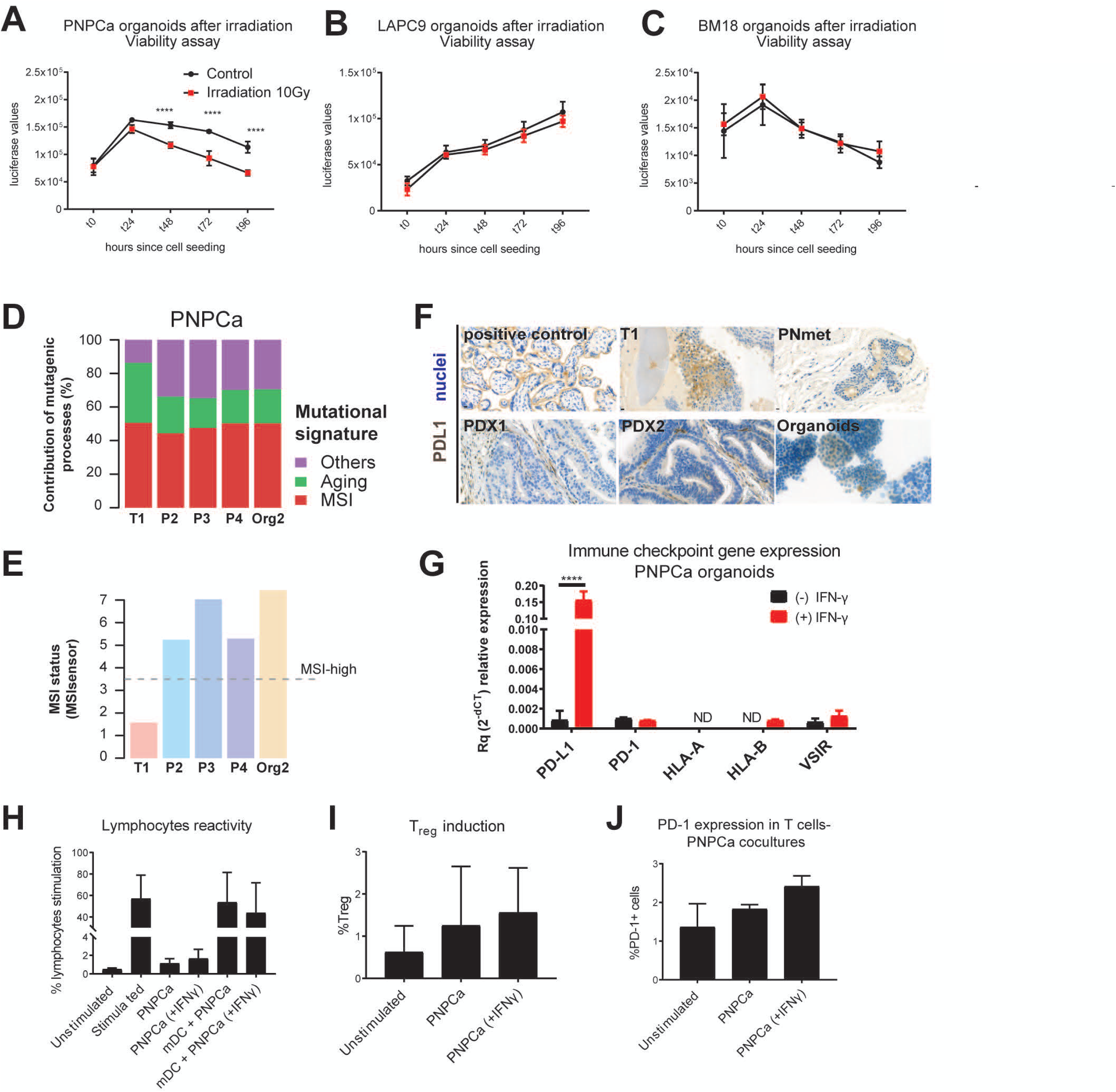
Correlation of genomic features and specific drug responses in organoid models. **A-C**. Time course of ATP-mediated luminescence viability assay following a single dose of 10Gy irradiation on organoids derived from PNPCa (**A**), LAPC9 (**B**) and BM18 (**C**) PDX tumors. **D**. Graph representing the percentage of contribution of specific mutagenic processes based on mutational signatures from PNPCa T1 (primary tumor), PDX passages (P2, P3, P4) and organoids (from P4 PDX). **E**. MSI status based on MSIsensor algorithm (https://github.com/ding-lab/msisensor), score ≥3.5 indicates as MSI-high. **F**. PD-L1 IHC staining on positive control (placenta tissue), primary T1 tumor, PNmet needle biopsy, PDX1 and PDX2 of the PNmet and cytosmear of PDX-organoids. **G**. Gene expression levels of immune markers based on RT-qPCR results on PNPCa organoids RNA at baseline (black bars) and after 48 hours exposure to IFN-γ (red bars). **H-J**. MLR assay showing lymphocyte reactivity, T_reg_ fraction and expression levels of surface PD-1, following coculture of PDX-derived PNPCa organoids with T cells and allogeneic, monocyte-derived dendritic cells (DCs).

The analysis of oncogenic signatures of the PNPCa demonstrated that the mutational landscape of the primary tumor, the PDX and organoid models was largely driven by mutational processes associated with signatures of microsatellite instability (MSI) (**Fig.3D**). The MSI phenotype is consistent with the observations that this tumor had a high mutation rate compared to other PCa [26], especially those with an overall flat copy number profile (**Sup.Fig.6A**) and an elevated proportion of small insertions and deletions [27] (**Sup.Fig.10A**). MSI status was further evaluated using the MSIsensor algorithm [28]. MSIsensor classified all the samples except the T1 tumor as MSI-H (**Fig.3E**, grey dotted line). To further confirm the MSI status of this latter sample we analyzed it with the Bethesda MSI test. Four out of the six loci of the Bethesda panel, were altered in the T1 tumor, confirming the tumor itself as MSI-H (**Sup.Fig.10B**).

Considering that MSI score is diagnostically used as a biomarker for immunotherapy response, we assessed the expression level and the functional activity of the PD-L1 antigen. While the primary tumor, the soft tissue metastasis and the PDX-derived organoids showed a moderate, epithelial-specific staining for the PD-L1 antigen, the PDX1 and PDX2 tissues revealed a loss of expression of this marker in the epithelial compartment (**Fig.3F**). Compared to the normal tissue N1, all the PDXs, Org2 and the primary tumor showed a reduction of transcript abundance of the major histocompatibility complex (HLA-A and HLA-B) and of galectin-9, the main ligand of the inhibitory receptor Tim-3. The increased level of expression of PD-L1 in the primary tumor compared to healthy tissue N1 is a feature that is not preserved in the PDX and in the organoid samples (**Sup.Fig.10C-D**). As both the *in vitro* systems adopted and the animal models used to maintain the PDXs lack the selective pressure of the immune system, we assessed the functional expression of these markers *in vitro* on PNPCa PDX-derived organoids. PNPCa organoids were cultured *in vitro* with 50 ng/ml IFN-γ for 48h before RNA extraction and immunological assays. Treatment with IFN-γ was sufficient to significantly (p<0.001) evoke the upregulation of PD-L1 and galectin-9, linked to immune evasion and of nectin-2, involved in NK-mediated binding of the cancer cells (**Fig.3G, Sup.10E**) [29, 30]. PNPCa PDX-derived organoids were then cocultured with CD3+ lymphocytes and allogeneic mature dendritic cells (mDC) in the setting of a mixed lymphocyte reaction (MLR). Despite the upregulation of checkpoint inhibitory antigens at the molecular level, PNPCa organoids did not modulate the proliferation of CD3+ lymphocytes, even after 48h pre-treatment with IFN-γ (**Fig.3H**). At the end of the coculture, lymphocytes were stained for the expression of the key regulatory T-cell markers CD4, CD25 and FoxP3 as well as for the expression of PD-1 to assess their tolerogenic status. Coculture with PNPCa organoids did not significantly increase the number of regulatory T cells nor the amount of PD-1 that was expressed on lymphocytes (**Fig.3I-J**). Organoid cultures may indeed provide a model for implementing *in vitro* immunologic responses and the efficacy of targeted therapies.

### Organoid drug response to standard-of-care and repurposing of FDA approved compounds on a medium-throughput automated screen

Progression of androgen-ablated patients to CRPC associated with acquisition of drug resistance is a frequent consequence of PCa. Functional assays are required to identify novel, effective drugs for these patients, particularly with the aim of adopting a precision medicine approach. In order to exploit the treatment naïve status of the PNPCa tumor, we set up a medium-throughput screening method using the robotic screening equipment (NEXUS Personalized Health Technologies, an ETH technology platform). We then tested the drug sensitivity profile of PNPCa-derived organoids to multiple PCa standard-of-care drugs as well as to different FDA-approved drugs with indications for other cancer types (drug repurposing). Compounds were assayed at the concentrations of 10, 1 and 0.1 uM (**SI Table 3**) and additional concentrations were used for the PCa standard-of-care compounds. Overall, the tested compounds targeted several distinct cellular processes and pathways, with a specific focus on signaling mediators including growth factor receptors and androgen response.

Cell density, positive controls and time of drug exposure were optimized for the PNPCa PDX-derived organoids **(Sup.Fig.11A-C)**. Once automatically seeded, the PNPCa PDX-derived cells were allowed to form organoids for 48h before adding the drugs and cell viability was assessed on-plate after 72h from initial drugs exposure (**Fig.4A**). The assay replicates (plates A to D) highly correlated within treatment groups as evidenced by the correlation plots (**Sup.Fig.11D**). PNPCa organoids showed resistance to the majority of tested drugs, however 14 drug compounds significantly reduced their viability (**Fig.4B**, (FDR<0.05)).

**Figure 4.**
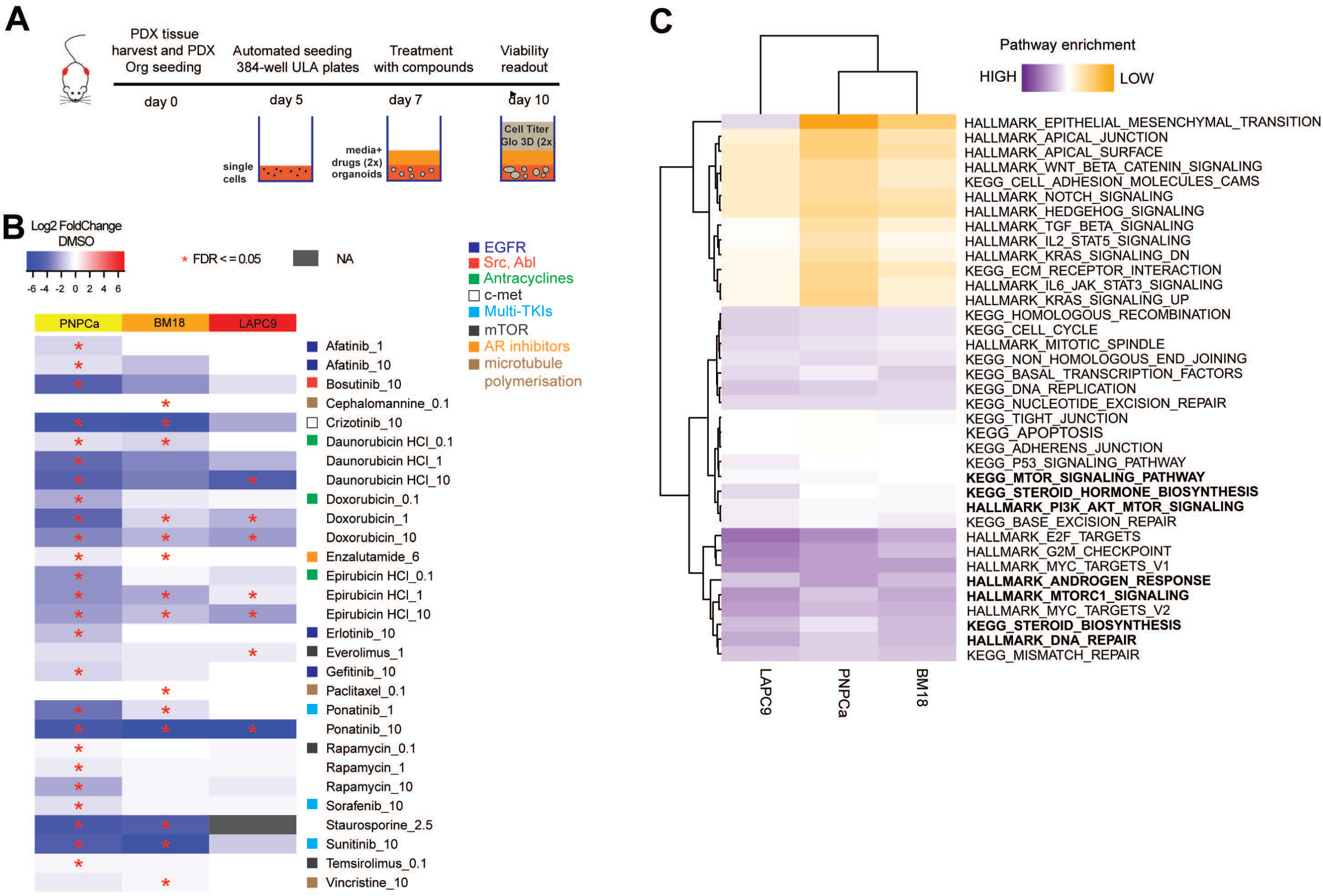
Drug sensitivity of organoids representing different stages and identification of novel compounds for repurposing use, based on medium-throughput organoid screens. **A**. Scheme of experimental protocol for organoid drug screens. **B**. Organoid drug screen heatmap of log2 fold change viability values (over DMSO vehicle control of each PDX model) for PNPCa (N=4), BM18 (N=3), LAPC9 (N=3). Statistically significant hits are indicated with an asterisk, FDR<=0.05. Staurosporine was used as positive control. ATP-mediated luciferase measurements, proportional to cell viability, were obtained after 48-72h treatment of organoids with the compounds, listed in alphabetical order with dose indication on the right, in uM. Negative log2 values (plotted in blue) indicate potential drug candidates with impact on cell viability. Medium-throughput automated drug screens, using selected FDA approved compounds, were performed at Nexus Theragnostics platform. **C**. ssGSEA of PDXs (PNPCa N=3, BM18 N=2, LAPC9 N=2) representing the enrichment score of selected pathways from Hallmarks and C2 Kegg (MSigDB) relative to non-carcinoma control tissue from PNPCa clinical sample (N1).

The most effective compounds were the AR inhibitor enzalutamide, EGFR/HER2 inhibitors (afatinib, erlotinib, gefitinib), mTOR inhibitors (rapamycin, temsirolimus), DNA synthesis inhibitors antracycline class (doxorubicin, daunorubicin, epirubicin), multi tyrosine kinase inhibitors (TKIs; sorafenib, ponatinib, sunitinib), the c-Met pathway inhibitor crizotinib and Scr-Abl inhibitors (bosutinib, ponatinib) (**Fig.4B, Table 1**). In order to generate a more comprehensive drug sensitivity profile, we applied the automated drug pipeline to organoids derived from two other advanced PCa PDXs, LAPC9 and BM18. The drug sensitivity profiles identified drugs that were exclusively effective in only one of the models as well as broadly effective drugs in all 3 PDXs (**Fig.4B, Sup.Fig.12)**. BM18 organoids were sensitive to drugs affecting DNA replication (doxorubicin), growth factor receptors and mTOR signaling (ponatinib, sunitinib and rapamycin, respectively), and were especially sensitive to the microtubule polymerization inhibitors (vincristine, cephalomannine, paclitaxel). LAPC9 organoids were mainly affected by signal transduction inhibitors and cell cycle/DNA replication inhibitors. In particular, LAPC9 organoids responded to only five compounds; antracyclines (same as BM18, PNPCa), TKI ponatinib and the mTORC1-targeting drug everolimus (**Fig.4B, Table 1**). Everolimus was the only compound to be effective exclusively on LAPC9 (**Fig.4B**). In order to assess drug sensitivity in a different functional assay, we performed the same drug treatments in *ex vivo* tissue slices from PNPCa PDX. We tested 13 compounds on PNPCa tissue slices including 12 of the effective compounds and docetaxel, which did not significantly affect organoid viability in the PNPCa drug screen. Of these, we were able to validate 11 out of the 13 tested compounds (**Sup.Fig.13A-B**). As performed for the PNPCa PDX, we tested a selection of drugs, both effective and ineffective in the organoid screening, on *ex vivo* tissue slices of LAPC9 and BM18 PDX. We validated the effectiveness of 7 out of 11 tested compounds on LAPC9 PDX and of 4 out of 5 tested compounds on BM18 PDX (**Sup.Fig.13C**).

**Table 1.** related to Fig.4. List of statistically significant, effective compounds, identified on PNPCa, BM18 and LAPC9 organoid drug screen, based on FDR value <0.05. Dose used (uM) and mechanism of action are indicated.

| List of effective compounds |  |  |  |  | TABLE 1 |
| --- | --- | --- | --- | --- | --- |
| Class | Compound | PNPCa | BM18 | LAPC9 | Target |
| AR | Enzalutamide | 6uM | 6uM | - | AR agonist |
| RTK inhibitors | Afatinib | 1uM<br>10uM | - | - | EGFR,HER2 |
|  | Bosutinib | 10uM | - | - | Scr, ABL |
|  | Crizotinib | 10uM | 10uM | - | c-Met, ALK, ROS1 |
|  | Erlotinib | 10uM | - | - | EGFR |
|  | Gefitinib | 10uM | - | - | EGFR |
|  | Ponatinib | 1uM<br>10uM | 1uM<br>10uM | 10uM | Abl,PDGFRA, VEGFR2,<br>FGFR1, Src |
|  | Sorafenib | 10uM | - | - | VEGFR, PDGFR, Raf |
|  | Sunitinib | 10uM | 10uM | - | VEGFR, PDGFR, c-Kit, Flt |
| Anthracyclines | Daunorubicin | 0.1uM<br>1uM<br>10uM | 0.1uM | 10uM | Telomerase |
|  | Doxorubicin | 0.1uM<br>1uM<br>10uM | 1uM<br>10uM | 1uM<br>10uM | Topoisomerase II |
|  | Epirubicin | 0.1uM<br>1uM<br>10uM | 1uM<br>10uM | 1uM<br>10uM | Topoisomerase II |
| mTOR inhibitors | Everolimus | - | - | 1uM | Selective mTORC1 |
|  | Temsirolimus | 0.1uM | - | - | FKBP12/mTORC1 |
|  | Rapamycin | 0.1uM<br>1uM<br>10uM | - | - | FKBP12/mTORC1 |
| Tubulin binding | Cephalomannine | - | 0.1uM | - | Microtubule stabilisation |
|  | Paclitaxel | - | 0.1uM | - | Microtubule stabilisation |
|  | Vincristine | - | 10uM | - | Microtubule stabilisation |

| List of effective compounds |  |  |  |  |  |
| --- | --- | --- | --- | --- | --- |
| Class | Compound | PNPCa | BM18 | LAPC9 | Target |
| AR | Enzalutamide | 6uM | 6uM | - | AR agonist |
| RTK inhibitors | Afatinib | 1uM<br>10uM | - | - | EGFR,HER2 |
|  | Bosutinib | 10uM | - | - | Scr, ABL |
|  | Crizotinib | 10uM | 10uM | - | c-Met, ALK, ROS1 |
|  | Erlotinib | 10uM | - | - | EGFR |
|  | Gefitinib | 10uM | - | - | EGFR |
|  | Ponatinib | 1uM<br>10uM | 1uM<br>10uM | 10uM | Abl,PDGFRa, VEGFR2, FGFR1, Src |
|  | Sorafenib | 10uM | - | - | VEGFR, PDGFR, Raf |
|  | Sunitinib | 10uM | 10uM | - | VEGFR, PDGFR, c-Kit, Flt |
| Anthracyclines | Daunorubicin | 0.1uM<br>1uM<br>10uM | 0.1uM | 10uM | Telomerase |
|  | Doxorubicin | 0.1uM<br>1uM<br>10uM | 1uM<br>10uM | 1uM<br>10uM | Topoisomerase II |
|  | Epirubicin | 0.1uM<br>1uM<br>10uM | 1uM<br>10uM | 1uM<br>10uM | Topoisomerase II |
| mTOR inhibitors | Everolimus | - | - | 1uM | Selective mTORC1 |
|  | Temsirolimus | 0.1uM | - | - | FKBP12/mTORC1 |
|  | Rapamycin | 0.1uM<br>1uM<br>10uM | - | - | FKBP12/mTORC1 |
| Tubulin binding | Cephalomannine | - | 0.1uM | - | Microtubule stabilisation |
|  | Paclitaxel | - | 0.1uM | - | Microtubule stabilisation |
|  | Vincristine | - | 10uM | - | Microtubule stabilisation |

To identify potential correlations between drug sensitivity and activity of biological processes, pathway enrichment analysis was performed. Based on pathway enrichment score (Hallmarks and C2 KEGG), ssGSEA of the 3 considered PDXs (PNPCa, BM18, LAPC9) revealed that PNPCa clustered with BM18, in line with the androgen dependency of these two models. Compared to the other PDX models, PNPCa organoids were selectively sensitive to EGFR inhibitors (gefitinib, erlotinib, afatinib), in concordance with the low enrichment score of KRAS signaling pathway (**Fig.4B-C**), as well as to other TKI inhibitors (bosutinib, crizotinig, sorafenib, sunitinib) and to the mTOR inhibitor temsirolimus. The androgen-independent LAPC9 model showed high enrichment of EMT, KRAS, JAK/STAT, WNT and NOTCH pathways, and was distinct from both BM18 and PNPCa models, which showed reduced expression of genes in these pathways. High enrichment of genes within the AR pathway, DNA repair, mTOR and p53 signaling among others, was found in all three PDX models (**Fig.4C**). The few effective compounds in BM18 and LAPC9 organoids, compared to PNPCa, is indicative of aggressive tumor phenotype and drug resistance. Antracyclines, mTOR inhibitors and ponatinib are commonly potent compounds on all tumor organoid models tested, and we sought to further investigate their efficacy on patient derived material.

### Defining a drug panel for therapy resistant PCa (PDXs and patient-derived material) for routine organoid screens and treatment decision

In order to develop a personalized treatment approach for PCa patients, we established patient-derived organoids (PDOs) from needle biopsies derived from radical prostatectomy and metastatic specimens. As a control for PDO formation efficiency, we also collected samples from macroscopically cancer-unaffected sites in radical prostatectomy specimens based on pathologist evaluation. On average, this matched control tissue formed fewer organoids compared to samples from malignant prostate cancer **(Fig.5A, “benign” and “tumor”**). PCa PDO showed two main morphological phenotypes *in vitro*: organoids with more acinar or cystic morphology characterized by an empty lumen delimited by a monolayer of cells, and organoids with an adenocarcinoma-like phenotype with an inner core of cells surrounded by a tightly packed outer layer (**Fig.5A, “acinar” and “adenocarcinoma”**). Although PDO cultures showed both inter- and intra-patient morphological heterogeneity, each PDO culture showed consistent morphology across passages. Tumorigenic potential of PCa organoids was assessed by *in vivo* intraprostatic injections (**Sup.Fig.14**, representative cases).

**Figure 5.**
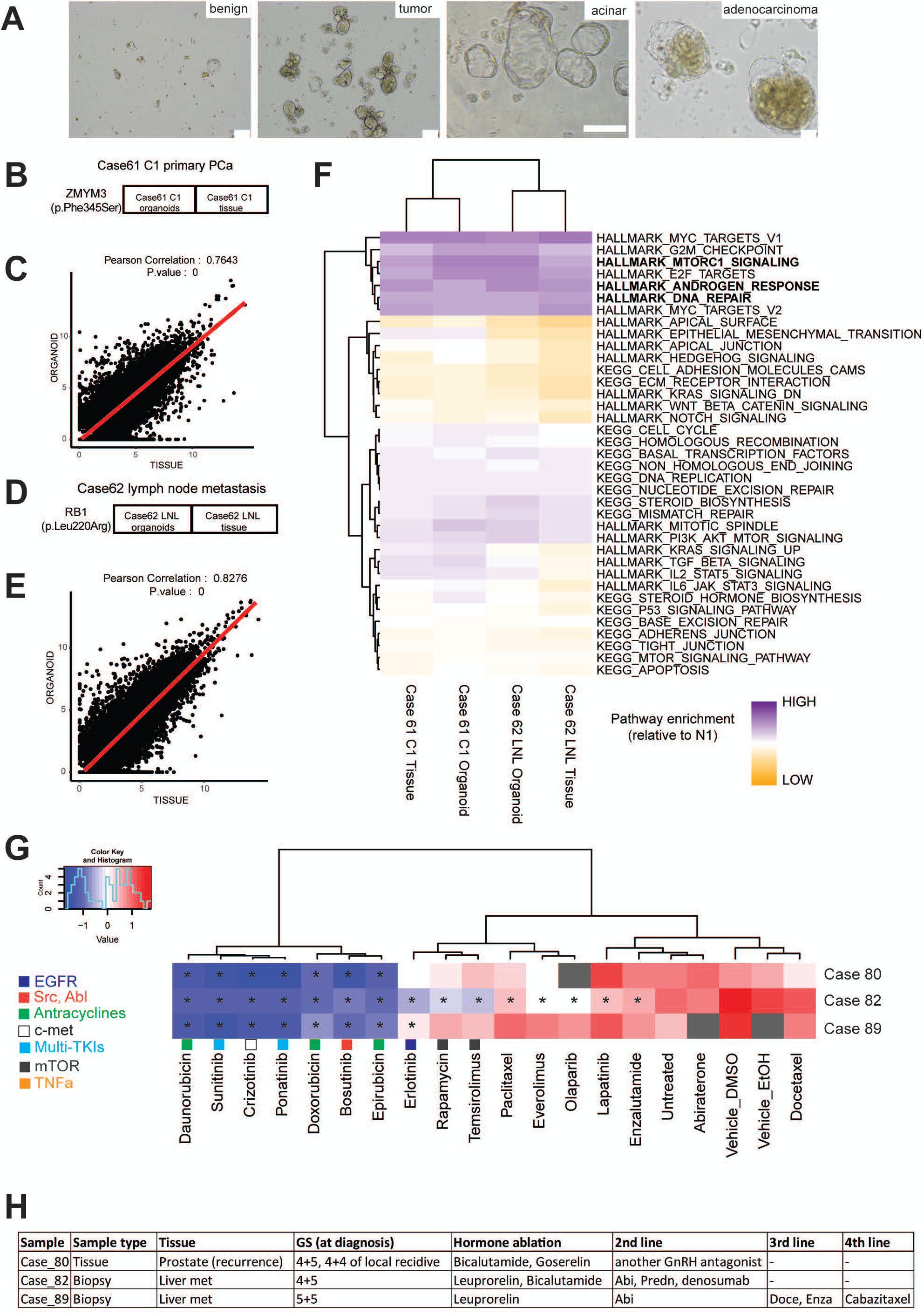
Patient derived organoids from therapy resistant advanced cases, showed sensitivity to multi tyrosine kinase inhibitors identified in the medium-throughput organoid screen. **A**. Representative brightfield images of patient derived organoids (PDOs) from PCa (“tumor”) and from cancer unaffected control area (“benign”) from the same patient (scale bar 100um). Representative images of PDO with an acinar or cystic morphology and with an adenocarcinoma-like morphology (“acinar” (scale bar 500um) and “adenocarcinoma” (scale bar 50um), respectively). **B-E**. Correlation plots of gene expression between tissue and organoids for PCa case 61 (**C**, Pearson correlation coefficient r=0.76) and PCa case 62 (**E**, Pearson correlation coefficient r=0.83). Somatic mutations in *ZMYM3* (Phe345Ser) and in *RB1* (Leu220Arg) were identified in the tissues and organoids of PCa cases 61 **(B)** and 62 **(D). F**. ssGSEA of PDOs and originating tissues, representing the enrichment score of selected pathways from Hallmarks and C2 Kegg (MSigDB) relative to non-carcinoma control tissue. **G**. Results of PDO drug screen assay on three advanced PCa cases (liver metastasis needle biopsies (PCa82,89) or tissue (PCa80). Heatmap represents z-scores from organoid viability assays, after 48h exposure of PDOs to the listed drugs. * p < 0.001; gray squares, data not available. **H**. Clinical features of three advances PCa cases, tested in organoid drug screens.

When sufficient material was available, patients’ blood, PCa tissues and matched organoids were subjected to targeted DNA and RNA sequencing using a panel of clinically relevant cancer-related genes. In primary PCa case 61 we detected a missense mutation (Phe345Ser) in the *ZMYM3* gene (**Fig.5B**), while in pelvic lymph node metastasis PCa case 62 we found a missense mutation (Leu220Arg) in the *RB1* gene (**Fig.5D**). In both cases, there was high correlation of the transcriptomic profiles between the PCa tissues and their matched PDOs (**Fig.5C,E**), including high enrichment for genes in Androgen Response, DNA repair, mTOR and MYC pathways (**Fig.5F**).

We selected a panel of 13 of the most effective compounds (based on statistical significance) resulting from the organoid drug screening performed on PDXs (**Fig.4B, Table 1**) to develop an *in vitro*, multi-drug assay on PDOs. Three advanced cases of PCa were screened for the indicated compound panel together with 4 PCa standard-of-care compounds (**Fig.5G**). One case (PCa case 80) was from a local recurrence after hormone ablation with bicalutamide and radiotherapy. In the other two cases, tissue was obtained from needle biopsies of liver metastasis in which both patients previously underwent hormone ablation therapy and were treated with at least one new generation antiandrogen abiraterone or enzalutamide (**Fig.5H**). While none of the tested cases showed sensitivity to the tested antiandrogens, in line with the clinical history of androgen deprivation, all tested samples were significantly sensitive to TKIs and to the tested anthracyclines (daunorubicin, doxorubicin and epirubicin) (**Fig.5G**). In particular, PCa case 82 exhibited a broader spectrum of sensitivity to the tested compounds (**Fig.5G**), while PCa case 89, which was previously treated with docetaxel and enzalutamide (**Fig.5H**), confirmed resistance to these compounds *in vitro*, endorsing the applicability of a personalized organoid screening approach to PCa patients.

## Discussion

Despite many improvements in the management of PCa in recent years, the increasing lack of therapeutic options during the disease course is a consequence of drug resistance acquisition and of limited number of experimental *in vivo* models that adequately recapitulate complex subtypes. In this study, we describe a novel, early onset and treatment-naïve PCa PNPCa PDX model. We additionally established organoid cultures from this PDX, comparing with other more advanced PCa PDXs (BM18, LAPC9), to develop an organoid-based drug screen pipeline. We then further adapted organoid cultures to patient-derived biopsy material, implementing a clinically-relevant, patient-tailored organoid drug screen.

PCa is generally a slow proliferating tumor with few key mutations and genetic alterations that are commonly found in all patients such as *TMPRSS-ERG, SPOP, FOXA1, PTEN* [31, 32]. Loss of *CHD1* heterozygosity is found in 15% PCa cases and drives prostate-specific cancer growth in transgenic mice [33]. PNPCa contains a frameshift mutation in the tumor suppressor gene *CHD1;* since *CHD1* deletions frequently co-occur with *SPOP* mutations [34], we compared the transcriptome of PNPCa PDX to signatures of C*HD1*-homozygous deletion cases and *SPOP*-mutated cases, which frequently occur together, along with other subtypes (*ETS* rearrangements and *FOXA1* mutations). *SPOP* is associated with DNA repair errors and a higher number of genomic rearrangements [35]. Although no *SPOP* mutation was identified in the PNPCa, there is a transcriptomic correlation with the profile of *SPOP-* mutated cases, and not with the *ERG*-mutated cases, as characterised in previous studies all exomes with *SPOP* mutations lacked *ERG* fusion [36, 37].

The PNPCa model displays MSI and a heterozygous frameshift mutation in the *BRCA2* gene and these events constitute two of the four genomic subtypes of metastatic CRPC based on a recent whole genome sequencing study [26]. *BRCA2* mutation carriers are at higher risk for rapid progression and poorer survival compared to mutated *BRCA1* carriers, highlighting the need for treatment stratification [38]. PNPCa organoids exhibited sensitivity to irradiation, a clinical treatment for patients harbouring *BRCA2* mutations [21]. Phase III clinical trials showed improved outcome of CRPC patients harboring *BRCA1/2* or *ATM* mutations when treated with olaparib compared to AR inhibitors (PROfound, NCT02987543).

Hypermutation and MSI are rare and sporadic features of PCa [39] that are associated with hereditary cancer predisposition. In PCa, high MSI is associated with poorly differentiated stage [40, 41] ranging from 1% of primary tumor cases to up to 12% metastatic cases [42]. To our knowledge, this is the only reported MSI-H case of an early metastasis retaining androgen sensitivity that has been modelled *in vivo*. MSI status has been proposed as a predictive biomarker of response to anti-PD1 immunotherapy with pembrolizumab with positive responses reported for patients with hypermutated MSI who have already been treated with enzalutamide [43, 44]. In 2017 the FDA approved anti-PD-1 treatment for patients with MSI or defective MMR mechanism for prostate [45] as well as other types of cancer, regardless of their original location. Among the patients with defined MSI-H/dMMR molecular phenotype, approximately 50% respond to anti-PD1/PDL1 immunotherapy [44]. Given the lack of biomarkers for CRPC and the higher MSI prevalence in metastatic cases compared to primary cases, screening of patients for MSI status during initial diagnosis could determine whether anti-PD1 treatment is the optimum treatment option. Although PD-L1 protein expression was low to undetectable in PNPCa, RNAseq revealed expression of PD-L1, PD-1 receptor and other immune checkpoint axis mediators. Upon stimulation with IFN-γ, organoids upregulated mRNA expression of PD-L1 proving suggesting they may be receptive to *ex vivo* immunomodulation and, possibly, to anti-PD-L1 immunotherapy. Despite expressing PD-L1, PNPCa organoids could not modulate an immune response *in vitro*, suggesting a multifactorial engagement of this immune checkpoint axis. PNPCa PDX and the original primary tumor showed a high degree of genetic and transcriptomic similarity, suggesting that the metastatic and primary tumor are not highly divergent biologically. Moreover, this molecular similarity was preserved in their derived organoids, supporting the use of organoid cultures for the development of near-patient assays.

Organoids derived from PDXs representing various disease stages were used for drug response profile characterisation, and for identification of potential drug candidates for non-responders to anti-androgens or chemotherapy, by screening compounds clinically used in other cancer types or diseases (drug repurposing [46]). We set up an automated medium-throughput screen of 74 FDA-approved compounds. In our established methodology, we eliminated extracellular matrix components, to facilitate drug availability and to simplify scaling up of organoid drug screens. Organoids were grown in suspension and any associated stromal cells were usually eliminated after one passage. The duration from initial organoid formation until readout of drug screens is two weeks, a time course that is suitable for clinical treatment decisions or treatment follow-up. In our assay, we reduced variability by performing organoid seeding, formation, drug treatment and readout, sequentially in the same culture vessel without transfer of organoids or medium exchange. In addition to the organoid systems, we also cross-validated drug responses using cultured *ex vivo* tissue slices derived from the established PDXs, based on our previously developed methodology [47].

AR-blocker enzalutamide was effective only in the androgen-dependent BM18 and PNPCa organoids, indicating that organoid drug response correlates with individual tumor phenotypes. A few drug classes were effective in all tested models: anthracyclins (often effective at multiple concentrations), TKIs and mTOR inhibitors, in line with the enrichment of PI3K/AKT and mTOR signaling pathway observed in RNAseq analysis. PNPCa was the only model showing sensitivity to EGFR-inhibitors. Deregulation of EGFR signaling is found in a subset of PCa cases, however EGFR inhibitors have showed limited effectiveness [48, 49]. Among the TKI inhibitors identified in our screen, ponatinib was broadly effective, in metastatic PCa PDX as well as in PDOs. While sorafenib and sunitinib have been tested in phase II/III clinical trials for CRPC [50, 51], ponatinib has not been yet investigated in PCa. Interestingly, both sorafenib and ponatinib were identified as CRPC candidate compounds based on a computational gene expression tool for drug identification [52]. Ponatinib inhibits the abnormally constitutive activation of FGFR, SRC, PDGFR and VEGFR which lead to oncogenic signals PI3K/AKT and RAS/RAF/ERK pathways in tumor cells and inhibits angiogenesis. Ponatinib treatment is provided to patients with in acute lymphoblastic leukaemia (PACE Trial [53]) and with acquired resistance to other TKIs [54] and currently its use in solid tumors is being investigated [55].

Proof of principle screens on patient-derived material revealed patient-specific responses and confirmed that organoids from CRPC patients, commonly treated with hormonal ablation, are insensitive to enzalutamide and abiraterone. One case of third- and fourth-line chemotherapy also exhibited lack of sensitivity to docetaxel. Our results highlight the applicability of patient-derived organoid drug screens to predict clinical outcome [20] and their correlation with genomic and transcriptomic features of the primary tumor, as shown in recent studies for lung [56], gastrointestinal cancer [20], hepatocellular carcinoma [57] and ovarian cancer [58]. The panel of drugs can be adapted depending on the individual patient profile e.g. previous treatments, histopathology and targetable genomic alterations. Organoid drug screens on primary or early metastasis PCa are required to assess functional correlation between drug sensitivity and therapy resistance/clonal selection. Additional steps to improve the assay for routine use include the reintroduction of stromal and immune cell types in organoid cultures [59] and the optimization of drug dosage in *vitro* assays to guide their clinical use [60].

In summary, we present a translational pipeline designed by cross-platform analysis of a novel, early-onset and treatment-naïve PCa xenograft model. The specific biologic and genetic landscape of this model may provide insights into tumor growth, metastasis and drug resistance profile at an earlier stage of the disease. Comparison of its drug response profile with those of more advanced PCa PDXs allowed the generation of a highly translational tool for the evaluation of drug response in PDO, thus supporting a precision medicine approach to clinical decision making.

## Materials & Methods

### Patient history

The patient presented with primary prostate cancer (Gleason 9) and underwent transurethral resection of the prostate (TUR-P) procedure. After 6 months, biopsy sampling was performed and the patient was diagnosed with a soft tissue metastasis; histopathology showed an infiltration of an adenocarcinoma from the prostate, PSA 91ng/ml. Orchiectomy was performed directly after biopsy sampling, thus the tumor was androgen-dependent at the time of collection. No biochemical relapse was observed up to 18 months since the diagnosis (PSA < 1ng/ml). All patients included in this study provided written informed consent (Cantonal Ethical approval KEK 06/03 and 2017-02295).

### Tumor sample preparation & xenograft surgery procedure

Needle biopsy from soft tissue metastasis was collected in Dulbecco`s MEM (Gibco, 61965-026) media containing primocin (InVivoGen). For xenograft implantation, needle biopsies were implanted subcutaneously in a Nod Scid Gamma male mouse, under aneasthesia (Domitor® 0.5mg/kg, Dormicum 5mg/kg, Fentanyl 0.05mg/kg). Animal license BE 55/16. Weekly subcutaneous injections of testosterone propionate dissolved in castor oil (Sigma, 86541-5G) were performed (2mg per dosage, 25G needle) starting 1 week after the surgery. For PDX passaging, serial subcutaneous implantations of tumor pieces into new recipients was performed. Abiraterone acetate (Selleckchem, S2246) treatment was administered once daily (ip) for 5 days per week over a duration of 4 weeks at 0.5 mmol/kg/d (5% benzyl alcohol and followed by 95% safflower oil solution).

### DNA isolation from organoids and tissue samples

For DNA extraction from organoids and tissue samples the DNeasy Blood and tissue kit (Qiagen, 69504) was used. DNA from FFPE material was extracted Maxwell® 16 LEV RNA FFPE Purification Kit (Promega, AS1260).

### RNA isolation from organoids and tissue samples

RNA isolation from organoids was performed using the PicoPure Arcturus (Thermo Scientific, KIT0204) kit method. Tissue RNA was extracted using standard protocol of Qiazol (Qiagen) tissue lysis by TissueLyser (2min, 20Hz). Quality of RNA was assessed by Bioanalyzer (Agilent). RNA from FFPE material was extracted using the Maxwell® 16 LEV RNA FFPE Purification Kit (Promega, AS1260).

### Tissue dissociation and Organoid culture

Tumor tissue is collected in Basis medium (Advanced DMEM F12 Serum Free medium (Thermo, 12634010) containing 10mM Hepes (Thermo,15630080), 2mM GlutaMAX supplement (Thermo, 35050061) and 100 μg/ml Primocin (InVivoGen, ant-pm-1)). After mechanical disruption the tissue is washed in Basis medium (220rcf, 5min) and incubated in enzyme mix for tissue dissociation (collagenase type II enzyme mix (Gibco, 17101-015, 5mg/ml dissolved in Basis medium, DNase: 15ug/ml (Roche, 10104159001) and 10 μM Y-27632-HCl Rock inhibitor (Selleckchem, S1049). Enzyme mix volume is adjusted so that the tissue volume does not exceed 1/10 of the total volume and tissue is incubated at 37°C for 1-2h with mixing every 20 minutes. After digestion of large pieces is complete, the suspension is passed through 100um cell strainer (Falcon®, VWR 734-0004) attached to a 50ml Falcon tube. Using a syringe rubber to crash tissue against the strainer & wash in 5ml basic medium (220rcf, 5min). Cell pellet is incubated in 5ml precooled EC lysis buffer (*150 mM NH*_*4*_*Cl, 10 mM KHCO*_*3*_, *0*.*1mM EDTA)*, incubated for 10 min, washed in equal volume of basis medium followed by centrifugation (220rcf, 5min). Pellet is resuspended in 2-5ml accutase™ (StemCell Technologies, 07920), depending on the sample amount; biopsies vs tissue and incubated for 10min at room temperature. The cell suspension is passed through 40um pore size strainer (Falcon®, VWR 734-0004), and the strainer is washed by adding 2ml of accutase on the strainer. Single cell suspension is counted to determine seeding density, and washed in 5ml of basis medium and spin down 220rcf, 5min. Cell pellet is reconstituted in organoid medium and seeded in ultra low attachment (ULA) plates; e.g. 30.000 cells per well in 96well plates with 100ul media, 100.000 cells per well of 24well plate with 750ul media, 300.000 to 500.000 cells per well of 6well plate (2ml). Plates are low attachment 96, 24 and 6-well (Corning, Costar #3471). Organoid culture media contains the following reagents: Basis medium containing 10μM Y-27632-HCl (Selleckchem, S1049), 5% fetal calf serum (Gibco #10270-106, LOT 42G7277K), 1x B-27 supplement (ThermoFisher, 17504044), 10mM Nicotinamide (Sigma, N0636 100G), 500ng/ml Rspondin (Peprotech, 120-38), 1.25mM N-acetyl-cysteine (Sigma, A9165), 10μM SB202190 (Selleckchem, S1077), 100ng/ml Noggin (Peprotech, 250-38), 500nM A83-01 (Tocris, 2939), 10nM Dihydrotestosterone DHT (Fluka Chemica, 10300), 10ng/ml Wnt3A, 50 ng/ml HGF (Peprotech, 100-39), 50ng/ml EGF, 10ng/ml FGF10 (Peprotech, 100-26), 1ng/ml FGF2 (Peprotech, 100-18B), 1μM PGE2 (Tocris, 2296). Media is prepared and kept at 4C for no longer than 7 days.

### Medium-throughput organoid drug screen at NEXUS Personalized Health Technologies automation platform

#### Compounds

A drug library was compiled based on predicted activity against prostate cancer (Selleck Chemicals, Houston TX, USA) as 96-well format sample storage tubes with drugs in 10mM concentration. Using a Tecan EVO 100 (Tecan AG, Männedorf, Switzerland), this drug library was aliquoted over 96-well plates (#651261, Greiner Bio-One, Kremsmünster, Austria) and further diluted to yield stock plates with a concentration of 10mM, 1mM and 0.1mM in DMSO (Sigma Aldrich, cat. D8418). After aliquotting, plates were sealed under argon gas using an Agilent PlateLoc (Agilent Technologies, Santa Clara CA, USA) with peelable aluminium heat-sealing foil (Agilent, cat. 24210-001) for 1s at 170°C. An overview of the purchased drugs and their known targets can be found in **SI Table 3**. Control molecules enzalutamide, docetaxel and doxorubicin were purchased at Sellechekchem (Lubio Science, Zürich, Switzerland, #S1250, #S1148, #S1208). Staurosporine was purchased at Toronto Research Chemicals (Toronto, Canada, #S685000).

#### Automated drug screening with prostate cancer organoids

Automated screening procedures were performed at NEXUS Personalized Health Technologies (ETH Zürich, Zürich, Switzerland) using an automated screening platform (HighRes Biosolutions, Beverley MA, USA). Prostate cancer organoids of BM18, LAPC9 or PNPCa origin were prepared and expanded from murine tumor tissue as for five to seven days to allow organoid formation using Costar ultra-low attachment plates (#3471, Corning, New York NY, USA). For the drug screens, organoids were dissociated into single cell suspension by both enzymatic (TrypLE incubation) and mechanical separation (22G needle), counted and seeded in ULA 384 well plates at appropriate cell density for each tumor model; 3’500 c/well (LAPC9, BM18) or 5’000 c/well (PNPCa). Cells were seeded 25uL per well in 384-well flat-bottom ultra-low attachment plates (#3827, Corning) using a BioTek EL406 with wide-bore 5uL peristaltic pump tubing (BioTek Instruments Inc., Winooski VT, USA). After cell seeding, plates were shaken for 2 minutes, incubated for 1 hour at room temperature and subsequently transferred to a 37°C incubator with 95% humidity and 5% CO_2_. 48 hours after cell seeding, 96-well plates (#651261, Greiner Bio-One) containing 1000-times concentrated compound stock solutions at differing concentrations (10mM, 1mM and 0.1mM) in DMSO were centrifuged at 250rcf for 10 seconds (HiG 5000, BioNex Solutions Inc., San Jose CA, USA) and desealed using a Brooks Xpeel (Brooks Life Sciences, Chelmsford MA, USA) and subsequently diluted 1:125 in culture medium and added to quadrant 1, 2 and 3 (respectively for 10mM, 1mM and 0.1mM stock plates) of deepwell 384-well plates (#781271, Greiner Bio-One). DMSO-stock solutions of control molecules (100% DMSO as negative control and as positive controls we included 1000-times concentrated docetaxel [30uM], enzalutamide [6mM] and doxorubicin [10mM] for LAPC9 and BM18 organoids, or staurosporine [2.5mM] instead of docetaxel for PNPCa organoids) were added to plate quadrant 4. An additional 1:1 dilution step was done prior to adding 20uL of diluted drugs to the cell culture plates (containing 20uL cell suspension after correction for evaporation). Compound dilutions were performed using automated liquid handling equipment (Tecan AG, Männedorf, Switzerland) in technical triplicate (LAPC9 and BM18) or technical quadruplicate (PNPCa). A Schematic representation of the compound dilution and addition procedure is shown in **Sup.Fig.15**. After compound exposure, compound DMSO stock plates were sealed under argon gas using an Agilent PlateLoc as described above, and organoid culture plates were transferred back to the 37°C incubator with a 95% humidity and 5% CO_2_ atmosphere. 48 hours after compound exposure, a CellTiter-Glo 3D assay (#G9682, Promega, Madison WI, USA) was used to measure ATP levels as a proxy for cell viability. This assay is lytic and thus maximized readout from all cells composing large organoid structures. The cell viability readout was performed according to manufacturer’s instructions using the automation equipment. Briefly, 40uL room-temperature CellTiter-Glo 3D reagent was added per well to the assay plates using an automated liquid handler (Tecan AG). Plates were subsequently shaken for 5 minutes on a BioTek EL406 and incubated in a temperature-controlled incubator at 22°C for 25 minutes. After incubation, luminescence was measured using a Tecan M1000 Pro plate reader (Tecan AG) with 1000ms integration time. Results were collected as spreadsheets and coupled to plate layouts.

Data have been normalized using the median of the negative control conditions (DMSO 0.1%) and values have been log2 transformed (each plate with its own internal negative control). For the statistical analysis the different plates have been considered as replicates. p-value and FDR were calculated for all drugs after removing DMSO and no-treated control conditions).

## Supporting information

Supplementary Figures with legends

## Acknowledgements

We thank all patients that participated in our study, all involved clinical personnel and study nurses. We would like to acknowledge Microscopy Facility of University of Bern, Viola Paradiso for the ion torrent sequencing. Source of Funding: Swiss National Science Foundation 31003A_169352; 310030_189149; UniBe Initiator Grant 2016; KWF 2015_7599; Novartis 17B076; SOCIBP SPHN Driver Project (SAM).

## Author contributions

S.K. and F.L.M. designed experiments, acquired data, interpreted data and wrote manuscript. M.R.D.F. performed bioinformatics data analysis and revised the manuscript. M.K., M.D.M., E.Z., J.G. and I.K, performed additional experiments on clinical samples and animal experiments. A.G., M.B. and J.P.T. performed bioinformatics data analysis and revised the manuscript. V.G provided the pathological evaluation. D.K acquired data and provided technical support. S.K., T.H.B and C.U.S generated the pipeline for FDA approved drug library tests, optimized the automation procedure for the PCa organoids, acquired data and wrote the manuscript. K.E. performed sequencing experiments. A.S., C.K.Y.N and S.P. performed bioinformatic data analysis. P.C.G. interpreted data and revised manuscript. M.S. provided clinical samples and revised manuscript. M.A.R. interpreted data and revised manuscript. G.N.T. provided clinical samples, interpreted data and revised manuscript. M.K.D.J. designed concept of the study and experiments, interpreted data and wrote manuscript.

## References

1. Ferlay, J., et al., Cancer incidence and mortality patterns in Europe: Estimates for 40 countries and 25 major cancers in 2018. European Journal of Cancer, 2018. 103: p. 356–387.

2. Beer, T.M., et al., Enzalutamide in Metastatic Prostate Cancer before Chemotherapy. New England Journal of Medicine, 2014. 371(5): p. 424–433.

3. de Bono, J.S., et al., Abiraterone and Increased Survival in Metastatic Prostate Cancer. New England Journal of Medicine, 2011. 364(21): p. 1995–2005.

4. Byrne, A.T., et al., Interrogating open issues in cancer precision medicine with patient-derived xenografts. Nature Reviews Cancer, 2017. 17(4): p. 254–268.

5. Wang, Y., et al., Development and characterization of efficient xenograft models for benign and malignant human prostate tissue. Prostate, 2005. 64(2): p. 149–59.

6. Wetterauer, C., et al., Early development of human lymphomas in a prostate cancer xenograft program using triple knock-out Immunocompromised mice. The Prostate, 2015. 75(6): p. 585–592.

7. Navone, N.M., et al., Movember GAP1 PDX project: An international collection of serially transplantable prostate cancer patient-derived xenograft (PDX) models. The Prostate, 2018. 78(16): p. 1262–1282.

8. Lin, D., et al., High Fidelity Patient-Derived Xenografts for Accelerating Prostate Cancer Discovery and Drug Development. Cancer Research, 2014. 74(4): p. 1272–1283.

9. Pauli, C., et al., Personalized In Vitro and In Vivo Cancer Models to Guide Precision Medicine. Cancer Discov, 2017. 7(5): p. 462–477.

10. Raheem, O., et al., A novel patient-derived intra-femoral xenograft model of bone metastatic prostate cancer that recapitulates mixed osteolytic and osteoblastic lesions. J Transl Med, 2011. 9: p. 185.

11. Williams, E.S., et al., Generation of Prostate Cancer Patient Derived Xenograft Models from Circulating Tumor Cells. J Vis Exp, 2015(105): p. 53182.

12. Puca, L., et al., Patient derived organoids to model rare prostate cancer phenotypes. Nat Commun, 2018. 9(1): p. 2404.

13. Klein, K.A., et al., Progression of metastatic human prostate cancer to androgen independence in immunodeficient SCID mice. Nature Medicine, 1997. 3(4): p. 402–408.

14. McCulloch, D.R., et al., BM18: A novel androgen-dependent human prostate cancer xenograft model derived from a bone metastasis. The Prostate, 2005. 65(1): p. 35–43.

15. Craft, N., et al., Evidence for Clonal Outgrowth of Androgen-independent Prostate Cancer Cells from Androgen-dependent Tumors through a Two-Step Process. Cancer Research, 1999. 59(19): p. 5030–5036.

16. Priolo, C., et al., Establishment and genomic characterization of mouse xenografts of human primary prostate tumors. Am J Pathol, 2010. 176(4): p. 1901–13.

17. Kondo, J. and M. Inoue, Application of Cancer Organoid Model for Drug Screening and Personalized Therapy. Cells, 2019. 8(5): p. 470.

18. Bleijs, M., et al., Xenograft and organoid model systems in cancer research. The EMBO Journal, 2019. 38(15): p. e101654.

19. Granat, L.M., et al., The promises and challenges of patient-derived tumor organoids in drug development and precision oncology. Animal Models and Experimental Medicine, 2019. 2(3): p. 150–161.

20. Vlachogiannis, G., et al., Patient-derived organoids model treatment response of metastatic gastrointestinal cancers. Science, 2018. 359(6378): p. 920–926.

21. Castro, E., et al., Germline BRCA Mutations Are Associated With Higher Risk of Nodal Involvement, Distant Metastasis, and Poor Survival Outcomes in Prostate Cancer. Journal of Clinical Oncology, 2013. 31(14): p. 1748–1757.

22. Risbridger, G.P., et al., Patient-derived Xenografts Reveal that Intraductal Carcinoma of the Prostate Is a Prominent Pathology in BRCA2 Mutation Carriers with Prostate Cancer and Correlates with Poor Prognosis. European Urology, 2015. 67(3): p. 496–503.

23. Taylor, R.A., et al., Germline BRCA2 mutations drive prostate cancers with distinct evolutionary trajectories. Nature Communications, 2017. 8(1): p. 13671.

24. Vesprini, D., et al., The therapeutic ratio is preserved for radiotherapy or cisplatin treatment in BRCA2-mutated prostate cancers. Canadian Urological Association journal = Journal de l’Association des urologues du Canada, 2011. 5(2): p. E31–E35.

25. Ma, Y., et al., Response to olaparib in metastatic castration-resistant prostate cancer with germline BRCA2 mutation: a case report. BMC medical genetics, 2018. 19(1): p. 185–185.

26. van Dessel, L.F., et al., The genomic landscape of metastatic castration-resistant prostate cancers using whole genome sequencing reveals multiple distinct genotypes with potential clinical impact. bioRxiv, 2019: p. 546051.

27. Alexandrov, L.B., et al., Signatures of mutational processes in human cancer. Nature, 2013. 500(7463): p. 415–421.

28. Niu, B., et al., MSIsensor: microsatellite instability detection using paired tumor-normal sequence data. Bioinformatics, 2014. 30(7): p. 1015–6.

29. Rabinovich, G.A. and J.R. Conejo-García, Shaping the Immune Landscape in Cancer by Galectin-Driven Regulatory Pathways. Journal of Molecular Biology, 2016. 428(16): p. 3266–3281.

30. O’Donnell, J.S., M.W.L. Teng, and M.J. Smyth, Cancer immunoediting and resistance to T cell-based immunotherapy. Nature Reviews Clinical Oncology, 2019. 16(3): p. 151–167.

31. Robinson, D., et al., Integrative Clinical Genomics of Advanced Prostate Cancer. Cell, 2015. 161(5): p. 1215–1228.

32. Grasso, C.S., et al., The mutational landscape of lethal castration-resistant prostate cancer. Nature, 2012. 487: p. 239.

33. Augello, M.A., et al., CHD1 Loss Alters AR Binding at Lineage-Specific Enhancers and Modulates Distinct Transcriptional Programs to Drive Prostate Tumorigenesis. Cancer Cell, 2019. 35(5): p. 817–819.

34. Rescigno, P., et al., Molecular and clinical implications of CHD1 loss and SPOP mutations in advanced prostate cancer. Journal of Clinical Oncology, 2018. 36(15_suppl): p. 5064–5064.

35. Boysen, G., et al., SPOP mutation leads to genomic instability in prostate cancer. Elife, 2015. 4.

36. Cancer Genome Atlas Research, N., The Molecular Taxonomy of Primary Prostate Cancer. Cell, 2015. 163(4): p. 1011–1025.

37. Barbieri, C.E., et al., Exome sequencing identifies recurrent SPOP, FOXA1 and MED12 mutations in prostate cancer. Nat Genet, 2012. 44(6): p. 685–9.

38. Narod, S.A., et al., Rapid progression of prostate cancer in men with a BRCA2 mutation. British Journal Of Cancer, 2008. 99: p. 371.

39. Prtilo, A., et al., Tissue microarray analysis of hMSH2 expression predicts outcome in men with prostate cancer. J Urol, 2005. 174(5): p. 1814-8; discussion 1818.

40. Uchida, T., et al., Microsatellite instability in prostate cancer. Oncogene, 1995. 10(5): p. 1019–1022.

41. Egawa, S., et al., Genomic Instability of Microsatellite Repeats in Prostate Cancer: Relationship to Clinicopathological Variables. Cancer Research, 1995. 55(11): p. 2418–2421.

42. Hempelmann, J.A., et al., Microsatellite instability in prostate cancer by PCR or next-generation sequencing. Journal for immunotherapy of cancer, 2018. 6(1): p. 29–29.

43. Graff, J.N., et al., Early evidence of anti-PD-1 activity in enzalutamide-resistant prostate cancer. Oncotarget, 2016. 7(33): p. 52810–52817.

44. Abida, W., et al., Analysis of the Prevalence of Microsatellite Instability in Prostate Cancer and Response to Immune Checkpoint Blockade. JAMA Oncology, 2019. 5(4): p. 471–478.

45. Nava Rodrigues, D., et al., Immunogenomic analyses associate immunological alterations with mismatch repair defects in prostate cancer. J Clin Invest, 2018. 128(10): p. 4441–4453.

46. Bertolini, F., V.P. Sukhatme, and G. Bouche, Drug repurposing in oncology—patient and health systems opportunities. Nature Reviews Clinical Oncology, 2015. 12: p. 732.

47. Karkampouna, S., et al., CRIPTO promotes an aggressive tumour phenotype and resistance to treatment in hepatocellular carcinoma. J Pathol, 2018. 245(3): p. 297–310.

48. Sridhar, S.S., et al., A multicenter phase II clinical trial of lapatinib (GW572016) in hormonally untreated advanced prostate cancer. Am J Clin Oncol, 2010. 33(6): p. 609–13.

49. Guerin, O., et al., EGFR Targeting in Hormone-Refractory Prostate Cancer: Current Appraisal and Prospects for Treatment. Pharmaceuticals (Basel), 2010. 3(7): p. 2238–2247.

50. Beardsley, E.K., et al., A phase II study of sorafenib in combination with bicalutamide in patients with chemotherapy-naive castration resistant prostate cancer. Invest New Drugs, 2012. 30(4): p. 1652–9.

51. Zurita, A.J., et al., Sunitinib in combination with docetaxel and prednisone in chemotherapy-naive patients with metastatic, castration-resistant prostate cancer: a phase 1/2 clinical trial. Ann Oncol, 2012. 23(3): p. 688–94.

52. Kim, I.W., J.H. Kim, and J.M. Oh, Screening of Drug Repositioning Candidates for Castration Resistant Prostate Cancer. Front Oncol, 2019. 9: p. 661.

53. Cortes, J.E., et al., Ponatinib efficacy and safety in Philadelphia chromosome-positive leukemia: final 5-year results of the phase 2 PACE trial. Blood, 2018. 132(4): p. 393–404.

54. Sanford, D., et al., Phase II trial of ponatinib in patients with chronic myeloid leukemia resistant to one previous tyrosine kinase inhibitor. Haematologica, 2015. 100(12): p. e494–5.

55. Tan, F.H., et al., Ponatinib: a novel multi-tyrosine kinase inhibitor against human malignancies. OncoTargets and therapy, 2019. 12: p. 635–645.

56. Kim, M., et al., Patient-derived lung cancer organoids as in vitro cancer models for therapeutic screening. Nature Communications, 2019. 10(1): p. 3991.

57. Nuciforo, S., et al., Organoid Models of Human Liver Cancers Derived from Tumor Needle Biopsies. Cell Reports, 2018. 24(5): p. 1363–1376.

58. Phan, N., et al., A simple high-throughput approach identifies actionable drug sensitivities in patient-derived tumor organoids. Communications Biology, 2019. 2(1): p. 78.

59. Richards, Z., et al., Prostate Stroma Increases the Viability and Maintains the Branching Phenotype of Human Prostate Organoids. iScience, 2019. 12: p. 304–317.

60. Liston, D.R. and M. Davis, Clinically Relevant Concentrations of Anticancer Drugs: A Guide for Nonclinical Studies. Clinical cancer research : an official journal of the American Association for Cancer Research, 2017. 23(14): p. 3489–3498.

