## Supplementary Figures with legends for "Patient-derived xenografts and organoids model therapy response in prostate cancer"

**Sup.Fig.1**

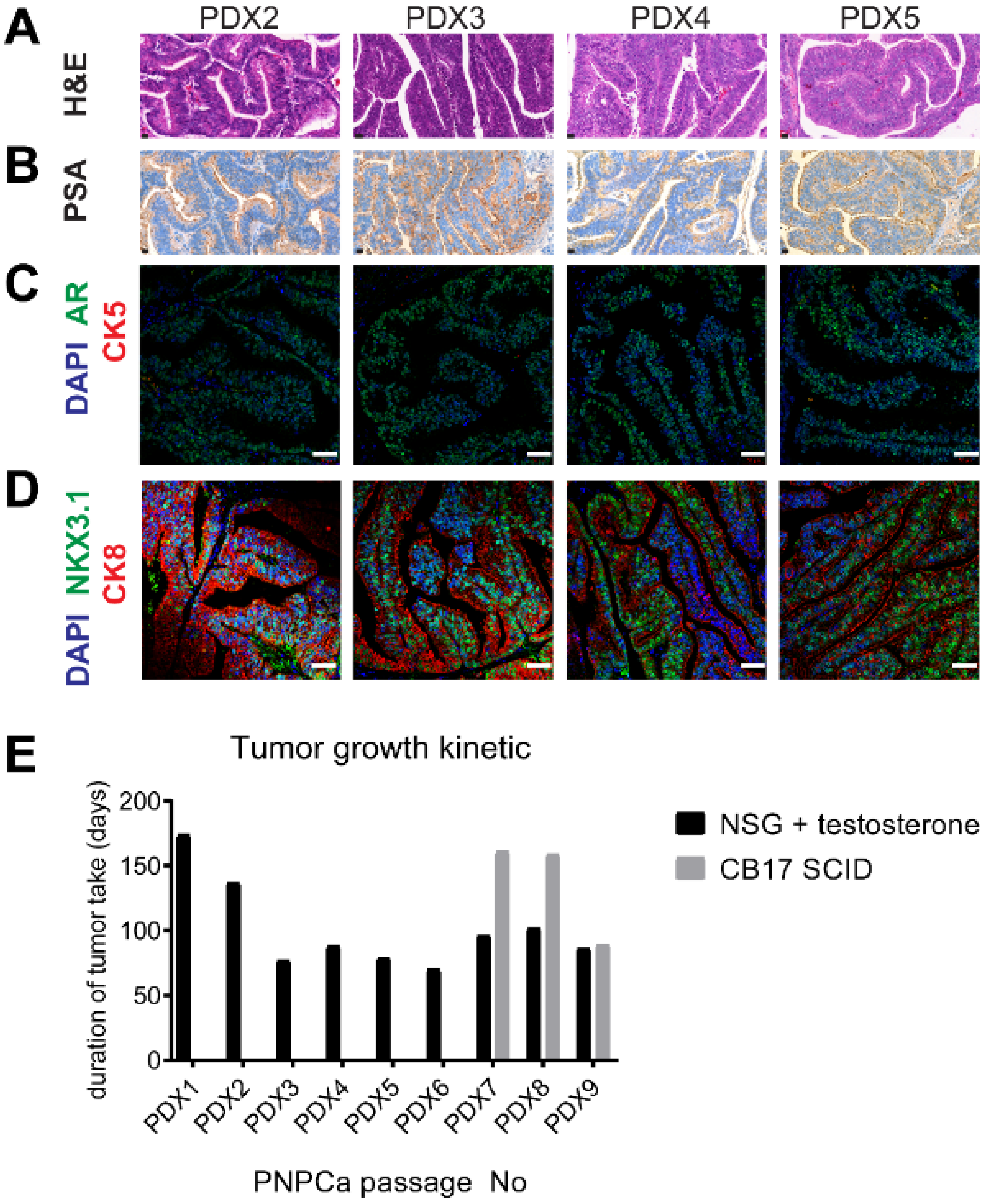

A
PDX2
PDX3
PDX4
PDX5
H&E
B
PSA
C
DAPI AR CK5
D
DAPI NKX3.1 CK8
E
Tumor growth kinetic
duration of tumor take (days)
200
150
100
50
0
PDX1
PDX2
PDX3
PDX4
PDX5
PDX6
PDX7
PDX8
PDX9
PNPCa passage No
NSG + testosterone
CB17 SCID

**Sup.Fig.2**

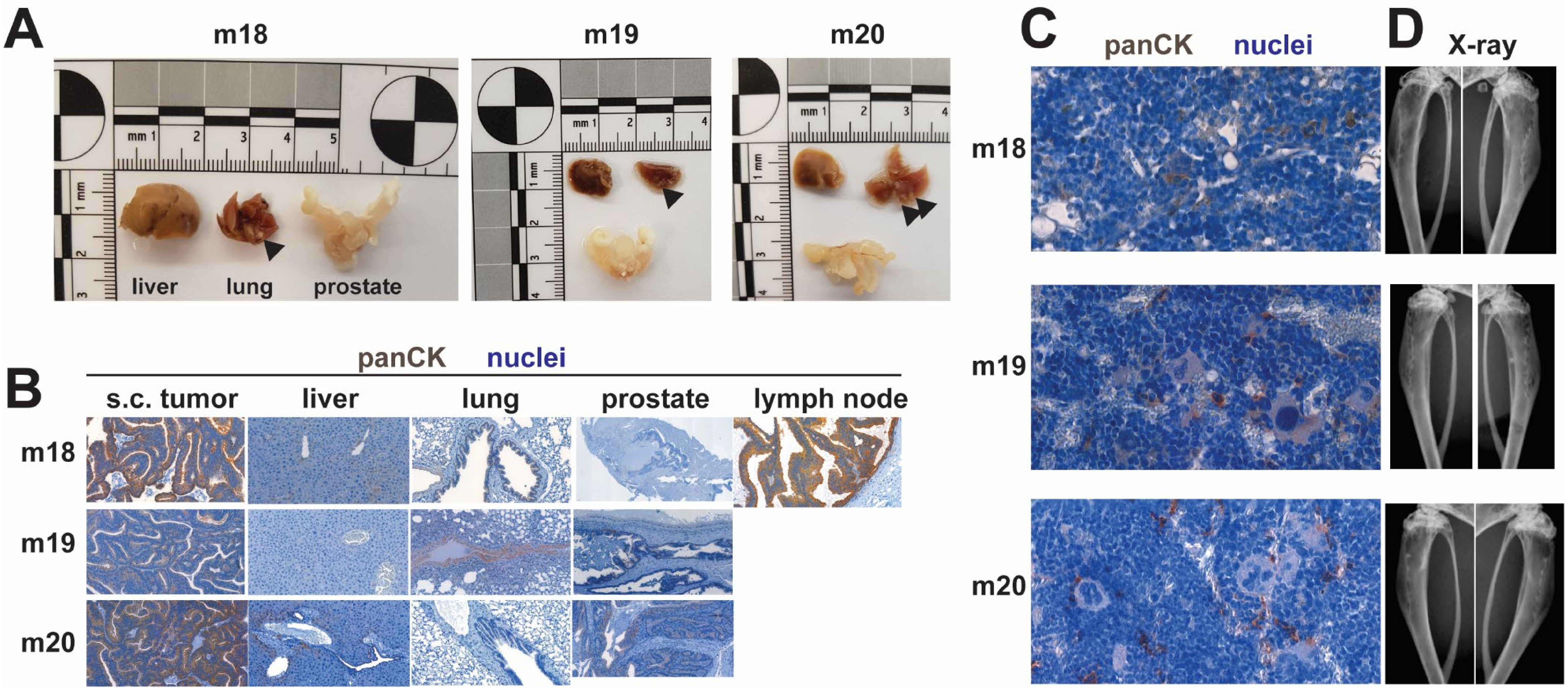
A
m18
m19
m20
liver
lung
prostate
B
panCK
nuclei
s.c. tumor
liver
lung
prostate
lymph node
m18
m19
m20
C
panCK
nuclei
m18
m19
m20
D
X-ray

**Sup.Fig.3**

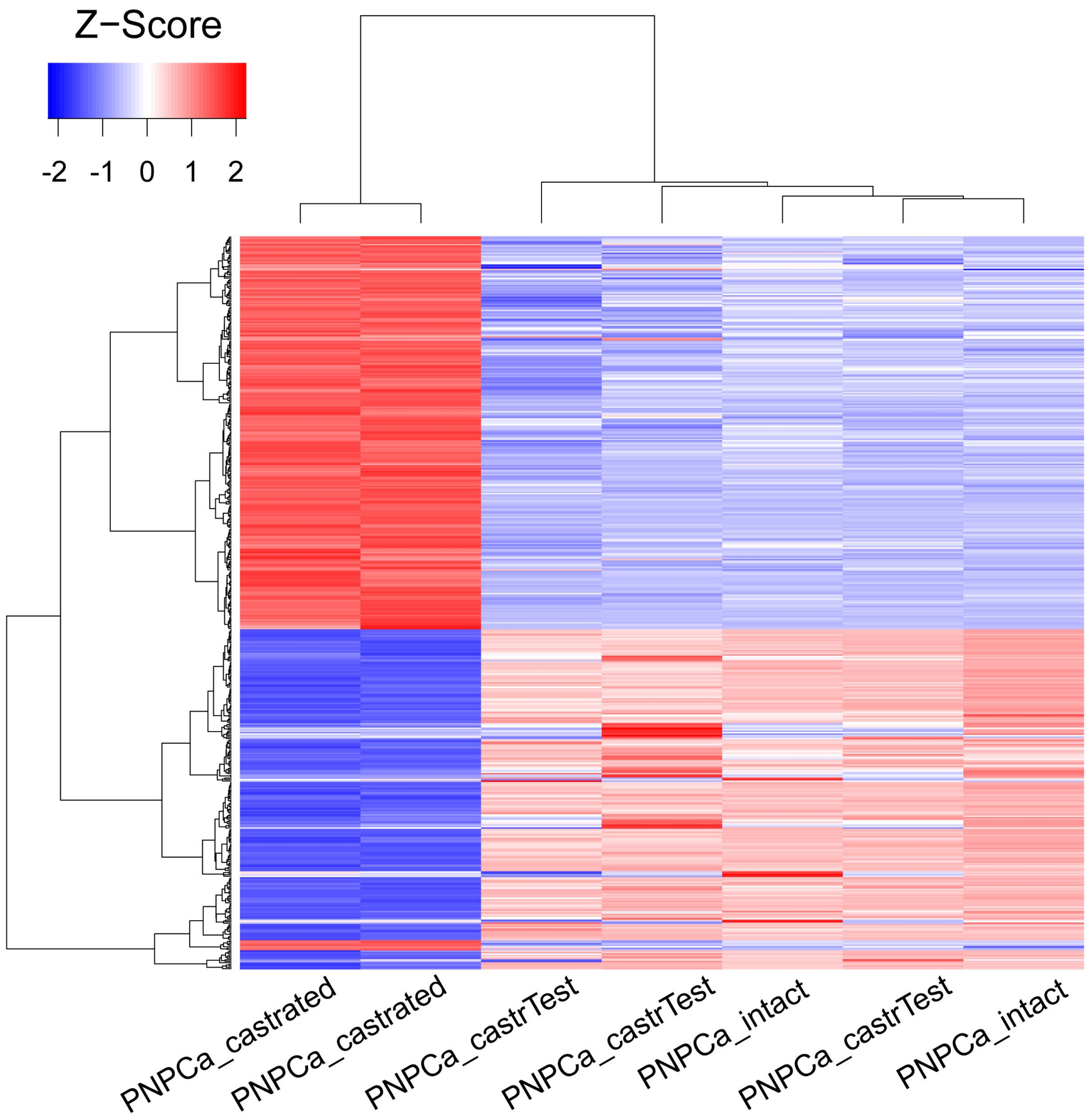
Z−Score
-2 -1 0 1 2
PNPCa_castrated
PNPCa_castrated
PNPCa_castrTest
PNPCa_castrTest
PNPCa_intact
PNPCa_castrTest
PNPCa_intact

**Sup.Fig.4**

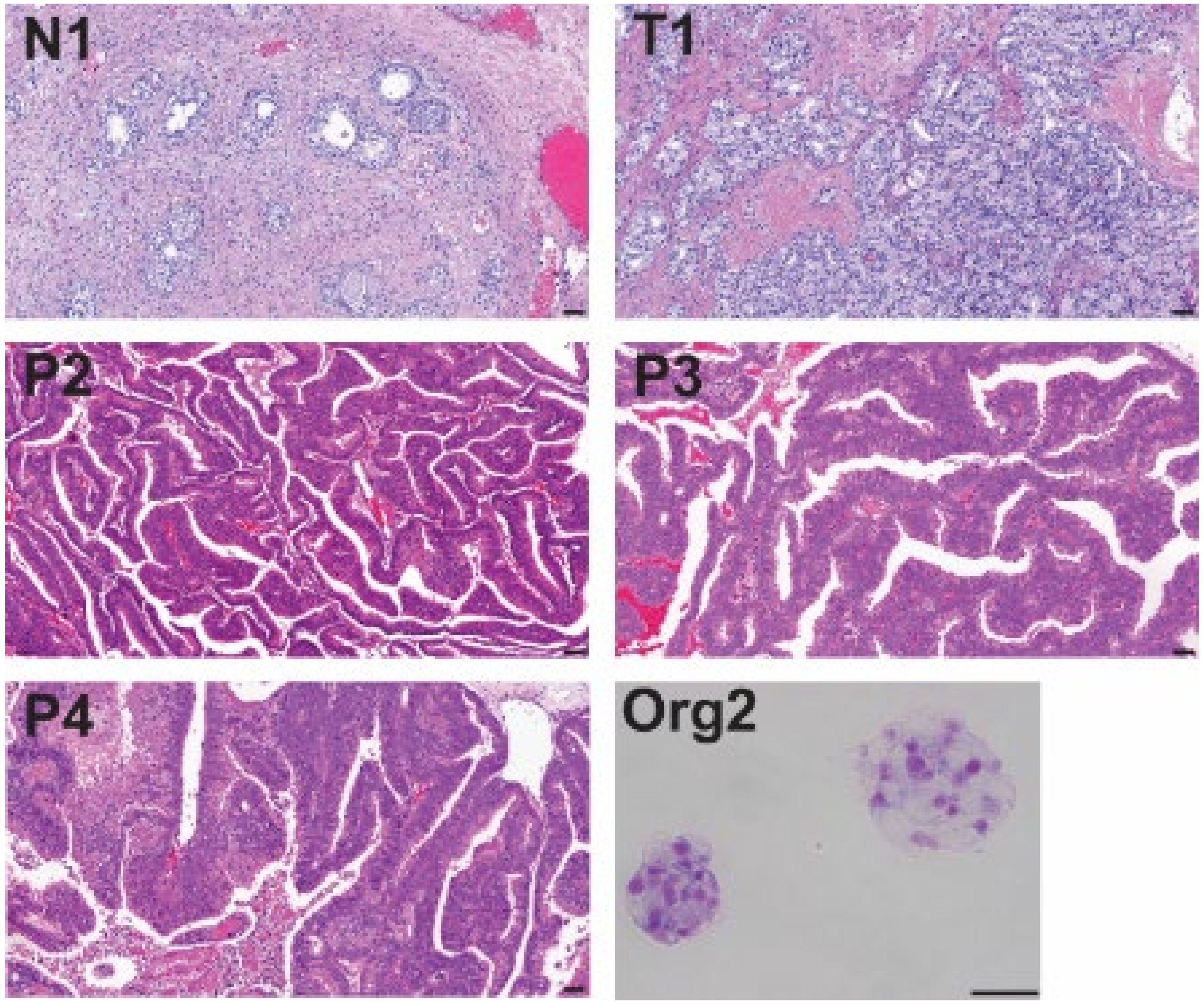

### Sup.Fig.5

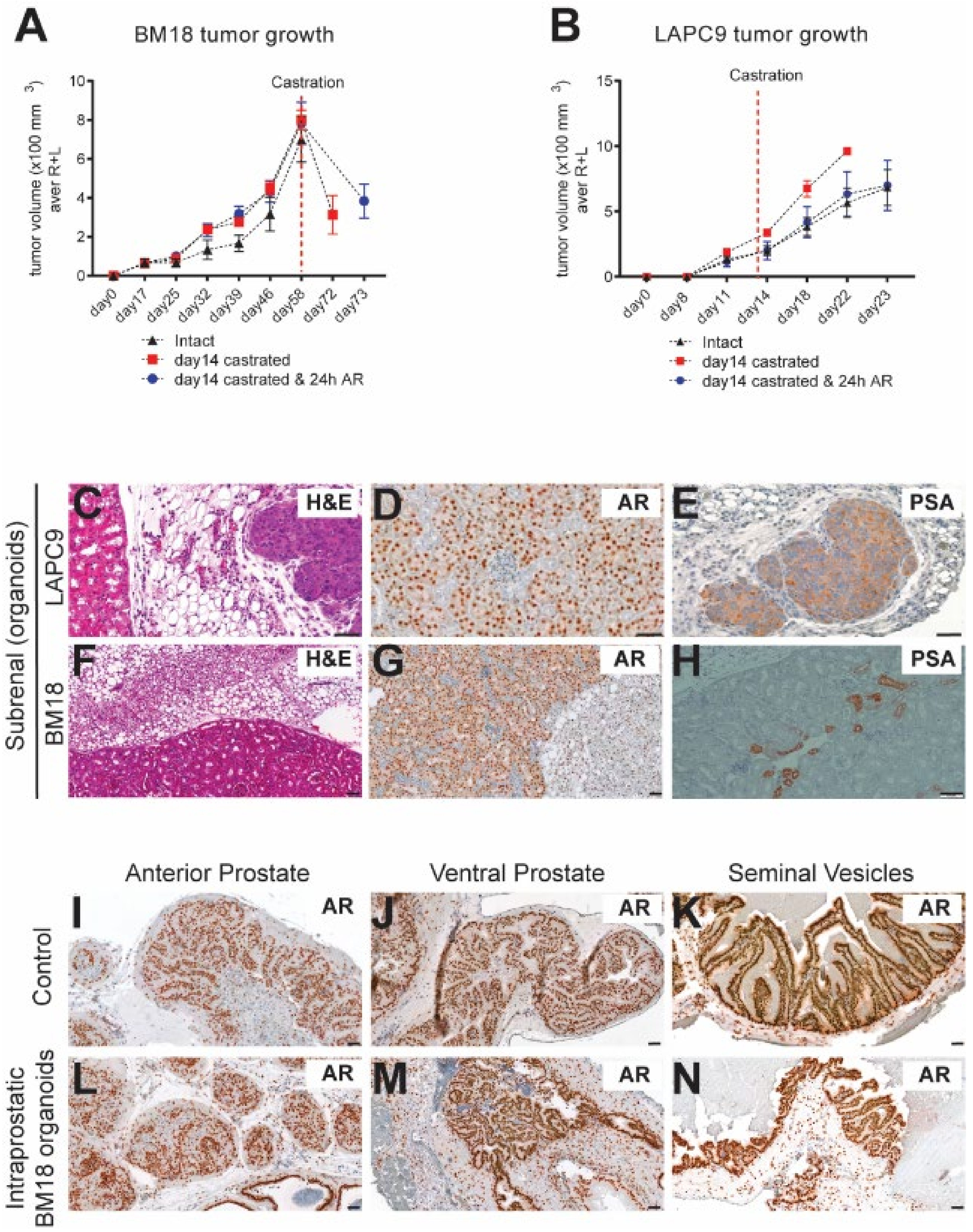

A
BM18 tumor growth
tumor volume (x100 mm 3) aver R+L
Castration
day0
day17
day25
day32
day39
day46
day58
day72
day73
Intact
day14 castrated
day14 castrated & 24h AR
B
LAPC9 tumor growth
tumor volume (x100 mm 3) aver R+L
Castration
day0
day8
day11
day14
day18
day22
day23
Intact
day14 castrated
day14 castrated & 24h AR
Subrenal (organoids)
LAPC9
BM18
C
H&E
D
AR
E
PSA
F
H&E
G
AR
H
PSA
Anterior Prostate
Ventral Prostate
Seminal Vesicles
Control
Intraprostatic BM18 organoids
I
AR
J
AR
K
AR
L
AR
M
AR
N
AR

#### Sup.Fig.6

**A**

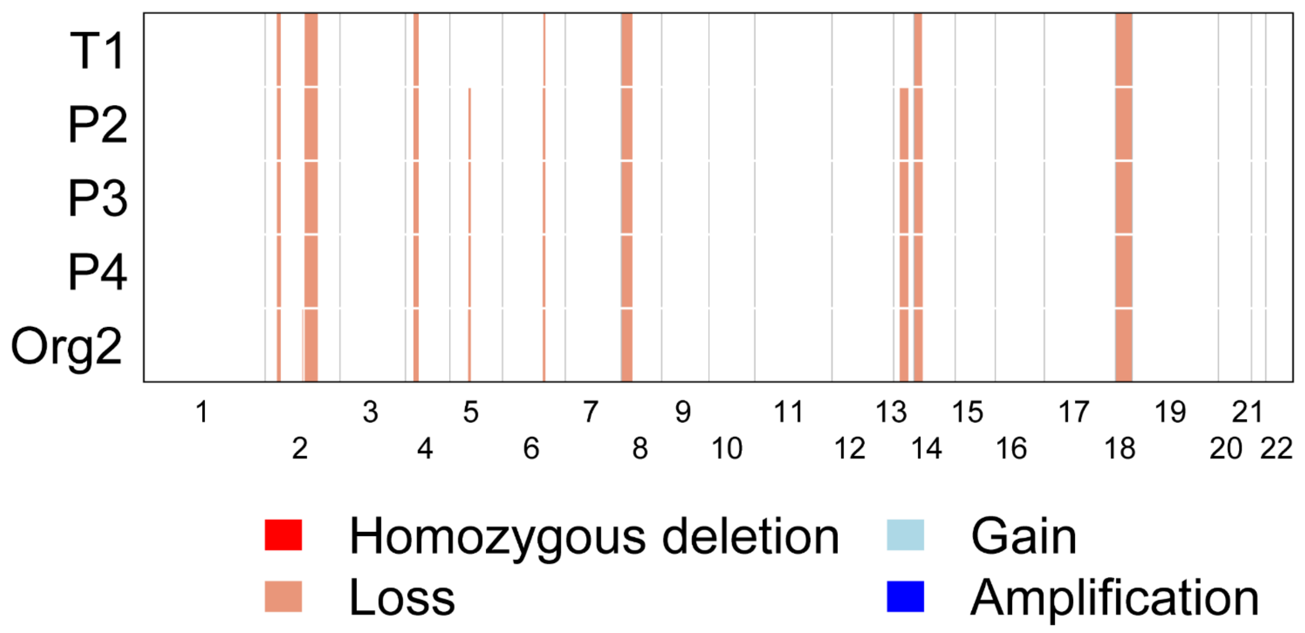
T1
P2
P3
P4
Org2
Homozygous deletion
Loss
Gain
Amplification

**B**

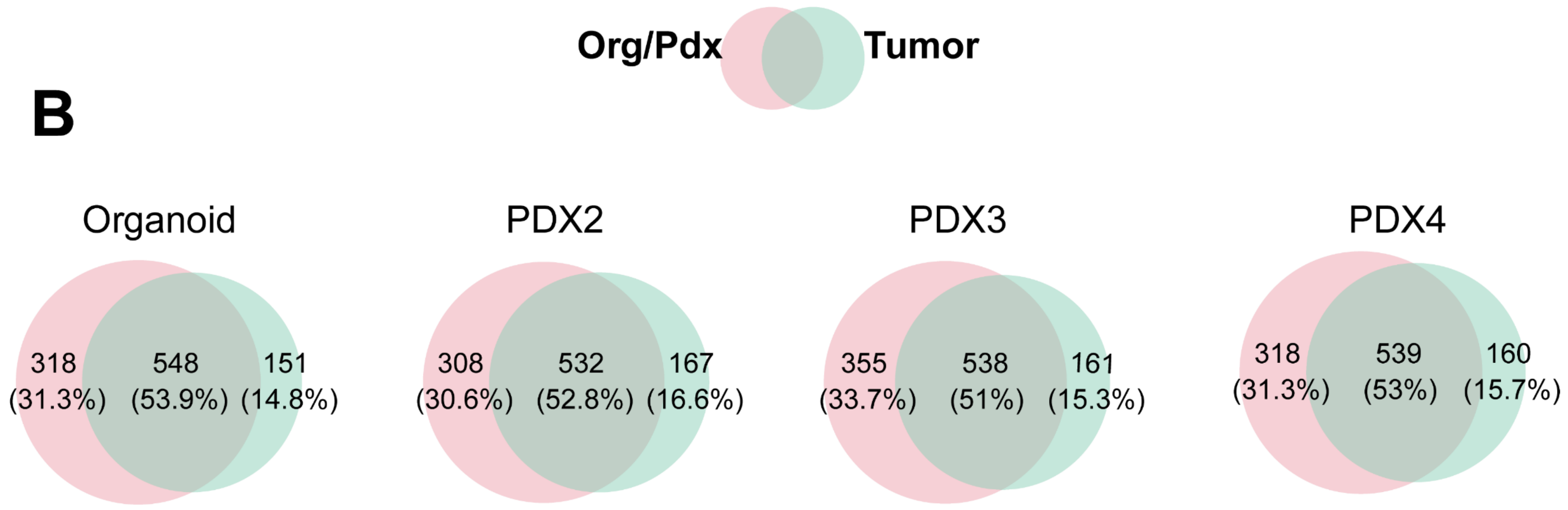
Org/Pdx
Tumor
Organoid
318 (31.3%)
548 (53.9%)
151 (14.8%)
PDX2
308 (30.6%)
532 (52.8%)
167 (16.6%)
PDX3
355 (33.7%)
538 (51%)
161 (15.3%)
PDX4
318 (31.3%)
539 (53%)
160 (15.7%)

**Sup.Fig.7**

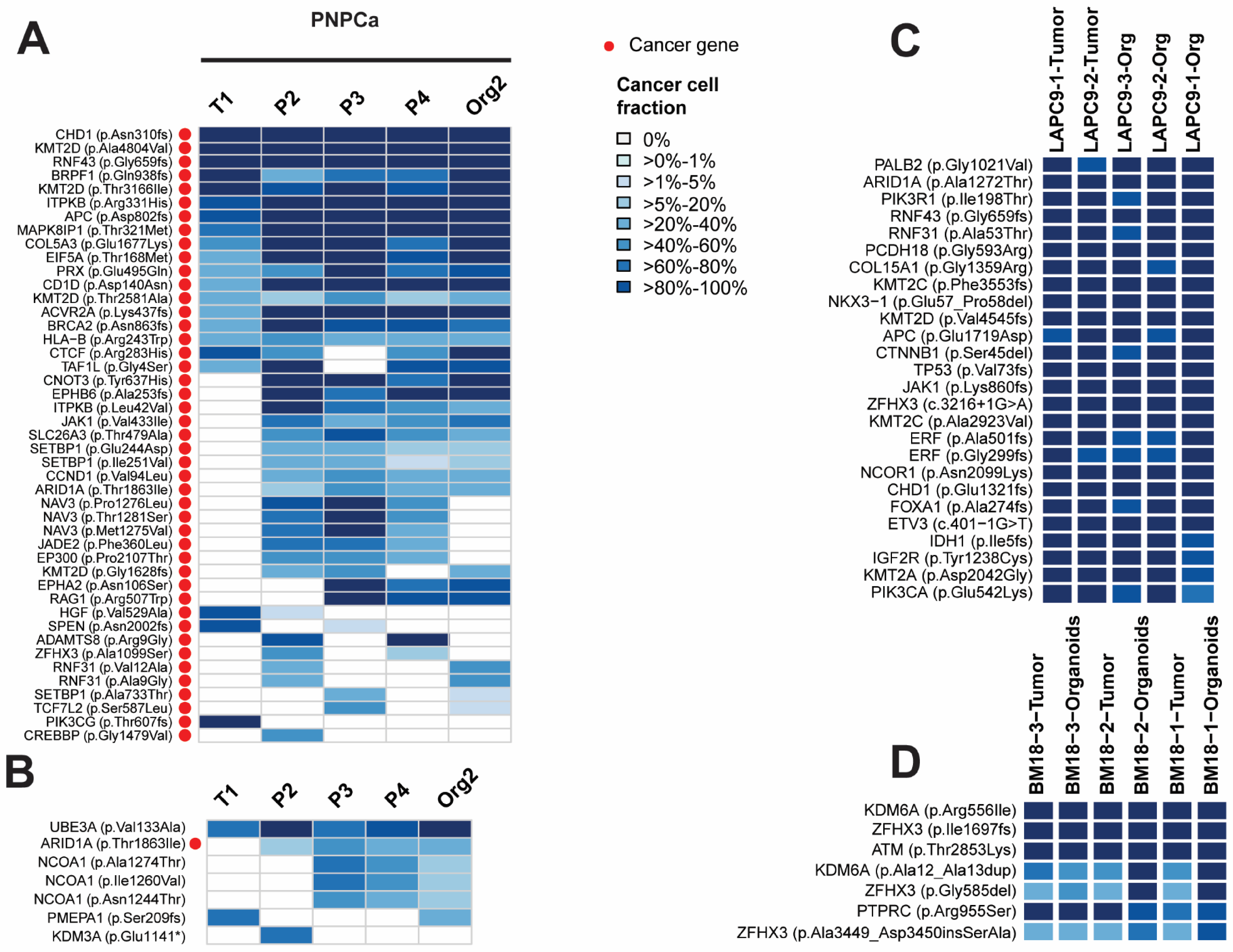
A
PNPCa
T1
P2
P3
P4
Org2
CHD1 (p.Asn310fs)
KMT2D (p.Ala4804Val)
RNF43 (p.Gly659fs)
BRPF1 (p.Gln938fs)
KMT2D (p.Thr3166Ile)
ITPKB (p.Arg331His)
APC (p.Asp802fs)
MAPK8IP1 (p.Thr321Met)
COL5A3 (p.Glu1677Lys)
EIF5A (p.Thr168Met)
PRX (p.Glu495Gln)
CD1D (p.Asp140Asn)
KMT2D (p.Thr2581Ala)
ACVR2A (p.Lys437fs)
BRCA2 (p.Asn863fs)
HLA−B (p.Arg243Trp)
CTCF (p.Arg283His)
TAF1L (p.Gly4Ser)
CNOT3 (p.Tyr637His)
EPHB6 (p.Ala253fs)
ITPKB (p.Leu42Val)
JAK1 (p.Val433Ile)
SLC26A3 (p.Thr479Ala)
SETBP1 (p.Glu244Asp)
SETBP1 (p.Ile251Val)
CCND1 (p.Val94Leu)
ARID1A (p.Thr1863Ile)
NAV3 (p.Pro1276Leu)
NAV3 (p.Thr1281Ser)
NAV3 (p.Met1275Val)
JADE2 (p.Phe360Leu)
EP300 (p.Pro2107Thr)
KMT2D (p.Gly1628fs)
EPHA2 (p.Asn106Ser)
RAG1 (p.Arg507Trp)
HGF (p.Val529Ala)
SPEN (p.Asn2002fs)
ADAMTS8 (p.Arg9Gly)
ZFHX3 (p.Ala1099Ser)
RNF31 (p.Val12Ala)
RNF31 (p.Ala9Gly)
SETBP1 (p.Ala733Thr)
TCF7L2 (p.Ser587Leu)
PIK3CG (p.Thr607fs)
CREBBP (p.Gly1479Val)
Cancer gene
Cancer cell fraction
0%
>0%-1%
>1%-5%
>5%-20%
>20%-40%
>40%-60%
>60%-80%
>80%-100%
B
T1
P2
P3
P4
Org2
UBE3A (p.Val133Ala)
ARID1A (p.Thr1863Ile)
NCOA1 (p.Ala1274Thr)
NCOA1 (p.Ile1260Val)
NCOA1 (p.Asn1244Thr)
PMEPA1 (p.Ser209fs)
KDM3A (p.Glu1141*)
C
LAPC9-1-Tumor
LAPC9-2-Tumor
LAPC9-3-Org
LAPC9-2-Org
LAPC9-1-Org
PALB2 (p.Gly1021Val)
ARID1A (p.Ala1272Thr)
PIK3R1 (p.Ile198Thr)
RNF43 (p.Gly659fs)
RNF31 (p.Ala53Thr)
PCDH18 (p.Gly593Arg)
COL15A1 (p.Gly1359Arg)
KMT2C (p.Phe3553fs)
NKX3−1 (p.Glu57_Pro58del)
KMT2D (p.Val4545fs)
APC (p.Glu1719Asp)
CTNNB1 (p.Ser45del)
TP53 (p.Val73fs)
JAK1 (p.Lys860fs)
ZFHX3 (c.3216+1G>A)
KMT2C (p.Ala2923Val)
ERF (p.Ala501fs)
ERF (p.Gly299fs)
NCOR1 (p.Asn2099Lys)
CHD1 (p.Glu1321fs)
FOXA1 (p.Ala274fs)
ETV3 (c.401−1G>T)
IDH1 (p.Ile5fs)
IGF2R (p.Tyr1238Cys)
KMT2A (p.Asp2042Gly)
PIK3CA (p.Glu542Lys)
D
BM18−3−Tumor
BM18−3−Organoids
BM18−2−Tumor
BM18−2−Organoids
BM18−1−Tumor
BM18−1−Organoids
KDM6A (p.Arg556Ile)
ZFHX3 (p.Ile1697fs)
ATM (p.Thr2853Lys)
KDM6A (p.Ala12_Ala13dup)
ZFHX3 (p.Gly585del)
PTPRC (p.Arg955Ser)
ZFHX3 (p.Ala3449_Asp3450insSerAla)

### Sup.Fig.8

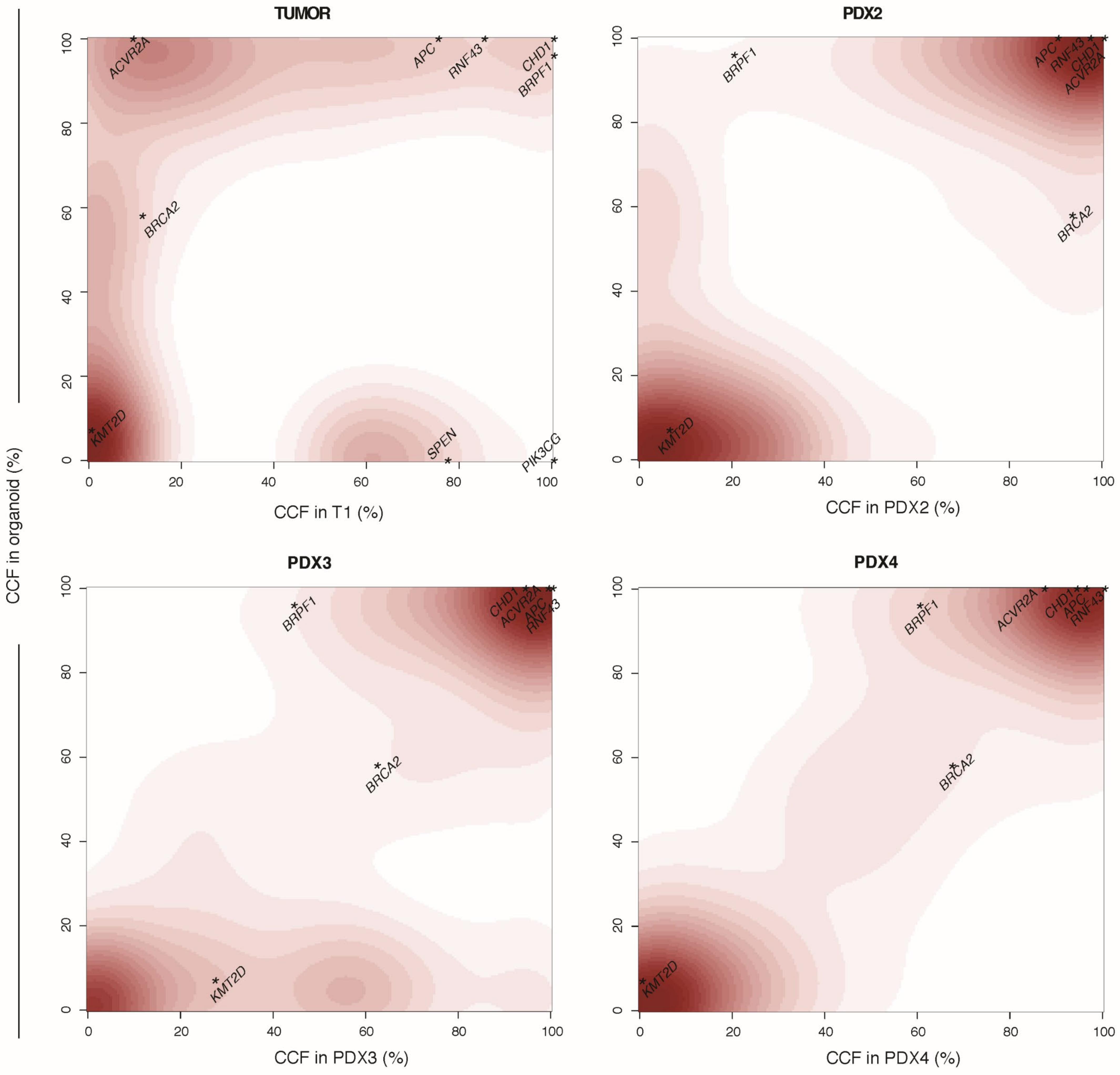
TUMOR
PDX2
PDX3
PDX4
CCF in organoid (%)
CCF in T1 (%)
CCF in PDX2 (%)
CCF in PDX3 (%)
CCF in PDX4 (%)
0
20
40
60
80
100
ACVR2A
APC
RNF43
CHD1
BRPF1
BRCA2
KMT2D
SPEN
PIK3CG

**Sup.Fig.9**

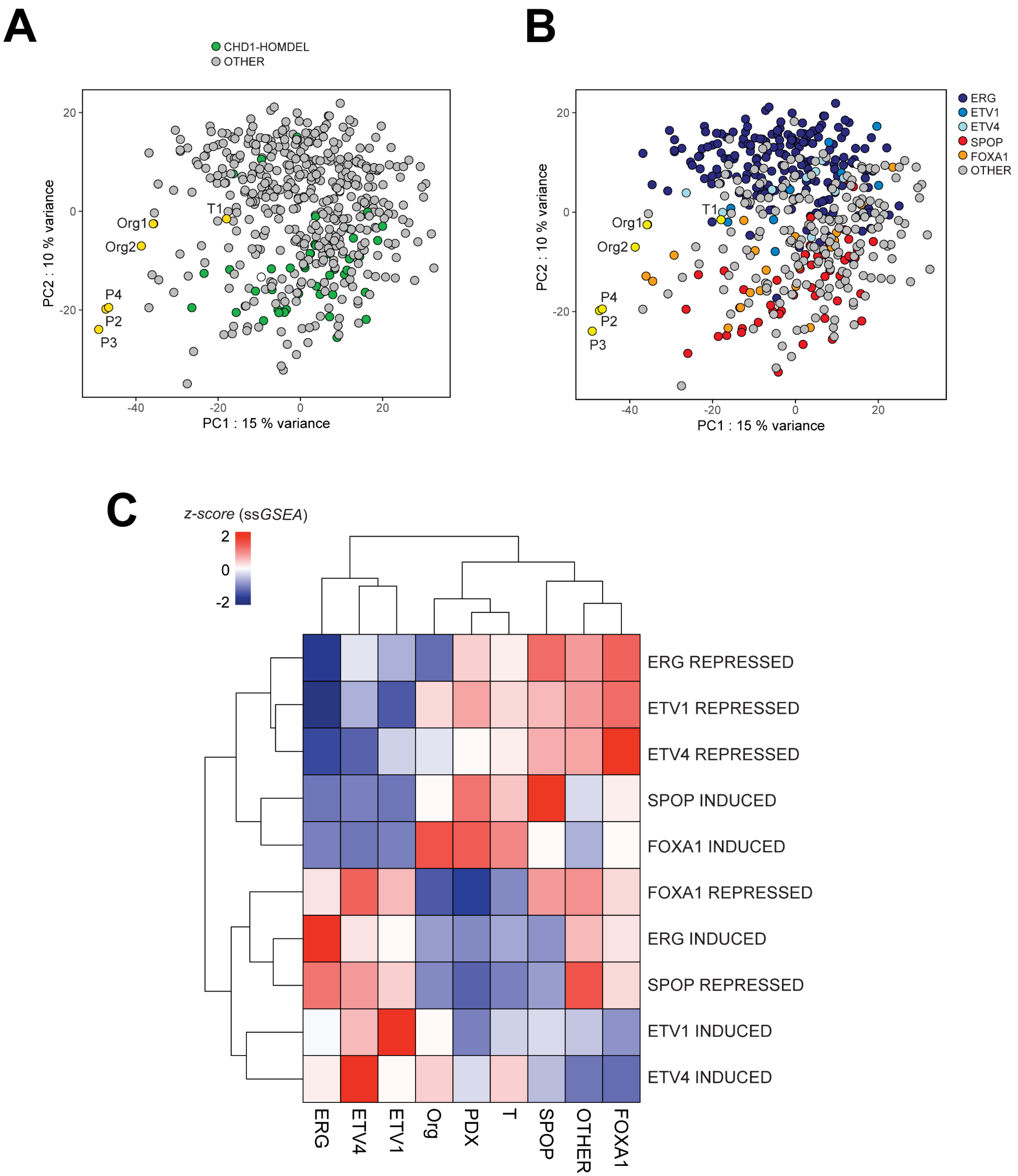

A
CHD1-HOMDEL
OTHER
PC2 : 10 % variance
PC1 : 15 % variance
Org1
Org2
T1
P4
P2
P3
B
ERG
ETV1
ETV4
SPOP
FOXA1
OTHER
C
z-score (ssGSEA)
2
0
-2
ERG REPRESSED
ETV1 REPRESSED
ETV4 REPRESSED
SPOP INDUCED
FOXA1 INDUCED
FOXA1 REPRESSED
ERG INDUCED
SPOP REPRESSED
ETV1 INDUCED
ETV4 INDUCED
ERG
ETV4
ETV1
Org
PDX
T
SPOP
OTHER
FOXA1

#### Sup.Fig.10

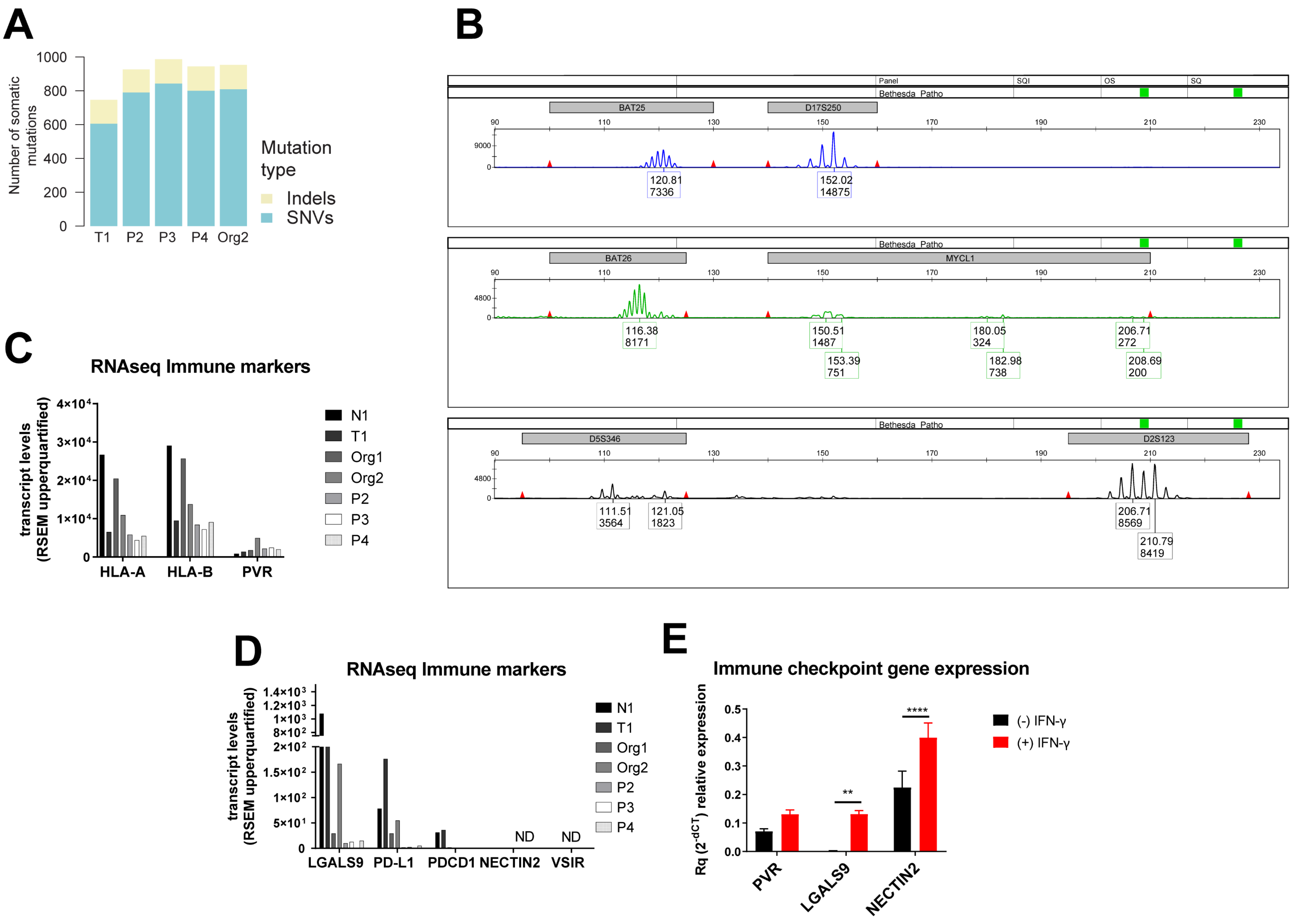

A
Number of somatic mutations
Mutation type
Indels
SNVs
T1
P2
P3
P4
Org2
B
Panel
SQI
OS
SQ
Bethesda_Patho
BAT25
D17S250
120.81
7336
152.02
14875
BAT26
MYCL1
116.38
8171
150.51
1487
153.39
751
180.05
324
182.98
738
206.71
272
208.69
200
D5S346
D2S123
111.51
3564
121.05
1823
206.71
8569
210.79
8419
C
RNAseq Immune markers
transcript levels (RSEM upperquartified)
HLA-A
HLA-B
PVR
N1
T1
Org1
Org2
P2
P3
P4
D
RNAseq Immune markers
transcript levels (RSEM upperquartified)
LGALS9
PD-L1
PDCD1
NECTIN2
VSIR
ND
ND
E
Immune checkpoint gene expression
Rq (2^-dCT) relative expression
PVR
LGALS9
NECTIN2
(-) IFN-γ
(+) IFN-γ

#### Sup.Fig.11

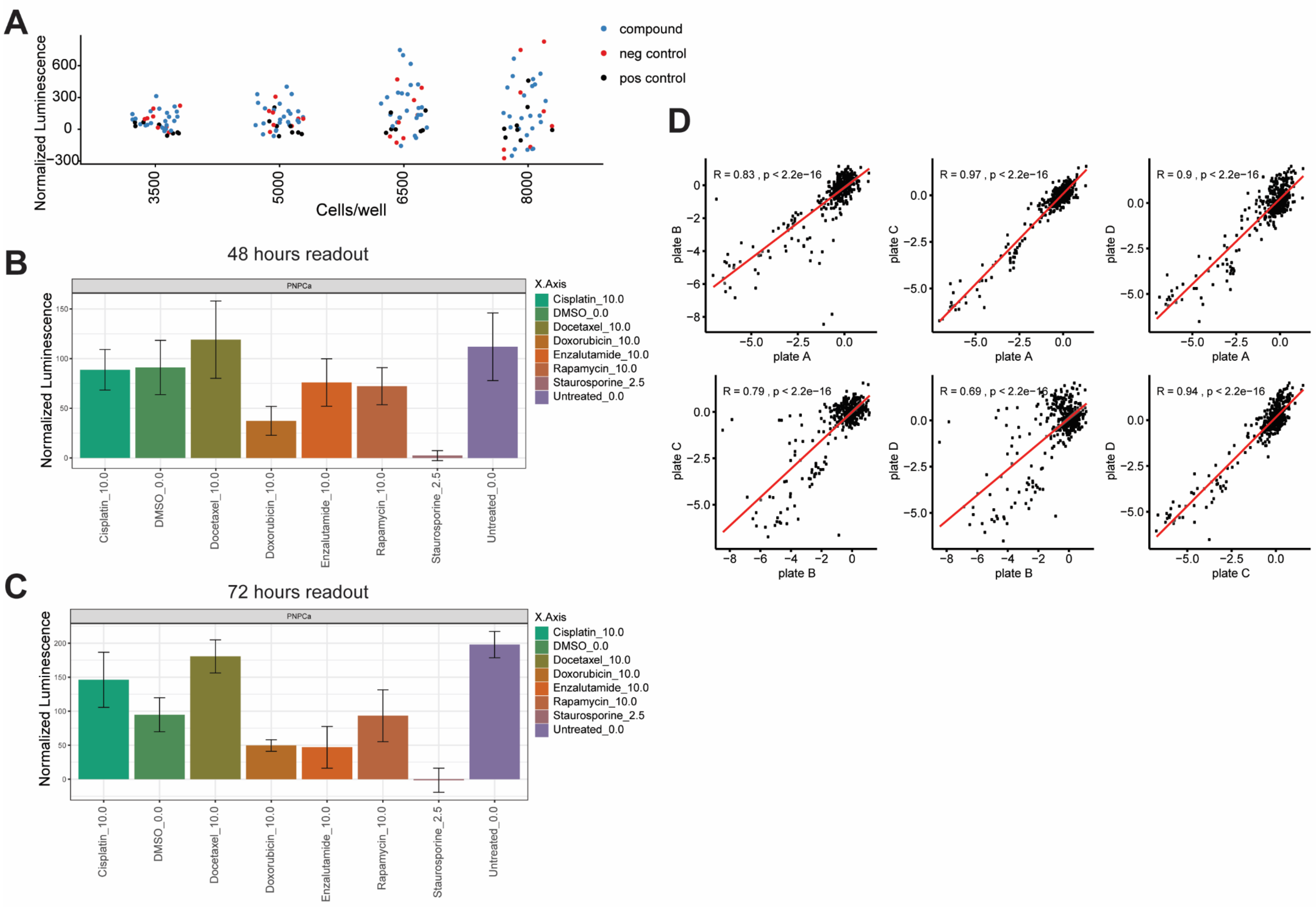

A
Normalized Luminescence
Cells/well
compound
neg control
pos control
B
48 hours readout
PNPCa
Normalized Luminescence
X.Axis
Cisplatin_10.0
DMSO_0.0
Docetaxel_10.0
Doxorubicin_10.0
Enzalutamide_10.0
Rapamycin_10.0
Staurosporine_2.5
Untreated_0.0
C
72 hours readout
PNPCa
Normalized Luminescence
X.Axis
Cisplatin_10.0
DMSO_0.0
Docetaxel_10.0
Doxorubicin_10.0
Enzalutamide_10.0
Rapamycin_10.0
Staurosporine_2.5
Untreated_0.0
D
R = 0.83 , p < 2.2e−16
R = 0.97 , p < 2.2e−16
R = 0.9 , p < 2.2e−16
R = 0.79 , p < 2.2e−16
R = 0.69 , p < 2.2e−16
R = 0.94 , p < 2.2e−16
plate A
plate B
plate C
plate D

### Sup.Fig.12

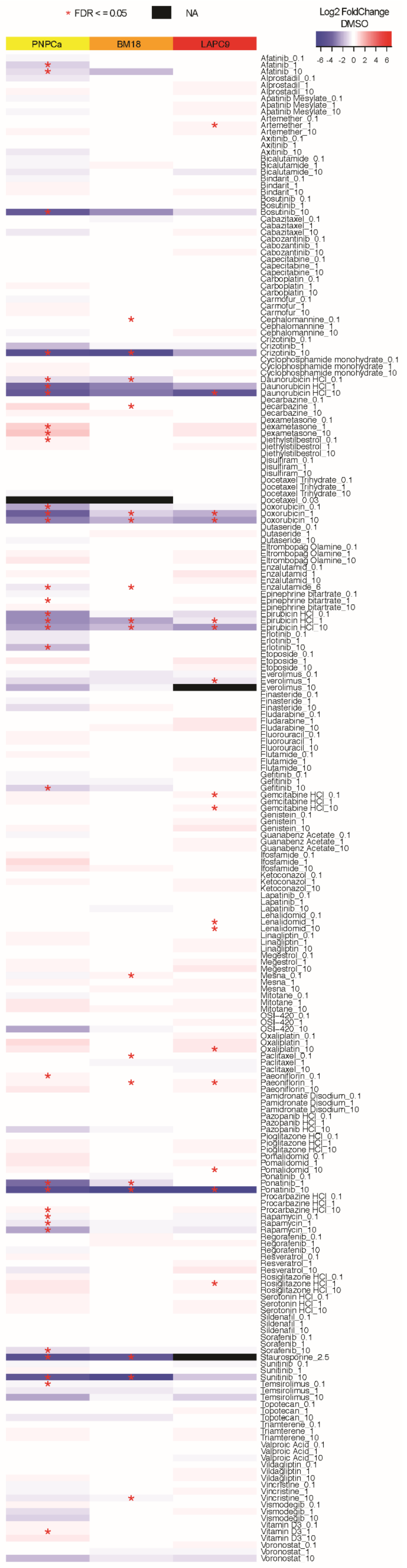

* FDR < = 0.05
NA
Log2 FoldChange
DMSO
-6 -4 -2 0 2 4 6
PNPCa
BM18
LAPC9
Afatinib_0.1
Afatinib_1
Afatinib_10
Alprostadil_0.1
Alprostadil_1
Alprostadil_10
Apatinib Mesylate_0.1
Apatinib Mesylate_1
Apatinib Mesylate_10
Artemether_0.1
Artemether_1
Artemether_10
Axitinib_0.1
Axitinib_1
Axitinib_10
Bicalutamide_0.1
Bicalutamide_1
Bicalutamide_10
Bindarit_0.1
Bindarit_1
Bindarit_10
Bosutinib_0.1
Bosutinib_1
Bosutinib_10
Cabazitaxel_0.1
Cabazitaxel_1
Cabazitaxel_10
Cabozantinib_0.1
Cabozantinib_1
Cabozantinib_10
Capecitabine_0.1
Capecitabine_1
Capecitabine_10
Carboplatin_0.1
Carboplatin_1
Carboplatin_10
Carmofur_0.1
Carmofur_1
Carmofur_10
Cephalomannine_0.1
Cephalomannine_1
Cephalomannine_10
Crizotinib_0.1
Crizotinib_1
Crizotinib_10
Cyclophosphamide monohydrate_0.1
Cyclophosphamide monohydrate_1
Cyclophosphamide monohydrate_10
Daunorubicin HCl_0.1
Daunorubicin HCl_1
Daunorubicin HCl_10
Decarbazine_0.1
Decarbazine_1
Decarbazine_10
Dexametasone_0.1
Dexametasone_1
Dexametasone_10
Diethylstilbestrol_0.1
Diethylstilbestrol_1
Diethylstilbestrol_10
Disulfiram_0.1
Disulfiram_1
Disulfiram_10
Docetaxel Trihydrate_0.1
Docetaxel Trihydrate_1
Docetaxel Trihydrate_10
Docetaxel_0.03
Doxorubicin_0.1
Doxorubicin_1
Doxorubicin_10
Dutaseride_0.1
Dutaseride_1
Dutaseride_10
Eltrombopag Olamine_0.1
Eltrombopag Olamine_1
Eltrombopag Olamine_10
Enzalutamid_0.1
Enzalutamid_1
Enzalutamid_10
Enzalutamide_6
Epinephrine bitartrate_0.1
Epinephrine bitartrate_1
Epinephrine bitartrate_10
Epirubicin HCl_0.1
Epirubicin HCl_1
Epirubicin HCl_10
Erlotinib_0.1
Erlotinib_1
Erlotinib_10
Etoposide_0.1
Etoposide_1
Etoposide_10
Everolimus_0.1
Everolimus_1
Everolimus_10
Finasteride_0.1
Finasteride_1
Finasteride_10
Fludarabine_0.1
Fludarabine_1
Fludarabine_10
Fluorouracil_0.1
Fluorouracil_1
Fluorouracil_10
Flutamide_0.1
Flutamide_1
Flutamide_10
Gefitinib_0.1
Gefitinib_1
Gefitinib_10
Gemcitabine HCl_0.1
Gemcitabine HCl_1
Gemcitabine HCl_10
Genistein_0.1
Genistein_1
Genistein_10
Guanabenz Acetate_0.1
Guanabenz Acetate_1
Guanabenz Acetate_10
Ifosfamide_0.1
Ifosfamide_1
Ifosfamide_10
Ketoconazol_0.1
Ketoconazol_1
Ketoconazol_10
Lapatinib_0.1
Lapatinib_1
Lapatinib_10
Lenalidomid_0.1
Lenalidomid_1
Lenalidomid_10
Linagliptin_0.1
Linagliptin_1
Linagliptin_10
Megestrol_0.1
Megestrol_1
Megestrol_10
Mesna_0.1
Mesna_1
Mesna_10
Mitotane_0.1
Mitotane_1
Mitotane_10
OSI−420_0.1
OSI−420_1
OSI−420_10
Oxaliplatin_0.1
Oxaliplatin_1
Oxaliplatin_10
Paclitaxel_0.1
Paclitaxel_1
Paclitaxel_10
Paeoniflorin_0.1
Paeoniflorin_1
Paeoniflorin_10
Pamidronate Disodium_0.1
Pamidronate Disodium_1
Pamidronate Disodium_10
Pazopanib HCl_0.1
Pazopanib HCl_1
Pazopanib HCl_10
Pioglitazone HCl_0.1
Pioglitazone HCl_1
Pioglitazone HCl_10
Pomalidomid_0.1
Pomalidomid_1
Pomalidomid_10
Ponatinib_0.1
Ponatinib_1
Ponatinib_10
Procarbazine HCl_0.1
Procarbazine HCl_1
Procarbazine HCl_10
Rapamycin_0.1
Rapamycin_1
Rapamycin_10
Regorafenib_0.1
Regorafenib_1
Regorafenib_10
Resveratrol_0.1
Resveratrol_1
Resveratrol_10
Rosiglitazone HCl_0.1
Rosiglitazone HCl_1
Rosiglitazone HCl_10
Serotonin HCl_0.1
Serotonin HCl_1
Serotonin HCl_10
Sildenafil_0.1
Sildenafil_1
Sildenafil_10
Sorafenib_0.1
Sorafenib_1
Sorafenib_10
Staurosporine_2.5
Sunitinib_0.1
Sunitinib_1
Sunitinib_10
Temsirolimus_0.1
Temsirolimus_1
Temsirolimus_10
Topotecan_0.1
Topotecan_1
Topotecan_10
Triamterene_0.1
Triamterene_1
Triamterene_10
Valproic Acid_0.1
Valproic Acid_1
Valproic Acid_10
Vildagliptin_0.1
Vildagliptin_1
Vildagliptin_10
Vincristine_0.1
Vincristine_1
Vincristine_10
Vismodegib_0.1
Vismodegib_1
Vismodegib_10
Vitamin D3_0.1
Vitamin D3_1
Vitamin D3_10
Voronostat_0.1
Voronostat_1
Voronostat_10

#### Sup.Fig.13

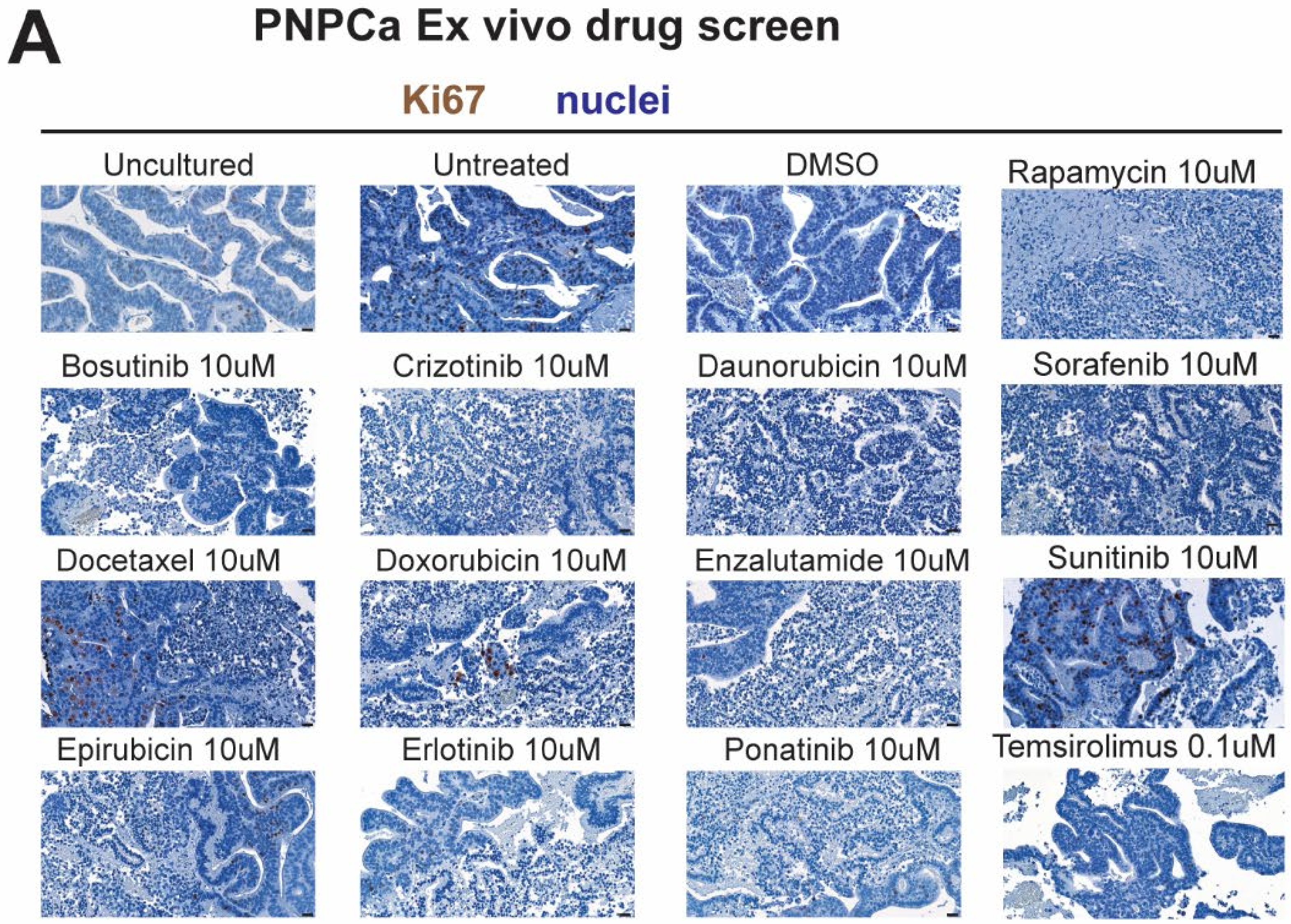

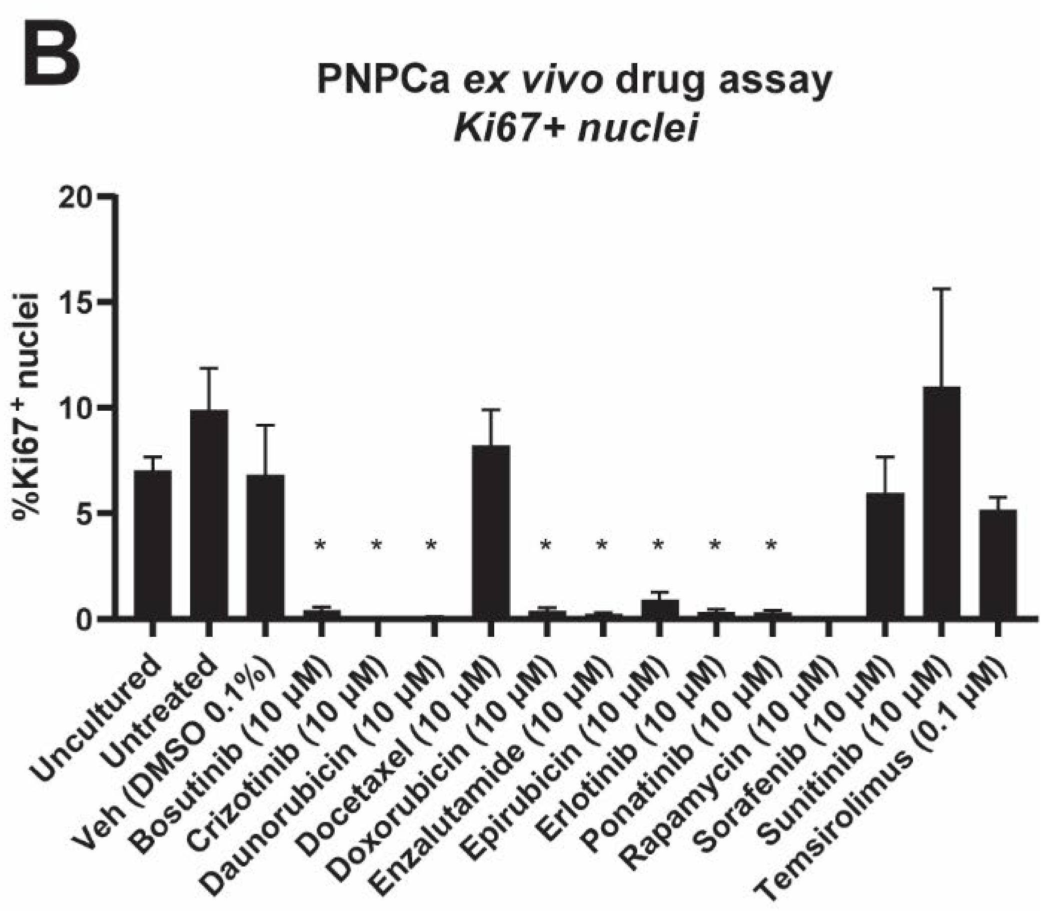

**C**

| Compound | LAPC9 | | | BM18 | | | PNPCa | | |
|---|---|---|---|---|---|---|---|---|---|
| | Dose | Ki67+ over total nuclei | q-value | Dose | Ki67+ over total nuclei | q-value | Dose | Ki67+ over total nuclei | q-value |
| Bosutinib | 10 µM | 0.3% | <0.0001 | | | | 10 µM | 0.4% | 0.049 |
| Crizotinib | 10 µM | 1.5% | 0.0283 | | | | 10 µM | 0.0% | 0.019 |
| Daunorubicin | 10 µM | 2.4% | 0.2335 | 1 µM | 5.1% | 0.0854 | 10 µM | 0.1% | 0.032 |
| Docetaxel | | | | | | | 10 µM | 8.2% | 0.999 |
| Doxorubicin | 1 µM | 0.0% | <0.0001 | 10 µM | 5.0% | 0.0317 | 10 µM | 0.4% | 0.023 |
| Enzalutamide | | | | 6 µM | 1.4% | 0.0007 | 10 µM | 0.2% | 0.026 |
| Epirubicin | 1 µM | 0.0% | <0.0001 | | | | 10 µM | 0.9% | 0.047 |
| Erlotinib | | | | 10 µM | 10.7% | 0.107 | 10 µM | 0.3% | 0.029 |
| Everolimus | 1 µM | 12.1% | 0.0283 | | | | | | |
| Paclitaxel | 1 µM | 1.7% | 0.2335 | | | | | | |
| Ponatinib | 10 µM | 0.5% | 0.0002 | | | | 10 µM | 0.3% | 0.021 |
| Rapamycin | 10 µM | 7.4% | 0.1726 | 10 µM | 10.5% | 0.1825 | 10 µM | 0.0% | 0.052 |
| Sorafenib | | | | | | | 10 µM | 4.8% | 0.999 |
| Sunitinib | 10 µM | 0.2% | <0.0001 | | | | 10 µM | 10.3% | 0.440 |
| Temsirolimus | 10 µM | 9.2% | 0.0707 | | | | 0.1 µM | 5.1% | 0.999 |

**Sup.Fig.14**

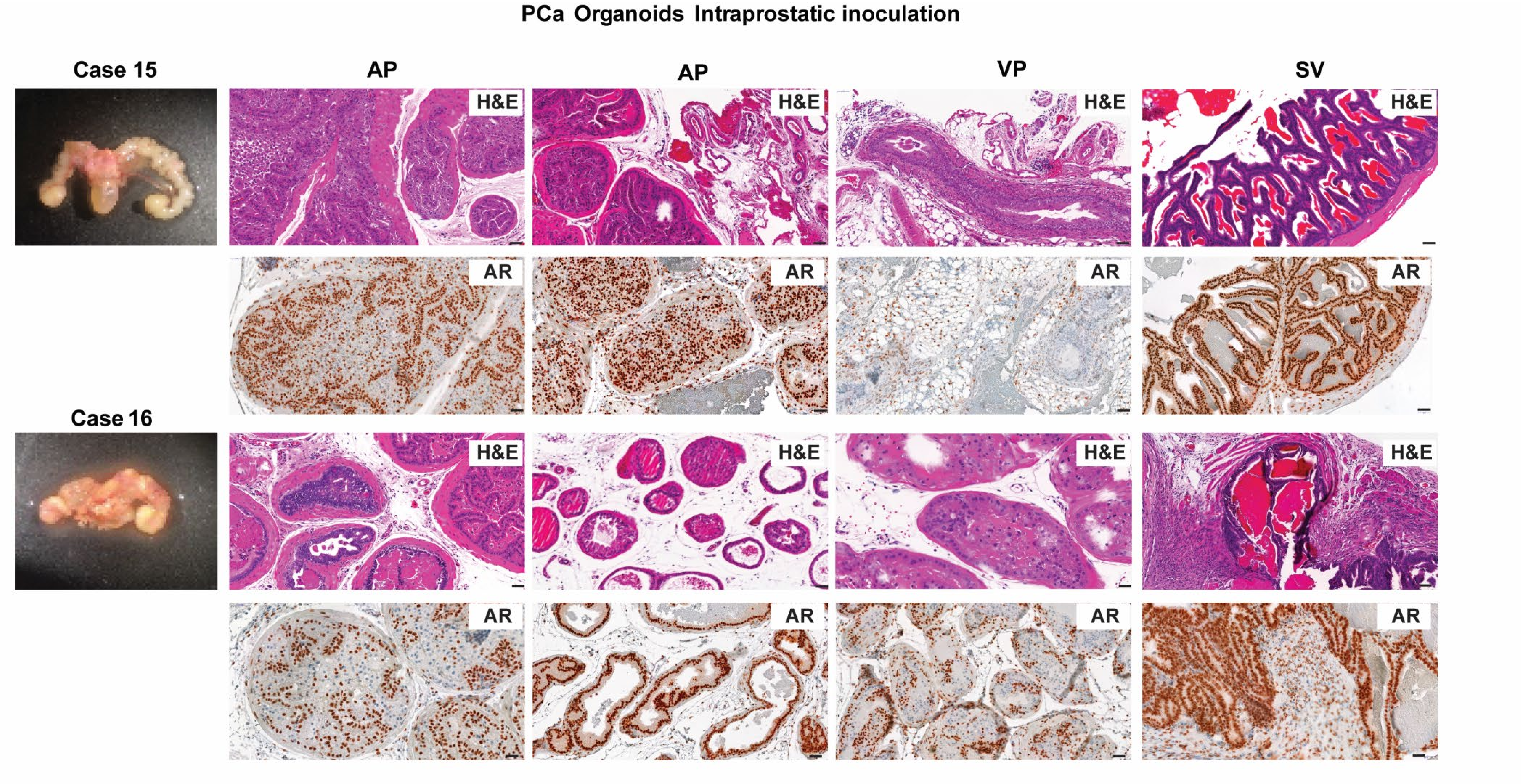

PCa Organoids Intraprostatic inoculation
Case 15
AP
AP
VP
SV
H&E
AR
Case 16

**Sup.Fig.15**

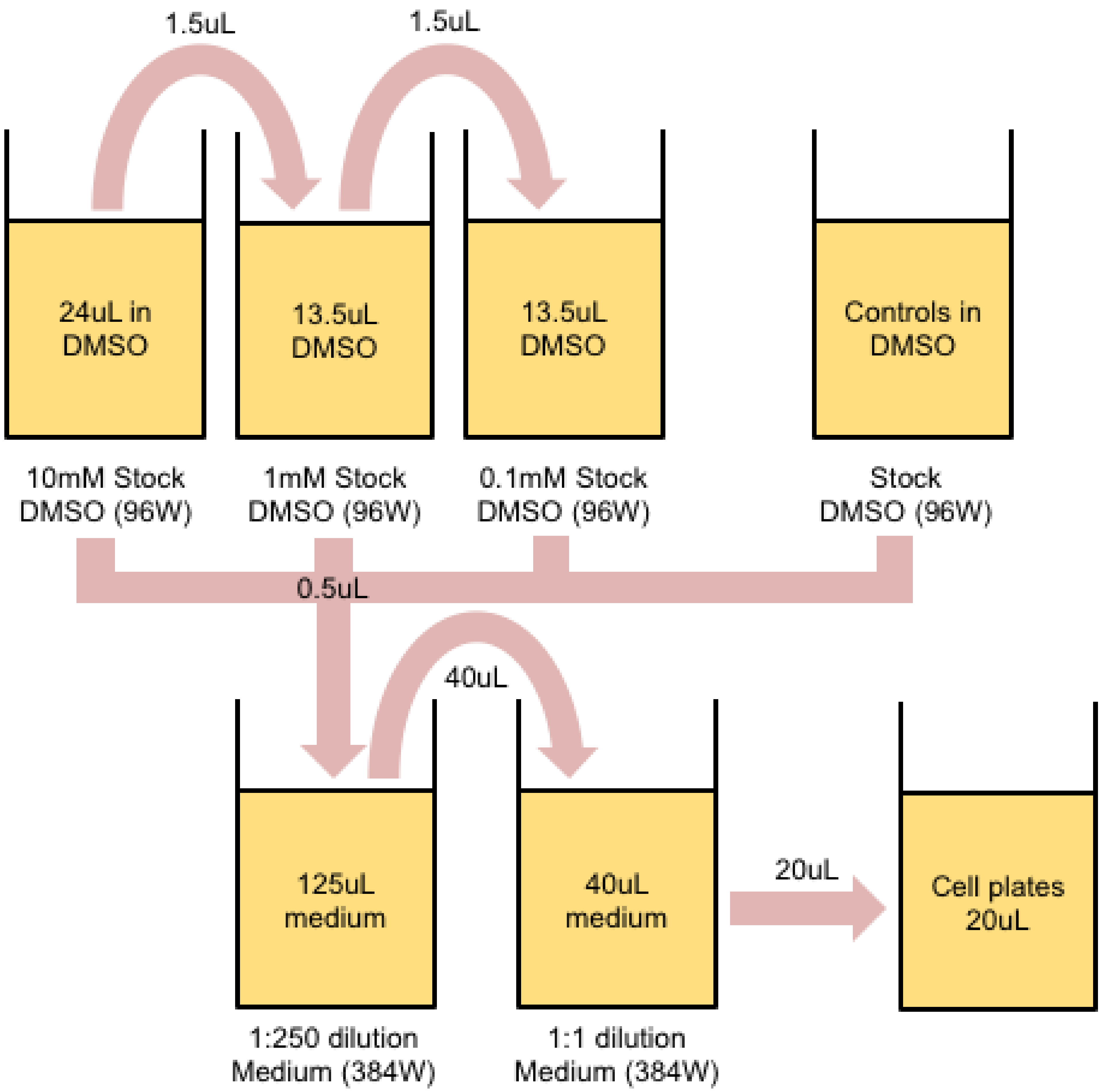
1.5uL
1.5uL
24uL in DMSO
13.5uL DMSO
13.5uL DMSO
Controls in DMSO
10mM Stock DMSO (96W)
1mM Stock DMSO (96W)
0.1mM Stock DMSO (96W)
Stock DMSO (96W)
0.5uL
40uL
125uL medium
40uL medium
20uL
Cell plates 20uL
1:250 dilution Medium (384W)
1:1 dilution Medium (384W)

#### Supplementary Figure Legends

**Sup. Figure 1. Maintenance of luminal epithelial morphology of the PDX and tumor growth kinetic in different genetic backgrounds, related to Fig.1**
**A.** Histological morphology of PNPCa PDX passages (PDX2-PDX5) as assessed by Hematoxylin and Eosin staining (H&E). Scale bars 20um. **B.** PSA protein expression. Scale bars 20um. **C.** Expression of AR (green), CK5 (red) assessed by immunofluorescence, DAPI (blue) marks the nuclei. Scale bars 50um. **D.** Expression of NKX3.1 (green), CK8 (red) assessed by immunofluorescence. Scale bars 50um. **E.** Duration of tumor take (days between subcutaneous implantation and tumor growth of ~1cm$^3$ in immunocompromised strains NOD-*scid* IL2Rgamma$^{null}$ (NSG). Testosterone supplementation was performed weekly to ensure grafting success with weekly testosterone injections. Starting from passage 7, the PDX tumor engrafted in less immunocompromised strain CB17 SCID with testosterone supplementation.

**Sup. Figure 2. Micrometastasis detection in PNPCa PDX, related to Fig.1**
**A.** Liver, lung and prostate tissues from NSG mice (N=3) with PDX subcutaneous tumors (Castrated-Test group). The duration of the time following castration was a total of 40 weeks. During that period, no spontaneous regrowth of tumor was observed, until supplemented with testosterone for seven weeks; endpoint collection of organs. **B.** Human panCK staining on subcutaneous PDX tumor (following testosterone), liver, lung, prostate (anterior lobe), lymph node. **C.** Human panCK staining on femur-tibia bones and **D.** X-ray of the left and right tibia of each mouse.

**Sup. Figure 3**. **RNAseq confirms testosterone readministration following castration reverses the transcriptomic changes of castration, and is similar to an intact tumor, related to Fig.1**
Heat-map of non-hierarchical clustering of the top 500 variable genes across all samples indicated ("Castrated-Test" group (N=3), all other treatment groups (N=2)).

**Sup. Figure 4. Histological morphology of PNPCa PDX and organoids, related to Fig.2**
Hematoxylin and Eosin staining of the samples used for exome sequencing; patient-derived material (primary tumor TUR-P ("T1") and non-carcinoma control "N1"), PDX passages from PNPCa met (p2, p3, p4) and PDX-derived organoids (Org2). Scale bars 50um.

**Sup. Figure 5. *In vivo* tumorigenicity of BM18 and LAPC9 organoids, related to Fig. 2**
**A.** Tumor growth kinetic of BM18 subcutaneous PDX tumors in response to castration and testosterone (DHT). **B.** Tumor growth kinetic of LAPC9 subcutaneous PDX tumors in response to castration and testosterone (DHT).
**C-H.** Subrenal implantation of LAPC9 organoids **(C-E)** and of BM18 organoids **(F-H)**; H&E staining (C,F), AR staining (D,G) and PSA staining (E,H) of the subrenal tumor growth. **I-N.** AR staining of control prostatic tissues (Anterior, Ventral Prostate and Seminal Vesicles) **(I-K)** and prostatic tissues from intraprostatic inoculation of BM18 organoids in the anterior prostate **(L-N)**.

**Sup.Figure 6. Genomic profile of PNPCa reveals high mutational load due to high microsatellite instability and BRCA2 mutation, related to Fig.2**
**A.** Genome-wide copy number profiles of genetic rearrangements and **B.** Venn diagram of the number of somatic mutations from whole exome sequencing of patient-derived material

(primary tumor TUR-P ("T1") and non-carcinoma control "N1"), PDX passages from PNPCa met (P2, P3, P4) and PDX (P4)-derived organoids (Org2).

**Sup. Figure 7. Somatic mutations in identified cancer genes in PCa PDX and PDX-derived organoids, related to Fig.2**

**A.** Heatmap of cancer cell fraction percentage of all identified somatic mutations in PNPCa, in known cancer genes; amino acid position of the mutation is indicated next to the gene symbol in parentheses. **B.** Heatmap indicating cancer cell fraction of all somatic mutations in genes of the AR signaling pathway. Cancer genes are indicated (red dot) in the PNPCa. **C.** Cancer cell fraction of mutated cancer genes of LAPC9 PDX tissue and organoids and **D.** of BM18 PDX and tissue and organoids.

**Sup. Figure 8. Density correlation plots of cancer cell fraction, related to Fig.2**

Correlation plots of cancer cell fraction (CCF, %) in PNPCa model; organoid Org2 sample (y axis), versus P2 (PDX2), T1 (Tumor), P3 (PDX3) and P4 (PDX4).

**Sup. Figure 9. Transcriptomic landscape of PNPCa is conserved among the PDX and organoids, with AR pathway enrichment, SPOP, FOXA1 and CHD1-like signatures, related to Fig.2**

**A.** Principal component analysis (PCA) of TCGA gene expression data from 480 primary PCa tumors, classified based on CHD1 homozygous deletion and **B.** genetic subtype (SPOP, FOXA1, ETS rearrangements; ERG, ETV1, ETV4). **C.** Z score of single sample gene set enrichment expression (ssGSEA) analysis among gene signature from the different genetic subgroups. N1, normal tissue; T1, primary tumor; P2-4, PNPCa PDX passage 2-4; Org1-2, PNPCa PDX-derived organoids passage 1-2. Others; cases with no mutations in SPOP, FOXA1, CHD1 and ETS groups. FOXA1 induced genes (n = 109), FOXA1 repressed genes (n = 183), ERG induced genes (n = 178), ERG repressed genes (n = 291), ETV1 induced genes (n = 9), ETV1 repressed genes (n = 25), ETV4 induced genes (n = 23), ETV1 repressed genes (n = 28).

**Sup. Figure 10. Confirmation of Microsatellite Instability of the primary tumor (T1), and immune marker expression, related to Fig.3**

**A.** Number of somatic mutations divided into single nucleotide variants (SNVs) and insertions-deletions (indels) of the PNPCa models. **B.** MSI testing based on the Bethesda panel, which consists of six loci BAT25, BAT26, MYCL1, D2S123, D5S346, and D17S250. Four out of the six loci contain repeats, classifying the tumor as MSI-high. Genomic DNA from the primary T1 tumor was obtained from high carcinoma- containing FFPE cores, as identified at pathological evaluation. **C,D.** Gene expression of markers related to immunosuppression (RNASeq data, RSEM transcript levels). **E.** Gene expression levels of immune markers based on RT-qPCR results on PNPCa organoids RNA after 48 hours treatment with IFN-γ.

**Sup. Figure 11. Optimization of Nexus pipeline for PNPCa PDX-derived organoids related to Fig. 4**

**A.** Luciferase values (viability assay ATP based, Cell Titer Glo 3D) on different cell densities of PNPCa organoids after 48 hours of seeding. **B-C**. Prescreen viability assays in response to standard-of-care compounds for assessment of positive controls and timing of drug exposure; 48 hours **(B)** and 72 hours **(C)**. **D.** Correlation plots of the log2 values between replicates obtained by luciferase measurements, proportional to cell viability, after 72h drug treatment.

**Sup. Figure 12. PNPCa, BM18 and LAPC9 organoid drug screen, related to Fig. 4**
Heatmap of viability values of all tested compounds in the NEXUS drug screen on PNPCa organoids (N=4) and BM18/LAPC9 organoids (N=3). Log2 Fold Change values relative to the DMSO control of each organoid model. Black segments, data not available.

**Sup. Figure 13. Drug compound validation in *ex vivo* tissue slice assay for proliferation effects, related to Fig.4**
**A.** Expression of proliferation marker Ki67 assessed by immunohistochemistry on PNPCa PDX tissue cultured *ex vivo* with selected drug compounds, previously used in the NEXUS organoid screen. Hematoxylin stains the nuclei. Scale bars 50um. **B.** Quantification of Ki67-positive cells. Different fields of view (min. 5 per condition) were used to reconstruct the section area (average), and the counts of positive nuclei was normalised over the total nuclei (%Ki67+ nuclei) per condition were calculated. Scale bars 50um. **C.** Table of tested and validated compounds in each PDX model. Percentage of Ki67+ cells and q value are represented.

**Sup. Figure 14. *In vivo* tumorigenicity of patient derived organoids, related to Fig. 5**
Representative images of H&E and AR staining of prostatic tissues from intraprostatic inoculation of primary PCa organoids (two representative cases). AP; anterior prostate, VP; ventral prostate, SV, seminal vesicles.

**Sup. Figure 15. Automated drug dilution and addition to target plates, related to Materials and Methods section**
Dilution steps of drug compounds from 96well plates to 384well plates performed at NEXUS Personalized Health Technologies.
