## Supplementary Material and methods for "Patient-derived xenografts and organoids model therapy response in prostate cancer"

**Organoid Maintenance, Passaging, Freezing and Thawing**

For first passage after isolation, organoids start forming within 2-3 days- up to 1 week and the media is changed after the first 2-4 days, depending on organoid density. Twice weekly organoids must be either refreshed/ split or media must be added. Eventually some cells, such as fibroblasts, will adhere to the plate bottom while the organoids remain in suspension and are collected by simply pipetting the supernatant into basis media. For passaging organoids when density is high or their size is >150um, the organoid suspension is collected in 15-ml Falcon tube, washed in basis medium in 220g, 3 min. If many floating non-viable single cells are present in the culture: centrifuge at low speed (10rcf) for 5 min, remove supernatant and wash with 4 ml Basis medium and centrifuge to pellet cells. Organoids are then incubated with TryPLE at 37°C for minimum 5 min to obtain single cells. When absolute single cell suspension is required for seeding exact No of cells use 22- or 23-gauge needle attached to a 1 mL syringe. Divide cell suspension in 2 tubes (expansion vs freezing). Spin down at 220rcf, 3 min, RT. After counting cells are seeded in 1:4 or 1:8 (depending on density) in fresh organoid media containing 10µM Y-27632-HCl inhibitor in ULA plates.

For cryopreservation, organoids are washed in basis media (220rcf, 3 min) and dissociated in TrypLE as mentioned before, resuspend in organoid freezing media (50% fetal calf serum, 40% Advanced F12/DMEM basis medium, 10%DMSO and aliquoted in cryovials (approximately 10E5 cells per vial).

**Immunofluorescence & Immunohistochemistry**

FFPE sections (4um) were deparaffinised and used for heat mediated antigen retrieval (citrate buffer pH 6, Vector labs). Sections were blocked for 30min, RT in 1%BSA in PBS-0.1%Tween20. The following antibodies were used:

| Dilution | Antibody | Company | Catalog No |
|---|---|---|---|
| 1to500 | **CK5/6** | Chemicon Milipore | MAB1620 |
| 1to500 | **AR** | abcam | ab133273 |
| 1to200 | **CK8** | Thermo Fisher | MA1-06318 |
| 1to250 | **Nkx3.1** | AthenaES | 314 |
| 1to500 | **Synaptophysin** | Invitrogen | 18-0130 |
| 1to1200 | **p63** | Abcam | ab124762 |

| | | | |
|---|---|---|---|
| 1to500 | **p63** | BD Pharmingen | 559951 |
| 1to400 | **Ki67** | Gene Tex | GTX16667 |
| 1to500 | **CD44** | BD Pharmingen | 550988 |
| 1to500 | **ALDH1A1** | LabForce/LSBio | LS-B2497/66673 |
| 1to250 | **panCK** | DAKO | M0821 |
| 1to500 | **PCNA** | Sigma Aldrich | P8825 |
| 1to500 | **cleaved caspase 3 (Asp175)** | Cell signaling | 9661 |
| 1to400 | **PD-L1** | Cell signaling, E1E3N clone | |
| 1to1000 | **PSA** | DAKO | A0562 |

For Immunohistochemistry, antigen retrieval was performed for 10 min in citrate buffer, followed by blocking of endogenous peroxidases in $H_2O_2$-$NaN_3$ and with swine serum for 15min. Primary antibody PSA was diluted 1to 3000 in 5%swine serum in DAKO antibody diluent. Secondary anti-rabbit antibody Envision HRP (DAKO) for 30min. Signal detection with AEC substrate (DAKO).

**Whole mount immunofluorescence staining of organoids**

Organoids in suspension were collected by pipetting P1000 into 15ml Falcon tube and washed with 250ul of 1xPBS (50rcf,3min). Following short fixation in 2% paraformaldehyde PFA at RT for 20 minutes, cells were spun down in 15ml falcon tube 50rcf, 3min. Organoids were washed 3x with 1xPBS/Glycine solution for 10 minutes each, gently rocking (2 rpm), spin 50rcf for 3 min to remove the supernatant. Buffer preparation 10x PBS/Glycine: 38.0g NaCl, 9.38g $Na_2HPO_4$ (sodium phosphate dibasic anhydrous), 2.07g $NaH_2PO_4$ (sodium phosphate monobasic), 37.5g glycine in 500ml total volume, adjust pH to 7.4 and filter sterilize. Organoids are then washed 3x with 1xIF Wash solution for 10 minutes each, gently rocking. Buffer preparation 10xIF wash: 38.0g NaCl, 9.38g $Na_2HPO_4$ (sodium phosphate dibasic anhydrous), 2.07g $NaH_2PO_4$ (sodium phosphate monobasic), 2.5g $NaN_3$, 5.0g Bovine Serum A (Fraction V), 10ml Triton X-100, 2.5ml Tween-20 in 500ml total volume, adjust pH to 7.4 and filter sterilize. Blocking is done in 1xIF wash buffer + 10% swine serum for 1 hour at RT. Aspirate block and incubate in primary antibody diluted in blocking buffer (300ul) in 1.5ml Eppendorf tube for overnight, gently rocking. For the washes IF wash is added and organoids were transferred to 15ml falcon tube for the washes.

Three washes with 1xIF Wash solution for 10 minutes each, gently rocking at RT. Spin 50rcf, 3min. Organoids were incubated with secondary antibodies AlexaFluor donkey anti-rabbit or anti-mouse (Invitrogen 1:250 dilution) in block buffer (1xIF wash + 10% swine serum) for 2 hours, gently rocking at RT. Two washes with IF Wash solution for 20 minutes each, gently rocking at RT. Incubate with DAPI (1ug/ml) in 1xPBS for 10 minutes, gently rocking at RT. Wash and resuspend in ~100ul PBS/IF wash. For imaging, organoids are transferred in 8well chamber slides (Nunc Labtek II Chamber #1.5 Coverglass system 155409). Antibodies used for whole mount staining are seen in table 1 (dilution 1:100).

**Viability assay Organoids-Cell Titer Glo 3D assay**

Organoids are dissociated in TrypLE, and seeded as single cells in ULA 96well plates (5000 cells in 100ul media, minimum 4 replicates per condition). After 1-3 days depending on the reformation of organoids, measure the volume of media left (after 48hours, remaining volume is 75ul). Organoid media is prepared containing the drug compounds (2x of final concentration). Add 75ul media plus drug treatments on organoids and incubated for 48hrs. Organoid suspension is transferred (100ul) into opaque 96-well plates for luciferase measurement (Thermo Fisher Nuna, 0.5ml capacity, 267350). Equal volume of Cell Titer 3D Glo (Promega) is added, using a plate reader, the plate is subjected for 5min orbital shaking RT and proceed with the guidelines of the assay. Luminescent signal is measured at Tecan plate Reader Infinite Pro 2000.

**Mixed Leukocyte Reaction (MLR) and regulatory T cells (T-reg) assay**

*Reagents and media*

Buffy coat-derived cells were cultured in complete RPMI medium consisting of RPMI-1640 (Sigma-Aldrich, Germany) supplemented with 10% heat-inactivated FBS (Thermo Fisher Scientific, Switzerland), 1% Glutamax supplement and 1% penicillin/streptomycin (both by Thermo Fisher Scientific). DC differentiation medium was prepared by adding 100 ng/mL granulocyte-macrophage colony-stimulation factor (GM-CSF, Miltenyi Biotec) and 100 ng/mL IL-4 (Peprotech Ltd, UK) to complete RPMI medium. MLR medium was prepared by adding to complete RPMI medium 1% ITS supplement (Thermo Fisher Scientific), 100 ug/ml R-spondin 1, 50ng/mL EGF, 10 ng/mL FGF-10, 10 ng/mL Wnt-3a and 1ng/mL FGF-2. FACS wash was prepared with 0.5% low endotoxin BSA (Sera Laboratories International, UK), 2mM EDTA (Sigma-Aldrich) in PBS pH 7.4.

*Cell Isolation and DC Generation*

Buffy coats were obtained from healthy donors and were used to isolate mononuclear cells (MNC) by gradient centrifugation (Lymphoprep; 1.077 g/mL; Axis-Shield, UK). After separation, CD14+ monocytes and CD3+ cells were purified from total MNC by magnetic separation columns (Miltenyi Biotec, Germany), according to manufacturer's instructions. Monocyte-derived DCs (moDCs) were generated by culturing CD14+ cells in DC differentiation medium for 5 days at 37°C in 5% CO2. After differentiation, moDC were maturated by incubation for 2 days in DC differentiation medium supplemented with IL-6 (100 ng/mL, Peprotech), TNFα (25 ng/mL, Miltenyi Biotech), IL-1β (30 ng/mL, Miltenyi Biotech), and 1 µg/mL PGE2 (Tocris).

*MLR and T-reg assay:* CD3+ cells were labelled with CellTrace Violet (Thermo Fisher Scientific) according to manufacturer's instructions at a final concentration of 5 uM and plated at 100.000 cells/well in 96-well ultra-low attachment plates (Corning). As a positive control CD3+ cells were cocultured with mature DC (1:10 to CD3+ cells) and, where indicated, PNPCa organoids-derived cells were added at a ratio of 1:2 to CD3+ cells; all cells were cultured in 200 ul MLR medium per well. Plate was protected from light and incubated for 5 days at 37°C in 5% CO2. As a negative control, stained and unstained CD3+ cells were cultured alone in MLR medium. For MLR assays, cells were collected at day 5, washed once in FACS wash and directly analysed by flowcytometry. For T-reg assay, cells were collected at day 5, washed in FACS wash and then stained at room temperature, protected from light, for 20 minutes with the following antibodies: APC-Violet 770-labeled anti-CD4, PE-Violet 770-labeled anti-CD127, PE-labeled anti CD25 (from Miltenyi Biotec), Brilliant Ultraviole 395-labeled anti-CD3, FITC-labeled anti PD-1, PerCP-Cy 5.5-labeled anti-CD8a (from BD). Cells were washed twice and then stained for FoxP3 with an APC-labeled anti-FoxP3 antibody (clone PCH101) using the FoxP3 Staining buffer Set (both from Thermo Fisher Scientific) and following manufacturer's instructions. For both MLR and T-reg assay, cells were analysed by flowcytometry with an LSR-II instrument (Becton Dickinson, BD, USA). Flowcytometry data were analysed with FlowJo v 10.5 for Windows.

**Whole exome sequencing**

DNA extracted from FFPE (original patient material), frozen tissue (PDXs) or organoids (300ng) for the PNPCa model were sequenced using whole-exome sequencing. Whole-exome capture libraries were constructed after sample-shearing, end repair, and phosphorylation and ligation to barcoded sequencing adaptors. Ligated DNA was subjected to HaloPlex Exome (Agilent) as previously described [1]. Sequencing was performed using Illumina HiSeq 2500 (2 × 100 bp).

Sample preparation and hybridization capture for BM18, LAPC9 were based on the SureSelectXT Low Input Automated Target Enrichment for Illumina Paired-End Mulitplexed Sequencing Version A0 (G9703-90010) using the Bravo Liquid Handling system. The Agilent SureSelectXT human all exon v7 capture library (5191-4006) was used. Given the lack of matched germline control, we sequenced additional whole blood DNA samples from eight individuals by the same workflow to serve as control. Clustering and DNA sequencing using the NovaSeq6000 was performed according to manufacturer's protocols. Mean amplicon coverages can be found in **SI Table 4.**

For the PNPCa model, sequence alignment was performed to the GRC37/hg19 reference using Burrows-Wheeler Aligner (BWA, v0.7.12) [2]. Somatic mutations and copy number alterations were determined according to the New York State Department of Health validated pipeline of EXaCT-1 clinical test [3], which accounts for the use of an amplicon-based WES assay, by comparison of each tumor sample with the matching control. The computational tool SPIA ensure that the samples from the same individual are analyzed [4].

For the BM18 and LAPC9 models, sequence reads were aligned to the reference human genome GRCh37 using BWA [2]. Local realignment, duplicate removal and base quality adjustment were performed using the Genome Analysis Toolkit (GATK, v3.6) [5] and Picard (http://broadinstitute.github.io/picard/). Somatic single nucleotide variants (SNVs) and small insertions and deletions (indels) were detected using MuTect (v1.1.4) [6] and Strelka (v1.0.15) [7], respectively, using the pool of normal as reference. We filtered out SNVs and indels outside of the target regions, those with variant allelic fraction (VAF) of <1% and/or those supported by <3 reads. We excluded variants for which the tumor VAF was <5 times that of the VAF of the pool of normal DNA sequenced and processed with the BM18 and LAPC9 samples, as well as all mutations found in the non-TCGA subset of the Exome Aggregation Consortium database v0.3.1 [8]. We further excluded variants identified in at least two of a panel of 123 non-tumoral samples [9] using the artifact detection mode of MuTect2 implemented in GATK.

To account for the presence of somatic mutations that may be present below the limit of sensitivity of somatic mutation callers, we used Genome Analysis Toolkit Unified Genotyper

[10] to interrogate the positions of all unique mutations in all tumor/organoids/PDX samples of a given tumor model to define the presence of additional mutations.

Allele-specific CNAs were identified using FACETS (v0.5.5) [11] as previously described [12], which performs a joint segmentation of the total and allelic copy ratio and infers allele-specific copy number states. Somatic mutations associated with the loss of the wild-type allele (i.e., loss of heterozygosity [LOH]) were identified as those were the lesser (minor) copy number state at the locus was 0. All mutations on chromosome X were considered to be associated with LOH. All gene amplifications and homozygous deletions were visually inspected using plots of raw log2 and allelic copy ratios. Copy number states were collapsed to the gene level based on the median values to coding gene resolution based on all coding genes retrieved from the Ensembl (release GRCh37.p13).

The cancer cell fraction (CCF) of each mutation on the autosomes was inferred using the number of reads supporting the reference and the alternate alleles, and the segmented log2 ratio from WES as input for ABSOLUTE (v1.0.6) [13]. Solutions from ABSOLUTE were manually reviewed as recommended [13, 14].

Cancer genes were annotated according to the cancer gene lists described by Kandoth et al. (127 significantly mutated genes) [15], Lawrence et al. (Cancer5000-S gene set) [16] and Armenia et al., (97 significantly mutated genes in PCa) [17]. Mutations affecting hotspot residues [18, 19] were annotated.

Decomposition of the mutational signature was performed using deconstructSigs [20] based on the set of 30 mutational signatures ("signature.cosmic," based on the signatures at https://cancer.sanger.ac.uk/cosmic/signatures) [21, 22]. Microsatellite instability detection was performed using MSIsensor. Score ≥3.5 was indicated as MSI-H according to the original publication [23].

**Targeted sequencing on the Ion torrent platform**

Targeted sequencing of the PCa organoids (PCa61, 62) was performed using a custom panel targeting the most frequently mutated genes in prostate cancers [24]. Somatic mutation calling was performed using PipeIT [25] which performs the initial variant calling step by Torrent Variant Caller (TVC, v5.0.3, Thermo Fisher Scientific) using low stringency variant calling parameters. PipeIT whitelists hotspot variants [19, 26] then filters out variants covered by less than 10 reads in either the tumor or the matched normal sample, supported by fewer than 8 reads or for which the tumor variant allele frequency (VAF) was <10 times that of the matched non-tumoral VAF. Whitelisted hotspot variants that did not pass the above read count and/or VAF filters were manually reviewed.

**RNA sequencing**

RNA extracted from FFPE (original patient material PNPCa), frozen tissue (PDXs) or organoids from PNPCa, BM18, LAPC9 and PCa cases (300ng) were subjected to RNA sequencing. Specimens were prepared for RNA sequencing using TruSeq RNA Library Preparation Kit v2 or riboZero as previously described [27]. RNA integrity was verified using the Agilent Bioanalyzer 2100 (Agilent Technologies). cDNA was synthesized from total RNA using Superscript III (Invitrogen). Sequencing was then performed on GAII, HiSeq 2000, or HiSeq 2500.

For the RNASeq, the NEBNext Ultra II Directional RNA Library Prep Kit for Illumina was used to process the sample(s). The sample preparation was performed according to the protocol "NEBNext Ultra II Directional RNA Library Prep Kit for Illumina" (NEB #E7760S/L). Briefly, mRNA was isolated from total RNA using the oligo-dT magnetic beads. After fragmentation of the mRNA, a cDNA synthesis was performed. This was used for ligation with the sequencing adapters and PCR amplification of the resulting product. The quality and yield after sample preparation was measured with the Fragment Analyzer. The size of the resulting products was consistent with the expected size distribution (a broad peak between 300-500 bp). Clustering and DNA sequencing using the NovaSeq6000 was performed according to manufacturer's protocols. A concentration of 1.1 nM of DNA was used. Image analysis, base calling, and quality check was performed with the Illumina data analysis pipeline RTA3.4.4 and Bcl2fastq v2.20.

Sequence reads were aligned using STAR two-pass to the human reference genome GRCh37 [28]. For the data analysis, RSEM was used to obtain FPKM (fragments per kilobase of exon model per million reads mapped) counts. We performed single sample enrichment analysis (ssGSEA) in the R package gsva, the C2 (canonical pathways), C6 (oncogenic signatures) and H (hallmark gene sets) [29]. The enrichment scores for each sample was normalized to the normal sample (N1). For annotated to multiple Ensembl genes, the one with highest expression was used for ssGSEA analysis.

Differential expression analysis was performed using the edgeR package [30]. Normalization was performed using the “TMM” (weighted trimmed mean) method and differential expression was assessed using the quasi-likelihood F-test, including a batch factor into the design matrix when required. Genes with FDR <0.05 and > 2-fold were considered significantly differentially expressed.

For gene set enrichment analysis (GSEA) we ranked genes based on T-statistics from differential expression analysis. Gene sets used were the C2 (KEGG and REACTOME gene sets), and H (hallmark gene sets) [29].

### Data Availability

Sequence data have been deposited at the European Genome-phenome Archive (EGA), which is hosted by the EBI and the CRG, under accession number (to be uploaded).

#### Fragment Analysis

MSI was assessed on the basis of the recommendations of the National Cancer Institute workshop on MSI, a panel of microsatellite loci (BAT25, BAT26, D2S123, D5S346, and D17S250) and two additional microsatellite markers (BAT40 and MYCL1) were used to determine MSI status [31]. PCR and fragment analysis were performed as previously described [32].

#### Transcriptomic signature profiling of SPOP, FOXA1, ETS genetic subgroups

For the analysis of transcriptomic signatures of the *SPOP*, *FOXA1* and ETS genetic subgroups [33], we retrieved gene-expression of 480 primary prostate cancer specimen from The Cancer Genome Atlas (TCGA). Raw-counts were retrieved using TCGABiolinks [34] package in R statistical environment. Normalization was performed using DESeq2 [35] pipeline, and data was ultimately transformed using variance stabilizing transformation. Genetic characterization of the tumors, in particular, mutations in *SPOP*, *FOXA1* and genetic rearrangements involving ETS transcription factors, was performed using annotations retrieved from cBioportal. We generated gene-signatures specific to each genetic alteration. To this purpose, we stratified the cohort in different subgroups (*FOXA1*, *SPOP*, *ERG*, *ETV1*, *ETV4*) and performed differential expression between these populations and all samples not affected by mutations in *SPOP/FOXA1* or ETS-rearrangements. We used a False Discovery Rate threshold of 0.001 and an absolute fold-change of 1. Gene-set activities were determined in individual patients using single sample gene-set enrichment analysis (ssGSEA) and subsequently averaged across groups.

### References


1. Rennert, H., et al., *Development and validation of a whole-exome sequencing test for simultaneous detection of point mutations, indels and copy-number alterations for precision cancer care.* Npj Genomic Medicine, 2016. **1**: p. 16019.
2. Li, H. and R. Durbin, *Fast and accurate short read alignment with Burrows-Wheeler transform.* Bioinformatics, 2009. **25**(14): p. 1754-60.
3. Rennert, H., et al., *Development and validation of a whole-exome sequencing test for simultaneous detection of point mutations, indels and copy-number alterations for precision cancer care.* NPJ Genom Med, 2016. **1**.
4. Demichelis, F., et al., *SNP panel identification assay (SPIA): a genetic-based assay for the identification of cell lines.* Nucleic Acids Res, 2008. **36**(7): p. 2446-56.
5. McKenna, A., et al., *The Genome Analysis Toolkit: a MapReduce framework for analyzing next-generation DNA sequencing data.* Genome research, 2010. **20**(9): p. 1297-1303.
6. Cibulskis, K., et al., *Sensitive detection of somatic point mutations in impure and heterogeneous cancer samples.* Nature Biotechnology, 2013. **31**: p. 213.
7. Saunders, C.T., et al., *Strelka: accurate somatic small-variant calling from sequenced tumor–normal sample pairs.* Bioinformatics, 2012. **28**(14): p. 1811-1817.
8. Lek, M., et al., *Analysis of protein-coding genetic variation in 60,706 humans.* Nature, 2016. **536**: p. 285.
9. Nuciforo, S., et al., *Organoid Models of Human Liver Cancers Derived from Tumor Needle Biopsies.* Cell Reports, 2018. **24**(5): p. 1363-1376.
10. McKenna, A., et al., *The Genome Analysis Toolkit: a MapReduce framework for analyzing next-generation DNA sequencing data.* Genome Res, 2010. **20**(9): p. 1297-303.
11. Shen, R. and V.E. Seshan, *FACETS: allele-specific copy number and clonal heterogeneity analysis tool for high-throughput DNA sequencing.* Nucleic acids research, 2016. **44**(16): p. e131-e131.
12. Piscuoglio, S., et al., *The Genomic Landscape of Male Breast Cancers.* Clinical cancer research : an official journal of the American Association for Cancer Research, 2016. **22**(16): p. 4045-4056.
13. Carter, S.L., et al., *Absolute quantification of somatic DNA alterations in human cancer.* Nature biotechnology, 2012. **30**(5): p. 413-421.
14. Landau, D.A., et al., *Evolution and impact of subclonal mutations in chronic lymphocytic leukemia.* Cell, 2013. **152**(4): p. 714-726.
15. Kandoth, C., et al., *Mutational landscape and significance across 12 major cancer types.* Nature, 2013. **502**(7471): p. 333-339.
16. Lawrence, M.S., et al., *Discovery and saturation analysis of cancer genes across 21 tumour types.* Nature, 2014. **505**(7484): p. 495-501.
17. Armenia, J., et al., *The long tail of oncogenic drivers in prostate cancer.* Nat Genet, 2018. **50**(5): p. 645-651.
18. Chang, M.T., et al., *Identifying recurrent mutations in cancer reveals widespread lineage diversity and mutational specificity.* Nature biotechnology, 2016. **34**(2): p. 155-163.
19. Gao, J., et al., *3D clusters of somatic mutations in cancer reveal numerous rare mutations as functional targets.* Genome medicine, 2017. **9**(1): p. 4-4.
20. Rosenthal, R., et al., *DeconstructSigs: delineating mutational processes in single tumors distinguishes DNA repair deficiencies and patterns of carcinoma evolution.* Genome biology, 2016. **17**: p. 31-31.
21. Nik-Zainal, S., et al., *Landscape of somatic mutations in 560 breast cancer whole-genome sequences.* Nature, 2016. **534**(7605): p. 47-54.
22. Alexandrov, L.B., et al., *Signatures of mutational processes in human cancer.* Nature, 2013. **500**(7463): p. 415-421.

23. Niu, B., et al., *MSIsensor: microsatellite instability detection using paired tumor-normal sequence data.* Bioinformatics, 2014. **30**(7): p. 1015-6.
24. Salami, S.S., et al., *Transcriptomic heterogeneity in multifocal prostate cancer.* JCI Insight, 2018. **3**(21).
25. Garofoli, A., et al., *PipeIT: A Singularity Container for Molecular Diagnostic Somatic Variant Calling on the Ion Torrent Next-Generation Sequencing Platform.* The Journal of Molecular Diagnostics, 2019. **21**(5): p. 884-894.
26. Chang, M.T., et al., *Identifying recurrent mutations in cancer reveals widespread lineage diversity and mutational specificity.* Nature Biotechnology, 2015. **34**: p. 155.
27. Beltran, H., et al., *Whole-Exome Sequencing of Metastatic Cancer and Biomarkers of Treatment Response.* JAMA Oncol, 2015. **1**(4): p. 466-74.
28. Dobin, A., et al., *STAR: ultrafast universal RNA-seq aligner.* Bioinformatics (Oxford, England), 2013. **29**(1): p. 15-21.
29. Liberzon, A., et al., *Molecular signatures database (MSigDB) 3.0.* Bioinformatics, 2011. **27**(12): p. 1739-40.
30. Nikolayeva, O. and M.D. Robinson, *edgeR for differential RNA-seq and ChIP-seq analysis: an application to stem cell biology.* Methods Mol Biol, 2014. **1150**: p. 45-79.
31. Umar, A., et al., *Revised Bethesda Guidelines for hereditary nonpolyposis colorectal cancer (Lynch syndrome) and microsatellite instability.* J Natl Cancer Inst, 2004. **96**(4): p. 261-8.
32. Kishore, S., et al., *3'-UTR poly(T/U) tract deletions and altered expression of EWSR1 are a hallmark of mismatch repair-deficient cancers.* Cancer Res, 2014. **74**(1): p. 224-34.
33. Barbieri, C.E., et al., *Exome sequencing identifies recurrent SPOP, FOXA1 and MED12 mutations in prostate cancer.* Nat Genet, 2012. **44**(6): p. 685-9.
34. Colaprico, A., et al., *TCGAbiolinks: an R/Bioconductor package for integrative analysis of TCGA data.* Nucleic Acids Research, 2015. **44**(8): p. e71-e71.
35. Love, M.I., W. Huber, and S. Anders, *Moderated estimation of fold change and dispersion for RNA-seq data with DESeq2.* Genome Biology, 2014. **15**(12): p. 550.
