## Supplementary Information Tables for "Patient-derived xenografts and organoids model therapy response in prostate cancer": S.Table 1. Tumor growth PNPCA 2way ANOVA castrated vs replaced.pdf

Compare each cell mean with the other cell mean in that row

| | |
|---|---|
| Number of | 1 |
| Number of | 43 |
| Alpha | 0.05 |

| Sidak's mu | Predicted (LS) mean diff. | 95.00% CI of diff. | Significant | Summary | Adjusted P Value |
|---|---|---|---|---|---|
| Castrated - Castrated - Testost. | | | | | |
| 0 | 4.00E-15 | -2.699 to 2.699 | No | ns | >0.9999 |
| 7 | 1.33E-15 | -2.699 to 2.699 | No | ns | >0.9999 |
| 14 | 8.88E-16 | -2.699 to 2.699 | No | ns | >0.9999 |
| 21 | 1.78E-15 | -2.699 to 2.699 | No | ns | >0.9999 |
| 28 | -8.88E-16 | -2.699 to 2.699 | No | ns | >0.9999 |
| 35 | -0.225 | -2.924 to 2.474 | No | ns | >0.9999 |
| 42 | -0.5 | -3.199 to 2.199 | No | ns | >0.9999 |
| 48 | -0.875 | -3.574 to 1.824 | No | ns | >0.9999 |
| 55 | -0.05 | -2.749 to 2.649 | No | ns | >0.9999 |
| 62 | -1.325 | -4.024 to 1.374 | No | ns | 0.993 |
| 67 | -1.325 | -4.024 to 1.374 | No | ns | 0.993 |
| 74 | -1.25 | -3.949 to 1.449 | No | ns | 0.9976 |
| 81 | -0.575 | -3.274 to 2.124 | No | ns | >0.9999 |
| 91 | -0.575 | -3.274 to 2.124 | No | ns | >0.9999 |
| 102 | 0.625 | -2.074 to 3.324 | No | ns | >0.9999 |
| 108 | -0.175 | -2.874 to 2.524 | No | ns | >0.9999 |
| 116 | 0.1667 | -2.532 to 2.865 | No | ns | >0.9999 |
| 123 | -0.625 | -3.324 to 2.074 | No | ns | >0.9999 |
| 130 | -0.5833 | -3.282 to 2.115 | No | ns | >0.9999 |
| 137 | -0.5 | -3.199 to 2.199 | No | ns | >0.9999 |
| 144 | -0.4167 | -3.115 to 2.282 | No | ns | >0.9999 |
| 151 | -0.125 | -2.824 to 2.574 | No | ns | >0.9999 |
| 158 | -1.78E-15 | -2.699 to 2.699 | No | ns | >0.9999 |
| 161 | 1.33E-15 | -2.699 to 2.699 | No | ns | >0.9999 |
| 165 | 0.3333 | -2.365 to 3.032 | No | ns | >0.9999 |
| 168 | 1.78E-15 | -2.699 to 2.699 | No | ns | >0.9999 |
| 172 | -0.25 | -2.949 to 2.449 | No | ns | >0.9999 |
| 175 | -1.33E-15 | -2.699 to 2.699 | No | ns | >0.9999 |
| 179 | -0.2083 | -2.907 to 2.490 | No | ns | >0.9999 |
| 186 | -0.625 | -3.324 to 2.074 | No | ns | >0.9999 |
| 189 | -0.625 | -3.324 to 2.074 | No | ns | >0.9999 |
| 196 | -0.8333 | -3.532 to 1.865 | No | ns | >0.9999 |
| 203 | -1.333 | -4.271 to 1.605 | No | ns | 0.9984 |
| 210 | -2.5 | -5.438 to 0.4379 | No | ns | 0.2179 |
| 217 | -3.333 | -6.271 to -0.3954 | Yes | * | 0.0105 |
| 224 | -3.667 | -6.605 to -0.7288 | Yes | ** | 0.0025 |
| 231 | -4 | -6.938 to -1.062 | Yes | *** | 0.0005 |
| 238 | -4.5 | -7.438 to -1.562 | Yes | **** | <0.0001 |
| 245 | -5.333 | -8.271 to -2.395 | Yes | **** | <0.0001 |
| 252 | -5 | -8.366 to -1.634 | Yes | **** | <0.0001 |
| 259 | -6.5 | -9.866 to -3.134 | Yes | **** | <0.0001 |
| 267 | -8 | -11.37 to -4.634 | Yes | **** | <0.0001 |
| 273 | -9.5 | -12.87 to -6.134 | Yes | **** | <0.0001 |

| Test detail | Predicted (LS) mean 1 | Predicted (LS) mean 2 | Predicted ( | SE of diff. | N1 | N2 | t | DF |
|---|---|---|---|---|---|---|---|---|
| Castrated - Castrated - Testost. | | | | | | | | |
| 0 | 8.88E-16 | -3.11E-15 | 4.00E-15 | 0.8244 | 5 | 4 | 4.85E-15 | 286 |
| 7 | 0 | -1.33E-15 | 1.33E-15 | 0.8244 | 5 | 4 | 1.62E-15 | 286 |
| 14 | 0 | -8.88E-16 | 8.88E-16 | 0.8244 | 5 | 4 | 1.08E-15 | 286 |
| 21 | 0 | -1.78E-15 | 1.78E-15 | 0.8244 | 5 | 4 | 2.16E-15 | 286 |
| 28 | -8.88E-16 | 0 | -8.88E-16 | 0.8244 | 5 | 4 | 1.08E-15 | 286 |
| 35 | 0.9 | 1.125 | -0.225 | 0.8244 | 5 | 4 | 0.2729 | 286 |
| 42 | 1.5 | 2 | -0.5 | 0.8244 | 5 | 4 | 0.6065 | 286 |
| 48 | 2.5 | 3.375 | -0.875 | 0.8244 | 5 | 4 | 1.061 | 286 |
| 55 | 3.7 | 3.75 | -0.05 | 0.8244 | 5 | 4 | 0.06065 | 286 |
| 62 | 4.3 | 5.625 | -1.325 | 0.8244 | 5 | 4 | 1.607 | 286 |
| 67 | 6.8 | 8.125 | -1.325 | 0.8244 | 5 | 4 | 1.607 | 286 |
| 74 | 6 | 7.25 | -1.25 | 0.8244 | 5 | 4 | 1.516 | 286 |
| 81 | 4.8 | 5.375 | -0.575 | 0.8244 | 5 | 4 | 0.6975 | 286 |
| 91 | 3.3 | 3.875 | -0.575 | 0.8244 | 5 | 4 | 0.6975 | 286 |
| 102 | 2.5 | 1.875 | 0.625 | 0.8244 | 5 | 4 | 0.7582 | 286 |
| 108 | 1.7 | 1.875 | -0.175 | 0.8244 | 5 | 4 | 0.2123 | 286 |
| 116 | 1.167 | 1 | 0.1667 | 0.8244 | 5 | 4 | 0.2022 | 286 |
| 123 | 0 | 0.625 | -0.625 | 0.8244 | 5 | 4 | 0.7582 | 286 |
| 130 | 0.6667 | 1.25 | -0.5833 | 0.8244 | 5 | 4 | 0.7076 | 286 |
| 137 | 0.5 | 1 | -0.5 | 0.8244 | 5 | 4 | 0.6065 | 286 |
| 144 | 0.3333 | 0.75 | -0.4167 | 0.8244 | 5 | 4 | 0.5054 | 286 |
| 151 | 1.33E-15 | 0.125 | -0.125 | 0.8244 | 5 | 4 | 0.1516 | 286 |
| 158 | -8.88E-16 | 8.88E-16 | -1.78E-15 | 0.8244 | 5 | 4 | 2.16E-15 | 286 |
| 161 | 0 | -1.33E-15 | 1.33E-15 | 0.8244 | 5 | 4 | 1.62E-15 | 286 |
| 165 | 0.3333 | -8.88E-16 | 0.3333 | 0.8244 | 5 | 4 | 0.4044 | 286 |
| 168 | 0 | -1.78E-15 | 1.78E-15 | 0.8244 | 5 | 4 | 2.16E-15 | 286 |
| 172 | 8.88E-16 | 0.25 | -0.25 | 0.8244 | 5 | 4 | 0.3033 | 286 |
| 175 | -1.33E-15 | 0 | -1.33E-15 | 0.8244 | 5 | 4 | 1.62E-15 | 286 |
| 179 | 0.1667 | 0.375 | -0.2083 | 0.8244 | 5 | 4 | 0.2527 | 286 |
| 186 | -1.33E-15 | 0.625 | -0.625 | 0.8244 | 5 | 4 | 0.7582 | 286 |
| 189 | -2.22E-15 | 0.625 | -0.625 | 0.8244 | 5 | 4 | 0.7582 | 286 |
| 196 | 0.5 | 1.333 | -0.8333 | 0.8244 | 5 | 4 | 1.011 | 286 |
| 203 | 8.88E-16 | 1.333 | -1.333 | 0.8974 | 5 | 3 | 1.486 | 286 |
| 210 | 0.3333 | 2.833 | -2.5 | 0.8974 | 5 | 3 | 2.786 | 286 |
| 217 | -1.11E-15 | 3.333 | -3.333 | 0.8974 | 5 | 3 | 3.714 | 286 |
| 224 | -8.88E-16 | 3.667 | -3.667 | 0.8974 | 5 | 3 | 4.086 | 286 |
| 231 | 0.1667 | 4.167 | -4 | 0.8974 | 5 | 3 | 4.457 | 286 |
| 238 | 0.1667 | 4.667 | -4.5 | 0.8974 | 5 | 3 | 5.014 | 286 |
| 245 | -1.78E-15 | 5.333 | -5.333 | 0.8974 | 5 | 3 | 5.943 | 286 |
| 252 | -8.88E-15 | 5 | -5 | 1.028 | 5 | 2 | 4.863 | 286 |
| 259 | -6.66E-15 | 6.5 | -6.5 | 1.028 | 5 | 2 | 6.322 | 286 |
| 267 | -7.11E-15 | 8 | -8 | 1.028 | 5 | 2 | 7.781 | 286 |
| 273 | 0 | 9.5 | -9.5 | 1.028 | 5 | 2 | 9.24 | 286 |
