## Supplementary Information Tables for "Patient-derived xenografts and organoids model therapy response in prostate cancer": S.Table 2. somatic mutations PNPCa.pdf

| TUMOR_SAMPLE | GENE | MUTATION | EFFECT | IMPACT | TUMOR VAF | NORMAL VAF | TUMOR DEPTH | NORMAL DEPTH | cancer_gene_census | kandoth | lawrence | prostate_smg |
|---|---|---|---|---|---|---|---|---|---|---|---|---|
| PM1548-SK1Org2 | B3GALT6 | p.Glu303Lys | missense_variant | MODERATE | 0.617647 | 0 | 34 | 86 | . | . | . | . |
| PM1548-SK1Org2 | SCNN1D | p.Arg344His | missense_variant | MODERATE | 0.460784 | 0 | 102 | 274 | . | . | . | . |
| PM1548-SK1Org2 | ATAD3B | p.Arg290Cys | missense_variant | MODERATE | 0.516129 | 0.00687285 | 155 | 291 | . | . | . | . |
| PM1548-SK1Org2 | C1orf86 | p.Pro72His | missense_variant | MODERATE | 0.597222 | 0 | 72 | 43 | . | . | . | . |
| PM1548-SK1Org2 | PRDM16 | p.Arg759Gln | missense_variant | MODERATE | 0.117647 | 0 | 85 | 40 | true | . | . | . |
| PM1548-SK1Org2 | DFFB | p.Tyr290Phe | missense_variant | MODERATE | 0.30086 | 0 | 349 | 203 | . | . | . | . |
| PM1548-SK1Org2 | NPHP4 | p.Arg488Gln | missense_variant | MODERATE | 0.361111 | 0 | 108 | 194 | . | . | . | . |
| PM1548-SK1Org2 | PLEKHG5 | p.Thr301fs | frameshift_variant | HIGH | 0.485075 | 0 | 134 | 105 | . | . | . | . |
| PM1548-SK1Org2 | ZBTB48 | p.Val626Ala | missense_variant | MODERATE | 0.256966 | 0 | 323 | 142 | . | . | . | . |
| PM1548-SK1Org2 | KLHL21 | p.Cys250fs | frameshift_variant | HIGH | 0.0588235 | 0 | 17 | 136 | . | . | . | . |
| PM1548-SK1Org2 | CAMTA1 | p.Ser510Thr | missense_variant | MODERATE | 0.00675676 | 0 | 148 | 185 | true | . | . | . |
| PM1548-SK1Org2 | PER3 | p.Ala803Ser | missense_variant | MODERATE | 0.00321543 | 0 | 311 | 110 | . | . | . | . |
| PM1548-SK1Org2 | SLC45A1 | p.Thr364Met | missense_variant | MODERATE | 0.646341 | 0 | 82 | 57 | . | . | . | . |
| PM1548-SK1Org2 | SLC45A1 | p.Tyr689* | stop_gained | HIGH | 0.531469 | 0 | 143 | 120 | . | . | . | . |
| PM1548-SK1Org2 | RERE | p.Lys83fs | frameshift_variant | HIGH | 0.320988 | 0 | 81 | 49 | . | . | . | . |
| PM1548-SK1Org2 | CORT | p.Gln133fs | frameshift_variant | HIGH | 0 | 0 | 141 | 48 | . | . | . | . |
| PM1548-SK1Org2 | EXOSC10 | p.Met1fs | frameshift_variant&start_lost | HIGH | 0.457944 | 0 | 107 | 61 | . | . | . | . |
| PM1548-SK1Org2 | FHAD1 | p.Ile953fs | frameshift_variant | HIGH | 0.492424 | 0 | 132 | 108 | . | . | . | . |
| PM1548-SK1Org2 | PLEKHM2 | p.Arg848Trp | missense_variant | MODERATE | 0 | 0 | 34 | 42 | . | . | . | . |
| PM1548-SK1Org2 | SPEN | p.Asn2002fs | frameshift_variant | HIGH | 0 | 0 | 310 | 173 | true | . | true | true |
| PM1548-SK1Org2 | EPHA2 | p.Asn106Ser | missense_variant | MODERATE | 0.303371 | 0 | 267 | 84 | . | . | true | . |
| PM1548-SK1Org2 | CROCC | p.Arg1590Cys | missense_variant | MODERATE | 0.318182 | 0 | 132 | 547 | . | . | . | . |
| PM1548-SK1Org2 | RCC2 | p.Gly465Arg | missense_variant | MODERATE | 0.494737 | 0 | 190 | 148 | . | . | . | . |
| PM1548-SK1Org2 | ARHGEF10L | p.Val965Met | missense_variant | MODERATE | 0 | 0 | 151 | 230 | . | . | . | . |
| PM1548-SK1Org2 | KLHDC7A | p.Ala652Val | missense_variant | MODERATE | 0.340659 | 0 | 91 | 60 | . | . | . | . |
| PM1548-SK1Org2 | RNF186 | p.Ile185Thr | missense_variant | MODERATE | 0 | 0 | 214 | 472 | . | . | . | . |
| PM1548-SK1Org2 | KIF17 | p.Arg196His | missense_variant | MODERATE | 0.537931 | 0 | 145 | 71 | . | . | . | . |
| PM1548-SK1Org2 | EIF4G3 | c.30+5101C>G | intron_variant | MODIFIER | 0 | 0 | 147 | 302 | . | . | . | . |
| PM1548-SK1Org2 | HSPG2 | p.Gly2259Asp | missense_variant | MODERATE | 0 | 0 | 147 | 55 | . | . | . | . |
| PM1548-SK1Org2 | EPHA8 | p.Ser87Asn | missense_variant | MODERATE | 0.0779221 | 0 | 231 | 190 | . | . | . | . |
| PM1548-SK1Org2 | AIM1L | n.26671971delA | intragenic_variant | MODIFIER | 0.0135135 | 0 | 148 | 55 | . | . | . | . |
| PM1548-SK1Org2 | ARID1A | p.Thr1863Ile | missense_variant | MODERATE | 0.0264901 | 0 | 151 | 201 | true | true | true | true |
| PM1548-SK1Org2 | PTPRU | p.Arg639Trp | missense_variant | MODERATE | 0 | 0 | 68 | 60 | . | . | . | . |
| PM1548-SK1Org2 | FABP3 | p.Thr54fs | frameshift_variant | HIGH | 0.497653 | 0 | 213 | 207 | . | . | . | . |
| PM1548-SK1Org2 | LCK | p.Arg222Cys | missense_variant | MODERATE | 0 | 0 | 267 | 146 | true | . | . | . |
| PM1548-SK1Org2 | KIAA1522 | p.Ala943Thr | missense_variant | MODERATE | 0.47651 | 0 | 149 | 198 | . | . | . | . |
| PM1548-SK1Org2 | ZNF362 | p.Ala105Val | missense_variant | MODERATE | 0 | 0 | 12 | 295 | . | . | . | . |
| PM1548-SK1Org2 | LSM10 | p.Ala2Val | missense_variant | MODERATE | 0.459547 | 0 | 309 | 400 | . | . | . | . |
| PM1548-SK1Org2 | CSF3R | p.Tyr508Cys | missense_variant | MODERATE | 0.0535714 | 0 | 112 | 167 | true | . | . | . |
| PM1548-SK1Org2 | EPHA10 | p.Pro330Ser | missense_variant | MODERATE | 0.589041 | 0 | 146 | 71 | . | . | . | . |
| PM1548-SK1Org2 | MACF1 | p.Gln6033Pro | missense_variant&splice_region_variant | MODERATE | 0 | 0 | 48 | 48 | . | . | . | . |
| PM1548-SK1Org2 | PABPC4 | p.Ser206Asn | missense_variant | MODERATE | 0.0512821 | 0 | 234 | 90 | . | . | . | . |
| PM1548-SK1Org2 | PABPC4 | p.Glu205Gly | missense_variant | MODERATE | 0.0594059 | 0 | 202 | 73 | . | . | . | . |
| PM1548-SK1Org2 | ZMPSTE24 | p.Leu362fs | frameshift_variant | HIGH | 0.45122 | 0 | 82 | 53 | . | . | . | . |
| PM1548-SK1Org2 | HIVEP3 | p.Pro1037Leu | missense_variant | MODERATE | 0.474074 | 0 | 135 | 35 | . | . | . | . |
| PM1548-SK1Org2 | MED8 | p.Asn295fs | frameshift_variant | HIGH | 0.627119 | 0 | 118 | 97 | . | . | . | . |
| PM1548-SK1Org2 | HYI | p.Ala55Ser | missense_variant | MODERATE | 0 | 0 | 35 | 158 | . | . | . | . |
| PM1548-SK1Org2 | PTPRF | p.Arg1669Trp | missense_variant | MODERATE | 0.476415 | 0 | 212 | 112 | . | . | . | . |
| PM1548-SK1Org2 | SLC6A9 | p.Arg662Gln | missense_variant | MODERATE | 0.536667 | 0 | 300 | 99 | . | . | . | . |
| PM1548-SK1Org2 | PTCH2 | p.Ala967Val | missense_variant | MODERATE | 0.00588235 | 0 | 170 | 194 | . | . | . | . |
| PM1548-SK1Org2 | PRDX1 | p.His10Tyr | missense_variant | MODERATE | 0 | 0 | 179 | 151 | . | . | . | . |
| PM1548-SK1Org2 | MAST2 | p.Arg1086Arg | synonymous_variant | LOW | 0.0277778 | 0 | 180 | 187 | . | . | . | . |
| PM1548-SK1Org2 | TAL1 | p.His86Leu | missense_variant | MODERATE | 0.128205 | 0 | 39 | 170 | true | . | . | . |
| PM1548-SK1Org2 | LRP8 | p.Asp50Gly | missense_variant | MODERATE | 0.365672 | 0 | 134 | 175 | . | . | . | . |
| PM1548-SK1Org2 | CDCP2 | p.Pro209fs | frameshift_variant | HIGH | 0.521739 | 0 | 230 | 107 | . | . | . | . |
| PM1548-SK1Org2 | MRPL37 | p.Asp142Glu | missense_variant | MODERATE | 0.0078125 | 0 | 128 | 91 | . | . | . | . |
| PM1548-SK1Org2 | CACHD1 | p.Glu211Lys | missense_variant | MODERATE | 0 | 0 | 151 | 147 | . | . | . | . |
| PM1548-SK1Org2 | JAK1 | p.Val433Ile | missense_variant | MODERATE | 0.252788 | 0 | 269 | 203 | true | . | . | true |
| PM1548-SK1Org2 | FPGT | p.Asp273Asn | missense_variant | MODERATE | 0 | 0 | 53 | 168 | . | . | . | . |
| PM1548-SK1Org2 | LPHN2 | p.Asn759Ile | missense_variant | MODERATE | 0.29703 | 0 | 101 | 85 | . | . | . | . |
| PM1548-SK1Org2 | SYDE2 | p.Gly14fs | frameshift_variant | HIGH | 0 | 0 | 48 | 141 | . | . | . | . |
| PM1548-SK1Org2 | CLCA1 | p.Arg676Trp | missense_variant | MODERATE | 0.433333 | 0 | 60 | 91 | . | . | . | . |
| PM1548-SK1Org2 | GBP6 | p.Gly77Asp | missense_variant | MODERATE | 0.262774 | 0 | 137 | 98 | . | . | . | . |
| PM1548-SK1Org2 | GFI1 | p.Arg348Met | missense_variant | MODERATE | 0.508403 | 0 | 238 | 270 | . | . | . | . |
| PM1548-SK1Org2 | ABCA4 | p.His423Gln | missense_variant | MODERATE | 0.0858896 | 0 | 163 | 257 | . | . | . | . |
| PM1548-SK1Org2 | ABCA4 | p.Glu422Asp | missense_variant | MODERATE | 0.0858896 | 0.00389105 | 163 | 257 | . | . | . | . |
| PM1548-SK1Org2 | LPPR4 | p.Ala343Thr | missense_variant | MODERATE | 0.411215 | 0 | 107 | 96 | . | . | . | . |
| PM1548-SK1Org2 | HIAT1 | p.Ala235Val | missense_variant&splice_region_variant | MODERATE | 0.576642 | 0 | 137 | 80 | . | . | . | . |
| PM1548-SK1Org2 | TRMT13 | p.Glu307fs | frameshift_variant | HIGH | 0 | 0 | 144 | 85 | . | . | . | . |
| PM1548-SK1Org2 | AKNAD1 | p.Leu577Met | missense_variant | MODERATE | 0.53125 | 0 | 288 | 457 | . | . | . | . |
| PM1548-SK1Org2 | CELSR2 | p.Asp1795Gly | missense_variant | MODERATE | 0.00215517 | 0 | 464 | 362 | . | . | . | . |
| PM1548-SK1Org2 | LAMTOR5 | p.Thr39Ala | missense_variant | MODERATE | 0 | 0 | 106 | 96 | . | . | . | . |
| PM1548-SK1Org2 | HIPK1 | p.Ala314Thr | missense_variant | MODERATE | 0.496644 | 0 | 149 | 273 | . | . | . | . |
| PM1548-SK1Org2 | TRIM33 | p.Val961Met | missense_variant | MODERATE | 0.341463 | 0 | 41 | 36 | true | . | . | . |
| PM1548-SK1Org2 | PDE4DIP | p.Gln2125Lys | missense_variant | MODERATE | 0.10559 | 0 | 322 | 86 | true | . | . | . |
| PM1548-SK1Org2 | PDE4DIP | p.Arg759* | stop_gained | HIGH | 0.229437 | 0 | 231 | 105 | true | . | . | . |
| PM1548-SK1Org2 | BCL9 | p.Pro517fs | frameshift_variant | HIGH | 0.484848 | 0 | 99 | 144 | true | . | . | . |
| PM1548-SK1Org2 | TARS2 | p.Tyr371* | stop_gained | HIGH | 0.232558 | 0 | 129 | 48 | . | . | . | . |
| PM1548-SK1Org2 | ADAMTSL4 | p.Gly780fs | frameshift_variant | HIGH | 0.532468 | 0 | 154 | 108 | . | . | . | . |
| PM1548-SK1Org2 | ZNF687 | p.Arg885His | missense_variant | MODERATE | 0.470899 | 0 | 189 | 322 | . | . | . | . |
| PM1548-SK1Org2 | PI4KB | p.Val202Met | missense_variant | MODERATE | 0 | 0 | 219 | 157 | . | . | . | . |
| PM1548-SK1Org2 | CELF3 | p.Val279Ile | missense_variant | MODERATE | 0.583333 | 0 | 108 | 53 | . | . | . | . |
| PM1548-SK1Org2 | DENND4B | p.Ala154fs | frameshift_variant | HIGH | 0.0915033 | 0 | 153 | 107 | . | . | . | . |
| PM1548-SK1Org2 | CREB3L4 | p.Val263fs | frameshift_variant | HIGH | 0.440559 | 0 | 143 | 42 | . | . | . | . |
| PM1548-SK1Org2 | EFNA4 | c.470-101delG | intron_variant | MODIFIER | 0 | 0 | 156 | 82 | . | . | . | . |
| PM1548-SK1Org2 | ASH1L | p.Asn1907Lys | missense_variant | MODERATE | 0 | 0 | 142 | 55 | . | . | . | . |
| PM1548-SK1Org2 | SMG5 | p.Gln846Lys | missense_variant | MODERATE | 0 | 0 | 72 | 128 | . | . | . | . |
| PM1548-SK1Org2 | CD1D | p.Asp140Asn | missense_variant | MODERATE | 0.484375 | 0 | 320 | 198 | . | . | true | . |
| PM1548-SK1Org2 | AIM2 | p.Lys342fs | frameshift_variant | HIGH | 0.476636 | 0 | 107 | 30 | . | . | . | . |
| PM1548-SK1Org2 | NIT1 | p.Arg261His | missense_variant | MODERATE | 0.489796 | 0 | 196 | 402 | . | . | . | . |
| PM1548-SK1Org2 | ILDR2 | p.Gly453Ala | missense_variant | MODERATE | 0.255319 | 0 | 47 | 327 | . | . | . | . |
| PM1548-SK1Org2 | ILDR2 | p.Arg415Trp | missense_variant | MODERATE | 0.407143 | 0 | 140 | 213 | . | . | . | . |
| PM1548-SK1Org2 | MYOC | p.Arg91Gln | missense_variant | MODERATE | 0 | 0 | 109 | 242 | . | . | . | . |
| PM1548-SK1Org2 | DNM3 | p.Glu7Asp | missense_variant | MODERATE | 0 | 0 | 82 | 77 | . | . | . | . |
| PM1548-SK1Org2 | DNM3 | p.Phe20Phe | synonymous_variant | LOW | 0.025641 | 0.0060241 | 156 | 166 | . | . | . | . |
| PM1548-SK1Org2 | DNM3 | p.His85Gln | missense_variant | MODERATE | 0.31746 | 0 | 63 | 56 | . | . | . | . |
| PM1548-SK1Org2 | ANKRD45 | p.Leu39fs | frameshift_variant | HIGH | 0.285714 | 0 | 105 | 229 | . | . | . | . |
| PM1548-SK1Org2 | ZBTB37 | p.Val284Ile | missense_variant | MODERATE | 0.0530973 | 0 | 113 | 152 | . | . | . | . |
| PM1548-SK1Org2 | ZBTB37 | p.Asn288Ser | missense_variant | MODERATE | 0.0504202 | 0 | 119 | 160 | . | . | . | . |
| PM1548-SK1Org2 | PAPPA2 | p.Thr1174fs | frameshift_variant | HIGH | 0.554217 | 0 | 83 | 36 | . | . | . | . |
| PM1548-SK1Org2 | CACNA1E | p.Asn2301Ser | missense_variant | MODERATE | 0.00458716 | 0 | 218 | 120 | . | . | . | . |
| PM1548-SK1Org2 | CACNA1E | p.Asn2302Asp | missense_variant | MODERATE | 0.0182648 | 0 | 219 | 119 | . | . | . | . |
| PM1548-SK1Org2 | CACNA1E | p.Asp2310Asp | synonymous_variant | LOW | 0.00497512 | 0 | 201 | 90 | . | . | . | . |
| PM1548-SK1Org2 | ZNF648 | p.Ser436Tyr | missense_variant | MODERATE | 0.0306122 | 0 | 196 | 389 | . | . | . | . |
| PM1548-SK1Org2 | ZNF648 | p.Ser436Pro | missense_variant | MODERATE | 0.0347826 | 0 | 230 | 439 | . | . | . | . |
| PM1548-SK1Org2 | ZNF648 | p.Pro423Ser | missense_variant | MODERATE | 0.0219298 | 0.00228311 | 228 | 438 | . | . | . | . |
| PM1548-SK1Org2 | C1orf27 | p.Thr381Ser | missense_variant | MODERATE | 0.38961 | 0 | 77 | 40 | . | . | . | . |
| PM1548-SK1Org2 | PLA2G4A | c.1033+2T>C | splice_donor_variant&intron_variant | HIGH | 0.604651 | 0 | 86 | 80 | . | . | . | . |
| PM1548-SK1Org2 | ASPM | p.Arg2700Gln | missense_variant | MODERATE | 0 | 0.00653595 | 81 | 153 | . | . | . | . |
| PM1548-SK1Org2 | DENND1B | c.83-20669G>A | intron_variant | MODIFIER | 0.467626 | 0 | 139 | 39 | . | . | . | . |
| PM1548-SK1Org2 | NR5A2 | p.Gly169Val | missense_variant | MODERATE | 0.514451 | 0 | 173 | 71 | . | . | . | . |
| PM1548-SK1Org2 | NAV1 | p.Thr719Ser | missense_variant | MODERATE | 0 | 0 | 113 | 176 | . | . | . | . |
| PM1548-SK1Org2 | NAV1 | p.Thr726Ser | missense_variant | MODERATE | 0.0251572 | 0.00930233 | 159 | 215 | . | . | . | . |
| PM1548-SK1Org2 | ATP2B4 | p.Asp867Asn | missense_variant&splice_region_variant | MODERATE | 0 | 0 | 95 | 121 | . | . | . | . |
| PM1548-SK1Org2 | NUAK2 | p.Arg378His | missense_variant | MODERATE | 0.46789 | 0 | 109 | 162 | . | . | . | . |
| PM1548-SK1Org2 | PFKFB2 | p.Glu261Ala | missense_variant | MODERATE | 0 | 0 | 143 | 81 | . | . | . | . |
| PM1548-SK1Org2 | CR2 | p.Thr209fs | frameshift_variant | HIGH | 0.233577 | 0 | 137 | 118 | . | . | . | . |
| PM1548-SK1Org2 | SMYD2 | p.Asn99fs | frameshift_variant | HIGH | 0.567164 | 0 | 268 | 180 | . | . | . | . |
| PM1548-SK1Org2 | HHIPL2 | p.Gly156Ser | missense_variant | MODERATE | 0.5 | 0 | 74 | 209 | . | . | . | . |
| PM1548-SK1Org2 | MIA3 | p.Arg1545Gln | missense_variant | MODERATE | 0.4875 | 0 | 160 | 38 | . | . | . | . |
| PM1548-SK1Org2 | LIN9 | p.Arg245Trp | missense_variant | MODERATE | 0.370968 | 0 | 62 | 86 | . | . | . | . |
| PM1548-SK1Org2 | ITPKB | p.Arg331His | missense_variant | MODERATE | 0.456204 | 0 | 274 | 395 | . | . | true | . |
| PM1548-SK1Org2 | ITPKB | p.Leu42Val | missense_variant | MODERATE | 0.0714286 | 0 | 14 | 35 | . | . | true | . |
| PM1548-SK1Org2 | SNAP47 | p.Phe343Leu | missense_variant | MODERATE | 0 | 0 | 180 | 78 | . | . | . | . |
| PM1548-SK1Org2 | IBA57 | p.Ala161Val | missense_variant | MODERATE | 0.270548 | 0 | 292 | 268 | . | . | . | . |
| PM1548-SK1Org2 | OBSCN | p.Tyr2035Phe | missense_variant | MODERATE | 0.0733138 | 0 | 341 | 508 | . | . | . | . |

| | | | | | | | | | | | | |
|---|---|---|---|---|---|---|---|---|---|---|---|---|
| PM1548-SK1Org2 | OBSCN | p.Gln2065Lys | missense_variant | MODERATE | 0.0266667 | 0 | 150 | 171 | . | . | . | . |
| PM1548-SK1Org2 | OBSCN | p.Cys2073Trp | missense_variant | MODERATE | 0 | 0 | 90 | 54 | . | . | . | . |
| PM1548-SK1Org2 | ACTA1 | p.Thr231Met | missense_variant | MODERATE | 0 | 0 | 98 | 311 | . | . | . | . |
| PM1548-SK1Org2 | GNPAT | p.Val551Met | missense_variant | MODERATE | 0.454545 | 0 | 165 | 79 | . | . | . | . |
| PM1548-SK1Org2 | DISC1 | p.Asp580Gly | missense_variant | MODERATE | 0.442804 | 0 | 271 | 167 | . | . | . | . |
| PM1548-SK1Org2 | GGPS1 | p.Val112Met | missense_variant | MODERATE | 0.00634921 | 0 | 315 | 405 | . | . | . | . |
| PM1548-SK1Org2 | NID1 | p.Pro918fs | frameshift_variant&splice_region_variant | HIGH | 0.416667 | 0 | 84 | 46 | . | . | . | . |
| PM1548-SK1Org2 | CEP170 | p.Glu1460Lys | missense_variant | MODERATE | 0.425197 | 0 | 127 | 32 | . | . | . | . |
| PM1548-SK1Org2 | ZNF496 | p.Arg606His | missense_variant | MODERATE | 0.494565 | 0 | 184 | 143 | . | . | . | . |
| PM1548-SK1Org2 | OR2T4 | p.Cys60Ser | missense_variant | MODERATE | 0.265306 | 0 | 49 | 107 | . | . | . | . |
| PM1548-SK1Org2 | OR2T6 | p.Tyr60His | missense_variant | MODERATE | 0.442623 | 0 | 122 | 68 | . | . | . | . |
| PM1548-SK1Org2 | DIP2C | p.Val1150Ile | missense_variant | MODERATE | 0 | 0 | 210 | 183 | . | . | . | . |
| PM1548-SK1Org2 | WDR37 | p.Pro169Ser | missense_variant | MODERATE | 0.512658 | 0 | 158 | 220 | . | . | . | . |
| PM1548-SK1Org2 | CALML5 | p.Glu102Lys | missense_variant | MODERATE | 0.479339 | 0 | 121 | 79 | . | . | . | . |
| PM1548-SK1Org2 | FRMD4A | p.Pro1005fs | frameshift_variant | HIGH | 0.00520833 | 0 | 192 | 118 | . | . | . | . |
| PM1548-SK1Org2 | CUBN | p.Trp3323Arg | missense_variant | MODERATE | 0.00425532 | 0 | 235 | 170 | . | . | . | . |
| PM1548-SK1Org2 | GPR158 | p.Arg448Gln | missense_variant | MODERATE | 0.545455 | 0 | 110 | 52 | . | . | . | . |
| PM1548-SK1Org2 | SVIL | p.Arg2075Cys | missense_variant | MODERATE | 0.48951 | 0 | 143 | 40 | . | . | . | . |
| PM1548-SK1Org2 | PARD3 | p.Arg1278Gly | missense_variant | MODERATE | 0 | 0 | 201 | 91 | . | . | . | . |
| PM1548-SK1Org2 | FRMPD2 | p.Pro226Leu | missense_variant | MODERATE | 0.48 | 0 | 200 | 178 | . | . | . | . |
| PM1548-SK1Org2 | PRKG1 | p.Asn601fs | frameshift_variant | HIGH | 0.425743 | 0 | 101 | 55 | . | . | . | . |
| PM1548-SK1Org2 | PBLD | p.Asp167Tyr | missense_variant | MODERATE | 0 | 0 | 133 | 186 | . | . | . | . |
| PM1548-SK1Org2 | DDX50 | p.Tyr39Cys | missense_variant | MODERATE | 0.341463 | 0 | 41 | 39 | . | . | . | . |
| PM1548-SK1Org2 | ADAMTS14 | p.Gly710Arg | missense_variant | MODERATE | 0.45977 | 0 | 87 | 78 | . | . | . | . |
| PM1548-SK1Org2 | DDIT4 | p.Ser46Cys | missense_variant | MODERATE | 0 | 0 | 192 | 263 | . | . | . | . |
| PM1548-SK1Org2 | DDIT4 | p.Thr47Ala | missense_variant | MODERATE | 0 | 0 | 192 | 263 | . | . | . | . |
| PM1548-SK1Org2 | KAT6B | p.Arg463His | missense_variant | MODERATE | 0.481013 | 0 | 158 | 104 | true | . | . | . |
| PM1548-SK1Org2 | KCNMA1 | p.Ala1039Val | missense_variant | MODERATE | 0.56044 | 0 | 91 | 73 | . | . | . | . |
| PM1548-SK1Org2 | ZMIZ1 | p.Asp1051fs | frameshift_variant | HIGH | 0.431818 | 0 | 176 | 67 | . | . | . | . |
| PM1548-SK1Org2 | MAT1A | p.Arg81Gln | missense_variant | MODERATE | 0.392157 | 0 | 153 | 225 | . | . | . | . |
| PM1548-SK1Org2 | TSPAN14 | p.Asp190Gly | missense_variant | MODERATE | 0 | 0 | 258 | 134 | . | . | . | . |
| PM1548-SK1Org2 | SH2D4B | p.Ala186Val | missense_variant | MODERATE | 0.493827 | 0 | 162 | 42 | . | . | . | . |
| PM1548-SK1Org2 | LRIT2 | p.Thr520fs | frameshift_variant | HIGH | 0.365169 | 0 | 178 | 224 | . | . | . | . |
| PM1548-SK1Org2 | CCSER2 | c.2326-13521C> | intron_variant | MODIFIER | 0.0588235 | 0 | 170 | 179 | . | . | . | . |
| PM1548-SK1Org2 | CYP26C1 | p.Ser70Leu | missense_variant | MODERATE | 0 | 0 | 37 | 149 | . | . | . | . |
| PM1548-SK1Org2 | MYOF | p.Arg726Gln | missense_variant | MODERATE | 0.617021 | 0 | 141 | 43 | . | . | . | . |
| PM1548-SK1Org2 | SORBS1 | p.Arg585Cys | missense_variant | MODERATE | 0.5 | 0 | 16 | 42 | . | . | . | . |
| PM1548-SK1Org2 | ALDH18A1 | p.Arg622Trp | missense_variant | MODERATE | 0.435294 | 0 | 170 | 86 | . | . | . | . |
| PM1548-SK1Org2 | ENTPD1 | p.Arg252Leu | missense_variant | MODERATE | 0 | 0 | 241 | 84 | . | . | . | . |
| PM1548-SK1Org2 | MORN4 | p.Leu202Ile | missense_variant | MODERATE | 0 | 0 | 146 | 87 | . | . | . | . |
| PM1548-SK1Org2 | PKD2L1 | p.Arg206Cys | missense_variant | MODERATE | 0.56129 | 0 | 155 | 130 | . | . | . | . |
| PM1548-SK1Org2 | MRPL43 | p.Asn66Ser | missense_variant | MODERATE | 0.529801 | 0 | 151 | 81 | . | . | . | . |
| PM1548-SK1Org2 | POLL | p.Arg419* | stop_gained | HIGH | 0 | 0 | 106 | 44 | . | . | . | . |
| PM1548-SK1Org2 | TAF5 | p.Phe374fs | frameshift_variant | HIGH | 0.666667 | 0 | 36 | 49 | . | . | . | . |
| PM1548-SK1Org2 | WDR96 | p.Val70Ala | missense_variant | MODERATE | 0.417582 | 0 | 91 | 59 | . | . | . | . |
| PM1548-SK1Org2 | SORCS3 | p.Lys1023Gln | missense_variant | MODERATE | 0.0201005 | 0 | 199 | 58 | . | . | . | . |
| PM1548-SK1Org2 | TCF7L2 | p.Ser587Leu | missense_variant | MODERATE | 0.00555556 | 0 | 180 | 171 | true | . | true | . |
| PM1548-SK1Org2 | ABLIM1 | p.Leu633Met | missense_variant | MODERATE | 0 | 0 | 220 | 266 | . | . | . | . |
| PM1548-SK1Org2 | GFRA1 | p.Ala173Val | missense_variant | MODERATE | 0.41 | 0 | 100 | 73 | . | . | . | . |
| PM1548-SK1Org2 | KIAA1598 | p.Arg482His | missense_variant | MODERATE | 0.362832 | 0 | 113 | 181 | true | . | . | . |
| PM1548-SK1Org2 | VAX1 | p.Cys200* | stop_gained | HIGH | 0 | 0 | 26 | 96 | . | . | . | . |
| PM1548-SK1Org2 | EIF3A | p.His257His | synonymous_variant | LOW | 0.0454545 | 0 | 264 | 131 | . | . | . | . |
| PM1548-SK1Org2 | ATE1 | p.Leu289Gln | missense_variant | MODERATE | 0.027027 | 0 | 74 | 77 | . | . | . | . |
| PM1548-SK1Org2 | PLEKHA1 | p.Arg9His | missense_variant | MODERATE | 0.294737 | 0 | 95 | 43 | . | . | . | . |
| PM1548-SK1Org2 | PLEKHA1 | p.Val404Met | missense_variant | MODERATE | 0.25 | 0 | 100 | 76 | . | . | . | . |
| PM1548-SK1Org2 | HMX3 | p.Pro23fs | frameshift_variant | HIGH | 0.560976 | 0 | 41 | 56 | . | . | . | . |
| PM1548-SK1Org2 | CLRN3 | p.Pro71Thr | missense_variant | MODERATE | 0 | 0.00613497 | 68 | 163 | . | . | . | . |
| PM1548-SK1Org2 | MKI67 | p.Arg2258* | stop_gained | HIGH | 0.516556 | 0 | 151 | 90 | . | . | . | . |
| PM1548-SK1Org2 | TTC40 | p.Thr1864Met | missense_variant | MODERATE | 0.490196 | 0 | 102 | 72 | . | . | . | . |
| PM1548-SK1Org2 | KNDC1 | p.Cys79fs | frameshift_variant | HIGH | 0.398601 | 0 | 143 | 162 | . | . | . | . |
| PM1548-SK1Org2 | PIDD | p.Ala727Glu | missense_variant | MODERATE | 0 | 0 | 65 | 108 | . | . | . | . |
| PM1548-SK1Org2 | MUC6 | p.Thr1861Ala | missense_variant | MODERATE | 0.213198 | 0 | 197 | 78 | . | . | . | . |
| PM1548-SK1Org2 | MUC6 | p.Ile1857Thr | missense_variant | MODERATE | 0.287234 | 0 | 188 | 78 | . | . | . | . |
| PM1548-SK1Org2 | MUC6 | p.His1752Gln | missense_variant | MODERATE | 0 | 0 | 103 | 31 | . | . | . | . |
| PM1548-SK1Org2 | MUC6 | p.Thr1711Ile | missense_variant | MODERATE | 0.0344828 | 0 | 319 | 51 | . | . | . | . |
| PM1548-SK1Org2 | MUC2 | p.Gly1331Arg | missense_variant | MODERATE | 0.357143 | 0 | 168 | 70 | . | . | . | . |
| PM1548-SK1Org2 | MUC5B | p.Ala110Thr | missense_variant | MODERATE | 0.4875 | 0 | 80 | 216 | . | . | . | . |
| PM1548-SK1Org2 | BRSK2 | p.Pro440fs | frameshift_variant | HIGH | 0.558011 | 0 | 181 | 593 | . | . | . | . |
| PM1548-SK1Org2 | TSPAN32 | p.Ala178Val | missense_variant | MODERATE | 0.533333 | 0 | 120 | 69 | . | . | . | . |
| PM1548-SK1Org2 | TSSC4 | p.Arg92Cys | missense_variant | MODERATE | 0.439353 | 0 | 371 | 524 | . | . | . | . |
| PM1548-SK1Org2 | OSBPL5 | p.Thr843Met | missense_variant | MODERATE | 0 | 0 | 129 | 106 | . | . | . | . |
| PM1548-SK1Org2 | STIM1 | p.Lys371Arg | missense_variant | MODERATE | 0.0138408 | 0 | 289 | 322 | . | . | . | . |
| PM1548-SK1Org2 | OR51E1 | p.Tyr63His | missense_variant | MODERATE | 0.17033 | 0 | 182 | 116 | . | . | . | . |
| PM1548-SK1Org2 | OR51A2 | p.Lys289Asn | missense_variant | MODERATE | 0.352459 | 0 | 122 | 30 | . | . | . | . |
| PM1548-SK1Org2 | SOX6 | p.Met803Ile | missense_variant | MODERATE | 0.0238095 | 0 | 84 | 33 | . | . | . | . |
| PM1548-SK1Org2 | SOX6 | p.Met803Thr | missense_variant | MODERATE | 0.0119048 | 0 | 84 | 33 | . | . | . | . |
| PM1548-SK1Org2 | SOX6 | p.Met803Val | missense_variant | MODERATE | 0.0116279 | 0 | 86 | 33 | . | . | . | . |
| PM1548-SK1Org2 | ABCC8 | p.Asp761Asn | missense_variant | MODERATE | 0.231214 | 0 | 346 | 231 | . | . | . | . |
| PM1548-SK1Org2 | LDHC | p.Lys5Glu | missense_variant | MODERATE | 0.446602 | 0 | 206 | 119 | . | . | . | . |
| PM1548-SK1Org2 | MRGPRX2 | p.Asn62Asn | synonymous_variant | LOW | 0.258242 | 0 | 182 | 83 | . | . | . | . |
| PM1548-SK1Org2 | ZDHHC13 | p.Gly591Asp | missense_variant | MODERATE | 0.254237 | 0 | 118 | 32 | . | . | . | . |
| PM1548-SK1Org2 | ANO3 | p.Leu273Pro | missense_variant | MODERATE | 0.261538 | 0 | 325 | 432 | . | . | . | . |
| PM1548-SK1Org2 | HIPK3 | p.Asp532Ala | missense_variant | MODERATE | 0.0538462 | 0 | 130 | 54 | . | . | . | . |
| PM1548-SK1Org2 | ABTB2 | p.Asp793Asn | missense_variant | MODERATE | 0.571429 | 0.00668896 | 224 | 299 | . | . | . | . |
| PM1548-SK1Org2 | RAG1 | p.Arg507Trp | missense_variant | MODERATE | 0.295775 | 0 | 142 | 86 | . | . | . | true |
| PM1548-SK1Org2 | TP53I11 | p.Leu62Arg | missense_variant | MODERATE | 0 | 0 | 71 | 34 | . | . | . | . |
| PM1548-SK1Org2 | MAPK8IP1 | p.Thr321Met | missense_variant | MODERATE | 0.545455 | 0 | 176 | 188 | . | true | . | . |
| PM1548-SK1Org2 | DGKZ | n.46369267G>A | intragenic_variant | MODIFIER | 0.347826 | 0 | 23 | 32 | . | . | . | . |
| PM1548-SK1Org2 | DGKZ | p.Asp574Asn | missense_variant | MODERATE | 0.563758 | 0 | 149 | 74 | . | . | . | . |
| PM1548-SK1Org2 | ACP2 | p.Arg231His | missense_variant | MODERATE | 0.513699 | 0 | 146 | 199 | . | . | . | . |
| PM1548-SK1Org2 | MYBPC3 | p.Arg654Ser | missense_variant | MODERATE | 0.00990099 | 0 | 202 | 97 | . | . | . | . |
| PM1548-SK1Org2 | OR5D14 | p.Tyr261Cys | missense_variant | MODERATE | 0.255474 | 0 | 137 | 70 | . | . | . | . |
| PM1548-SK1Org2 | OR5W2 | p.Ala29Ser | missense_variant | MODERATE | 0.513514 | 0 | 111 | 99 | . | . | . | . |
| PM1548-SK1Org2 | APLNR | p.Ala161Thr | missense_variant | MODERATE | 0.288235 | 0 | 170 | 160 | . | . | . | . |
| PM1548-SK1Org2 | OR10Q1 | p.Thr287Ile | missense_variant | MODERATE | 0.490909 | 0 | 110 | 89 | . | . | . | . |
| PM1548-SK1Org2 | DTX4 | p.Lys396fs | frameshift_variant | HIGH | 0.535032 | 0 | 157 | 34 | . | . | . | . |
| PM1548-SK1Org2 | DTX4 | p.Gly510fs | frameshift_variant | HIGH | 0.529183 | 0.00719424 | 257 | 139 | . | . | . | . |
| PM1548-SK1Org2 | DAK | p.Arg380Gln | missense_variant | MODERATE | 0.560185 | 0 | 216 | 316 | . | . | . | . |
| PM1548-SK1Org2 | FEN1 | p.Arg332Cys | missense_variant | MODERATE | 0 | 0 | 180 | 80 | . | . | . | . |
| PM1548-SK1Org2 | BEST1 | p.Ser413Asn | missense_variant | MODERATE | 0.456376 | 0 | 149 | 159 | . | . | . | . |
| PM1548-SK1Org2 | AHNAK | p.Ser4908Leu | missense_variant | MODERATE | 0.421875 | 0 | 64 | 39 | . | . | . | . |
| PM1548-SK1Org2 | AHNAK | p.His4772Asn | missense_variant | MODERATE | 0 | 0 | 80 | 54 | . | . | . | . |
| PM1548-SK1Org2 | AHNAK | p.Gly4561Asp | missense_variant | MODERATE | 0.0277778 | 0 | 144 | 206 | . | . | . | . |
| PM1548-SK1Org2 | AHNAK | p.Val4560Leu | missense_variant | MODERATE | 0.025974 | 0 | 154 | 203 | . | . | . | . |
| PM1548-SK1Org2 | AHNAK | p.Val3640Ile | missense_variant | MODERATE | 0.152174 | 0 | 184 | 44 | . | . | . | . |
| PM1548-SK1Org2 | AHNAK | p.Asn3637Lys | missense_variant | MODERATE | 0.119318 | 0 | 176 | 44 | . | . | . | . |
| PM1548-SK1Org2 | AHNAK | p.Lys2706fs | frameshift_variant | HIGH | 0.509804 | 0 | 51 | 43 | . | . | . | . |
| PM1548-SK1Org2 | AHNAK | p.Thr1986Ala | missense_variant | MODERATE | 0 | 0 | 297 | 324 | . | . | . | . |
| PM1548-SK1Org2 | CHRM1 | p.Pro319fs | frameshift_variant | HIGH | 0 | 0 | 106 | 31 | . | . | . | . |
| PM1548-SK1Org2 | SLC22A6 | p.Val240Met | missense_variant | MODERATE | 0 | 0 | 186 | 227 | . | . | . | . |
| PM1548-SK1Org2 | SLC22A25 | p.Asn62Asp | missense_variant | MODERATE | 0 | 0 | 179 | 110 | . | . | . | . |
| PM1548-SK1Org2 | C11orf95 | n.1245C>T | non_coding_exon_variant | MODIFIER | 0 | 0 | 5 | 34 | . | . | . | . |
| PM1548-SK1Org2 | NRXN2 | p.Asp277Asn | missense_variant | MODERATE | 0.179775 | 0 | 178 | 229 | . | . | . | . |
| PM1548-SK1Org2 | SNX15 | p.Leu10Met | missense_variant | MODERATE | 0 | 0 | 44 | 37 | . | . | . | . |
| PM1548-SK1Org2 | RBM4 | p.Tyr303His | missense_variant | MODERATE | 0.0233463 | 0 | 257 | 269 | . | . | . | . |
| PM1548-SK1Org2 | SPTBN2 | p.Cys1393Ser | missense_variant | MODERATE | 0.0115607 | 0 | 173 | 300 | . | . | . | . |
| PM1548-SK1Org2 | SPTBN2 | p.Arg465Gln | missense_variant | MODERATE | 0.340351 | 0 | 285 | 212 | . | . | . | . |
| PM1548-SK1Org2 | LRP5 | p.Ala724Val | missense_variant | MODERATE | 0.445783 | 0 | 83 | 227 | . | . | . | . |
| PM1548-SK1Org2 | IGHMBP2 | p.Pro809Ser | missense_variant | MODERATE | 0.657895 | 0 | 76 | 132 | . | . | . | . |
| PM1548-SK1Org2 | TPCN2 | p.Gly492Val | missense_variant | MODERATE | 0 | 0 | 41 | 21 | . | . | . | . |
| PM1548-SK1Org2 | CCND1 | p.Val94Leu | missense_variant | MODERATE | 0.0731707 | 0 | 82 | 259 | true | true | true | . |
| PM1548-SK1Org2 | INPPL1 | p.Leu957Phe | missense_variant | MODERATE | 0.0134228 | 0 | 149 | 30 | . | . | . | . |
| PM1548-SK1Org2 | PDE2A | p.His99fs | frameshift_variant | HIGH | 0.554545 | 0 | 220 | 161 | . | . | . | . |
| PM1548-SK1Org2 | RNF169 | p.Glu131* | stop_gained | HIGH | 0 | 0 | 17 | 52 | . | . | . | . |
| PM1548-SK1Org2 | TENM4 | p.Ala235Thr | missense_variant | MODERATE | 0.439394 | 0 | 132 | 201 | . | . | . | . |
| PM1548-SK1Org2 | PCF11 | p.Leu745Leu | synonymous_variant | LOW | 0 | 0 | 86 | 203 | . | . | . | . |
| PM1548-SK1Org2 | PCF11 | p.Ser756Gly | missense_variant | MODERATE | 0 | 0 | 115 | 215 | . | . | . | . |

| | | | | | | | | | | | | |
|---|---|---|---|---|---|---|---|---|---|---|---|---|
| PM1548-SK1Org2 | PCF11 | p.Arg833Trp | missense_variant | MODERATE | 0.355828 | 0 | 163 | 258 | . | . | . | . |
| PM1548-SK1Org2 | TYR | p.Gly191fs | frameshift_variant | HIGH | 0.492308 | 0 | 195 | 94 | . | . | . | . |
| PM1548-SK1Org2 | FAT3 | p.Ala2495Ser | missense_variant | MODERATE | 0 | 0 | 319 | 229 | . | . | . | . |
| PM1548-SK1Org2 | PANX1 | p.Ser250Thr | missense_variant | MODERATE | 0.496241 | 0 | 266 | 235 | . | . | . | . |
| PM1548-SK1Org2 | MTMR2 | p.Lys451Glu | missense_variant | MODERATE | 0 | 0 | 133 | 77 | . | . | . | . |
| PM1548-SK1Org2 | MTMR2 | p.Asp436Asn | missense_variant | MODERATE | 0.0196078 | 0 | 102 | 75 | . | . | . | . |
| PM1548-SK1Org2 | MTMR2 | p.Leu433Val | missense_variant | MODERATE | 0 | 0 | 130 | 282 | . | . | . | . |
| PM1548-SK1Org2 | MMP3 | p.Asp200Gly | missense_variant | MODERATE | 0 | 0 | 93 | 91 | . | . | . | . |
| PM1548-SK1Org2 | DYNC2H1 | p.Tyr3523His | missense_variant | MODERATE | 0.473118 | 0 | 93 | 62 | . | . | . | . |
| PM1548-SK1Org2 | CASP1 | p.Phe262Tyr | missense_variant | MODERATE | 0.441489 | 0 | 188 | 177 | . | . | . | . |
| PM1548-SK1Org2 | KBTBD3 | p.Lys478fs | frameshift_variant | HIGH | 0.4 | 0 | 40 | 50 | . | . | . | . |
| PM1548-SK1Org2 | NXPE1 | p.Thr169Ile | missense_variant | MODERATE | 0 | 0 | 65 | 97 | . | . | . | . |
| PM1548-SK1Org2 | NXPE1 | p.Met157Ile | missense_variant | MODERATE | 0.043956 | 0 | 91 | 97 | . | . | . | . |
| PM1548-SK1Org2 | NXPE1 | p.Met157Thr | missense_variant | MODERATE | 0.043956 | 0 | 91 | 96 | . | . | . | . |
| PM1548-SK1Org2 | NXPE1 | p.Thr149Lys | missense_variant | MODERATE | 0.025 | 0 | 120 | 118 | . | . | . | . |
| PM1548-SK1Org2 | DSCAML1 | p.Val1762Ile | missense_variant | MODERATE | 0.47449 | 0 | 196 | 226 | . | . | . | . |
| PM1548-SK1Org2 | DSCAML1 | p.Asp949fs | frameshift_variant | HIGH | 0.414365 | 0 | 181 | 244 | . | . | . | . |
| PM1548-SK1Org2 | ATP5L | p.Leu84Trp | missense_variant | MODERATE | 0.0814815 | 0 | 135 | 125 | . | . | . | . |
| PM1548-SK1Org2 | ATP5L | p.Val89Ile | missense_variant | MODERATE | 0.0729927 | 0 | 137 | 140 | . | . | . | . |
| PM1548-SK1Org2 | ATP5L | p.Ile99Val | missense_variant | MODERATE | 0.0437956 | 0 | 137 | 141 | . | . | . | . |
| PM1548-SK1Org2 | PHLDB1 | p.Ala932Ser | missense_variant | MODERATE | 0.0330579 | 0 | 121 | 223 | . | . | . | . |
| PM1548-SK1Org2 | PHLDB1 | p.Pro938Leu | missense_variant | MODERATE | 0.017094 | 0 | 117 | 220 | . | . | . | . |
| PM1548-SK1Org2 | CBL | p.Ser237Leu | missense_variant | MODERATE | 0.533333 | 0 | 90 | 128 | true | . | . | . |
| PM1548-SK1Org2 | CBL | p.Pro504Gln | missense_variant | MODERATE | 0.00478469 | 0 | 209 | 111 | true | . | . | . |
| PM1548-SK1Org2 | CRTAM | p.Leu60fs | frameshift_variant | HIGH | 0.1875 | 0 | 32 | 170 | . | . | . | . |
| PM1548-SK1Org2 | OR10G8 | p.Lys265Glu | missense_variant | MODERATE | 0.111628 | 0 | 215 | 232 | . | . | . | . |
| PM1548-SK1Org2 | OR10G8 | p.Lys265Asn | missense_variant | MODERATE | 0.166667 | 0 | 144 | 193 | . | . | . | . |
| PM1548-SK1Org2 | OR8G2 | p.Val2Gly | missense_variant | MODERATE | 0.488889 | 0 | 45 | 100 | . | . | . | . |
| PM1548-SK1Org2 | ROBO4 | p.Arg776His | missense_variant | MODERATE | 0.573333 | 0 | 225 | 263 | . | . | . | . |
| PM1548-SK1Org2 | ADAMTS8 | p.Arg9Gly | missense_variant | MODERATE | 0 | 0 | 6 | 48 | . | . | . | true |
| PM1548-SK1Org2 | IQSEC3 | p.Gln1075Lys | missense_variant | MODERATE | 0 | 0 | 76 | 59 | . | . | . | . |
| PM1548-SK1Org2 | FOXM1 | p.Leu334Ser | missense_variant | MODERATE | 0.013245 | 0 | 151 | 177 | . | . | . | . |
| PM1548-SK1Org2 | FOXM1 | p.Asp328Glu | missense_variant | MODERATE | 0.012987 | 0 | 154 | 203 | . | . | . | . |
| PM1548-SK1Org2 | VWF | p.Ala865Asp | missense_variant | MODERATE | 0.24186 | 0 | 215 | 226 | . | . | . | . |
| PM1548-SK1Org2 | IFFO1 | p.Val126Met | missense_variant | MODERATE | 0.489796 | 0 | 49 | 191 | . | . | . | . |
| PM1548-SK1Org2 | PHB2 | p.Arg270Arg | synonymous_variant | LOW | 0.027027 | 0 | 296 | 405 | . | . | . | . |
| PM1548-SK1Org2 | PHB2 | p.Asn269Asn | synonymous_variant | LOW | 0.027027 | 0 | 296 | 404 | . | . | . | . |
| PM1548-SK1Org2 | TAS2R10 | p.Ser41Thr | missense_variant | MODERATE | 0.496774 | 0 | 155 | 57 | . | . | . | . |
| PM1548-SK1Org2 | TAS2R31 | p.Ala177Thr | missense_variant | MODERATE | 0.110092 | 0 | 109 | 47 | . | . | . | . |
| PM1548-SK1Org2 | PLCZ1 | p.Gly133Val | missense_variant | MODERATE | 0 | 0 | 69 | 42 | . | . | . | . |
| PM1548-SK1Org2 | PDE3A | p.Glu171fs | frameshift_variant | HIGH | 0.208333 | 0 | 72 | 56 | . | . | . | . |
| PM1548-SK1Org2 | TM7SF3 | c.869-699C>G | intron_variant | MODIFIER | 0.470588 | 0 | 85 | 125 | . | . | . | . |
| PM1548-SK1Org2 | DDX11 | p.Asp935Asn | missense_variant | MODERATE | 0.542373 | 0 | 59 | 35 | . | . | . | . |
| PM1548-SK1Org2 | COL2A1 | p.Asp682Asn | missense_variant | MODERATE | 0.49569 | 0 | 232 | 53 | true | . | . | . |
| PM1548-SK1Org2 | KMT2D | p.Ala4804Val | missense_variant | MODERATE | 0.46988 | 0.0060241 | 249 | 166 | true | true | true | true |
| PM1548-SK1Org2 | KMT2D | p.Leu4214Leu | synonymous_variant | LOW | 0.0078125 | 0 | 256 | 376 | true | true | true | true |
| PM1548-SK1Org2 | KMT2D | p.Thr3166Ile | missense_variant | MODERATE | 0.530612 | 0 | 147 | 193 | true | true | true | true |
| PM1548-SK1Org2 | KMT2D | p.Thr2581Ala | missense_variant | MODERATE | 0.0305344 | 0 | 131 | 134 | true | true | true | true |
| PM1548-SK1Org2 | KMT2D | p.Gly1628fs | frameshift_variant | HIGH | 0.0316206 | 0 | 253 | 136 | true | true | true | true |
| PM1548-SK1Org2 | PRPH | p.Ala348Thr | missense_variant | MODERATE | 0.482517 | 0 | 143 | 109 | . | . | . | . |
| PM1548-SK1Org2 | FAM186B | p.Leu494Met | missense_variant | MODERATE | 0.53304 | 0 | 227 | 412 | . | . | . | . |
| PM1548-SK1Org2 | ANKRD33 | p.Ala173Val | missense_variant | MODERATE | 0.394958 | 0 | 119 | 241 | . | . | . | . |
| PM1548-SK1Org2 | KRT74 | p.Gln440His | missense_variant | MODERATE | 0 | 0 | 40 | 34 | . | . | . | . |
| PM1548-SK1Org2 | PCBP2 | p.Pro175fs | frameshift_variant | HIGH | 0.571429 | 0 | 91 | 35 | . | . | . | . |
| PM1548-SK1Org2 | HNRNPA1 | p.Thr61Ala | missense_variant | MODERATE | 0.507937 | 0 | 189 | 168 | . | . | . | . |
| PM1548-SK1Org2 | ITGA5 | p.Arg193His | missense_variant | MODERATE | 0.220183 | 0 | 109 | 104 | . | . | . | . |
| PM1548-SK1Org2 | MYO1A | p.Arg559Cys | missense_variant | MODERATE | 0.486607 | 0 | 224 | 190 | . | . | . | . |
| PM1548-SK1Org2 | NAB2 | p.Pro388Thr | missense_variant | MODERATE | 0.00497512 | 0 | 201 | 78 | true | . | . | . |
| PM1548-SK1Org2 | NAB2 | p.Pro388His | missense_variant | MODERATE | 0.00497512 | 0 | 201 | 78 | true | . | . | . |
| PM1548-SK1Org2 | R3HDM2 | p.Arg956His | missense_variant | MODERATE | 0 | 0 | 109 | 33 | . | . | . | . |
| PM1548-SK1Org2 | CTDSP2 | p.Glu259Asp | missense_variant | MODERATE | 0.00904977 | 0 | 221 | 90 | . | . | . | . |
| PM1548-SK1Org2 | CTDSP2 | p.Ala258Thr | missense_variant | MODERATE | 0.00909091 | 0 | 220 | 90 | . | . | . | . |
| PM1548-SK1Org2 | CTDSP2 | p.Ile251Val | missense_variant | MODERATE | 0.01 | 0 | 200 | 94 | . | . | . | . |
| PM1548-SK1Org2 | PTPRB | p.Val1992Ala | missense_variant | MODERATE | 0.0111732 | 0 | 179 | 51 | true | . | . | . |
| PM1548-SK1Org2 | NAV3 | p.Met1275Val | missense_variant | MODERATE | 0 | 0 | 92 | 67 | . | true | . | . |
| PM1548-SK1Org2 | NAV3 | p.Pro1276Leu | missense_variant | MODERATE | 0 | 0 | 111 | 130 | . | true | . | . |
| PM1548-SK1Org2 | NAV3 | p.Thr1281Ser | missense_variant | MODERATE | 0 | 0 | 130 | 146 | . | true | . | . |
| PM1548-SK1Org2 | TMTC2 | p.Gly143Arg | missense_variant | MODERATE | 0.535714 | 0 | 84 | 67 | . | . | . | . |
| PM1548-SK1Org2 | ALX1 | p.Thr68Ser | missense_variant | MODERATE | 0.419355 | 0 | 217 | 191 | . | . | . | . |
| PM1548-SK1Org2 | FGD6 | p.Met965fs | frameshift_variant | HIGH | 0.492958 | 0 | 71 | 56 | . | . | . | . |
| PM1548-SK1Org2 | APAF1 | p.Gln578Arg | missense_variant | MODERATE | 0 | 0 | 153 | 126 | . | . | . | . |
| PM1548-SK1Org2 | SLC41A2 | p.Pro266Ser | missense_variant | MODERATE | 0.402439 | 0 | 82 | 65 | . | . | . | . |
| PM1548-SK1Org2 | PRDM4 | p.Glu313Glu | synonymous_variant | LOW | 0.0429448 | 0 | 163 | 274 | . | . | . | . |
| PM1548-SK1Org2 | TMEM119 | p.Thr81fs | frameshift_variant | HIGH | 0.209524 | 0 | 210 | 297 | . | . | . | . |
| PM1548-SK1Org2 | FOXN4 | p.Ala462Thr | missense_variant | MODERATE | 0.531429 | 0.00490196 | 175 | 204 | . | . | . | . |
| PM1548-SK1Org2 | FAM222A | p.Pro392Thr | missense_variant | MODERATE | 0.0689655 | 0 | 87 | 422 | . | . | . | . |
| PM1548-SK1Org2 | FAM222A | p.Ser400Ala | missense_variant | MODERATE | 0.0576923 | 0 | 104 | 423 | . | . | . | . |
| PM1548-SK1Org2 | TCHP | p.Ala371Thr | missense_variant | MODERATE | 0.572327 | 0 | 159 | 45 | . | . | . | . |
| PM1548-SK1Org2 | SH2B3 | p.Glu523fs | frameshift_variant | HIGH | 0 | 0 | 174 | 158 | true | . | . | . |
| PM1548-SK1Org2 | HECTD4 | p.Ala3707Val | missense_variant | MODERATE | 0.6 | 0 | 90 | 62 | . | . | . | . |
| PM1548-SK1Org2 | TPCN1 | p.Leu234Arg | missense_variant | MODERATE | 0.530806 | 0 | 211 | 134 | . | . | . | . |
| PM1548-SK1Org2 | RBM19 | p.Asn533Asn | synonymous_variant | LOW | 0.483333 | 0 | 120 | 250 | . | . | . | . |
| PM1548-SK1Org2 | RAB35 | n.120541738C>T | intragenic_variant | MODIFIER | 0.464135 | 0 | 237 | 243 | . | . | . | . |
| PM1548-SK1Org2 | GCN1L1 | p.Ser2359Pro | missense_variant | MODERATE | 0.502857 | 0 | 175 | 83 | . | . | . | . |
| PM1548-SK1Org2 | SETD1B | p.His8fs | frameshift_variant | HIGH | 0.0606061 | 0 | 99 | 141 | . | . | . | . |
| PM1548-SK1Org2 | SETD1B | p.Pro1361fs | frameshift_variant | HIGH | 0.517544 | 0 | 114 | 190 | . | . | . | . |
| PM1548-SK1Org2 | ZCCHC8 | p.Gly551Asp | missense_variant | MODERATE | 0.2375 | 0 | 80 | 34 | true | . | . | . |
| PM1548-SK1Org2 | ABCB9 | p.His709Gln | missense_variant | MODERATE | 0.0123839 | 0 | 323 | 333 | . | . | . | . |
| PM1548-SK1Org2 | ABCB9 | p.His709Arg | missense_variant | MODERATE | 0.0124224 | 0 | 322 | 333 | . | . | . | . |
| PM1548-SK1Org2 | ABCB9 | p.Lys694Arg | missense_variant | MODERATE | 0.0150376 | 0 | 266 | 301 | . | . | . | . |
| PM1548-SK1Org2 | PITPNM2 | p.Arg749Trp | missense_variant | MODERATE | 0.50838 | 0 | 179 | 160 | . | . | . | . |
| PM1548-SK1Org2 | SBNO1 | p.Arg1283Cys | missense_variant&splice_region_variant | MODERATE | 0.510791 | 0 | 139 | 244 | . | . | . | . |
| PM1548-SK1Org2 | DNAH10 | p.Pro3133His | missense_variant | MODERATE | 0.301676 | 0 | 179 | 220 | . | . | . | . |
| PM1548-SK1Org2 | TMEM132B | p.Ala297Thr | missense_variant | MODERATE | 0 | 0 | 139 | 79 | . | . | . | . |
| PM1548-SK1Org2 | GPR133 | p.Gly731Arg | missense_variant | MODERATE | 0.464286 | 0 | 196 | 110 | . | . | . | . |
| PM1548-SK1Org2 | FBRSL1 | p.Glu251Asp | missense_variant | MODERATE | 0 | 0 | 34 | 153 | . | . | . | . |
| PM1548-SK1Org2 | MPHOSPH8 | p.Val686Ile | missense_variant | MODERATE | 0 | 0 | 314 | 75 | . | . | . | . |
| PM1548-SK1Org2 | GJA3 | p.Ala213Thr | missense_variant | MODERATE | 0.269231 | 0 | 52 | 151 | . | . | . | . |
| PM1548-SK1Org2 | LATS2 | p.Pro41Gln | missense_variant | MODERATE | 0.526066 | 0 | 211 | 38 | . | . | . | . |
| PM1548-SK1Org2 | CDK8 | p.Asn372Thr | missense_variant | MODERATE | 0.0363636 | 0 | 165 | 249 | . | . | . | . |
| PM1548-SK1Org2 | POLR1D | n.28240022G>A | intragenic_variant | MODIFIER | 0.542857 | 0 | 210 | 101 | . | . | . | . |
| PM1548-SK1Org2 | FRY | p.Gln2970His | missense_variant | MODERATE | 0.00990099 | 0 | 101 | 39 | . | . | . | . |
| PM1548-SK1Org2 | BRCA2 | p.Asn863fs | frameshift_variant | HIGH | 0.276596 | 0 | 94 | 84 | true | true | . | true |
| PM1548-SK1Org2 | STARD13 | p.Arg155His | missense_variant | MODERATE | 0 | 0 | 142 | 32 | . | . | . | . |
| PM1548-SK1Org2 | MAB21L1 | p.Ser103Ser | synonymous_variant | LOW | 0.446809 | 0 | 94 | 99 | . | . | . | . |
| PM1548-SK1Org2 | FOXO1 | p.Ser152Thr | missense_variant | MODERATE | 0.0909091 | 0 | 22 | 140 | true | . | . | . |
| PM1548-SK1Org2 | KBTBD7 | p.Tyr195Phe | missense_variant | MODERATE | 0.0467836 | 0 | 171 | 644 | . | . | . | . |
| PM1548-SK1Org2 | GPALPP1 | p.Pro37Arg | missense_variant | MODERATE | 1 | 0 | 61 | 103 | . | . | . | . |
| PM1548-SK1Org2 | FAM124A | p.Ser257Pro | missense_variant | MODERATE | 0 | 0 | 108 | 101 | . | . | . | . |
| PM1548-SK1Org2 | NEK3 | p.Asn343Lys | missense_variant | MODERATE | 0.0384615 | 0 | 26 | 44 | . | . | . | . |
| PM1548-SK1Org2 | DACH1 | p.Arg196Arg | synonymous_variant | LOW | 0.243902 | 0 | 41 | 115 | . | . | . | . |
| PM1548-SK1Org2 | ABCC4 | p.Ile1219fs | frameshift_variant | HIGH | 0 | 0 | 68 | 130 | . | . | . | . |
| PM1548-SK1Org2 | FGF14 | p.Asp33Gly | missense_variant | MODERATE | 0.0410959 | 0 | 219 | 97 | . | . | . | . |
| PM1548-SK1Org2 | CCDC168 | p.Met4045fs | frameshift_variant | HIGH | 0.462585 | 0 | 147 | 95 | . | . | . | . |
| PM1548-SK1Org2 | ARHGEF7 | p.Thr23fs | frameshift_variant | HIGH | 0.186667 | 0 | 75 | 189 | . | . | . | . |
| PM1548-SK1Org2 | MCF2L | p.Arg590Gln | missense_variant | MODERATE | 0.5 | 0 | 72 | 34 | . | . | . | . |
| PM1548-SK1Org2 | CDC16 | p.Asn557fs | frameshift_variant | HIGH | 0 | 0 | 218 | 90 | . | . | . | . |
| PM1548-SK1Org2 | UPF3A | p.Asp129Glu | missense_variant | MODERATE | 0.392857 | 0 | 56 | 60 | . | . | . | . |
| PM1548-SK1Org2 | OR11G2 | p.Phe141Phe | synonymous_variant | LOW | 0.0533981 | 0 | 206 | 117 | . | . | . | . |
| PM1548-SK1Org2 | ACIN1 | p.Glu192Asp | missense_variant | MODERATE | 0.0444444 | 0 | 90 | 129 | . | . | . | . |
| PM1548-SK1Org2 | ACIN1 | p.Glu192Lys | missense_variant | MODERATE | 0.0444444 | 0 | 90 | 129 | . | . | . | . |
| PM1548-SK1Org2 | PABPN1 | c.*5dupA | frameshift_variant&stop_retained_variant | HIGH | 0 | 0 | 40 | 92 | . | . | . | . |
| PM1548-SK1Org2 | RNF31 | p.Ala9Gly | missense_variant | MODERATE | 0.222222 | 0 | 18 | 104 | . | . | . | true |
| PM1548-SK1Org2 | RNF31 | p.Val12Ala | missense_variant | MODERATE | 0.222222 | 0 | 18 | 104 | . | . | . | true |
| PM1548-SK1Org2 | LTB4R2 | p.Ala92Val | missense_variant | MODERATE | 0.266667 | 0 | 30 | 173 | . | . | . | . |
| PM1548-SK1Org2 | FOXG1 | p.Glu173Asp | missense_variant | MODERATE | 0.119048 | 0 | 42 | 31 | . | . | . | . |
| PM1548-SK1Org2 | CLEC14A | p.Ser404Asn | missense_variant | MODERATE | 0 | 0 | 190 | 138 | . | . | . | . |

| | | | | | | | | | | | | |
|---|---|---|---|---|---|---|---|---|---|---|---|---|
| PM1548-SK1Org2 | KTN1 | p.Phe1182Val | missense_variant | MODERATE | 0.596774 | 0 | 62 | 114 | true | . | . | . |
| PM1548-SK1Org2 | KCNH5 | p.Arg333His | missense_variant | MODERATE | 0.655172 | 0 | 29 | 68 | . | . | . | . |
| PM1548-SK1Org2 | ZBTB25 | p.His371Asn | missense_variant | MODERATE | 0 | 0 | 137 | 47 | . | . | . | . |
| PM1548-SK1Org2 | HSPA2 | p.Gly138fs | frameshift_variant | HIGH | 0.190647 | 0 | 278 | 148 | . | . | . | . |
| PM1548-SK1Org2 | HSPA2 | p.Phe220Phe | synonymous_variant | LOW | 0.05 | 0 | 100 | 96 | . | . | . | . |
| PM1548-SK1Org2 | FUT8 | p.Arg249Cys | missense_variant | MODERATE | 0.55144 | 0 | 243 | 316 | . | . | . | . |
| PM1548-SK1Org2 | TMEM229B | p.Leu37Phe | missense_variant | MODERATE | 0 | 0 | 190 | 483 | . | . | . | . |
| PM1548-SK1Org2 | ZFP36L1 | p.Gly271fs | frameshift_variant | HIGH | 0.427273 | 0 | 220 | 215 | . | . | . | . |
| PM1548-SK1Org2 | ELMSAN1 | p.Arg739Cys | missense_variant | MODERATE | 0.5 | 0 | 142 | 46 | . | . | . | . |
| PM1548-SK1Org2 | ELMSAN1 | p.Gln36fs | frameshift_variant | HIGH | 0.310345 | 0 | 145 | 43 | . | . | . | . |
| PM1548-SK1Org2 | VRTN | p.Gly328fs | frameshift_variant | HIGH | 0.298343 | 0 | 181 | 58 | . | . | . | . |
| PM1548-SK1Org2 | GALC | p.Arg685His | missense_variant | MODERATE | 0 | 0 | 61 | 31 | . | . | . | . |
| PM1548-SK1Org2 | PTPN21 | p.Ile849fs | frameshift_variant | HIGH | 0.555556 | 0 | 72 | 156 | . | . | . | . |
| PM1548-SK1Org2 | ZC3H14 | p.Asn577Asp | missense_variant | MODERATE | 0.00952381 | 0 | 210 | 222 | . | . | . | . |
| PM1548-SK1Org2 | CCDC88C | p.Lys818Glu | missense_variant | MODERATE | 0 | 0 | 130 | 69 | . | . | . | . |
| PM1548-SK1Org2 | FBLN5 | p.Asp357Asn | missense_variant | MODERATE | 0.344262 | 0 | 183 | 147 | . | . | . | . |
| PM1548-SK1Org2 | UNC79 | p.Arg746Gln | missense_variant | MODERATE | 0.579439 | 0 | 107 | 30 | . | . | . | . |
| PM1548-SK1Org2 | SYNE3 | p.Ile83Val | missense_variant | MODERATE | 0.497537 | 0 | 203 | 217 | . | . | . | . |
| PM1548-SK1Org2 | SETD3 | p.Phe388Leu | missense_variant | MODERATE | 0.383721 | 0 | 86 | 32 | . | . | . | . |
| PM1548-SK1Org2 | TECPR2 | p.Gln498fs | frameshift_variant | HIGH | 0.475728 | 0 | 103 | 55 | . | . | . | . |
| PM1548-SK1Org2 | BRF1 | p.Ala129Ser | missense_variant | MODERATE | 0 | 0 | 157 | 135 | . | . | . | . |
| PM1548-SK1Org2 | CYFIP1 | p.Arg1085Leu | missense_variant | MODERATE | 0 | 0 | 193 | 334 | . | . | . | . |
| PM1548-SK1Org2 | MAGEL2 | p.Ala11Val | missense_variant | MODERATE | 0.287554 | 0 | 233 | 317 | . | . | . | . |
| PM1548-SK1Org2 | PAR5 | n.2021_2022del | intron_variant | MODIFIER | 0.108434 | 0 | 83 | 165 | . | . | . | . |
| PM1548-SK1Org2 | UBE3A | p.Val133Ala | missense_variant | MODERATE | 0.388889 | 0 | 54 | 79 | . | . | . | . |
| PM1548-SK1Org2 | ACTC1 | p.Ala137Thr | missense_variant | MODERATE | 0.522936 | 0 | 109 | 175 | . | . | . | . |
| PM1548-SK1Org2 | SPRED1 | p.Arg332His | missense_variant | MODERATE | 0.475728 | 0.0070922 | 103 | 141 | . | . | . | . |
| PM1548-SK1Org2 | CASC5 | p.Asp178Gly | missense_variant | MODERATE | 0.180851 | 0 | 94 | 58 | true | . | . | . |
| PM1548-SK1Org2 | SPTBN5 | p.Arg1943His | missense_variant | MODERATE | 0.216418 | 0 | 134 | 148 | . | . | . | . |
| PM1548-SK1Org2 | SPTBN5 | p.Val1842Gly | missense_variant | MODERATE | 0.0615385 | 0 | 65 | 87 | . | . | . | . |
| PM1548-SK1Org2 | GANC | p.Ala394Val | missense_variant | MODERATE | 0 | 0 | 206 | 106 | . | . | . | . |
| PM1548-SK1Org2 | GANC | p.Gly404Asp | missense_variant | MODERATE | 0 | 0 | 207 | 105 | . | . | . | . |
| PM1548-SK1Org2 | TTBK2 | p.Ala197Thr | missense_variant | MODERATE | 0.587156 | 0 | 109 | 69 | . | . | . | . |
| PM1548-SK1Org2 | CCNDBP1 | p.Asp237Asn | missense_variant | MODERATE | 0.289256 | 0 | 121 | 44 | . | . | . | . |
| PM1548-SK1Org2 | TGM7 | p.Ser305Phe | missense_variant | MODERATE | 0.606557 | 0 | 61 | 54 | . | . | . | . |
| PM1548-SK1Org2 | ZSCAN29 | p.Arg33Trp | missense_variant | MODERATE | 0 | 0 | 149 | 209 | . | . | . | . |
| PM1548-SK1Org2 | SLC28A2 | p.Gly128Asp | missense_variant | MODERATE | 0.198529 | 0 | 136 | 169 | . | . | . | . |
| PM1548-SK1Org2 | MYO5A | p.Arg831His | missense_variant | MODERATE | 0.382114 | 0 | 123 | 78 | true | . | . | . |
| PM1548-SK1Org2 | TCF12 | p.Val183fs | frameshift_variant | HIGH | 0.165354 | 0 | 127 | 200 | true | . | . | . |
| PM1548-SK1Org2 | LIPC | p.Phe286fs | frameshift_variant | HIGH | 0.632653 | 0 | 49 | 89 | . | . | . | . |
| PM1548-SK1Org2 | OAZ2 | p.Ser92Ser | synonymous_variant | LOW | 0.0512821 | 0 | 156 | 163 | . | . | . | . |
| PM1548-SK1Org2 | OAZ2 | p.Glu87Gly | missense_variant | MODERATE | 0 | 0 | 121 | 155 | . | . | . | . |
| PM1548-SK1Org2 | OAZ2 | p.Glu84Val | missense_variant | MODERATE | 0 | 0 | 264 | 248 | . | . | . | . |
| PM1548-SK1Org2 | OAZ2 | p.Val79Gly | missense_variant | MODERATE | 0 | 0 | 260 | 233 | . | . | . | . |
| PM1548-SK1Org2 | SPG21 | p.Ala282Val | missense_variant | MODERATE | 0.46 | 0 | 50 | 32 | . | . | . | . |
| PM1548-SK1Org2 | CILP | p.Asn930Asn | synonymous_variant | LOW | 0.0474453 | 0.00877193 | 274 | 114 | . | . | . | . |
| PM1548-SK1Org2 | IGDCC3 | p.Pro358Leu | missense_variant | MODERATE | 0 | 0 | 249 | 237 | . | . | . | . |
| PM1548-SK1Org2 | NOX5 | p.His470fs | frameshift_variant | HIGH | 0.169643 | 0 | 112 | 164 | . | . | . | . |
| PM1548-SK1Org2 | MYO9A | p.Cys1767Ser | missense_variant | MODERATE | 0.123457 | 0 | 162 | 186 | . | . | . | . |
| PM1548-SK1Org2 | NPTN | p.Arg334Trp | missense_variant | MODERATE | 0.587302 | 0 | 126 | 86 | . | . | . | . |
| PM1548-SK1Org2 | ISLR2 | p.Val74Ala | missense_variant | MODERATE | 0.424 | 0 | 125 | 331 | . | . | . | . |
| PM1548-SK1Org2 | C15orf39 | p.Ala475Val | missense_variant | MODERATE | 0.490566 | 0.00393701 | 265 | 254 | . | . | . | . |
| PM1548-SK1Org2 | ETFA | p.Thr45Arg | missense_variant | MODERATE | 0.0826446 | 0 | 121 | 214 | . | . | . | . |
| PM1548-SK1Org2 | ETFA | p.Thr45Ala | missense_variant | MODERATE | 0.0826446 | 0 | 121 | 214 | . | . | . | . |
| PM1548-SK1Org2 | PSTPIP1 | p.Ala280fs | frameshift_variant&splice_region_variant | HIGH | 0.544248 | 0 | 226 | 438 | . | . | . | . |
| PM1548-SK1Org2 | PEAK1 | p.Ala36Thr | missense_variant | MODERATE | 0.138298 | 0 | 188 | 189 | . | . | . | . |
| PM1548-SK1Org2 | CHRNB4 | p.Gly261Asp | missense_variant | MODERATE | 0.480198 | 0 | 202 | 56 | . | . | . | . |
| PM1548-SK1Org2 | MESDC1 | p.Ser224Asn | missense_variant | MODERATE | 0.43038 | 0 | 79 | 161 | . | . | . | . |
| PM1548-SK1Org2 | IL16 | p.Thr417Met | missense_variant | MODERATE | 0 | 0 | 117 | 34 | . | . | . | . |
| PM1548-SK1Org2 | BTBD1 | p.Arg236Gln | missense_variant | MODERATE | 0.00564972 | 0 | 177 | 77 | . | . | . | . |
| PM1548-SK1Org2 | ZNF592 | p.Ala1232Val | missense_variant | MODERATE | 0.592593 | 0 | 81 | 35 | . | . | . | . |
| PM1548-SK1Org2 | AEN | p.Ala194Val | missense_variant | MODERATE | 0 | 0 | 218 | 122 | . | . | . | . |
| PM1548-SK1Org2 | IQGAP1 | p.Met1231fs | frameshift_variant | HIGH | 0.41958 | 0 | 143 | 170 | . | . | . | . |
| PM1548-SK1Org2 | FES | c.1531-47G>A | intron_variant | MODIFIER | 0.456989 | 0 | 186 | 35 | . | . | . | . |
| PM1548-SK1Org2 | MAN2A2 | p.Pro139Leu | missense_variant | MODERATE | 0.490826 | 0 | 436 | 490 | . | . | . | . |
| PM1548-SK1Org2 | RCCD1 | p.Arg87Ser | missense_variant | MODERATE | 0 | 0 | 91 | 63 | . | . | . | . |
| PM1548-SK1Org2 | ADAMTS17 | p.Cys921Tyr | missense_variant | MODERATE | 0.244681 | 0 | 188 | 58 | . | . | . | . |
| PM1548-SK1Org2 | WDR90 | p.Ala1738Thr | missense_variant | MODERATE | 0.487437 | 0 | 199 | 185 | . | . | . | . |
| PM1548-SK1Org2 | RPUSD1 | p.Arg255Gln | missense_variant | MODERATE | 0.487879 | 0 | 330 | 198 | . | . | . | . |
| PM1548-SK1Org2 | SOX8 | p.Pro11Thr | missense_variant | MODERATE | 0 | 0 | 21 | 62 | . | . | . | . |
| PM1548-SK1Org2 | SOX8 | p.Asp27Asn | missense_variant | MODERATE | 0 | 0 | 28 | 120 | . | . | . | . |
| PM1548-SK1Org2 | CACNA1H | p.Arg1892His | missense_variant | MODERATE | 0.177083 | 0 | 192 | 231 | . | . | . | . |
| PM1548-SK1Org2 | BAIAP3 | p.Ala740Asp | missense_variant | MODERATE | 0.025641 | 0 | 39 | 90 | . | . | . | . |
| PM1548-SK1Org2 | IFT140 | p.Ala1330Val | missense_variant | MODERATE | 0 | 0 | 88 | 188 | . | . | . | . |
| PM1548-SK1Org2 | NME3 | p.Ala20Ser | missense_variant | MODERATE | 0 | 0 | 16 | 162 | . | . | . | . |
| PM1548-SK1Org2 | PRSS33 | p.Leu174fs | frameshift_variant | HIGH | 0 | 0 | 188 | 199 | . | . | . | . |
| PM1548-SK1Org2 | CCDC64B | p.Arg144Gln | missense_variant | MODERATE | 0 | 0.00254453 | 159 | 393 | . | . | . | . |
| PM1548-SK1Org2 | CREBBP | p.Gly1479Val | missense_variant | MODERATE | 0 | 0 | 189 | 193 | true | . | true | . |
| PM1548-SK1Org2 | CORO7 | p.Thr230Ala | missense_variant | MODERATE | 0.221053 | 0 | 95 | 172 | . | . | . | . |
| PM1548-SK1Org2 | GLYR1 | p.Ser148Thr | missense_variant | MODERATE | 0 | 0 | 133 | 42 | . | . | . | . |
| PM1548-SK1Org2 | TVP23A | p.Phe140fs | frameshift_variant | HIGH | 0.548387 | 0 | 155 | 182 | . | . | . | . |
| PM1548-SK1Org2 | CIITA | p.Arg580His | missense_variant | MODERATE | 0.438272 | 0 | 162 | 335 | true | . | . | . |
| PM1548-SK1Org2 | RMI2 | p.Ile130Val | missense_variant | MODERATE | 0.0322581 | 0 | 124 | 171 | true | . | . | . |
| PM1548-SK1Org2 | TNFRSF17 | p.Pro145Gln | missense_variant | MODERATE | 0.275229 | 0 | 109 | 38 | true | . | . | . |
| PM1548-SK1Org2 | CDR2 | p.Gln130Arg | missense_variant | MODERATE | 0.462585 | 0 | 147 | 85 | . | . | . | . |
| PM1548-SK1Org2 | USP31 | p.Gly1267Arg | missense_variant | MODERATE | 0.44898 | 0 | 98 | 32 | . | . | . | . |
| PM1548-SK1Org2 | COG7 | p.Pro67Ala | missense_variant | MODERATE | 0.3875 | 0 | 80 | 71 | . | . | . | . |
| PM1548-SK1Org2 | GTF3C1 | p.Glu1761Gly | missense_variant | MODERATE | 0.0103627 | 0 | 193 | 99 | . | . | . | . |
| PM1548-SK1Org2 | ATXN2L | c.3140-64G>T | intron_variant | MODIFIER | 0.0213904 | 0 | 187 | 124 | . | . | . | . |
| PM1548-SK1Org2 | ATP2A1 | p.Gly229Ser | missense_variant | MODERATE | 0.0330579 | 0 | 242 | 168 | . | . | . | . |
| PM1548-SK1Org2 | KIF22 | p.Ile592fs | frameshift_variant | HIGH | 0.451852 | 0 | 135 | 127 | . | . | . | . |
| PM1548-SK1Org2 | PRR14 | p.Arg96Gln | missense_variant | MODERATE | 0.507519 | 0 | 266 | 271 | . | . | . | . |
| PM1548-SK1Org2 | ZNF629 | p.Asp434Asn | missense_variant | MODERATE | 0.496855 | 0 | 159 | 34 | . | . | . | . |
| PM1548-SK1Org2 | HSD3B7 | p.Val361Ile | missense_variant | MODERATE | 0.474359 | 0 | 156 | 165 | . | . | . | . |
| PM1548-SK1Org2 | KAT8 | p.Val438Ala | missense_variant | MODERATE | 0.48913 | 0 | 92 | 67 | . | . | . | . |
| PM1548-SK1Org2 | FUS | c.1295+1G>A | splice_donor_variant&intron_variant | HIGH | 0.442396 | 0 | 217 | 234 | true | . | . | . |
| PM1548-SK1Org2 | PHKB | p.Tyr52His | missense_variant | MODERATE | 0 | 0 | 115 | 155 | . | . | . | . |
| PM1548-SK1Org2 | LONP2 | p.Gly681Cys | missense_variant | MODERATE | 0.560976 | 0 | 123 | 276 | . | . | . | . |
| PM1548-SK1Org2 | N4BP1 | p.Arg640Cys | missense_variant | MODERATE | 0.277778 | 0 | 72 | 94 | . | . | . | . |
| PM1548-SK1Org2 | SALL1 | p.Ser1318Asn | missense_variant | MODERATE | 0.505495 | 0 | 182 | 197 | . | . | . | . |
| PM1548-SK1Org2 | SALL1 | p.His756Asp | missense_variant | MODERATE | 0.452381 | 0 | 42 | 53 | . | . | . | . |
| PM1548-SK1Org2 | LPCAT2 | c.1216-7923dup | intron_variant | MODIFIER | 0.449153 | 0 | 118 | 56 | . | . | . | . |
| PM1548-SK1Org2 | SLC6A2 | p.Gly591Asp | missense_variant | MODERATE | 0.54185 | 0 | 227 | 342 | . | . | . | . |
| PM1548-SK1Org2 | MMP15 | p.Ala603Asp | missense_variant | MODERATE | 0 | 0 | 257 | 388 | . | . | . | . |
| PM1548-SK1Org2 | CDH11 | p.Ala166Thr | missense_variant | MODERATE | 0.0909091 | 0 | 121 | 177 | true | . | . | . |
| PM1548-SK1Org2 | CDH11 | p.Thr163Ile | missense_variant | MODERATE | 0.0983607 | 0 | 122 | 177 | true | . | . | . |
| PM1548-SK1Org2 | CDH11 | p.His161His | synonymous_variant | LOW | 0.126316 | 0 | 95 | 175 | true | . | . | . |
| PM1548-SK1Org2 | CDH11 | p.Val149Asp | missense_variant | MODERATE | 0.245399 | 0 | 163 | 62 | true | . | . | . |
| PM1548-SK1Org2 | CDH11 | p.Val147Asp | missense_variant | MODERATE | 0.287671 | 0 | 146 | 55 | true | . | . | . |
| PM1548-SK1Org2 | CDH11 | p.Glu140Gly | missense_variant | MODERATE | 0.145455 | 0 | 165 | 80 | true | . | . | . |
| PM1548-SK1Org2 | CDH11 | p.Arg137Gln | missense_variant | MODERATE | 0.149701 | 0 | 167 | 79 | true | . | . | . |
| PM1548-SK1Org2 | TK2 | p.Asn114Asn | synonymous_variant | LOW | 0.426471 | 0 | 136 | 102 | . | . | . | . |
| PM1548-SK1Org2 | B3GNT9 | p.Asp221Asn | missense_variant | MODERATE | 0.117647 | 0 | 85 | 208 | . | . | . | . |
| PM1548-SK1Org2 | LRRC29 | p.Gln205His | missense_variant | MODERATE | 0.108949 | 0 | 257 | 214 | . | . | . | . |
| PM1548-SK1Org2 | KCTD19 | p.Gly313Arg | missense_variant | MODERATE | 0.511765 | 0 | 170 | 109 | . | . | . | . |
| PM1548-SK1Org2 | TPPP3 | p.Ser49Ala | missense_variant | MODERATE | 0.00526316 | 0 | 190 | 59 | . | . | . | . |
| PM1548-SK1Org2 | CTCF | p.Arg283His | missense_variant | MODERATE | 0.602094 | 0 | 191 | 194 | true | true | true | . |
| PM1548-SK1Org2 | CDH3 | p.Asp264Glu | missense_variant | MODERATE | 0 | 0 | 212 | 403 | . | . | . | . |
| PM1548-SK1Org2 | WWP2 | p.Arg633Gln | missense_variant | MODERATE | 0.426087 | 0 | 115 | 88 | . | . | . | . |
| PM1548-SK1Org2 | PMFBP1 | p.Glu205fs | frameshift_variant | HIGH | 0.433099 | 0 | 284 | 212 | . | . | . | . |
| PM1548-SK1Org2 | ZFHX3 | p.Ala1099Ser | missense_variant | MODERATE | 0 | 0 | 101 | 40 | true | . | true | true |
| PM1548-SK1Org2 | RFWD3 | p.Val155Ala | missense_variant | MODERATE | 0.322917 | 0 | 96 | 69 | . | . | . | . |
| PM1548-SK1Org2 | PKD1L2 | n.81164209C>T | intragenic_variant | MODIFIER | 0.507143 | 0 | 140 | 77 | . | . | . | . |
| PM1548-SK1Org2 | TAF1C | p.Pro707fs | frameshift_variant | HIGH | 0.45614 | 0 | 114 | 205 | . | . | . | . |
| PM1548-SK1Org2 | PIEZO1 | p.Pro1061Leu | missense_variant | MODERATE | 0.52459 | 0 | 122 | 171 | . | . | . | . |
| PM1548-SK1Org2 | PIEZO1 | p.Met1002Ile | missense_variant | MODERATE | 0.568182 | 0 | 88 | 33 | . | . | . | . |
| PM1548-SK1Org2 | GALNS | p.Thr109Thr | synonymous_variant | LOW | 0.269841 | 0 | 126 | 187 | . | . | . | . |

| | | | | | | | | | | | | |
|---|---|---|---|---|---|---|---|---|---|---|---|---|
| PM1548-SK1Org2 | ANKRD11 | p.Asp2130Asn | missense_variant | MODERATE | 0 | 0 | 45 | 42 | . | . | . | . |
| PM1548-SK1Org2 | GEMIN4 | p.Arg229Gln | missense_variant | MODERATE | 0.489362 | 0 | 94 | 146 | . | . | . | . |
| PM1548-SK1Org2 | RNMTL1 | p.Met143fs | frameshift_variant | HIGH | 0.207317 | 0 | 82 | 36 | . | . | . | . |
| PM1548-SK1Org2 | PRPF8 | p.Arg1057Gln | missense_variant | MODERATE | 0.652406 | 0 | 187 | 146 | . | . | . | . |
| PM1548-SK1Org2 | HIC1 | p.Pro693Leu | missense_variant | MODERATE | 0.0526316 | 0.0126582 | 38 | 79 | . | . | . | . |
| PM1548-SK1Org2 | SMG6 | p.Arg603His | missense_variant | MODERATE | 0.196347 | 0.00434783 | 219 | 230 | . | . | . | . |
| PM1548-SK1Org2 | CTNS | p.Asp74Asn | missense_variant | MODERATE | 0.477612 | 0.00314465 | 335 | 318 | . | . | . | . |
| PM1548-SK1Org2 | ITGAE | p.Arg113Gln | missense_variant | MODERATE | 0.52381 | 0 | 210 | 94 | . | . | . | . |
| PM1548-SK1Org2 | ZZEF1 | p.Arg194Trp | missense_variant | MODERATE | 0.466019 | 0 | 103 | 95 | . | . | . | . |
| PM1548-SK1Org2 | KIF1C | p.Ser511Phe | missense_variant | MODERATE | 0.394366 | 0 | 71 | 140 | . | . | . | . |
| PM1548-SK1Org2 | USP6 | p.Pro181Leu | missense_variant&splice_region_variant | MODERATE | 0.633333 | 0 | 60 | 73 | true | . | . | . |
| PM1548-SK1Org2 | DVL2 | p.Arg442Cys | missense_variant | MODERATE | 0.520958 | 0 | 167 | 34 | . | . | . | . |
| PM1548-SK1Org2 | EIF5A | p.Thr168Met | missense_variant | MODERATE | 0.509434 | 0 | 212 | 86 | . | . | true | . |
| PM1548-SK1Org2 | DNAH2 | p.Gly4241fs | frameshift_variant | HIGH | 0.49505 | 0 | 202 | 74 | . | . | . | . |
| PM1548-SK1Org2 | ALOX12B | p.Ala355Thr | missense_variant | MODERATE | 0.491429 | 0 | 175 | 295 | . | . | . | . |
| PM1548-SK1Org2 | GLP2R | p.Ala515Thr | missense_variant | MODERATE | 0.545455 | 0.00334448 | 198 | 299 | . | . | . | . |
| PM1548-SK1Org2 | MYH13 | p.Glu984Lys | missense_variant | MODERATE | 0.442105 | 0 | 95 | 72 | . | . | . | . |
| PM1548-SK1Org2 | MYH8 | p.Gly186Arg | missense_variant | MODERATE | 0.494444 | 0 | 180 | 81 | . | . | . | . |
| PM1548-SK1Org2 | MYH4 | p.Tyr424Cys | missense_variant | MODERATE | 0.331288 | 0 | 163 | 298 | . | . | . | . |
| PM1548-SK1Org2 | DNAH9 | p.Arg1542Met | missense_variant | MODERATE | 0 | 0 | 244 | 342 | . | . | . | . |
| PM1548-SK1Org2 | DNAH9 | p.Ala2514Thr | missense_variant | MODERATE | 0.492308 | 0 | 195 | 99 | . | . | . | . |
| PM1548-SK1Org2 | ARHGAP44 | p.Ala495Thr | missense_variant | MODERATE | 0.448276 | 0 | 116 | 64 | . | . | . | . |
| PM1548-SK1Org2 | FLCN | p.His429fs | frameshift_variant | HIGH | 0.0125786 | 0 | 159 | 269 | true | . | . | . |
| PM1548-SK1Org2 | FLCN | p.Thr174Ala | missense_variant | MODERATE | 0.00970874 | 0 | 103 | 227 | true | . | . | . |
| PM1548-SK1Org2 | TOM1L2 | p.Met482fs | frameshift_variant | HIGH | 0.169014 | 0 | 142 | 63 | . | . | . | . |
| PM1548-SK1Org2 | KIAA0100 | p.Arg689* | stop_gained | HIGH | 0.221122 | 0 | 303 | 219 | . | . | . | . |
| PM1548-SK1Org2 | SUPT6H | p.Ile1356Leu | missense_variant | MODERATE | 0.262136 | 0 | 206 | 211 | . | . | . | . |
| PM1548-SK1Org2 | PHF12 | p.Leu393fs | frameshift_variant | HIGH | 0.493088 | 0 | 217 | 60 | . | . | . | . |
| PM1548-SK1Org2 | SEZ6 | p.Gly660Asp | missense_variant | MODERATE | 0.503876 | 0 | 258 | 83 | . | . | . | . |
| PM1548-SK1Org2 | ANKRD13B | p.Arg233Gln | missense_variant | MODERATE | 0 | 0 | 252 | 98 | . | . | . | . |
| PM1548-SK1Org2 | FNDC8 | p.Asn57Ser | missense_variant | MODERATE | 0.625 | 0 | 32 | 149 | . | . | . | . |
| PM1548-SK1Org2 | NLE1 | p.Val10Ala | missense_variant | MODERATE | 0.276074 | 0 | 163 | 97 | . | . | . | . |
| PM1548-SK1Org2 | SRCIN1 | p.Leu460Phe | missense_variant | MODERATE | 0.366667 | 0 | 30 | 52 | . | . | . | . |
| PM1548-SK1Org2 | PCGF2 | p.Asp157Asn | missense_variant | MODERATE | 0.510121 | 0 | 247 | 53 | . | . | . | . |
| PM1548-SK1Org2 | STAC2 | p.Cys141Cys | synonymous_variant | LOW | 0.308036 | 0 | 224 | 250 | . | . | . | . |
| PM1548-SK1Org2 | GRB7 | p.Arg552Gln | missense_variant | MODERATE | 0 | 0 | 113 | 159 | . | . | . | . |
| PM1548-SK1Org2 | NR1D1 | p.Pro593Pro | synonymous_variant | LOW | 0.257426 | 0 | 101 | 103 | . | . | . | . |
| PM1548-SK1Org2 | MSL1 | p.Leu372Ile | missense_variant | MODERATE | 0.101449 | 0 | 69 | 43 | . | . | . | . |
| PM1548-SK1Org2 | WIPF2 | p.Ala39Thr | missense_variant | MODERATE | 0.542857 | 0 | 105 | 66 | . | . | . | . |
| PM1548-SK1Org2 | KRT25 | p.Ala91Thr | missense_variant | MODERATE | 0.6 | 0 | 105 | 148 | . | . | . | . |
| PM1548-SK1Org2 | KRT15 | p.Arg398Trp | missense_variant | MODERATE | 0.546512 | 0 | 172 | 53 | . | . | . | . |
| PM1548-SK1Org2 | KRT16 | p.Gly243Ser | missense_variant | MODERATE | 0 | 0 | 145 | 130 | . | . | . | . |
| PM1548-SK1Org2 | DHX58 | p.Arg224Cys | missense_variant | MODERATE | 0.306011 | 0 | 183 | 87 | . | . | . | . |
| PM1548-SK1Org2 | PLEKHH3 | p.Pro318Thr | missense_variant | MODERATE | 0.257895 | 0 | 190 | 120 | . | . | . | . |
| PM1548-SK1Org2 | ETV4 | p.Gln13Arg | missense_variant | MODERATE | 0 | 0 | 90 | 215 | true | . | . | . |
| PM1548-SK1Org2 | MPP3 | p.Arg265Gln | missense_variant | MODERATE | 0.569444 | 0 | 72 | 78 | . | . | . | . |
| PM1548-SK1Org2 | ATXN7L3 | p.Ile246Phe | missense_variant | MODERATE | 0.527273 | 0 | 110 | 37 | . | . | . | . |
| PM1548-SK1Org2 | ITGA2B | p.Asn330Ser | missense_variant | MODERATE | 0.342466 | 0 | 73 | 60 | . | . | . | . |
| PM1548-SK1Org2 | ARHGAP27 | p.Leu247fs | frameshift_variant | HIGH | 0.310345 | 0 | 145 | 82 | . | . | . | . |
| PM1548-SK1Org2 | WNT3 | p.Ala223Ser | missense_variant | MODERATE | 0 | 0 | 189 | 125 | . | . | . | . |
| PM1548-SK1Org2 | KPNB1 | p.Ile686Leu | missense_variant | MODERATE | 0.00909091 | 0 | 220 | 143 | . | . | . | . |
| PM1548-SK1Org2 | KPNB1 | p.Ile686Met | missense_variant | MODERATE | 0.00909091 | 0 | 220 | 143 | . | . | . | . |
| PM1548-SK1Org2 | SP6 | p.Ala138Thr | missense_variant | MODERATE | 0 | 0 | 132 | 221 | . | . | . | . |
| PM1548-SK1Org2 | PPP1R9B | p.Glu24Glu | synonymous_variant | LOW | 0 | 0 | 12 | 79 | . | . | . | . |
| PM1548-SK1Org2 | CUEDC1 | p.Arg71Cys | missense_variant | MODERATE | 0.561873 | 0 | 299 | 107 | . | . | . | . |
| PM1548-SK1Org2 | LPO | p.Arg82Cys | missense_variant | MODERATE | 0.352941 | 0 | 119 | 404 | . | . | . | . |
| PM1548-SK1Org2 | RNF43 | p.Gly659fs | frameshift_variant | HIGH | 0.525 | 0 | 40 | 76 | true | . | . | true |
| PM1548-SK1Org2 | TBX2 | p.Leu460fs | frameshift_variant | HIGH | 0.459259 | 0 | 135 | 331 | . | . | . | . |
| PM1548-SK1Org2 | MRC2 | p.Arg490Cys | missense_variant | MODERATE | 0 | 0 | 107 | 51 | . | . | . | . |
| PM1548-SK1Org2 | CCDC47 | p.Ser303Pro | missense_variant | MODERATE | 0 | 0 | 115 | 131 | . | . | . | . |
| PM1548-SK1Org2 | FTSJ3 | p.Met493Ile | missense_variant | MODERATE | 0.410714 | 0 | 112 | 61 | . | . | . | . |
| PM1548-SK1Org2 | ABCA10 | p.Tyr1399His | missense_variant | MODERATE | 0.398551 | 0 | 138 | 335 | . | . | . | . |
| PM1548-SK1Org2 | SLC16A5 | p.Gly93Cys | missense_variant | MODERATE | 0 | 0 | 95 | 58 | . | . | . | . |
| PM1548-SK1Org2 | ITGB4 | p.Pro305Leu | missense_variant | MODERATE | 0.507042 | 0 | 142 | 78 | . | . | . | . |
| PM1548-SK1Org2 | ITGB4 | p.Pro723Ser | missense_variant | MODERATE | 0.150754 | 0 | 199 | 167 | . | . | . | . |
| PM1548-SK1Org2 | H3F3B | p.Ala8Asp | missense_variant | MODERATE | 0.117647 | 0 | 51 | 88 | true | . | . | . |
| PM1548-SK1Org2 | EVPL | p.Arg266Trp | missense_variant | MODERATE | 0.307692 | 0 | 13 | 157 | . | . | . | . |
| PM1548-SK1Org2 | RHBDF2 | p.Ala354Thr | missense_variant | MODERATE | 0 | 0 | 146 | 138 | . | . | . | . |
| PM1548-SK1Org2 | TNRC6C | p.Arg1160His | missense_variant | MODERATE | 0.418919 | 0 | 74 | 184 | . | . | . | . |
| PM1548-SK1Org2 | TMC6 | p.His31His | synonymous_variant | LOW | 0.605556 | 0 | 180 | 491 | . | . | . | . |
| PM1548-SK1Org2 | CCDC40 | p.Gly929Ser | missense_variant | MODERATE | 0 | 0 | 8 | 34 | . | . | . | . |
| PM1548-SK1Org2 | GAA | p.Gly828Asp | missense_variant&splice_region_variant | MODERATE | 0.0637255 | 0 | 204 | 76 | . | . | . | . |
| PM1548-SK1Org2 | ENDOV | p.Pro259Leu | missense_variant | MODERATE | 0.410256 | 0 | 78 | 82 | . | . | . | . |
| PM1548-SK1Org2 | RPTOR | p.Arg899* | stop_gained | HIGH | 0.453488 | 0 | 86 | 71 | . | . | . | . |
| PM1548-SK1Org2 | SIRT7 | p.Thr181Met | missense_variant | MODERATE | 0.266409 | 0 | 259 | 217 | . | . | . | . |
| PM1548-SK1Org2 | OGFOD3 | p.Ser127Ser | splice_region_variant&synonymous_variant | LOW | 0.534483 | 0 | 232 | 110 | . | . | . | . |
| PM1548-SK1Org2 | TXNDC2 | p.Ile284Leu | missense_variant | MODERATE | 0 | 0 | 156 | 275 | . | . | . | . |
| PM1548-SK1Org2 | TXNDC2 | p.Ser285Pro | missense_variant | MODERATE | 0 | 0 | 156 | 275 | . | . | . | . |
| PM1548-SK1Org2 | TXNDC2 | p.Pro288Leu | missense_variant | MODERATE | 0 | 0.00816327 | 145 | 245 | . | . | . | . |
| PM1548-SK1Org2 | ANKRD62 | p.Gln745His | missense_variant | MODERATE | 1 | 0 | 74 | 165 | . | . | . | . |
| PM1548-SK1Org2 | ZNF521 | p.Cys754Tyr | missense_variant | MODERATE | 0.594937 | 0 | 79 | 57 | true | . | . | . |
| PM1548-SK1Org2 | KCTD1 | p.Asn124His | missense_variant | MODERATE | 0.222222 | 0 | 9 | 69 | . | . | . | . |
| PM1548-SK1Org2 | DSG4 | p.Thr1001Met | missense_variant | MODERATE | 1 | 0 | 94 | 104 | . | . | . | . |
| PM1548-SK1Org2 | CCDC178 | p.Arg827Met | missense_variant | MODERATE | 0 | 0 | 148 | 192 | . | . | . | . |
| PM1548-SK1Org2 | INO80C | p.Gly216Ser | missense_variant | MODERATE | 0 | 0 | 53 | 89 | . | . | . | . |
| PM1548-SK1Org2 | SETBP1 | p.Glu244Asp | missense_variant | MODERATE | 0.0384615 | 0 | 78 | 133 | true | true | . | . |
| PM1548-SK1Org2 | SETBP1 | p.Ile251Val | missense_variant | MODERATE | 0.0196078 | 0 | 102 | 201 | true | true | . | . |
| PM1548-SK1Org2 | SETBP1 | p.Ala733Thr | missense_variant | MODERATE | 0.0106383 | 0 | 94 | 315 | true | true | . | . |
| PM1548-SK1Org2 | SIGLEC15 | p.Ala96Thr | missense_variant | MODERATE | 0 | 0 | 9 | 175 | . | . | . | . |
| PM1548-SK1Org2 | SIGLEC15 | p.Arg99Pro | missense_variant | MODERATE | 0 | 0 | 9 | 176 | . | . | . | . |
| PM1548-SK1Org2 | SKOR2 | p.Asp736Glu | missense_variant | MODERATE | 0.0714286 | 0 | 56 | 157 | . | . | . | . |
| PM1548-SK1Org2 | SKOR2 | p.Gly720Glu | missense_variant | MODERATE | 0.0416667 | 0 | 48 | 134 | . | . | . | . |
| PM1548-SK1Org2 | SMAD7 | p.Pro55Ala | missense_variant | MODERATE | 0.25 | 0 | 36 | 32 | . | . | . | . |
| PM1548-SK1Org2 | DSEL | p.Arg1009His | missense_variant | MODERATE | 1 | 0 | 87 | 67 | . | . | . | . |
| PM1548-SK1Org2 | FAM69C | p.Gly48Cys | missense_variant | MODERATE | 0 | 0 | 25 | 170 | . | . | . | . |
| PM1548-SK1Org2 | TSHZ1 | p.Ala570Val | missense_variant | MODERATE | 0 | 0 | 137 | 279 | . | . | . | . |
| PM1548-SK1Org2 | TSHZ1 | p.Tyr775Cys | missense_variant | MODERATE | 1 | 0 | 98 | 122 | . | . | . | . |
| PM1548-SK1Org2 | ZNF516 | p.Gly232del | disruptive_inframe_deletion | MODERATE | 0 | 0 | 95 | 279 | . | . | . | . |
| PM1548-SK1Org2 | ZNF516 | p.Pro229_Gly23( | inframe_insertion | MODERATE | 0 | 0 | 97 | 280 | . | . | . | . |
| PM1548-SK1Org2 | GALR1 | p.Leu46Met | missense_variant | MODERATE | 0.148936 | 0 | 47 | 102 | . | . | . | . |
| PM1548-SK1Org2 | GALR1 | p.Lys62Lys | synonymous_variant | LOW | 0.136842 | 0 | 95 | 218 | . | . | . | . |
| PM1548-SK1Org2 | SALL3 | p.Pro1105fs | frameshift_variant | HIGH | 1 | 0 | 18 | 200 | . | . | . | . |
| PM1548-SK1Org2 | PPAP2C | n.291320T>C | intragenic_variant | MODIFIER | 0.463415 | 0 | 82 | 59 | . | . | . | . |
| PM1548-SK1Org2 | LPPR3 | p.Gly646Ser | missense_variant | MODERATE | 1 | 0 | 4 | 120 | . | . | . | . |
| PM1548-SK1Org2 | CFD | p.Arg199Cys | missense_variant | MODERATE | 0.438776 | 0 | 196 | 193 | . | . | . | . |
| PM1548-SK1Org2 | ARID3A | p.Arg503Cys | missense_variant | MODERATE | 0.35 | 0 | 40 | 124 | . | . | . | . |
| PM1548-SK1Org2 | APC2 | p.Asp501Asn | missense_variant | MODERATE | 0.537688 | 0 | 199 | 282 | . | . | . | . |
| PM1548-SK1Org2 | PCSK4 | p.Gly57Asp | missense_variant | MODERATE | 0.231818 | 0 | 220 | 340 | . | . | . | . |
| PM1548-SK1Org2 | ADAMTSL5 | p.Ala343fs | frameshift_variant | HIGH | 0.395349 | 0 | 258 | 366 | . | . | . | . |
| PM1548-SK1Org2 | ADAMTSL5 | p.Pro341Ala | missense_variant | MODERATE | 0.0438596 | 0 | 228 | 313 | . | . | . | . |
| PM1548-SK1Org2 | MEX3D | p.Gly262Asp | missense_variant | MODERATE | 0.578313 | 0 | 83 | 217 | . | . | . | . |
| PM1548-SK1Org2 | TCF3 | p.Gly247fs | frameshift_variant | HIGH | 0 | 0 | 157 | 71 | true | . | . | . |
| PM1548-SK1Org2 | C19orf35 | p.Arg325Cys | missense_variant | MODERATE | 0 | 0 | 65 | 53 | . | . | . | . |
| PM1548-SK1Org2 | NCLN | p.Thr70Met | missense_variant | MODERATE | 0.578947 | 0 | 95 | 140 | . | . | . | . |
| PM1548-SK1Org2 | DOHH | p.Arg286Trp | missense_variant | MODERATE | 0.294118 | 0.00684932 | 204 | 146 | . | . | . | . |
| PM1548-SK1Org2 | EEF2 | p.Pro738Ser | missense_variant | MODERATE | 0.488235 | 0 | 170 | 230 | . | . | . | . |
| PM1548-SK1Org2 | PIAS4 | p.Tyr265Tyr | synonymous_variant | LOW | 0.409091 | 0 | 110 | 240 | . | . | . | . |
| PM1548-SK1Org2 | MAP2K2 | p.Lys51Arg | missense_variant | MODERATE | 0 | 0 | 198 | 75 | true | . | . | . |
| PM1548-SK1Org2 | MAP2K2 | p.Lys51Gln | missense_variant | MODERATE | 0 | 0 | 198 | 75 | true | . | . | . |
| PM1548-SK1Org2 | CREB3L3 | p.Val140Ala | missense_variant | MODERATE | 0 | 0 | 94 | 205 | . | . | . | . |
| PM1548-SK1Org2 | UBXN6 | p.Thr337Met | missense_variant | MODERATE | 0 | 0 | 148 | 93 | . | . | . | . |
| PM1548-SK1Org2 | PLIN4 | p.Val592Met | missense_variant | MODERATE | 0 | 0 | 140 | 40 | . | . | . | . |
| PM1548-SK1Org2 | PLIN5 | p.Thr456Ile | missense_variant | MODERATE | 0.573333 | 0 | 75 | 37 | . | . | . | . |
| PM1548-SK1Org2 | AC005594.3 | n.381G>A | non_coding_exon_variant | MODIFIER | 0.488889 | 0.0037037 | 180 | 270 | . | . | . | . |
| PM1548-SK1Org2 | DPP9 | p.Arg125Trp | missense_variant | MODERATE | 0.0508475 | 0 | 59 | 40 | . | . | . | . |
| PM1548-SK1Org2 | KDM4B | p.Arg618Cys | missense_variant | MODERATE | 0 | 0 | 50 | 167 | . | . | . | . |
| PM1548-SK1Org2 | KDM4B | p.Glu1013Lys | missense_variant | MODERATE | 0.254545 | 0 | 110 | 113 | . | . | . | . |

| | | | | | | | | | | | | |
|---|---|---|---|---|---|---|---|---|---|---|---|---|
| PM1548-SK1Org2 | DENND1C | p.Thr165fs | frameshift_variant | HIGH | 0.443709 | 0 | 151 | 34 | . | . | . | . |
| PM1548-SK1Org2 | PEX11G | p.Arg124Trp | missense_variant | MODERATE | 0.456 | 0 | 125 | 176 | . | . | . | . |
| PM1548-SK1Org2 | CTD-3193O13.9 | p.Thr838Thr | synonymous_variant | LOW | 0.436464 | 0 | 181 | 174 | . | . | . | . |
| PM1548-SK1Org2 | COL5A3 | p.Glu1677Lys | missense_variant | MODERATE | 0.46 | 0 | 100 | 264 | . | . | true | true |
| PM1548-SK1Org2 | ZNF653 | p.Asp247Gly | missense_variant | MODERATE | 0.210526 | 0 | 133 | 166 | . | . | . | . |
| PM1548-SK1Org2 | ELOF1 | p.Arg41Cys | missense_variant | MODERATE | 0.546218 | 0 | 238 | 108 | . | . | . | . |
| PM1548-SK1Org2 | MRI1 | p.His330fs | frameshift_variant | HIGH | 0.487395 | 0 | 119 | 82 | . | . | . | . |
| PM1548-SK1Org2 | RFX1 | p.Val600Ile | missense_variant | MODERATE | 0.126214 | 0 | 206 | 347 | . | . | . | . |
| PM1548-SK1Org2 | PRKACA | p.Ala219Val | missense_variant | MODERATE | 0.456897 | 0 | 116 | 200 | true | . | . | . |
| PM1548-SK1Org2 | TECR | p.Val85Met | missense_variant | MODERATE | 0.506173 | 0.00641026 | 81 | 156 | . | . | . | . |
| PM1548-SK1Org2 | EMR2 | p.Pro294Thr | missense_variant | MODERATE | 0.102128 | 0 | 235 | 117 | . | . | . | . |
| PM1548-SK1Org2 | EMR2 | p.Tyr292Ser | missense_variant | MODERATE | 0.106383 | 0 | 235 | 117 | . | . | . | . |
| PM1548-SK1Org2 | KLF2 | p.Ala238fs | frameshift_variant | HIGH | 0.4 | 0 | 30 | 137 | . | . | . | . |
| PM1548-SK1Org2 | NWD1 | p.Arg788Cys | missense_variant | MODERATE | 0.474359 | 0 | 156 | 334 | . | . | . | . |
| PM1548-SK1Org2 | BST2 | p.Ile120fs | frameshift_variant | HIGH | 0.462121 | 0 | 132 | 64 | . | . | . | . |
| PM1548-SK1Org2 | FCHO1 | p.Ala286Val | missense_variant&splice_region_variant | MODERATE | 0.471223 | 0 | 278 | 272 | . | . | . | . |
| PM1548-SK1Org2 | DDX49 | p.Arg310Gln | missense_variant&splice_region_variant | MODERATE | 0.441026 | 0 | 195 | 300 | . | . | . | . |
| PM1548-SK1Org2 | SUGP2 | p.Cys490Tyr | missense_variant | MODERATE | 0.6 | 0 | 130 | 81 | . | . | . | . |
| PM1548-SK1Org2 | NR2C2AP | p.Lys28fs | frameshift_variant | HIGH | 0.416667 | 0 | 132 | 183 | . | . | . | . |
| PM1548-SK1Org2 | NCAN | p.Arg256Gln | missense_variant | MODERATE | 0.423423 | 0 | 111 | 124 | . | . | . | . |
| PM1548-SK1Org2 | NDUFA13 | p.Trp131Leu | missense_variant | MODERATE | 0 | 0 | 63 | 265 | . | . | . | . |
| PM1548-SK1Org2 | ZNF728 | p.Pro380Ser | missense_variant | MODERATE | 0.387755 | 0 | 49 | 32 | . | . | . | . |
| PM1548-SK1Org2 | ZNF728 | p.Arg323Pro | missense_variant | MODERATE | 0.154472 | 0 | 123 | 48 | . | . | . | . |
| PM1548-SK1Org2 | ZNF728 | p.Arg323Trp | missense_variant | MODERATE | 0.154472 | 0 | 123 | 48 | . | . | . | . |
| PM1548-SK1Org2 | ZNF507 | p.Asp475Gly | missense_variant | MODERATE | 0.528926 | 0 | 121 | 42 | . | . | . | . |
| PM1548-SK1Org2 | KIAA0355 | p.Val801Met | missense_variant | MODERATE | 0.0536913 | 0.0060241 | 149 | 166 | . | . | . | . |
| PM1548-SK1Org2 | GRAMD1A | p.Gly605Ser | missense_variant | MODERATE | 0.107143 | 0 | 112 | 68 | . | . | . | . |
| PM1548-SK1Org2 | RBM42 | p.Pro117Arg | missense_variant | MODERATE | 0.327327 | 0 | 333 | 129 | . | . | . | . |
| PM1548-SK1Org2 | ARHGAP33 | p.Gln144fs | frameshift_variant | HIGH | 0.51938 | 0 | 387 | 430 | . | . | . | . |
| PM1548-SK1Org2 | ZNF850 | p.Ala575Thr | missense_variant | MODERATE | 0 | 0 | 90 | 89 | . | . | . | . |
| PM1548-SK1Org2 | ZNF850 | p.Ser572Ala | missense_variant | MODERATE | 0 | 0 | 96 | 133 | . | . | . | . |
| PM1548-SK1Org2 | ZNF850 | p.Val554Ile | missense_variant | MODERATE | 0 | 0 | 52 | 86 | . | . | . | . |
| PM1548-SK1Org2 | DPF1 | p.Cys58Cys | synonymous_variant | LOW | 0 | 0 | 85 | 394 | . | . | . | . |
| PM1548-SK1Org2 | DPF1 | p.Cys50Cys | synonymous_variant | LOW | 0 | 0 | 84 | 391 | . | . | . | . |
| PM1548-SK1Org2 | YIF1B | p.Phe39Phe | synonymous_variant | LOW | 0.498721 | 0 | 391 | 475 | . | . | . | . |
| PM1548-SK1Org2 | RYR1 | p.Val3218Met | missense_variant | MODERATE | 0.00666667 | 0 | 150 | 114 | . | . | . | . |
| PM1548-SK1Org2 | FCGBP | p.Ala5241Val | missense_variant | MODERATE | 0.454054 | 0 | 185 | 284 | . | . | . | . |
| PM1548-SK1Org2 | PRX | p.Glu495Gln | missense_variant | MODERATE | 0.318182 | 0 | 44 | 97 | . | true | . | . |
| PM1548-SK1Org2 | B3GNT8 | p.Glu333Ala | missense_variant | MODERATE | 0.366492 | 0 | 191 | 219 | . | . | . | . |
| PM1548-SK1Org2 | ARHGEF1 | p.Arg137Trp | missense_variant | MODERATE | 0.590551 | 0 | 127 | 130 | . | . | . | . |
| PM1548-SK1Org2 | ZNF227 | p.Gly390Ser | missense_variant | MODERATE | 0.634615 | 0 | 104 | 73 | . | . | . | . |
| PM1548-SK1Org2 | PVRL2 | p.Pro53Leu | missense_variant | MODERATE | 0.509434 | 0 | 106 | 119 | . | . | . | . |
| PM1548-SK1Org2 | MARK4 | p.Thr651Thr | synonymous_variant | LOW | 0.183267 | 0 | 251 | 352 | . | . | . | . |
| PM1548-SK1Org2 | KLC3 | p.Arg320Cys | missense_variant | MODERATE | 0.518519 | 0 | 108 | 102 | . | . | . | . |
| PM1548-SK1Org2 | PPP1R13L | p.Leu397Pro | missense_variant | MODERATE | 0.00655738 | 0 | 305 | 478 | . | . | . | . |
| PM1548-SK1Org2 | GIPR | p.Gly144Ser | missense_variant | MODERATE | 0.382353 | 0 | 68 | 35 | . | . | . | . |
| PM1548-SK1Org2 | STRN4 | p.Ala484Val | missense_variant | MODERATE | 0.54321 | 0 | 162 | 141 | . | . | . | . |
| PM1548-SK1Org2 | FKRP | p.Ala193Thr | missense_variant | MODERATE | 0 | 0 | 25 | 98 | . | . | . | . |
| PM1548-SK1Org2 | SLC1A5 | p.Ala534Thr | missense_variant | MODERATE | 0.295775 | 0 | 71 | 44 | . | . | . | . |
| PM1548-SK1Org2 | ZC3H4 | p.Arg894His | missense_variant | MODERATE | 0 | 0 | 116 | 254 | . | . | . | . |
| PM1548-SK1Org2 | ZC3H4 | p.Gly668Ser | missense_variant | MODERATE | 0.470588 | 0 | 204 | 220 | . | . | . | . |
| PM1548-SK1Org2 | ZC3H4 | p.Lys500Arg | missense_variant | MODERATE | 0 | 0 | 156 | 129 | . | . | . | . |
| PM1548-SK1Org2 | ZNF541 | p.Arg214His | missense_variant | MODERATE | 0 | 0 | 72 | 53 | . | . | . | . |
| PM1548-SK1Org2 | GLTSCR1 | p.Ser1126fs | frameshift_variant | HIGH | 0.456 | 0 | 125 | 148 | . | . | . | . |
| PM1548-SK1Org2 | CRX | p.Ala35Thr | missense_variant&splice_region_variant | MODERATE | 0.5 | 0 | 184 | 73 | . | . | . | . |
| PM1548-SK1Org2 | PLEKHA4 | p.Arg225Trp | missense_variant | MODERATE | 0.483146 | 0 | 89 | 36 | . | . | . | . |
| PM1548-SK1Org2 | BAX | p.Glu41fs | frameshift_variant | HIGH | 0.00416667 | 0 | 240 | 76 | . | . | . | . |
| PM1548-SK1Org2 | FLT3LG | p.Ser118fs | frameshift_variant | HIGH | 0.243478 | 0 | 115 | 82 | . | . | . | . |
| PM1548-SK1Org2 | SCAF1 | p.Gly651fs | frameshift_variant | HIGH | 0.530612 | 0 | 49 | 93 | . | . | . | . |
| PM1548-SK1Org2 | ASPDH | p.Glu177Lys | missense_variant | MODERATE | 0.458182 | 0 | 275 | 225 | . | . | . | . |
| PM1548-SK1Org2 | LRRC4B | p.Gly50fs | frameshift_variant | HIGH | 0.571429 | 0 | 42 | 58 | . | . | . | . |
| PM1548-SK1Org2 | ZNF611 | p.Met226Thr | missense_variant | MODERATE | 0 | 0 | 53 | 80 | . | . | . | . |
| PM1548-SK1Org2 | CNOT3 | p.Tyr637His | missense_variant | MODERATE | 0.542373 | 0 | 118 | 41 | true | . | . | true |
| PM1548-SK1Org2 | LILRB3 | c.355+1720T>A | intron_variant | MODIFIER | 0.00995025 | 0 | 201 | 187 | . | . | . | . |
| PM1548-SK1Org2 | LILRB3 | c.355+1720T>A | intron_variant | MODIFIER | 0.00995025 | 0 | 201 | 187 | . | . | . | . |
| PM1548-SK1Org2 | LILRB3 | c.355+1720T>A | intron_variant | MODIFIER | 0.00990099 | 0 | 202 | 184 | . | . | . | . |
| PM1548-SK1Org2 | LILRB3 | c.355+1720T>A | intron_variant | MODIFIER | 0.00990099 | 0 | 202 | 184 | . | . | . | . |
| PM1548-SK1Org2 | EPS8L1 | p.Arg608Cys | missense_variant | MODERATE | 0.515625 | 0 | 64 | 167 | . | . | . | . |
| PM1548-SK1Org2 | SBK2 | p.Ala115Thr | missense_variant | MODERATE | 0.488449 | 0 | 303 | 186 | . | . | . | . |
| PM1548-SK1Org2 | ZNF579 | p.Arg101Cys | missense_variant | MODERATE | 0.493333 | 0 | 75 | 216 | . | . | . | . |
| PM1548-SK1Org2 | ZNF865 | p.Leu610Phe | missense_variant | MODERATE | 0.0141844 | 0 | 141 | 212 | . | . | . | . |
| PM1548-SK1Org2 | ZNF865 | p.Leu632Leu | synonymous_variant | LOW | 0.0192308 | 0 | 104 | 209 | . | . | . | . |
| PM1548-SK1Org2 | ZNF71 | p.Gln343Gln | synonymous_variant | LOW | 0.502041 | 0 | 245 | 303 | . | . | . | . |
| PM1548-SK1Org2 | TRIM28 | p.Arg492Cys | missense_variant | MODERATE | 0.52439 | 0 | 164 | 142 | . | . | . | . |
| PM1548-SK1Org2 | ACP1 | p.His67Asn | missense_variant | MODERATE | 0 | 0 | 36 | 36 | . | . | . | . |
| PM1548-SK1Org2 | PXDN | p.Val850Met | missense_variant | MODERATE | 0 | 0 | 274 | 308 | . | . | . | . |
| PM1548-SK1Org2 | ROCK2 | p.Glu1060Gly | missense_variant | MODERATE | 0 | 0 | 97 | 93 | . | . | . | . |
| PM1548-SK1Org2 | FAM228B | p.Asp82Gly | missense_variant | MODERATE | 0.49359 | 0 | 156 | 174 | . | . | . | . |
| PM1548-SK1Org2 | NCOA1 | p.Asn1244Thr | missense_variant | MODERATE | 0.016 | 0 | 250 | 206 | true | . | . | . |
| PM1548-SK1Org2 | NCOA1 | p.Ile1260Val | missense_variant | MODERATE | 0.018797 | 0 | 266 | 204 | true | . | . | . |
| PM1548-SK1Org2 | NCOA1 | p.Ala1274Thr | missense_variant | MODERATE | 0.0110497 | 0 | 181 | 87 | true | . | . | . |
| PM1548-SK1Org2 | ADCY3 | p.Thr182Met | missense_variant | MODERATE | 0.58296 | 0 | 223 | 84 | . | . | . | . |
| PM1548-SK1Org2 | TRIM54 | p.Arg97Gln | missense_variant | MODERATE | 0.432432 | 0 | 222 | 313 | . | . | . | . |
| PM1548-SK1Org2 | TRIM54 | p.Glu131Asp | missense_variant | MODERATE | 0.048913 | 0 | 184 | 142 | . | . | . | . |
| PM1548-SK1Org2 | HEATR5B | p.Lys48fs | frameshift_variant | HIGH | 0 | 0 | 39 | 85 | . | . | . | . |
| PM1548-SK1Org2 | RMDN2 | p.Arg130His | missense_variant | MODERATE | 0.25 | 0 | 132 | 120 | . | . | . | . |
| PM1548-SK1Org2 | SOS1 | p.Asn15Lys | missense_variant | MODERATE | 0.573248 | 0 | 157 | 93 | . | . | . | . |
| PM1548-SK1Org2 | FSHR | p.Arg569Lys | missense_variant | MODERATE | 0.0909091 | 0 | 66 | 177 | . | . | . | . |
| PM1548-SK1Org2 | FSHR | p.Ser566Arg | missense_variant | MODERATE | 0.0983607 | 0 | 61 | 133 | . | . | . | . |
| PM1548-SK1Org2 | FSHR | p.Asn558Asn | synonymous_variant | LOW | 0.0722892 | 0 | 83 | 139 | . | . | . | . |
| PM1548-SK1Org2 | FSHR | p.Ile550Thr | missense_variant | MODERATE | 0.0560748 | 0 | 107 | 160 | . | . | . | . |
| PM1548-SK1Org2 | CCDC88A | p.Pro1470Ser | missense_variant | MODERATE | 0.418994 | 0 | 179 | 211 | . | . | . | . |
| PM1548-SK1Org2 | CCDC85A | p.His295Tyr | missense_variant | MODERATE | 0.253521 | 0 | 71 | 101 | . | . | . | . |
| PM1548-SK1Org2 | CCDC85A | p.Pro359Leu | missense_variant | MODERATE | 0.365079 | 0 | 63 | 45 | . | . | . | . |
| PM1548-SK1Org2 | FANCL | c.555+2T>C | splice_donor_variant&intron_variant | HIGH | 0.416667 | 0 | 96 | 47 | . | . | . | . |
| PM1548-SK1Org2 | OTX1 | p.Arg39Trp | missense_variant | MODERATE | 0.644444 | 0 | 135 | 144 | . | . | . | . |
| PM1548-SK1Org2 | AFTPH | p.Asp198fs | frameshift_variant | HIGH | 0 | 0 | 195 | 159 | . | . | . | . |
| PM1548-SK1Org2 | NAGK | p.Lys90Gln | missense_variant | MODERATE | 0.0147601 | 0 | 271 | 344 | . | . | . | . |
| PM1548-SK1Org2 | NAGK | p.Asn100Asp | missense_variant | MODERATE | 0.015873 | 0 | 252 | 271 | . | . | . | . |
| PM1548-SK1Org2 | FBXO41 | p.Arg594Gln | missense_variant | MODERATE | 0 | 0 | 203 | 69 | . | . | . | . |
| PM1548-SK1Org2 | TET3 | p.Pro1195Leu | missense_variant | MODERATE | 0.546763 | 0 | 139 | 50 | . | . | . | . |
| PM1548-SK1Org2 | C2orf81 | p.Ala344Val | missense_variant | MODERATE | 0.148148 | 0 | 54 | 154 | . | . | . | . |
| PM1548-SK1Org2 | TCF7L1 | p.His246Pro | missense_variant | MODERATE | 0.0491803 | 0.00719424 | 183 | 139 | . | . | . | . |
| PM1548-SK1Org2 | KDM3A | p.Glu1141* | stop_gained | HIGH | 0 | 0 | 203 | 166 | . | . | . | . |
| PM1548-SK1Org2 | ZNF514 | p.Gly396Glu | missense_variant | MODERATE | 0.526316 | 0 | 209 | 187 | . | . | . | . |
| PM1548-SK1Org2 | NCAPH | p.Gln453* | stop_gained&splice_region_variant | HIGH | 0.530435 | 0 | 115 | 53 | . | . | . | . |
| PM1548-SK1Org2 | ZAP70 | p.Ala387Val | missense_variant | MODERATE | 0.261538 | 0 | 195 | 293 | . | . | . | . |
| PM1548-SK1Org2 | LONRF2 | p.Asp406Gly | missense_variant | MODERATE | 0 | 0 | 90 | 174 | . | . | . | . |
| PM1548-SK1Org2 | CHST10 | p.Asn138Asn | synonymous_variant | LOW | 0.465753 | 0 | 146 | 236 | . | . | . | . |
| PM1548-SK1Org2 | IL18R1 | p.Pro527fs | frameshift_variant | HIGH | 0.376623 | 0 | 77 | 255 | . | . | . | . |
| PM1548-SK1Org2 | FBLN7 | c.670+86C>T | intron_variant | MODIFIER | 0.578313 | 0 | 83 | 109 | . | . | . | . |
| PM1548-SK1Org2 | FBLN7 | p.Arg381Cys | missense_variant | MODERATE | 0.388601 | 0 | 193 | 87 | . | . | . | . |
| PM1548-SK1Org2 | ZC3H6 | p.Arg99fs | frameshift_variant | HIGH | 0.440367 | 0 | 109 | 90 | . | . | . | . |
| PM1548-SK1Org2 | POLR1B | p.Cys893Tyr | missense_variant | MODERATE | 0.337209 | 0 | 86 | 125 | . | . | . | . |
| PM1548-SK1Org2 | IL36B | c.261+1014delT | intron_variant | MODIFIER | 0.485437 | 0 | 206 | 101 | . | . | . | . |
| PM1548-SK1Org2 | TMEM185B | p.Pro341fs | frameshift_variant | HIGH | 0.270588 | 0 | 170 | 229 | . | . | . | . |
| PM1548-SK1Org2 | CLASP1 | p.Ala1225Val | missense_variant | MODERATE | 0.186207 | 0 | 145 | 85 | . | . | . | . |
| PM1548-SK1Org2 | CYP27C1 | p.Ala103Thr | missense_variant | MODERATE | 0.59434 | 0 | 106 | 179 | . | . | . | . |
| PM1548-SK1Org2 | ACVR2A | p.Lys437fs | frameshift_variant | HIGH | 1 | 0 | 57 | 199 | . | true | . | . |
| PM1548-SK1Org2 | NEB | p.Phe2843Tyr | missense_variant | MODERATE | 0.1 | 0 | 120 | 178 | . | . | . | . |
| PM1548-SK1Org2 | TBR1 | p.Gly599Ser | missense_variant | MODERATE | 1 | 0 | 4 | 40 | . | . | . | . |
| PM1548-SK1Org2 | SCN3A | p.Ala1768Val | missense_variant | MODERATE | 0.938462 | 0 | 65 | 50 | . | . | . | . |
| PM1548-SK1Org2 | TLK1 | p.Gln18Arg | missense_variant | MODERATE | 0.125 | 0 | 64 | 529 | . | . | . | . |
| PM1548-SK1Org2 | ITGA6 | p.Val266Leu | missense_variant | MODERATE | 0.0470588 | 0 | 85 | 156 | . | . | . | . |
| PM1548-SK1Org2 | SP3 | p.Val461Ile | missense_variant | MODERATE | 0.0375 | 0 | 160 | 262 | . | . | . | . |
| PM1548-SK1Org2 | EVX2 | p.Gly438Ser | missense_variant | MODERATE | 1 | 0 | 58 | 165 | . | . | . | . |
| PM1548-SK1Org2 | EVX2 | p.Asp320Glu | missense_variant | MODERATE | 0 | 0 | 15 | 48 | . | . | . | . |

| | | | | | | | | | | | | |
|---|---|---|---|---|---|---|---|---|---|---|---|---|
| PM1548-SK1Org2 | CCDC141 | p.Ala1030Val | missense_variant | MODERATE | 0.333333 | 0 | 72 | 69 | . | . | . | . |
| PM1548-SK1Org2 | PDE1A | p.Ala371Val | missense_variant | MODERATE | 1 | 0 | 65 | 75 | . | . | . | . |
| PM1548-SK1Org2 | FSIP2 | p.Arg829Cys | missense_variant | MODERATE | 0.363636 | 0 | 110 | 51 | . | . | . | . |
| PM1548-SK1Org2 | DNAH7 | p.Ala3252Glu | missense_variant | MODERATE | 0.0625 | 0 | 48 | 64 | . | . | . | . |
| PM1548-SK1Org2 | GTF3C3 | p.Pro248Leu | missense_variant | MODERATE | 0.0174672 | 0 | 229 | 201 | . | . | . | . |
| PM1548-SK1Org2 | AOX1 | p.Phe1162Ser | missense_variant | MODERATE | 0.179856 | 0 | 139 | 36 | . | . | . | . |
| PM1548-SK1Org2 | PIKFYVE | p.Leu419Leu | synonymous_variant | LOW | 0.0327869 | 0 | 183 | 195 | . | . | . | . |
| PM1548-SK1Org2 | MAP2 | p.Gly1784Ser | missense_variant | MODERATE | 0.00980392 | 0 | 204 | 93 | . | . | . | . |
| PM1548-SK1Org2 | SMARCAL1 | p.Arg499Gln | missense_variant | MODERATE | 0.561644 | 0 | 146 | 137 | . | . | . | . |
| PM1548-SK1Org2 | AAMP | p.Glu63Asp | missense_variant | MODERATE | 0.0218978 | 0.00606061 | 274 | 165 | . | . | . | . |
| PM1548-SK1Org2 | ZNF142 | p.Ala335Pro | missense_variant | MODERATE | 0 | 0 | 234 | 317 | . | . | . | . |
| PM1548-SK1Org2 | CYP27A1 | p.Ser203Leu | missense_variant | MODERATE | 0.481132 | 0 | 212 | 38 | . | . | . | . |
| PM1548-SK1Org2 | WNT6 | p.Asp283Val | missense_variant | MODERATE | 0.530303 | 0 | 66 | 120 | . | . | . | . |
| PM1548-SK1Org2 | FAM134A | p.Asp391Val | missense_variant | MODERATE | 0.477064 | 0 | 327 | 285 | . | . | . | . |
| PM1548-SK1Org2 | DNPEP | p.Arg191Gln | missense_variant | MODERATE | 0 | 0 | 420 | 296 | . | . | . | . |
| PM1548-SK1Org2 | CHPF | p.Arg721Trp | missense_variant | MODERATE | 0.15 | 0 | 140 | 74 | . | . | . | . |
| PM1548-SK1Org2 | WDFY1 | p.Ala75Asp | missense_variant | MODERATE | 0 | 0 | 170 | 181 | . | . | . | . |
| PM1548-SK1Org2 | SERPINE2 | p.Met296Val | missense_variant | MODERATE | 0.423077 | 0 | 182 | 110 | . | . | . | . |
| PM1548-SK1Org2 | B3GNT7 | p.Gly302Asp | missense_variant | MODERATE | 0.573529 | 0 | 136 | 58 | . | . | . | . |
| PM1548-SK1Org2 | DGKD | p.Arg935Trp | missense_variant | MODERATE | 0 | 0 | 89 | 67 | . | . | . | . |
| PM1548-SK1Org2 | ACKR3 | p.Val42Ala | missense_variant | MODERATE | 0.536842 | 0 | 190 | 137 | true | . | . | . |
| PM1548-SK1Org2 | COL6A3 | p.Gly1714Glu | missense_variant | MODERATE | 0.527559 | 0 | 127 | 48 | . | . | . | . |
| PM1548-SK1Org2 | COL6A3 | p.Ile1609Asn | missense_variant | MODERATE | 0.454106 | 0 | 207 | 113 | . | . | . | . |
| PM1548-SK1Org2 | ING5 | p.Arg106His | missense_variant | MODERATE | 0 | 0 | 231 | 105 | . | . | . | . |
| PM1548-SK1Org2 | TBC1D20 | p.Pro71Ser | missense_variant | MODERATE | 0 | 0 | 77 | 49 | . | . | . | . |
| PM1548-SK1Org2 | VPS16 | p.Thr818Met | missense_variant | MODERATE | 0.543624 | 0 | 149 | 238 | . | . | . | . |
| PM1548-SK1Org2 | ATRN | p.Ala122Thr | missense_variant | MODERATE | 0.013986 | 0.00645161 | 143 | 155 | . | . | . | . |
| PM1548-SK1Org2 | SLC23A2 | p.Ile412fs | frameshift_variant | HIGH | 0.422222 | 0 | 90 | 149 | . | . | . | . |
| PM1548-SK1Org2 | BMP2 | p.Ala10Val | missense_variant | MODERATE | 0 | 0 | 61 | 162 | . | . | . | . |
| PM1548-SK1Org2 | PLCB1 | p.Arg768Gln | missense_variant | MODERATE | 0 | 0 | 72 | 91 | . | . | . | . |
| PM1548-SK1Org2 | JAG1 | p.Gly537Ser | missense_variant | MODERATE | 0 | 0 | 138 | 184 | . | . | . | . |
| PM1548-SK1Org2 | CSRP2BP | p.Trp173* | stop_gained | HIGH | 0.268519 | 0 | 108 | 168 | . | . | . | . |
| PM1548-SK1Org2 | CRNKL1 | p.Phe352Leu | missense_variant | MODERATE | 0.334764 | 0 | 233 | 71 | . | . | . | . |
| PM1548-SK1Org2 | PLAGL2 | p.Arg180His | missense_variant | MODERATE | 0.477778 | 0 | 180 | 145 | . | . | . | . |
| PM1548-SK1Org2 | BPIFA2 | p.Arg221His | missense_variant | MODERATE | 0.452381 | 0 | 210 | 61 | . | . | . | . |
| PM1548-SK1Org2 | NECAB3 | p.Arg278fs | frameshift_variant | HIGH | 0.380435 | 0 | 92 | 39 | . | . | . | . |
| PM1548-SK1Org2 | CEP250 | p.Lys787Glu | missense_variant | MODERATE | 0.00564972 | 0 | 177 | 64 | . | . | . | . |
| PM1548-SK1Org2 | SOGA1 | p.Pro974Ser | missense_variant&splice_region_variant | MODERATE | 0.111475 | 0 | 305 | 259 | . | . | . | . |
| PM1548-SK1Org2 | PLCG1 | p.Ile942Met | missense_variant | MODERATE | 0.109589 | 0 | 73 | 138 | true | . | . | . |
| PM1548-SK1Org2 | ZSWIM3 | p.Arg225Gln | missense_variant | MODERATE | 0.492857 | 0 | 280 | 55 | . | . | . | . |
| PM1548-SK1Org2 | NFATC2 | p.Phe158Tyr | missense_variant | MODERATE | 0.166667 | 0 | 60 | 59 | true | . | . | . |
| PM1548-SK1Org2 | NFATC2 | p.Gly22fs | frameshift_variant | HIGH | 0.313131 | 0 | 99 | 146 | true | . | . | . |
| PM1548-SK1Org2 | SALL4 | p.Phe391Leu | missense_variant | MODERATE | 0.210145 | 0 | 138 | 33 | . | . | . | . |
| PM1548-SK1Org2 | ZFP64 | n.50701484A>G | intragenic_variant | MODIFIER | 0.6 | 0 | 40 | 142 | . | . | . | . |
| PM1548-SK1Org2 | DOK5 | p.Val169Ala | missense_variant | MODERATE | 0.415 | 0 | 200 | 338 | . | . | . | . |
| PM1548-SK1Org2 | DOK5 | p.Ala267Asp | missense_variant | MODERATE | 0.296703 | 0 | 182 | 145 | . | . | . | . |
| PM1548-SK1Org2 | TFAP2C | p.Ala354Val | missense_variant | MODERATE | 0.505682 | 0 | 176 | 269 | . | . | . | . |
| PM1548-SK1Org2 | PMEPA1 | p.Ser209fs | frameshift_variant | HIGH | 0.0380952 | 0 | 105 | 249 | . | . | . | . |
| PM1548-SK1Org2 | NELFCD | p.Val321Ile | missense_variant | MODERATE | 0.498024 | 0 | 253 | 73 | . | . | . | . |
| PM1548-SK1Org2 | ZNF831 | p.Leu434Met | missense_variant | MODERATE | 0.464286 | 0 | 84 | 129 | . | . | . | . |
| PM1548-SK1Org2 | HELZ2 | p.Ala639Thr | missense_variant | MODERATE | 0.213592 | 0 | 103 | 139 | . | . | . | . |
| PM1548-SK1Org2 | RTEL1 | p.Arg729Cys | missense_variant | MODERATE | 0.598214 | 0 | 112 | 260 | . | . | . | . |
| PM1548-SK1Org2 | TNFRSF6B | p.Leu297Phe | missense_variant | MODERATE | 0 | 0 | 107 | 277 | . | . | . | . |
| PM1548-SK1Org2 | SLC2A4RG | p.Arg82fs | frameshift_variant | HIGH | 0.452381 | 0 | 42 | 206 | . | . | . | . |
| PM1548-SK1Org2 | PRPF6 | p.Arg317His | missense_variant | MODERATE | 0.268722 | 0 | 227 | 86 | . | . | . | . |
| PM1548-SK1Org2 | LTN1 | p.Gly1705Val | missense_variant | MODERATE | 0.430464 | 0 | 151 | 36 | . | . | . | . |
| PM1548-SK1Org2 | TIAM1 | p.His1010Tyr | missense_variant | MODERATE | 0.248663 | 0 | 374 | 418 | . | . | . | . |
| PM1548-SK1Org2 | URB1 | p.Pro2036Leu | missense_variant | MODERATE | 0.484472 | 0 | 161 | 109 | . | . | . | . |
| PM1548-SK1Org2 | PAXBP1 | p.Pro161Thr | missense_variant | MODERATE | 0 | 0 | 36 | 40 | . | . | . | . |
| PM1548-SK1Org2 | PAXBP1 | p.Ser105Arg | missense_variant | MODERATE | 0.0344828 | 0 | 29 | 158 | . | . | . | . |
| PM1548-SK1Org2 | GART | p.Glu297Gln | missense_variant | MODERATE | 0.148148 | 0 | 81 | 160 | . | . | . | . |
| PM1548-SK1Org2 | SON | p.Thr790Ser | missense_variant | MODERATE | 0.0328947 | 0 | 304 | 177 | . | . | . | . |
| PM1548-SK1Org2 | SON | c.6657+163T>C | intron_variant | MODIFIER | 0.470968 | 0 | 155 | 33 | . | . | . | . |
| PM1548-SK1Org2 | ITSN1 | p.Met768Thr | missense_variant | MODERATE | 0.313559 | 0 | 118 | 46 | . | . | . | true |
| PM1548-SK1Org2 | SLC5A3 | p.Arg252Gln | missense_variant | MODERATE | 0.549618 | 0 | 131 | 44 | . | . | . | . |
| PM1548-SK1Org2 | UMODL1 | p.Gly1319Ser | missense_variant | MODERATE | 0.442211 | 0 | 199 | 131 | . | . | . | . |
| PM1548-SK1Org2 | CRYAA | p.Ile61Val | missense_variant | MODERATE | 0.421296 | 0 | 216 | 344 | . | . | . | . |
| PM1548-SK1Org2 | SLC19A1 | p.Asp488Gly | missense_variant | MODERATE | 0.211538 | 0 | 52 | 131 | . | . | . | . |
| PM1548-SK1Org2 | FTCD | p.Ile187Thr | missense_variant | MODERATE | 0.39726 | 0 | 73 | 175 | . | . | . | . |
| PM1548-SK1Org2 | LSS | p.Thr457Met | missense_variant | MODERATE | 0.393064 | 0 | 173 | 295 | . | . | . | . |
| PM1548-SK1Org2 | CECR2 | p.Ser425Ser | synonymous_variant | LOW | 0.0727273 | 0 | 55 | 131 | . | . | . | . |
| PM1548-SK1Org2 | CECR2 | p.Pro428Pro | synonymous_variant | LOW | 0.0727273 | 0 | 55 | 130 | . | . | . | . |
| PM1548-SK1Org2 | CECR2 | p.Tyr431Tyr | synonymous_variant | LOW | 0.0727273 | 0 | 55 | 130 | . | . | . | . |
| PM1548-SK1Org2 | ATP6V1E1 | p.Leu128Met | missense_variant | MODERATE | 0.537445 | 0 | 227 | 148 | . | . | . | . |
| PM1548-SK1Org2 | CLTCL1 | p.Ala1567Thr | missense_variant | MODERATE | 0.495627 | 0 | 343 | 279 | true | . | . | . |
| PM1548-SK1Org2 | SCARF2 | p.Arg335Cys | missense_variant | MODERATE | 0 | 0 | 82 | 100 | . | . | . | . |
| PM1548-SK1Org2 | SLC7A4 | p.Ala256Thr | missense_variant | MODERATE | 0.548673 | 0 | 226 | 223 | . | . | . | . |
| PM1548-SK1Org2 | IGLL1 | p.Arg60Gln | missense_variant | MODERATE | 0 | 0 | 165 | 81 | . | . | . | . |
| PM1548-SK1Org2 | C22orf43 | p.Ser153Gly | missense_variant | MODERATE | 0 | 0 | 118 | 97 | . | . | . | . |
| PM1548-SK1Org2 | CABIN1 | p.Pro585Gln | missense_variant | MODERATE | 0 | 0 | 37 | 59 | . | . | . | . |
| PM1548-SK1Org2 | ASPHD2 | p.Arg358Trp | missense_variant | MODERATE | 0.460177 | 0 | 226 | 411 | . | . | . | . |
| PM1548-SK1Org2 | TFIP11 | p.Arg39* | stop_gained | HIGH | 0.539326 | 0 | 178 | 103 | . | . | . | . |
| PM1548-SK1Org2 | RASL10A | p.Arg108Trp | missense_variant | MODERATE | 0 | 0 | 206 | 108 | . | . | . | . |
| PM1548-SK1Org2 | AP1B1 | p.Ala655Asp | missense_variant | MODERATE | 0 | 0 | 96 | 31 | . | . | . | . |
| PM1548-SK1Org2 | LIMK2 | p.Asn477Ser | missense_variant | MODERATE | 0.041958 | 0 | 143 | 257 | . | . | . | . |
| PM1548-SK1Org2 | SFI1 | p.Thr710Ser | missense_variant | MODERATE | 0.36478 | 0 | 159 | 103 | . | . | . | . |
| PM1548-SK1Org2 | CSF2RB | p.Gln642fs | frameshift_variant | HIGH | 0.506329 | 0 | 158 | 133 | . | . | . | . |
| PM1548-SK1Org2 | C1QTNF6 | p.Met229Thr | missense_variant | MODERATE | 0.642857 | 0 | 14 | 20 | . | . | . | . |
| PM1548-SK1Org2 | GRAP2 | p.Val129Ile | missense_variant | MODERATE | 0.456 | 0 | 125 | 162 | . | . | . | . |
| PM1548-SK1Org2 | MKL1 | p.Pro138Leu | missense_variant | MODERATE | 0.578431 | 0 | 102 | 100 | true | . | . | . |
| PM1548-SK1Org2 | EP300 | p.Pro2107Thr | missense_variant | MODERATE | 0 | 0 | 216 | 94 | true | true | true | . |
| PM1548-SK1Org2 | ZC3H7B | p.Arg908Cys | missense_variant | MODERATE | 0.597561 | 0 | 164 | 73 | . | . | . | . |
| PM1548-SK1Org2 | PARVB | c.211+24725C>A | intron_variant | MODIFIER | 0.47619 | 0 | 42 | 165 | . | . | . | . |
| PM1548-SK1Org2 | CELSR1 | p.Val1411Gly | missense_variant | MODERATE | 0.241379 | 0 | 174 | 333 | . | . | . | . |
| PM1548-SK1Org2 | CELSR1 | p.Glu1159Lys | missense_variant | MODERATE | 0.596774 | 0 | 124 | 186 | . | . | . | . |
| PM1548-SK1Org2 | TUBGCP6 | p.Cys1239Trp | missense_variant | MODERATE | 0 | 0.00947867 | 130 | 211 | . | . | . | . |
| PM1548-SK1Org2 | TUBGCP6 | p.Ser1237Pro | missense_variant | MODERATE | 0 | 0 | 211 | 238 | . | . | . | . |
| PM1548-SK1Org2 | MIOX | p.Asp78Gly | missense_variant | MODERATE | 0 | 0 | 95 | 102 | . | . | . | . |
| PM1548-SK1Org2 | MAPK8IP2 | p.Thr518Thr | synonymous_variant | LOW | 0.168831 | 0 | 77 | 166 | . | . | . | . |
| PM1548-SK1Org2 | ARSA | c.684+1G>A | splice_donor_variant&intron_variant | HIGH | 0.470588 | 0 | 34 | 160 | . | . | . | . |
| PM1548-SK1Org2 | SETD5 | p.Arg1193* | stop_gained | HIGH | 0.556818 | 0 | 176 | 243 | . | . | . | . |
| PM1548-SK1Org2 | CPNE9 | p.Ser284Ser | synonymous_variant | LOW | 0.046875 | 0 | 128 | 197 | . | . | . | . |
| PM1548-SK1Org2 | CPNE9 | p.Asp473Asp | synonymous_variant | LOW | 0.484615 | 0 | 260 | 85 | . | . | . | . |
| PM1548-SK1Org2 | BRPF1 | p.Gln938fs | frameshift_variant | HIGH | 0.459459 | 0 | 111 | 171 | . | . | . | true |
| PM1548-SK1Org2 | OGG1 | p.Ala85Thr | missense_variant | MODERATE | 0.485507 | 0 | 276 | 334 | . | . | . | . |
| PM1548-SK1Org2 | CRELD1 | p.Arg220* | stop_gained | HIGH | 0.436242 | 0 | 298 | 387 | . | . | . | . |
| PM1548-SK1Org2 | PRRT3 | p.His679Arg | missense_variant | MODERATE | 0.211268 | 0 | 71 | 166 | . | . | . | . |
| PM1548-SK1Org2 | CAND2 | p.Arg489His | missense_variant | MODERATE | 0.5 | 0 | 272 | 331 | . | . | . | . |
| PM1548-SK1Org2 | SLC6A6 | p.Phe392fs | frameshift_variant | HIGH | 0.504065 | 0 | 123 | 115 | . | . | . | . |
| PM1548-SK1Org2 | NR2C2 | p.Phe582fs | frameshift_variant | HIGH | 0.547945 | 0 | 146 | 75 | . | . | . | . |
| PM1548-SK1Org2 | SH3BP5 | p.Thr119Ala | missense_variant | MODERATE | 0.503333 | 0 | 300 | 345 | . | . | . | . |
| PM1548-SK1Org2 | PLCL2 | p.Gln628Arg | missense_variant | MODERATE | 0 | 0 | 27 | 118 | . | . | . | . |
| PM1548-SK1Org2 | SLC4A7 | p.Ile1097Val | missense_variant | MODERATE | 0.284314 | 0.00657895 | 102 | 152 | . | . | . | . |
| PM1548-SK1Org2 | CLASP2 | p.Cys10Cys | synonymous_variant | LOW | 0.333333 | 0.00520833 | 45 | 192 | . | . | . | . |
| PM1548-SK1Org2 | SCN10A | p.Val378Leu | missense_variant | MODERATE | 0.015873 | 0.00809717 | 189 | 247 | . | . | . | . |
| PM1548-SK1Org2 | TTC21A | p.Ser332Thr | missense_variant | MODERATE | 0.125749 | 0 | 167 | 40 | . | . | . | . |
| PM1548-SK1Org2 | SNRK | p.Pro598Thr | missense_variant | MODERATE | 0 | 0 | 42 | 93 | . | . | . | . |
| PM1548-SK1Org2 | SLC6A20 | p.Asn103Lys | missense_variant | MODERATE | 0 | 0 | 69 | 172 | . | . | . | . |
| PM1548-SK1Org2 | SLC6A20 | p.Gly66Trp | missense_variant | MODERATE | 0.127119 | 0.00847458 | 118 | 118 | . | . | . | . |
| PM1548-SK1Org2 | FYCO1 | p.Lys501del | inframe_deletion | MODERATE | 0.554622 | 0 | 238 | 171 | . | . | . | . |
| PM1548-SK1Org2 | ALS2CL | p.Asn384Lys | missense_variant | MODERATE | 0.538136 | 0 | 236 | 77 | . | . | . | . |
| PM1548-SK1Org2 | DHX30 | n.47859525delC | intragenic_variant | MODIFIER | 0.448905 | 0 | 274 | 119 | . | . | . | . |
| PM1548-SK1Org2 | UCN2 | p.Thr27Pro | missense_variant | MODERATE | 0.0382979 | 0 | 235 | 135 | . | . | . | . |
| PM1548-SK1Org2 | CELSR3 | p.Pro1425Ser | missense_variant | MODERATE | 0 | 0 | 107 | 95 | . | . | . | . |
| PM1548-SK1Org2 | SLC25A20 | p.Ala227Val | missense_variant | MODERATE | 0.512346 | 0 | 162 | 125 | . | . | . | . |
| PM1548-SK1Org2 | KLHDC8B | p.Gly38fs | frameshift_variant | HIGH | 0.42439 | 0 | 205 | 198 | . | . | . | . |
| PM1548-SK1Org2 | RRP9 | p.Ala342Thr | missense_variant | MODERATE | 0.552995 | 0 | 217 | 161 | . | . | . | . |

| | | | | | | | | | | | | |
|---|---|---|---|---|---|---|---|---|---|---|---|---|
| PM1548-SK1Org2 | DNAH1 | p.Leu2072Ile | missense_variant | MODERATE | 0.555556 | 0 | 117 | 99 | . | . | . | . |
| PM1548-SK1Org2 | FAM208A | p.Ala227Val | missense_variant | MODERATE | 0.4 | 0 | 110 | 80 | . | . | . | . |
| PM1548-SK1Org2 | DNAH12 | p.Pro1035Ala | missense_variant | MODERATE | 0 | 0 | 184 | 189 | . | . | . | . |
| PM1548-SK1Org2 | PDE12 | p.Ala293Ser | missense_variant | MODERATE | 0 | 0 | 204 | 150 | . | . | . | . |
| PM1548-SK1Org2 | PDE12 | p.Leu294Phe | missense_variant | MODERATE | 0 | 0 | 204 | 150 | . | . | . | . |
| PM1548-SK1Org2 | ADAMTS9 | p.Pro596His | missense_variant | MODERATE | 0 | 0 | 194 | 126 | . | . | . | . |
| PM1548-SK1Org2 | ADAMTS9 | p.Ser593Gly | missense_variant | MODERATE | 0 | 0 | 213 | 130 | . | . | . | . |
| PM1548-SK1Org2 | ROBO1 | p.Tyr1009* | stop_gained | HIGH | 0 | 0 | 44 | 54 | . | . | . | . |
| PM1548-SK1Org2 | OR5K4 | p.Val51Ile | missense_variant | MODERATE | 0.003003 | 0 | 333 | 442 | . | . | . | . |
| PM1548-SK1Org2 | OR5K4 | p.Arg53Pro | missense_variant | MODERATE | 0.003003 | 0 | 333 | 441 | . | . | . | . |
| PM1548-SK1Org2 | OR5K4 | p.Leu56His | missense_variant | MODERATE | 0.0026178 | 0 | 382 | 510 | . | . | . | . |
| PM1548-SK1Org2 | DCBLD2 | p.Thr718Met | missense_variant | MODERATE | 0.470356 | 0 | 253 | 81 | . | . | . | . |
| PM1548-SK1Org2 | TFG | p.Ser213Pro | missense_variant | MODERATE | 0.508 | 0 | 250 | 76 | true | . | . | . |
| PM1548-SK1Org2 | CBLB | p.Pro823Ser | missense_variant | MODERATE | 0.188679 | 0 | 159 | 275 | true | . | . | . |
| PM1548-SK1Org2 | BOC | p.Pro593Leu | missense_variant | MODERATE | 0.0285714 | 0.00197628 | 210 | 506 | . | . | . | . |
| PM1548-SK1Org2 | BOC | p.His601Gln | missense_variant | MODERATE | 0 | 0 | 140 | 255 | . | . | . | . |
| PM1548-SK1Org2 | KIAA2018 | p.Asn1016fs | frameshift_variant | HIGH | 0.0634921 | 0 | 63 | 36 | . | . | . | . |
| PM1548-SK1Org2 | KIAA1407 | p.Glu683Gly | missense_variant | MODERATE | 0.432432 | 0 | 74 | 85 | . | . | . | . |
| PM1548-SK1Org2 | LSAMP | p.Val277Ile | missense_variant | MODERATE | 0.297297 | 0 | 74 | 91 | . | . | . | . |
| PM1548-SK1Org2 | ARHGAP31 | p.Lys857Glu | missense_variant | MODERATE | 0.530612 | 0 | 49 | 33 | . | . | . | . |
| PM1548-SK1Org2 | FSTL1 | p.Val13Ala | missense_variant | MODERATE | 0.25 | 0 | 76 | 223 | . | . | . | . |
| PM1548-SK1Org2 | EAF2 | p.Thr112fs | frameshift_variant | HIGH | 0.39726 | 0 | 73 | 115 | . | . | . | . |
| PM1548-SK1Org2 | ILDR1 | p.Ala76Val | missense_variant&splice_region_variant | MODERATE | 0.246201 | 0 | 329 | 159 | . | . | . | . |
| PM1548-SK1Org2 | MYLK | p.Ala1684Thr | missense_variant | MODERATE | 0.486111 | 0 | 216 | 98 | . | . | . | . |
| PM1548-SK1Org2 | ALDH1L1 | p.Glu439fs | frameshift_variant | HIGH | 0.551724 | 0 | 87 | 55 | . | . | . | . |
| PM1548-SK1Org2 | CHCHD6 | p.Ser220Leu | missense_variant | MODERATE | 0 | 0 | 161 | 164 | . | . | . | . |
| PM1548-SK1Org2 | C3orf27 | p.Asp85fs | frameshift_variant | HIGH | 0.365385 | 0 | 260 | 229 | . | . | . | . |
| PM1548-SK1Org2 | TMCC1 | p.Arg620Ser | missense_variant | MODERATE | 0 | 0 | 110 | 62 | . | . | . | . |
| PM1548-SK1Org2 | CDV3 | p.Gly226Asp | missense_variant | MODERATE | 0.447552 | 0 | 143 | 44 | . | . | . | . |
| PM1548-SK1Org2 | CDV3 | p.Asp228Glu | missense_variant | MODERATE | 0.39548 | 0 | 177 | 55 | . | . | . | . |
| PM1548-SK1Org2 | AMOTL2 | p.Arg292Cys | missense_variant | MODERATE | 0 | 0 | 126 | 43 | . | . | . | . |
| PM1548-SK1Org2 | EPHB1 | p.Ala82fs | frameshift_variant | HIGH | 0.428571 | 0 | 245 | 293 | . | . | . | . |
| PM1548-SK1Org2 | SLC35G2 | p.Asn97fs | frameshift_variant | HIGH | 0.574627 | 0 | 134 | 31 | . | . | . | . |
| PM1548-SK1Org2 | ESYT3 | p.Val238Met | missense_variant | MODERATE | 0.466667 | 0 | 195 | 78 | . | . | . | . |
| PM1548-SK1Org2 | MED12L | p.Ser970Leu | missense_variant | MODERATE | 0.256579 | 0 | 152 | 121 | . | . | . | . |
| PM1548-SK1Org2 | DHX36 | p.Met154fs | frameshift_variant | HIGH | 0 | 0 | 33 | 44 | . | . | . | . |
| PM1548-SK1Org2 | CCNL1 | p.Ile354Val | missense_variant | MODERATE | 0 | 0 | 32 | 120 | . | . | . | . |
| PM1548-SK1Org2 | PPM1L | p.Glu106Glu | synonymous_variant | LOW | 0.0212766 | 0 | 188 | 275 | . | . | . | . |
| PM1548-SK1Org2 | SLITRK3 | p.Arg759Cys | missense_variant | MODERATE | 0.507937 | 0 | 189 | 99 | . | . | . | . |
| PM1548-SK1Org2 | ZBBX | p.Ile496fs | frameshift_variant | HIGH | 0.482759 | 0 | 58 | 76 | . | . | . | . |
| PM1548-SK1Org2 | FNDC3B | p.Glu687Gly | missense_variant | MODERATE | 0 | 0 | 85 | 96 | . | . | . | . |
| PM1548-SK1Org2 | NCEH1 | p.Ser105fs | frameshift_variant | HIGH | 0.165563 | 0 | 151 | 228 | . | . | . | . |
| PM1548-SK1Org2 | KLHL24 | p.Asn555fs | frameshift_variant | HIGH | 0.473282 | 0 | 131 | 55 | . | . | . | . |
| PM1548-SK1Org2 | CHRD | p.Arg76Cys | missense_variant | MODERATE | 0.516129 | 0 | 93 | 91 | . | . | . | . |
| PM1548-SK1Org2 | TP63 | p.Thr368Ala | missense_variant | MODERATE | 0.0496454 | 0 | 141 | 121 | . | . | . | . |
| PM1548-SK1Org2 | LEPREL1 | p.Asp204Glu | missense_variant | MODERATE | 0 | 0 | 266 | 177 | . | . | . | . |
| PM1548-SK1Org2 | CCDC50 | p.Ala343Ser | missense_variant | MODERATE | 0.015873 | 0 | 63 | 55 | . | . | . | . |
| PM1548-SK1Org2 | MUC20 | p.Gly514Arg | missense_variant | MODERATE | 0.0152672 | 0 | 131 | 228 | . | . | . | . |
| PM1548-SK1Org2 | LRCH3 | p.Leu49Val | missense_variant | MODERATE | 0 | 0.00369004 | 118 | 271 | . | . | . | . |
| PM1548-SK1Org2 | GAK | p.Ala690Val | missense_variant | MODERATE | 0 | 0 | 140 | 111 | . | . | . | . |
| PM1548-SK1Org2 | DGKQ | p.Thr299Met | missense_variant | MODERATE | 0.5 | 0 | 150 | 253 | . | . | . | . |
| PM1548-SK1Org2 | SLC26A1 | p.Ala624Thr | missense_variant | MODERATE | 0 | 0 | 195 | 466 | . | . | . | . |
| PM1548-SK1Org2 | POLN | p.Tyr686Cys | missense_variant | MODERATE | 0 | 0 | 159 | 37 | . | . | . | . |
| PM1548-SK1Org2 | HTT | p.Ala2259Thr | missense_variant&splice_region_variant | MODERATE | 0.507937 | 0 | 63 | 52 | . | . | . | . |
| PM1548-SK1Org2 | PPP2R2C | p.Gly397fs | frameshift_variant | HIGH | 0 | 0 | 124 | 188 | . | . | . | . |
| PM1548-SK1Org2 | SH3TC1 | p.Glu203Gly | missense_variant | MODERATE | 0.49345 | 0.00213675 | 229 | 468 | . | . | . | . |
| PM1548-SK1Org2 | GPR78 | p.Val209Ile | missense_variant | MODERATE | 0.636364 | 0 | 55 | 34 | . | . | . | . |
| PM1548-SK1Org2 | KCTD8 | p.Leu62Leu | synonymous_variant | LOW | 0.03125 | 0 | 64 | 213 | . | . | . | . |
| PM1548-SK1Org2 | YTHDC1 | p.Ser684Asn | missense_variant | MODERATE | 0.577778 | 0 | 135 | 59 | . | . | . | . |
| PM1548-SK1Org2 | ANKRD17 | p.Ile2241Val | missense_variant | MODERATE | 0.0106952 | 0 | 187 | 147 | . | . | . | . |
| PM1548-SK1Org2 | PF4 | p.Ala4Thr | missense_variant | MODERATE | 0.260563 | 0 | 142 | 40 | . | . | . | . |
| PM1548-SK1Org2 | PARM1 | p.Ser82Pro | missense_variant | MODERATE | 0 | 0 | 241 | 388 | . | . | . | . |
| PM1548-SK1Org2 | CDKL2 | p.Pro452Ser | missense_variant | MODERATE | 0 | 0 | 60 | 51 | . | . | . | . |
| PM1548-SK1Org2 | SHROOM3 | p.Lys1017del | inframe_deletion | MODERATE | 0.563636 | 0 | 55 | 97 | . | . | . | . |
| PM1548-SK1Org2 | BMP3 | p.Lys358fs | frameshift_variant | HIGH | 0.507812 | 0 | 128 | 124 | . | . | . | . |
| PM1548-SK1Org2 | PRKG2 | p.Val461Phe | missense_variant | MODERATE | 0 | 0 | 104 | 44 | . | . | . | . |
| PM1548-SK1Org2 | SEC31A | p.Ile462fs | frameshift_variant | HIGH | 0.828947368 | 0 | 152 | 87 | . | . | . | . |
| PM1548-SK1Org2 | SEC31A | p.Lys461fs | frameshift_variant | HIGH | 0.026315789 | 0 | 152 | 87 | . | . | . | . |
| PM1548-SK1Org2 | SEC31A | p.Gln422Pro | missense_variant | MODERATE | 0.196629 | 0 | 178 | 143 | . | . | . | . |
| PM1548-SK1Org2 | AGPAT9 | p.Met372fs | frameshift_variant | HIGH | 0.397516 | 0 | 161 | 70 | . | . | . | . |
| PM1548-SK1Org2 | NAP1L5 | p.Lys108fs | frameshift_variant | HIGH | 0.494595 | 0 | 370 | 237 | . | . | . | . |
| PM1548-SK1Org2 | CASP6 | p.Pro33Leu | missense_variant | MODERATE | 0.283019 | 0.00719424 | 212 | 278 | . | . | . | . |
| PM1548-SK1Org2 | PITX2 | c.411+284C>T | intron_variant | MODIFIER | 0.535714 | 0 | 112 | 59 | . | . | . | . |
| PM1548-SK1Org2 | ALPK1 | p.Tyr1216fs | frameshift_variant | HIGH | 0.333333 | 0 | 36 | 44 | . | . | . | . |
| PM1548-SK1Org2 | TNIP3 | p.Ala59Thr | missense_variant | MODERATE | 0.361111 | 0 | 72 | 33 | . | . | . | . |
| PM1548-SK1Org2 | PCDH10 | p.Tyr324Cys | missense_variant | MODERATE | 0.459016 | 0 | 61 | 49 | . | . | . | . |
| PM1548-SK1Org2 | SETD7 | p.Leu331Ile | missense_variant | MODERATE | 0.529126 | 0 | 206 | 328 | . | . | . | . |
| PM1548-SK1Org2 | FREM3 | p.Arg502* | stop_gained | HIGH | 0 | 0 | 125 | 34 | . | . | . | . |
| PM1548-SK1Org2 | NEK1 | p.Thr137Asn | missense_variant | MODERATE | 0.0405405 | 0 | 74 | 84 | . | . | . | . |
| PM1548-SK1Org2 | AHRR | p.Arg72His | missense_variant | MODERATE | 0.417808 | 0 | 146 | 75 | . | . | . | . |
| PM1548-SK1Org2 | EXOC3 | p.Val576Met | missense_variant | MODERATE | 0.614458 | 0 | 166 | 107 | . | . | . | . |
| PM1548-SK1Org2 | LPCAT1 | p.Val411Ile | missense_variant | MODERATE | 0 | 0 | 96 | 136 | . | . | . | . |
| PM1548-SK1Org2 | IRX2 | p.Tyr110Cys | missense_variant | MODERATE | 0.43662 | 0 | 71 | 189 | . | . | . | . |
| PM1548-SK1Org2 | NSUN2 | p.Ala653Thr | missense_variant&splice_region_variant | MODERATE | 0.173077 | 0 | 104 | 83 | . | . | . | . |
| PM1548-SK1Org2 | SRD5A1 | p.Met162Ile | missense_variant | MODERATE | 0.617886 | 0 | 123 | 54 | . | . | . | . |
| PM1548-SK1Org2 | SEMA5A | p.Ile679Thr | missense_variant | MODERATE | 0 | 0 | 153 | 101 | . | . | . | . |
| PM1548-SK1Org2 | NIPBL | p.Glu1060Glu | synonymous_variant | LOW | 0.0357143 | 0 | 56 | 121 | . | . | . | . |
| PM1548-SK1Org2 | CARD6 | p.Gln393Glu | missense_variant | MODERATE | 0.00623053 | 0 | 321 | 371 | . | . | . | . |
| PM1548-SK1Org2 | CARD6 | p.Gln400His | missense_variant | MODERATE | 0.00671141 | 0 | 298 | 367 | . | . | . | . |
| PM1548-SK1Org2 | CARD6 | p.Gln402* | stop_gained | HIGH | 0.00668896 | 0 | 299 | 367 | . | . | . | . |
| PM1548-SK1Org2 | CARD6 | p.Gln402Arg | missense_variant | MODERATE | 0.00743494 | 0 | 269 | 337 | . | . | . | . |
| PM1548-SK1Org2 | CARD6 | p.Gln402His | missense_variant | MODERATE | 0.00746269 | 0 | 268 | 337 | . | . | . | . |
| PM1548-SK1Org2 | C6 | p.Ala2Val | missense_variant | MODERATE | 0.363636 | 0 | 110 | 129 | . | . | . | . |
| PM1548-SK1Org2 | SNX18 | n.53839150G>A | intragenic_variant | MODIFIER | 0.461078 | 0 | 167 | 117 | . | . | . | . |
| PM1548-SK1Org2 | GPX8 | p.Phe125fs | frameshift_variant | HIGH | 0.456044 | 0 | 182 | 228 | . | . | . | . |
| PM1548-SK1Org2 | UTP15 | c.169-826A>G | intron_variant | MODIFIER | 0 | 0 | 139 | 55 | . | . | . | . |
| PM1548-SK1Org2 | TTC37 | p.Ala1407Thr | missense_variant | MODERATE | 1 | 0 | 46 | 32 | . | . | . | . |
| PM1548-SK1Org2 | CHD1 | p.Asn310fs | frameshift_variant | HIGH | 1 | 0 | 90 | 201 | . | . | . | true |
| PM1548-SK1Org2 | PAM | p.Arg260Gln | missense_variant | MODERATE | 0 | 0 | 96 | 107 | . | . | . | . |
| PM1548-SK1Org2 | GIN1 | p.Leu345Pro | missense_variant | MODERATE | 0.407407 | 0 | 81 | 72 | . | . | . | . |
| PM1548-SK1Org2 | APC | p.Asp802fs | frameshift_variant&stop_gained | HIGH | 0.594406 | 0 | 143 | 125 | true | true | true | true |
| PM1548-SK1Org2 | MCC | p.Arg800Lys | missense_variant | MODERATE | 0.360656 | 0 | 61 | 65 | . | . | . | . |
| PM1548-SK1Org2 | FTMT | p.Ala90Val | missense_variant | MODERATE | 0.496063 | 0 | 127 | 105 | . | . | . | . |
| PM1548-SK1Org2 | FBN2 | p.Leu2853Arg | missense_variant | MODERATE | 0.443515 | 0 | 239 | 91 | . | . | . | . |
| PM1548-SK1Org2 | FNIP1 | p.Leu90Met | missense_variant | MODERATE | 0 | 0 | 27 | 76 | . | . | . | . |
| PM1548-SK1Org2 | JADE2 | p.Phe360Leu | missense_variant | MODERATE | 0 | 0 | 208 | 343 | . | . | . | true |
| PM1548-SK1Org2 | FBXL21 | n.956C>T | non_coding_exon_variant | MODIFIER | 0.444444 | 0 | 54 | 86 | . | . | . | . |
| PM1548-SK1Org2 | KDM3B | p.Arg921Trp | missense_variant | MODERATE | 0 | 0 | 164 | 178 | . | . | . | . |
| PM1548-SK1Org2 | PSD2 | p.Arg574His | missense_variant | MODERATE | 0.52381 | 0.00649351 | 105 | 154 | . | . | . | . |
| PM1548-SK1Org2 | PCDHA3 | p.Glu558Val | missense_variant | MODERATE | 0.00529101 | 0 | 189 | 71 | . | . | . | . |
| PM1548-SK1Org2 | PCDHA5 | p.Ala56Thr | missense_variant | MODERATE | 0.285714 | 0 | 84 | 32 | . | . | . | . |
| PM1548-SK1Org2 | PCDHA5 | p.Arg64Arg | synonymous_variant | LOW | 0.201439 | 0 | 139 | 38 | . | . | . | . |
| PM1548-SK1Org2 | PCDHA5 | p.Gly69Asp | missense_variant | MODERATE | 0.170732 | 0 | 164 | 45 | . | . | . | . |
| PM1548-SK1Org2 | PCDHA8 | p.Ala57Thr | missense_variant | MODERATE | 0.0215054 | 0 | 186 | 46 | . | . | . | . |
| PM1548-SK1Org2 | PCDHA13 | p.Glu558Val | missense_variant | MODERATE | 0 | 0 | 122 | 106 | . | . | . | . |
| PM1548-SK1Org2 | PCDHAC2 | p.Arg543Trp | missense_variant | MODERATE | 0.505376 | 0 | 93 | 153 | . | . | . | . |
| PM1548-SK1Org2 | PCDHAC2 | c.2565+37G>A | intron_variant | MODIFIER | 0.5 | 0 | 70 | 61 | . | . | . | . |
| PM1548-SK1Org2 | PCDHGA1 | p.Ser370Gly | missense_variant | MODERATE | 0.455357 | 0 | 224 | 385 | . | . | . | . |
| PM1548-SK1Org2 | PCDHGA9 | p.Val422Ala | missense_variant | MODERATE | 0 | 0 | 39 | 94 | . | . | . | . |
| PM1548-SK1Org2 | PCDH1 | p.Ala203Thr | missense_variant | MODERATE | 0.483051 | 0 | 236 | 253 | . | . | . | . |
| PM1548-SK1Org2 | PCDH12 | p.Gly10Glu | missense_variant | MODERATE | 0.233129 | 0 | 163 | 44 | . | . | . | . |
| PM1548-SK1Org2 | CSF1R | p.Glu955Lys | missense_variant | MODERATE | 0.383721 | 0 | 86 | 65 | . | . | . | . |
| PM1548-SK1Org2 | TIMD4 | p.Ala333Val | missense_variant | MODERATE | 0.490854 | 0 | 328 | 229 | . | . | . | . |
| PM1548-SK1Org2 | ATP10B | p.Arg115Lys | missense_variant | MODERATE | 0.257576 | 0 | 198 | 155 | . | . | . | . |
| PM1548-SK1Org2 | UBTD2 | p.Thr160Ile | missense_variant | MODERATE | 0.478261 | 0 | 138 | 33 | . | . | . | . |
| PM1548-SK1Org2 | GPRIN1 | p.Pro806fs | frameshift_variant | HIGH | 0 | 0 | 20 | 165 | . | . | . | . |
| PM1548-SK1Org2 | GPRIN1 | p.Pro805Thr | missense_variant | MODERATE | 0 | 0 | 20 | 165 | . | . | . | . |

| | | | | | | | | | | | | |
|---|---|---|---|---|---|---|---|---|---|---|---|---|
| PM1548-SK1Org2 | GPRIN1 | p.Ser803fs | frameshift_variant | HIGH | 0 | 0 | 20 | 165 | . | . | . | . |
| PM1548-SK1Org2 | GPRIN1 | p.Glu801Asp | missense_variant | MODERATE | 0 | 0 | 20 | 165 | . | . | . | . |
| PM1548-SK1Org2 | UIMC1 | p.Phe165Cys | missense_variant | MODERATE | 0.0288462 | 0 | 208 | 180 | . | . | . | . |
| PM1548-SK1Org2 | UIMC1 | p.Ile156Val | missense_variant&splice_region_variant | MODERATE | 0.0339806 | 0 | 206 | 259 | . | . | . | . |
| PM1548-SK1Org2 | FGFR4 | p.Pro568fs | frameshift_variant | HIGH | 0.39375 | 0 | 160 | 173 | true | . | . | . |
| PM1548-SK1Org2 | FAM193B | p.Pro761Ser | missense_variant | MODERATE | 0 | 0 | 212 | 84 | . | . | . | . |
| PM1548-SK1Org2 | GRM6 | p.Ala93Thr | missense_variant | MODERATE | 0.56 | 0 | 25 | 91 | . | . | . | . |
| PM1548-SK1Org2 | RREB1 | p.Val754Met | missense_variant | MODERATE | 0.347826 | 0 | 46 | 159 | . | . | . | . |
| PM1548-SK1Org2 | HIST1H4C | p.Val82Ala | missense_variant | MODERATE | 0.428571 | 0 | 112 | 187 | . | . | . | . |
| PM1548-SK1Org2 | HIST1H2AM | p.Leu52Met | missense_variant | MODERATE | 0.0181818 | 0 | 110 | 43 | . | . | . | . |
| PM1548-SK1Org2 | TRIM27 | p.Ala24Val | missense_variant | MODERATE | 0 | 0 | 62 | 147 | true | . | . | . |
| PM1548-SK1Org2 | OR2H1 | p.Ile295Leu | missense_variant | MODERATE | 0.471429 | 0 | 140 | 181 | . | . | . | . |
| PM1548-SK1Org2 | HLA-F | p.Leu396fs | frameshift_variant | HIGH | 0.546053 | 0 | 152 | 38 | . | . | . | . |
| PM1548-SK1Org2 | HLA-G | p.Arg285Ser | missense_variant | MODERATE | 0.0625 | 0 | 192 | 42 | . | . | . | . |
| PM1548-SK1Org2 | TRIM26 | p.Glu241Lys | missense_variant | MODERATE | 0.586667 | 0 | 150 | 200 | . | . | . | . |
| PM1548-SK1Org2 | MDC1 | p.Gln1431Arg | missense_variant | MODERATE | 0.00333333 | 0 | 300 | 163 | . | . | . | . |
| PM1548-SK1Org2 | MUC22 | p.Ile1250Thr | missense_variant | MODERATE | 0.0574713 | 0 | 87 | 195 | . | . | . | . |
| PM1548-SK1Org2 | HLA-C | p.Arg243Trp | missense_variant | MODERATE | 0 | 0 | 20 | 40 | . | . | . | . |
| PM1548-SK1Org2 | HLA-B | p.Arg243Trp | missense_variant | MODERATE | 0.0581395 | 0.00879765 | 172 | 341 | . | . | true | . |
| PM1548-SK1Org2 | VWA7 | p.Thr26fs | frameshift_variant | HIGH | 0.207746 | 0 | 284 | 239 | . | . | . | . |
| PM1548-SK1Org2 | CFB | p.Asp1161fs | frameshift_variant | HIGH | 0.48731 | 0 | 197 | 133 | . | . | . | . |
| PM1548-SK1Org2 | RGL2 | p.Glu372fs | frameshift_variant | HIGH | 0.390625 | 0 | 256 | 190 | . | . | . | . |
| PM1548-SK1Org2 | DNAH8 | p.Tyr1705Cys | missense_variant | MODERATE | 0 | 0 | 152 | 68 | . | . | . | . |
| PM1548-SK1Org2 | KLHDC3 | p.Arg67Cys | missense_variant | MODERATE | 0.492308 | 0 | 130 | 262 | . | . | . | . |
| PM1548-SK1Org2 | PKHD1 | p.Arg880His | missense_variant | MODERATE | 0.582677 | 0 | 127 | 253 | . | . | . | . |
| PM1548-SK1Org2 | DST | p.Thr1051fs | frameshift_variant | HIGH | 0.428571 | 0 | 217 | 228 | . | . | . | . |
| PM1548-SK1Org2 | KIAA1586 | p.Arg617Leu | missense_variant | MODERATE | 0 | 0 | 68 | 71 | . | . | . | . |
| PM1548-SK1Org2 | ZNF451 | p.Phe193Leu | missense_variant&splice_region_variant | MODERATE | 0.553398 | 0 | 103 | 32 | . | . | . | . |
| PM1548-SK1Org2 | COL12A1 | p.Pro2509Thr | missense_variant | MODERATE | 0 | 0 | 140 | 157 | . | . | . | . |
| PM1548-SK1Org2 | SENP6 | p.Ser627Gly | missense_variant | MODERATE | 0 | 0 | 15 | 66 | . | . | . | . |
| PM1548-SK1Org2 | ME1 | c.78+1G>A | splice_donor_variant&intron_variant | HIGH | 0.503676 | 0 | 272 | 306 | . | . | . | . |
| PM1548-SK1Org2 | PRSS35 | p.Arg265* | stop_gained | HIGH | 0.398148 | 0 | 108 | 41 | . | . | . | . |
| PM1548-SK1Org2 | ANKRD6 | p.Ala390Val | missense_variant | MODERATE | 0.295082 | 0 | 61 | 31 | . | . | . | . |
| PM1548-SK1Org2 | MDN1 | p.Thr2167Lys | missense_variant | MODERATE | 0.0571429 | 0 | 70 | 72 | . | . | . | . |
| PM1548-SK1Org2 | MMS22L | p.Gly37Glu | missense_variant | MODERATE | 0.502326 | 0 | 215 | 237 | . | . | . | . |
| PM1548-SK1Org2 | ATG5 | p.Asp149fs | frameshift_variant | HIGH | 0 | 0 | 97 | 95 | . | . | . | . |
| PM1548-SK1Org2 | RFPL4B | p.Arg203Cys | missense_variant | MODERATE | 0.36 | 0 | 50 | 78 | . | . | . | . |
| PM1548-SK1Org2 | KPNA5 | p.Phe383Ser | missense_variant | MODERATE | 0.306931 | 0 | 101 | 67 | . | . | . | . |
| PM1548-SK1Org2 | FAM184A | p.Glu79Asp | missense_variant | MODERATE | 0.296053 | 0 | 152 | 52 | . | . | . | . |
| PM1548-SK1Org2 | TBC1D32 | p.Ile27Asn | missense_variant | MODERATE | 0.464286 | 0 | 84 | 41 | . | . | . | . |
| PM1548-SK1Org2 | HSF2 | p.Arg196His | missense_variant | MODERATE | 0.285714 | 0 | 35 | 76 | . | . | . | . |
| PM1548-SK1Org2 | KIAA0408 | p.Phe313Leu | missense_variant | MODERATE | 0 | 0 | 115 | 109 | . | . | . | . |
| PM1548-SK1Org2 | SOGA3 | p.Ala214fs | frameshift_variant | HIGH | 0.146341 | 0 | 41 | 41 | . | . | . | . |
| PM1548-SK1Org2 | ENPP1 | c.240+2T>C | splice_donor_variant&intron_variant | HIGH | 0.404762 | 0 | 84 | 63 | . | . | . | . |
| PM1548-SK1Org2 | CTGF | p.Ser227Phe | missense_variant | MODERATE | 0.00606061 | 0 | 165 | 36 | . | . | . | . |
| PM1548-SK1Org2 | CTGF | p.Ala226Thr | missense_variant | MODERATE | 0.00507614 | 0 | 197 | 46 | . | . | . | . |
| PM1548-SK1Org2 | CTGF | p.Ile195Met | missense_variant | MODERATE | 0.010101 | 0 | 99 | 61 | . | . | . | . |
| PM1548-SK1Org2 | ADGB | p.Asp890Val | missense_variant | MODERATE | 0.148148 | 0 | 27 | 47 | . | . | . | . |
| PM1548-SK1Org2 | SYNE1 | p.Pro1307Ser | missense_variant | MODERATE | 0.497487 | 0 | 199 | 79 | . | . | . | . |
| PM1548-SK1Org2 | ARID1B | p.Pro1578Ser | missense_variant | MODERATE | 0 | 0 | 183 | 48 | true | . | . | . |
| PM1548-SK1Org2 | ARID1B | p.Ser1617Thr | missense_variant | MODERATE | 0 | 0 | 165 | 59 | true | . | . | . |
| PM1548-SK1Org2 | TULP4 | p.Pro1275Thr | missense_variant | MODERATE | 0.0187793 | 0 | 213 | 171 | . | . | . | . |
| PM1548-SK1Org2 | TULP4 | p.Asp1298Asn | missense_variant | MODERATE | 0.0626781 | 0.0078125 | 351 | 256 | . | . | . | . |
| PM1548-SK1Org2 | TULP4 | p.Asp1298Glu | missense_variant | MODERATE | 0.0968661 | 0 | 351 | 255 | . | . | . | . |
| PM1548-SK1Org2 | SMOC2 | p.Ala54Thr | missense_variant | MODERATE | 0.462366 | 0 | 93 | 36 | . | . | . | . |
| PM1548-SK1Org2 | GPR146 | p.Pro248Gln | missense_variant | MODERATE | 0 | 0 | 35 | 46 | . | . | . | . |
| PM1548-SK1Org2 | ELFN1 | p.Arg389Cys | missense_variant | MODERATE | 0.451613 | 0 | 93 | 43 | . | . | . | . |
| PM1548-SK1Org2 | SDK1 | p.Pro1727fs | frameshift_variant | HIGH | 0.489583 | 0 | 192 | 269 | . | . | . | . |
| PM1548-SK1Org2 | PMS2 | p.Thr277Ala | missense_variant | MODERATE | 0.140351 | 0 | 57 | 38 | true | . | . | . |
| PM1548-SK1Org2 | MIOS | p.Leu213His | missense_variant | MODERATE | 0.431818 | 0 | 88 | 44 | . | . | . | . |
| PM1548-SK1Org2 | ETV1 | p.Asn37Thr | missense_variant | MODERATE | 0.140845 | 0 | 71 | 72 | true | . | . | . |
| PM1548-SK1Org2 | MPP6 | p.Lys306fs | frameshift_variant | HIGH | 0 | 0 | 77 | 33 | . | . | . | . |
| PM1548-SK1Org2 | HERPUD2 | p.Val266Ala | missense_variant | MODERATE | 0.306667 | 0 | 225 | 59 | . | . | . | . |
| PM1548-SK1Org2 | POU6F2 | p.Glu562Ala | missense_variant | MODERATE | 0.256757 | 0 | 222 | 225 | . | . | . | . |
| PM1548-SK1Org2 | MYO1G | p.Ala779Val | missense_variant | MODERATE | 0.152047 | 0 | 171 | 358 | . | . | . | . |
| PM1548-SK1Org2 | VWC2 | p.Ser3Arg | missense_variant | MODERATE | 0.117647 | 0 | 17 | 64 | . | . | . | . |
| PM1548-SK1Org2 | GRB10 | p.Ala349Thr | missense_variant | MODERATE | 0.455479 | 0 | 292 | 372 | . | . | . | . |
| PM1548-SK1Org2 | WBSCR22 | p.Pro8Gln | missense_variant | MODERATE | 0 | 0 | 28 | 35 | . | . | . | . |
| PM1548-SK1Org2 | CLIP2 | p.Arg771Gln | missense_variant | MODERATE | 0.311111 | 0 | 180 | 150 | . | . | . | . |
| PM1548-SK1Org2 | TMEM60 | p.Ala78fs | frameshift_variant | HIGH | 0.483871 | 0 | 186 | 31 | . | . | . | . |
| PM1548-SK1Org2 | GNAI1 | p.Cys214Phe | missense_variant | MODERATE | 0 | 0 | 160 | 196 | . | . | . | . |
| PM1548-SK1Org2 | HGF | p.Val529Ala | missense_variant | MODERATE | 0 | 0 | 237 | 125 | . | true | . | . |
| PM1548-SK1Org2 | ABCB1 | p.Tyr1165* | stop_gained | HIGH | 0.46875 | 0 | 64 | 32 | . | . | . | . |
| PM1548-SK1Org2 | ABCB1 | c.3490-1G>C | splice_acceptor_variant&intron_variant | HIGH | 0.482759 | 0 | 58 | 32 | . | . | . | . |
| PM1548-SK1Org2 | ABCB1 | p.Ser196Ala | missense_variant | MODERATE | 0 | 0 | 65 | 56 | . | . | . | . |
| PM1548-SK1Org2 | MTERF | p.Ala221Thr | missense_variant | MODERATE | 0.47619 | 0 | 84 | 211 | . | . | . | . |
| PM1548-SK1Org2 | AKAP9 | p.Ile1377Thr | missense_variant | MODERATE | 0.490066 | 0 | 151 | 76 | true | . | . | . |
| PM1548-SK1Org2 | ASB4 | p.Thr229Met | missense_variant | MODERATE | 0.558333 | 0 | 120 | 168 | . | . | . | . |
| PM1548-SK1Org2 | LMTK2 | p.Asp236fs | frameshift_variant | HIGH | 0.430962 | 0 | 239 | 299 | . | . | . | . |
| PM1548-SK1Org2 | LMTK2 | p.Leu1130Ser | missense_variant | MODERATE | 0.487179 | 0 | 234 | 72 | . | . | . | . |
| PM1548-SK1Org2 | TRRAP | p.Asn1675Thr | missense_variant | MODERATE | 0.0314961 | 0 | 127 | 117 | true | . | . | . |
| PM1548-SK1Org2 | TRRAP | p.Arg1978Trp | missense_variant | MODERATE | 0.180328 | 0 | 183 | 221 | true | . | . | . |
| PM1548-SK1Org2 | TRRAP | p.Pro2508Ala | missense_variant | MODERATE | 0.045045 | 0 | 222 | 385 | true | . | . | . |
| PM1548-SK1Org2 | GIGYF1 | p.Leu580Pro | missense_variant | MODERATE | 0.0625 | 0 | 128 | 69 | . | . | . | . |
| PM1548-SK1Org2 | SLC12A9 | p.Leu559fs | frameshift_variant | HIGH | 0.551724 | 0 | 58 | 56 | . | . | . | . |
| PM1548-SK1Org2 | MOGAT3 | p.Asn82Lys | missense_variant | MODERATE | 0 | 0 | 244 | 150 | . | . | . | . |
| PM1548-SK1Org2 | SH2B2 | p.Ala74Ala | synonymous_variant | LOW | 0.571429 | 0 | 42 | 70 | . | . | . | . |
| PM1548-SK1Org2 | LRWD1 | p.Leu87Phe | missense_variant | MODERATE | 0 | 0 | 95 | 78 | . | . | . | . |
| PM1548-SK1Org2 | FBXL13 | p.Cys375Ser | missense_variant | MODERATE | 0 | 0 | 110 | 94 | . | . | . | . |
| PM1548-SK1Org2 | RELN | p.Arg2363His | missense_variant | MODERATE | 0.455814 | 0 | 215 | 39 | . | . | . | . |
| PM1548-SK1Org2 | ATXN7L1 | p.Val629Phe | missense_variant | MODERATE | 0 | 0 | 39 | 193 | . | . | . | . |
| PM1548-SK1Org2 | PIK3CG | p.Thr607fs | frameshift_variant | HIGH | 0 | 0 | 98 | 44 | . | true | . | . |
| PM1548-SK1Org2 | CBLL1 | p.Val277Ala | missense_variant | MODERATE | 0.00645161 | 0 | 155 | 200 | . | . | . | . |
| PM1548-SK1Org2 | SLC26A3 | p.Thr479Ala | missense_variant | MODERATE | 0.0336134 | 0 | 119 | 195 | . | . | true | . |
| PM1548-SK1Org2 | FOXP2 | p.Gly98Val | missense_variant | MODERATE | 0 | 0 | 182 | 125 | . | . | . | . |
| PM1548-SK1Org2 | PTPRZ1 | p.Pro751Leu | missense_variant | MODERATE | 0.413333 | 0 | 75 | 64 | . | . | . | . |
| PM1548-SK1Org2 | SND1 | p.Ala331Pro | missense_variant | MODERATE | 0.00645161 | 0 | 310 | 236 | true | . | . | . |
| PM1548-SK1Org2 | CEP41 | p.Asp152Gly | missense_variant | MODERATE | 0.110465 | 0 | 172 | 235 | . | . | . | . |
| PM1548-SK1Org2 | TSGA13 | p.Lys151fs | frameshift_variant | HIGH | 0 | 0 | 128 | 195 | . | . | . | . |
| PM1548-SK1Org2 | TTC26 | p.Ala221Thr | missense_variant | MODERATE | 0.568345 | 0 | 139 | 82 | . | . | . | . |
| PM1548-SK1Org2 | MKRN1 | c.186-2435G>C | intron_variant | MODIFIER | 0.431034 | 0 | 58 | 136 | . | . | . | . |
| PM1548-SK1Org2 | MGAM | p.Asn1603Asp | missense_variant | MODERATE | 0.511278 | 0 | 133 | 85 | . | . | . | . |
| PM1548-SK1Org2 | EPHB6 | p.Ala253fs | frameshift_variant | HIGH | 0.39834 | 0 | 241 | 105 | . | true | . | . |
| PM1548-SK1Org2 | ZNF775 | p.Gly65fs | frameshift_variant | HIGH | 0 | 0 | 114 | 50 | . | . | . | . |
| PM1548-SK1Org2 | FASTK | p.Ile345Asn | missense_variant | MODERATE | 0.214286 | 0 | 294 | 260 | . | . | . | . |
| PM1548-SK1Org2 | GALNT11 | p.Leu188Pro | missense_variant | MODERATE | 0.403846 | 0 | 156 | 157 | . | . | . | . |
| PM1548-SK1Org2 | TNKS | p.His859Asn | missense_variant | MODERATE | 0 | 0 | 34 | 75 | . | . | . | . |
| PM1548-SK1Org2 | RP1L1 | p.Gln1044fs | frameshift_variant | HIGH | 1 | 0 | 24 | 101 | . | . | . | . |
| PM1548-SK1Org2 | XKR6 | p.Arg80Lys | missense_variant | MODERATE | 0.269231 | 0 | 104 | 127 | . | . | . | . |
| PM1548-SK1Org2 | PCM1 | p.Met1321Val | missense_variant | MODERATE | 0.186047 | 0 | 43 | 137 | true | . | . | . |
| PM1548-SK1Org2 | SORBS3 | p.Thr100Ala | missense_variant | MODERATE | 0 | 0 | 136 | 140 | . | . | . | . |
| PM1548-SK1Org2 | LOXL2 | p.Pro40Leu | missense_variant | MODERATE | 1 | 0 | 44 | 224 | . | . | . | . |
| PM1548-SK1Org2 | NEFM | p.Leu5Leu | synonymous_variant | LOW | 0.12 | 0 | 25 | 41 | . | . | . | . |
| PM1548-SK1Org2 | NRG1 | p.Gly39Glu | missense_variant | MODERATE | 0.121951 | 0 | 82 | 55 | true | . | . | . |
| PM1548-SK1Org2 | NRG1 | p.Pro44Ser | missense_variant | MODERATE | 0.112903 | 0 | 62 | 58 | true | . | . | . |
| PM1548-SK1Org2 | NRG1 | p.Ala74Pro | missense_variant | MODERATE | 0.147059 | 0 | 34 | 35 | true | . | . | . |
| PM1548-SK1Org2 | NRG1 | c.1488-469C>G | intron_variant | MODIFIER | 0.0327869 | 0 | 61 | 107 | true | . | . | . |
| PM1548-SK1Org2 | ADAM2 | p.Arg3His | missense_variant | MODERATE | 0 | 0 | 57 | 64 | . | . | . | . |
| PM1548-SK1Org2 | DKK4 | p.Asp195Asn | missense_variant | MODERATE | 0.46281 | 0 | 121 | 81 | . | . | . | . |
| PM1548-SK1Org2 | MCM4 | p.Thr629fs | frameshift_variant | HIGH | 0.481081 | 0 | 185 | 85 | . | . | . | . |
| PM1548-SK1Org2 | PXDNL | p.Ala353Thr | missense_variant | MODERATE | 0.553191 | 0 | 188 | 174 | . | . | . | . |
| PM1548-SK1Org2 | RP1 | p.Tyr834fs | frameshift_variant | HIGH | 0.507937 | 0 | 63 | 39 | . | . | . | . |
| PM1548-SK1Org2 | YTHDF3 | p.Gln323Gln | synonymous_variant | LOW | 0.027248 | 0 | 367 | 589 | . | . | . | . |
| PM1548-SK1Org2 | C8orf46 | p.Thr78Ile | missense_variant | MODERATE | 0 | 0 | 40 | 109 | . | . | . | . |
| PM1548-SK1Org2 | ZFHX4 | p.Ala2905Val | missense_variant | MODERATE | 0.421488 | 0 | 121 | 42 | . | . | . | . |
| PM1548-SK1Org2 | HEY1 | p.Val187Ala | missense_variant | MODERATE | 0.0505618 | 0 | 178 | 281 | true | . | . | . |
| PM1548-SK1Org2 | HEY1 | p.Thr186Ser | missense_variant | MODERATE | 0.0576923 | 0 | 156 | 272 | true | . | . | . |

| | | | | | | | | | | | | |
|---|---|---|---|---|---|---|---|---|---|---|---|---|
| PM1548-SK1Org2 | VPS13B | p.Asn2064Thr | missense_variant | MODERATE | 0.229008 | 0 | 131 | 75 | . | . | . | . |
| PM1548-SK1Org2 | TRPS1 | p.Arg297Cys | missense_variant | MODERATE | 0.488095 | 0 | 168 | 259 | . | . | . | . |
| PM1548-SK1Org2 | TRPS1 | p.Asn265Asn | synonymous_variant | LOW | 0.0408163 | 0 | 147 | 334 | . | . | . | . |
| PM1548-SK1Org2 | EXT1 | p.Val66Phe | missense_variant | MODERATE | 0.0397727 | 0 | 176 | 88 | true | . | . | . |
| PM1548-SK1Org2 | TBC1D31 | p.Ala208Thr | missense_variant | MODERATE | 0.413636 | 0 | 220 | 184 | . | . | . | . |
| PM1548-SK1Org2 | ATAD2 | p.Glu119fs | frameshift_variant | HIGH | 0.34375 | 0 | 64 | 107 | . | . | . | . |
| PM1548-SK1Org2 | ZNF572 | p.Ser55Pro | missense_variant | MODERATE | 0.601307 | 0 | 153 | 39 | . | . | . | . |
| PM1548-SK1Org2 | MYC | p.Gly175Ser | missense_variant | MODERATE | 0.021978 | 0 | 91 | 270 | true | . | . | . |
| PM1548-SK1Org2 | ASAP1 | p.Glu919Lys | missense_variant | MODERATE | 0.378571 | 0 | 140 | 184 | . | . | . | . |
| PM1548-SK1Org2 | WISP1 | p.Ile175Met | missense_variant | MODERATE | 0.25641 | 0 | 195 | 119 | . | . | . | . |
| PM1548-SK1Org2 | KHDRBS3 | p.Tyr316Tyr | splice_region_variant&synonymous_variant | LOW | 0.504348 | 0 | 115 | 229 | . | . | . | . |
| PM1548-SK1Org2 | COL22A1 | p.Thr125Met | missense_variant | MODERATE | 0.508197 | 0 | 61 | 58 | . | . | . | . |
| PM1548-SK1Org2 | LY6D | p.Ala99Thr | missense_variant | MODERATE | 0.168367 | 0.00956938 | 196 | 209 | . | . | . | . |
| PM1548-SK1Org2 | TOP1MT | p.Pro193Leu | missense_variant | MODERATE | 0 | 0.00595238 | 60 | 168 | . | . | . | . |
| PM1548-SK1Org2 | ZC3H3 | p.Gly101fs | frameshift_variant | HIGH | 0.27957 | 0 | 93 | 100 | . | . | . | . |
| PM1548-SK1Org2 | GSDMD | p.Glu451Ala | missense_variant | MODERATE | 0.473282 | 0 | 131 | 215 | . | . | . | . |
| PM1548-SK1Org2 | MAPK15 | p.Pro493Leu | missense_variant | MODERATE | 0.478764 | 0 | 259 | 366 | . | . | . | . |
| PM1548-SK1Org2 | SCRIB | p.Cys22Trp | missense_variant | MODERATE | 0.338235 | 0 | 68 | 81 | . | . | . | . |
| PM1548-SK1Org2 | PLEC | p.Arg3540Gln | missense_variant | MODERATE | 0 | 0 | 44 | 130 | . | . | . | . |
| PM1548-SK1Org2 | PLEC | p.Val648Met | missense_variant | MODERATE | 0 | 0 | 49 | 166 | . | . | . | . |
| PM1548-SK1Org2 | CYHR1 | n.145689659du | intragenic_variant | MODIFIER | 0.152 | 0 | 125 | 56 | . | . | . | . |
| PM1548-SK1Org2 | ZNF7 | p.Val136Ile | missense_variant | MODERATE | 0 | 0 | 44 | 154 | . | . | . | . |
| PM1548-SK1Org2 | KANK1 | p.Thr192Ser | missense_variant | MODERATE | 0.00632911 | 0 | 158 | 65 | . | . | . | . |
| PM1548-SK1Org2 | DMRT3 | p.Leu385Leu | synonymous_variant | LOW | 0.0280374 | 0 | 214 | 221 | . | . | . | . |
| PM1548-SK1Org2 | RLN2 | p.His8fs | frameshift_variant | HIGH | 0.29646 | 0 | 226 | 225 | . | . | . | . |
| PM1548-SK1Org2 | IFNA4 | p.Ala74Thr | missense_variant | MODERATE | 0.285714 | 0 | 42 | 51 | . | . | . | . |
| PM1548-SK1Org2 | TAF1L | p.Gly4Ser | missense_variant | MODERATE | 0.311688 | 0 | 77 | 329 | . | . | . | true |
| PM1548-SK1Org2 | BAG1 | p.Glu321Lys | missense_variant | MODERATE | 0.463768 | 0 | 207 | 192 | . | . | . | . |
| PM1548-SK1Org2 | PRSS3 | p.Ser102Tyr | missense_variant | MODERATE | 0.125 | 0 | 232 | 102 | . | . | . | . |
| PM1548-SK1Org2 | CD72 | p.Ala111Thr | missense_variant | MODERATE | 0.505338 | 0 | 281 | 227 | . | . | . | . |
| PM1548-SK1Org2 | RGP1 | p.Arg328Trp | missense_variant | MODERATE | 0.506098 | 0 | 164 | 168 | . | . | . | . |
| PM1548-SK1Org2 | ANKRD18A | p.Gly900Ala | missense_variant | MODERATE | 0.421875 | 0 | 64 | 119 | . | . | . | . |
| PM1548-SK1Org2 | ANKRD18A | p.Ala897Val | missense_variant | MODERATE | 0.421875 | 0 | 64 | 119 | . | . | . | . |
| PM1548-SK1Org2 | CNTNAP3 | p.Arg841Ser | missense_variant | MODERATE | 0 | 0 | 45 | 45 | . | . | . | . |
| PM1548-SK1Org2 | TMC1 | p.Ala755Pro | missense_variant&splice_region_variant | MODERATE | 0.300813 | 0 | 123 | 45 | . | . | . | . |
| PM1548-SK1Org2 | RMI1 | p.Thr325Ala | missense_variant | MODERATE | 0.546392 | 0 | 291 | 482 | . | . | . | . |
| PM1548-SK1Org2 | NAA35 | p.*726* | stop_retained_variant | LOW | 0 | 0 | 202 | 360 | . | . | . | . |
| PM1548-SK1Org2 | SPATA31E1 | p.Arg436His | missense_variant | MODERATE | 0.54 | 0 | 100 | 212 | . | . | . | . |
| PM1548-SK1Org2 | ROR2 | p.Ala307Thr | missense_variant | MODERATE | 0 | 0 | 231 | 69 | . | . | . | . |
| PM1548-SK1Org2 | NOL8 | p.Arg430Lys | missense_variant | MODERATE | 0.0434783 | 0 | 92 | 112 | . | . | . | . |
| PM1548-SK1Org2 | NOL8 | p.Leu427Gln | missense_variant | MODERATE | 0.0510204 | 0 | 98 | 113 | . | . | . | . |
| PM1548-SK1Org2 | IPPK | p.Phe34Ser | missense_variant | MODERATE | 0.543779 | 0 | 217 | 158 | . | . | . | . |
| PM1548-SK1Org2 | SUSD3 | c.89-6671G>T | intron_variant | MODIFIER | 0 | 0 | 43 | 26 | . | . | . | . |
| PM1548-SK1Org2 | WNK2 | p.His758fs | frameshift_variant | HIGH | 0.429688 | 0 | 128 | 277 | . | . | . | . |
| PM1548-SK1Org2 | C9orf3 | p.Ala207Thr | missense_variant | MODERATE | 0.25 | 0 | 164 | 105 | . | . | . | . |
| PM1548-SK1Org2 | PTCH1 | p.Thr1064Met | missense_variant | MODERATE | 0.283582 | 0 | 201 | 167 | true | . | . | . |
| PM1548-SK1Org2 | ZNF510 | p.Arg592Gly | missense_variant | MODERATE | 0 | 0 | 156 | 140 | . | . | . | . |
| PM1548-SK1Org2 | CCDC180 | p.Met439Ile | missense_variant | MODERATE | 0.511628 | 0 | 129 | 95 | . | . | . | . |
| PM1548-SK1Org2 | GABBR2 | p.Arg622* | stop_gained | HIGH | 0.565217 | 0 | 69 | 79 | . | . | . | . |
| PM1548-SK1Org2 | ANKS6 | p.Ala872Val | missense_variant | MODERATE | 0.258929 | 0 | 112 | 240 | . | . | . | . |
| PM1548-SK1Org2 | SLC44A1 | p.Arg437His | missense_variant | MODERATE | 0.433333 | 0 | 90 | 71 | . | . | . | . |
| PM1548-SK1Org2 | SVEP1 | p.Ile1787Thr | missense_variant | MODERATE | 0.433735 | 0 | 83 | 79 | . | . | . | . |
| PM1548-SK1Org2 | TNC | p.Lys1068Asn | missense_variant | MODERATE | 0.190476 | 0 | 105 | 37 | . | . | . | . |
| PM1548-SK1Org2 | BRINP1 | p.Arg106His | missense_variant | MODERATE | 0.429688 | 0 | 128 | 219 | . | . | . | . |
| PM1548-SK1Org2 | FBXW2 | p.Arg385His | missense_variant | MODERATE | 0.0347826 | 0 | 115 | 162 | . | . | . | . |
| PM1548-SK1Org2 | FBXW2 | p.Ile377Val | missense_variant | MODERATE | 0.0322581 | 0 | 124 | 161 | . | . | . | . |
| PM1548-SK1Org2 | DAB2IP | p.His1025fs | frameshift_variant | HIGH | 0.421569 | 0 | 102 | 32 | . | . | . | . |
| PM1548-SK1Org2 | LRSAM1 | p.Val385Ile | missense_variant | MODERATE | 0.467742 | 0 | 124 | 115 | . | . | . | . |
| PM1548-SK1Org2 | SH2D3C | p.Arg821His | missense_variant | MODERATE | 0.375 | 0 | 176 | 143 | . | . | . | . |
| PM1548-SK1Org2 | C9orf114 | p.Val77Ile | missense_variant | MODERATE | 0.447205 | 0 | 161 | 277 | . | . | . | . |
| PM1548-SK1Org2 | C9orf50 | p.Ala252Val | missense_variant | MODERATE | 0.444444 | 0 | 189 | 411 | . | . | . | . |
| PM1548-SK1Org2 | NUP214 | p.Arg887His | missense_variant | MODERATE | 0.537736 | 0 | 106 | 67 | true | . | . | . |
| PM1548-SK1Org2 | ADAMTS13 | p.Arg7fs | frameshift_variant | HIGH | 0.509653 | 0 | 259 | 247 | . | . | . | . |
| PM1548-SK1Org2 | SARDH | p.Ala819Val | missense_variant | MODERATE | 0.495935 | 0 | 123 | 41 | . | . | . | . |
| PM1548-SK1Org2 | SARDH | p.Arg280Trp | missense_variant | MODERATE | 0 | 0 | 143 | 43 | . | . | . | . |
| PM1548-SK1Org2 | LHX3 | p.Ala180Thr | missense_variant | MODERATE | 0.397727 | 0 | 88 | 204 | . | . | . | . |
| PM1548-SK1Org2 | MAMDC4 | p.Gln963His | missense_variant | MODERATE | 0.536082 | 0 | 97 | 91 | . | . | . | . |
| PM1548-SK1Org2 | AL807752.1 | p.Trp56fs | frameshift_variant | HIGH | 0.647059 | 0 | 34 | 54 | . | . | . | . |
| PM1548-SK1Org2 | TPRN | p.Asn499Asp | missense_variant | MODERATE | 0.113208 | 0 | 424 | 335 | . | . | . | . |
| PM1548-SK1Org2 | NSMF | p.Arg366Cys | missense_variant | MODERATE | 0.55303 | 0 | 132 | 90 | . | . | . | . |
| PM1548-SK1Org2 | PNPLA7 | p.Leu178Met | missense_variant | MODERATE | 0 | 0 | 174 | 199 | . | . | . | . |
| PM1548-SK1Org2 | HDHD1 | p.Arg149Cys | missense_variant | MODERATE | 1 | 0 | 60 | 104 | . | . | . | . |
| PM1548-SK1Org2 | EGFL6 | p.Trp5* | stop_gained | HIGH | 0 | 0 | 46 | 81 | . | . | . | . |
| PM1548-SK1Org2 | YY2 | p.Glu5Val | missense_variant | MODERATE | 0.484848 | 0 | 99 | 51 | . | . | . | . |
| PM1548-SK1Org2 | TMEM47 | p.Cys104Cys | synonymous_variant | LOW | 0.0238095 | 0 | 84 | 132 | . | . | . | . |
| PM1548-SK1Org2 | USP27X | p.Arg155His | missense_variant | MODERATE | 0 | 0 | 130 | 112 | . | . | . | . |
| PM1548-SK1Org2 | KDM5C | p.Glu1003Asp | missense_variant | MODERATE | 0 | 0 | 75 | 54 | true | true | true | . |
| PM1548-SK1Org2 | IQSEC2 | p.Arg612Cys | missense_variant | MODERATE | 0 | 0 | 73 | 42 | . | . | . | . |
| PM1548-SK1Org2 | FAM120C | p.Gln36_Gln37de | disruptive_inframe_deletion | MODERATE | 0.95082 | 0 | 61 | 126 | . | . | . | . |
| PM1548-SK1Org2 | MAGED2 | p.Glu53Asp | missense_variant | MODERATE | 0 | 0 | 50 | 102 | . | . | . | . |
| PM1548-SK1Org2 | TRO | p.Ala1132Thr | missense_variant | MODERATE | 0 | 0 | 35 | 69 | . | . | . | . |
| PM1548-SK1Org2 | DLG3 | p.Leu333Ile | missense_variant | MODERATE | 1 | 0 | 63 | 34 | . | . | . | . |
| PM1548-SK1Org2 | ARMCX4 | p.Tyr2123fs | frameshift_variant | HIGH | 0 | 0 | 124 | 78 | . | . | . | . |
| PM1548-SK1Org2 | PAK3 | p.Ala466Val | missense_variant | MODERATE | 0.944444 | 0 | 18 | 37 | . | . | . | . |
| PM1548-SK1Org2 | ZCCHC16 | p.Arg297Cys | missense_variant | MODERATE | 0.495726 | 0 | 117 | 98 | . | . | . | . |
| PM1548-SK1Org2 | AVPR2 | p.Ala60Val | missense_variant | MODERATE | 0 | 0 | 41 | 59 | . | . | . | . |
| PM1548-SK1P2 | B3GALT6 | p.Glu303Lys | missense_variant | MODERATE | 0.432735 | 0 | 446 | 86 | . | . | . | . |
| PM1548-SK1P2 | SCNN1D | p.Arg344His | missense_variant | MODERATE | 0.459627 | 0 | 322 | 274 | . | . | . | . |
| PM1548-SK1P2 | ATAD3B | p.Arg290Cys | missense_variant | MODERATE | 0.463576 | 0.00687285 | 906 | 291 | . | . | . | . |
| PM1548-SK1P2 | C1orf86 | p.Pro72His | missense_variant | MODERATE | 0.408669 | 0 | 323 | 43 | . | . | . | . |
| PM1548-SK1P2 | PRDM16 | p.Arg759Gln | missense_variant | MODERATE | 0 | 0 | 12 | 40 | true | . | . | . |
| PM1548-SK1P2 | DFFB | p.Tyr290Phe | missense_variant | MODERATE | 0.0357143 | 0 | 28 | 203 | . | . | . | . |
| PM1548-SK1P2 | NPHP4 | p.Arg488Gln | missense_variant | MODERATE | 0.533686 | 0 | 757 | 194 | . | . | . | . |
| PM1548-SK1P2 | PLEKHG5 | p.Thr301fs | frameshift_variant | HIGH | 0.505628 | 0 | 1155 | 106 | . | . | . | . |
| PM1548-SK1P2 | ZBTB48 | p.Val626Ala | missense_variant | MODERATE | 0.0170455 | 0 | 528 | 143 | . | . | . | . |
| PM1548-SK1P2 | KLHL21 | p.Cys250fs | frameshift_variant | HIGH | 0.458217 | 0 | 718 | 134 | . | . | . | . |
| PM1548-SK1P2 | CAMTA1 | p.Ser510Thr | missense_variant | MODERATE | 0.0141844 | 0 | 141 | 185 | true | . | . | . |
| PM1548-SK1P2 | PER3 | p.Ala803Ser | missense_variant | MODERATE | 0.139535 | 0.00900901 | 43 | 111 | . | . | . | . |
| PM1548-SK1P2 | SLC45A1 | p.Thr364Met | missense_variant | MODERATE | 0.571429 | 0 | 7 | 56 | . | . | . | . |
| PM1548-SK1P2 | SLC45A1 | p.Tyr689* | stop_gained | HIGH | 0.777778 | 0 | 9 | 120 | . | . | . | . |
| PM1548-SK1P2 | RERE | p.Lys83fs | frameshift_variant | HIGH | 0 | 0 | 2 | 49 | . | . | . | . |
| PM1548-SK1P2 | CORT | p.Gln133fs | frameshift_variant | HIGH | 0.0121951 | 0 | 82 | 48 | . | . | . | . |
| PM1548-SK1P2 | EXOSC10 | p.Met1fs | frameshift_variant&start_lost | HIGH | 0.506977 | 0 | 215 | 61 | . | . | . | . |
| PM1548-SK1P2 | FHAD1 | p.Ile953fs | frameshift_variant | HIGH | 0.510345 | 0 | 145 | 108 | . | . | . | . |
| PM1548-SK1P2 | PLEKHM2 | p.Arg848Trp | missense_variant | MODERATE | 0 | 0 | 38 | 42 | . | . | . | . |
| PM1548-SK1P2 | SPEN | p.Asn2002fs | frameshift_variant | HIGH | 0 | 0 | 843 | 173 | true | . | true | true |
| PM1548-SK1P2 | EPHA2 | p.Asn106Ser | missense_variant | MODERATE | 0 | 0 | 33 | 84 | . | . | true | . |
| PM1548-SK1P2 | CROCC | p.Arg1590Cys | missense_variant | MODERATE | 0.332203 | 0 | 295 | 547 | . | . | . | . |
| PM1548-SK1P2 | RCC2 | p.Gly465Arg | missense_variant | MODERATE | 0.727273 | 0 | 44 | 148 | . | . | . | . |
| PM1548-SK1P2 | ARHGEF10L | p.Val965Met | missense_variant | MODERATE | 0.00084674 | 0 | 1181 | 230 | . | . | . | . |
| PM1548-SK1P2 | KLHDC7A | p.Ala652Val | missense_variant | MODERATE | 0.473684 | 0 | 19 | 60 | . | . | . | . |
| PM1548-SK1P2 | RNF186 | p.Ile185Thr | missense_variant | MODERATE | 0.00306748 | 0 | 326 | 472 | . | . | . | . |
| PM1548-SK1P2 | KIF17 | p.Arg196His | missense_variant | MODERATE | 0.386364 | 0 | 44 | 71 | . | . | . | . |
| PM1548-SK1P2 | EIF4G3 | c.30+5101C>G | intron_variant | MODIFIER | 0.12 | 0 | 25 | 302 | . | . | . | . |
| PM1548-SK1P2 | HSPG2 | p.Gly2259Asp | missense_variant | MODERATE | 0 | 0 | 11 | 55 | . | . | . | . |
| PM1548-SK1P2 | EPHA8 | p.Ser87Asn | missense_variant | MODERATE | 0.114286 | 0 | 35 | 189 | . | . | . | . |
| PM1548-SK1P2 | AIM1L | n.26671971delA | intragenic_variant | MODIFIER | 0 | 0 | 9 | 55 | . | . | . | . |
| PM1548-SK1P2 | ARID1A | p.Thr1863Ile | missense_variant | MODERATE | 0.021544 | 0 | 557 | 201 | true | true | true | true |
| PM1548-SK1P2 | PTPRU | p.Arg639Trp | missense_variant | MODERATE | 0 | 0 | 385 | 60 | . | . | . | . |
| PM1548-SK1P2 | FABP3 | p.Thr54fs | frameshift_variant | HIGH | 0.363636 | 0 | 33 | 207 | . | . | . | . |
| PM1548-SK1P2 | LCK | p.Arg222Cys | missense_variant | MODERATE | 0 | 0 | 60 | 146 | true | . | . | . |
| PM1548-SK1P2 | KIAA1522 | p.Ala943Thr | missense_variant | MODERATE | 0.517483 | 0 | 143 | 198 | . | . | . | . |
| PM1548-SK1P2 | ZNF362 | p.Ala105Val | missense_variant | MODERATE | 0 | 0 | 118 | 295 | . | . | . | . |
| PM1548-SK1P2 | LSM10 | p.Ala2Val | missense_variant | MODERATE | 0.518519 | 0 | 54 | 400 | . | . | . | . |
| PM1548-SK1P2 | CSF3R | p.Tyr508Cys | missense_variant | MODERATE | 0 | 0 | 506 | 167 | true | . | . | . |
| PM1548-SK1P2 | EPHA10 | p.Pro330Ser | missense_variant | MODERATE | 0.420382 | 0 | 157 | 71 | . | . | . | . |
| PM1548-SK1P2 | PABPC4 | p.Ser206Asn | missense_variant | MODERATE | 0.171429 | 0 | 35 | 90 | . | . | . | . |

| | | | | | | | | | | | | |
|---|---|---|---|---|---|---|---|---|---|---|---|---|
| PM1548-SK1P2 | PABPC4 | p.Glu205Gly | missense_variant | MODERATE | 0.2 | 0 | 30 | 73 | . | . | . | . |
| PM1548-SK1P2 | ZMPSTE24 | p.Leu362fs | frameshift_variant | HIGH | 0.5 | 0 | 28 | 53 | . | . | . | . |
| PM1548-SK1P2 | HIVEP3 | p.Pro1037Leu | missense_variant | MODERATE | 0.125 | 0 | 16 | 35 | . | . | . | . |
| PM1548-SK1P2 | MED8 | p.Asn295fs | frameshift_variant | HIGH | 0.615741 | 0 | 216 | 97 | . | . | . | . |
| PM1548-SK1P2 | HYI | p.Ala55Ser | missense_variant | MODERATE | 0.000809717 | 0 | 1235 | 158 | . | . | . | . |
| PM1548-SK1P2 | PTPRF | p.Arg1669Trp | missense_variant | MODERATE | 0.517815 | 0 | 421 | 112 | . | . | . | . |
| PM1548-SK1P2 | SLC6A9 | p.Arg662Gln | missense_variant | MODERATE | 0.496183 | 0 | 262 | 99 | . | . | . | . |
| PM1548-SK1P2 | PTCH2 | p.Ala967Val | missense_variant | MODERATE | 0 | 0 | 183 | 194 | . | . | . | . |
| PM1548-SK1P2 | PRDX1 | p.His10Tyr | missense_variant | MODERATE | 0 | 0 | 31 | 151 | . | . | . | . |
| PM1548-SK1P2 | MAST2 | p.Arg1086Arg | synonymous_variant | LOW | 0.0540541 | 0 | 74 | 187 | . | . | . | . |
| PM1548-SK1P2 | TAL1 | p.His86Leu | missense_variant | MODERATE | 0 | 0 | 221 | 170 | true | . | . | . |
| PM1548-SK1P2 | LRP8 | p.Asp50Gly | missense_variant | MODERATE | 0.391727 | 0 | 411 | 175 | . | . | . | . |
| PM1548-SK1P2 | CDCP2 | p.Pro209fs | frameshift_variant | HIGH | 0.2 | 0 | 10 | 107 | . | . | . | . |
| PM1548-SK1P2 | MRPL37 | p.Asp142Glu | missense_variant | MODERATE | 0.169014 | 0 | 71 | 90 | . | . | . | . |
| PM1548-SK1P2 | CACHD1 | p.Glu211Lys | missense_variant | MODERATE | 0.133333 | 0 | 30 | 147 | . | . | . | . |
| PM1548-SK1P2 | JAK1 | p.Val433Ile | missense_variant | MODERATE | 0.292398 | 0 | 171 | 203 | true | . | . | true |
| PM1548-SK1P2 | FPGT | p.Asp273Asn | missense_variant | MODERATE | 0 | 0 | 3 | 168 | . | . | . | . |
| PM1548-SK1P2 | LPHN2 | p.Asn759Ile | missense_variant | MODERATE | 1 | 0 | 2 | 85 | . | . | . | . |
| PM1548-SK1P2 | SYDE2 | p.Gly14fs | frameshift_variant | HIGH | 0 | 0 | 281 | 141 | . | . | . | . |
| PM1548-SK1P2 | CLCA1 | p.Arg676Trp | missense_variant | MODERATE | 0.166667 | 0 | 6 | 91 | . | . | . | . |
| PM1548-SK1P2 | GBP6 | p.Gly77Asp | missense_variant | MODERATE | 0.25 | 0 | 4 | 98 | . | . | . | . |
| PM1548-SK1P2 | GFI1 | p.Arg348Met | missense_variant | MODERATE | 0.47033 | 0 | 455 | 270 | . | . | . | . |
| PM1548-SK1P2 | ABCA4 | p.His423Gln | missense_variant | MODERATE | 0 | 0 | 46 | 257 | . | . | . | . |
| PM1548-SK1P2 | ABCA4 | p.Glu422Asp | missense_variant | MODERATE | 0 | 0.00389105 | 46 | 257 | . | . | . | . |
| PM1548-SK1P2 | LPPR4 | p.Ala343Thr | missense_variant | MODERATE | 0.448276 | 0 | 29 | 97 | . | . | . | . |
| PM1548-SK1P2 | HIAT1 | p.Ala235Val | missense_variant&splice_region_variant | MODERATE | 0.676471 | 0 | 68 | 80 | . | . | . | . |
| PM1548-SK1P2 | TRMT13 | p.Glu307fs | frameshift_variant | HIGH | 0 | 0 | 25 | 85 | . | . | . | . |
| PM1548-SK1P2 | AKNAD1 | p.Leu577Met | missense_variant | MODERATE | 0.35 | 0 | 20 | 459 | . | . | . | . |
| PM1548-SK1P2 | CELSR2 | p.Asp1795Gly | missense_variant | MODERATE | 0.125 | 0 | 32 | 362 | . | . | . | . |
| PM1548-SK1P2 | LAMTOR5 | p.Thr39Ala | missense_variant | MODERATE | 0 | 0 | 199 | 96 | . | . | . | . |
| PM1548-SK1P2 | HIPK1 | p.Ala314Thr | missense_variant | MODERATE | 0.666667 | 0 | 6 | 273 | . | . | . | . |
| PM1548-SK1P2 | TRIM33 | p.Val961Met | missense_variant | MODERATE | 0.333333 | 0 | 6 | 37 | true | . | . | . |
| PM1548-SK1P2 | PDE4DIP | p.Gln2125Lys | missense_variant | MODERATE | 0.3 | 0 | 20 | 86 | true | . | . | . |
| PM1548-SK1P2 | PDE4DIP | p.Arg759* | stop_gained | HIGH | 0.287234 | 0 | 94 | 104 | true | . | . | . |
| PM1548-SK1P2 | BCL9 | p.Pro517fs | frameshift_variant | HIGH | 0.424658 | 0 | 219 | 142 | true | . | . | . |
| PM1548-SK1P2 | TARS2 | p.Tyr371* | stop_gained | HIGH | 0 | 0 | 40 | 48 | . | . | . | . |
| PM1548-SK1P2 | ADAMTSL4 | p.Gly780fs | frameshift_variant | HIGH | 0.549045 | 0 | 785 | 110 | . | . | . | . |
| PM1548-SK1P2 | ZNF687 | p.Arg885His | missense_variant | MODERATE | 0.49422 | 0 | 692 | 322 | . | . | . | . |
| PM1548-SK1P2 | PI4KB | p.Val202Met | missense_variant | MODERATE | 0 | 0 | 6 | 157 | . | . | . | . |
| PM1548-SK1P2 | CELF3 | p.Val279Ile | missense_variant | MODERATE | 0.423077 | 0 | 26 | 54 | . | . | . | . |
| PM1548-SK1P2 | DENND4B | p.Ala154fs | frameshift_variant | HIGH | 0.342105 | 0 | 456 | 106 | . | . | . | . |
| PM1548-SK1P2 | CREB3L4 | p.Val263fs | frameshift_variant | HIGH | 0.430108 | 0 | 93 | 42 | . | . | . | . |
| PM1548-SK1P2 | EFNA4 | c.470-101delG | intron_variant | MODIFIER | 0 | 0 | 9 | 82 | . | . | . | . |
| PM1548-SK1P2 | ASH1L | p.Asn1907Lys | missense_variant | MODERATE | 0 | 0 | 59 | 55 | . | . | . | . |
| PM1548-SK1P2 | SMG5 | p.Gln846Lys | missense_variant | MODERATE | 0 | 0 | 73 | 128 | . | . | . | . |
| PM1548-SK1P2 | CD1D | p.Asp140Asn | missense_variant | MODERATE | 0.625 | 0 | 16 | 198 | . | . | true | . |
| PM1548-SK1P2 | AIM2 | p.Lys342fs | frameshift_variant | HIGH | 0.6 | 0 | 10 | 33 | . | . | . | . |
| PM1548-SK1P2 | NIT1 | p.Arg261His | missense_variant | MODERATE | 0.292434 | 0 | 489 | 402 | . | . | . | . |
| PM1548-SK1P2 | ILDR2 | p.Gly453Ala | missense_variant | MODERATE | 0 | 0 | 134 | 328 | . | . | . | . |
| PM1548-SK1P2 | ILDR2 | p.Arg415Trp | missense_variant | MODERATE | 0.449713 | 0 | 1392 | 213 | . | . | . | . |
| PM1548-SK1P2 | MYOC | p.Arg91Gln | missense_variant | MODERATE | 0 | 0 | 526 | 242 | . | . | . | . |
| PM1548-SK1P2 | DNM3 | p.Glu7Asp | missense_variant | MODERATE | 0 | 0 | 50 | 77 | . | . | . | . |
| PM1548-SK1P2 | DNM3 | p.Phe20Phe | synonymous_variant | LOW | 0.0520833 | 0.0060241 | 96 | 166 | . | . | . | . |
| PM1548-SK1P2 | DNM3 | p.His85Gln | missense_variant | MODERATE | 0 | 0 | 44 | 56 | . | . | . | . |
| PM1548-SK1P2 | ANKRD45 | p.Leu39fs | frameshift_variant | HIGH | 0.373626 | 0 | 91 | 229 | . | . | . | . |
| PM1548-SK1P2 | ZBTB37 | p.Val284Ile | missense_variant | MODERATE | 0.0172414 | 0 | 58 | 152 | . | . | . | . |
| PM1548-SK1P2 | ZBTB37 | p.Asn288Ser | missense_variant | MODERATE | 0.047619 | 0 | 42 | 160 | . | . | . | . |
| PM1548-SK1P2 | PAPPA2 | p.Thr1174fs | frameshift_variant | HIGH | 0.666667 | 0 | 3 | 36 | . | . | . | . |
| PM1548-SK1P2 | CACNA1E | p.Asn2301Ser | missense_variant | MODERATE | 0 | 0 | 49 | 120 | . | . | . | . |
| PM1548-SK1P2 | CACNA1E | p.Asn2302Asp | missense_variant | MODERATE | 0 | 0 | 49 | 119 | . | . | . | . |
| PM1548-SK1P2 | CACNA1E | p.Asp2310Asp | synonymous_variant | LOW | 0 | 0 | 48 | 90 | . | . | . | . |
| PM1548-SK1P2 | ZNF648 | p.Ser436Tyr | missense_variant | MODERATE | 0.111111 | 0 | 54 | 413 | . | . | . | . |
| PM1548-SK1P2 | ZNF648 | p.Ser436Pro | missense_variant | MODERATE | 0.103448 | 0 | 58 | 439 | . | . | . | . |
| PM1548-SK1P2 | ZNF648 | p.Pro423Ser | missense_variant | MODERATE | 0.103448 | 0 | 58 | 436 | . | . | . | . |
| PM1548-SK1P2 | C1orf27 | p.Thr381Ser | missense_variant | MODERATE | 0 | 0 | 2 | 10 | . | . | . | . |
| PM1548-SK1P2 | PLA2G4A | c.1033+2T>C | splice_donor_variant&intron_variant | HIGH | 0.7 | 0 | 10 | 80 | . | . | . | . |
| PM1548-SK1P2 | ASPM | p.Arg2700Gln | missense_variant | MODERATE | 0 | 0.00653595 | 60 | 153 | . | . | . | . |
| PM1548-SK1P2 | DENND1B | c.83-20669G>A | intron_variant | MODIFIER | 0.466667 | 0 | 15 | 39 | . | . | . | . |
| PM1548-SK1P2 | NR5A2 | p.Gly169Val | missense_variant | MODERATE | 0.4 | 0 | 15 | 71 | . | . | . | . |
| PM1548-SK1P2 | NAV1 | p.Thr719Ser | missense_variant | MODERATE | 0.0162602 | 0 | 246 | 176 | . | . | . | . |
| PM1548-SK1P2 | NAV1 | p.Thr726Ser | missense_variant | MODERATE | 0.0160643 | 0.00930233 | 249 | 215 | . | . | . | . |
| PM1548-SK1P2 | ATP2B4 | p.Asp867Asn | missense_variant&splice_region_variant | MODERATE | 0.1 | 0 | 30 | 121 | . | . | . | . |
| PM1548-SK1P2 | NUAK2 | p.Arg378His | missense_variant | MODERATE | 0.262097 | 0 | 248 | 162 | . | . | . | . |
| PM1548-SK1P2 | PFKFB2 | p.Glu261Ala | missense_variant | MODERATE | 0 | 0 | 76 | 81 | . | . | . | . |
| PM1548-SK1P2 | CR2 | p.Thr209fs | frameshift_variant | HIGH | 0.6 | 0 | 10 | 118 | . | . | . | . |
| PM1548-SK1P2 | SMYD2 | p.Asn99fs | frameshift_variant | HIGH | 0.708333 | 0 | 24 | 181 | . | . | . | . |
| PM1548-SK1P2 | HHIPL2 | p.Gly156Ser | missense_variant | MODERATE | 0.234375 | 0 | 64 | 209 | . | . | . | . |
| PM1548-SK1P2 | MIA3 | p.Arg1545Gln | missense_variant | MODERATE | 0.363636 | 0 | 11 | 39 | . | . | . | . |
| PM1548-SK1P2 | LIN9 | p.Arg245Trp | missense_variant | MODERATE | 0 | 0 | 1 | 86 | . | . | . | . |
| PM1548-SK1P2 | ITPKB | p.Arg331His | missense_variant | MODERATE | 0.426451 | 0 | 741 | 395 | . | . | true | . |
| PM1548-SK1P2 | ITPKB | p.Leu42Val | missense_variant | MODERATE | 0.414938 | 0 | 241 | 34 | . | . | true | . |
| PM1548-SK1P2 | SNAP47 | p.Phe343Leu | missense_variant | MODERATE | 0.112676 | 0 | 71 | 77 | . | . | . | . |
| PM1548-SK1P2 | IBA57 | p.Ala161Val | missense_variant | MODERATE | 0.0169492 | 0 | 59 | 268 | . | . | . | . |
| PM1548-SK1P2 | OBSCN | p.Tyr2035Phe | missense_variant | MODERATE | 0.0141643 | 0 | 353 | 508 | . | . | . | . |
| PM1548-SK1P2 | OBSCN | p.Gln2065Lys | missense_variant | MODERATE | 0.0106383 | 0 | 94 | 171 | . | . | . | . |
| PM1548-SK1P2 | OBSCN | p.Cys2073Trp | missense_variant | MODERATE | 0 | 0 | 32 | 54 | . | . | . | . |
| PM1548-SK1P2 | ACTA1 | p.Thr231Met | missense_variant | MODERATE | 0 | 0 | 12 | 311 | . | . | . | . |
| PM1548-SK1P2 | GNPAT | p.Val551Met | missense_variant | MODERATE | 0.35 | 0 | 20 | 71 | . | . | . | . |
| PM1548-SK1P2 | DISC1 | p.Asp580Gly | missense_variant | MODERATE | 0.5 | 0 | 24 | 167 | . | . | . | . |
| PM1548-SK1P2 | GGPS1 | p.Val112Met | missense_variant | MODERATE | 0 | 0 | 54 | 405 | . | . | . | . |
| PM1548-SK1P2 | NID1 | p.Pro918fs | frameshift_variant&splice_region_variant | HIGH | 0.363636 | 0 | 33 | 47 | . | . | . | . |
| PM1548-SK1P2 | CEP170 | p.Glu1460Lys | missense_variant | MODERATE | 0.333333 | 0 | 6 | 32 | . | . | . | . |
| PM1548-SK1P2 | ZNF496 | p.Arg606His | missense_variant | MODERATE | 0.536913 | 0 | 298 | 143 | . | . | . | . |
| PM1548-SK1P2 | OR2T4 | p.Cys60Ser | missense_variant | MODERATE | 0 | 0 | 1 | 107 | . | . | . | . |
| PM1548-SK1P2 | OR2T6 | p.Tyr60His | missense_variant | MODERATE | 0.716981 | 0 | 53 | 68 | . | . | . | . |
| PM1548-SK1P2 | DIP2C | p.Val1150Ile | missense_variant | MODERATE | 0.171429 | 0 | 35 | 183 | . | . | . | . |
| PM1548-SK1P2 | WDR37 | p.Pro169Ser | missense_variant | MODERATE | 0.314815 | 0 | 54 | 220 | . | . | . | . |
| PM1548-SK1P2 | CALML5 | p.Glu102Lys | missense_variant | MODERATE | 0.546917 | 0 | 373 | 79 | . | . | . | . |
| PM1548-SK1P2 | FRMD4A | p.Pro1005fs | frameshift_variant | HIGH | 0 | 0 | 11 | 118 | . | . | . | . |
| PM1548-SK1P2 | CUBN | p.Trp3323Arg | missense_variant | MODERATE | 0 | 0 | 24 | 170 | . | . | . | . |
| PM1548-SK1P2 | GPR158 | p.Arg448Gln | missense_variant | MODERATE | 0.333333 | 0 | 12 | 52 | . | . | . | . |
| PM1548-SK1P2 | SVIL | p.Arg2075Cys | missense_variant | MODERATE | 0.5 | 0 | 10 | 40 | . | . | . | . |
| PM1548-SK1P2 | PARD3 | p.Arg1278Gly | missense_variant | MODERATE | 0 | 0 | 31 | 91 | . | . | . | . |
| PM1548-SK1P2 | FRMPD2 | p.Pro226Leu | missense_variant | MODERATE | 0.5 | 0 | 4 | 178 | . | . | . | . |
| PM1548-SK1P2 | PRKG1 | p.Asn601fs | frameshift_variant | HIGH | 0.320755 | 0 | 53 | 56 | . | . | . | . |
| PM1548-SK1P2 | PBLD | p.Asp167Tyr | missense_variant | MODERATE | 0.129032 | 0 | 31 | 185 | . | . | . | . |
| PM1548-SK1P2 | DDX50 | p.Tyr39Cys | missense_variant | MODERATE | 0.363636 | 0 | 11 | 39 | . | . | . | . |
| PM1548-SK1P2 | ADAMTS14 | p.Gly710Arg | missense_variant | MODERATE | 0.714286 | 0 | 42 | 78 | . | . | . | . |
| PM1548-SK1P2 | DDIT4 | p.Ser46Cys | missense_variant | MODERATE | 0.167939 | 0 | 131 | 262 | . | . | . | . |
| PM1548-SK1P2 | DDIT4 | p.Thr47Ala | missense_variant | MODERATE | 0.172932 | 0 | 133 | 262 | . | . | . | . |
| PM1548-SK1P2 | KAT6B | p.Arg463His | missense_variant | MODERATE | 0.547368 | 0 | 95 | 104 | true | . | . | . |
| PM1548-SK1P2 | KCNMA1 | p.Ala1039Val | missense_variant | MODERATE | 0.125 | 0 | 8 | 74 | . | . | . | . |
| PM1548-SK1P2 | ZMIZ1 | p.Asp1051fs | frameshift_variant | HIGH | 0.343284 | 0 | 67 | 71 | . | . | . | . |
| PM1548-SK1P2 | MAT1A | p.Arg81Gln | missense_variant | MODERATE | 0.25 | 0 | 36 | 225 | . | . | . | . |
| PM1548-SK1P2 | TSPAN14 | p.Asp190Gly | missense_variant | MODERATE | 0.114286 | 0 | 35 | 133 | . | . | . | . |
| PM1548-SK1P2 | SH2D4B | p.Ala186Val | missense_variant | MODERATE | 0.244898 | 0 | 49 | 42 | . | . | . | . |
| PM1548-SK1P2 | LRIT2 | p.Thr520fs | frameshift_variant | HIGH | 0.209302 | 0 | 43 | 223 | . | . | . | . |
| PM1548-SK1P2 | CCSER2 | c.2326-13521C> | intron_variant | MODIFIER | 0 | 0 | 17 | 179 | . | . | . | . |
| PM1548-SK1P2 | CYP26C1 | p.Ser70Leu | missense_variant | MODERATE | 0 | 0 | 172 | 149 | . | . | . | . |
| PM1548-SK1P2 | MYOF | p.Arg726Gln | missense_variant | MODERATE | 0.6 | 0 | 5 | 43 | . | . | . | . |
| PM1548-SK1P2 | SORBS1 | p.Arg585Cys | missense_variant | MODERATE | 0.153846 | 0 | 13 | 42 | . | . | . | . |
| PM1548-SK1P2 | ALDH18A1 | p.Arg622Trp | missense_variant | MODERATE | 0.733333 | 0 | 15 | 86 | . | . | . | . |
| PM1548-SK1P2 | ENTPD1 | p.Arg252Leu | missense_variant | MODERATE | 0.111111 | 0 | 36 | 84 | . | . | . | . |
| PM1548-SK1P2 | MORN4 | p.Leu202Ile | missense_variant | MODERATE | 0.111111 | 0 | 36 | 86 | . | . | . | . |
| PM1548-SK1P2 | PKD2L1 | p.Arg206Cys | missense_variant | MODERATE | 0.454106 | 0 | 207 | 130 | . | . | . | . |
| PM1548-SK1P2 | MRPL43 | p.Asn66Ser | missense_variant | MODERATE | 0.556034 | 0 | 232 | 81 | . | . | . | . |
| PM1548-SK1P2 | POLL | p.Arg419* | stop_gained | HIGH | 0 | 0 | 18 | 44 | . | . | . | . |

| | | | | | | | | | | | | |
|---|---|---|---|---|---|---|---|---|---|---|---|---|
| PM1548-SK1P2 | TAF5 | p.Phe374fs | frameshift_variant | HIGH | 0.555556 | 0 | 9 | 50 | . | . | . | . |
| PM1548-SK1P2 | WDR96 | p.Val70Ala | missense_variant | MODERATE | 0.333333 | 0 | 6 | 59 | . | . | . | . |
| PM1548-SK1P2 | SORCS3 | p.Lys1023Gln | missense_variant | MODERATE | 0 | 0 | 23 | 58 | . | . | . | . |
| PM1548-SK1P2 | TCF7L2 | p.Ser587Leu | missense_variant | MODERATE | 0 | 0 | 167 | 171 | true | . | true | . |
| PM1548-SK1P2 | ABLIM1 | p.Leu633Met | missense_variant | MODERATE | 0.114286 | 0 | 35 | 265 | . | . | . | . |
| PM1548-SK1P2 | GFRA1 | p.Ala173Val | missense_variant | MODERATE | 0.25 | 0 | 16 | 73 | . | . | . | . |
| PM1548-SK1P2 | KIAA1598 | p.Arg482His | missense_variant | MODERATE | 0 | 0 | 20 | 182 | true | . | . | . |
| PM1548-SK1P2 | VAX1 | p.Cys200* | stop_gained | HIGH | 0 | 0 | 602 | 96 | . | . | . | . |
| PM1548-SK1P2 | EIF3A | p.His257His | synonymous_variant | LOW | 0.105263 | 0 | 38 | 130 | . | . | . | . |
| PM1548-SK1P2 | ATE1 | p.Leu289Gln | missense_variant | MODERATE | 0 | 0 | 13 | 77 | . | . | . | . |
| PM1548-SK1P2 | PLEKHA1 | p.Arg9His | missense_variant | MODERATE | 1 | 0 | 4 | 43 | . | . | . | . |
| PM1548-SK1P2 | PLEKHA1 | p.Val404Met | missense_variant | MODERATE | 0.285714 | 0 | 28 | 76 | . | . | . | . |
| PM1548-SK1P2 | HMX3 | p.Pro23fs | frameshift_variant | HIGH | 0.856618 | 0 | 272 | 61 | . | . | . | . |
| PM1548-SK1P2 | CLRN3 | p.Pro71Thr | missense_variant | MODERATE | 0.113636 | 0.00613497 | 44 | 163 | . | . | . | . |
| PM1548-SK1P2 | MKI67 | p.Arg2258* | stop_gained | HIGH | 0.571429 | 0 | 14 | 90 | . | . | . | . |
| PM1548-SK1P2 | TTC40 | p.Thr1864Met | missense_variant | MODERATE | 0.594595 | 0 | 37 | 72 | . | . | . | . |
| PM1548-SK1P2 | KNDC1 | p.Cys79fs | frameshift_variant | HIGH | 0.47619 | 0 | 21 | 162 | . | . | . | . |
| PM1548-SK1P2 | PIDD | p.Ala727Glu | missense_variant | MODERATE | 0 | 0 | 450 | 108 | . | . | . | . |
| PM1548-SK1P2 | MUC6 | p.Thr1861Ala | missense_variant | MODERATE | 0.631579 | 0 | 19 | 78 | . | . | . | . |
| PM1548-SK1P2 | MUC6 | p.Ile1857Thr | missense_variant | MODERATE | 0.5625 | 0 | 16 | 78 | . | . | . | . |
| PM1548-SK1P2 | MUC6 | p.His1752Gln | missense_variant | MODERATE | 0.109091 | 0 | 55 | 31 | . | . | . | . |
| PM1548-SK1P2 | MUC6 | p.Thr1711Ile | missense_variant | MODERATE | 0.225 | 0 | 40 | 51 | . | . | . | . |
| PM1548-SK1P2 | MUC2 | p.Gly1331Arg | missense_variant | MODERATE | 0.202381 | 0 | 84 | 70 | . | . | . | . |
| PM1548-SK1P2 | MUC5B | p.Ala110Thr | missense_variant | MODERATE | 0.580097 | 0 | 824 | 216 | . | . | . | . |
| PM1548-SK1P2 | BRSK2 | p.Pro440fs | frameshift_variant | HIGH | 0.468456 | 0 | 2235 | 592 | . | . | . | . |
| PM1548-SK1P2 | TSPAN32 | p.Ala178Val | missense_variant | MODERATE | 0.417476 | 0 | 103 | 69 | . | . | . | . |
| PM1548-SK1P2 | TSSC4 | p.Arg92Cys | missense_variant | MODERATE | 0.502648 | 0 | 2266 | 524 | . | . | . | . |
| PM1548-SK1P2 | OSBPL5 | p.Thr843Met | missense_variant | MODERATE | 0 | 0 | 4 | 106 | . | . | . | . |
| PM1548-SK1P2 | STIM1 | p.Lys371Arg | missense_variant | MODERATE | 0.0645161 | 0 | 62 | 322 | . | . | . | . |
| PM1548-SK1P2 | OR51E1 | p.Tyr63His | missense_variant | MODERATE | 0.526132 | 0 | 287 | 116 | . | . | . | . |
| PM1548-SK1P2 | OR51A2 | p.Lys289Asn | missense_variant | MODERATE | 0 | 0 | 11 | 30 | . | . | . | . |
| PM1548-SK1P2 | SOX6 | p.Met803Ile | missense_variant | MODERATE | 0.125 | 0 | 40 | 33 | . | . | . | . |
| PM1548-SK1P2 | SOX6 | p.Met803Thr | missense_variant | MODERATE | 0.125 | 0 | 40 | 33 | . | . | . | . |
| PM1548-SK1P2 | SOX6 | p.Met803Val | missense_variant | MODERATE | 0.121951 | 0 | 41 | 33 | . | . | . | . |
| PM1548-SK1P2 | ABCC8 | p.Asp761Asn | missense_variant | MODERATE | 0.025 | 0 | 40 | 231 | . | . | . | . |
| PM1548-SK1P2 | LDHC | p.Lys5Glu | missense_variant | MODERATE | 0.3 | 0 | 10 | 119 | . | . | . | . |
| PM1548-SK1P2 | MRGPRX2 | p.Asn62Asn | synonymous_variant | LOW | 0 | 0 | 39 | 83 | . | . | . | . |
| PM1548-SK1P2 | ZDHHC13 | p.Gly591Asp | missense_variant | MODERATE | 0 | 0 | 2 | 33 | . | . | . | . |
| PM1548-SK1P2 | ANO3 | p.Leu273Pro | missense_variant | MODERATE | 0 | 0 | 43 | 435 | . | . | . | . |
| PM1548-SK1P2 | HIPK3 | p.Asp532Ala | missense_variant | MODERATE | 0.0434783 | 0 | 69 | 54 | . | . | . | . |
| PM1548-SK1P2 | ABTB2 | p.Asp793Asn | missense_variant | MODERATE | 0.533898 | 0.00668896 | 118 | 299 | . | . | . | . |
| PM1548-SK1P2 | RAG1 | p.Arg507Trp | missense_variant | MODERATE | 0 | 0 | 87 | 86 | . | . | . | true |
| PM1548-SK1P2 | TP53I11 | p.Leu62Arg | missense_variant | MODERATE | 0.0035461 | 0 | 282 | 34 | . | . | . | . |
| PM1548-SK1P2 | MAPK8IP1 | p.Thr321Met | missense_variant | MODERATE | 0.592727 | 0 | 275 | 188 | . | true | . | . |
| PM1548-SK1P2 | DGKZ | n.46369267G>A | intragenic_variant | MODIFIER | 0 | 0 | 47 | 32 | . | . | . | . |
| PM1548-SK1P2 | DGKZ | p.Asp574Asn | missense_variant | MODERATE | 0.155556 | 0 | 90 | 74 | . | . | . | . |
| PM1548-SK1P2 | ACP2 | p.Arg231His | missense_variant | MODERATE | 0.441805 | 0 | 421 | 199 | . | . | . | . |
| PM1548-SK1P2 | MYBPC3 | p.Arg654Ser | missense_variant | MODERATE | 0 | 0 | 20 | 97 | . | . | . | . |
| PM1548-SK1P2 | OR5D14 | p.Tyr261Cys | missense_variant | MODERATE | 0.166667 | 0 | 12 | 71 | . | . | . | . |
| PM1548-SK1P2 | OR5W2 | p.Ala29Ser | missense_variant | MODERATE | 0.384615 | 0 | 13 | 99 | . | . | . | . |
| PM1548-SK1P2 | APLNR | p.Ala161Thr | missense_variant | MODERATE | 0.642857 | 0 | 28 | 160 | . | . | . | . |
| PM1548-SK1P2 | OR10Q1 | p.Thr287Ile | missense_variant | MODERATE | 0.8125 | 0 | 16 | 90 | . | . | . | . |
| PM1548-SK1P2 | DTX4 | p.Lys396fs | frameshift_variant | HIGH | 0.421053 | 0 | 57 | 34 | . | . | . | . |
| PM1548-SK1P2 | DTX4 | p.Gly510fs | frameshift_variant | HIGH | 0.52381 | 0.00719424 | 42 | 139 | . | . | . | . |
| PM1548-SK1P2 | DAK | p.Arg380Gln | missense_variant | MODERATE | 0.340395 | 0 | 708 | 316 | . | . | . | . |
| PM1548-SK1P2 | FEN1 | p.Arg332Cys | missense_variant | MODERATE | 0.114286 | 0 | 35 | 79 | . | . | . | . |
| PM1548-SK1P2 | BEST1 | p.Ser413Asn | missense_variant | MODERATE | 0.481481 | 0 | 27 | 159 | . | . | . | . |
| PM1548-SK1P2 | AHNAK | p.Ser4908Leu | missense_variant | MODERATE | 0.26306 | 0 | 536 | 39 | . | . | . | . |
| PM1548-SK1P2 | AHNAK | p.His4772Asn | missense_variant | MODERATE | 0 | 0 | 103 | 54 | . | . | . | . |
| PM1548-SK1P2 | AHNAK | p.Gly4561Asp | missense_variant | MODERATE | 0.0582011 | 0 | 378 | 206 | . | . | . | . |
| PM1548-SK1P2 | AHNAK | p.Val4560Leu | missense_variant | MODERATE | 0.0633245 | 0 | 379 | 203 | . | . | . | . |
| PM1548-SK1P2 | AHNAK | p.Val3640Ile | missense_variant | MODERATE | 0.184615 | 0 | 65 | 44 | . | . | . | . |
| PM1548-SK1P2 | AHNAK | p.Asn3637Lys | missense_variant | MODERATE | 0.145161 | 0 | 62 | 44 | . | . | . | . |
| PM1548-SK1P2 | AHNAK | p.Lys2706fs | frameshift_variant | HIGH | 0.56213 | 0 | 169 | 43 | . | . | . | . |
| PM1548-SK1P2 | AHNAK | p.Thr1986Ala | missense_variant | MODERATE | 0.111111 | 0 | 72 | 323 | . | . | . | . |
| PM1548-SK1P2 | CHRM1 | p.Pro319fs | frameshift_variant | HIGH | 0 | 0 | 8 | 31 | . | . | . | . |
| PM1548-SK1P2 | SLC22A6 | p.Val240Met | missense_variant | MODERATE | 0 | 0 | 24 | 227 | . | . | . | . |
| PM1548-SK1P2 | SLC22A25 | p.Asn62Asp | missense_variant | MODERATE | 0 | 0 | 74 | 110 | . | . | . | . |
| PM1548-SK1P2 | C11orf95 | n.1245C>T | non_coding_exon_variant | MODIFIER | 0.490196 | 0 | 51 | 34 | . | . | . | . |
| PM1548-SK1P2 | NRXN2 | p.Asp277Asn | missense_variant | MODERATE | 0 | 0 | 115 | 229 | . | . | . | . |
| PM1548-SK1P2 | SNX15 | p.Leu10Met | missense_variant | MODERATE | 0 | 0 | 17 | 37 | . | . | . | . |
| PM1548-SK1P2 | RBM4 | p.Tyr303His | missense_variant | MODERATE | 0.3 | 0 | 20 | 269 | . | . | . | . |
| PM1548-SK1P2 | SPTBN2 | p.Cys1393Ser | missense_variant | MODERATE | 0.0364583 | 0 | 384 | 300 | . | . | . | . |
| PM1548-SK1P2 | SPTBN2 | p.Arg465Gln | missense_variant | MODERATE | 0.017094 | 0 | 117 | 212 | . | . | . | . |
| PM1548-SK1P2 | LRP5 | p.Ala724Val | missense_variant | MODERATE | 1 | 0 | 1 | 227 | . | . | . | . |
| PM1548-SK1P2 | IGHMBP2 | p.Pro809Ser | missense_variant | MODERATE | 0.537778 | 0 | 225 | 132 | . | . | . | . |
| PM1548-SK1P2 | TPCN2 | p.Gly492Val | missense_variant | MODERATE | 0.131579 | 0 | 38 | 32 | . | . | . | . |
| PM1548-SK1P2 | CCND1 | p.Val94Leu | missense_variant | MODERATE | 0.0487805 | 0 | 533 | 259 | true | true | true | . |
| PM1548-SK1P2 | INPPL1 | p.Leu957Phe | missense_variant | MODERATE | 0.288889 | 0 | 45 | 30 | . | . | . | . |
| PM1548-SK1P2 | PDE2A | p.His99fs | frameshift_variant | HIGH | 0.565149 | 0 | 637 | 161 | . | . | . | . |
| PM1548-SK1P2 | RNF169 | p.Glu131* | stop_gained | HIGH | 0 | 0 | 239 | 52 | . | . | . | . |
| PM1548-SK1P2 | TENM4 | p.Ala235Thr | missense_variant | MODERATE | 0.6 | 0 | 65 | 201 | . | . | . | . |
| PM1548-SK1P2 | PCF11 | p.Leu745Leu | synonymous_variant | LOW | 0.0103627 | 0 | 193 | 203 | . | . | . | . |
| PM1548-SK1P2 | PCF11 | p.Ser756Gly | missense_variant | MODERATE | 0.0102564 | 0 | 195 | 215 | . | . | . | . |
| PM1548-SK1P2 | PCF11 | p.Arg833Trp | missense_variant | MODERATE | 0.386918 | 0 | 902 | 258 | . | . | . | . |
| PM1548-SK1P2 | TYR | p.Gly191fs | frameshift_variant | HIGH | 0.5 | 0 | 6 | 94 | . | . | . | . |
| PM1548-SK1P2 | FAT3 | p.Ala2495Ser | missense_variant | MODERATE | 0.1 | 0 | 40 | 228 | . | . | . | . |
| PM1548-SK1P2 | PANX1 | p.Ser250Thr | missense_variant | MODERATE | 0.166667 | 0 | 12 | 236 | . | . | . | . |
| PM1548-SK1P2 | MTMR2 | p.Lys451Glu | missense_variant | MODERATE | 0.181818 | 0 | 33 | 77 | . | . | . | . |
| PM1548-SK1P2 | MTMR2 | p.Asp436Asn | missense_variant | MODERATE | 0.1875 | 0 | 32 | 75 | . | . | . | . |
| PM1548-SK1P2 | MTMR2 | p.Leu433Val | missense_variant | MODERATE | 0.162162 | 0 | 37 | 282 | . | . | . | . |
| PM1548-SK1P2 | MMP3 | p.Asp200Gly | missense_variant | MODERATE | 0.102564 | 0 | 39 | 91 | . | . | . | . |
| PM1548-SK1P2 | DYNC2H1 | p.Tyr3523His | missense_variant | MODERATE | 0.5 | 0 | 4 | 62 | . | . | . | . |
| PM1548-SK1P2 | CASP1 | p.Phe262Tyr | missense_variant | MODERATE | 0.888889 | 0 | 9 | 177 | . | . | . | . |
| PM1548-SK1P2 | KBTBD3 | p.Lys478fs | frameshift_variant | HIGH | 0.484848 | 0 | 33 | 50 | . | . | . | . |
| PM1548-SK1P2 | NXPE1 | p.Thr169Ile | missense_variant | MODERATE | 0.125 | 0 | 16 | 97 | . | . | . | . |
| PM1548-SK1P2 | NXPE1 | p.Met157Ile | missense_variant | MODERATE | 0.625 | 0 | 16 | 97 | . | . | . | . |
| PM1548-SK1P2 | NXPE1 | p.Met157Thr | missense_variant | MODERATE | 0.625 | 0 | 16 | 96 | . | . | . | . |
| PM1548-SK1P2 | NXPE1 | p.Thr149Lys | missense_variant | MODERATE | 0.714286 | 0 | 14 | 118 | . | . | . | . |
| PM1548-SK1P2 | DSCAML1 | p.Val1762Ile | missense_variant | MODERATE | 0.494624 | 0 | 93 | 226 | . | . | . | . |
| PM1548-SK1P2 | DSCAML1 | p.Asp949fs | frameshift_variant | HIGH | 0.59799 | 0 | 597 | 245 | . | . | . | . |
| PM1548-SK1P2 | ATP5L | p.Leu84Trp | missense_variant | MODERATE | 0.0677966 | 0 | 59 | 125 | . | . | . | . |
| PM1548-SK1P2 | ATP5L | p.Val89Ile | missense_variant | MODERATE | 0.0322581 | 0 | 62 | 140 | . | . | . | . |
| PM1548-SK1P2 | ATP5L | p.Ile99Val | missense_variant | MODERATE | 0.031746 | 0 | 63 | 141 | . | . | . | . |
| PM1548-SK1P2 | PHLDB1 | p.Ala932Ser | missense_variant | MODERATE | 0.147059 | 0 | 68 | 223 | . | . | . | . |
| PM1548-SK1P2 | PHLDB1 | p.Pro938Leu | missense_variant | MODERATE | 0.114286 | 0 | 70 | 219 | . | . | . | . |
| PM1548-SK1P2 | CBL | p.Ser237Leu | missense_variant | MODERATE | 0.5 | 0 | 10 | 128 | true | . | . | . |
| PM1548-SK1P2 | CBL | p.Pro504Gln | missense_variant | MODERATE | 0 | 0 | 29 | 111 | true | . | . | . |
| PM1548-SK1P2 | CRTAM | p.Leu60fs | frameshift_variant | HIGH | 0.451282 | 0 | 195 | 170 | . | . | . | . |
| PM1548-SK1P2 | OR10G8 | p.Lys265Glu | missense_variant | MODERATE | 0.107784 | 0 | 167 | 231 | . | . | . | . |
| PM1548-SK1P2 | OR10G8 | p.Lys265Asn | missense_variant | MODERATE | 0.110429 | 0 | 163 | 192 | . | . | . | . |
| PM1548-SK1P2 | ROBO4 | p.Arg776His | missense_variant | MODERATE | 0.480519 | 0 | 77 | 263 | . | . | . | . |
| PM1548-SK1P2 | ADAMTS8 | p.Arg9Gly | missense_variant | MODERATE | 0.367816 | 0 | 87 | 48 | . | . | . | true |
| PM1548-SK1P2 | IQSEC3 | p.Gln1075Lys | missense_variant | MODERATE | 0 | 0 | 307 | 59 | . | . | . | . |
| PM1548-SK1P2 | FOXM1 | p.Leu334Ser | missense_variant | MODERATE | 0.117647 | 0 | 68 | 177 | . | . | . | . |
| PM1548-SK1P2 | FOXM1 | p.Asp328Glu | missense_variant | MODERATE | 0.121212 | 0 | 66 | 203 | . | . | . | . |
| PM1548-SK1P2 | VWF | p.Ala865Asp | missense_variant | MODERATE | 0.016835 | 0 | 297 | 226 | . | . | . | . |
| PM1548-SK1P2 | IFFO1 | p.Val126Met | missense_variant | MODERATE | 0.445026 | 0 | 382 | 191 | . | . | . | . |
| PM1548-SK1P2 | PHB2 | p.Arg270Arg | synonymous_variant | LOW | 0.10929 | 0 | 183 | 402 | . | . | . | . |
| PM1548-SK1P2 | PHB2 | p.Asn269Asn | synonymous_variant | LOW | 0.10989 | 0 | 182 | 400 | . | . | . | . |
| PM1548-SK1P2 | TAS2R10 | p.Ser41Thr | missense_variant | MODERATE | 0.333333 | 0 | 18 | 57 | . | . | . | . |
| PM1548-SK1P2 | TAS2R31 | p.Ala177Thr | missense_variant | MODERATE | 0 | 0 | 8 | 47 | . | . | . | . |
| PM1548-SK1P2 | PLCZ1 | p.Gly133Val | missense_variant | MODERATE | 0 | 0 | 2 | 42 | . | . | . | . |
| PM1548-SK1P2 | PDE3A | p.Glu171fs | frameshift_variant | HIGH | 0.485714 | 0 | 105 | 56 | . | . | . | . |
| PM1548-SK1P2 | TM7SF3 | c.869-699C>G | intron_variant | MODIFIER | 0.423077 | 0 | 52 | 125 | . | . | . | . |
| PM1548-SK1P2 | DDX11 | p.Asp935Asn | missense_variant | MODERATE | 0.666667 | 0 | 9 | 35 | . | . | . | . |
| PM1548-SK1P2 | COL2A1 | p.Asp682Asn | missense_variant | MODERATE | 0.554913 | 0 | 519 | 53 | true | . | . | . |

| | | | | | | | | | | | | |
|---|---|---|---|---|---|---|---|---|---|---|---|---|
| PM1548-SK1P2 | KMT2D | p.Ala4804Val | missense_variant | MODERATE | 0.431034 | 0.0060241 | 58 | 166 | true | true | true | true |
| PM1548-SK1P2 | KMT2D | p.Leu4214Leu | synonymous_variant | LOW | 0.101695 | 0 | 59 | 373 | true | true | true | true |
| PM1548-SK1P2 | KMT2D | p.Thr3166Ile | missense_variant | MODERATE | 0.327982 | 0 | 436 | 193 | true | true | true | true |
| PM1548-SK1P2 | KMT2D | p.Thr2581Ala | missense_variant | MODERATE | 0.0163132 | 0 | 613 | 134 | true | true | true | true |
| PM1548-SK1P2 | KMT2D | p.Gly1628fs | frameshift_variant | HIGH | 0.0283688 | 0 | 141 | 136 | true | true | true | true |
| PM1548-SK1P2 | PRPH | p.Ala348Thr | missense_variant | MODERATE | 0.518987 | 0 | 79 | 109 | . | . | . | . |
| PM1548-SK1P2 | FAM186B | p.Leu494Met | missense_variant | MODERATE | 0.548896 | 0 | 317 | 412 | . | . | . | . |
| PM1548-SK1P2 | ANKRD33 | p.Ala173Val | missense_variant | MODERATE | 0.15625 | 0 | 480 | 241 | . | . | . | . |
| PM1548-SK1P2 | KRT74 | p.Gln440His | missense_variant | MODERATE | 0.105263 | 0 | 38 | 34 | . | . | . | . |
| PM1548-SK1P2 | PCBP2 | p.Pro175fs | frameshift_variant | HIGH | 0.25 | 0 | 16 | 35 | . | . | . | . |
| PM1548-SK1P2 | HNRNPA1 | p.Thr61Ala | missense_variant | MODERATE | 0.558824 | 0 | 34 | 168 | . | . | . | . |
| PM1548-SK1P2 | ITGA5 | p.Arg193His | missense_variant | MODERATE | 0.478873 | 0 | 71 | 104 | . | . | . | . |
| PM1548-SK1P2 | MYO1A | p.Arg559Cys | missense_variant | MODERATE | 0.546154 | 0 | 520 | 190 | . | . | . | . |
| PM1548-SK1P2 | NAB2 | p.Pro388Thr | missense_variant | MODERATE | 0.0425532 | 0 | 47 | 78 | true | . | . | . |
| PM1548-SK1P2 | NAB2 | p.Pro388His | missense_variant | MODERATE | 0.0425532 | 0 | 47 | 78 | true | . | . | . |
| PM1548-SK1P2 | R3HDM2 | p.Arg956His | missense_variant | MODERATE | 0 | 0 | 25 | 33 | . | . | . | . |
| PM1548-SK1P2 | CTDSP2 | p.Glu259Asp | missense_variant | MODERATE | 0.133333 | 0 | 15 | 90 | . | . | . | . |
| PM1548-SK1P2 | CTDSP2 | p.Ala258Thr | missense_variant | MODERATE | 0.133333 | 0 | 15 | 90 | . | . | . | . |
| PM1548-SK1P2 | CTDSP2 | p.Ile251Val | missense_variant | MODERATE | 0.181818 | 0 | 11 | 94 | . | . | . | . |
| PM1548-SK1P2 | PTPRB | p.Val1992Ala | missense_variant | MODERATE | 0.0769231 | 0 | 26 | 51 | true | . | . | . |
| PM1548-SK1P2 | NAV3 | p.Met1275Val | missense_variant | MODERATE | 0.277778 | 0 | 18 | 67 | . | true | . | . |
| PM1548-SK1P2 | NAV3 | p.Pro1276Leu | missense_variant | MODERATE | 0.30303 | 0 | 33 | 130 | . | true | . | . |
| PM1548-SK1P2 | NAV3 | p.Thr1281Ser | missense_variant | MODERATE | 0.285714 | 0 | 35 | 144 | . | true | . | . |
| PM1548-SK1P2 | TMTC2 | p.Gly143Arg | missense_variant | MODERATE | 0.552632 | 0 | 38 | 67 | . | . | . | . |
| PM1548-SK1P2 | ALX1 | p.Thr68Ser | missense_variant | MODERATE | 0.5 | 0 | 4 | 191 | . | . | . | . |
| PM1548-SK1P2 | FGD6 | p.Met965fs | frameshift_variant | HIGH | 0.833333 | 0 | 6 | 56 | . | . | . | . |
| PM1548-SK1P2 | APAF1 | p.Gln578Arg | missense_variant | MODERATE | 0.0392157 | 0 | 51 | 126 | . | . | . | . |
| PM1548-SK1P2 | SLC41A2 | p.Pro266Ser | missense_variant | MODERATE | 1 | 0 | 2 | 65 | . | . | . | . |
| PM1548-SK1P2 | PRDM4 | p.Glu313Glu | synonymous_variant | LOW | 0.0666667 | 0 | 120 | 274 | . | . | . | . |
| PM1548-SK1P2 | TMEM119 | p.Thr81fs | frameshift_variant | HIGH | 0.40604 | 0 | 1192 | 297 | . | . | . | . |
| PM1548-SK1P2 | FOXN4 | p.Ala462Thr | missense_variant | MODERATE | 0.444444 | 0.00492611 | 27 | 203 | . | . | . | . |
| PM1548-SK1P2 | FAM222A | p.Pro392Thr | missense_variant | MODERATE | 0.0357143 | 0 | 140 | 422 | . | . | . | . |
| PM1548-SK1P2 | FAM222A | p.Ser400Ala | missense_variant | MODERATE | 0.0342466 | 0 | 146 | 423 | . | . | . | . |
| PM1548-SK1P2 | TCHP | p.Ala371Thr | missense_variant | MODERATE | 0.375 | 0 | 8 | 45 | . | . | . | . |
| PM1548-SK1P2 | SH2B3 | p.Glu523fs | frameshift_variant | HIGH | 0 | 0 | 452 | 158 | true | . | . | . |
| PM1548-SK1P2 | HECTD4 | p.Ala3707Val | missense_variant | MODERATE | 0.333333 | 0 | 18 | 63 | . | . | . | . |
| PM1548-SK1P2 | TPCN1 | p.Leu234Arg | missense_variant | MODERATE | 0.258741 | 0 | 286 | 134 | . | . | . | . |
| PM1548-SK1P2 | RBM19 | p.Asn533Asn | synonymous_variant | LOW | 0.488806 | 0 | 268 | 249 | . | . | . | . |
| PM1548-SK1P2 | RAB35 | n.120541738C>T | intragenic_variant | MODIFIER | 0.495327 | 0 | 321 | 243 | . | . | . | . |
| PM1548-SK1P2 | GCN1L1 | p.Ser2359Pro | missense_variant | MODERATE | 0.222222 | 0 | 9 | 85 | . | . | . | . |
| PM1548-SK1P2 | SETD1B | p.His8fs | frameshift_variant | HIGH | 0.0204082 | 0 | 49 | 141 | . | . | . | . |
| PM1548-SK1P2 | SETD1B | p.Pro1361fs | frameshift_variant | HIGH | 0.641791 | 0 | 134 | 190 | . | . | . | . |
| PM1548-SK1P2 | ZCCHC8 | p.Gly551Asp | missense_variant | MODERATE | 0 | 0 | 10 | 34 | true | . | . | . |
| PM1548-SK1P2 | ABCB9 | p.His709Gln | missense_variant | MODERATE | 0.0928571 | 0 | 140 | 333 | . | . | . | . |
| PM1548-SK1P2 | ABCB9 | p.His709Arg | missense_variant | MODERATE | 0.0928571 | 0 | 140 | 333 | . | . | . | . |
| PM1548-SK1P2 | ABCB9 | p.Lys694Arg | missense_variant | MODERATE | 0.0921986 | 0 | 141 | 301 | . | . | . | . |
| PM1548-SK1P2 | PITPNM2 | p.Arg749Trp | missense_variant | MODERATE | 0.325 | 0 | 160 | 160 | . | . | . | . |
| PM1548-SK1P2 | SBNO1 | p.Arg1283Cys | missense_variant&splice_region_variant | MODERATE | 0.365385 | 0 | 52 | 244 | . | . | . | . |
| PM1548-SK1P2 | DNAH10 | p.Pro3133His | missense_variant | MODERATE | 0.388889 | 0.00452489 | 18 | 221 | . | . | . | . |
| PM1548-SK1P2 | TMEM132B | p.Ala297Thr | missense_variant | MODERATE | 0 | 0 | 5 | 79 | . | . | . | . |
| PM1548-SK1P2 | GPR133 | p.Gly731Arg | missense_variant | MODERATE | 0.388889 | 0 | 18 | 110 | . | . | . | . |
| PM1548-SK1P2 | FBRSL1 | p.Glu251Asp | missense_variant | MODERATE | 0 | 0 | 1136 | 153 | . | . | . | . |
| PM1548-SK1P2 | MPHOSPH8 | p.Val686Ile | missense_variant | MODERATE | 0 | 0 | 71 | 75 | . | . | . | . |
| PM1548-SK1P2 | GJA3 | p.Ala213Thr | missense_variant | MODERATE | 0.0361446 | 0 | 249 | 153 | . | . | . | . |
| PM1548-SK1P2 | LATS2 | p.Pro41Gln | missense_variant | MODERATE | 0.466667 | 0.025641 | 15 | 39 | . | . | . | . |
| PM1548-SK1P2 | CDK8 | p.Asn372Thr | missense_variant | MODERATE | 0 | 0 | 13 | 249 | . | . | . | . |
| PM1548-SK1P2 | POLR1D | n.28240022G>A | intragenic_variant | MODIFIER | 0.492795 | 0 | 347 | 101 | . | . | . | . |
| PM1548-SK1P2 | FRY | p.Gln2970His | missense_variant | MODERATE | 0.181818 | 0 | 22 | 39 | . | . | . | . |
| PM1548-SK1P2 | BRCA2 | p.Asn863fs | frameshift_variant | HIGH | 0.466667 | 0 | 45 | 85 | true | true | . | true |
| PM1548-SK1P2 | STARD13 | p.Arg155His | missense_variant | MODERATE | 0 | 0 | 19 | 32 | . | . | . | . |
| PM1548-SK1P2 | MAB21L1 | p.Ser103Ser | synonymous_variant | LOW | 0.375 | 0 | 8 | 101 | . | . | . | . |
| PM1548-SK1P2 | FOXO1 | p.Ser152Thr | missense_variant | MODERATE | 0 | 0 | 126 | 140 | true | . | . | . |
| PM1548-SK1P2 | KBTBD7 | p.Tyr195Phe | missense_variant | MODERATE | 0.102564 | 0 | 39 | 644 | . | . | . | . |
| PM1548-SK1P2 | GPALPP1 | p.Pro37Arg | missense_variant | MODERATE | 1 | 0 | 1 | 103 | . | . | . | . |
| PM1548-SK1P2 | FAM124A | p.Ser257Pro | missense_variant | MODERATE | 0 | 0 | 3 | 101 | . | . | . | . |
| PM1548-SK1P2 | NEK3 | p.Asn343Lys | missense_variant | MODERATE | 0 | 0 | 18 | 44 | . | . | . | . |
| PM1548-SK1P2 | DACH1 | p.Arg196Arg | synonymous_variant | LOW | 0.0526316 | 0 | 19 | 115 | . | . | . | . |
| PM1548-SK1P2 | ABCC4 | p.Ile1219fs | frameshift_variant | HIGH | 0 | 0 | 63 | 130 | . | . | . | . |
| PM1548-SK1P2 | FGF14 | p.Asp33Gly | missense_variant | MODERATE | 0.133333 | 0 | 15 | 97 | . | . | . | . |
| PM1548-SK1P2 | CCDC168 | p.Met4045fs | frameshift_variant | HIGH | 0.44898 | 0 | 49 | 94 | . | . | . | . |
| PM1548-SK1P2 | ARHGEF7 | p.Thr23fs | frameshift_variant | HIGH | 0.0100334 | 0 | 598 | 189 | . | . | . | . |
| PM1548-SK1P2 | MCF2L | p.Arg590Gln | missense_variant | MODERATE | 0.461538 | 0 | 13 | 34 | . | . | . | . |
| PM1548-SK1P2 | CDC16 | p.Asn557fs | frameshift_variant | HIGH | 0 | 0 | 10 | 90 | . | . | . | . |
| PM1548-SK1P2 | UPF3A | p.Asp129Glu | missense_variant | MODERATE | 0.538721 | 0 | 297 | 60 | . | . | . | . |
| PM1548-SK1P2 | OR11G2 | p.Phe141Phe | synonymous_variant | LOW | 0.1 | 0 | 20 | 117 | . | . | . | . |
| PM1548-SK1P2 | ACIN1 | p.Glu192Asp | missense_variant | MODERATE | 0.129032 | 0 | 31 | 128 | . | . | . | . |
| PM1548-SK1P2 | ACIN1 | p.Glu192Lys | missense_variant | MODERATE | 0.129032 | 0 | 31 | 129 | . | . | . | . |
| PM1548-SK1P2 | PABPN1 | c.*5dupA | frameshift_variant&stop_retained_variant | HIGH | 0 | 0 | 8 | 92 | . | . | . | . |
| PM1548-SK1P2 | RNF31 | p.Ala9Gly | missense_variant | MODERATE | 0.105263 | 0 | 38 | 103 | . | . | . | true |
| PM1548-SK1P2 | RNF31 | p.Val12Ala | missense_variant | MODERATE | 0.105263 | 0 | 38 | 104 | . | . | . | true |
| PM1548-SK1P2 | LTB4R2 | p.Ala92Val | missense_variant | MODERATE | 0.105422 | 0 | 332 | 173 | . | . | . | . |
| PM1548-SK1P2 | FOXG1 | p.Glu173Asp | missense_variant | MODERATE | 0.131313 | 0 | 99 | 31 | . | . | . | . |
| PM1548-SK1P2 | CLEC14A | p.Ser404Asn | missense_variant | MODERATE | 0 | 0 | 51 | 138 | . | . | . | . |
| PM1548-SK1P2 | KTN1 | p.Phe1182Val | missense_variant | MODERATE | 0.565217 | 0 | 23 | 114 | true | . | . | . |
| PM1548-SK1P2 | KCNH5 | p.Arg333His | missense_variant | MODERATE | 0.621622 | 0 | 37 | 68 | . | . | . | . |
| PM1548-SK1P2 | ZBTB25 | p.His371Asn | missense_variant | MODERATE | 0.125 | 0 | 32 | 47 | . | . | . | . |
| PM1548-SK1P2 | HSPA2 | p.Gly138fs | frameshift_variant | HIGH | 0.465517 | 0 | 232 | 147 | . | . | . | . |
| PM1548-SK1P2 | HSPA2 | p.Phe220Phe | synonymous_variant | LOW | 0.0531915 | 0 | 94 | 96 | . | . | . | . |
| PM1548-SK1P2 | FUT8 | p.Arg249Cys | missense_variant | MODERATE | 0.466667 | 0 | 15 | 330 | . | . | . | . |
| PM1548-SK1P2 | TMEM229B | p.Leu37Phe | missense_variant | MODERATE | 0.00952381 | 0 | 210 | 483 | . | . | . | . |
| PM1548-SK1P2 | ZFP36L1 | p.Gly271fs | frameshift_variant | HIGH | 0.564202 | 0 | 257 | 216 | . | . | . | . |
| PM1548-SK1P2 | ELMSAN1 | p.Arg739Cys | missense_variant | MODERATE | 0.464912 | 0 | 114 | 46 | . | . | . | . |
| PM1548-SK1P2 | ELMSAN1 | p.Gln36fs | frameshift_variant | HIGH | 0.48731 | 0 | 197 | 44 | . | . | . | . |
| PM1548-SK1P2 | VRTN | p.Gly328fs | frameshift_variant | HIGH | 0 | 0 | 17 | 58 | . | . | . | . |
| PM1548-SK1P2 | GALC | p.Arg685His | missense_variant | MODERATE | 0 | 0 | 25 | 31 | . | . | . | . |
| PM1548-SK1P2 | PTPN21 | p.Ile849fs | frameshift_variant | HIGH | 0.5 | 0 | 4 | 156 | . | . | . | . |
| PM1548-SK1P2 | ZC3H14 | p.Asn577Asp | missense_variant | MODERATE | 0.285714 | 0 | 28 | 222 | . | . | . | . |
| PM1548-SK1P2 | CCDC88C | p.Lys818Glu | missense_variant | MODERATE | 0.133333 | 0 | 30 | 69 | . | . | . | . |
| PM1548-SK1P2 | FBLN5 | p.Asp357Asn | missense_variant | MODERATE | 0.5 | 0 | 8 | 147 | . | . | . | . |
| PM1548-SK1P2 | UNC79 | p.Arg746Gln | missense_variant | MODERATE | 0.25 | 0 | 12 | 31 | . | . | . | . |
| PM1548-SK1P2 | SYNE3 | p.Ile83Val | missense_variant | MODERATE | 0.392523 | 0 | 107 | 217 | . | . | . | . |
| PM1548-SK1P2 | TECPR2 | p.Gln498fs | frameshift_variant | HIGH | 0 | 0 | 1 | 55 | . | . | . | . |
| PM1548-SK1P2 | BRF1 | p.Ala129Ser | missense_variant | MODERATE | 0.277778 | 0 | 36 | 135 | . | . | . | . |
| PM1548-SK1P2 | CYFIP1 | p.Arg1085Leu | missense_variant | MODERATE | 0.111111 | 0 | 36 | 330 | . | . | . | . |
| PM1548-SK1P2 | MAGEL2 | p.Ala11Val | missense_variant | MODERATE | 0.356204 | 0 | 685 | 317 | . | . | . | . |
| PM1548-SK1P2 | PAR5 | n.2021_2022del | intron_variant | MODIFIER | 0.28 | 0 | 25 | 165 | . | . | . | . |
| PM1548-SK1P2 | UBE3A | p.Val133Ala | missense_variant | MODERATE | 0.666667 | 0 | 6 | 79 | . | . | . | . |
| PM1548-SK1P2 | ACTC1 | p.Ala137Thr | missense_variant | MODERATE | 1 | 0 | 11 | 73 | . | . | . | . |
| PM1548-SK1P2 | SPRED1 | p.Arg332His | missense_variant | MODERATE | 0.375 | 0.0070922 | 24 | 141 | . | . | . | . |
| PM1548-SK1P2 | CASC5 | p.Asp178Gly | missense_variant | MODERATE | 0 | 0 | 9 | 58 | true | . | . | . |
| PM1548-SK1P2 | SPTBN5 | p.Arg1943His | missense_variant | MODERATE | 0.431373 | 0 | 51 | 148 | . | . | . | . |
| PM1548-SK1P2 | SPTBN5 | p.Val1842Gly | missense_variant | MODERATE | 0.1 | 0 | 30 | 86 | . | . | . | . |
| PM1548-SK1P2 | GANC | p.Ala394Val | missense_variant | MODERATE | 0 | 0 | 18 | 106 | . | . | . | . |
| PM1548-SK1P2 | GANC | p.Gly404Asp | missense_variant | MODERATE | 0 | 0 | 4 | 105 | . | . | . | . |
| PM1548-SK1P2 | TTBK2 | p.Ala197Thr | missense_variant | MODERATE | 1 | 0 | 1 | 69 | . | . | . | . |
| PM1548-SK1P2 | CCNDBP1 | p.Asp237Asn | missense_variant | MODERATE | 0 | 0 | 9 | 44 | . | . | . | . |
| PM1548-SK1P2 | TGM7 | p.Ser305Phe | missense_variant | MODERATE | 0.296296 | 0 | 27 | 54 | . | . | . | . |
| PM1548-SK1P2 | ZSCAN29 | p.Arg33Trp | missense_variant | MODERATE | 0 | 0 | 45 | 209 | . | . | . | . |
| PM1548-SK1P2 | SLC28A2 | p.Gly128Asp | missense_variant | MODERATE | 0.215385 | 0 | 65 | 169 | . | . | . | . |
| PM1548-SK1P2 | MYO5A | p.Arg831His | missense_variant | MODERATE | 0.549327 | 0 | 446 | 78 | true | . | . | . |
| PM1548-SK1P2 | TCF12 | p.Val183fs | frameshift_variant | HIGH | 0 | 0 | 6 | 200 | true | . | . | . |
| PM1548-SK1P2 | LIPC | p.Phe286fs | frameshift_variant | HIGH | 0.55 | 0 | 20 | 89 | . | . | . | . |
| PM1548-SK1P2 | OAZ2 | p.Ser92Ser | synonymous_variant | LOW | 0.146341 | 0 | 41 | 163 | . | . | . | . |
| PM1548-SK1P2 | OAZ2 | p.Glu87Gly | missense_variant | MODERATE | 0.142857 | 0 | 42 | 159 | . | . | . | . |
| PM1548-SK1P2 | OAZ2 | p.Glu84Val | missense_variant | MODERATE | 0.146341 | 0 | 41 | 247 | . | . | . | . |
| PM1548-SK1P2 | OAZ2 | p.Val79Gly | missense_variant | MODERATE | 0.102564 | 0 | 39 | 233 | . | . | . | . |
| PM1548-SK1P2 | SPG21 | p.Ala282Val | missense_variant | MODERATE | 0.363636 | 0 | 11 | 32 | . | . | . | . |

| | | | | | | | | | | | | |
|---|---|---|---|---|---|---|---|---|---|---|---|---|
| PM1548-SK1P2 | CILP | p.Asn930Asn | synonymous_variant | LOW | 0.0512821 | 0.00877193 | 39 | 114 | . | . | . | . |
| PM1548-SK1P2 | IGDCC3 | p.Pro358Leu | missense_variant | MODERATE | 0 | 0 | 491 | 237 | . | . | . | . |
| PM1548-SK1P2 | NOX5 | p.His470fs | frameshift_variant | HIGH | 0.317073 | 0 | 41 | 164 | . | . | . | . |
| PM1548-SK1P2 | MYO9A | p.Cys1767Ser | missense_variant | MODERATE | 0.0444444 | 0 | 45 | 186 | . | . | . | . |
| PM1548-SK1P2 | NPTN | p.Arg334Trp | missense_variant | MODERATE | 0.262295 | 0 | 61 | 86 | . | . | . | . |
| PM1548-SK1P2 | ISLR2 | p.Val74Ala | missense_variant | MODERATE | 0.519685 | 0 | 381 | 331 | . | . | . | . |
| PM1548-SK1P2 | C15orf39 | p.Ala475Val | missense_variant | MODERATE | 0.446593 | 0.00393701 | 1086 | 254 | . | . | . | . |
| PM1548-SK1P2 | ETFA | p.Thr45Arg | missense_variant | MODERATE | 0.0740741 | 0 | 54 | 214 | . | . | . | . |
| PM1548-SK1P2 | ETFA | p.Thr45Ala | missense_variant | MODERATE | 0.0740741 | 0 | 54 | 214 | . | . | . | . |
| PM1548-SK1P2 | PSTPIP1 | p.Ala280fs | frameshift_variant&splice_region_variant | HIGH | 0.485618 | 0 | 591 | 440 | . | . | . | . |
| PM1548-SK1P2 | PEAK1 | p.Ala36Thr | missense_variant | MODERATE | 0 | 0 | 22 | 189 | . | . | . | . |
| PM1548-SK1P2 | CHRNB4 | p.Gly261Asp | missense_variant | MODERATE | 0.4 | 0 | 15 | 56 | . | . | . | . |
| PM1548-SK1P2 | MESDC1 | p.Ser224Asn | missense_variant | MODERATE | 0.510417 | 0 | 768 | 161 | . | . | . | . |
| PM1548-SK1P2 | IL16 | p.Thr417Met | missense_variant | MODERATE | 0 | 0 | 20 | 34 | . | . | . | . |
| PM1548-SK1P2 | BTBD1 | p.Arg236Gln | missense_variant | MODERATE | 0 | 0 | 7 | 77 | . | . | . | . |
| PM1548-SK1P2 | ZNF592 | p.Ala1232Val | missense_variant | MODERATE | 0.5 | 0 | 2 | 35 | . | . | . | . |
| PM1548-SK1P2 | AEN | p.Ala194Val | missense_variant | MODERATE | 0 | 0 | 15 | 122 | . | . | . | . |
| PM1548-SK1P2 | IQGAP1 | p.Met1231fs | frameshift_variant | HIGH | 0.482866 | 0 | 321 | 170 | . | . | . | . |
| PM1548-SK1P2 | FES | c.1531-47G>A | intron_variant | MODIFIER | 0.375502 | 0 | 498 | 35 | . | . | . | . |
| PM1548-SK1P2 | MAN2A2 | p.Pro139Leu | missense_variant | MODERATE | 0.605263 | 0 | 266 | 490 | . | . | . | . |
| PM1548-SK1P2 | ADAMTS17 | p.Cys921Tyr | missense_variant | MODERATE | 0.047619 | 0 | 21 | 58 | . | . | . | . |
| PM1548-SK1P2 | WDR90 | p.Ala1738Thr | missense_variant | MODERATE | 0.495763 | 0 | 944 | 185 | . | . | . | . |
| PM1548-SK1P2 | RPUSD1 | p.Arg255Gln | missense_variant | MODERATE | 0.507227 | 0 | 761 | 198 | . | . | . | . |
| PM1548-SK1P2 | SOX8 | p.Pro11Thr | missense_variant | MODERATE | 0 | 0 | 253 | 62 | . | . | . | . |
| PM1548-SK1P2 | SOX8 | p.Asp27Asn | missense_variant | MODERATE | 0 | 0 | 494 | 120 | . | . | . | . |
| PM1548-SK1P2 | CACNA1H | p.Arg1892His | missense_variant | MODERATE | 0.267884 | 0 | 657 | 231 | . | . | . | . |
| PM1548-SK1P2 | BAIAP3 | p.Ala740Asp | missense_variant | MODERATE | 0 | 0 | 49 | 90 | . | . | . | . |
| PM1548-SK1P2 | IFT140 | p.Ala1330Val | missense_variant | MODERATE | 0 | 0 | 767 | 188 | . | . | . | . |
| PM1548-SK1P2 | NME3 | p.Ala20Ser | missense_variant | MODERATE | 0 | 0 | 620 | 162 | . | . | . | . |
| PM1548-SK1P2 | PRSS33 | p.Leu174fs | frameshift_variant | HIGH | 0 | 0 | 606 | 199 | . | . | . | . |
| PM1548-SK1P2 | CCDC64B | p.Arg144Gln | missense_variant | MODERATE | 0 | 0.00254453 | 1211 | 393 | . | . | . | . |
| PM1548-SK1P2 | CREBBP | p.Gly1479Val | missense_variant | MODERATE | 0.121212 | 0 | 33 | 193 | true | . | true | . |
| PM1548-SK1P2 | CORO7 | p.Thr230Ala | missense_variant | MODERATE | 0.537037 | 0 | 486 | 171 | . | . | . | . |
| PM1548-SK1P2 | GLYR1 | p.Ser148Thr | missense_variant | MODERATE | 0.208955 | 0 | 67 | 42 | . | . | . | . |
| PM1548-SK1P2 | TVP23A | p.Phe140fs | frameshift_variant | HIGH | 0.457859 | 0 | 439 | 176 | . | . | . | . |
| PM1548-SK1P2 | CIITA | p.Arg580His | missense_variant | MODERATE | 0.526917 | 0 | 613 | 335 | true | . | . | . |
| PM1548-SK1P2 | RMI2 | p.Ile130Val | missense_variant | MODERATE | 0.105263 | 0 | 38 | 171 | true | . | . | . |
| PM1548-SK1P2 | TNFRSF17 | p.Pro145Gln | missense_variant | MODERATE | 0.36 | 0 | 25 | 38 | true | . | . | . |
| PM1548-SK1P2 | CDR2 | p.Gln130Arg | missense_variant | MODERATE | 0.392857 | 0 | 28 | 85 | . | . | . | . |
| PM1548-SK1P2 | USP31 | p.Gly1267Arg | missense_variant | MODERATE | 0 | 0 | 2 | 32 | . | . | . | . |
| PM1548-SK1P2 | COG7 | p.Pro67Ala | missense_variant | MODERATE | 0.297872 | 0 | 47 | 71 | . | . | . | . |
| PM1548-SK1P2 | GTF3C1 | p.Glu1761Gly | missense_variant | MODERATE | 0.117021 | 0 | 94 | 99 | . | . | . | . |
| PM1548-SK1P2 | ATXN2L | c.3140-64G>T | intron_variant | MODIFIER | 0.190476 | 0 | 21 | 124 | . | . | . | . |
| PM1548-SK1P2 | ATP2A1 | p.Gly229Ser | missense_variant | MODERATE | 0.03 | 0 | 100 | 168 | . | . | . | . |
| PM1548-SK1P2 | KIF22 | p.Ile592fs | frameshift_variant | HIGH | 0.454545 | 0 | 44 | 129 | . | . | . | . |
| PM1548-SK1P2 | PRR14 | p.Arg96Gln | missense_variant | MODERATE | 0.433333 | 0 | 30 | 271 | . | . | . | . |
| PM1548-SK1P2 | ZNF629 | p.Asp434Asn | missense_variant | MODERATE | 0.348214 | 0 | 112 | 34 | . | . | . | . |
| PM1548-SK1P2 | HSD3B7 | p.Val361Ile | missense_variant | MODERATE | 0.588889 | 0 | 90 | 165 | . | . | . | . |
| PM1548-SK1P2 | KAT8 | p.Val438Ala | missense_variant | MODERATE | 0.526316 | 0 | 152 | 67 | . | . | . | . |
| PM1548-SK1P2 | FUS | c.1295+1G>A | splice_donor_variant&intron_variant | HIGH | 0.454545 | 0 | 44 | 234 | true | . | . | . |
| PM1548-SK1P2 | PHKB | p.Tyr52His | missense_variant | MODERATE | 0 | 0 | 151 | 155 | . | . | . | . |
| PM1548-SK1P2 | LONP2 | p.Gly681Cys | missense_variant | MODERATE | 0 | 0 | 9 | 276 | . | . | . | . |
| PM1548-SK1P2 | N4BP1 | p.Arg640Cys | missense_variant | MODERATE | 0 | 0 | 2 | 94 | . | . | . | . |
| PM1548-SK1P2 | SALL1 | p.Ser1318Asn | missense_variant | MODERATE | 0.52 | 0 | 25 | 197 | . | . | . | . |
| PM1548-SK1P2 | SALL1 | p.His756Asp | missense_variant | MODERATE | 0.52381 | 0 | 21 | 53 | . | . | . | . |
| PM1548-SK1P2 | LPCAT2 | c.1216-7923dup | intron_variant | MODIFIER | 0.28 | 0 | 25 | 56 | . | . | . | . |
| PM1548-SK1P2 | SLC6A2 | p.Gly591Asp | missense_variant | MODERATE | 0.557252 | 0 | 131 | 342 | . | . | . | . |
| PM1548-SK1P2 | MMP15 | p.Ala603Asp | missense_variant | MODERATE | 0.170732 | 0.00512821 | 41 | 390 | . | . | . | . |
| PM1548-SK1P2 | CDH11 | p.Ala166Thr | missense_variant | MODERATE | 0.0285714 | 0 | 35 | 177 | true | . | . | . |
| PM1548-SK1P2 | CDH11 | p.Thr163Ile | missense_variant | MODERATE | 0.0285714 | 0 | 35 | 177 | true | . | . | . |
| PM1548-SK1P2 | CDH11 | p.His161His | synonymous_variant | LOW | 0.0294118 | 0 | 34 | 175 | true | . | . | . |
| PM1548-SK1P2 | CDH11 | p.Val149Asp | missense_variant | MODERATE | 0.166667 | 0 | 36 | 62 | true | . | . | . |
| PM1548-SK1P2 | CDH11 | p.Val147Asp | missense_variant | MODERATE | 0.205882 | 0 | 34 | 55 | true | . | . | . |
| PM1548-SK1P2 | CDH11 | p.Glu140Gly | missense_variant | MODERATE | 0.2 | 0 | 20 | 80 | true | . | . | . |
| PM1548-SK1P2 | CDH11 | p.Arg137Gln | missense_variant | MODERATE | 0.263158 | 0 | 19 | 79 | true | . | . | . |
| PM1548-SK1P2 | TK2 | p.Asn114Asn | synonymous_variant | LOW | 1 | 0 | 9 | 102 | . | . | . | . |
| PM1548-SK1P2 | B3GNT9 | p.Asp221Asn | missense_variant | MODERATE | 0.235294 | 0 | 17 | 208 | . | . | . | . |
| PM1548-SK1P2 | LRRC29 | p.Gln205His | missense_variant | MODERATE | 0.313043 | 0 | 115 | 215 | . | . | . | . |
| PM1548-SK1P2 | KCTD19 | p.Gly313Arg | missense_variant | MODERATE | 0.428962 | 0 | 366 | 109 | . | . | . | . |
| PM1548-SK1P2 | TPPP3 | p.Ser49Ala | missense_variant | MODERATE | 0.04 | 0 | 25 | 59 | . | . | . | . |
| PM1548-SK1P2 | CTCF | p.Arg283His | missense_variant | MODERATE | 0.125 | 0 | 16 | 196 | true | true | true | . |
| PM1548-SK1P2 | CDH3 | p.Asp264Glu | missense_variant | MODERATE | 0 | 0 | 42 | 403 | . | . | . | . |
| PM1548-SK1P2 | WWP2 | p.Arg633Gln | missense_variant | MODERATE | 0 | 0 | 3 | 88 | . | . | . | . |
| PM1548-SK1P2 | PMFBP1 | p.Glu205fs | frameshift_variant | HIGH | 0.222222 | 0 | 18 | 212 | . | . | . | . |
| PM1548-SK1P2 | ZFHX3 | p.Ala1099Ser | missense_variant | MODERATE | 0.135135 | 0 | 37 | 40 | true | . | true | true |
| PM1548-SK1P2 | RFWD3 | p.Val155Ala | missense_variant | MODERATE | 0.0220994 | 0 | 181 | 69 | . | . | . | . |
| PM1548-SK1P2 | PKD1L2 | n.81164209C>T | intragenic_variant | MODIFIER | 0.693333 | 0 | 75 | 77 | . | . | . | . |
| PM1548-SK1P2 | TAF1C | p.Pro707fs | frameshift_variant | HIGH | 0.564417 | 0 | 326 | 204 | . | . | . | . |
| PM1548-SK1P2 | PIEZO1 | p.Pro1061Leu | missense_variant | MODERATE | 0.469799 | 0 | 447 | 171 | . | . | . | . |
| PM1548-SK1P2 | PIEZO1 | p.Met1002Ile | missense_variant | MODERATE | 0.520408 | 0 | 98 | 33 | . | . | . | . |
| PM1548-SK1P2 | GALNS | p.Thr109Thr | synonymous_variant | LOW | 0.0169492 | 0 | 59 | 187 | . | . | . | . |
| PM1548-SK1P2 | ANKRD11 | p.Asp2130Asn | missense_variant | MODERATE | 0 | 0 | 16 | 42 | . | . | . | . |
| PM1548-SK1P2 | GEMIN4 | p.Arg229Gln | missense_variant | MODERATE | 0.4 | 0 | 5 | 149 | . | . | . | . |
| PM1548-SK1P2 | RNMTL1 | p.Met143fs | frameshift_variant | HIGH | 0.164384 | 0 | 73 | 36 | . | . | . | . |
| PM1548-SK1P2 | PRPF8 | p.Arg1057Gln | missense_variant | MODERATE | 0.504098 | 0 | 244 | 146 | . | . | . | . |
| PM1548-SK1P2 | HIC1 | p.Pro693Leu | missense_variant | MODERATE | 0.00421941 | 0.0126582 | 237 | 79 | . | . | . | . |
| PM1548-SK1P2 | SMG6 | p.Arg603His | missense_variant | MODERATE | 0.154545 | 0.00434783 | 110 | 230 | . | . | . | . |
| PM1548-SK1P2 | CTNS | p.Asp74Asn | missense_variant | MODERATE | 0.203704 | 0.00314465 | 54 | 318 | . | . | . | . |
| PM1548-SK1P2 | ITGAE | p.Arg113Gln | missense_variant | MODERATE | 0.444444 | 0 | 90 | 94 | . | . | . | . |
| PM1548-SK1P2 | ZZEF1 | p.Arg194Trp | missense_variant | MODERATE | 0.5 | 0 | 2 | 95 | . | . | . | . |
| PM1548-SK1P2 | KIF1C | p.Ser511Phe | missense_variant | MODERATE | 0.875 | 0 | 8 | 140 | . | . | . | . |
| PM1548-SK1P2 | USP6 | p.Pro181Leu | missense_variant&splice_region_variant | MODERATE | 0 | 0 | 6 | 73 | true | . | . | . |
| PM1548-SK1P2 | DVL2 | p.Arg442Cys | missense_variant | MODERATE | 0.531599 | 0 | 269 | 34 | . | . | . | . |
| PM1548-SK1P2 | EIF5A | p.Thr168Met | missense_variant | MODERATE | 0.580645 | 0 | 62 | 86 | . | . | true | . |
| PM1548-SK1P2 | DNAH2 | p.Gly4241fs | frameshift_variant | HIGH | 0.487805 | 0 | 41 | 94 | . | . | . | . |
| PM1548-SK1P2 | ALOX12B | p.Ala355Thr | missense_variant | MODERATE | 0.354633 | 0 | 313 | 295 | . | . | . | . |
| PM1548-SK1P2 | GLP2R | p.Ala515Thr | missense_variant | MODERATE | 0.566667 | 0.00334448 | 30 | 299 | . | . | . | . |
| PM1548-SK1P2 | MYH13 | p.Glu984Lys | missense_variant | MODERATE | 0.390625 | 0 | 64 | 72 | . | . | . | . |
| PM1548-SK1P2 | MYH8 | p.Gly186Arg | missense_variant | MODERATE | 0.315789 | 0 | 19 | 81 | . | . | . | . |
| PM1548-SK1P2 | MYH4 | p.Tyr424Cys | missense_variant | MODERATE | 0 | 0 | 85 | 298 | . | . | . | . |
| PM1548-SK1P2 | DNAH9 | p.Arg1542Met | missense_variant | MODERATE | 0.108108 | 0 | 37 | 342 | . | . | . | . |
| PM1548-SK1P2 | DNAH9 | p.Ala2514Thr | missense_variant | MODERATE | 0.24183 | 0 | 153 | 99 | . | . | . | . |
| PM1548-SK1P2 | ARHGAP44 | p.Ala495Thr | missense_variant | MODERATE | 0.25 | 0 | 4 | 53 | . | . | . | . |
| PM1548-SK1P2 | FLCN | p.His429fs | frameshift_variant | HIGH | 0.113527 | 0 | 414 | 269 | true | . | . | . |
| PM1548-SK1P2 | FLCN | p.Thr174Ala | missense_variant | MODERATE | 0.016129 | 0 | 310 | 227 | true | . | . | . |
| PM1548-SK1P2 | TOM1L2 | p.Met482fs | frameshift_variant | HIGH | 0.402985 | 0 | 67 | 83 | . | . | . | . |
| PM1548-SK1P2 | KIAA0100 | p.Arg689* | stop_gained | HIGH | 0.0252101 | 0 | 119 | 219 | . | . | . | . |
| PM1548-SK1P2 | SUPT6H | p.Ile1356Leu | missense_variant | MODERATE | 0 | 0.00465116 | 16 | 215 | . | . | . | . |
| PM1548-SK1P2 | PHF12 | p.Leu393fs | frameshift_variant | HIGH | 0.369231 | 0 | 130 | 61 | . | . | . | . |
| PM1548-SK1P2 | SEZ6 | p.Gly660Asp | missense_variant | MODERATE | 0.666667 | 0 | 18 | 83 | . | . | . | . |
| PM1548-SK1P2 | ANKRD13B | p.Arg233Gln | missense_variant | MODERATE | 0.0175439 | 0 | 57 | 98 | . | . | . | . |
| PM1548-SK1P2 | FNDC8 | p.Asn57Ser | missense_variant | MODERATE | 0.666667 | 0 | 6 | 150 | . | . | . | . |
| PM1548-SK1P2 | NLE1 | p.Val10Ala | missense_variant | MODERATE | 0 | 0 | 156 | 97 | . | . | . | . |
| PM1548-SK1P2 | SRCIN1 | p.Leu460Phe | missense_variant | MODERATE | 0 | 0 | 2 | 53 | . | . | . | . |
| PM1548-SK1P2 | PCGF2 | p.Asp157Asn | missense_variant | MODERATE | 0.545455 | 0 | 11 | 53 | . | . | . | . |
| PM1548-SK1P2 | STAC2 | p.Cys141Cys | synonymous_variant | LOW | 0 | 0 | 17 | 250 | . | . | . | . |
| PM1548-SK1P2 | GRB7 | p.Arg552Gln | missense_variant | MODERATE | 0 | 0 | 1083 | 159 | . | . | . | . |
| PM1548-SK1P2 | NR1D1 | p.Pro593Pro | synonymous_variant | LOW | 0.609005 | 0.00961538 | 422 | 104 | . | . | . | . |
| PM1548-SK1P2 | MSL1 | p.Leu372Ile | missense_variant | MODERATE | 0 | 0 | 8 | 43 | . | . | . | . |
| PM1548-SK1P2 | WIPF2 | p.Ala39Thr | missense_variant | MODERATE | 0.443038 | 0 | 79 | 66 | . | . | . | . |
| PM1548-SK1P2 | KRT25 | p.Ala91Thr | missense_variant | MODERATE | 0.321429 | 0 | 28 | 148 | . | . | . | . |
| PM1548-SK1P2 | KRT15 | p.Arg398Trp | missense_variant | MODERATE | 0.633333 | 0 | 30 | 53 | . | . | . | . |
| PM1548-SK1P2 | KRT16 | p.Gly243Ser | missense_variant | MODERATE | 0.454545 | 0 | 22 | 130 | . | . | . | . |
| PM1548-SK1P2 | DHX58 | p.Arg224Cys | missense_variant | MODERATE | 0.227273 | 0 | 22 | 91 | . | . | . | . |
| PM1548-SK1P2 | PLEKHH3 | p.Pro318Thr | missense_variant | MODERATE | 0.125 | 0 | 8 | 121 | . | . | . | . |
| PM1548-SK1P2 | ETV4 | p.Gln13Arg | missense_variant | MODERATE | 0.00998336 | 0 | 601 | 215 | true | . | . | . |
| PM1548-SK1P2 | MPP3 | p.Arg265Gln | missense_variant | MODERATE | 0.52 | 0 | 75 | 78 | . | . | . | . |

| | | | | | | | | | | | | |
|---|---|---|---|---|---|---|---|---|---|---|---|---|
| PM1548-SK1P2 | ATXN7L3 | p.Ile246Phe | missense_variant | MODERATE | 0.4375 | 0 | 48 | 37 | . | . | . | . |
| PM1548-SK1P2 | ITGA2B | p.Asn330Ser | missense_variant | MODERATE | 0.5 | 0 | 10 | 60 | . | . | . | . |
| PM1548-SK1P2 | ARHGAP27 | p.Leu247fs | frameshift_variant | HIGH | 0.510526 | 0 | 190 | 82 | . | . | . | . |
| PM1548-SK1P2 | WNT3 | p.Ala223Ser | missense_variant | MODERATE | 0 | 0 | 21 | 125 | . | . | . | . |
| PM1548-SK1P2 | KPNB1 | p.Ile686Leu | missense_variant | MODERATE | 0.105263 | 0 | 38 | 143 | . | . | . | . |
| PM1548-SK1P2 | KPNB1 | p.Ile686Met | missense_variant | MODERATE | 0.105263 | 0 | 38 | 143 | . | . | . | . |
| PM1548-SK1P2 | SP6 | p.Ala138Thr | missense_variant | MODERATE | 0 | 0 | 573 | 221 | . | . | . | . |
| PM1548-SK1P2 | PPP1R9B | p.Glu24Glu | synonymous_variant | LOW | 0.00802139 | 0 | 748 | 79 | . | . | . | . |
| PM1548-SK1P2 | CUEDC1 | p.Arg71Cys | missense_variant | MODERATE | 0.366197 | 0 | 71 | 107 | . | . | . | . |
| PM1548-SK1P2 | LPO | p.Arg82Cys | missense_variant | MODERATE | 0.290323 | 0 | 31 | 404 | . | . | . | . |
| PM1548-SK1P2 | RNF43 | p.Gly659fs | frameshift_variant | HIGH | 0.484375 | 0 | 256 | 76 | true | . | . | true |
| PM1548-SK1P2 | TBX2 | p.Leu460fs | frameshift_variant | HIGH | 0.582938 | 0 | 211 | 333 | . | . | . | . |
| PM1548-SK1P2 | MRC2 | p.Arg490Cys | missense_variant | MODERATE | 0 | 0 | 31 | 51 | . | . | . | . |
| PM1548-SK1P2 | CCDC47 | p.Ser303Pro | missense_variant | MODERATE | 0.117647 | 0 | 34 | 131 | . | . | . | . |
| PM1548-SK1P2 | FTSJ3 | p.Met493Ile | missense_variant | MODERATE | 0.666667 | 0 | 3 | 61 | . | . | . | . |
| PM1548-SK1P2 | ABCA10 | p.Tyr1399His | missense_variant | MODERATE | 0.36 | 0 | 25 | 336 | . | . | . | . |
| PM1548-SK1P2 | SLC16A5 | p.Gly93Cys | missense_variant | MODERATE | 0 | 0 | 9 | 58 | . | . | . | . |
| PM1548-SK1P2 | ITGB4 | p.Pro305Leu | missense_variant | MODERATE | 0.4 | 0 | 15 | 78 | . | . | . | . |
| PM1548-SK1P2 | ITGB4 | p.Pro723Ser | missense_variant | MODERATE | 0 | 0 | 579 | 167 | . | . | . | . |
| PM1548-SK1P2 | H3F3B | p.Ala8Asp | missense_variant | MODERATE | 0 | 0 | 5 | 88 | true | . | . | . |
| PM1548-SK1P2 | EVPL | p.Arg266Trp | missense_variant | MODERATE | 0 | 0 | 106 | 157 | . | . | . | . |
| PM1548-SK1P2 | RHBDF2 | p.Ala354Thr | missense_variant | MODERATE | 0 | 0 | 75 | 138 | . | . | . | . |
| PM1548-SK1P2 | TNRC6C | p.Arg1160His | missense_variant | MODERATE | 0.444444 | 0 | 45 | 184 | . | . | . | . |
| PM1548-SK1P2 | TMC6 | p.His31His | synonymous_variant | LOW | 0.481967 | 0 | 305 | 491 | . | . | . | . |
| PM1548-SK1P2 | GAA | p.Gly828Asp | missense_variant&splice_region_variant | MODERATE | 0 | 0 | 65 | 76 | . | . | . | . |
| PM1548-SK1P2 | ENDOV | p.Pro259Leu | missense_variant | MODERATE | 0.35 | 0 | 80 | 82 | . | . | . | . |
| PM1548-SK1P2 | RPTOR | p.Arg899* | stop_gained | HIGH | 0.166667 | 0 | 12 | 72 | . | . | . | . |
| PM1548-SK1P2 | SIRT7 | p.Thr181Met | missense_variant | MODERATE | 0.25 | 0 | 44 | 217 | . | . | . | . |
| PM1548-SK1P2 | OGFOD3 | p.Ser127Ser | splice_region_variant&synonymous_variant | LOW | 0.5 | 0 | 26 | 110 | . | . | . | . |
| PM1548-SK1P2 | TXNDC2 | p.Ile284Leu | missense_variant | MODERATE | 0.108108 | 0 | 37 | 273 | . | . | . | . |
| PM1548-SK1P2 | TXNDC2 | p.Ser285Pro | missense_variant | MODERATE | 0.111111 | 0 | 36 | 274 | . | . | . | . |
| PM1548-SK1P2 | TXNDC2 | p.Pro288Leu | missense_variant | MODERATE | 0.121212 | 0.00411523 | 33 | 243 | . | . | . | . |
| PM1548-SK1P2 | ANKRD62 | p.Gln745His | missense_variant | MODERATE | 0.75 | 0 | 8 | 165 | . | . | . | . |
| PM1548-SK1P2 | ZNF521 | p.Cys754Tyr | missense_variant | MODERATE | 0.823529 | 0 | 17 | 57 | true | . | . | . |
| PM1548-SK1P2 | KCTD1 | p.Asn124His | missense_variant | MODERATE | 0.09375 | 0 | 224 | 69 | . | . | . | . |
| PM1548-SK1P2 | DSG4 | p.Thr1001Met | missense_variant | MODERATE | 0.976744 | 0 | 43 | 104 | . | . | . | . |
| PM1548-SK1P2 | CCDC178 | p.Arg827Met | missense_variant | MODERATE | 0 | 0 | 75 | 192 | . | . | . | . |
| PM1548-SK1P2 | INO80C | p.Gly216Ser | missense_variant | MODERATE | 0 | 0 | 93 | 89 | . | . | . | . |
| PM1548-SK1P2 | SETBP1 | p.Glu244Asp | missense_variant | MODERATE | 0.166667 | 0 | 30 | 133 | true | true | . | . |
| PM1548-SK1P2 | SETBP1 | p.Ile251Val | missense_variant | MODERATE | 0.157895 | 0 | 38 | 199 | true | true | . | . |
| PM1548-SK1P2 | SETBP1 | p.Ala733Thr | missense_variant | MODERATE | 0 | 0 | 20 | 315 | true | true | . | . |
| PM1548-SK1P2 | SIGLEC15 | p.Ala96Thr | missense_variant | MODERATE | 0.193548 | 0 | 62 | 175 | . | . | . | . |
| PM1548-SK1P2 | SIGLEC15 | p.Arg99Pro | missense_variant | MODERATE | 0.193548 | 0 | 62 | 175 | . | . | . | . |
| PM1548-SK1P2 | SKOR2 | p.Asp736Glu | missense_variant | MODERATE | 0.0327869 | 0 | 183 | 157 | . | . | . | . |
| PM1548-SK1P2 | SKOR2 | p.Gly720Glu | missense_variant | MODERATE | 0.0357143 | 0 | 168 | 134 | . | . | . | . |
| PM1548-SK1P2 | SMAD7 | p.Pro55Ala | missense_variant | MODERATE | 0 | 0 | 8 | 32 | . | . | . | . |
| PM1548-SK1P2 | DSEL | p.Arg1009His | missense_variant | MODERATE | 1 | 0 | 7 | 67 | . | . | . | . |
| PM1548-SK1P2 | FAM69C | p.Gly48Cys | missense_variant | MODERATE | 0.125 | 0 | 32 | 170 | . | . | . | . |
| PM1548-SK1P2 | TSHZ1 | p.Ala570Val | missense_variant | MODERATE | 0 | 0 | 138 | 279 | . | . | . | . |
| PM1548-SK1P2 | TSHZ1 | p.Tyr775Cys | missense_variant | MODERATE | 1 | 0.00806452 | 10 | 124 | . | . | . | . |
| PM1548-SK1P2 | ZNF516 | p.Gly232del | disruptive_inframe_deletion | MODERATE | 0.0810811 | 0 | 37 | 279 | . | . | . | . |
| PM1548-SK1P2 | ZNF516 | p.Pro229_Gly23( | inframe_insertion | MODERATE | 0.0882353 | 0 | 34 | 280 | . | . | . | . |
| PM1548-SK1P2 | GALR1 | p.Leu46Met | missense_variant | MODERATE | 0.0454545 | 0 | 242 | 102 | . | . | . | . |
| PM1548-SK1P2 | GALR1 | p.Lys62Lys | synonymous_variant | LOW | 0.05 | 0 | 380 | 218 | . | . | . | . |
| PM1548-SK1P2 | SALL3 | p.Pro1105fs | frameshift_variant | HIGH | 1 | 0 | 1 | 200 | . | . | . | . |
| PM1548-SK1P2 | PPAP2C | n.291320T>C | intragenic_variant | MODIFIER | 0.338235 | 0 | 68 | 59 | . | . | . | . |
| PM1548-SK1P2 | LPPR3 | p.Gly646Ser | missense_variant | MODERATE | 0.555556 | 0 | 45 | 117 | . | . | . | . |
| PM1548-SK1P2 | CFD | p.Arg199Cys | missense_variant | MODERATE | 0.483965 | 0 | 343 | 193 | . | . | . | . |
| PM1548-SK1P2 | ARID3A | p.Arg503Cys | missense_variant | MODERATE | 0.5 | 0 | 8 | 124 | . | . | . | . |
| PM1548-SK1P2 | APC2 | p.Asp501Asn | missense_variant | MODERATE | 0.498069 | 0 | 1295 | 282 | . | . | . | . |
| PM1548-SK1P2 | PCSK4 | p.Gly57Asp | missense_variant | MODERATE | 0.0152355 | 0 | 1444 | 340 | . | . | . | . |
| PM1548-SK1P2 | ADAMTSL5 | p.Ala343fs | frameshift_variant | HIGH | 0.292517 | 0 | 735 | 366 | . | . | . | . |
| PM1548-SK1P2 | ADAMTSL5 | p.Pro341Ala | missense_variant | MODERATE | 0.336384 | 0 | 437 | 312 | . | . | . | . |
| PM1548-SK1P2 | MEX3D | p.Gly262Asp | missense_variant | MODERATE | 0.8 | 0 | 10 | 217 | . | . | . | . |
| PM1548-SK1P2 | TCF3 | p.Gly247fs | frameshift_variant | HIGH | 0 | 0 | 101 | 71 | true | . | . | . |
| PM1548-SK1P2 | C19orf35 | p.Arg325Cys | missense_variant | MODERATE | 0 | 0 | 77 | 53 | . | . | . | . |
| PM1548-SK1P2 | NCLN | p.Thr70Met | missense_variant | MODERATE | 0.552632 | 0 | 646 | 140 | . | . | . | . |
| PM1548-SK1P2 | DOHH | p.Arg286Trp | missense_variant | MODERATE | 0.00294985 | 0.0136054 | 339 | 147 | . | . | . | . |
| PM1548-SK1P2 | EEF2 | p.Pro738Ser | missense_variant | MODERATE | 0.489362 | 0 | 235 | 230 | . | . | . | . |
| PM1548-SK1P2 | PIAS4 | p.Tyr265Tyr | synonymous_variant | LOW | 0.522353 | 0 | 850 | 240 | . | . | . | . |
| PM1548-SK1P2 | MAP2K2 | p.Lys51Arg | missense_variant | MODERATE | 0.0714286 | 0 | 28 | 75 | true | . | . | . |
| PM1548-SK1P2 | MAP2K2 | p.Lys51Gln | missense_variant | MODERATE | 0.0714286 | 0 | 28 | 75 | true | . | . | . |
| PM1548-SK1P2 | CREB3L3 | p.Val140Ala | missense_variant | MODERATE | 0 | 0 | 561 | 205 | . | . | . | . |
| PM1548-SK1P2 | UBXN6 | p.Thr337Met | missense_variant | MODERATE | 0.0208333 | 0 | 48 | 93 | . | . | . | . |
| PM1548-SK1P2 | PLIN4 | p.Val592Met | missense_variant | MODERATE | 0.302326 | 0 | 43 | 40 | . | . | . | . |
| PM1548-SK1P2 | PLIN5 | p.Thr456Ile | missense_variant | MODERATE | 0.557377 | 0 | 61 | 37 | . | . | . | . |
| PM1548-SK1P2 | AC005594.3 | n.381G>A | non_coding_exon_variant | MODIFIER | 0.461783 | 0.0037037 | 314 | 270 | . | . | . | . |
| PM1548-SK1P2 | DPP9 | p.Arg125Trp | missense_variant | MODERATE | 0.268116 | 0 | 414 | 40 | . | . | . | . |
| PM1548-SK1P2 | KDM4B | p.Arg618Cys | missense_variant | MODERATE | 0 | 0 | 400 | 167 | . | . | . | . |
| PM1548-SK1P2 | KDM4B | p.Glu1013Lys | missense_variant | MODERATE | 0 | 0 | 1393 | 113 | . | . | . | . |
| PM1548-SK1P2 | DENND1C | p.Thr165fs | frameshift_variant | HIGH | 0.206897 | 0 | 29 | 34 | . | . | . | . |
| PM1548-SK1P2 | PEX11G | p.Arg124Trp | missense_variant | MODERATE | 0.479757 | 0 | 988 | 176 | . | . | . | . |
| PM1548-SK1P2 | CTD-3193O13.9 | p.Thr838Thr | synonymous_variant | LOW | 0.35 | 0 | 40 | 174 | . | . | . | . |
| PM1548-SK1P2 | COL5A3 | p.Glu1677Lys | missense_variant | MODERATE | 0.5 | 0 | 8 | 264 | . | . | true | true |
| PM1548-SK1P2 | ZNF653 | p.Asp247Gly | missense_variant | MODERATE | 0.475 | 0 | 40 | 166 | . | . | . | . |
| PM1548-SK1P2 | ELOF1 | p.Arg41Cys | missense_variant | MODERATE | 0.462783 | 0 | 309 | 108 | . | . | . | . |
| PM1548-SK1P2 | MRI1 | p.His330fs | frameshift_variant | HIGH | 0.347826 | 0 | 23 | 82 | . | . | . | . |
| PM1548-SK1P2 | RFX1 | p.Val600Ile | missense_variant | MODERATE | 0.463453 | 0 | 643 | 345 | . | . | . | . |
| PM1548-SK1P2 | PRKACA | p.Ala219Val | missense_variant | MODERATE | 0.947368 | 0 | 19 | 150 | true | . | . | . |
| PM1548-SK1P2 | TECR | p.Val85Met | missense_variant | MODERATE | 0.408759 | 0.00641026 | 137 | 156 | . | . | . | . |
| PM1548-SK1P2 | EMR2 | p.Pro294Thr | missense_variant | MODERATE | 0.102362 | 0 | 127 | 117 | . | . | . | . |
| PM1548-SK1P2 | EMR2 | p.Tyr292Ser | missense_variant | MODERATE | 0.117117 | 0 | 111 | 114 | . | . | . | . |
| PM1548-SK1P2 | KLF2 | p.Ala238fs | frameshift_variant | HIGH | 0.445498 | 0 | 422 | 137 | . | . | . | . |
| PM1548-SK1P2 | NWD1 | p.Arg788Cys | missense_variant | MODERATE | 0.603774 | 0 | 106 | 334 | . | . | . | . |
| PM1548-SK1P2 | BST2 | p.Ile120fs | frameshift_variant | HIGH | 0.1 | 0 | 10 | 64 | . | . | . | . |
| PM1548-SK1P2 | FCHO1 | p.Ala286Val | missense_variant&splice_region_variant | MODERATE | 0.48913 | 0 | 184 | 272 | . | . | . | . |
| PM1548-SK1P2 | DDX49 | p.Arg310Gln | missense_variant&splice_region_variant | MODERATE | 0.454798 | 0 | 719 | 300 | . | . | . | . |
| PM1548-SK1P2 | SUGP2 | p.Cys490Tyr | missense_variant | MODERATE | 0.818182 | 0 | 11 | 81 | . | . | . | . |
| PM1548-SK1P2 | NR2C2AP | p.Lys28fs | frameshift_variant | HIGH | 0.44856 | 0 | 243 | 183 | . | . | . | . |
| PM1548-SK1P2 | NCAN | p.Arg256Gln | missense_variant | MODERATE | 0.46988 | 0 | 83 | 124 | . | . | . | . |
| PM1548-SK1P2 | NDUFA13 | p.Trp131Leu | missense_variant | MODERATE | 0.12 | 0 | 50 | 264 | . | . | . | . |
| PM1548-SK1P2 | ZNF728 | p.Pro380Ser | missense_variant | MODERATE | 0 | 0 | 15 | 32 | . | . | . | . |
| PM1548-SK1P2 | ZNF728 | p.Arg323Pro | missense_variant | MODERATE | 0 | 0 | 2 | 48 | . | . | . | . |
| PM1548-SK1P2 | ZNF728 | p.Arg323Trp | missense_variant | MODERATE | 0 | 0 | 2 | 48 | . | . | . | . |
| PM1548-SK1P2 | ZNF507 | p.Asp475Gly | missense_variant | MODERATE | 0.666667 | 0 | 3 | 42 | . | . | . | . |
| PM1548-SK1P2 | KIAA0355 | p.Val801Met | missense_variant | MODERATE | 0.100372 | 0 | 269 | 184 | . | . | . | . |
| PM1548-SK1P2 | GRAMD1A | p.Gly605Ser | missense_variant | MODERATE | 0.132353 | 0 | 68 | 68 | . | . | . | . |
| PM1548-SK1P2 | RBM42 | p.Pro117Arg | missense_variant | MODERATE | 0 | 0 | 90 | 129 | . | . | . | . |
| PM1548-SK1P2 | ARHGAP33 | p.Gln144fs | frameshift_variant | HIGH | 0.512492 | 0 | 1641 | 431 | . | . | . | . |
| PM1548-SK1P2 | ZNF850 | p.Ala575Thr | missense_variant | MODERATE | 0.2 | 0 | 30 | 89 | . | . | . | . |
| PM1548-SK1P2 | ZNF850 | p.Ser572Ala | missense_variant | MODERATE | 0.147059 | 0 | 34 | 133 | . | . | . | . |
| PM1548-SK1P2 | ZNF850 | p.Val554Ile | missense_variant | MODERATE | 0 | 0 | 25 | 86 | . | . | . | . |
| PM1548-SK1P2 | DPF1 | p.Cys58Cys | synonymous_variant | LOW | 0.013986 | 0 | 143 | 394 | . | . | . | . |
| PM1548-SK1P2 | DPF1 | p.Cys50Cys | synonymous_variant | LOW | 0.0141844 | 0 | 141 | 391 | . | . | . | . |
| PM1548-SK1P2 | YIF1B | p.Phe39Phe | synonymous_variant | LOW | 0.429907 | 0 | 107 | 480 | . | . | . | . |
| PM1548-SK1P2 | RYR1 | p.Val3218Met | missense_variant | MODERATE | 0 | 0 | 1158 | 114 | . | . | . | . |
| PM1548-SK1P2 | FCGBP | p.Ala5241Val | missense_variant | MODERATE | 0.486957 | 0 | 230 | 284 | . | . | . | . |
| PM1548-SK1P2 | PRX | p.Glu495Gln | missense_variant | MODERATE | 0.162162 | 0 | 37 | 130 | . | true | . | . |
| PM1548-SK1P2 | B3GNT8 | p.Glu333Ala | missense_variant | MODERATE | 0.37284 | 0 | 405 | 219 | . | . | . | . |
| PM1548-SK1P2 | ARHGEF1 | p.Arg137Trp | missense_variant | MODERATE | 0.487705 | 0 | 244 | 130 | . | . | . | . |
| PM1548-SK1P2 | ZNF227 | p.Gly390Ser | missense_variant | MODERATE | 0.75 | 0 | 8 | 73 | . | . | . | . |
| PM1548-SK1P2 | PVRL2 | p.Pro53Leu | missense_variant | MODERATE | 0.425234 | 0 | 214 | 119 | . | . | . | . |
| PM1548-SK1P2 | MARK4 | p.Thr651Thr | synonymous_variant | LOW | 0 | 0 | 66 | 352 | . | . | . | . |
| PM1548-SK1P2 | KLC3 | p.Arg320Cys | missense_variant | MODERATE | 0.551724 | 0 | 145 | 102 | . | . | . | . |
| PM1548-SK1P2 | PPP1R13L | p.Leu397Pro | missense_variant | MODERATE | 0.135593 | 0 | 59 | 477 | . | . | . | . |
| PM1548-SK1P2 | GIPR | p.Gly144Ser | missense_variant | MODERATE | 0.428571 | 0 | 7 | 35 | . | . | . | . |
| PM1548-SK1P2 | STRN4 | p.Ala484Val | missense_variant | MODERATE | 0.540984 | 0 | 122 | 141 | . | . | . | . |

| | | | | | | | | | | | | |
|---|---|---|---|---|---|---|---|---|---|---|---|---|
| PM1548-SK1P2 | FKRP | p.Ala193Thr | missense_variant | MODERATE | 0 | 0 | 785 | 98 | . | . | . | . |
| PM1548-SK1P2 | SLC1A5 | p.Ala534Thr | missense_variant | MODERATE | 0.363636 | 0 | 11 | 44 | . | . | . | . |
| PM1548-SK1P2 | ZC3H4 | p.Arg894His | missense_variant | MODERATE | 0 | 0 | 564 | 254 | . | . | . | . |
| PM1548-SK1P2 | ZC3H4 | p.Gly668Ser | missense_variant | MODERATE | 0.130435 | 0 | 23 | 225 | . | . | . | . |
| PM1548-SK1P2 | ZC3H4 | p.Lys500Arg | missense_variant | MODERATE | 0.125 | 0 | 32 | 128 | . | . | . | . |
| PM1548-SK1P2 | ZNF541 | p.Arg214His | missense_variant | MODERATE | 0 | 0 | 27 | 53 | . | . | . | . |
| PM1548-SK1P2 | GLTSCR1 | p.Ser1126fs | frameshift_variant | HIGH | 0.489218 | 0 | 742 | 146 | . | . | . | . |
| PM1548-SK1P2 | CRX | p.Ala35Thr | missense_variant&splice_region_variant | MODERATE | 0.47619 | 0 | 231 | 73 | . | . | . | . |
| PM1548-SK1P2 | PLEKHA4 | p.Arg225Trp | missense_variant | MODERATE | 0.564706 | 0 | 85 | 36 | . | . | . | . |
| PM1548-SK1P2 | BAX | p.Glu41fs | frameshift_variant | HIGH | 0.00306748 | 0 | 326 | 76 | . | . | . | . |
| PM1548-SK1P2 | FLT3LG | p.Ser118fs | frameshift_variant | HIGH | 0.512195 | 0 | 41 | 82 | . | . | . | . |
| PM1548-SK1P2 | SCAF1 | p.Gly651fs | frameshift_variant | HIGH | 0.493902 | 0 | 164 | 92 | . | . | . | . |
| PM1548-SK1P2 | ASPDH | p.Glu177Lys | missense_variant | MODERATE | 0.434783 | 0 | 23 | 226 | . | . | . | . |
| PM1548-SK1P2 | LRRC4B | p.Gly50fs | frameshift_variant | HIGH | 0.5 | 0 | 18 | 59 | . | . | . | . |
| PM1548-SK1P2 | CNOT3 | p.Tyr637His | missense_variant | MODERATE | 0.777778 | 0 | 9 | 41 | true | . | . | true |
| PM1548-SK1P2 | LILRB3 | c.355+1720T>A | intron_variant | MODIFIER | 0.3125 | 0 | 48 | 186 | . | . | . | . |
| PM1548-SK1P2 | LILRB3 | c.355+1720T>A | intron_variant | MODIFIER | 0.3125 | 0 | 48 | 187 | . | . | . | . |
| PM1548-SK1P2 | LILRB3 | c.355+1720T>A | intron_variant | MODIFIER | 0.3125 | 0 | 48 | 184 | . | . | . | . |
| PM1548-SK1P2 | LILRB3 | c.355+1720T>A | intron_variant | MODIFIER | 0.3125 | 0 | 48 | 184 | . | . | . | . |
| PM1548-SK1P2 | EPS8L1 | p.Arg608Cys | missense_variant | MODERATE | 0.33913 | 0 | 115 | 167 | . | . | . | . |
| PM1548-SK1P2 | SBK2 | p.Ala115Thr | missense_variant | MODERATE | 0.415162 | 0 | 277 | 186 | . | . | . | . |
| PM1548-SK1P2 | ZNF579 | p.Arg101Cys | missense_variant | MODERATE | 0.334177 | 0 | 395 | 216 | . | . | . | . |
| PM1548-SK1P2 | ZNF865 | p.Leu610Phe | missense_variant | MODERATE | 0.0344828 | 0 | 174 | 212 | . | . | . | . |
| PM1548-SK1P2 | ZNF865 | p.Leu632Leu | synonymous_variant | LOW | 0.037234 | 0 | 188 | 209 | . | . | . | . |
| PM1548-SK1P2 | ZNF71 | p.Gln343Gln | synonymous_variant | LOW | 0.540441 | 0 | 544 | 303 | . | . | . | . |
| PM1548-SK1P2 | TRIM28 | p.Arg492Cys | missense_variant | MODERATE | 0.410256 | 0 | 39 | 142 | . | . | . | . |
| PM1548-SK1P2 | PXDN | p.Val850Met | missense_variant | MODERATE | 0 | 0 | 1004 | 308 | . | . | . | . |
| PM1548-SK1P2 | ROCK2 | p.Glu1060Gly | missense_variant | MODERATE | 0.108108 | 0 | 37 | 93 | . | . | . | . |
| PM1548-SK1P2 | FAM228B | p.Asp82Gly | missense_variant | MODERATE | 0.333333 | 0 | 9 | 174 | . | . | . | . |
| PM1548-SK1P2 | NCOA1 | p.Asn1244Thr | missense_variant | MODERATE | 0 | 0 | 51 | 206 | true | . | . | . |
| PM1548-SK1P2 | NCOA1 | p.Ile1260Val | missense_variant | MODERATE | 0 | 0 | 61 | 204 | true | . | . | . |
| PM1548-SK1P2 | NCOA1 | p.Ala1274Thr | missense_variant | MODERATE | 0 | 0 | 23 | 87 | true | . | . | . |
| PM1548-SK1P2 | ADCY3 | p.Thr182Met | missense_variant | MODERATE | 0.62605 | 0 | 238 | 84 | . | . | . | . |
| PM1548-SK1P2 | TRIM54 | p.Arg97Gln | missense_variant | MODERATE | 0.419355 | 0 | 31 | 313 | . | . | . | . |
| PM1548-SK1P2 | TRIM54 | p.Glu131Asp | missense_variant | MODERATE | 0.0344828 | 0 | 29 | 142 | . | . | . | . |
| PM1548-SK1P2 | HEATR5B | p.Lys48fs | frameshift_variant | HIGH | 0 | 0 | 90 | 85 | . | . | . | . |
| PM1548-SK1P2 | RMDN2 | p.Arg130His | missense_variant | MODERATE | 0.038961 | 0 | 77 | 121 | . | . | . | . |
| PM1548-SK1P2 | SOS1 | p.Asn15Lys | missense_variant | MODERATE | 0.494366 | 0 | 710 | 93 | . | . | . | . |
| PM1548-SK1P2 | FSHR | p.Arg569Lys | missense_variant | MODERATE | 0.111675 | 0 | 197 | 177 | . | . | . | . |
| PM1548-SK1P2 | FSHR | p.Ser566Arg | missense_variant | MODERATE | 0.111111 | 0 | 198 | 133 | . | . | . | . |
| PM1548-SK1P2 | FSHR | p.Asn558Asn | synonymous_variant | LOW | 0.11 | 0 | 200 | 139 | . | . | . | . |
| PM1548-SK1P2 | FSHR | p.Ile550Thr | missense_variant | MODERATE | 0.121387 | 0 | 173 | 159 | . | . | . | . |
| PM1548-SK1P2 | CCDC88A | p.Pro1470Ser | missense_variant | MODERATE | 0.382353 | 0 | 102 | 211 | . | . | . | . |
| PM1548-SK1P2 | CCDC85A | p.His295Tyr | missense_variant | MODERATE | 0 | 0 | 6 | 103 | . | . | . | . |
| PM1548-SK1P2 | CCDC85A | p.Pro359Leu | missense_variant | MODERATE | 1 | 0 | 3 | 45 | . | . | . | . |
| PM1548-SK1P2 | FANCL | c.555+2T>C | splice_donor_variant&intron_variant | HIGH | 0.0625 | 0 | 16 | 47 | . | . | . | . |
| PM1548-SK1P2 | OTX1 | p.Arg39Trp | missense_variant | MODERATE | 0.464286 | 0 | 28 | 146 | . | . | . | . |
| PM1548-SK1P2 | AFTPH | p.Asp198fs | frameshift_variant | HIGH | 0.125 | 0 | 32 | 158 | . | . | . | . |
| PM1548-SK1P2 | NAGK | p.Lys90Gln | missense_variant | MODERATE | 0 | 0 | 82 | 344 | . | . | . | . |
| PM1548-SK1P2 | NAGK | p.Asn100Asp | missense_variant | MODERATE | 0 | 0 | 80 | 271 | . | . | . | . |
| PM1548-SK1P2 | FBXO41 | p.Arg594Gln | missense_variant | MODERATE | 0 | 0 | 10 | 69 | . | . | . | . |
| PM1548-SK1P2 | TET3 | p.Pro1195Leu | missense_variant | MODERATE | 0.384615 | 0 | 13 | 50 | . | . | . | . |
| PM1548-SK1P2 | C2orf81 | p.Ala344Val | missense_variant | MODERATE | 0.00606061 | 0 | 495 | 154 | . | . | . | . |
| PM1548-SK1P2 | TCF7L1 | p.His246Pro | missense_variant | MODERATE | 0.147059 | 0.0075188 | 68 | 133 | . | . | . | . |
| PM1548-SK1P2 | KDM3A | p.Glu1141* | stop_gained | HIGH | 0.228571 | 0 | 35 | 166 | . | . | . | . |
| PM1548-SK1P2 | ZNF514 | p.Gly396Glu | missense_variant | MODERATE | 0.322222 | 0 | 90 | 187 | . | . | . | . |
| PM1548-SK1P2 | NCAPH | p.Gln453* | stop_gained&splice_region_variant | HIGH | 0.727273 | 0 | 11 | 53 | . | . | . | . |
| PM1548-SK1P2 | ZAP70 | p.Ala387Val | missense_variant | MODERATE | 0.381356 | 0 | 118 | 293 | . | . | . | . |
| PM1548-SK1P2 | LONRF2 | p.Asp406Gly | missense_variant | MODERATE | 0 | 0 | 7 | 174 | . | . | . | . |
| PM1548-SK1P2 | CHST10 | p.Asn138Asn | synonymous_variant | LOW | 0.475884 | 0 | 311 | 237 | . | . | . | . |
| PM1548-SK1P2 | IL18R1 | p.Pro527fs | frameshift_variant | HIGH | 0.408867 | 0 | 203 | 255 | . | . | . | . |
| PM1548-SK1P2 | FBLN7 | c.670+86C>T | intron_variant | MODIFIER | 0.555147 | 0 | 272 | 109 | . | . | . | . |
| PM1548-SK1P2 | FBLN7 | p.Arg381Cys | missense_variant | MODERATE | 0.378378 | 0 | 37 | 87 | . | . | . | . |
| PM1548-SK1P2 | ZC3H6 | p.Arg99fs | frameshift_variant | HIGH | 0.583333 | 0 | 12 | 90 | . | . | . | . |
| PM1548-SK1P2 | POLR1B | p.Cys893Tyr | missense_variant | MODERATE | 0.0357143 | 0 | 28 | 127 | . | . | . | . |
| PM1548-SK1P2 | IL36B | c.261+1014delT | intron_variant | MODIFIER | 0.333333 | 0 | 6 | 101 | . | . | . | . |
| PM1548-SK1P2 | TMEM185B | p.Pro341fs | frameshift_variant | HIGH | 0.418605 | 0 | 43 | 231 | . | . | . | . |
| PM1548-SK1P2 | CLASP1 | p.Ala1225Val | missense_variant | MODERATE | 0.545455 | 0 | 33 | 85 | . | . | . | . |
| PM1548-SK1P2 | CYP27C1 | p.Ala103Thr | missense_variant | MODERATE | 0.6 | 0 | 15 | 179 | . | . | . | . |
| PM1548-SK1P2 | ACVR2A | p.Lys437fs | frameshift_variant | HIGH | 1 | 0 | 31 | 205 | . | true | . | . |
| PM1548-SK1P2 | NEB | p.Phe2843Tyr | missense_variant | MODERATE | 0.0980392 | 0 | 51 | 178 | . | . | . | . |
| PM1548-SK1P2 | TBR1 | p.Gly599Ser | missense_variant | MODERATE | 0.982609 | 0 | 115 | 40 | . | . | . | . |
| PM1548-SK1P2 | TLK1 | p.Gln18Arg | missense_variant | MODERATE | 0.0898876 | 0 | 89 | 529 | . | . | . | . |
| PM1548-SK1P2 | ITGA6 | p.Val266Leu | missense_variant | MODERATE | 0 | 0 | 28 | 156 | . | . | . | . |
| PM1548-SK1P2 | SP3 | p.Val461Ile | missense_variant | MODERATE | 0.153846 | 0 | 117 | 261 | . | . | . | . |
| PM1548-SK1P2 | EVX2 | p.Gly438Ser | missense_variant | MODERATE | 1 | 0 | 293 | 165 | . | . | . | . |
| PM1548-SK1P2 | EVX2 | p.Asp320Glu | missense_variant | MODERATE | 0 | 0 | 3 | 48 | . | . | . | . |
| PM1548-SK1P2 | CCDC141 | p.Ala1030Val | missense_variant | MODERATE | 0.176471 | 0 | 34 | 69 | . | . | . | . |
| PM1548-SK1P2 | PDE1A | p.Ala371Val | missense_variant | MODERATE | 0.904762 | 0 | 21 | 75 | . | . | . | . |
| PM1548-SK1P2 | FSIP2 | p.Arg829Cys | missense_variant | MODERATE | 0.25 | 0 | 12 | 51 | . | . | . | . |
| PM1548-SK1P2 | DNAH7 | p.Ala3252Glu | missense_variant | MODERATE | 0.102041 | 0 | 49 | 64 | . | . | . | . |
| PM1548-SK1P2 | GTF3C3 | p.Pro248Leu | missense_variant | MODERATE | 0 | 0 | 11 | 201 | . | . | . | . |
| PM1548-SK1P2 | AOX1 | p.Phe1162Ser | missense_variant | MODERATE | 0.323077 | 0 | 65 | 36 | . | . | . | . |
| PM1548-SK1P2 | PIKFYVE | p.Leu419Leu | synonymous_variant | LOW | 0.0909091 | 0 | 22 | 195 | . | . | . | . |
| PM1548-SK1P2 | SMARCAL1 | p.Arg499Gln | missense_variant | MODERATE | 0.630282 | 0 | 284 | 137 | . | . | . | . |
| PM1548-SK1P2 | AAMP | p.Glu63Asp | missense_variant | MODERATE | 0 | 0.00606061 | 58 | 165 | . | . | . | . |
| PM1548-SK1P2 | ZNF142 | p.Ala335Pro | missense_variant | MODERATE | 0.176471 | 0 | 34 | 317 | . | . | . | . |
| PM1548-SK1P2 | CYP27A1 | p.Ser203Leu | missense_variant | MODERATE | 0.272727 | 0 | 11 | 39 | . | . | . | . |
| PM1548-SK1P2 | WNT6 | p.Asp283Val | missense_variant | MODERATE | 0.471429 | 0 | 700 | 120 | . | . | . | . |
| PM1548-SK1P2 | FAM134A | p.Asp391Val | missense_variant | MODERATE | 0.515152 | 0 | 66 | 285 | . | . | . | . |
| PM1548-SK1P2 | DNPEP | p.Arg191Gln | missense_variant | MODERATE | 0 | 0 | 7 | 296 | . | . | . | . |
| PM1548-SK1P2 | CHPF | p.Arg721Trp | missense_variant | MODERATE | 0.456107 | 0 | 524 | 73 | . | . | . | . |
| PM1548-SK1P2 | WDFY1 | p.Ala75Asp | missense_variant | MODERATE | 0.115942 | 0 | 69 | 181 | . | . | . | . |
| PM1548-SK1P2 | B3GNT7 | p.Gly302Asp | missense_variant | MODERATE | 0.444444 | 0 | 9 | 60 | . | . | . | . |
| PM1548-SK1P2 | DGKD | p.Arg935Trp | missense_variant | MODERATE | 0 | 0 | 15 | 67 | . | . | . | . |
| PM1548-SK1P2 | ACKR3 | p.Val42Ala | missense_variant | MODERATE | 0.574468 | 0 | 47 | 137 | true | . | . | . |
| PM1548-SK1P2 | COL6A3 | p.Gly1714Glu | missense_variant | MODERATE | 0.5 | 0 | 10 | 50 | . | . | . | . |
| PM1548-SK1P2 | COL6A3 | p.Ile1609Asn | missense_variant | MODERATE | 0.4375 | 0 | 16 | 113 | . | . | . | . |
| PM1548-SK1P2 | ING5 | p.Arg106His | missense_variant | MODERATE | 0 | 0 | 102 | 105 | . | . | . | . |
| PM1548-SK1P2 | TBC1D20 | p.Pro71Ser | missense_variant | MODERATE | 0.117647 | 0 | 34 | 48 | . | . | . | . |
| PM1548-SK1P2 | VPS16 | p.Thr818Met | missense_variant | MODERATE | 0.425633 | 0 | 632 | 238 | . | . | . | . |
| PM1548-SK1P2 | ATRN | p.Ala122Thr | missense_variant | MODERATE | 0.0460829 | 0.00645161 | 217 | 155 | . | . | . | . |
| PM1548-SK1P2 | SLC23A2 | p.Ile412fs | frameshift_variant | HIGH | 0.5 | 0 | 6 | 150 | . | . | . | . |
| PM1548-SK1P2 | BMP2 | p.Ala10Val | missense_variant | MODERATE | 0.0565217 | 0 | 230 | 162 | . | . | . | . |
| PM1548-SK1P2 | PLCB1 | p.Arg768Gln | missense_variant | MODERATE | 0 | 0 | 5 | 91 | . | . | . | . |
| PM1548-SK1P2 | JAG1 | p.Gly537Ser | missense_variant | MODERATE | 0.133333 | 0 | 30 | 184 | . | . | . | . |
| PM1548-SK1P2 | CSRP2BP | p.Trp173* | stop_gained | HIGH | 0 | 0 | 4 | 168 | . | . | . | . |
| PM1548-SK1P2 | CRNKL1 | p.Phe352Leu | missense_variant | MODERATE | 0.666667 | 0 | 12 | 71 | . | . | . | . |
| PM1548-SK1P2 | PLAGL2 | p.Arg180His | missense_variant | MODERATE | 0.350575 | 0 | 174 | 145 | . | . | . | . |
| PM1548-SK1P2 | BPIFA2 | p.Arg221His | missense_variant | MODERATE | 0.636364 | 0 | 11 | 61 | . | . | . | . |
| PM1548-SK1P2 | NECAB3 | p.Arg278fs | frameshift_variant | HIGH | 0 | 0 | 2 | 39 | . | . | . | . |
| PM1548-SK1P2 | CEP250 | p.Lys787Glu | missense_variant | MODERATE | 0.111111 | 0 | 36 | 63 | . | . | . | . |
| PM1548-SK1P2 | SOGA1 | p.Pro974Ser | missense_variant&splice_region_variant | MODERATE | 0 | 0 | 47 | 259 | . | . | . | . |
| PM1548-SK1P2 | PLCG1 | p.Ile942Met | missense_variant | MODERATE | 0 | 0 | 202 | 140 | true | . | . | . |
| PM1548-SK1P2 | ZSWIM3 | p.Arg225Gln | missense_variant | MODERATE | 0.454545 | 0 | 33 | 55 | . | . | . | . |
| PM1548-SK1P2 | NFATC2 | p.Phe158Tyr | missense_variant | MODERATE | 0 | 0 | 118 | 59 | true | . | . | . |
| PM1548-SK1P2 | NFATC2 | p.Gly22fs | frameshift_variant | HIGH | 0.0197628 | 0 | 253 | 146 | true | . | . | . |
| PM1548-SK1P2 | SALL4 | p.Phe391Leu | missense_variant | MODERATE | 0.472973 | 0 | 74 | 33 | . | . | . | . |
| PM1548-SK1P2 | ZFP64 | n.50701484A>G | intragenic_variant | MODIFIER | 0.117834 | 0 | 314 | 142 | . | . | . | . |
| PM1548-SK1P2 | DOK5 | p.Val169Ala | missense_variant | MODERATE | 0.163636 | 0 | 110 | 338 | . | . | . | . |
| PM1548-SK1P2 | DOK5 | p.Ala267Asp | missense_variant | MODERATE | 0 | 0 | 304 | 146 | . | . | . | . |
| PM1548-SK1P2 | TFAP2C | p.Ala354Val | missense_variant | MODERATE | 0.484848 | 0 | 33 | 269 | . | . | . | . |
| PM1548-SK1P2 | PMEPA1 | p.Ser209fs | frameshift_variant | HIGH | 0 | 0 | 1241 | 249 | . | . | . | . |
| PM1548-SK1P2 | NELFCD | p.Val321Ile | missense_variant | MODERATE | 0.592593 | 0 | 27 | 74 | . | . | . | . |
| PM1548-SK1P2 | ZNF831 | p.Leu434Met | missense_variant | MODERATE | 0.395586 | 0 | 589 | 129 | . | . | . | . |
| PM1548-SK1P2 | HELZ2 | p.Ala639Thr | missense_variant | MODERATE | 0.48802 | 0 | 793 | 137 | . | . | . | . |
| PM1548-SK1P2 | RTEL1 | p.Arg729Cys | missense_variant | MODERATE | 0.540373 | 0 | 322 | 260 | . | . | . | . |

| | | | | | | | | | | | | |
|---|---|---|---|---|---|---|---|---|---|---|---|---|
| PM1548-SK1P2 | TNFRSF6B | p.Leu297Phe | missense_variant | MODERATE | 0 | 0 | 728 | 277 | . | . | . | . |
| PM1548-SK1P2 | SLC2A4RG | p.Arg82fs | frameshift_variant | HIGH | 0.44186 | 0 | 43 | 203 | . | . | . | . |
| PM1548-SK1P2 | PRPF6 | p.Arg317His | missense_variant | MODERATE | 0 | 0 | 32 | 86 | . | . | . | . |
| PM1548-SK1P2 | LTN1 | p.Gly1705Val | missense_variant | MODERATE | 0.2 | 0 | 15 | 36 | . | . | . | . |
| PM1548-SK1P2 | TIAM1 | p.His1010Tyr | missense_variant | MODERATE | 0.441176 | 0 | 68 | 479 | . | . | . | . |
| PM1548-SK1P2 | URB1 | p.Pro2036Leu | missense_variant | MODERATE | 0.25 | 0 | 4 | 14 | . | . | . | . |
| PM1548-SK1P2 | PAXBP1 | p.Pro161Thr | missense_variant | MODERATE | 0 | 0 | 5 | 40 | . | . | . | . |
| PM1548-SK1P2 | PAXBP1 | p.Ser105Arg | missense_variant | MODERATE | 0 | 0 | 449 | 158 | . | . | . | . |
| PM1548-SK1P2 | GART | p.Glu297Gln | missense_variant | MODERATE | 0 | 0 | 49 | 160 | . | . | . | . |
| PM1548-SK1P2 | SON | p.Thr790Ser | missense_variant | MODERATE | 0.0909091 | 0 | 11 | 177 | . | . | . | . |
| PM1548-SK1P2 | SON | c.6657+163T>C | intron_variant | MODIFIER | 0.35 | 0 | 60 | 33 | . | . | . | . |
| PM1548-SK1P2 | ITSN1 | p.Met768Thr | missense_variant | MODERATE | 0.5 | 0 | 6 | 46 | . | . | . | true |
| PM1548-SK1P2 | SLC5A3 | p.Arg252Gln | missense_variant | MODERATE | 0.686275 | 0 | 51 | 44 | . | . | . | . |
| PM1548-SK1P2 | UMODL1 | p.Gly1319Ser | missense_variant | MODERATE | 0.484536 | 0 | 97 | 131 | . | . | . | . |
| PM1548-SK1P2 | CRYAA | p.Ile61Val | missense_variant | MODERATE | 0.422917 | 0 | 480 | 344 | . | . | . | . |
| PM1548-SK1P2 | SLC19A1 | p.Asp488Gly | missense_variant | MODERATE | 0.409091 | 0 | 264 | 130 | . | . | . | . |
| PM1548-SK1P2 | FTCD | p.Ile187Thr | missense_variant | MODERATE | 0.296296 | 0 | 81 | 175 | . | . | . | . |
| PM1548-SK1P2 | LSS | p.Thr457Met | missense_variant | MODERATE | 0.477352 | 0 | 287 | 295 | . | . | . | . |
| PM1548-SK1P2 | CECR2 | p.Ser425Ser | synonymous_variant | LOW | 0.0571429 | 0 | 35 | 131 | . | . | . | . |
| PM1548-SK1P2 | CECR2 | p.Pro428Pro | synonymous_variant | LOW | 0.0606061 | 0 | 33 | 130 | . | . | . | . |
| PM1548-SK1P2 | CECR2 | p.Tyr431Tyr | synonymous_variant | LOW | 0.0606061 | 0 | 33 | 130 | . | . | . | . |
| PM1548-SK1P2 | ATP6V1E1 | p.Leu128Met | missense_variant | MODERATE | 0.275862 | 0 | 29 | 148 | . | . | . | . |
| PM1548-SK1P2 | CLTCL1 | p.Ala1567Thr | missense_variant | MODERATE | 0.556745 | 0 | 467 | 279 | true | . | . | . |
| PM1548-SK1P2 | SCARF2 | p.Arg335Cys | missense_variant | MODERATE | 0 | 0 | 6 | 100 | . | . | . | . |
| PM1548-SK1P2 | SLC7A4 | p.Ala256Thr | missense_variant | MODERATE | 0.234899 | 0 | 149 | 223 | . | . | . | . |
| PM1548-SK1P2 | IGLL1 | p.Arg60Gln | missense_variant | MODERATE | 0 | 0 | 17 | 81 | . | . | . | . |
| PM1548-SK1P2 | C22orf43 | p.Ser153Gly | missense_variant | MODERATE | 0 | 0 | 5 | 97 | . | . | . | . |
| PM1548-SK1P2 | ASPHD2 | p.Arg358Trp | missense_variant | MODERATE | 0.429688 | 0 | 128 | 411 | . | . | . | . |
| PM1548-SK1P2 | TFIP11 | p.Arg39* | stop_gained | HIGH | 0.466667 | 0 | 285 | 103 | . | . | . | . |
| PM1548-SK1P2 | RASL10A | p.Arg108Trp | missense_variant | MODERATE | 0.108108 | 0 | 37 | 108 | . | . | . | . |
| PM1548-SK1P2 | AP1B1 | p.Ala655Asp | missense_variant | MODERATE | 0 | 0 | 12 | 31 | . | . | . | . |
| PM1548-SK1P2 | LIMK2 | p.Asn477Ser | missense_variant | MODERATE | 0.0424671 | 0 | 989 | 257 | . | . | . | . |
| PM1548-SK1P2 | SFI1 | p.Thr710Ser | missense_variant | MODERATE | 0 | 0 | 8 | 104 | . | . | . | . |
| PM1548-SK1P2 | CSF2RB | p.Gln642fs | frameshift_variant | HIGH | 0.570159 | 0 | 1575 | 133 | . | . | . | . |
| PM1548-SK1P2 | C1QTNF6 | p.Met229Thr | missense_variant | MODERATE | 0.519757 | 0 | 329 | 41 | . | . | . | . |
| PM1548-SK1P2 | GRAP2 | p.Val129Ile | missense_variant | MODERATE | 0.410256 | 0 | 117 | 162 | . | . | . | . |
| PM1548-SK1P2 | MKL1 | p.Pro138Leu | missense_variant | MODERATE | 0.161871 | 0 | 278 | 100 | true | . | . | . |
| PM1548-SK1P2 | EP300 | p.Pro2107Thr | missense_variant | MODERATE | 0.153846 | 0 | 13 | 94 | true | true | true | . |
| PM1548-SK1P2 | ZC3H7B | p.Arg908Cys | missense_variant | MODERATE | 0.635514 | 0 | 107 | 73 | . | . | . | . |
| PM1548-SK1P2 | PARVB | c.211+24725C>A | intron_variant | MODIFIER | 0.491525 | 0 | 472 | 165 | . | . | . | . |
| PM1548-SK1P2 | CELSR1 | p.Val1411Gly | missense_variant | MODERATE | 0.296188 | 0 | 341 | 333 | . | . | . | . |
| PM1548-SK1P2 | CELSR1 | p.Glu1159Lys | missense_variant | MODERATE | 0.410256 | 0 | 39 | 186 | . | . | . | . |
| PM1548-SK1P2 | TUBGCP6 | p.Cys1239Trp | missense_variant | MODERATE | 0.388489 | 0.00947867 | 139 | 211 | . | . | . | . |
| PM1548-SK1P2 | TUBGCP6 | p.Ser1237Pro | missense_variant | MODERATE | 0.148148 | 0.00420168 | 135 | 238 | . | . | . | . |
| PM1548-SK1P2 | MIOX | p.Asp78Gly | missense_variant | MODERATE | 0 | 0 | 264 | 102 | . | . | . | . |
| PM1548-SK1P2 | MAPK8IP2 | p.Thr518Thr | synonymous_variant | LOW | 0.0277325 | 0 | 613 | 166 | . | . | . | . |
| PM1548-SK1P2 | ARSA | c.684+1G>A | splice_donor_variant&intron_variant | HIGH | 0.531969 | 0 | 391 | 160 | . | . | . | . |
| PM1548-SK1P2 | SETD5 | p.Arg1193* | stop_gained | HIGH | 0.567797 | 0 | 118 | 243 | . | . | . | . |
| PM1548-SK1P2 | CPNE9 | p.Ser284Ser | synonymous_variant | LOW | 0.0625 | 0 | 64 | 197 | . | . | . | . |
| PM1548-SK1P2 | CPNE9 | p.Asp473Asp | synonymous_variant | LOW | 0.43956 | 0 | 182 | 85 | . | . | . | . |
| PM1548-SK1P2 | BRPF1 | p.Gln938fs | frameshift_variant | HIGH | 0.1 | 0 | 60 | 218 | . | . | . | true |
| PM1548-SK1P2 | OGG1 | p.Ala85Thr | missense_variant | MODERATE | 0.592593 | 0.00289855 | 27 | 345 | . | . | . | . |
| PM1548-SK1P2 | CRELD1 | p.Arg220* | stop_gained | HIGH | 0.23913 | 0 | 598 | 387 | . | . | . | . |
| PM1548-SK1P2 | PRRT3 | p.His679Arg | missense_variant | MODERATE | 0.468085 | 0 | 517 | 166 | . | . | . | . |
| PM1548-SK1P2 | CAND2 | p.Arg489His | missense_variant | MODERATE | 0.282609 | 0 | 46 | 331 | . | . | . | . |
| PM1548-SK1P2 | SLC6A6 | p.Phe392fs | frameshift_variant | HIGH | 0.475763 | 0 | 557 | 115 | . | . | . | . |
| PM1548-SK1P2 | NR2C2 | p.Phe582fs | frameshift_variant | HIGH | 0.381313 | 0 | 396 | 75 | . | . | . | . |
| PM1548-SK1P2 | SH3BP5 | p.Thr119Ala | missense_variant | MODERATE | 0.430556 | 0 | 504 | 345 | . | . | . | . |
| PM1548-SK1P2 | PLCL2 | p.Gln628Arg | missense_variant | MODERATE | 0.1875 | 0 | 16 | 118 | . | . | . | . |
| PM1548-SK1P2 | SLC4A7 | p.Ile1097Val | missense_variant | MODERATE | 0.0344828 | 0.00657895 | 29 | 152 | . | . | . | . |
| PM1548-SK1P2 | CLASP2 | p.Cys10Cys | synonymous_variant | LOW | 0.482102 | 0.00520833 | 1732 | 192 | . | . | . | . |
| PM1548-SK1P2 | SCN10A | p.Val378Leu | missense_variant | MODERATE | 0.113208 | 0.00816327 | 53 | 245 | . | . | . | . |
| PM1548-SK1P2 | TTC21A | p.Ser332Thr | missense_variant | MODERATE | 0 | 0 | 22 | 40 | . | . | . | . |
| PM1548-SK1P2 | SNRK | p.Pro598Thr | missense_variant | MODERATE | 0.00884956 | 0 | 226 | 93 | . | . | . | . |
| PM1548-SK1P2 | SLC6A20 | p.Asn103Lys | missense_variant | MODERATE | 0 | 0 | 490 | 172 | . | . | . | . |
| PM1548-SK1P2 | SLC6A20 | p.Gly66Trp | missense_variant | MODERATE | 0.262295 | 0.00847458 | 61 | 118 | . | . | . | . |
| PM1548-SK1P2 | FYCO1 | p.Lys501del | inframe_deletion | MODERATE | 0.481675 | 0 | 191 | 171 | . | . | . | . |
| PM1548-SK1P2 | ALS2CL | p.Asn384Lys | missense_variant | MODERATE | 0.5 | 0 | 12 | 77 | . | . | . | . |
| PM1548-SK1P2 | DHX30 | n.47859525delC | intragenic_variant | MODIFIER | 0.433333 | 0 | 300 | 118 | . | . | . | . |
| PM1548-SK1P2 | UCN2 | p.Thr27Pro | missense_variant | MODERATE | 0.144781 | 0 | 297 | 134 | . | . | . | . |
| PM1548-SK1P2 | CELSR3 | p.Pro1425Ser | missense_variant | MODERATE | 0 | 0 | 259 | 95 | . | . | . | . |
| PM1548-SK1P2 | SLC25A20 | p.Ala227Val | missense_variant | MODERATE | 0.655172 | 0 | 29 | 126 | . | . | . | . |
| PM1548-SK1P2 | KLHDC8B | p.Gly38fs | frameshift_variant | HIGH | 0.116279 | 0 | 86 | 199 | . | . | . | . |
| PM1548-SK1P2 | RRP9 | p.Ala342Thr | missense_variant | MODERATE | 0.375723 | 0 | 346 | 161 | . | . | . | . |
| PM1548-SK1P2 | DNAH1 | p.Leu2072Ile | missense_variant | MODERATE | 0.416667 | 0 | 24 | 99 | . | . | . | . |
| PM1548-SK1P2 | FAM208A | p.Ala227Val | missense_variant | MODERATE | 0.25 | 0 | 8 | 81 | . | . | . | . |
| PM1548-SK1P2 | DNAH12 | p.Pro1035Ala | missense_variant | MODERATE | 0.136364 | 0 | 44 | 189 | . | . | . | . |
| PM1548-SK1P2 | PDE12 | p.Ala293Ser | missense_variant | MODERATE | 0 | 0 | 39 | 150 | . | . | . | . |
| PM1548-SK1P2 | PDE12 | p.Leu294Phe | missense_variant | MODERATE | 0 | 0 | 38 | 150 | . | . | . | . |
| PM1548-SK1P2 | ADAMTS9 | p.Pro596His | missense_variant | MODERATE | 0 | 0 | 9 | 126 | . | . | . | . |
| PM1548-SK1P2 | ADAMTS9 | p.Ser593Gly | missense_variant | MODERATE | 0 | 0 | 10 | 130 | . | . | . | . |
| PM1548-SK1P2 | ROBO1 | p.Tyr1009* | stop_gained | HIGH | 0 | 0 | 32 | 54 | . | . | . | . |
| PM1548-SK1P2 | OR5K4 | p.Val51Ile | missense_variant | MODERATE | 0.107143 | 0 | 28 | 442 | . | . | . | . |
| PM1548-SK1P2 | OR5K4 | p.Arg53Pro | missense_variant | MODERATE | 0.107143 | 0 | 28 | 441 | . | . | . | . |
| PM1548-SK1P2 | OR5K4 | p.Leu56His | missense_variant | MODERATE | 0.0714286 | 0 | 42 | 510 | . | . | . | . |
| PM1548-SK1P2 | DCBLD2 | p.Thr718Met | missense_variant | MODERATE | 0.456311 | 0 | 206 | 81 | . | . | . | . |
| PM1548-SK1P2 | TFG | p.Ser213Pro | missense_variant | MODERATE | 0.4375 | 0 | 32 | 76 | true | . | . | . |
| PM1548-SK1P2 | CBLB | p.Pro823Ser | missense_variant | MODERATE | 0.333333 | 0 | 9 | 275 | true | . | . | . |
| PM1548-SK1P2 | BOC | p.Pro593Leu | missense_variant | MODERATE | 0.104167 | 0.00199203 | 96 | 502 | . | . | . | . |
| PM1548-SK1P2 | BOC | p.His601Gln | missense_variant | MODERATE | 0 | 0 | 74 | 255 | . | . | . | . |
| PM1548-SK1P2 | KIAA2018 | p.Asn1016fs | frameshift_variant | HIGH | 0.0697674 | 0 | 86 | 36 | . | . | . | . |
| PM1548-SK1P2 | KIAA1407 | p.Glu683Gly | missense_variant | MODERATE | 0.333333 | 0 | 9 | 85 | . | . | . | . |
| PM1548-SK1P2 | LSAMP | p.Val277Ile | missense_variant | MODERATE | 0.503759 | 0 | 133 | 91 | . | . | . | . |
| PM1548-SK1P2 | ARHGAP31 | p.Lys857Glu | missense_variant | MODERATE | 0 | 0 | 1 | 33 | . | . | . | . |
| PM1548-SK1P2 | FSTL1 | p.Val13Ala | missense_variant | MODERATE | 0.319149 | 0 | 141 | 223 | . | . | . | . |
| PM1548-SK1P2 | EAF2 | p.Thr112fs | frameshift_variant | HIGH | 0.4375 | 0 | 16 | 119 | . | . | . | . |
| PM1548-SK1P2 | ILDR1 | p.Ala76Val | missense_variant&splice_region_variant | MODERATE | 0 | 0 | 12 | 159 | . | . | . | . |
| PM1548-SK1P2 | MYLK | p.Ala1684Thr | missense_variant | MODERATE | 0.271028 | 0 | 107 | 98 | . | . | . | . |
| PM1548-SK1P2 | ALDH1L1 | p.Glu439fs | frameshift_variant | HIGH | 0.507853 | 0 | 191 | 55 | . | . | . | . |
| PM1548-SK1P2 | CHCHD6 | p.Ser220Leu | missense_variant | MODERATE | 0.00784314 | 0 | 255 | 164 | . | . | . | . |
| PM1548-SK1P2 | C3orf27 | p.Asp85fs | frameshift_variant | HIGH | 0.420168 | 0 | 119 | 229 | . | . | . | . |
| PM1548-SK1P2 | TMCC1 | p.Arg620Ser | missense_variant | MODERATE | 0.1 | 0 | 40 | 62 | . | . | . | . |
| PM1548-SK1P2 | CDV3 | p.Gly226Asp | missense_variant | MODERATE | 0 | 0 | 24 | 44 | . | . | . | . |
| PM1548-SK1P2 | CDV3 | p.Asp228Glu | missense_variant | MODERATE | 0 | 0 | 24 | 55 | . | . | . | . |
| PM1548-SK1P2 | AMOTL2 | p.Arg292Cys | missense_variant | MODERATE | 0 | 0 | 24 | 43 | . | . | . | . |
| PM1548-SK1P2 | EPHB1 | p.Ala82fs | frameshift_variant | HIGH | 0.0633484 | 0 | 221 | 300 | . | . | . | . |
| PM1548-SK1P2 | SLC35G2 | p.Asn97fs | frameshift_variant | HIGH | 0 | 0 | 1 | 31 | . | . | . | . |
| PM1548-SK1P2 | ESYT3 | p.Val238Met | missense_variant | MODERATE | 0.551331 | 0 | 263 | 78 | . | . | . | . |
| PM1548-SK1P2 | MED12L | p.Ser970Leu | missense_variant | MODERATE | 0 | 0 | 12 | 123 | . | . | . | . |
| PM1548-SK1P2 | DHX36 | p.Met154fs | frameshift_variant | HIGH | 0 | 0 | 6 | 44 | . | . | . | . |
| PM1548-SK1P2 | CCNL1 | p.Ile354Val | missense_variant | MODERATE | 0 | 0 | 33 | 120 | . | . | . | . |
| PM1548-SK1P2 | PPM1L | p.Glu106Glu | synonymous_variant | LOW | 0.444444 | 0 | 9 | 275 | . | . | . | . |
| PM1548-SK1P2 | SLITRK3 | p.Arg759Cys | missense_variant | MODERATE | 0.542373 | 0 | 59 | 99 | . | . | . | . |
| PM1548-SK1P2 | ZBBX | p.Ile496fs | frameshift_variant | HIGH | 0.3125 | 0 | 16 | 76 | . | . | . | . |
| PM1548-SK1P2 | FNDC3B | p.Glu687Gly | missense_variant | MODERATE | 0 | 0 | 1 | 96 | . | . | . | . |
| PM1548-SK1P2 | NCEH1 | p.Ser105fs | frameshift_variant | HIGH | 0 | 0 | 32 | 228 | . | . | . | . |
| PM1548-SK1P2 | KLHL24 | p.Asn555fs | frameshift_variant | HIGH | 0 | 0 | 3 | 55 | . | . | . | . |
| PM1548-SK1P2 | CHRD | p.Arg76Cys | missense_variant | MODERATE | 0.641026 | 0 | 78 | 91 | . | . | . | . |
| PM1548-SK1P2 | TP63 | p.Thr368Ala | missense_variant | MODERATE | 0.246753 | 0 | 77 | 121 | . | . | . | . |
| PM1548-SK1P2 | LEPREL1 | p.Asp204Glu | missense_variant | MODERATE | 0.125 | 0 | 48 | 176 | . | . | . | . |
| PM1548-SK1P2 | CCDC50 | p.Ala343Ser | missense_variant | MODERATE | 0.117647 | 0 | 34 | 159 | . | . | . | . |
| PM1548-SK1P2 | MUC20 | p.Gly514Arg | missense_variant | MODERATE | 0.285714 | 0 | 28 | 228 | . | . | . | . |
| PM1548-SK1P2 | LRCH3 | p.Leu49Val | missense_variant | MODERATE | 0.139394 | 0 | 165 | 266 | . | . | . | . |
| PM1548-SK1P2 | GAK | p.Ala690Val | missense_variant | MODERATE | 0 | 0 | 132 | 111 | . | . | . | . |
| PM1548-SK1P2 | DGKQ | p.Thr299Met | missense_variant | MODERATE | 0.445759 | 0 | 507 | 253 | . | . | . | . |
| PM1548-SK1P2 | SLC26A1 | p.Ala624Thr | missense_variant | MODERATE | 0.00242718 | 0 | 412 | 466 | . | . | . | . |
| PM1548-SK1P2 | POLN | p.Tyr686Cys | missense_variant | MODERATE | 0 | 0 | 87 | 37 | . | . | . | . |

| | | | | | | | | | | | | |
|---|---|---|---|---|---|---|---|---|---|---|---|---|
| PM1548-SK1P2 | HTT | p.Ala2259Thr | missense_variant&splice_region_variant | MODERATE | 0.666667 | 0 | 3 | 52 | . | . | . | . |
| PM1548-SK1P2 | PPP2R2C | p.Gly397fs | frameshift_variant | HIGH | 0 | 0 | 50 | 188 | . | . | . | . |
| PM1548-SK1P2 | SH3TC1 | p.Glu203Gly | missense_variant | MODERATE | 0.291667 | 0.0021322 | 24 | 469 | . | . | . | . |
| PM1548-SK1P2 | GPR78 | p.Val209Ile | missense_variant | MODERATE | 0.574803 | 0 | 127 | 34 | . | . | . | . |
| PM1548-SK1P2 | KCTD8 | p.Leu62Leu | synonymous_variant | LOW | 0.00990099 | 0 | 202 | 213 | . | . | . | . |
| PM1548-SK1P2 | YTHDC1 | p.Ser684Asn | missense_variant | MODERATE | 0.333333 | 0 | 6 | 59 | . | . | . | . |
| PM1548-SK1P2 | ANKRD17 | p.Ile2241Val | missense_variant | MODERATE | 0 | 0 | 12 | 147 | . | . | . | . |
| PM1548-SK1P2 | PF4 | p.Ala4Thr | missense_variant | MODERATE | 0.423077 | 0 | 26 | 40 | . | . | . | . |
| PM1548-SK1P2 | PARM1 | p.Ser82Pro | missense_variant | MODERATE | 0.105263 | 0 | 38 | 387 | . | . | . | . |
| PM1548-SK1P2 | CDKL2 | p.Pro452Ser | missense_variant | MODERATE | 0 | 0 | 4 | 51 | . | . | . | . |
| PM1548-SK1P2 | SHROOM3 | p.Lys1017del | inframe_deletion | MODERATE | 0.517906 | 0 | 363 | 97 | . | . | . | . |
| PM1548-SK1P2 | BMP3 | p.Lys358fs | frameshift_variant | HIGH | 0.625 | 0 | 8 | 124 | . | . | . | . |
| PM1548-SK1P2 | PRKG2 | p.Val461Phe | missense_variant | MODERATE | 0 | 0 | 2 | 44 | . | . | . | . |
| PM1548-SK1P2 | SEC31A | p.Ile462fs | frameshift_variant | HIGH | 1 | 0 | 12 | 87 | . | . | . | . |
| PM1548-SK1P2 | SEC31A | p.Lys461fs | frameshift_variant | HIGH | 0 | 0 | 12 | 87 | . | . | . | . |
| PM1548-SK1P2 | SEC31A | p.Gln422Pro | missense_variant | MODERATE | 0.294118 | 0 | 34 | 143 | . | . | . | . |
| PM1548-SK1P2 | AGPAT9 | p.Met372fs | frameshift_variant | HIGH | 0.430769 | 0 | 65 | 70 | . | . | . | . |
| PM1548-SK1P2 | NAP1L5 | p.Lys108fs | frameshift_variant | HIGH | 0.456954 | 0 | 151 | 237 | . | . | . | . |
| PM1548-SK1P2 | CASP6 | p.Pro33Leu | missense_variant | MODERATE | 0 | 0.00719424 | 20 | 278 | . | . | . | . |
| PM1548-SK1P2 | PITX2 | c.411+284C>T | intron_variant | MODIFIER | 0.519348 | 0 | 491 | 59 | . | . | . | . |
| PM1548-SK1P2 | TNIP3 | p.Ala59Thr | missense_variant | MODERATE | 1 | 0 | 1 | 33 | . | . | . | . |
| PM1548-SK1P2 | PCDH10 | p.Tyr324Cys | missense_variant | MODERATE | 0.947917 | 0 | 192 | 49 | . | . | . | . |
| PM1548-SK1P2 | SETD7 | p.Leu331Ile | missense_variant | MODERATE | 0.422434 | 0 | 419 | 328 | . | . | . | . |
| PM1548-SK1P2 | FREM3 | p.Arg502* | stop_gained | HIGH | 0 | 0 | 11 | 34 | . | . | . | . |
| PM1548-SK1P2 | NEK1 | p.Thr137Asn | missense_variant | MODERATE | 0.0526316 | 0 | 19 | 84 | . | . | . | . |
| PM1548-SK1P2 | AHRR | p.Arg72His | missense_variant | MODERATE | 0.399361 | 0 | 313 | 75 | . | . | . | . |
| PM1548-SK1P2 | EXOC3 | p.Val576Met | missense_variant | MODERATE | 0.833333 | 0 | 12 | 107 | . | . | . | . |
| PM1548-SK1P2 | LPCAT1 | p.Val411Ile | missense_variant | MODERATE | 0 | 0 | 299 | 136 | . | . | . | . |
| PM1548-SK1P2 | IRX2 | p.Tyr110Cys | missense_variant | MODERATE | 0.514654 | 0 | 853 | 189 | . | . | . | . |
| PM1548-SK1P2 | NSUN2 | p.Ala653Thr | missense_variant&splice_region_variant | MODERATE | 0.428571 | 0 | 35 | 83 | . | . | . | . |
| PM1548-SK1P2 | SRD5A1 | p.Met162Ile | missense_variant | MODERATE | 0.5 | 0 | 40 | 54 | . | . | . | . |
| PM1548-SK1P2 | SEMA5A | p.Ile679Thr | missense_variant | MODERATE | 0.1 | 0 | 40 | 101 | . | . | . | . |
| PM1548-SK1P2 | NIPBL | p.Glu1060Glu | synonymous_variant | LOW | 0 | 0 | 12 | 121 | . | . | . | . |
| PM1548-SK1P2 | CARD6 | p.Gln393Glu | missense_variant | MODERATE | 0 | 0 | 25 | 371 | . | . | . | . |
| PM1548-SK1P2 | CARD6 | p.Gln400His | missense_variant | MODERATE | 0 | 0 | 24 | 367 | . | . | . | . |
| PM1548-SK1P2 | CARD6 | p.Gln402* | stop_gained | HIGH | 0 | 0 | 24 | 367 | . | . | . | . |
| PM1548-SK1P2 | CARD6 | p.Gln402Arg | missense_variant | MODERATE | 0 | 0 | 23 | 337 | . | . | . | . |
| PM1548-SK1P2 | CARD6 | p.Gln402His | missense_variant | MODERATE | 0 | 0 | 23 | 337 | . | . | . | . |
| PM1548-SK1P2 | C6 | p.Ala2Val | missense_variant | MODERATE | 0.173077 | 0 | 52 | 129 | . | . | . | . |
| PM1548-SK1P2 | SNX18 | n.53839150G>A | intragenic_variant | MODIFIER | 0.333333 | 0 | 15 | 117 | . | . | . | . |
| PM1548-SK1P2 | GPX8 | p.Phe125fs | frameshift_variant | HIGH | 0.3125 | 0 | 32 | 228 | . | . | . | . |
| PM1548-SK1P2 | UTP15 | c.169-826A>G | intron_variant | MODIFIER | 0.133333 | 0 | 30 | 55 | . | . | . | . |
| PM1548-SK1P2 | TTC37 | p.Ala1407Thr | missense_variant | MODERATE | 1 | 0 | 3 | 33 | . | . | . | . |
| PM1548-SK1P2 | CHD1 | p.Asn310fs | frameshift_variant | HIGH | 1 | 0 | 45 | 201 | . | . | . | true |
| PM1548-SK1P2 | PAM | p.Arg260Gln | missense_variant | MODERATE | 0 | 0 | 22 | 107 | . | . | . | . |
| PM1548-SK1P2 | GIN1 | p.Leu345Pro | missense_variant | MODERATE | 0.0833333 | 0 | 12 | 72 | . | . | . | . |
| PM1548-SK1P2 | APC | p.Asp802fs | frameshift_variant&stop_gained | HIGH | 0.448598 | 0 | 214 | 125 | true | true | true | true |
| PM1548-SK1P2 | MCC | p.Arg800Lys | missense_variant | MODERATE | 0 | 0 | 4 | 65 | . | . | . | . |
| PM1548-SK1P2 | FTMT | p.Ala90Val | missense_variant | MODERATE | 0.53202 | 0 | 406 | 105 | . | . | . | . |
| PM1548-SK1P2 | FBN2 | p.Leu2853Arg | missense_variant | MODERATE | 0.409836 | 0 | 61 | 91 | . | . | . | . |
| PM1548-SK1P2 | FNIP1 | p.Leu90Met | missense_variant | MODERATE | 0 | 0 | 3 | 76 | . | . | . | . |
| PM1548-SK1P2 | JADE2 | p.Phe360Leu | missense_variant | MODERATE | 0.226667 | 0 | 150 | 343 | . | . | . | true |
| PM1548-SK1P2 | FBXL21 | n.956C>T | non_coding_exon_variant | MODIFIER | 0.5 | 0 | 6 | 13 | . | . | . | . |
| PM1548-SK1P2 | KDM3B | p.Arg921Trp | missense_variant | MODERATE | 0 | 0 | 23 | 178 | . | . | . | . |
| PM1548-SK1P2 | PSD2 | p.Arg574His | missense_variant | MODERATE | 0.380597 | 0.0147059 | 536 | 136 | . | . | . | . |
| PM1548-SK1P2 | PCDHA3 | p.Glu558Val | missense_variant | MODERATE | 0.134615 | 0 | 52 | 71 | . | . | . | . |
| PM1548-SK1P2 | PCDHA5 | p.Ala56Thr | missense_variant | MODERATE | 0.264151 | 0 | 53 | 32 | . | . | . | . |
| PM1548-SK1P2 | PCDHA5 | p.Arg64Arg | synonymous_variant | LOW | 0.254545 | 0 | 55 | 38 | . | . | . | . |
| PM1548-SK1P2 | PCDHA5 | p.Gly69Asp | missense_variant | MODERATE | 0.224138 | 0 | 58 | 47 | . | . | . | . |
| PM1548-SK1P2 | PCDHA8 | p.Ala57Thr | missense_variant | MODERATE | 0.0869565 | 0 | 46 | 46 | . | . | . | . |
| PM1548-SK1P2 | PCDHA13 | p.Glu558Val | missense_variant | MODERATE | 0.285714 | 0 | 21 | 106 | . | . | . | . |
| PM1548-SK1P2 | PCDHAC2 | p.Arg543Trp | missense_variant | MODERATE | 0.583333 | 0 | 24 | 153 | . | . | . | . |
| PM1548-SK1P2 | PCDHAC2 | c.2565+37G>A | intron_variant | MODIFIER | 0.470199 | 0 | 151 | 61 | . | . | . | . |
| PM1548-SK1P2 | PCDHGA1 | p.Ser370Gly | missense_variant | MODERATE | 0.638889 | 0 | 36 | 385 | . | . | . | . |
| PM1548-SK1P2 | PCDHGA9 | p.Val422Ala | missense_variant | MODERATE | 0 | 0 | 5 | 94 | . | . | . | . |
| PM1548-SK1P2 | PCDH1 | p.Ala203Thr | missense_variant | MODERATE | 0.443378 | 0 | 521 | 253 | . | . | . | . |
| PM1548-SK1P2 | PCDH12 | p.Gly10Glu | missense_variant | MODERATE | 0.3 | 0 | 50 | 44 | . | . | . | . |
| PM1548-SK1P2 | CSF1R | p.Glu955Lys | missense_variant | MODERATE | 0.541667 | 0 | 24 | 65 | . | . | . | . |
| PM1548-SK1P2 | TIMD4 | p.Ala333Val | missense_variant | MODERATE | 0.36 | 0 | 25 | 229 | . | . | . | . |
| PM1548-SK1P2 | ATP10B | p.Arg115Lys | missense_variant | MODERATE | 0.0112045 | 0 | 357 | 157 | . | . | . | . |
| PM1548-SK1P2 | UBTD2 | p.Thr160Ile | missense_variant | MODERATE | 0.428571 | 0 | 7 | 33 | . | . | . | . |
| PM1548-SK1P2 | GPRIN1 | p.Pro806fs | frameshift_variant | HIGH | 0.0773481 | 0 | 181 | 165 | . | . | . | . |
| PM1548-SK1P2 | GPRIN1 | p.Pro805Thr | missense_variant | MODERATE | 0.0777778 | 0 | 180 | 165 | . | . | . | . |
| PM1548-SK1P2 | GPRIN1 | p.Ser803fs | frameshift_variant | HIGH | 0.0782123 | 0 | 179 | 165 | . | . | . | . |
| PM1548-SK1P2 | GPRIN1 | p.Glu801Asp | missense_variant | MODERATE | 0.0782123 | 0 | 179 | 165 | . | . | . | . |
| PM1548-SK1P2 | UIMC1 | p.Phe165Cys | missense_variant | MODERATE | 0.171429 | 0 | 35 | 220 | . | . | . | . |
| PM1548-SK1P2 | UIMC1 | p.Ile156Val | missense_variant&splice_region_variant | MODERATE | 0.133333 | 0 | 45 | 259 | . | . | . | . |
| PM1548-SK1P2 | FGFR4 | p.Pro568fs | frameshift_variant | HIGH | 0.105263 | 0 | 19 | 174 | true | . | . | . |
| PM1548-SK1P2 | FAM193B | p.Pro761Ser | missense_variant | MODERATE | 0.126582 | 0 | 79 | 84 | . | . | . | . |
| PM1548-SK1P2 | GRM6 | p.Ala93Thr | missense_variant | MODERATE | 0.490566 | 0 | 318 | 91 | . | . | . | . |
| PM1548-SK1P2 | RREB1 | p.Val754Met | missense_variant | MODERATE | 0.0165017 | 0 | 1515 | 159 | . | . | . | . |
| PM1548-SK1P2 | HIST1H4C | p.Val82Ala | missense_variant | MODERATE | 0.142857 | 0 | 14 | 187 | . | . | . | . |
| PM1548-SK1P2 | HIST1H2AM | p.Leu52Met | missense_variant | MODERATE | 0.0487805 | 0 | 41 | 43 | . | . | . | . |
| PM1548-SK1P2 | TRIM27 | p.Ala24Val | missense_variant | MODERATE | 0.033195 | 0 | 241 | 147 | true | . | . | . |
| PM1548-SK1P2 | OR2H1 | p.Ile295Leu | missense_variant | MODERATE | 0.434783 | 0 | 23 | 181 | . | . | . | . |
| PM1548-SK1P2 | HLA-F | p.Leu396fs | frameshift_variant | HIGH | 0.454545 | 0 | 11 | 38 | . | . | . | . |
| PM1548-SK1P2 | HLA-G | p.Arg285Ser | missense_variant | MODERATE | 0.1 | 0 | 10 | 42 | . | . | . | . |
| PM1548-SK1P2 | TRIM26 | p.Glu241Lys | missense_variant | MODERATE | 0.285714 | 0 | 42 | 200 | . | . | . | . |
| PM1548-SK1P2 | MDC1 | p.Gln1431Arg | missense_variant | MODERATE | 0.175676 | 0 | 74 | 161 | . | . | . | . |
| PM1548-SK1P2 | MUC22 | p.Ile1250Thr | missense_variant | MODERATE | 0.129032 | 0 | 31 | 194 | . | . | . | . |
| PM1548-SK1P2 | HLA-C | p.Arg243Trp | missense_variant | MODERATE | 0.2 | 0 | 10 | 40 | . | . | . | . |
| PM1548-SK1P2 | HLA-B | p.Arg243Trp | missense_variant | MODERATE | 0.12963 | 0.00591716 | 162 | 338 | . | . | true | . |
| PM1548-SK1P2 | VWA7 | p.Thr26fs | frameshift_variant | HIGH | 0.516667 | 0 | 180 | 236 | . | . | . | . |
| PM1548-SK1P2 | CFB | p.Asp1161fs | frameshift_variant | HIGH | 0.43956 | 0 | 91 | 131 | . | . | . | . |
| PM1548-SK1P2 | RGL2 | p.Glu372fs | frameshift_variant | HIGH | 0.107623 | 0 | 223 | 192 | . | . | . | . |
| PM1548-SK1P2 | DNAH8 | p.Tyr1705Cys | missense_variant | MODERATE | 0.129032 | 0 | 31 | 68 | . | . | . | . |
| PM1548-SK1P2 | KLHDC3 | p.Arg67Cys | missense_variant | MODERATE | 0.0909091 | 0 | 11 | 263 | . | . | . | . |
| PM1548-SK1P2 | PKHD1 | p.Arg880His | missense_variant | MODERATE | 0.333333 | 0 | 30 | 253 | . | . | . | . |
| PM1548-SK1P2 | DST | p.Thr1051fs | frameshift_variant | HIGH | 0.384615 | 0 | 13 | 228 | . | . | . | . |
| PM1548-SK1P2 | KIAA1586 | p.Arg617Leu | missense_variant | MODERATE | 0.102564 | 0 | 39 | 71 | . | . | . | . |
| PM1548-SK1P2 | ZNF451 | p.Phe193Leu | missense_variant&splice_region_variant | MODERATE | 0.5 | 0 | 2 | 32 | . | . | . | . |
| PM1548-SK1P2 | COL12A1 | p.Pro2509Thr | missense_variant | MODERATE | 0.108108 | 0 | 37 | 155 | . | . | . | . |
| PM1548-SK1P2 | SENP6 | p.Ser627Gly | missense_variant | MODERATE | 0.0140845 | 0 | 71 | 66 | . | . | . | . |
| PM1548-SK1P2 | ME1 | c.78+1G>A | splice_donor_variant&intron_variant | HIGH | 0.387409 | 0 | 413 | 306 | . | . | . | . |
| PM1548-SK1P2 | ANKRD6 | p.Ala390Val | missense_variant | MODERATE | 0 | 0 | 5 | 31 | . | . | . | . |
| PM1548-SK1P2 | MDN1 | p.Thr2167Lys | missense_variant | MODERATE | 0 | 0 | 1 | 72 | . | . | . | . |
| PM1548-SK1P2 | MMS22L | p.Gly37Glu | missense_variant | MODERATE | 0.823529 | 0 | 17 | 240 | . | . | . | . |
| PM1548-SK1P2 | ATG5 | p.Asp149fs | frameshift_variant | HIGH | 0 | 0 | 135 | 95 | . | . | . | . |
| PM1548-SK1P2 | RFPL4B | p.Arg203Cys | missense_variant | MODERATE | 0.25 | 0 | 8 | 78 | . | . | . | . |
| PM1548-SK1P2 | KPNA5 | p.Phe383Ser | missense_variant | MODERATE | 0 | 0 | 8 | 67 | . | . | . | . |
| PM1548-SK1P2 | FAM184A | p.Glu79Asp | missense_variant | MODERATE | 0 | 0 | 35 | 52 | . | . | . | . |
| PM1548-SK1P2 | TBC1D32 | p.Ile27Asn | missense_variant | MODERATE | 0.666667 | 0 | 12 | 42 | . | . | . | . |
| PM1548-SK1P2 | HSF2 | p.Arg196His | missense_variant | MODERATE | 0.545455 | 0 | 11 | 76 | . | . | . | . |
| PM1548-SK1P2 | KIAA0408 | p.Phe313Leu | missense_variant | MODERATE | 0.104167 | 0 | 48 | 108 | . | . | . | . |
| PM1548-SK1P2 | SOGA3 | p.Ala214fs | frameshift_variant | HIGH | 0 | 0 | 17 | 41 | . | . | . | . |
| PM1548-SK1P2 | ENPP1 | c.240+2T>C | splice_donor_variant&intron_variant | HIGH | 0.545455 | 0 | 99 | 63 | . | . | . | . |
| PM1548-SK1P2 | CTGF | p.Ser227Phe | missense_variant | MODERATE | 0.0571429 | 0 | 35 | 36 | . | . | . | . |
| PM1548-SK1P2 | CTGF | p.Ala226Thr | missense_variant | MODERATE | 0.0526316 | 0 | 38 | 46 | . | . | . | . |
| PM1548-SK1P2 | CTGF | p.Ile195Met | missense_variant | MODERATE | 0.0606061 | 0 | 33 | 61 | . | . | . | . |
| PM1548-SK1P2 | ADGB | p.Asp890Val | missense_variant | MODERATE | 0 | 0 | 14 | 47 | . | . | . | . |
| PM1548-SK1P2 | SYNE1 | p.Pro1307Ser | missense_variant | MODERATE | 0.818182 | 0 | 22 | 79 | . | . | . | . |
| PM1548-SK1P2 | ARID1B | p.Pro1578Ser | missense_variant | MODERATE | 0.0571429 | 0 | 35 | 48 | true | . | . | . |
| PM1548-SK1P2 | ARID1B | p.Ser1617Thr | missense_variant | MODERATE | 0.0266667 | 0 | 75 | 59 | true | . | . | . |
| PM1548-SK1P2 | TULP4 | p.Pro1275Thr | missense_variant | MODERATE | 0.105263 | 0 | 19 | 171 | . | . | . | . |
| PM1548-SK1P2 | TULP4 | p.Asp1298Asn | missense_variant | MODERATE | 0 | 0.0078125 | 27 | 256 | . | . | . | . |
| PM1548-SK1P2 | TULP4 | p.Asp1298Glu | missense_variant | MODERATE | 0.0740741 | 0 | 27 | 255 | . | . | . | . |
| PM1548-SK1P2 | SMOC2 | p.Ala54Thr | missense_variant | MODERATE | 0.787879 | 0 | 33 | 36 | . | . | . | . |

| | | | | | | | | | | | | |
|---|---|---|---|---|---|---|---|---|---|---|---|---|
| PM1548-SK1P2 | GPR146 | p.Pro248Gln | missense_variant | MODERATE | 0 | 0 | 105 | 46 | . | . | . | . |
| PM1548-SK1P2 | ELFN1 | p.Arg389Cys | missense_variant | MODERATE | 0.482818 | 0 | 582 | 43 | . | . | . | . |
| PM1548-SK1P2 | SDK1 | p.Pro1727fs | frameshift_variant | HIGH | 0.45977 | 0 | 522 | 269 | . | . | . | . |
| PM1548-SK1P2 | PMS2 | p.Thr277Ala | missense_variant | MODERATE | 0.5 | 0 | 4 | 38 | true | . | . | . |
| PM1548-SK1P2 | MIOS | p.Leu213His | missense_variant | MODERATE | 1 | 0 | 1 | 44 | . | . | . | . |
| PM1548-SK1P2 | ETV1 | p.Asn37Thr | missense_variant | MODERATE | 0 | 0 | 23 | 72 | true | . | . | . |
| PM1548-SK1P2 | MPP6 | p.Lys306fs | frameshift_variant | HIGH | 0 | 0 | 226 | 33 | . | . | . | . |
| PM1548-SK1P2 | HERPUD2 | p.Val266Ala | missense_variant | MODERATE | 0.046875 | 0 | 64 | 59 | . | . | . | . |
| PM1548-SK1P2 | POU6F2 | p.Glu562Ala | missense_variant | MODERATE | 0.0135135 | 0 | 518 | 226 | . | . | . | . |
| PM1548-SK1P2 | MYO1G | p.Ala779Val | missense_variant | MODERATE | 0.473272 | 0 | 767 | 358 | . | . | . | . |
| PM1548-SK1P2 | VWC2 | p.Ser3Arg | missense_variant | MODERATE | 0.102041 | 0 | 49 | 64 | . | . | . | . |
| PM1548-SK1P2 | GRB10 | p.Ala349Thr | missense_variant | MODERATE | 0.675676 | 0 | 37 | 372 | . | . | . | . |
| PM1548-SK1P2 | WBSCR22 | p.Pro8Gln | missense_variant | MODERATE | 0 | 0 | 4 | 35 | . | . | . | . |
| PM1548-SK1P2 | CLIP2 | p.Arg771Gln | missense_variant | MODERATE | 0.00917431 | 0 | 654 | 150 | . | . | . | . |
| PM1548-SK1P2 | TMEM60 | p.Ala78fs | frameshift_variant | HIGH | 0.428571 | 0 | 7 | 31 | . | . | . | . |
| PM1548-SK1P2 | GNAI1 | p.Cys214Phe | missense_variant | MODERATE | 0.263158 | 0 | 38 | 194 | . | . | . | . |
| PM1548-SK1P2 | HGF | p.Val529Ala | missense_variant | MODERATE | 0.00636943 | 0 | 157 | 125 | . | true | . | . |
| PM1548-SK1P2 | ABCB1 | p.Tyr1165* | stop_gained | HIGH | 0.722222 | 0 | 18 | 32 | . | . | . | . |
| PM1548-SK1P2 | ABCB1 | c.3490-1G>C | splice_acceptor_variant&intron_variant | HIGH | 0.722222 | 0 | 18 | 32 | . | . | . | . |
| PM1548-SK1P2 | ABCB1 | p.Ser196Ala | missense_variant | MODERATE | 0.117647 | 0 | 51 | 55 | . | . | . | . |
| PM1548-SK1P2 | MTERF | p.Ala221Thr | missense_variant | MODERATE | 0 | 0 | 2 | 212 | . | . | . | . |
| PM1548-SK1P2 | AKAP9 | p.Ile1377Thr | missense_variant | MODERATE | 0 | 0 | 3 | 76 | true | . | . | . |
| PM1548-SK1P2 | ASB4 | p.Thr229Met | missense_variant | MODERATE | 0.416667 | 0 | 12 | 168 | . | . | . | . |
| PM1548-SK1P2 | LMTK2 | p.Asp236fs | frameshift_variant | HIGH | 0.25 | 0 | 8 | 300 | . | . | . | . |
| PM1548-SK1P2 | LMTK2 | p.Leu1130Ser | missense_variant | MODERATE | 0.571429 | 0 | 21 | 72 | . | . | . | . |
| PM1548-SK1P2 | TRRAP | p.Asn1675Thr | missense_variant | MODERATE | 0.0567376 | 0 | 141 | 117 | true | . | . | . |
| PM1548-SK1P2 | TRRAP | p.Arg1978Trp | missense_variant | MODERATE | 0 | 0 | 90 | 221 | true | . | . | . |
| PM1548-SK1P2 | TRRAP | p.Pro2508Ala | missense_variant | MODERATE | 0.153846 | 0 | 117 | 384 | true | . | . | . |
| PM1548-SK1P2 | GIGYF1 | p.Leu580Pro | missense_variant | MODERATE | 0.146939 | 0 | 245 | 70 | . | . | . | . |
| PM1548-SK1P2 | SLC12A9 | p.Leu559fs | frameshift_variant | HIGH | 0.538462 | 0 | 13 | 56 | . | . | . | . |
| PM1548-SK1P2 | MOGAT3 | p.Asn82Lys | missense_variant | MODERATE | 0.12 | 0 | 50 | 150 | . | . | . | . |
| PM1548-SK1P2 | SH2B2 | p.Ala74Ala | synonymous_variant | LOW | 0.553691 | 0 | 298 | 70 | . | . | . | . |
| PM1548-SK1P2 | LRWD1 | p.Leu87Phe | missense_variant | MODERATE | 0.102564 | 0 | 39 | 78 | . | . | . | . |
| PM1548-SK1P2 | FBXL13 | p.Cys375Ser | missense_variant | MODERATE | 0 | 0 | 9 | 94 | . | . | . | . |
| PM1548-SK1P2 | RELN | p.Arg2363His | missense_variant | MODERATE | 0.393939 | 0 | 99 | 39 | . | . | . | . |
| PM1548-SK1P2 | ATXN7L1 | p.Val629Phe | missense_variant | MODERATE | 0 | 0 | 2 | 193 | . | . | . | . |
| PM1548-SK1P2 | PIK3CG | p.Thr607fs | frameshift_variant | HIGH | 0 | 0 | 14 | 44 | . | true | . | . |
| PM1548-SK1P2 | CBLL1 | p.Val277Ala | missense_variant | MODERATE | 0.105263 | 0 | 38 | 198 | . | . | . | . |
| PM1548-SK1P2 | SLC26A3 | p.Thr479Ala | missense_variant | MODERATE | 0.196721 | 0 | 61 | 194 | . | . | true | . |
| PM1548-SK1P2 | FOXP2 | p.Gly98Val | missense_variant | MODERATE | 0.133333 | 0.00793651 | 30 | 126 | . | . | . | . |
| PM1548-SK1P2 | PTPRZ1 | p.Pro751Leu | missense_variant | MODERATE | 0 | 0 | 2 | 64 | . | . | . | . |
| PM1548-SK1P2 | SND1 | p.Ala331Pro | missense_variant | MODERATE | 0.0223464 | 0 | 179 | 236 | true | . | . | . |
| PM1548-SK1P2 | CEP41 | p.Asp152Gly | missense_variant | MODERATE | 0.132812 | 0 | 128 | 235 | . | . | . | . |
| PM1548-SK1P2 | TSGA13 | p.Lys151fs | frameshift_variant | HIGH | 0 | 0 | 2 | 195 | . | . | . | . |
| PM1548-SK1P2 | TTC26 | p.Ala221Thr | missense_variant | MODERATE | 0.325581 | 0 | 43 | 82 | . | . | . | . |
| PM1548-SK1P2 | MKRN1 | c.186-2435G>C | intron_variant | MODIFIER | 0.8 | 0 | 15 | 136 | . | . | . | . |
| PM1548-SK1P2 | MGAM | p.Asn1603Asp | missense_variant | MODERATE | 0.363636 | 0 | 55 | 85 | . | . | . | . |
| PM1548-SK1P2 | EPHB6 | p.Ala253fs | frameshift_variant | HIGH | 0.625 | 0 | 24 | 107 | . | true | . | . |
| PM1548-SK1P2 | ZNF775 | p.Gly65fs | frameshift_variant | HIGH | 0 | 0 | 4 | 50 | . | . | . | . |
| PM1548-SK1P2 | FASTK | p.Ile345Asn | missense_variant | MODERATE | 0 | 0 | 37 | 260 | . | . | . | . |
| PM1548-SK1P2 | GALNT11 | p.Leu188Pro | missense_variant | MODERATE | 0 | 0 | 7 | 157 | . | . | . | . |
| PM1548-SK1P2 | TNKS | p.His859Asn | missense_variant | MODERATE | 0 | 0 | 33 | 75 | . | . | . | . |
| PM1548-SK1P2 | RP1L1 | p.Gln1044fs | frameshift_variant | HIGH | 1 | 0 | 456 | 101 | . | . | . | . |
| PM1548-SK1P2 | XKR6 | p.Arg80Lys | missense_variant | MODERATE | 0 | 0 | 2 | 127 | . | . | . | . |
| PM1548-SK1P2 | PCM1 | p.Met1321Val | missense_variant | MODERATE | 0 | 0 | 1 | 138 | true | . | . | . |
| PM1548-SK1P2 | SORBS3 | p.Thr100Ala | missense_variant | MODERATE | 0 | 0 | 125 | 140 | . | . | . | . |
| PM1548-SK1P2 | LOXL2 | p.Pro40Leu | missense_variant | MODERATE | 0.978947 | 0 | 95 | 224 | . | . | . | . |
| PM1548-SK1P2 | NEFM | p.Leu5Leu | synonymous_variant | LOW | 0 | 0 | 31 | 41 | . | . | . | . |
| PM1548-SK1P2 | NRG1 | p.Gly39Glu | missense_variant | MODERATE | 0 | 0 | 59 | 55 | true | . | . | . |
| PM1548-SK1P2 | NRG1 | p.Pro44Ser | missense_variant | MODERATE | 0 | 0 | 48 | 58 | true | . | . | . |
| PM1548-SK1P2 | NRG1 | p.Ala74Pro | missense_variant | MODERATE | 0 | 0 | 4 | 35 | true | . | . | . |
| PM1548-SK1P2 | NRG1 | c.1488-469C>G | intron_variant | MODIFIER | 0.190476 | 0 | 21 | 107 | true | . | . | . |
| PM1548-SK1P2 | ADAM2 | p.Arg3His | missense_variant | MODERATE | 0 | 0 | 58 | 64 | . | . | . | . |
| PM1548-SK1P2 | DKK4 | p.Asp195Asn | missense_variant | MODERATE | 0.355932 | 0 | 118 | 81 | . | . | . | . |
| PM1548-SK1P2 | MCM4 | p.Thr629fs | frameshift_variant | HIGH | 0.4375 | 0 | 32 | 85 | . | . | . | . |
| PM1548-SK1P2 | PXDNL | p.Ala353Thr | missense_variant | MODERATE | 0.666667 | 0 | 3 | 175 | . | . | . | . |
| PM1548-SK1P2 | RP1 | p.Tyr834fs | frameshift_variant | HIGH | 0.307692 | 0 | 13 | 39 | . | . | . | . |
| PM1548-SK1P2 | YTHDF3 | p.Gln323Gln | synonymous_variant | LOW | 0 | 0 | 35 | 589 | . | . | . | . |
| PM1548-SK1P2 | C8orf46 | p.Thr78Ile | missense_variant | MODERATE | 0 | 0 | 56 | 109 | . | . | . | . |
| PM1548-SK1P2 | ZFHX4 | p.Ala2905Val | missense_variant | MODERATE | 0.375 | 0 | 8 | 42 | . | . | . | . |
| PM1548-SK1P2 | HEY1 | p.Val187Ala | missense_variant | MODERATE | 0.0144231 | 0 | 416 | 281 | true | . | . | . |
| PM1548-SK1P2 | HEY1 | p.Thr186Ser | missense_variant | MODERATE | 0.0145278 | 0 | 413 | 272 | true | . | . | . |
| PM1548-SK1P2 | VPS13B | p.Asn2064Thr | missense_variant | MODERATE | 0.545455 | 0 | 22 | 75 | . | . | . | . |
| PM1548-SK1P2 | TRPS1 | p.Arg297Cys | missense_variant | MODERATE | 0.351648 | 0 | 91 | 259 | . | . | . | . |
| PM1548-SK1P2 | TRPS1 | p.Asn265Asn | synonymous_variant | LOW | 0.131579 | 0 | 38 | 334 | . | . | . | . |
| PM1548-SK1P2 | EXT1 | p.Val66Phe | missense_variant | MODERATE | 0 | 0 | 29 | 88 | true | . | . | . |
| PM1548-SK1P2 | TBC1D31 | p.Ala208Thr | missense_variant | MODERATE | 0.573427 | 0 | 143 | 184 | . | . | . | . |
| PM1548-SK1P2 | ATAD2 | p.Glu119fs | frameshift_variant | HIGH | 0.466667 | 0 | 30 | 108 | . | . | . | . |
| PM1548-SK1P2 | ZNF572 | p.Ser55Pro | missense_variant | MODERATE | 0.671875 | 0 | 64 | 39 | . | . | . | . |
| PM1548-SK1P2 | MYC | p.Gly175Ser | missense_variant | MODERATE | 0 | 0 | 145 | 270 | true | . | . | . |
| PM1548-SK1P2 | ASAP1 | p.Glu919Lys | missense_variant | MODERATE | 0.0810811 | 0 | 37 | 187 | . | . | . | . |
| PM1548-SK1P2 | WISP1 | p.Ile175Met | missense_variant | MODERATE | 0.029703 | 0 | 101 | 119 | . | . | . | . |
| PM1548-SK1P2 | KHDRBS3 | p.Tyr316Tyr | splice_region_variant&synonymous_variant | LOW | 0.363636 | 0 | 33 | 229 | . | . | . | . |
| PM1548-SK1P2 | COL22A1 | p.Thr125Met | missense_variant | MODERATE | 0.75 | 0 | 4 | 59 | . | . | . | . |
| PM1548-SK1P2 | LY6D | p.Ala99Thr | missense_variant | MODERATE | 0.238245 | 0.00956938 | 638 | 209 | . | . | . | . |
| PM1548-SK1P2 | TOP1MT | p.Pro193Leu | missense_variant | MODERATE | 0 | 0.00595238 | 30 | 168 | . | . | . | . |
| PM1548-SK1P2 | ZC3H3 | p.Gly101fs | frameshift_variant | HIGH | 0.333333 | 0 | 6 | 100 | . | . | . | . |
| PM1548-SK1P2 | GSDMD | p.Glu451Ala | missense_variant | MODERATE | 0.27186 | 0 | 629 | 215 | . | . | . | . |
| PM1548-SK1P2 | MAPK15 | p.Pro493Leu | missense_variant | MODERATE | 0.327945 | 0 | 433 | 366 | . | . | . | . |
| PM1548-SK1P2 | SCRIB | p.Cys22Trp | missense_variant | MODERATE | 0.390523 | 0 | 612 | 81 | . | . | . | . |
| PM1548-SK1P2 | PLEC | p.Arg3540Gln | missense_variant | MODERATE | 0 | 0 | 363 | 130 | . | . | . | . |
| PM1548-SK1P2 | PLEC | p.Val648Met | missense_variant | MODERATE | 0 | 0 | 65 | 166 | . | . | . | . |
| PM1548-SK1P2 | CYHR1 | n.145689659dup | intragenic_variant | MODIFIER | 0.141892 | 0 | 148 | 56 | . | . | . | . |
| PM1548-SK1P2 | ZNF7 | p.Val136Ile | missense_variant | MODERATE | 0 | 0 | 2 | 154 | . | . | . | . |
| PM1548-SK1P2 | KANK1 | p.Thr192Ser | missense_variant | MODERATE | 0 | 0 | 38 | 65 | . | . | . | . |
| PM1548-SK1P2 | DMRT3 | p.Leu385Leu | synonymous_variant | LOW | 0.102564 | 0 | 39 | 220 | . | . | . | . |
| PM1548-SK1P2 | RLN2 | p.His8fs | frameshift_variant | HIGH | 0.369231 | 0 | 65 | 225 | . | . | . | . |
| PM1548-SK1P2 | IFNA4 | p.Ala74Thr | missense_variant | MODERATE | 0 | 0 | 4 | 51 | . | . | . | . |
| PM1548-SK1P2 | TAF1L | p.Gly4Ser | missense_variant | MODERATE | 0.5 | 0 | 8 | 329 | . | . | . | true |
| PM1548-SK1P2 | BAG1 | p.Glu321Lys | missense_variant | MODERATE | 0.541379 | 0 | 290 | 192 | . | . | . | . |
| PM1548-SK1P2 | PRSS3 | p.Ser102Tyr | missense_variant | MODERATE | 0.0714286 | 0 | 28 | 102 | . | . | . | . |
| PM1548-SK1P2 | CD72 | p.Ala111Thr | missense_variant | MODERATE | 0.505263 | 0 | 95 | 227 | . | . | . | . |
| PM1548-SK1P2 | RGP1 | p.Arg328Trp | missense_variant | MODERATE | 0.391304 | 0 | 23 | 169 | . | . | . | . |
| PM1548-SK1P2 | ANKRD18A | p.Gly900Ala | missense_variant | MODERATE | 0.0666667 | 0 | 15 | 119 | . | . | . | . |
| PM1548-SK1P2 | ANKRD18A | p.Ala897Val | missense_variant | MODERATE | 0.125 | 0 | 16 | 119 | . | . | . | . |
| PM1548-SK1P2 | CNTNAP3 | p.Arg841Ser | missense_variant | MODERATE | 0 | 0 | 2 | 45 | . | . | . | . |
| PM1548-SK1P2 | TMC1 | p.Ala755Pro | missense_variant&splice_region_variant | MODERATE | 0 | 0 | 11 | 45 | . | . | . | . |
| PM1548-SK1P2 | RMI1 | p.Thr325Ala | missense_variant | MODERATE | 0.457286 | 0 | 199 | 482 | . | . | . | . |
| PM1548-SK1P2 | NAA35 | p.*726* | stop_retained_variant | LOW | 0 | 0 | 19 | 360 | . | . | . | . |
| PM1548-SK1P2 | SPATA31E1 | p.Arg436His | missense_variant | MODERATE | 0.238095 | 0 | 21 | 212 | . | . | . | . |
| PM1548-SK1P2 | ROR2 | p.Ala307Thr | missense_variant | MODERATE | 0 | 0 | 45 | 69 | . | . | . | . |
| PM1548-SK1P2 | NOL8 | p.Arg430Lys | missense_variant | MODERATE | 0.111111 | 0 | 36 | 110 | . | . | . | . |
| PM1548-SK1P2 | NOL8 | p.Leu427Gln | missense_variant | MODERATE | 0.147059 | 0 | 34 | 112 | . | . | . | . |
| PM1548-SK1P2 | IPPK | p.Phe34Ser | missense_variant | MODERATE | 0.416667 | 0 | 12 | 158 | . | . | . | . |
| PM1548-SK1P2 | SUSD3 | c.89-6671G>T | intron_variant | MODIFIER | 0.133333 | 0 | 45 | 76 | . | . | . | . |
| PM1548-SK1P2 | WNK2 | p.His758fs | frameshift_variant | HIGH | 0.507163 | 0 | 349 | 278 | . | . | . | . |
| PM1548-SK1P2 | C9orf3 | p.Ala207Thr | missense_variant | MODERATE | 0.027972 | 0 | 143 | 105 | . | . | . | . |
| PM1548-SK1P2 | PTCH1 | p.Thr1064Met | missense_variant | MODERATE | 0.30303 | 0 | 33 | 167 | true | . | . | . |
| PM1548-SK1P2 | ZNF510 | p.Arg592Gly | missense_variant | MODERATE | 0.117647 | 0 | 34 | 140 | . | . | . | . |
| PM1548-SK1P2 | CCDC180 | p.Met439Ile | missense_variant | MODERATE | 0.472973 | 0 | 74 | 95 | . | . | . | . |
| PM1548-SK1P2 | GABBR2 | p.Arg622* | stop_gained | HIGH | 0.545455 | 0 | 11 | 80 | . | . | . | . |
| PM1548-SK1P2 | ANKS6 | p.Ala872Val | missense_variant | MODERATE | 0 | 0.00413223 | 96 | 242 | . | . | . | . |
| PM1548-SK1P2 | SLC44A1 | p.Arg437His | missense_variant | MODERATE | 0.4 | 0 | 5 | 72 | . | . | . | . |
| PM1548-SK1P2 | SVEP1 | p.Ile1787Thr | missense_variant | MODERATE | 0.5 | 0 | 8 | 79 | . | . | . | . |
| PM1548-SK1P2 | TNC | p.Lys1068Asn | missense_variant | MODERATE | 0 | 0 | 71 | 37 | . | . | . | . |
| PM1548-SK1P2 | BRINP1 | p.Arg106His | missense_variant | MODERATE | 0.5 | 0 | 12 | 219 | . | . | . | . |
| PM1548-SK1P2 | FBXW2 | p.Arg385His | missense_variant | MODERATE | 0 | 0 | 51 | 162 | . | . | . | . |

| | | | | | | | | | | | | |
|---|---|---|---|---|---|---|---|---|---|---|---|---|
| PM1548-SK1P2 | FBXW2 | p.Ile377Val | missense_variant | MODERATE | 0 | 0 | 52 | 161 | . | . | . | . |
| PM1548-SK1P2 | DAB2IP | p.His1025fs | frameshift_variant | HIGH | 0.509091 | 0 | 55 | 33 | . | . | . | . |
| PM1548-SK1P2 | LRSAM1 | p.Val385Ile | missense_variant | MODERATE | 0.380952 | 0 | 21 | 115 | . | . | . | . |
| PM1548-SK1P2 | SH2D3C | p.Arg821His | missense_variant | MODERATE | 0.466292 | 0 | 356 | 143 | . | . | . | . |
| PM1548-SK1P2 | C9orf114 | p.Val77Ile | missense_variant | MODERATE | 0.429936 | 0 | 628 | 277 | . | . | . | . |
| PM1548-SK1P2 | C9orf50 | p.Ala252Val | missense_variant | MODERATE | 0.529412 | 0 | 561 | 411 | . | . | . | . |
| PM1548-SK1P2 | NUP214 | p.Arg887His | missense_variant | MODERATE | 0.4 | 0 | 20 | 67 | true | . | . | . |
| PM1548-SK1P2 | ADAMTS13 | p.Arg7fs | frameshift_variant | HIGH | 0.618321 | 0 | 262 | 248 | . | . | . | . |
| PM1548-SK1P2 | SARDH | p.Ala819Val | missense_variant | MODERATE | 0 | 0 | 2 | 41 | . | . | . | . |
| PM1548-SK1P2 | SARDH | p.Arg280Trp | missense_variant | MODERATE | 0 | 0 | 6 | 43 | . | . | . | . |
| PM1548-SK1P2 | LHX3 | p.Ala180Thr | missense_variant | MODERATE | 0.526082 | 0 | 901 | 204 | . | . | . | . |
| PM1548-SK1P2 | MAMDC4 | p.Gln963His | missense_variant | MODERATE | 0.432781 | 0 | 543 | 91 | . | . | . | . |
| PM1548-SK1P2 | AL807752.1 | p.Trp56fs | frameshift_variant | HIGH | 0.428571 | 0 | 42 | 54 | . | . | . | . |
| PM1548-SK1P2 | TPRN | p.Asn499Asp | missense_variant | MODERATE | 0.00113766 | 0 | 879 | 335 | . | . | . | . |
| PM1548-SK1P2 | NSMF | p.Arg366Cys | missense_variant | MODERATE | 0.46747 | 0 | 415 | 90 | . | . | . | . |
| PM1548-SK1P2 | PNPLA7 | p.Leu178Met | missense_variant | MODERATE | 0.121212 | 0 | 33 | 198 | . | . | . | . |
| PM1548-SK1P2 | HDHD1 | p.Arg149Cys | missense_variant | MODERATE | 1 | 0 | 3 | 104 | . | . | . | . |
| PM1548-SK1P2 | EGFL6 | p.Trp5* | stop_gained | HIGH | 0 | 0 | 45 | 81 | . | . | . | . |
| PM1548-SK1P2 | YY2 | p.Glu5Val | missense_variant | MODERATE | 0.0421053 | 0 | 190 | 38 | . | . | . | . |
| PM1548-SK1P2 | TMEM47 | p.Cys104Cys | synonymous_variant | LOW | 0.2 | 0 | 10 | 132 | . | . | . | . |
| PM1548-SK1P2 | USP27X | p.Arg155His | missense_variant | MODERATE | 0 | 0 | 24 | 112 | . | . | . | . |
| PM1548-SK1P2 | KDM5C | p.Glu1003Asp | missense_variant | MODERATE | 0 | 0 | 72 | 54 | true | true | true | . |
| PM1548-SK1P2 | IQSEC2 | p.Arg612Cys | missense_variant | MODERATE | 0 | 0 | 123 | 42 | . | . | . | . |
| PM1548-SK1P2 | FAM120C | p.Gln36_Gln37de | disruptive_inframe_deletion | MODERATE | 1 | 0 | 49 | 126 | . | . | . | . |
| PM1548-SK1P2 | MAGED2 | p.Glu53Asp | missense_variant | MODERATE | 0 | 0 | 241 | 102 | . | . | . | . |
| PM1548-SK1P2 | TRO | p.Ala1132Thr | missense_variant | MODERATE | 0.102564 | 0 | 39 | 69 | . | . | . | . |
| PM1548-SK1P2 | DLG3 | p.Leu333Ile | missense_variant | MODERATE | 0.833333 | 0 | 6 | 34 | . | . | . | . |
| PM1548-SK1P2 | ARMCX4 | p.Tyr2123fs | frameshift_variant | HIGH | 0 | 0 | 49 | 78 | . | . | . | . |
| PM1548-SK1P2 | PAK3 | p.Ala466Val | missense_variant | MODERATE | 1 | 0 | 33 | 37 | . | . | . | . |
| PM1548-SK1P2 | ZCCHC16 | p.Arg297Cys | missense_variant | MODERATE | 0.929577 | 0 | 71 | 98 | . | . | . | . |
| PM1548-SK1P2 | AVPR2 | p.Ala60Val | missense_variant | MODERATE | 0 | 0 | 5 | 59 | . | . | . | . |
| PM1548-SK1P3 | B3GALT6 | p.Glu303Lys | missense_variant | MODERATE | 0.358974 | 0 | 117 | 86 | . | . | . | . |
| PM1548-SK1P3 | SCNN1D | p.Arg344His | missense_variant | MODERATE | 0.431452 | 0 | 248 | 274 | . | . | . | . |
| PM1548-SK1P3 | ATAD3B | p.Arg290Cys | missense_variant | MODERATE | 0.466863 | 0.00687285 | 679 | 291 | . | . | . | . |
| PM1548-SK1P3 | C1orf86 | p.Pro72His | missense_variant | MODERATE | 0.525822 | 0 | 213 | 43 | . | . | . | . |
| PM1548-SK1P3 | PRDM16 | p.Arg759Gln | missense_variant | MODERATE | 0 | 0 | 11 | 40 | true | . | . | . |
| PM1548-SK1P3 | DFFB | p.Tyr290Phe | missense_variant | MODERATE | 0.488372 | 0 | 43 | 203 | . | . | . | . |
| PM1548-SK1P3 | NPHP4 | p.Arg488Gln | missense_variant | MODERATE | 0.519704 | 0 | 406 | 194 | . | . | . | . |
| PM1548-SK1P3 | PLEKHG5 | p.Thr301fs | frameshift_variant | HIGH | 0.501479 | 0 | 676 | 105 | . | . | . | . |
| PM1548-SK1P3 | ZBTB48 | p.Val626Ala | missense_variant | MODERATE | 0.326923 | 0 | 520 | 142 | . | . | . | . |
| PM1548-SK1P3 | KLHL21 | p.Cys250fs | frameshift_variant | HIGH | 0.378947 | 0 | 95 | 136 | . | . | . | . |
| PM1548-SK1P3 | CAMTA1 | p.Ser510Thr | missense_variant | MODERATE | 0.12963 | 0 | 108 | 185 | true | . | . | . |
| PM1548-SK1P3 | PER3 | p.Ala803Ser | missense_variant | MODERATE | 0 | 0 | 76 | 110 | . | . | . | . |
| PM1548-SK1P3 | SLC45A1 | p.Thr364Met | missense_variant | MODERATE | 0.545455 | 0 | 11 | 56 | . | . | . | . |
| PM1548-SK1P3 | SLC45A1 | p.Tyr689* | stop_gained | HIGH | 0.863636 | 0 | 22 | 120 | . | . | . | . |
| PM1548-SK1P3 | RERE | p.Lys83fs | frameshift_variant | HIGH | 1 | 0 | 2 | 49 | . | . | . | . |
| PM1548-SK1P3 | CORT | p.Gln133fs | frameshift_variant | HIGH | 0 | 0 | 48 | 48 | . | . | . | . |
| PM1548-SK1P3 | EXOSC10 | p.Met1fs | frameshift_variant&start_lost | HIGH | 0.521368 | 0 | 234 | 61 | . | . | . | . |
| PM1548-SK1P3 | FHAD1 | p.Ile953fs | frameshift_variant | HIGH | 0.510204 | 0 | 196 | 108 | . | . | . | . |
| PM1548-SK1P3 | PLEKHM2 | p.Arg848Trp | missense_variant | MODERATE | 0 | 0 | 21 | 42 | . | . | . | . |
| PM1548-SK1P3 | SPEN | p.Asn2002fs | frameshift_variant | HIGH | 0.00166389 | 0 | 601 | 173 | true | . | true | true |
| PM1548-SK1P3 | EPHA2 | p.Asn106Ser | missense_variant | MODERATE | 0.515152 | 0 | 33 | 84 | . | . | true | . |
| PM1548-SK1P3 | CROCC | p.Arg1590Cys | missense_variant | MODERATE | 0.220859 | 0 | 326 | 548 | . | . | . | . |
| PM1548-SK1P3 | RCC2 | p.Gly465Arg | missense_variant | MODERATE | 0.694444 | 0 | 72 | 148 | . | . | . | . |
| PM1548-SK1P3 | ARHGEF10L | p.Val965Met | missense_variant | MODERATE | 0.00297324 | 0 | 1009 | 230 | . | . | . | . |
| PM1548-SK1P3 | KLHDC7A | p.Ala652Val | missense_variant | MODERATE | 0 | 0 | 12 | 60 | . | . | . | . |
| PM1548-SK1P3 | RNF186 | p.Ile185Thr | missense_variant | MODERATE | 0 | 0 | 369 | 472 | . | . | . | . |
| PM1548-SK1P3 | KIF17 | p.Arg196His | missense_variant | MODERATE | 0.375 | 0 | 40 | 71 | . | . | . | . |
| PM1548-SK1P3 | EIF4G3 | c.30+5101C>G | intron_variant | MODIFIER | 0.368421 | 0 | 38 | 302 | . | . | . | . |
| PM1548-SK1P3 | HSPG2 | p.Gly2259Asp | missense_variant | MODERATE | 0 | 0 | 8 | 55 | . | . | . | . |
| PM1548-SK1P3 | EPHA8 | p.Ser87Asn | missense_variant | MODERATE | 0 | 0 | 51 | 190 | . | . | . | . |
| PM1548-SK1P3 | AIM1L | n.26671971delA | intragenic_variant | MODIFIER | 0.205882 | 0 | 34 | 55 | . | . | . | . |
| PM1548-SK1P3 | ARID1A | p.Thr1863Ile | missense_variant | MODERATE | 0.105793 | 0 | 397 | 201 | true | true | true | true |
| PM1548-SK1P3 | PTPRU | p.Arg639Trp | missense_variant | MODERATE | 0.008 | 0 | 125 | 60 | . | . | . | . |
| PM1548-SK1P3 | FABP3 | p.Thr54fs | frameshift_variant | HIGH | 0.608696 | 0 | 23 | 207 | . | . | . | . |
| PM1548-SK1P3 | LCK | p.Arg222Cys | missense_variant | MODERATE | 0 | 0 | 123 | 146 | true | . | . | . |
| PM1548-SK1P3 | KIAA1522 | p.Ala943Thr | missense_variant | MODERATE | 0.589744 | 0 | 195 | 198 | . | . | . | . |
| PM1548-SK1P3 | ZNF362 | p.Ala105Val | missense_variant | MODERATE | 0 | 0 | 15 | 295 | . | . | . | . |
| PM1548-SK1P3 | LSM10 | p.Ala2Val | missense_variant | MODERATE | 0.414634 | 0 | 82 | 400 | . | . | . | . |
| PM1548-SK1P3 | CSF3R | p.Tyr508Cys | missense_variant | MODERATE | 0 | 0 | 229 | 167 | true | . | . | . |
| PM1548-SK1P3 | EPHA10 | p.Pro330Ser | missense_variant | MODERATE | 0.498208 | 0 | 279 | 71 | . | . | . | . |
| PM1548-SK1P3 | MACF1 | p.Gln6033Pro | missense_variant&splice_region_variant | MODERATE | 0 | 0 | 3 | 48 | . | . | . | . |
| PM1548-SK1P3 | PABPC4 | p.Ser206Asn | missense_variant | MODERATE | 0.0540541 | 0 | 37 | 90 | . | . | . | . |
| PM1548-SK1P3 | PABPC4 | p.Glu205Gly | missense_variant | MODERATE | 0.0540541 | 0 | 37 | 73 | . | . | . | . |
| PM1548-SK1P3 | ZMPSTE24 | p.Leu362fs | frameshift_variant | HIGH | 0.724138 | 0 | 29 | 53 | . | . | . | . |
| PM1548-SK1P3 | HIVEP3 | p.Pro1037Leu | missense_variant | MODERATE | 0.485714 | 0 | 35 | 35 | . | . | . | . |
| PM1548-SK1P3 | MED8 | p.Asn295fs | frameshift_variant | HIGH | 0.699681 | 0 | 313 | 97 | . | . | . | . |
| PM1548-SK1P3 | HYI | p.Ala55Ser | missense_variant | MODERATE | 0 | 0 | 263 | 158 | . | . | . | . |
| PM1548-SK1P3 | PTPRF | p.Arg1669Trp | missense_variant | MODERATE | 0.507605 | 0 | 526 | 112 | . | . | . | . |
| PM1548-SK1P3 | SLC6A9 | p.Arg662Gln | missense_variant | MODERATE | 0.343478 | 0 | 230 | 99 | . | . | . | . |
| PM1548-SK1P3 | PTCH2 | p.Ala967Val | missense_variant | MODERATE | 0 | 0 | 223 | 194 | . | . | . | . |
| PM1548-SK1P3 | PRDX1 | p.His10Tyr | missense_variant | MODERATE | 0.390244 | 0 | 41 | 149 | . | . | . | . |
| PM1548-SK1P3 | MAST2 | p.Arg1086Arg | synonymous_variant | LOW | 0.270833 | 0 | 96 | 243 | . | . | . | . |
| PM1548-SK1P3 | TAL1 | p.His86Leu | missense_variant | MODERATE | 0 | 0 | 132 | 170 | true | . | . | . |
| PM1548-SK1P3 | LRP8 | p.Asp50Gly | missense_variant | MODERATE | 0.406072 | 0 | 527 | 175 | . | . | . | . |
| PM1548-SK1P3 | CDCP2 | p.Pro209fs | frameshift_variant | HIGH | 0.217391 | 0 | 23 | 107 | . | . | . | . |
| PM1548-SK1P3 | MRPL37 | p.Asp142Glu | missense_variant | MODERATE | 0.0819672 | 0 | 122 | 91 | . | . | . | . |
| PM1548-SK1P3 | CACHD1 | p.Glu211Lys | missense_variant | MODERATE | 0 | 0 | 52 | 147 | . | . | . | . |
| PM1548-SK1P3 | JAK1 | p.Val433Ile | missense_variant | MODERATE | 0.0753012 | 0 | 332 | 203 | true | . | . | true |
| PM1548-SK1P3 | FPGT | p.Asp273Asn | missense_variant | MODERATE | 0 | 0 | 5 | 168 | . | . | . | . |
| PM1548-SK1P3 | LPHN2 | p.Asn759Ile | missense_variant | MODERATE | 0.333333 | 0 | 6 | 85 | . | . | . | . |
| PM1548-SK1P3 | SYDE2 | p.Gly14fs | frameshift_variant | HIGH | 0 | 0 | 80 | 141 | . | . | . | . |
| PM1548-SK1P3 | CLCA1 | p.Arg676Trp | missense_variant | MODERATE | 0.8 | 0 | 5 | 91 | . | . | . | . |
| PM1548-SK1P3 | GBP6 | p.Gly77Asp | missense_variant | MODERATE | 0 | 0 | 8 | 98 | . | . | . | . |
| PM1548-SK1P3 | GFI1 | p.Arg348Met | missense_variant | MODERATE | 0.486891 | 0 | 534 | 270 | . | . | . | . |
| PM1548-SK1P3 | ABCA4 | p.His423Gln | missense_variant | MODERATE | 0.269231 | 0 | 52 | 257 | . | . | . | . |
| PM1548-SK1P3 | ABCA4 | p.Glu422Asp | missense_variant | MODERATE | 0.269231 | 0 | 52 | 256 | . | . | . | . |
| PM1548-SK1P3 | LPPR4 | p.Ala343Thr | missense_variant | MODERATE | 0.540541 | 0 | 74 | 96 | . | . | . | . |
| PM1548-SK1P3 | HIAT1 | p.Ala235Val | missense_variant&splice_region_variant | MODERATE | 0.58 | 0 | 50 | 80 | . | . | . | . |
| PM1548-SK1P3 | TRMT13 | p.Glu307fs | frameshift_variant | HIGH | 0 | 0 | 58 | 85 | . | . | . | . |
| PM1548-SK1P3 | AKNAD1 | p.Leu577Met | missense_variant | MODERATE | 0.391304 | 0 | 23 | 459 | . | . | . | . |
| PM1548-SK1P3 | CELSR2 | p.Asp1795Gly | missense_variant | MODERATE | 0 | 0 | 39 | 362 | . | . | . | . |
| PM1548-SK1P3 | LAMTOR5 | p.Thr39Ala | missense_variant | MODERATE | 0 | 0 | 137 | 96 | . | . | . | . |
| PM1548-SK1P3 | HIPK1 | p.Ala314Thr | missense_variant | MODERATE | 0.5 | 0 | 4 | 273 | . | . | . | . |
| PM1548-SK1P3 | TRIM33 | p.Val961Met | missense_variant | MODERATE | 0.2 | 0 | 5 | 37 | true | . | . | . |
| PM1548-SK1P3 | PDE4DIP | p.Gln2125Lys | missense_variant | MODERATE | 0.205882 | 0 | 68 | 86 | true | . | . | . |
| PM1548-SK1P3 | PDE4DIP | p.Arg759* | stop_gained | HIGH | 0.290801 | 0 | 337 | 104 | true | . | . | . |
| PM1548-SK1P3 | BCL9 | p.Pro517fs | frameshift_variant | HIGH | 0.459259 | 0 | 135 | 142 | true | . | . | . |
| PM1548-SK1P3 | TARS2 | p.Tyr371* | stop_gained | HIGH | 0.170213 | 0 | 47 | 48 | . | . | . | . |
| PM1548-SK1P3 | ADAMTSL4 | p.Gly780fs | frameshift_variant | HIGH | 0.580153 | 0 | 131 | 108 | . | . | . | . |
| PM1548-SK1P3 | ZNF687 | p.Arg885His | missense_variant | MODERATE | 0.490358 | 0 | 726 | 322 | . | . | . | . |
| PM1548-SK1P3 | PI4KB | p.Val202Met | missense_variant | MODERATE | 0 | 0 | 19 | 157 | . | . | . | . |
| PM1548-SK1P3 | CELF3 | p.Val279Ile | missense_variant | MODERATE | 0.583333 | 0 | 24 | 54 | . | . | . | . |
| PM1548-SK1P3 | DENND4B | p.Ala154fs | frameshift_variant | HIGH | 0.0729483 | 0 | 329 | 107 | . | . | . | . |
| PM1548-SK1P3 | CREB3L4 | p.Val263fs | frameshift_variant | HIGH | 0.339506 | 0 | 162 | 42 | . | . | . | . |
| PM1548-SK1P3 | EFNA4 | c.470-101delG | intron_variant | MODIFIER | 0 | 0 | 20 | 82 | . | . | . | . |
| PM1548-SK1P3 | ASH1L | p.Asn1907Lys | missense_variant | MODERATE | 0 | 0 | 71 | 55 | . | . | . | . |
| PM1548-SK1P3 | SMG5 | p.Gln846Lys | missense_variant | MODERATE | 0 | 0 | 37 | 128 | . | . | . | . |
| PM1548-SK1P3 | CD1D | p.Asp140Asn | missense_variant | MODERATE | 0.62963 | 0 | 27 | 198 | . | . | true | . |
| PM1548-SK1P3 | AIM2 | p.Lys342fs | frameshift_variant | HIGH | 0.411765 | 0 | 17 | 33 | . | . | . | . |
| PM1548-SK1P3 | NIT1 | p.Arg261His | missense_variant | MODERATE | 0.34 | 0 | 450 | 402 | . | . | . | . |
| PM1548-SK1P3 | ILDR2 | p.Gly453Ala | missense_variant | MODERATE | 0.0434783 | 0 | 46 | 328 | . | . | . | . |
| PM1548-SK1P3 | ILDR2 | p.Arg415Trp | missense_variant | MODERATE | 0.395683 | 0 | 417 | 213 | . | . | . | . |
| PM1548-SK1P3 | MYOC | p.Arg91Gln | missense_variant | MODERATE | 0 | 0 | 426 | 242 | . | . | . | . |
| PM1548-SK1P3 | DNM3 | p.Glu7Asp | missense_variant | MODERATE | 0.439024 | 0 | 41 | 77 | . | . | . | . |
| PM1548-SK1P3 | DNM3 | p.Phe20Phe | synonymous_variant | LOW | 0.397436 | 0 | 78 | 163 | . | . | . | . |
| PM1548-SK1P3 | DNM3 | p.His85Gln | missense_variant | MODERATE | 0.385827 | 0 | 127 | 56 | . | . | . | . |

| | | | | | | | | | | | | |
|---|---|---|---|---|---|---|---|---|---|---|---|---|
| PM1548-SK1P3 | ANKRD45 | p.Leu39fs | frameshift_variant | HIGH | 0.38961 | 0 | 77 | 229 | . | . | . | . |
| PM1548-SK1P3 | ZBTB37 | p.Val284Ile | missense_variant | MODERATE | 0.285714 | 0 | 49 | 152 | . | . | . | . |
| PM1548-SK1P3 | ZBTB37 | p.Asn288Ser | missense_variant | MODERATE | 0.333333 | 0 | 42 | 160 | . | . | . | . |
| PM1548-SK1P3 | PAPPA2 | p.Thr1174fs | frameshift_variant | HIGH | 0.272727 | 0 | 11 | 36 | . | . | . | . |
| PM1548-SK1P3 | CACNA1E | p.Asn2301Ser | missense_variant | MODERATE | 0.313131 | 0 | 99 | 119 | . | . | . | . |
| PM1548-SK1P3 | CACNA1E | p.Asn2302Asp | missense_variant | MODERATE | 0.316327 | 0 | 98 | 119 | . | . | . | . |
| PM1548-SK1P3 | CACNA1E | p.Asp2310Asp | synonymous_variant | LOW | 0.297872 | 0 | 94 | 89 | . | . | . | . |
| PM1548-SK1P3 | ZNF648 | p.Ser436Tyr | missense_variant | MODERATE | 0.25 | 0 | 40 | 413 | . | . | . | . |
| PM1548-SK1P3 | ZNF648 | p.Ser436Pro | missense_variant | MODERATE | 0.222222 | 0 | 45 | 439 | . | . | . | . |
| PM1548-SK1P3 | ZNF648 | p.Pro423Ser | missense_variant | MODERATE | 0.227273 | 0.00228311 | 44 | 438 | . | . | . | . |
| PM1548-SK1P3 | C1orf27 | p.Thr381Ser | missense_variant | MODERATE | 0.583333 | 0 | 12 | 10 | . | . | . | . |
| PM1548-SK1P3 | PLA2G4A | c.1033+2T>C | splice_donor_variant&intron_variant | HIGH | 0.26087 | 0 | 23 | 80 | . | . | . | . |
| PM1548-SK1P3 | ASPM | p.Arg2700Gln | missense_variant | MODERATE | 0 | 0.00653595 | 113 | 153 | . | . | . | . |
| PM1548-SK1P3 | DENND1B | c.83-20669G>A | intron_variant | MODIFIER | 0.392857 | 0 | 28 | 39 | . | . | . | . |
| PM1548-SK1P3 | NR5A2 | p.Gly169Val | missense_variant | MODERATE | 0.842105 | 0 | 19 | 71 | . | . | . | . |
| PM1548-SK1P3 | NAV1 | p.Thr719Ser | missense_variant | MODERATE | 0.276042 | 0 | 192 | 173 | . | . | . | . |
| PM1548-SK1P3 | NAV1 | p.Thr726Ser | missense_variant | MODERATE | 0.276923 | 0 | 195 | 212 | . | . | . | . |
| PM1548-SK1P3 | ATP2B4 | p.Asp867Asn | missense_variant&splice_region_variant | MODERATE | 0 | 0 | 27 | 121 | . | . | . | . |
| PM1548-SK1P3 | NUAK2 | p.Arg378His | missense_variant | MODERATE | 0.270833 | 0 | 144 | 162 | . | . | . | . |
| PM1548-SK1P3 | PFKFB2 | p.Glu261Ala | missense_variant | MODERATE | 0 | 0 | 81 | 81 | . | . | . | . |
| PM1548-SK1P3 | CR2 | p.Thr209fs | frameshift_variant | HIGH | 0.25 | 0 | 28 | 118 | . | . | . | . |
| PM1548-SK1P3 | SMYD2 | p.Asn99fs | frameshift_variant | HIGH | 0.592593 | 0 | 27 | 181 | . | . | . | . |
| PM1548-SK1P3 | HHIPL2 | p.Gly156Ser | missense_variant | MODERATE | 0.583333 | 0 | 12 | 109 | . | . | . | . |
| PM1548-SK1P3 | MIA3 | p.Arg1545Gln | missense_variant | MODERATE | 0.44 | 0 | 25 | 39 | . | . | . | . |
| PM1548-SK1P3 | LIN9 | p.Arg245Trp | missense_variant | MODERATE | 0.5 | 0 | 6 | 86 | . | . | . | . |
| PM1548-SK1P3 | ITPKB | p.Arg331His | missense_variant | MODERATE | 0.442688 | 0 | 253 | 395 | . | . | true | . |
| PM1548-SK1P3 | ITPKB | p.Leu42Val | missense_variant | MODERATE | 0.228571 | 0 | 70 | 35 | . | . | true | . |
| PM1548-SK1P3 | SNAP47 | p.Phe343Leu | missense_variant | MODERATE | 0 | 0 | 154 | 78 | . | . | . | . |
| PM1548-SK1P3 | IBA57 | p.Ala161Val | missense_variant | MODERATE | 0.329114 | 0 | 79 | 268 | . | . | . | . |
| PM1548-SK1P3 | OBSCN | p.Tyr2035Phe | missense_variant | MODERATE | 0.267974 | 0 | 459 | 507 | . | . | . | . |
| PM1548-SK1P3 | OBSCN | p.Gln2065Lys | missense_variant | MODERATE | 0.284672 | 0 | 274 | 171 | . | . | . | . |
| PM1548-SK1P3 | OBSCN | p.Cys2073Trp | missense_variant | MODERATE | 0.616162 | 0 | 99 | 54 | . | . | . | . |
| PM1548-SK1P3 | ACTA1 | p.Thr231Met | missense_variant | MODERATE | 0 | 0 | 18 | 311 | . | . | . | . |
| PM1548-SK1P3 | GNPAT | p.Val551Met | missense_variant | MODERATE | 0.30303 | 0 | 33 | 79 | . | . | . | . |
| PM1548-SK1P3 | DISC1 | p.Asp580Gly | missense_variant | MODERATE | 0.611111 | 0 | 36 | 167 | . | . | . | . |
| PM1548-SK1P3 | GGPS1 | p.Val112Met | missense_variant | MODERATE | 0.4 | 0 | 65 | 405 | . | . | . | . |
| PM1548-SK1P3 | NID1 | p.Pro918fs | frameshift_variant&splice_region_variant | HIGH | 0.482143 | 0 | 56 | 46 | . | . | . | . |
| PM1548-SK1P3 | CEP170 | p.Glu1460Lys | missense_variant | MODERATE | 0.166667 | 0 | 12 | 32 | . | . | . | . |
| PM1548-SK1P3 | ZNF496 | p.Arg606His | missense_variant | MODERATE | 0.483051 | 0 | 118 | 143 | . | . | . | . |
| PM1548-SK1P3 | OR2T4 | p.Cys60Ser | missense_variant | MODERATE | 0 | 0 | 5 | 107 | . | . | . | . |
| PM1548-SK1P3 | OR2T6 | p.Tyr60His | missense_variant | MODERATE | 0.58 | 0 | 100 | 68 | . | . | . | . |
| PM1548-SK1P3 | DIP2C | p.Val1150Ile | missense_variant | MODERATE | 0 | 0 | 49 | 183 | . | . | . | . |
| PM1548-SK1P3 | WDR37 | p.Pro169Ser | missense_variant | MODERATE | 0.322581 | 0 | 62 | 220 | . | . | . | . |
| PM1548-SK1P3 | CALML5 | p.Glu102Lys | missense_variant | MODERATE | 0.615385 | 0 | 39 | 79 | . | . | . | . |
| PM1548-SK1P3 | FRMD4A | p.Pro1005fs | frameshift_variant | HIGH | 0 | 0 | 21 | 118 | . | . | . | . |
| PM1548-SK1P3 | CUBN | p.Trp3323Arg | missense_variant | MODERATE | 0 | 0 | 44 | 170 | . | . | . | . |
| PM1548-SK1P3 | GPR158 | p.Arg448Gln | missense_variant | MODERATE | 0.272727 | 0 | 22 | 52 | . | . | . | . |
| PM1548-SK1P3 | SVIL | p.Arg2075Cys | missense_variant | MODERATE | 0.461538 | 0 | 13 | 40 | . | . | . | . |
| PM1548-SK1P3 | PARD3 | p.Arg1278Gly | missense_variant | MODERATE | 0 | 0 | 73 | 91 | . | . | . | . |
| PM1548-SK1P3 | FRMPD2 | p.Pro226Leu | missense_variant | MODERATE | 0.777778 | 0 | 63 | 178 | . | . | . | . |
| PM1548-SK1P3 | PRKG1 | p.Asn601fs | frameshift_variant | HIGH | 0.345238 | 0 | 84 | 55 | . | . | . | . |
| PM1548-SK1P3 | PBLD | p.Asp167Tyr | missense_variant | MODERATE | 0 | 0 | 56 | 186 | . | . | . | . |
| PM1548-SK1P3 | DDX50 | p.Tyr39Cys | missense_variant | MODERATE | 0.4 | 0 | 15 | 39 | . | . | . | . |
| PM1548-SK1P3 | ADAMTS14 | p.Gly710Arg | missense_variant | MODERATE | 0.511111 | 0 | 45 | 78 | . | . | . | . |
| PM1548-SK1P3 | DDIT4 | p.Ser46Cys | missense_variant | MODERATE | 0.172414 | 0 | 87 | 263 | . | . | . | . |
| PM1548-SK1P3 | DDIT4 | p.Thr47Ala | missense_variant | MODERATE | 0.172414 | 0 | 87 | 263 | . | . | . | . |
| PM1548-SK1P3 | KAT6B | p.Arg463His | missense_variant | MODERATE | 0.479839 | 0 | 248 | 104 | true | . | . | . |
| PM1548-SK1P3 | KCNMA1 | p.Ala1039Val | missense_variant | MODERATE | 0.4 | 0 | 15 | 74 | . | . | . | . |
| PM1548-SK1P3 | ZMIZ1 | p.Asp1051fs | frameshift_variant | HIGH | 0.463768 | 0 | 69 | 67 | . | . | . | . |
| PM1548-SK1P3 | MAT1A | p.Arg81Gln | missense_variant | MODERATE | 0.333333 | 0 | 21 | 225 | . | . | . | . |
| PM1548-SK1P3 | TSPAN14 | p.Asp190Gly | missense_variant | MODERATE | 0 | 0 | 44 | 134 | . | . | . | . |
| PM1548-SK1P3 | SH2D4B | p.Ala186Val | missense_variant | MODERATE | 0.325843 | 0 | 89 | 42 | . | . | . | . |
| PM1548-SK1P3 | LRIT2 | p.Thr520fs | frameshift_variant | HIGH | 0.273381 | 0 | 139 | 224 | . | . | . | . |
| PM1548-SK1P3 | CCSER2 | c.2326-13521C> | intron_variant | MODIFIER | 0.258065 | 0 | 31 | 176 | . | . | . | . |
| PM1548-SK1P3 | CYP26C1 | p.Ser70Leu | missense_variant | MODERATE | 0 | 0 | 84 | 149 | . | . | . | . |
| PM1548-SK1P3 | MYOF | p.Arg726Gln | missense_variant | MODERATE | 0.6 | 0 | 25 | 43 | . | . | . | . |
| PM1548-SK1P3 | SORBS1 | p.Arg585Cys | missense_variant | MODERATE | 0.642857 | 0 | 28 | 42 | . | . | . | . |
| PM1548-SK1P3 | ALDH18A1 | p.Arg622Trp | missense_variant | MODERATE | 0.636364 | 0 | 22 | 86 | . | . | . | . |
| PM1548-SK1P3 | ENTPD1 | p.Arg252Leu | missense_variant | MODERATE | 0 | 0 | 68 | 84 | . | . | . | . |
| PM1548-SK1P3 | MORN4 | p.Leu202Ile | missense_variant | MODERATE | 0 | 0 | 33 | 87 | . | . | . | . |
| PM1548-SK1P3 | PKD2L1 | p.Arg206Cys | missense_variant | MODERATE | 0.468619 | 0 | 239 | 130 | . | . | . | . |
| PM1548-SK1P3 | MRPL43 | p.Asn66Ser | missense_variant | MODERATE | 0.437326 | 0 | 359 | 81 | . | . | . | . |
| PM1548-SK1P3 | POLL | p.Arg419* | stop_gained | HIGH | 0 | 0 | 23 | 44 | . | . | . | . |
| PM1548-SK1P3 | TAF5 | p.Phe374fs | frameshift_variant | HIGH | 0.285714 | 0 | 14 | 50 | . | . | . | . |
| PM1548-SK1P3 | WDR96 | p.Val70Ala | missense_variant | MODERATE | 0 | 0 | 3 | 59 | . | . | . | . |
| PM1548-SK1P3 | SORCS3 | p.Lys1023Gln | missense_variant | MODERATE | 0.285714 | 0 | 42 | 58 | . | . | . | . |
| PM1548-SK1P3 | TCF7L2 | p.Ser587Leu | missense_variant | MODERATE | 0.108333 | 0 | 120 | 171 | true | . | true | . |
| PM1548-SK1P3 | ABLIM1 | p.Leu633Met | missense_variant | MODERATE | 0 | 0 | 97 | 266 | . | . | . | . |
| PM1548-SK1P3 | GFRA1 | p.Ala173Val | missense_variant | MODERATE | 0.75 | 0 | 4 | 73 | . | . | . | . |
| PM1548-SK1P3 | KIAA1598 | p.Arg482His | missense_variant | MODERATE | 0.292683 | 0 | 41 | 181 | true | . | . | . |
| PM1548-SK1P3 | VAX1 | p.Cys200* | stop_gained | HIGH | 0 | 0 | 160 | 96 | . | . | . | . |
| PM1548-SK1P3 | EIF3A | p.His257His | synonymous_variant | LOW | 0.0754717 | 0 | 53 | 131 | . | . | . | . |
| PM1548-SK1P3 | ATE1 | p.Leu289Gln | missense_variant | MODERATE | 0.444444 | 0 | 36 | 77 | . | . | . | . |
| PM1548-SK1P3 | PLEKHA1 | p.Arg9His | missense_variant | MODERATE | 0.136364 | 0 | 22 | 43 | . | . | . | . |
| PM1548-SK1P3 | PLEKHA1 | p.Val404Met | missense_variant | MODERATE | 0.114754 | 0 | 61 | 76 | . | . | . | . |
| PM1548-SK1P3 | HMX3 | p.Pro23fs | frameshift_variant | HIGH | 0.637931 | 0 | 116 | 56 | . | . | . | . |
| PM1548-SK1P3 | CLRN3 | p.Pro71Thr | missense_variant | MODERATE | 0.0346821 | 0.00613497 | 173 | 163 | . | . | . | . |
| PM1548-SK1P3 | MKI67 | p.Arg2258* | stop_gained | HIGH | 0.542857 | 0 | 35 | 90 | . | . | . | . |
| PM1548-SK1P3 | TTC40 | p.Thr1864Met | missense_variant | MODERATE | 0.589286 | 0 | 56 | 72 | . | . | . | . |
| PM1548-SK1P3 | KNDC1 | p.Cys79fs | frameshift_variant | HIGH | 0.347826 | 0 | 23 | 162 | . | . | . | . |
| PM1548-SK1P3 | PIDD | p.Ala727Glu | missense_variant | MODERATE | 0 | 0 | 188 | 108 | . | . | . | . |
| PM1548-SK1P3 | MUC6 | p.Thr1861Ala | missense_variant | MODERATE | 0.714286 | 0 | 63 | 78 | . | . | . | . |
| PM1548-SK1P3 | MUC6 | p.Ile1857Thr | missense_variant | MODERATE | 0.761905 | 0 | 63 | 78 | . | . | . | . |
| PM1548-SK1P3 | MUC6 | p.His1752Gln | missense_variant | MODERATE | 0.0355987 | 0 | 309 | 31 | . | . | . | . |
| PM1548-SK1P3 | MUC6 | p.Thr1711Ile | missense_variant | MODERATE | 0.125 | 0 | 128 | 51 | . | . | . | . |
| PM1548-SK1P3 | MUC2 | p.Gly1331Arg | missense_variant | MODERATE | 0.554054 | 0 | 74 | 70 | . | . | . | . |
| PM1548-SK1P3 | MUC5B | p.Ala110Thr | missense_variant | MODERATE | 0.658537 | 0 | 246 | 216 | . | . | . | . |
| PM1548-SK1P3 | BRSK2 | p.Pro440fs | frameshift_variant | HIGH | 0.428449 | 0 | 1167 | 593 | . | . | . | . |
| PM1548-SK1P3 | TSPAN32 | p.Ala178Val | missense_variant | MODERATE | 0.443902 | 0 | 205 | 69 | . | . | . | . |
| PM1548-SK1P3 | TSSC4 | p.Arg92Cys | missense_variant | MODERATE | 0.483704 | 0 | 1350 | 524 | . | . | . | . |
| PM1548-SK1P3 | OSBPL5 | p.Thr843Met | missense_variant | MODERATE | 0 | 0 | 19 | 106 | . | . | . | . |
| PM1548-SK1P3 | STIM1 | p.Lys371Arg | missense_variant | MODERATE | 0.340206 | 0 | 97 | 319 | . | . | . | . |
| PM1548-SK1P3 | OR51E1 | p.Tyr63His | missense_variant | MODERATE | 0.144522 | 0 | 429 | 116 | . | . | . | . |
| PM1548-SK1P3 | OR51A2 | p.Lys289Asn | missense_variant | MODERATE | 0.025641 | 0 | 39 | 30 | . | . | . | . |
| PM1548-SK1P3 | SOX6 | p.Met803Ile | missense_variant | MODERATE | 0.105263 | 0 | 76 | 33 | . | . | . | . |
| PM1548-SK1P3 | SOX6 | p.Met803Thr | missense_variant | MODERATE | 0.0921053 | 0 | 76 | 33 | . | . | . | . |
| PM1548-SK1P3 | SOX6 | p.Met803Val | missense_variant | MODERATE | 0.0921053 | 0 | 76 | 33 | . | . | . | . |
| PM1548-SK1P3 | ABCC8 | p.Asp761Asn | missense_variant | MODERATE | 0.361702 | 0 | 94 | 230 | . | . | . | . |
| PM1548-SK1P3 | LDHC | p.Lys5Glu | missense_variant | MODERATE | 0.428571 | 0 | 14 | 119 | . | . | . | . |
| PM1548-SK1P3 | MRGPRX2 | p.Asn62Asn | synonymous_variant | LOW | 0.397849 | 0 | 93 | 83 | . | . | . | . |
| PM1548-SK1P3 | ZDHHC13 | p.Gly591Asp | missense_variant | MODERATE | 0 | 0 | 2 | 33 | . | . | . | . |
| PM1548-SK1P3 | ANO3 | p.Leu273Pro | missense_variant | MODERATE | 0.5625 | 0 | 64 | 432 | . | . | . | . |
| PM1548-SK1P3 | HIPK3 | p.Asp532Ala | missense_variant | MODERATE | 0.0290456 | 0 | 241 | 54 | . | . | . | . |
| PM1548-SK1P3 | ABTB2 | p.Asp793Asn | missense_variant | MODERATE | 0.470852 | 0.00668896 | 223 | 299 | . | . | . | . |
| PM1548-SK1P3 | RAG1 | p.Arg507Trp | missense_variant | MODERATE | 0.453704 | 0 | 216 | 86 | . | . | . | true |
| PM1548-SK1P3 | TP53I11 | p.Leu62Arg | missense_variant | MODERATE | 0.00148368 | 0 | 674 | 34 | . | . | . | . |
| PM1548-SK1P3 | MAPK8IP1 | p.Thr321Met | missense_variant | MODERATE | 0.63745 | 0 | 251 | 188 | . | true | . | . |
| PM1548-SK1P3 | DGKZ | n.46369267G>A | intragenic_variant | MODIFIER | 0 | 0 | 31 | 32 | . | . | . | . |
| PM1548-SK1P3 | DGKZ | p.Asp574Asn | missense_variant | MODERATE | 0.481013 | 0 | 79 | 74 | . | . | . | . |
| PM1548-SK1P3 | ACP2 | p.Arg231His | missense_variant | MODERATE | 0.451264 | 0 | 277 | 199 | . | . | . | . |
| PM1548-SK1P3 | MYBPC3 | p.Arg654Ser | missense_variant | MODERATE | 0.266667 | 0 | 30 | 97 | . | . | . | . |
| PM1548-SK1P3 | OR5D14 | p.Tyr261Cys | missense_variant | MODERATE | 0.3 | 0 | 10 | 71 | . | . | . | . |
| PM1548-SK1P3 | OR5W2 | p.Ala29Ser | missense_variant | MODERATE | 0.305556 | 0 | 36 | 99 | . | . | . | . |
| PM1548-SK1P3 | APLNR | p.Ala161Thr | missense_variant | MODERATE | 0.157143 | 0 | 70 | 160 | . | . | . | . |
| PM1548-SK1P3 | OR10Q1 | p.Thr287Ile | missense_variant | MODERATE | 0.555556 | 0 | 27 | 90 | . | . | . | . |
| PM1548-SK1P3 | DTX4 | p.Lys396fs | frameshift_variant | HIGH | 0.462025 | 0 | 158 | 34 | . | . | . | . |
| PM1548-SK1P3 | DTX4 | p.Gly510fs | frameshift_variant | HIGH | 0.397849 | 0.00719424 | 93 | 139 | . | . | . | . |

| | | | | | | | | | | | | |
|---|---|---|---|---|---|---|---|---|---|---|---|---|
| PM1548-SK1P3 | DAK | p.Arg380Gln | missense_variant | MODERATE | 0.369963 | 0 | 546 | 316 | . | . | . | . |
| PM1548-SK1P3 | FEN1 | p.Arg332Cys | missense_variant | MODERATE | 0 | 0 | 37 | 80 | . | . | . | . |
| PM1548-SK1P3 | BEST1 | p.Ser413Asn | missense_variant | MODERATE | 0.734375 | 0 | 64 | 159 | . | . | . | . |
| PM1548-SK1P3 | AHNAK | p.Ser4908Leu | missense_variant | MODERATE | 0.498008 | 0 | 502 | 39 | . | . | . | . |
| PM1548-SK1P3 | AHNAK | p.His4772Asn | missense_variant | MODERATE | 0 | 0 | 172 | 54 | . | . | . | . |
| PM1548-SK1P3 | AHNAK | p.Gly4561Asp | missense_variant | MODERATE | 0.304762 | 0 | 315 | 206 | . | . | . | . |
| PM1548-SK1P3 | AHNAK | p.Val4560Leu | missense_variant | MODERATE | 0.298413 | 0 | 315 | 203 | . | . | . | . |
| PM1548-SK1P3 | AHNAK | p.Val3640Ile | missense_variant | MODERATE | 0.634615 | 0 | 52 | 44 | . | . | . | . |
| PM1548-SK1P3 | AHNAK | p.Asn3637Lys | missense_variant | MODERATE | 0.55814 | 0 | 43 | 44 | . | . | . | . |
| PM1548-SK1P3 | AHNAK | p.Lys2706fs | frameshift_variant | HIGH | 0.544803 | 0 | 279 | 43 | . | . | . | . |
| PM1548-SK1P3 | AHNAK | p.Thr1986Ala | missense_variant | MODERATE | 0.135802 | 0 | 81 | 324 | . | . | . | . |
| PM1548-SK1P3 | CHRM1 | p.Pro319fs | frameshift_variant | HIGH | 0 | 0 | 5 | 31 | . | . | . | . |
| PM1548-SK1P3 | SLC22A6 | p.Val240Met | missense_variant | MODERATE | 0.263158 | 0 | 38 | 226 | . | . | . | . |
| PM1548-SK1P3 | SLC22A25 | p.Asn62Asp | missense_variant | MODERATE | 0 | 0 | 77 | 110 | . | . | . | . |
| PM1548-SK1P3 | C11orf95 | n.1245C>T | non_coding_exon_variant | MODIFIER | 0.166667 | 0 | 12 | 34 | . | . | . | . |
| PM1548-SK1P3 | NRXN2 | p.Asp277Asn | missense_variant | MODERATE | 0.27451 | 0 | 51 | 222 | . | . | . | . |
| PM1548-SK1P3 | SNX15 | p.Leu10Met | missense_variant | MODERATE | 0 | 0 | 12 | 37 | . | . | . | . |
| PM1548-SK1P3 | RBM4 | p.Tyr303His | missense_variant | MODERATE | 0.285714 | 0 | 35 | 269 | . | . | . | . |
| PM1548-SK1P3 | SPTBN2 | p.Cys1393Ser | missense_variant | MODERATE | 0.276423 | 0 | 123 | 299 | . | . | . | . |
| PM1548-SK1P3 | SPTBN2 | p.Arg465Gln | missense_variant | MODERATE | 0.254902 | 0 | 51 | 212 | . | . | . | . |
| PM1548-SK1P3 | LRP5 | p.Ala724Val | missense_variant | MODERATE | 1 | 0 | 5 | 227 | . | . | . | . |
| PM1548-SK1P3 | IGHMBP2 | p.Pro809Ser | missense_variant | MODERATE | 0.492958 | 0 | 71 | 132 | . | . | . | . |
| PM1548-SK1P3 | TPCN2 | p.Gly492Val | missense_variant | MODERATE | 0 | 0 | 48 | 21 | . | . | . | . |
| PM1548-SK1P3 | CCND1 | p.Val94Leu | missense_variant | MODERATE | 0.194514 | 0 | 401 | 258 | true | true | true | . |
| PM1548-SK1P3 | INPPL1 | p.Leu957Phe | missense_variant | MODERATE | 0.285714 | 0 | 56 | 30 | . | . | . | . |
| PM1548-SK1P3 | PDE2A | p.His99fs | frameshift_variant | HIGH | 0.615789 | 0 | 760 | 160 | . | . | . | . |
| PM1548-SK1P3 | RNF169 | p.Glu131* | stop_gained | HIGH | 0 | 0 | 22 | 52 | . | . | . | . |
| PM1548-SK1P3 | TENM4 | p.Ala235Thr | missense_variant | MODERATE | 0.644068 | 0 | 59 | 201 | . | . | . | . |
| PM1548-SK1P3 | PCF11 | p.Leu745Leu | synonymous_variant | LOW | 0.347222 | 0 | 144 | 203 | . | . | . | . |
| PM1548-SK1P3 | PCF11 | p.Ser756Gly | missense_variant | MODERATE | 0.324503 | 0 | 151 | 215 | . | . | . | . |
| PM1548-SK1P3 | PCF11 | p.Arg833Trp | missense_variant | MODERATE | 0.467444 | 0 | 1029 | 258 | . | . | . | . |
| PM1548-SK1P3 | TYR | p.Gly191fs | frameshift_variant | HIGH | 0.588235 | 0 | 17 | 94 | . | . | . | . |
| PM1548-SK1P3 | FAT3 | p.Ala2495Ser | missense_variant | MODERATE | 0.0169492 | 0 | 118 | 229 | . | . | . | . |
| PM1548-SK1P3 | PANX1 | p.Ser250Thr | missense_variant | MODERATE | 0.310345 | 0 | 29 | 236 | . | . | . | . |
| PM1548-SK1P3 | MTMR2 | p.Lys451Glu | missense_variant | MODERATE | 0.0886076 | 0 | 158 | 77 | . | . | . | . |
| PM1548-SK1P3 | MTMR2 | p.Asp436Asn | missense_variant | MODERATE | 0.125828 | 0 | 151 | 75 | . | . | . | . |
| PM1548-SK1P3 | MTMR2 | p.Leu433Val | missense_variant | MODERATE | 0.0802469 | 0 | 162 | 282 | . | . | . | . |
| PM1548-SK1P3 | MMP3 | p.Asp200Gly | missense_variant | MODERATE | 0 | 0 | 55 | 91 | . | . | . | . |
| PM1548-SK1P3 | DYNC2H1 | p.Tyr3523His | missense_variant | MODERATE | 0.73913 | 0 | 46 | 62 | . | . | . | . |
| PM1548-SK1P3 | CASP1 | p.Phe262Tyr | missense_variant | MODERATE | 0.294118 | 0 | 17 | 177 | . | . | . | . |
| PM1548-SK1P3 | KBTBD3 | p.Lys478fs | frameshift_variant | HIGH | 0.535354 | 0 | 99 | 50 | . | . | . | . |
| PM1548-SK1P3 | NXPE1 | p.Thr169Ile | missense_variant | MODERATE | 0.416667 | 0 | 48 | 97 | . | . | . | . |
| PM1548-SK1P3 | NXPE1 | p.Met157Ile | missense_variant | MODERATE | 0.787234 | 0 | 47 | 97 | . | . | . | . |
| PM1548-SK1P3 | NXPE1 | p.Met157Thr | missense_variant | MODERATE | 0.791667 | 0 | 48 | 95 | . | . | . | . |
| PM1548-SK1P3 | NXPE1 | p.Thr149Lys | missense_variant | MODERATE | 0.72 | 0 | 50 | 118 | . | . | . | . |
| PM1548-SK1P3 | DSCAML1 | p.Val1762Ile | missense_variant | MODERATE | 0.447917 | 0 | 96 | 226 | . | . | . | . |
| PM1548-SK1P3 | DSCAML1 | p.Asp949fs | frameshift_variant | HIGH | 0.616822 | 0 | 428 | 244 | . | . | . | . |
| PM1548-SK1P3 | ATP5L | p.Leu84Trp | missense_variant | MODERATE | 0.333333 | 0 | 120 | 125 | . | . | . | . |
| PM1548-SK1P3 | ATP5L | p.Val89Ile | missense_variant | MODERATE | 0.322581 | 0 | 124 | 139 | . | . | . | . |
| PM1548-SK1P3 | ATP5L | p.Ile99Val | missense_variant | MODERATE | 0.325203 | 0 | 123 | 141 | . | . | . | . |
| PM1548-SK1P3 | PHLDB1 | p.Ala932Ser | missense_variant | MODERATE | 0.0277778 | 0 | 72 | 223 | . | . | . | . |
| PM1548-SK1P3 | PHLDB1 | p.Pro938Leu | missense_variant | MODERATE | 0.028169 | 0 | 71 | 220 | . | . | . | . |
| PM1548-SK1P3 | CBL | p.Ser237Leu | missense_variant | MODERATE | 0.25 | 0 | 16 | 128 | true | . | . | . |
| PM1548-SK1P3 | CBL | p.Pro504Gln | missense_variant | MODERATE | 0.108108 | 0 | 74 | 111 | true | . | . | . |
| PM1548-SK1P3 | CRTAM | p.Leu60fs | frameshift_variant | HIGH | 0.462085 | 0 | 422 | 169 | . | . | . | . |
| PM1548-SK1P3 | OR10G8 | p.Lys265Glu | missense_variant | MODERATE | 0.211454 | 0 | 227 | 232 | . | . | . | . |
| PM1548-SK1P3 | OR10G8 | p.Lys265Asn | missense_variant | MODERATE | 0.219731 | 0 | 223 | 193 | . | . | . | . |
| PM1548-SK1P3 | OR8G2 | p.Val2Gly | missense_variant | MODERATE | 0 | 0 | 3 | 1 | . | . | . | . |
| PM1548-SK1P3 | ROBO4 | p.Arg776His | missense_variant | MODERATE | 0.517483 | 0 | 143 | 263 | . | . | . | . |
| PM1548-SK1P3 | ADAMTS8 | p.Arg9Gly | missense_variant | MODERATE | 0 | 0 | 11 | 48 | . | . | . | true |
| PM1548-SK1P3 | IQSEC3 | p.Gln1075Lys | missense_variant | MODERATE | 0 | 0 | 106 | 59 | . | . | . | . |
| PM1548-SK1P3 | FOXM1 | p.Leu334Ser | missense_variant | MODERATE | 0.178571 | 0 | 56 | 177 | . | . | . | . |
| PM1548-SK1P3 | FOXM1 | p.Asp328Glu | missense_variant | MODERATE | 0.204082 | 0 | 49 | 203 | . | . | . | . |
| PM1548-SK1P3 | VWF | p.Ala865Asp | missense_variant | MODERATE | 0.363309 | 0 | 278 | 226 | . | . | . | . |
| PM1548-SK1P3 | IFFO1 | p.Val126Met | missense_variant | MODERATE | 0.451327 | 0 | 113 | 191 | . | . | . | . |
| PM1548-SK1P3 | PHB2 | p.Arg270Arg | synonymous_variant | LOW | 0.133779 | 0 | 299 | 405 | . | . | . | . |
| PM1548-SK1P3 | PHB2 | p.Asn269Asn | synonymous_variant | LOW | 0.133779 | 0 | 299 | 404 | . | . | . | . |
| PM1548-SK1P3 | TAS2R10 | p.Ser41Thr | missense_variant | MODERATE | 0.584906 | 0 | 53 | 57 | . | . | . | . |
| PM1548-SK1P3 | TAS2R31 | p.Ala177Thr | missense_variant | MODERATE | 0 | 0 | 15 | 47 | . | . | . | . |
| PM1548-SK1P3 | PLCZ1 | p.Gly133Val | missense_variant | MODERATE | 0 | 0 | 22 | 42 | . | . | . | . |
| PM1548-SK1P3 | PDE3A | p.Glu171fs | frameshift_variant | HIGH | 0.111111 | 0 | 54 | 56 | . | . | . | . |
| PM1548-SK1P3 | TM7SF3 | c.869-699C>G | intron_variant | MODIFIER | 0.612903 | 0 | 93 | 125 | . | . | . | . |
| PM1548-SK1P3 | DDX11 | p.Asp935Asn | missense_variant | MODERATE | 0.428571 | 0 | 7 | 35 | . | . | . | . |
| PM1548-SK1P3 | COL2A1 | p.Asp682Asn | missense_variant | MODERATE | 0.598291 | 0 | 468 | 53 | true | . | . | . |
| PM1548-SK1P3 | KMT2D | p.Ala4804Val | missense_variant | MODERATE | 0.580645 | 0.0060241 | 62 | 166 | true | true | true | true |
| PM1548-SK1P3 | KMT2D | p.Leu4214Leu | synonymous_variant | LOW | 0.272152 | 0 | 158 | 373 | true | true | true | true |
| PM1548-SK1P3 | KMT2D | p.Thr3166Ile | missense_variant | MODERATE | 0.420513 | 0 | 195 | 193 | true | true | true | true |
| PM1548-SK1P3 | KMT2D | p.Thr2581Ala | missense_variant | MODERATE | 0.104348 | 0 | 345 | 133 | true | true | true | true |
| PM1548-SK1P3 | KMT2D | p.Gly1628fs | frameshift_variant | HIGH | 0.133333 | 0 | 60 | 136 | true | true | true | true |
| PM1548-SK1P3 | PRPH | p.Ala348Thr | missense_variant | MODERATE | 0.55625 | 0 | 160 | 109 | . | . | . | . |
| PM1548-SK1P3 | FAM186B | p.Leu494Met | missense_variant | MODERATE | 0.526846 | 0 | 298 | 412 | . | . | . | . |
| PM1548-SK1P3 | ANKRD33 | p.Ala173Val | missense_variant | MODERATE | 0.370787 | 0 | 445 | 241 | . | . | . | . |
| PM1548-SK1P3 | KRT74 | p.Gln440His | missense_variant | MODERATE | 0 | 0 | 6 | 34 | . | . | . | . |
| PM1548-SK1P3 | PCBP2 | p.Pro175fs | frameshift_variant | HIGH | 0.545455 | 0 | 11 | 35 | . | . | . | . |
| PM1548-SK1P3 | HNRNPA1 | p.Thr61Ala | missense_variant | MODERATE | 0.404762 | 0 | 42 | 168 | . | . | . | . |
| PM1548-SK1P3 | ITGA5 | p.Arg193His | missense_variant | MODERATE | 0.110236 | 0 | 127 | 104 | . | . | . | . |
| PM1548-SK1P3 | MYO1A | p.Arg559Cys | missense_variant | MODERATE | 0.415094 | 0 | 424 | 190 | . | . | . | . |
| PM1548-SK1P3 | NAB2 | p.Pro388Thr | missense_variant | MODERATE | 0.136986 | 0 | 73 | 78 | true | . | . | . |
| PM1548-SK1P3 | NAB2 | p.Pro388His | missense_variant | MODERATE | 0.136986 | 0 | 73 | 78 | true | . | . | . |
| PM1548-SK1P3 | R3HDM2 | p.Arg956His | missense_variant | MODERATE | 0 | 0 | 16 | 33 | . | . | . | . |
| PM1548-SK1P3 | CTDSP2 | p.Glu259Asp | missense_variant | MODERATE | 0.355556 | 0 | 45 | 90 | . | . | . | . |
| PM1548-SK1P3 | CTDSP2 | p.Ala258Thr | missense_variant | MODERATE | 0.354167 | 0 | 48 | 88 | . | . | . | . |
| PM1548-SK1P3 | CTDSP2 | p.Ile251Val | missense_variant | MODERATE | 0.45 | 0 | 40 | 94 | . | . | . | . |
| PM1548-SK1P3 | PTPRB | p.Val1992Ala | missense_variant | MODERATE | 0.50495 | 0 | 101 | 50 | true | . | . | . |
| PM1548-SK1P3 | NAV3 | p.Met1275Val | missense_variant | MODERATE | 0.459016 | 0 | 61 | 67 | . | true | . | . |
| PM1548-SK1P3 | NAV3 | p.Pro1276Leu | missense_variant | MODERATE | 0.57 | 0 | 100 | 130 | . | true | . | . |
| PM1548-SK1P3 | NAV3 | p.Thr1281Ser | missense_variant | MODERATE | 0.564356 | 0 | 101 | 144 | . | true | . | . |
| PM1548-SK1P3 | TMTC2 | p.Gly143Arg | missense_variant | MODERATE | 0.515152 | 0 | 33 | 67 | . | . | . | . |
| PM1548-SK1P3 | ALX1 | p.Thr68Ser | missense_variant | MODERATE | 0.25 | 0 | 4 | 191 | . | . | . | . |
| PM1548-SK1P3 | FGD6 | p.Met965fs | frameshift_variant | HIGH | 0.764706 | 0 | 17 | 56 | . | . | . | . |
| PM1548-SK1P3 | APAF1 | p.Gln578Arg | missense_variant | MODERATE | 0.25 | 0 | 68 | 126 | . | . | . | . |
| PM1548-SK1P3 | PRDM4 | p.Glu313Glu | synonymous_variant | LOW | 0.263158 | 0 | 76 | 274 | . | . | . | . |
| PM1548-SK1P3 | TMEM119 | p.Thr81fs | frameshift_variant | HIGH | 0.127119 | 0 | 1652 | 297 | . | . | . | . |
| PM1548-SK1P3 | FOXN4 | p.Ala462Thr | missense_variant | MODERATE | 0.536585 | 0.00490196 | 41 | 204 | . | . | . | . |
| PM1548-SK1P3 | FAM222A | p.Pro392Thr | missense_variant | MODERATE | 0.305085 | 0 | 59 | 422 | . | . | . | . |
| PM1548-SK1P3 | FAM222A | p.Ser400Ala | missense_variant | MODERATE | 0.276923 | 0 | 65 | 420 | . | . | . | . |
| PM1548-SK1P3 | TCHP | p.Ala371Thr | missense_variant | MODERATE | 0.333333 | 0 | 3 | 45 | . | . | . | . |
| PM1548-SK1P3 | SH2B3 | p.Glu523fs | frameshift_variant | HIGH | 0 | 0 | 440 | 158 | true | . | . | . |
| PM1548-SK1P3 | HECTD4 | p.Ala3707Val | missense_variant | MODERATE | 0.517241 | 0 | 29 | 63 | . | . | . | . |
| PM1548-SK1P3 | TPCN1 | p.Leu234Arg | missense_variant | MODERATE | 0.304527 | 0 | 243 | 134 | . | . | . | . |
| PM1548-SK1P3 | RBM19 | p.Asn533Asn | synonymous_variant | LOW | 0.491228 | 0 | 171 | 250 | . | . | . | . |
| PM1548-SK1P3 | RAB35 | n.120541738C>T | intragenic_variant | MODIFIER | 0.493056 | 0 | 432 | 243 | . | . | . | . |
| PM1548-SK1P3 | GCN1L1 | p.Ser2359Pro | missense_variant | MODERATE | 0.466667 | 0 | 15 | 85 | . | . | . | . |
| PM1548-SK1P3 | SETD1B | p.His8fs | frameshift_variant | HIGH | 0 | 0 | 18 | 141 | . | . | . | . |
| PM1548-SK1P3 | SETD1B | p.Pro1361fs | frameshift_variant | HIGH | 0.586022 | 0 | 186 | 190 | . | . | . | . |
| PM1548-SK1P3 | ZCCHC8 | p.Gly551Asp | missense_variant | MODERATE | 0.619048 | 0 | 21 | 34 | true | . | . | . |
| PM1548-SK1P3 | ABCB9 | p.His709Gln | missense_variant | MODERATE | 0.304348 | 0 | 115 | 341 | . | . | . | . |
| PM1548-SK1P3 | ABCB9 | p.His709Arg | missense_variant | MODERATE | 0.301724 | 0 | 116 | 330 | . | . | . | . |
| PM1548-SK1P3 | ABCB9 | p.Lys694Arg | missense_variant | MODERATE | 0.319328 | 0 | 119 | 299 | . | . | . | . |
| PM1548-SK1P3 | PITPNM2 | p.Arg749Trp | missense_variant | MODERATE | 0.46875 | 0 | 96 | 160 | . | . | . | . |
| PM1548-SK1P3 | SBNO1 | p.Arg1283Cys | missense_variant&splice_region_variant | MODERATE | 0.307018 | 0 | 114 | 244 | . | . | . | . |
| PM1548-SK1P3 | DNAH10 | p.Pro3133His | missense_variant | MODERATE | 0.382353 | 0.00452489 | 34 | 221 | . | . | . | . |
| PM1548-SK1P3 | TMEM132B | p.Ala297Thr | missense_variant | MODERATE | 0 | 0 | 15 | 79 | . | . | . | . |
| PM1548-SK1P3 | GPR133 | p.Gly731Arg | missense_variant | MODERATE | 0.736842 | 0 | 19 | 110 | . | . | . | . |
| PM1548-SK1P3 | FBRSL1 | p.Glu251Asp | missense_variant | MODERATE | 0 | 0 | 209 | 153 | . | . | . | . |
| PM1548-SK1P3 | MPHOSPH8 | p.Val686Ile | missense_variant | MODERATE | 0 | 0 | 59 | 75 | . | . | . | . |
| PM1548-SK1P3 | GJA3 | p.Ala213Thr | missense_variant | MODERATE | 0.464286 | 0 | 84 | 151 | . | . | . | . |

| | | | | | | | | | | | | |
|---|---|---|---|---|---|---|---|---|---|---|---|---|
| PM1548-SK1P3 | LATS2 | p.Pro41Gln | missense_variant | MODERATE | 0.608696 | 0.025641 | 23 | 39 | . | . | . | . |
| PM1548-SK1P3 | CDK8 | p.Asn372Thr | missense_variant | MODERATE | 0.424242 | 0 | 33 | 248 | . | . | . | . |
| PM1548-SK1P3 | POLR1D | n.28240022G>A | intragenic_variant | MODIFIER | 0.519588 | 0 | 485 | 101 | . | . | . | . |
| PM1548-SK1P3 | FRY | p.Gln2970His | missense_variant | MODERATE | 0.490566 | 0 | 53 | 39 | . | . | . | . |
| PM1548-SK1P3 | BRCA2 | p.Asn863fs | frameshift_variant | HIGH | 0.308642 | 0 | 81 | 84 | true | true | . | true |
| PM1548-SK1P3 | STARD13 | p.Arg155His | missense_variant | MODERATE | 0.266667 | 0 | 30 | 32 | . | . | . | . |
| PM1548-SK1P3 | MAB21L1 | p.Ser103Ser | synonymous_variant | LOW | 0.4 | 0 | 10 | 101 | . | . | . | . |
| PM1548-SK1P3 | FOXO1 | p.Ser152Thr | missense_variant | MODERATE | 0.111111 | 0 | 72 | 139 | true | . | . | . |
| PM1548-SK1P3 | KBTBD7 | p.Tyr195Phe | missense_variant | MODERATE | 0.261538 | 0 | 65 | 644 | . | . | . | . |
| PM1548-SK1P3 | GPALPP1 | p.Pro37Arg | missense_variant | MODERATE | 1 | 0 | 6 | 103 | . | . | . | . |
| PM1548-SK1P3 | FAM124A | p.Ser257Pro | missense_variant | MODERATE | 0 | 0 | 7 | 101 | . | . | . | . |
| PM1548-SK1P3 | NEK3 | p.Asn343Lys | missense_variant | MODERATE | 0 | 0 | 26 | 44 | . | . | . | . |
| PM1548-SK1P3 | DACH1 | p.Arg196Arg | synonymous_variant | LOW | 0.352941 | 0 | 34 | 114 | . | . | . | . |
| PM1548-SK1P3 | ABCC4 | p.Ile1219fs | frameshift_variant | HIGH | 0 | 0 | 277 | 130 | . | . | . | . |
| PM1548-SK1P3 | FGF14 | p.Asp33Gly | missense_variant | MODERATE | 0.607143 | 0 | 56 | 97 | . | . | . | . |
| PM1548-SK1P3 | CCDC168 | p.Met4045fs | frameshift_variant | HIGH | 0.389937 | 0 | 159 | 95 | . | . | . | . |
| PM1548-SK1P3 | ARHGEF7 | p.Thr23fs | frameshift_variant | HIGH | 0.281776 | 0 | 653 | 188 | . | . | . | . |
| PM1548-SK1P3 | MCF2L | p.Arg590Gln | missense_variant | MODERATE | 0.333333 | 0 | 27 | 34 | . | . | . | . |
| PM1548-SK1P3 | CDC16 | p.Asn557fs | frameshift_variant | HIGH | 0 | 0 | 23 | 90 | . | . | . | . |
| PM1548-SK1P3 | UPF3A | p.Asp129Glu | missense_variant | MODERATE | 0.467492 | 0 | 323 | 60 | . | . | . | . |
| PM1548-SK1P3 | OR11G2 | p.Phe141Phe | synonymous_variant | LOW | 0.409836 | 0 | 61 | 116 | . | . | . | . |
| PM1548-SK1P3 | ACIN1 | p.Glu192Asp | missense_variant | MODERATE | 0.111111 | 0 | 63 | 129 | . | . | . | . |
| PM1548-SK1P3 | ACIN1 | p.Glu192Lys | missense_variant | MODERATE | 0.111111 | 0 | 63 | 129 | . | . | . | . |
| PM1548-SK1P3 | PABPN1 | c.*5dupA | frameshift_variant&stop_retained_variant | HIGH | 0.275 | 0 | 40 | 91 | . | . | . | . |
| PM1548-SK1P3 | RNF31 | p.Ala9Gly | missense_variant | MODERATE | 0 | 0 | 58 | 104 | . | . | . | true |
| PM1548-SK1P3 | RNF31 | p.Val12Ala | missense_variant | MODERATE | 0 | 0 | 58 | 104 | . | . | . | true |
| PM1548-SK1P3 | LTB4R2 | p.Ala92Val | missense_variant | MODERATE | 0.188889 | 0 | 90 | 173 | . | . | . | . |
| PM1548-SK1P3 | FOXG1 | p.Glu173Asp | missense_variant | MODERATE | 0.222222 | 0 | 108 | 31 | . | . | . | . |
| PM1548-SK1P3 | CLEC14A | p.Ser404Asn | missense_variant | MODERATE | 0 | 0 | 82 | 138 | . | . | . | . |
| PM1548-SK1P3 | KTN1 | p.Phe1182Val | missense_variant | MODERATE | 0.585366 | 0 | 41 | 114 | true | . | . | . |
| PM1548-SK1P3 | KCNH5 | p.Arg333His | missense_variant | MODERATE | 0.581395 | 0 | 43 | 68 | . | . | . | . |
| PM1548-SK1P3 | ZBTB25 | p.His371Asn | missense_variant | MODERATE | 0 | 0 | 39 | 47 | . | . | . | . |
| PM1548-SK1P3 | HSPA2 | p.Gly138fs | frameshift_variant | HIGH | 0.160256 | 0 | 312 | 148 | . | . | . | . |
| PM1548-SK1P3 | HSPA2 | p.Phe220Phe | synonymous_variant | LOW | 0.25 | 0 | 88 | 96 | . | . | . | . |
| PM1548-SK1P3 | FUT8 | p.Arg249Cys | missense_variant | MODERATE | 0.428571 | 0 | 49 | 316 | . | . | . | . |
| PM1548-SK1P3 | TMEM229B | p.Leu37Phe | missense_variant | MODERATE | 0.479167 | 0 | 192 | 482 | . | . | . | . |
| PM1548-SK1P3 | ZFP36L1 | p.Gly271fs | frameshift_variant | HIGH | 0.484337 | 0 | 415 | 215 | . | . | . | . |
| PM1548-SK1P3 | ELMSAN1 | p.Arg739Cys | missense_variant | MODERATE | 0.494845 | 0 | 194 | 46 | . | . | . | . |
| PM1548-SK1P3 | ELMSAN1 | p.Gln36fs | frameshift_variant | HIGH | 0.598639 | 0 | 147 | 43 | . | . | . | . |
| PM1548-SK1P3 | VRTN | p.Gly328fs | frameshift_variant | HIGH | 0.230769 | 0 | 13 | 58 | . | . | . | . |
| PM1548-SK1P3 | GALC | p.Arg685His | missense_variant | MODERATE | 0 | 0 | 27 | 31 | . | . | . | . |
| PM1548-SK1P3 | PTPN21 | p.Ile849fs | frameshift_variant | HIGH | 0.5 | 0 | 8 | 156 | . | . | . | . |
| PM1548-SK1P3 | ZC3H14 | p.Asn577Asp | missense_variant | MODERATE | 0.328358 | 0 | 67 | 222 | . | . | . | . |
| PM1548-SK1P3 | CCDC88C | p.Lys818Glu | missense_variant | MODERATE | 0 | 0 | 55 | 69 | . | . | . | . |
| PM1548-SK1P3 | FBLN5 | p.Asp357Asn | missense_variant | MODERATE | 0.333333 | 0 | 18 | 147 | . | . | . | . |
| PM1548-SK1P3 | UNC79 | p.Arg746Gln | missense_variant | MODERATE | 0.666667 | 0 | 15 | 31 | . | . | . | . |
| PM1548-SK1P3 | SYNE3 | p.Ile83Val | missense_variant | MODERATE | 0.525 | 0 | 80 | 217 | . | . | . | . |
| PM1548-SK1P3 | SETD3 | p.Phe388Leu | missense_variant | MODERATE | 0.25 | 0 | 4 | 32 | . | . | . | . |
| PM1548-SK1P3 | TECPR2 | p.Gln498fs | frameshift_variant | HIGH | 0.25 | 0 | 4 | 55 | . | . | . | . |
| PM1548-SK1P3 | BRF1 | p.Ala129Ser | missense_variant | MODERATE | 0 | 0 | 29 | 135 | . | . | . | . |
| PM1548-SK1P3 | CYFIP1 | p.Arg1085Leu | missense_variant | MODERATE | 0 | 0 | 16 | 334 | . | . | . | . |
| PM1548-SK1P3 | MAGEL2 | p.Ala11Val | missense_variant | MODERATE | 0.105333 | 0 | 750 | 323 | . | . | . | . |
| PM1548-SK1P3 | PAR5 | n.2021_2022del | intron_variant | MODIFIER | 0.25 | 0 | 100 | 165 | . | . | . | . |
| PM1548-SK1P3 | UBE3A | p.Val133Ala | missense_variant | MODERATE | 0.272727 | 0 | 11 | 79 | . | . | . | . |
| PM1548-SK1P3 | ACTC1 | p.Ala137Thr | missense_variant | MODERATE | 0.4 | 0 | 15 | 73 | . | . | . | . |
| PM1548-SK1P3 | SPRED1 | p.Arg332His | missense_variant | MODERATE | 0.622222 | 0.0070922 | 45 | 141 | . | . | . | . |
| PM1548-SK1P3 | CASC5 | p.Asp178Gly | missense_variant | MODERATE | 0.258065 | 0 | 31 | 58 | true | . | . | . |
| PM1548-SK1P3 | SPTBN5 | p.Arg1943His | missense_variant | MODERATE | 0.0727273 | 0 | 55 | 148 | . | . | . | . |
| PM1548-SK1P3 | SPTBN5 | p.Val1842Gly | missense_variant | MODERATE | 0.149123 | 0 | 114 | 87 | . | . | . | . |
| PM1548-SK1P3 | GANC | p.Ala394Val | missense_variant | MODERATE | 0.282051 | 0 | 39 | 106 | . | . | . | . |
| PM1548-SK1P3 | GANC | p.Gly404Asp | missense_variant | MODERATE | 0.25 | 0 | 52 | 160 | . | . | . | . |
| PM1548-SK1P3 | TTBK2 | p.Ala197Thr | missense_variant | MODERATE | 0.166667 | 0 | 6 | 69 | . | . | . | . |
| PM1548-SK1P3 | CCNDBP1 | p.Asp237Asn | missense_variant | MODERATE | 0.2 | 0 | 10 | 44 | . | . | . | . |
| PM1548-SK1P3 | TGM7 | p.Ser305Phe | missense_variant | MODERATE | 0.230769 | 0 | 26 | 54 | . | . | . | . |
| PM1548-SK1P3 | ZSCAN29 | p.Arg33Trp | missense_variant | MODERATE | 0 | 0 | 58 | 209 | . | . | . | . |
| PM1548-SK1P3 | SLC28A2 | p.Gly128Asp | missense_variant | MODERATE | 0.105263 | 0 | 57 | 169 | . | . | . | . |
| PM1548-SK1P3 | MYO5A | p.Arg831His | missense_variant | MODERATE | 0.506562 | 0 | 381 | 78 | true | . | . | . |
| PM1548-SK1P3 | TCF12 | p.Val183fs | frameshift_variant | HIGH | 0.222222 | 0 | 18 | 200 | true | . | . | . |
| PM1548-SK1P3 | LIPC | p.Phe286fs | frameshift_variant | HIGH | 0.333333 | 0 | 3 | 89 | . | . | . | . |
| PM1548-SK1P3 | OAZ2 | p.Ser92Ser | synonymous_variant | LOW | 0.115942 | 0 | 69 | 163 | . | . | . | . |
| PM1548-SK1P3 | OAZ2 | p.Glu87Gly | missense_variant | MODERATE | 0.123077 | 0 | 65 | 155 | . | . | . | . |
| PM1548-SK1P3 | OAZ2 | p.Glu84Val | missense_variant | MODERATE | 0.121212 | 0 | 66 | 248 | . | . | . | . |
| PM1548-SK1P3 | OAZ2 | p.Val79Gly | missense_variant | MODERATE | 0.118644 | 0 | 59 | 233 | . | . | . | . |
| PM1548-SK1P3 | SPG21 | p.Ala282Val | missense_variant | MODERATE | 0.5 | 0 | 16 | 32 | . | . | . | . |
| PM1548-SK1P3 | CILP | p.Asn930Asn | synonymous_variant | LOW | 0.538462 | 0 | 78 | 111 | . | . | . | . |
| PM1548-SK1P3 | IGDCC3 | p.Pro358Leu | missense_variant | MODERATE | 0 | 0 | 376 | 237 | . | . | . | . |
| PM1548-SK1P3 | NOX5 | p.His470fs | frameshift_variant | HIGH | 0.115385 | 0 | 52 | 164 | . | . | . | . |
| PM1548-SK1P3 | MYO9A | p.Cys1767Ser | missense_variant | MODERATE | 0.38 | 0 | 100 | 186 | . | . | . | . |
| PM1548-SK1P3 | NPTN | p.Arg334Trp | missense_variant | MODERATE | 0.279412 | 0 | 68 | 86 | . | . | . | . |
| PM1548-SK1P3 | ISLR2 | p.Val74Ala | missense_variant | MODERATE | 0.52381 | 0 | 105 | 331 | . | . | . | . |
| PM1548-SK1P3 | C15orf39 | p.Ala475Val | missense_variant | MODERATE | 0.470528 | 0.00393701 | 984 | 254 | . | . | . | . |
| PM1548-SK1P3 | ETFA | p.Thr45Arg | missense_variant | MODERATE | 0.25641 | 0 | 39 | 214 | . | . | . | . |
| PM1548-SK1P3 | ETFA | p.Thr45Ala | missense_variant | MODERATE | 0.25641 | 0 | 39 | 213 | . | . | . | . |
| PM1548-SK1P3 | PSTPIP1 | p.Ala280fs | frameshift_variant&splice_region_variant | HIGH | 0.396714 | 0 | 426 | 438 | . | . | . | . |
| PM1548-SK1P3 | PEAK1 | p.Ala36Thr | missense_variant | MODERATE | 0.272727 | 0 | 44 | 186 | . | . | . | . |
| PM1548-SK1P3 | CHRNB4 | p.Gly261Asp | missense_variant | MODERATE | 0.619048 | 0 | 21 | 56 | . | . | . | . |
| PM1548-SK1P3 | MESDC1 | p.Ser224Asn | missense_variant | MODERATE | 0.653333 | 0 | 75 | 161 | . | . | . | . |
| PM1548-SK1P3 | IL16 | p.Thr417Met | missense_variant | MODERATE | 0 | 0 | 38 | 34 | . | . | . | . |
| PM1548-SK1P3 | BTBD1 | p.Arg236Gln | missense_variant | MODERATE | 0 | 0 | 19 | 77 | . | . | . | . |
| PM1548-SK1P3 | ZNF592 | p.Ala1232Val | missense_variant | MODERATE | 0.4 | 0 | 5 | 35 | . | . | . | . |
| PM1548-SK1P3 | AEN | p.Ala194Val | missense_variant | MODERATE | 0 | 0 | 23 | 122 | . | . | . | . |
| PM1548-SK1P3 | IQGAP1 | p.Met1231fs | frameshift_variant | HIGH | 0.519298 | 0 | 285 | 170 | . | . | . | . |
| PM1548-SK1P3 | FES | c.1531-47G>A | intron_variant | MODIFIER | 0.383253 | 0 | 621 | 35 | . | . | . | . |
| PM1548-SK1P3 | MAN2A2 | p.Pro139Leu | missense_variant | MODERATE | 0.488142 | 0 | 506 | 490 | . | . | . | . |
| PM1548-SK1P3 | RCCD1 | p.Arg87Ser | missense_variant | MODERATE | 0 | 0 | 3 | 63 | . | . | . | . |
| PM1548-SK1P3 | ADAMTS17 | p.Cys921Tyr | missense_variant | MODERATE | 0.404762 | 0 | 42 | 57 | . | . | . | . |
| PM1548-SK1P3 | WDR90 | p.Ala1738Thr | missense_variant | MODERATE | 0.459893 | 0 | 374 | 185 | . | . | . | . |
| PM1548-SK1P3 | RPUSD1 | p.Arg255Gln | missense_variant | MODERATE | 0.536278 | 0 | 317 | 198 | . | . | . | . |
| PM1548-SK1P3 | SOX8 | p.Pro11Thr | missense_variant | MODERATE | 0 | 0 | 77 | 62 | . | . | . | . |
| PM1548-SK1P3 | SOX8 | p.Asp27Asn | missense_variant | MODERATE | 0 | 0 | 123 | 120 | . | . | . | . |
| PM1548-SK1P3 | CACNA1H | p.Arg1892His | missense_variant | MODERATE | 0.0457666 | 0 | 437 | 231 | . | . | . | . |
| PM1548-SK1P3 | BAIAP3 | p.Ala740Asp | missense_variant | MODERATE | 0 | 0 | 27 | 90 | . | . | . | . |
| PM1548-SK1P3 | IFT140 | p.Ala1330Val | missense_variant | MODERATE | 0 | 0 | 320 | 188 | . | . | . | . |
| PM1548-SK1P3 | NME3 | p.Ala20Ser | missense_variant | MODERATE | 0 | 0 | 118 | 162 | . | . | . | . |
| PM1548-SK1P3 | PRSS33 | p.Leu174fs | frameshift_variant | HIGH | 0 | 0 | 243 | 199 | . | . | . | . |
| PM1548-SK1P3 | CCDC64B | p.Arg144Gln | missense_variant | MODERATE | 0.00129199 | 0.00254453 | 774 | 393 | . | . | . | . |
| PM1548-SK1P3 | CREBBP | p.Gly1479Val | missense_variant | MODERATE | 0 | 0 | 57 | 193 | true | . | true | . |
| PM1548-SK1P3 | CORO7 | p.Thr230Ala | missense_variant | MODERATE | 0.164794 | 0 | 267 | 172 | . | . | . | . |
| PM1548-SK1P3 | GLYR1 | p.Ser148Thr | missense_variant | MODERATE | 0.257143 | 0 | 70 | 42 | . | . | . | . |
| PM1548-SK1P3 | TVP23A | p.Phe140fs | frameshift_variant | HIGH | 0.439252 | 0 | 321 | 182 | . | . | . | . |
| PM1548-SK1P3 | CIITA | p.Arg580His | missense_variant | MODERATE | 0.557769 | 0 | 502 | 335 | true | . | . | . |
| PM1548-SK1P3 | RMI2 | p.Ile130Val | missense_variant | MODERATE | 0 | 0 | 39 | 171 | true | . | . | . |
| PM1548-SK1P3 | TNFRSF17 | p.Pro145Gln | missense_variant | MODERATE | 0 | 0 | 29 | 38 | true | . | . | . |
| PM1548-SK1P3 | CDR2 | p.Gln130Arg | missense_variant | MODERATE | 0.44186 | 0 | 43 | 85 | . | . | . | . |
| PM1548-SK1P3 | USP31 | p.Gly1267Arg | missense_variant | MODERATE | 0.2 | 0 | 5 | 32 | . | . | . | . |
| PM1548-SK1P3 | COG7 | p.Pro67Ala | missense_variant | MODERATE | 0.88 | 0 | 25 | 71 | . | . | . | . |
| PM1548-SK1P3 | GTF3C1 | p.Glu1761Gly | missense_variant | MODERATE | 0.121951 | 0 | 82 | 99 | . | . | . | . |
| PM1548-SK1P3 | ATXN2L | c.3140-64G>T | intron_variant | MODIFIER | 0.389831 | 0 | 59 | 124 | . | . | . | . |
| PM1548-SK1P3 | ATP2A1 | p.Gly229Ser | missense_variant | MODERATE | 0.270073 | 0 | 137 | 167 | . | . | . | . |
| PM1548-SK1P3 | KIF22 | p.Ile592fs | frameshift_variant | HIGH | 0.555556 | 0 | 72 | 127 | . | . | . | . |
| PM1548-SK1P3 | PRR14 | p.Arg96Gln | missense_variant | MODERATE | 0.730769 | 0 | 52 | 271 | . | . | . | . |
| PM1548-SK1P3 | ZNF629 | p.Asp434Asn | missense_variant | MODERATE | 0.534314 | 0 | 204 | 34 | . | . | . | . |
| PM1548-SK1P3 | HSD3B7 | p.Val361Ile | missense_variant | MODERATE | 0.49505 | 0 | 101 | 165 | . | . | . | . |
| PM1548-SK1P3 | KAT8 | p.Val438Ala | missense_variant | MODERATE | 0.5 | 0 | 86 | 67 | . | . | . | . |
| PM1548-SK1P3 | FUS | c.1295+1G>A | splice_donor_variant&intron_variant | HIGH | 0.447059 | 0 | 85 | 234 | true | . | . | . |
| PM1548-SK1P3 | PHKB | p.Tyr52His | missense_variant | MODERATE | 0 | 0 | 403 | 155 | . | . | . | . |
| PM1548-SK1P3 | LONP2 | p.Gly681Cys | missense_variant | MODERATE | 0.25 | 0 | 24 | 276 | . | . | . | . |

| | | | | | | | | | | | | |
|---|---|---|---|---|---|---|---|---|---|---|---|---|
| PM1548-SK1P3 | N4BP1 | p.Arg640Cys | missense_variant | MODERATE | 0 | 0 | 3 | 94 | . | . | . | . |
| PM1548-SK1P3 | SALL1 | p.Ser1318Asn | missense_variant | MODERATE | 0.227273 | 0 | 22 | 197 | . | . | . | . |
| PM1548-SK1P3 | SALL1 | p.His756Asp | missense_variant | MODERATE | 0.166667 | 0 | 18 | 53 | . | . | . | . |
| PM1548-SK1P3 | LPCAT2 | c.1216-7923dup | intron_variant | MODIFIER | 0.342857 | 0 | 35 | 56 | . | . | . | . |
| PM1548-SK1P3 | SLC6A2 | p.Gly591Asp | missense_variant | MODERATE | 0.472222 | 0 | 216 | 342 | . | . | . | . |
| PM1548-SK1P3 | MMP15 | p.Ala603Asp | missense_variant | MODERATE | 0 | 0 | 62 | 388 | . | . | . | . |
| PM1548-SK1P3 | CDH11 | p.Ala166Thr | missense_variant | MODERATE | 0.315789 | 0 | 95 | 176 | true | . | . | . |
| PM1548-SK1P3 | CDH11 | p.Thr163Ile | missense_variant | MODERATE | 0.417582 | 0 | 91 | 177 | true | . | . | . |
| PM1548-SK1P3 | CDH11 | p.His161His | synonymous_variant | LOW | 0.425532 | 0 | 94 | 175 | true | . | . | . |
| PM1548-SK1P3 | CDH11 | p.Val149Asp | missense_variant | MODERATE | 0.487923 | 0 | 207 | 62 | true | . | . | . |
| PM1548-SK1P3 | CDH11 | p.Val147Asp | missense_variant | MODERATE | 0.591346 | 0 | 208 | 55 | true | . | . | . |
| PM1548-SK1P3 | CDH11 | p.Glu140Gly | missense_variant | MODERATE | 0.504425 | 0 | 113 | 80 | true | . | . | . |
| PM1548-SK1P3 | CDH11 | p.Arg137Gln | missense_variant | MODERATE | 0.60177 | 0 | 113 | 79 | true | . | . | . |
| PM1548-SK1P3 | TK2 | p.Asn114Asn | synonymous_variant | LOW | 0.65 | 0 | 20 | 102 | . | . | . | . |
| PM1548-SK1P3 | B3GNT9 | p.Asp221Asn | missense_variant | MODERATE | 0.333333 | 0 | 30 | 208 | . | . | . | . |
| PM1548-SK1P3 | LRRC29 | p.Gln205His | missense_variant | MODERATE | 0.04 | 0 | 100 | 214 | . | . | . | . |
| PM1548-SK1P3 | KCTD19 | p.Gly313Arg | missense_variant | MODERATE | 0.475694 | 0 | 288 | 109 | . | . | . | . |
| PM1548-SK1P3 | TPPP3 | p.Ser49Ala | missense_variant | MODERATE | 0.263158 | 0 | 38 | 59 | . | . | . | . |
| PM1548-SK1P3 | CTCF | p.Arg283His | missense_variant | MODERATE | 0 | 0 | 7 | 196 | true | true | true | . |
| PM1548-SK1P3 | CDH3 | p.Asp264Glu | missense_variant | MODERATE | 0.25641 | 0 | 39 | 401 | . | . | . | . |
| PM1548-SK1P3 | WWP2 | p.Arg633Gln | missense_variant | MODERATE | 0.454545 | 0 | 11 | 88 | . | . | . | . |
| PM1548-SK1P3 | PMFBP1 | p.Glu205fs | frameshift_variant | HIGH | 0.109091 | 0 | 55 | 212 | . | . | . | . |
| PM1548-SK1P3 | ZFHX3 | p.Ala1099Ser | missense_variant | MODERATE | 0 | 0 | 54 | 40 | true | . | true | true |
| PM1548-SK1P3 | RFWD3 | p.Val155Ala | missense_variant | MODERATE | 0.312121 | 0 | 330 | 69 | . | . | . | . |
| PM1548-SK1P3 | PKD1L2 | n.81164209C>T | intragenic_variant | MODIFIER | 0.68 | 0 | 50 | 77 | . | . | . | . |
| PM1548-SK1P3 | TAF1C | p.Pro707fs | frameshift_variant | HIGH | 0.585635 | 0 | 181 | 205 | . | . | . | . |
| PM1548-SK1P3 | PIEZO1 | p.Pro1061Leu | missense_variant | MODERATE | 0.5 | 0 | 356 | 171 | . | . | . | . |
| PM1548-SK1P3 | PIEZO1 | p.Met1002Ile | missense_variant | MODERATE | 0.32 | 0 | 25 | 33 | . | . | . | . |
| PM1548-SK1P3 | GALNS | p.Thr109Thr | synonymous_variant | LOW | 0.454545 | 0 | 77 | 187 | . | . | . | . |
| PM1548-SK1P3 | ANKRD11 | p.Asp2130Asn | missense_variant | MODERATE | 0 | 0 | 24 | 42 | . | . | . | . |
| PM1548-SK1P3 | GEMIN4 | p.Arg229Gln | missense_variant | MODERATE | 0.526316 | 0 | 19 | 149 | . | . | . | . |
| PM1548-SK1P3 | RNMTL1 | p.Met143fs | frameshift_variant | HIGH | 0.21519 | 0 | 79 | 36 | . | . | . | . |
| PM1548-SK1P3 | PRPF8 | p.Arg1057Gln | missense_variant | MODERATE | 0.534884 | 0 | 473 | 146 | . | . | . | . |
| PM1548-SK1P3 | HIC1 | p.Pro693Leu | missense_variant | MODERATE | 0 | 0.0126582 | 19 | 79 | . | . | . | . |
| PM1548-SK1P3 | SMG6 | p.Arg603His | missense_variant | MODERATE | 0.081761 | 0.00434783 | 159 | 230 | . | . | . | . |
| PM1548-SK1P3 | CTNS | p.Asp74Asn | missense_variant | MODERATE | 0.492754 | 0.00314465 | 69 | 318 | . | . | . | . |
| PM1548-SK1P3 | ITGAE | p.Arg113Gln | missense_variant | MODERATE | 0.424242 | 0 | 99 | 94 | . | . | . | . |
| PM1548-SK1P3 | ZZEF1 | p.Arg194Trp | missense_variant | MODERATE | 0.333333 | 0 | 12 | 95 | . | . | . | . |
| PM1548-SK1P3 | KIF1C | p.Ser511Phe | missense_variant | MODERATE | 0.457143 | 0 | 35 | 140 | . | . | . | . |
| PM1548-SK1P3 | USP6 | p.Pro181Leu | missense_variant&splice_region_variant | MODERATE | 0.4 | 0 | 10 | 73 | true | . | . | . |
| PM1548-SK1P3 | DVL2 | p.Arg442Cys | missense_variant | MODERATE | 0.449115 | 0 | 452 | 34 | . | . | . | . |
| PM1548-SK1P3 | EIF5A | p.Thr168Met | missense_variant | MODERATE | 0.48 | 0 | 100 | 86 | . | . | true | . |
| PM1548-SK1P3 | DNAH2 | p.Gly4241fs | frameshift_variant | HIGH | 0.443396 | 0 | 106 | 74 | . | . | . | . |
| PM1548-SK1P3 | ALOX12B | p.Ala355Thr | missense_variant | MODERATE | 0.393873 | 0 | 457 | 295 | . | . | . | . |
| PM1548-SK1P3 | GLP2R | p.Ala515Thr | missense_variant | MODERATE | 0 | 0.00983607 | 18 | 305 | . | . | . | . |
| PM1548-SK1P3 | MYH13 | p.Glu984Lys | missense_variant | MODERATE | 0.481481 | 0 | 189 | 72 | . | . | . | . |
| PM1548-SK1P3 | MYH8 | p.Gly186Arg | missense_variant | MODERATE | 0.575758 | 0 | 33 | 81 | . | . | . | . |
| PM1548-SK1P3 | MYH4 | p.Tyr424Cys | missense_variant | MODERATE | 0.462264 | 0 | 106 | 298 | . | . | . | . |
| PM1548-SK1P3 | DNAH9 | p.Arg1542Met | missense_variant | MODERATE | 0 | 0 | 49 | 342 | . | . | . | . |
| PM1548-SK1P3 | DNAH9 | p.Ala2514Thr | missense_variant | MODERATE | 0.295082 | 0 | 61 | 99 | . | . | . | . |
| PM1548-SK1P3 | ARHGAP44 | p.Ala495Thr | missense_variant | MODERATE | 0.714286 | 0 | 7 | 53 | . | . | . | . |
| PM1548-SK1P3 | FLCN | p.His429fs | frameshift_variant | HIGH | 0.0594595 | 0 | 185 | 269 | true | . | . | . |
| PM1548-SK1P3 | FLCN | p.Thr174Ala | missense_variant | MODERATE | 0.131387 | 0 | 137 | 225 | true | . | . | . |
| PM1548-SK1P3 | TOM1L2 | p.Met482fs | frameshift_variant | HIGH | 0.141732 | 0 | 127 | 63 | . | . | . | . |
| PM1548-SK1P3 | KIAA0100 | p.Arg689* | stop_gained | HIGH | 0.431694 | 0 | 183 | 219 | . | . | . | . |
| PM1548-SK1P3 | SUPT6H | p.Ile1356Leu | missense_variant | MODERATE | 0.333333 | 0 | 30 | 211 | . | . | . | . |
| PM1548-SK1P3 | PHF12 | p.Leu393fs | frameshift_variant | HIGH | 0.410646 | 0 | 263 | 60 | . | . | . | . |
| PM1548-SK1P3 | SEZ6 | p.Gly660Asp | missense_variant | MODERATE | 0.533333 | 0 | 15 | 83 | . | . | . | . |
| PM1548-SK1P3 | ANKRD13B | p.Arg233Gln | missense_variant | MODERATE | 0 | 0 | 112 | 98 | . | . | . | . |
| PM1548-SK1P3 | FNDC8 | p.Asn57Ser | missense_variant | MODERATE | 0 | 0 | 2 | 150 | . | . | . | . |
| PM1548-SK1P3 | NLE1 | p.Val10Ala | missense_variant | MODERATE | 0.414894 | 0 | 94 | 97 | . | . | . | . |
| PM1548-SK1P3 | SRCIN1 | p.Leu460Phe | missense_variant | MODERATE | 0.285714 | 0 | 7 | 53 | . | . | . | . |
| PM1548-SK1P3 | PCGF2 | p.Asp157Asn | missense_variant | MODERATE | 0.482759 | 0 | 29 | 53 | . | . | . | . |
| PM1548-SK1P3 | STAC2 | p.Cys141Cys | synonymous_variant | LOW | 0.407407 | 0 | 27 | 250 | . | . | . | . |
| PM1548-SK1P3 | GRB7 | p.Arg552Gln | missense_variant | MODERATE | 0 | 0 | 510 | 159 | . | . | . | . |
| PM1548-SK1P3 | NR1D1 | p.Pro593Pro | synonymous_variant | LOW | 0.191419 | 0.00961538 | 303 | 104 | . | . | . | . |
| PM1548-SK1P3 | MSL1 | p.Leu372Ile | missense_variant | MODERATE | 0.315789 | 0 | 38 | 43 | . | . | . | . |
| PM1548-SK1P3 | WIPF2 | p.Ala39Thr | missense_variant | MODERATE | 0.385714 | 0 | 70 | 66 | . | . | . | . |
| PM1548-SK1P3 | KRT25 | p.Ala91Thr | missense_variant | MODERATE | 0.403846 | 0 | 52 | 148 | . | . | . | . |
| PM1548-SK1P3 | KRT15 | p.Arg398Trp | missense_variant | MODERATE | 0.5 | 0 | 28 | 53 | . | . | . | . |
| PM1548-SK1P3 | KRT16 | p.Gly243Ser | missense_variant | MODERATE | 0.358974 | 0 | 39 | 131 | . | . | . | . |
| PM1548-SK1P3 | DHX58 | p.Arg224Cys | missense_variant | MODERATE | 0.4 | 0 | 90 | 87 | . | . | . | . |
| PM1548-SK1P3 | PLEKHH3 | p.Pro318Thr | missense_variant | MODERATE | 0.1875 | 0 | 16 | 121 | . | . | . | . |
| PM1548-SK1P3 | ETV4 | p.Gln13Arg | missense_variant | MODERATE | 0.1 | 0 | 240 | 215 | true | . | . | . |
| PM1548-SK1P3 | MPP3 | p.Arg265Gln | missense_variant | MODERATE | 0.545455 | 0 | 22 | 79 | . | . | . | . |
| PM1548-SK1P3 | ATXN7L3 | p.Ile246Phe | missense_variant | MODERATE | 0.234375 | 0 | 64 | 37 | . | . | . | . |
| PM1548-SK1P3 | ITGA2B | p.Asn330Ser | missense_variant | MODERATE | 0.166667 | 0 | 6 | 60 | . | . | . | . |
| PM1548-SK1P3 | ARHGAP27 | p.Leu247fs | frameshift_variant | HIGH | 0.552632 | 0 | 152 | 82 | . | . | . | . |
| PM1548-SK1P3 | WNT3 | p.Ala223Ser | missense_variant | MODERATE | 0.27907 | 0 | 43 | 124 | . | . | . | . |
| PM1548-SK1P3 | KPNB1 | p.Ile686Leu | missense_variant | MODERATE | 0 | 0 | 63 | 143 | . | . | . | . |
| PM1548-SK1P3 | KPNB1 | p.Ile686Met | missense_variant | MODERATE | 0 | 0 | 63 | 143 | . | . | . | . |
| PM1548-SK1P3 | SP6 | p.Ala138Thr | missense_variant | MODERATE | 0 | 0 | 273 | 221 | . | . | . | . |
| PM1548-SK1P3 | PPP1R9B | p.Glu24Glu | synonymous_variant | LOW | 0.25641 | 0 | 39 | 79 | . | . | . | . |
| PM1548-SK1P3 | CUEDC1 | p.Arg71Cys | missense_variant | MODERATE | 0.402778 | 0 | 72 | 107 | . | . | . | . |
| PM1548-SK1P3 | LPO | p.Arg82Cys | missense_variant | MODERATE | 0.344828 | 0 | 58 | 404 | . | . | . | . |
| PM1548-SK1P3 | RNF43 | p.Gly659fs | frameshift_variant | HIGH | 0.497854 | 0 | 233 | 76 | true | . | . | true |
| PM1548-SK1P3 | TBX2 | p.Leu460fs | frameshift_variant | HIGH | 0.481481 | 0 | 54 | 331 | . | . | . | . |
| PM1548-SK1P3 | MRC2 | p.Arg490Cys | missense_variant | MODERATE | 0 | 0 | 16 | 51 | . | . | . | . |
| PM1548-SK1P3 | CCDC47 | p.Ser303Pro | missense_variant | MODERATE | 0 | 0 | 11 | 131 | . | . | . | . |
| PM1548-SK1P3 | FTSJ3 | p.Met493Ile | missense_variant | MODERATE | 0.333333 | 0 | 15 | 61 | . | . | . | . |
| PM1548-SK1P3 | ABCA10 | p.Tyr1399His | missense_variant | MODERATE | 0.393939 | 0 | 33 | 335 | . | . | . | . |
| PM1548-SK1P3 | SLC16A5 | p.Gly93Cys | missense_variant | MODERATE | 0 | 0 | 5 | 58 | . | . | . | . |
| PM1548-SK1P3 | ITGB4 | p.Pro305Leu | missense_variant | MODERATE | 0.636364 | 0 | 11 | 78 | . | . | . | . |
| PM1548-SK1P3 | ITGB4 | p.Pro723Ser | missense_variant | MODERATE | 0.104225 | 0 | 355 | 167 | . | . | . | . |
| PM1548-SK1P3 | H3F3B | p.Ala8Asp | missense_variant | MODERATE | 0 | 0 | 22 | 88 | true | . | . | . |
| PM1548-SK1P3 | EVPL | p.Arg266Trp | missense_variant | MODERATE | 0.5 | 0 | 26 | 157 | . | . | . | . |
| PM1548-SK1P3 | RHBDF2 | p.Ala354Thr | missense_variant | MODERATE | 0 | 0 | 35 | 138 | . | . | . | . |
| PM1548-SK1P3 | TNRC6C | p.Arg1160His | missense_variant | MODERATE | 0.339286 | 0 | 56 | 184 | . | . | . | . |
| PM1548-SK1P3 | TMC6 | p.His31His | synonymous_variant | LOW | 0.546296 | 0 | 216 | 491 | . | . | . | . |
| PM1548-SK1P3 | CCDC40 | p.Gly929Ser | missense_variant | MODERATE | 0 | 0 | 2 | 34 | . | . | . | . |
| PM1548-SK1P3 | GAA | p.Gly828Asp | missense_variant&splice_region_variant | MODERATE | 0.0273973 | 0 | 146 | 76 | . | . | . | . |
| PM1548-SK1P3 | ENDOV | p.Pro259Leu | missense_variant | MODERATE | 0.508333 | 0 | 120 | 82 | . | . | . | . |
| PM1548-SK1P3 | RPTOR | p.Arg899* | stop_gained | HIGH | 0.230769 | 0 | 13 | 72 | . | . | . | . |
| PM1548-SK1P3 | SIRT7 | p.Thr181Met | missense_variant | MODERATE | 0.0731707 | 0 | 41 | 217 | . | . | . | . |
| PM1548-SK1P3 | OGFOD3 | p.Ser127Ser | splice_region_variant&synonymous_variant | LOW | 0.571429 | 0 | 28 | 110 | . | . | . | . |
| PM1548-SK1P3 | TXNDC2 | p.Ile284Leu | missense_variant | MODERATE | 0 | 0 | 53 | 275 | . | . | . | . |
| PM1548-SK1P3 | TXNDC2 | p.Ser285Pro | missense_variant | MODERATE | 0 | 0 | 52 | 275 | . | . | . | . |
| PM1548-SK1P3 | TXNDC2 | p.Pro288Leu | missense_variant | MODERATE | 0 | 0.00816327 | 47 | 245 | . | . | . | . |
| PM1548-SK1P3 | ANKRD62 | p.Gln745His | missense_variant | MODERATE | 1 | 0 | 15 | 165 | . | . | . | . |
| PM1548-SK1P3 | ZNF521 | p.Cys754Tyr | missense_variant | MODERATE | 0.241379 | 0 | 58 | 57 | true | . | . | . |
| PM1548-SK1P3 | KCTD1 | p.Asn124His | missense_variant | MODERATE | 0.451613 | 0 | 31 | 69 | . | . | . | . |
| PM1548-SK1P3 | DSG4 | p.Thr1001Met | missense_variant | MODERATE | 0.981818 | 0 | 110 | 104 | . | . | . | . |
| PM1548-SK1P3 | CCDC178 | p.Arg827Met | missense_variant | MODERATE | 0 | 0 | 74 | 192 | . | . | . | . |
| PM1548-SK1P3 | INO80C | p.Gly216Ser | missense_variant | MODERATE | 0 | 0 | 142 | 89 | . | . | . | . |
| PM1548-SK1P3 | SETBP1 | p.Glu244Asp | missense_variant | MODERATE | 0.137255 | 0 | 51 | 133 | true | true | . | . |
| PM1548-SK1P3 | SETBP1 | p.Ile251Val | missense_variant | MODERATE | 0.0864198 | 0 | 81 | 201 | true | true | . | . |
| PM1548-SK1P3 | SETBP1 | p.Ala733Thr | missense_variant | MODERATE | 0.135135 | 0 | 37 | 313 | true | true | . | . |
| PM1548-SK1P3 | SIGLEC15 | p.Ala96Thr | missense_variant | MODERATE | 0.363636 | 0 | 11 | 175 | . | . | . | . |
| PM1548-SK1P3 | SIGLEC15 | p.Arg99Pro | missense_variant | MODERATE | 0.363636 | 0 | 11 | 176 | . | . | . | . |
| PM1548-SK1P3 | SKOR2 | p.Asp736Glu | missense_variant | MODERATE | 0.300971 | 0 | 206 | 157 | . | . | . | . |
| PM1548-SK1P3 | SKOR2 | p.Gly720Glu | missense_variant | MODERATE | 0.282609 | 0 | 184 | 134 | . | . | . | . |
| PM1548-SK1P3 | SMAD7 | p.Pro55Ala | missense_variant | MODERATE | 0.444444 | 0 | 9 | 32 | . | . | . | . |
| PM1548-SK1P3 | DSEL | p.Arg1009His | missense_variant | MODERATE | 1 | 0 | 13 | 67 | . | . | . | . |
| PM1548-SK1P3 | FAM69C | p.Gly48Cys | missense_variant | MODERATE | 0 | 0 | 13 | 170 | . | . | . | . |
| PM1548-SK1P3 | TSHZ1 | p.Ala570Val | missense_variant | MODERATE | 0 | 0 | 70 | 279 | . | . | . | . |
| PM1548-SK1P3 | TSHZ1 | p.Tyr775Cys | missense_variant | MODERATE | 0.944444 | 0.00806452 | 18 | 124 | . | . | . | . |
| PM1548-SK1P3 | ZNF516 | p.Gly232del | disruptive_inframe_deletion | MODERATE | 0.483871 | 0 | 31 | 279 | . | . | . | . |

| | | | | | | | | | | | | |
|---|---|---|---|---|---|---|---|---|---|---|---|---|
| PM1548-SK1P3 | ZNF516 | p.Pro229_Gly23( | inframe_insertion | MODERATE | 0.466667 | 0 | 30 | 280 | . | . | . | . |
| PM1548-SK1P3 | GALR1 | p.Leu46Met | missense_variant | MODERATE | 0.303318 | 0 | 211 | 102 | . | . | . | . |
| PM1548-SK1P3 | GALR1 | p.Lys62Lys | synonymous_variant | LOW | 0.326316 | 0 | 380 | 212 | . | . | . | . |
| PM1548-SK1P3 | SALL3 | p.Pro1105fs | frameshift_variant | HIGH | 1 | 0 | 18 | 200 | . | . | . | . |
| PM1548-SK1P3 | PPAP2C | n.291320T>C | intragenic_variant | MODIFIER | 0.0555556 | 0 | 18 | 60 | . | . | . | . |
| PM1548-SK1P3 | LPPR3 | p.Gly646Ser | missense_variant | MODERATE | 0.333333 | 0 | 9 | 120 | . | . | . | . |
| PM1548-SK1P3 | CFD | p.Arg199Cys | missense_variant | MODERATE | 0.496774 | 0 | 155 | 193 | . | . | . | . |
| PM1548-SK1P3 | ARID3A | p.Arg503Cys | missense_variant | MODERATE | 1 | 0 | 4 | 124 | . | . | . | . |
| PM1548-SK1P3 | APC2 | p.Asp501Asn | missense_variant | MODERATE | 0.5 | 0 | 796 | 282 | . | . | . | . |
| PM1548-SK1P3 | PCSK4 | p.Gly57Asp | missense_variant | MODERATE | 0.366142 | 0 | 1016 | 340 | . | . | . | . |
| PM1548-SK1P3 | ADAMTSL5 | p.Ala343fs | frameshift_variant | HIGH | 0.231959 | 0 | 970 | 366 | . | . | . | . |
| PM1548-SK1P3 | ADAMTSL5 | p.Pro341Ala | missense_variant | MODERATE | 0.375723 | 0 | 519 | 312 | . | . | . | . |
| PM1548-SK1P3 | MEX3D | p.Gly262Asp | missense_variant | MODERATE | 0.444444 | 0 | 9 | 217 | . | . | . | . |
| PM1548-SK1P3 | TCF3 | p.Gly247fs | frameshift_variant | HIGH | 0 | 0 | 66 | 71 | true | . | . | . |
| PM1548-SK1P3 | C19orf35 | p.Arg325Cys | missense_variant | MODERATE | 0 | 0 | 92 | 53 | . | . | . | . |
| PM1548-SK1P3 | NCLN | p.Thr70Met | missense_variant | MODERATE | 0.559322 | 0 | 295 | 140 | . | . | . | . |
| PM1548-SK1P3 | DOHH | p.Arg286Trp | missense_variant | MODERATE | 0.45 | 0.0136054 | 100 | 147 | . | . | . | . |
| PM1548-SK1P3 | EEF2 | p.Pro738Ser | missense_variant | MODERATE | 0.408654 | 0 | 208 | 230 | . | . | . | . |
| PM1548-SK1P3 | PIAS4 | p.Tyr265Tyr | synonymous_variant | LOW | 0.433387 | 0 | 623 | 240 | . | . | . | . |
| PM1548-SK1P3 | MAP2K2 | p.Lys51Arg | missense_variant | MODERATE | 0.315789 | 0 | 38 | 75 | true | . | . | . |
| PM1548-SK1P3 | MAP2K2 | p.Lys51Gln | missense_variant | MODERATE | 0.307692 | 0 | 39 | 75 | true | . | . | . |
| PM1548-SK1P3 | CREB3L3 | p.Val140Ala | missense_variant | MODERATE | 0 | 0 | 385 | 205 | . | . | . | . |
| PM1548-SK1P3 | UBXN6 | p.Thr337Met | missense_variant | MODERATE | 0 | 0 | 97 | 93 | . | . | . | . |
| PM1548-SK1P3 | PLIN4 | p.Val592Met | missense_variant | MODERATE | 0.112782 | 0 | 133 | 40 | . | . | . | . |
| PM1548-SK1P3 | PLIN5 | p.Thr456Ile | missense_variant | MODERATE | 0.540541 | 0 | 37 | 37 | . | . | . | . |
| PM1548-SK1P3 | AC005594.3 | n.381G>A | non_coding_exon_variant | MODIFIER | 0.520833 | 0.0037037 | 240 | 270 | . | . | . | . |
| PM1548-SK1P3 | DPP9 | p.Arg125Trp | missense_variant | MODERATE | 0.0591716 | 0 | 338 | 40 | . | . | . | . |
| PM1548-SK1P3 | KDM4B | p.Arg618Cys | missense_variant | MODERATE | 0 | 0 | 115 | 167 | . | . | . | . |
| PM1548-SK1P3 | KDM4B | p.Glu1013Lys | missense_variant | MODERATE | 0.37148 | 0 | 1101 | 111 | . | . | . | . |
| PM1548-SK1P3 | DENND1C | p.Thr165fs | frameshift_variant | HIGH | 0.333333 | 0 | 30 | 34 | . | . | . | . |
| PM1548-SK1P3 | PEX11G | p.Arg124Trp | missense_variant | MODERATE | 0.46383 | 0 | 235 | 176 | . | . | . | . |
| PM1548-SK1P3 | CTD-3193O13.9 | p.Thr838Thr | synonymous_variant | LOW | 0.297297 | 0 | 37 | 174 | . | . | . | . |
| PM1548-SK1P3 | COL5A3 | p.Glu1677Lys | missense_variant | MODERATE | 0.485714 | 0 | 35 | 264 | . | . | true | true |
| PM1548-SK1P3 | ZNF653 | p.Asp247Gly | missense_variant | MODERATE | 0.28 | 0 | 25 | 166 | . | . | . | . |
| PM1548-SK1P3 | ELOF1 | p.Arg41Cys | missense_variant | MODERATE | 0.471311 | 0 | 244 | 108 | . | . | . | . |
| PM1548-SK1P3 | MRI1 | p.His330fs | frameshift_variant | HIGH | 0.475 | 0 | 40 | 82 | . | . | . | . |
| PM1548-SK1P3 | RFX1 | p.Val600Ile | missense_variant | MODERATE | 0.11435 | 0 | 446 | 347 | . | . | . | . |
| PM1548-SK1P3 | PRKACA | p.Ala219Val | missense_variant | MODERATE | 0.516129 | 0 | 31 | 200 | true | . | . | . |
| PM1548-SK1P3 | TECR | p.Val85Met | missense_variant | MODERATE | 0.558824 | 0.00641026 | 68 | 156 | . | . | . | . |
| PM1548-SK1P3 | EMR2 | p.Pro294Thr | missense_variant | MODERATE | 0.119403 | 0 | 134 | 117 | . | . | . | . |
| PM1548-SK1P3 | EMR2 | p.Tyr292Ser | missense_variant | MODERATE | 0.135593 | 0 | 118 | 117 | . | . | . | . |
| PM1548-SK1P3 | KLF2 | p.Ala238fs | frameshift_variant | HIGH | 0.545455 | 0 | 99 | 137 | . | . | . | . |
| PM1548-SK1P3 | NWD1 | p.Arg788Cys | missense_variant | MODERATE | 0.669903 | 0 | 103 | 334 | . | . | . | . |
| PM1548-SK1P3 | BST2 | p.Ile120fs | frameshift_variant | HIGH | 0.833333 | 0 | 12 | 64 | . | . | . | . |
| PM1548-SK1P3 | FCHO1 | p.Ala286Val | missense_variant&splice_region_variant | MODERATE | 0.39604 | 0 | 101 | 272 | . | . | . | . |
| PM1548-SK1P3 | DDX49 | p.Arg310Gln | missense_variant&splice_region_variant | MODERATE | 0.450882 | 0 | 794 | 300 | . | . | . | . |
| PM1548-SK1P3 | SUGP2 | p.Cys490Tyr | missense_variant | MODERATE | 0.047619 | 0 | 21 | 81 | . | . | . | . |
| PM1548-SK1P3 | NR2C2AP | p.Lys28fs | frameshift_variant | HIGH | 0.485294 | 0 | 340 | 183 | . | . | . | . |
| PM1548-SK1P3 | NCAN | p.Arg256Gln | missense_variant | MODERATE | 0.515152 | 0 | 132 | 124 | . | . | . | . |
| PM1548-SK1P3 | NDUFA13 | p.Trp131Leu | missense_variant | MODERATE | 0 | 0 | 29 | 265 | . | . | . | . |
| PM1548-SK1P3 | ZNF728 | p.Pro380Ser | missense_variant | MODERATE | 0.208333 | 0 | 24 | 32 | . | . | . | . |
| PM1548-SK1P3 | ZNF728 | p.Arg323Pro | missense_variant | MODERATE | 0.217391 | 0 | 23 | 48 | . | . | . | . |
| PM1548-SK1P3 | ZNF728 | p.Arg323Trp | missense_variant | MODERATE | 0.217391 | 0 | 23 | 48 | . | . | . | . |
| PM1548-SK1P3 | ZNF507 | p.Asp475Gly | missense_variant | MODERATE | 0.714286 | 0 | 7 | 42 | . | . | . | . |
| PM1548-SK1P3 | KIAA0355 | p.Val801Met | missense_variant | MODERATE | 0.307054 | 0 | 241 | 184 | . | . | . | . |
| PM1548-SK1P3 | GRAMD1A | p.Gly605Ser | missense_variant | MODERATE | 0.0743802 | 0 | 121 | 68 | . | . | . | . |
| PM1548-SK1P3 | RBM42 | p.Pro117Arg | missense_variant | MODERATE | 0.455357 | 0 | 112 | 129 | . | . | . | . |
| PM1548-SK1P3 | ARHGAP33 | p.Gln144fs | frameshift_variant | HIGH | 0.524668 | 0 | 2108 | 430 | . | . | . | . |
| PM1548-SK1P3 | ZNF850 | p.Ala575Thr | missense_variant | MODERATE | 0.0138889 | 0 | 72 | 89 | . | . | . | . |
| PM1548-SK1P3 | ZNF850 | p.Ser572Ala | missense_variant | MODERATE | 0.0113636 | 0 | 88 | 133 | . | . | . | . |
| PM1548-SK1P3 | ZNF850 | p.Val554Ile | missense_variant | MODERATE | 0.258065 | 0 | 62 | 85 | . | . | . | . |
| PM1548-SK1P3 | DPF1 | p.Cys58Cys | synonymous_variant | LOW | 0.255319 | 0 | 47 | 394 | . | . | . | . |
| PM1548-SK1P3 | DPF1 | p.Cys50Cys | synonymous_variant | LOW | 0.25 | 0 | 48 | 391 | . | . | . | . |
| PM1548-SK1P3 | YIF1B | p.Phe39Phe | synonymous_variant | LOW | 0.330935 | 0 | 139 | 480 | . | . | . | . |
| PM1548-SK1P3 | RYR1 | p.Val3218Met | missense_variant | MODERATE | 0 | 0 | 149 | 114 | . | . | . | . |
| PM1548-SK1P3 | FCGBP | p.Ala5241Val | missense_variant | MODERATE | 0.326797 | 0 | 153 | 284 | . | . | . | . |
| PM1548-SK1P3 | PRX | p.Glu495Gln | missense_variant | MODERATE | 0.438596 | 0 | 57 | 97 | . | true | . | . |
| PM1548-SK1P3 | B3GNT8 | p.Glu333Ala | missense_variant | MODERATE | 0.533333 | 0 | 540 | 219 | . | . | . | . |
| PM1548-SK1P3 | ARHGEF1 | p.Arg137Trp | missense_variant | MODERATE | 0.521127 | 0 | 142 | 130 | . | . | . | . |
| PM1548-SK1P3 | ZNF227 | p.Gly390Ser | missense_variant | MODERATE | 0.333333 | 0 | 15 | 73 | . | . | . | . |
| PM1548-SK1P3 | PVRL2 | p.Pro53Leu | missense_variant | MODERATE | 0.432432 | 0 | 74 | 119 | . | . | . | . |
| PM1548-SK1P3 | MARK4 | p.Thr651Thr | synonymous_variant | LOW | 0.0990991 | 0 | 111 | 352 | . | . | . | . |
| PM1548-SK1P3 | KLC3 | p.Arg320Cys | missense_variant | MODERATE | 0.686869 | 0 | 99 | 102 | . | . | . | . |
| PM1548-SK1P3 | PPP1R13L | p.Leu397Pro | missense_variant | MODERATE | 0.03125 | 0 | 96 | 478 | . | . | . | . |
| PM1548-SK1P3 | GIPR | p.Gly144Ser | missense_variant | MODERATE | 1 | 0 | 2 | 35 | . | . | . | . |
| PM1548-SK1P3 | STRN4 | p.Ala484Val | missense_variant | MODERATE | 0.54386 | 0 | 228 | 141 | . | . | . | . |
| PM1548-SK1P3 | FKRP | p.Ala193Thr | missense_variant | MODERATE | 0.00206186 | 0 | 485 | 98 | . | . | . | . |
| PM1548-SK1P3 | SLC1A5 | p.Ala534Thr | missense_variant | MODERATE | 0.125 | 0 | 8 | 44 | . | . | . | . |
| PM1548-SK1P3 | ZC3H4 | p.Arg894His | missense_variant | MODERATE | 0 | 0 | 459 | 254 | . | . | . | . |
| PM1548-SK1P3 | ZC3H4 | p.Gly668Ser | missense_variant | MODERATE | 0.347826 | 0 | 23 | 225 | . | . | . | . |
| PM1548-SK1P3 | ZC3H4 | p.Lys500Arg | missense_variant | MODERATE | 0 | 0 | 58 | 129 | . | . | . | . |
| PM1548-SK1P3 | ZNF541 | p.Arg214His | missense_variant | MODERATE | 0 | 0 | 19 | 53 | . | . | . | . |
| PM1548-SK1P3 | GLTSCR1 | p.Ser1126fs | frameshift_variant | HIGH | 0.499051 | 0 | 527 | 148 | . | . | . | . |
| PM1548-SK1P3 | CRX | p.Ala35Thr | missense_variant&splice_region_variant | MODERATE | 0.44186 | 0 | 430 | 73 | . | . | . | . |
| PM1548-SK1P3 | PLEKHA4 | p.Arg225Trp | missense_variant | MODERATE | 0.433333 | 0 | 30 | 36 | . | . | . | . |
| PM1548-SK1P3 | BAX | p.Glu41fs | frameshift_variant | HIGH | 0.0104167 | 0 | 192 | 76 | . | . | . | . |
| PM1548-SK1P3 | FLT3LG | p.Ser118fs | frameshift_variant | HIGH | 0.418182 | 0 | 55 | 82 | . | . | . | . |
| PM1548-SK1P3 | SCAF1 | p.Gly651fs | frameshift_variant | HIGH | 0.473684 | 0 | 76 | 93 | . | . | . | . |
| PM1548-SK1P3 | ASPDH | p.Glu177Lys | missense_variant | MODERATE | 0.666667 | 0 | 36 | 225 | . | . | . | . |
| PM1548-SK1P3 | LRRC4B | p.Gly50fs | frameshift_variant | HIGH | 0.574074 | 0 | 54 | 58 | . | . | . | . |
| PM1548-SK1P3 | ZNF611 | p.Met226Thr | missense_variant | MODERATE | 0 | 0 | 4 | 80 | . | . | . | . |
| PM1548-SK1P3 | CNOT3 | p.Tyr637His | missense_variant | MODERATE | 0.5 | 0 | 18 | 41 | true | . | . | true |
| PM1548-SK1P3 | LILRB3 | c.355+1720T>A | intron_variant | MODIFIER | 0.220339 | 0 | 59 | 187 | . | . | . | . |
| PM1548-SK1P3 | LILRB3 | c.355+1720T>A | intron_variant | MODIFIER | 0.220339 | 0 | 59 | 187 | . | . | . | . |
| PM1548-SK1P3 | LILRB3 | c.355+1720T>A | intron_variant | MODIFIER | 0.216667 | 0 | 60 | 184 | . | . | . | . |
| PM1548-SK1P3 | LILRB3 | c.355+1720T>A | intron_variant | MODIFIER | 0.216667 | 0 | 60 | 184 | . | . | . | . |
| PM1548-SK1P3 | EPS8L1 | p.Arg608Cys | missense_variant | MODERATE | 0.388889 | 0 | 36 | 167 | . | . | . | . |
| PM1548-SK1P3 | SBK2 | p.Ala115Thr | missense_variant | MODERATE | 0.425743 | 0 | 202 | 186 | . | . | . | . |
| PM1548-SK1P3 | ZNF579 | p.Arg101Cys | missense_variant | MODERATE | 0.602273 | 0 | 176 | 216 | . | . | . | . |
| PM1548-SK1P3 | ZNF865 | p.Leu610Phe | missense_variant | MODERATE | 0.27907 | 0 | 86 | 220 | . | . | . | . |
| PM1548-SK1P3 | ZNF865 | p.Leu632Leu | synonymous_variant | LOW | 0.346154 | 0 | 78 | 209 | . | . | . | . |
| PM1548-SK1P3 | ZNF71 | p.Gln343Gln | synonymous_variant | LOW | 0.448567 | 0 | 1186 | 303 | . | . | . | . |
| PM1548-SK1P3 | TRIM28 | p.Arg492Cys | missense_variant | MODERATE | 0.446809 | 0 | 47 | 142 | . | . | . | . |
| PM1548-SK1P3 | PXDN | p.Val850Met | missense_variant | MODERATE | 0 | 0 | 875 | 308 | . | . | . | . |
| PM1548-SK1P3 | ROCK2 | p.Glu1060Gly | missense_variant | MODERATE | 0 | 0 | 24 | 93 | . | . | . | . |
| PM1548-SK1P3 | FAM228B | p.Asp82Gly | missense_variant | MODERATE | 0.464286 | 0 | 28 | 174 | . | . | . | . |
| PM1548-SK1P3 | NCOA1 | p.Asn1244Thr | missense_variant | MODERATE | 0.172043 | 0 | 93 | 206 | true | . | . | . |
| PM1548-SK1P3 | NCOA1 | p.Ile1260Val | missense_variant | MODERATE | 0.24031 | 0 | 129 | 203 | true | . | . | . |
| PM1548-SK1P3 | NCOA1 | p.Ala1274Thr | missense_variant | MODERATE | 0.266667 | 0 | 60 | 84 | true | . | . | . |
| PM1548-SK1P3 | ADCY3 | p.Thr182Met | missense_variant | MODERATE | 0.581281 | 0 | 203 | 84 | . | . | . | . |
| PM1548-SK1P3 | TRIM54 | p.Arg97Gln | missense_variant | MODERATE | 0.542857 | 0 | 35 | 313 | . | . | . | . |
| PM1548-SK1P3 | TRIM54 | p.Glu131Asp | missense_variant | MODERATE | 0.25 | 0 | 44 | 141 | . | . | . | . |
| PM1548-SK1P3 | HEATR5B | p.Lys48fs | frameshift_variant | HIGH | 0 | 0 | 199 | 85 | . | . | . | . |
| PM1548-SK1P3 | RMDN2 | p.Arg130His | missense_variant | MODERATE | 0.252033 | 0 | 123 | 120 | . | . | . | . |
| PM1548-SK1P3 | SOS1 | p.Asn15Lys | missense_variant | MODERATE | 0.471642 | 0 | 335 | 93 | . | . | . | . |
| PM1548-SK1P3 | FSHR | p.Arg569Lys | missense_variant | MODERATE | 0.266667 | 0 | 165 | 177 | . | . | . | . |
| PM1548-SK1P3 | FSHR | p.Ser566Arg | missense_variant | MODERATE | 0.266667 | 0 | 165 | 133 | . | . | . | . |
| PM1548-SK1P3 | FSHR | p.Asn558Asn | synonymous_variant | LOW | 0.268293 | 0 | 164 | 139 | . | . | . | . |
| PM1548-SK1P3 | FSHR | p.Ile550Thr | missense_variant | MODERATE | 0.255319 | 0 | 141 | 159 | . | . | . | . |
| PM1548-SK1P3 | CCDC88A | p.Pro1470Ser | missense_variant | MODERATE | 0.41844 | 0 | 141 | 211 | . | . | . | . |
| PM1548-SK1P3 | CCDC85A | p.His295Tyr | missense_variant | MODERATE | 0.363636 | 0 | 11 | 103 | . | . | . | . |
| PM1548-SK1P3 | CCDC85A | p.Pro359Leu | missense_variant | MODERATE | 1 | 0 | 1 | 45 | . | . | . | . |
| PM1548-SK1P3 | FANCL | c.555+2T>C | splice_donor_variant&intron_variant | HIGH | 0.433962 | 0 | 53 | 47 | . | . | . | . |
| PM1548-SK1P3 | OTX1 | p.Arg39Trp | missense_variant | MODERATE | 0.333333 | 0 | 18 | 146 | . | . | . | . |
| PM1548-SK1P3 | AFTPH | p.Asp198fs | frameshift_variant | HIGH | 0 | 0 | 49 | 159 | . | . | . | . |
| PM1548-SK1P3 | NAGK | p.Lys90Gln | missense_variant | MODERATE | 0.277778 | 0 | 108 | 340 | . | . | . | . |
| PM1548-SK1P3 | NAGK | p.Asn100Asp | missense_variant | MODERATE | 0.272727 | 0 | 110 | 271 | . | . | . | . |

| | | | | | | | | | | | | |
|---|---|---|---|---|---|---|---|---|---|---|---|---|
| PM1548-SK1P3 | FBXO41 | p.Arg594Gln | missense_variant | MODERATE | 0 | 0 | 23 | 69 | . | . | . | . |
| PM1548-SK1P3 | TET3 | p.Pro1195Leu | missense_variant | MODERATE | 0.357143 | 0 | 14 | 50 | . | . | . | . |
| PM1548-SK1P3 | C2orf81 | p.Ala344Val | missense_variant | MODERATE | 0.481481 | 0 | 189 | 234 | . | . | . | . |
| PM1548-SK1P3 | TCF7L1 | p.His246Pro | missense_variant | MODERATE | 0.138298 | 0.00719424 | 94 | 139 | . | . | . | . |
| PM1548-SK1P3 | KDM3A | p.Glu1141* | stop_gained | HIGH | 0 | 0 | 83 | 166 | . | . | . | . |
| PM1548-SK1P3 | ZNF514 | p.Gly396Glu | missense_variant | MODERATE | 0.304878 | 0 | 82 | 187 | . | . | . | . |
| PM1548-SK1P3 | NCAPH | p.Gln453* | stop_gained&splice_region_variant | HIGH | 0.111111 | 0 | 9 | 53 | . | . | . | . |
| PM1548-SK1P3 | ZAP70 | p.Ala387Val | missense_variant | MODERATE | 0.315789 | 0 | 57 | 293 | . | . | . | . |
| PM1548-SK1P3 | LONRF2 | p.Asp406Gly | missense_variant | MODERATE | 0 | 0 | 23 | 174 | . | . | . | . |
| PM1548-SK1P3 | CHST10 | p.Asn138Asn | synonymous_variant | LOW | 0.518664 | 0 | 509 | 236 | . | . | . | . |
| PM1548-SK1P3 | IL18R1 | p.Pro527fs | frameshift_variant | HIGH | 0.386503 | 0 | 163 | 255 | . | . | . | . |
| PM1548-SK1P3 | FBLN7 | c.670+86C>T | intron_variant | MODIFIER | 0.580838 | 0 | 167 | 109 | . | . | . | . |
| PM1548-SK1P3 | FBLN7 | p.Arg381Cys | missense_variant | MODERATE | 0.307692 | 0 | 26 | 32 | . | . | . | . |
| PM1548-SK1P3 | ZC3H6 | p.Arg99fs | frameshift_variant | HIGH | 0.325 | 0 | 40 | 90 | . | . | . | . |
| PM1548-SK1P3 | POLR1B | p.Cys893Tyr | missense_variant | MODERATE | 0.310345 | 0 | 58 | 125 | . | . | . | . |
| PM1548-SK1P3 | IL36B | c.261+1014delT | intron_variant | MODIFIER | 0 | 0 | 4 | 101 | . | . | . | . |
| PM1548-SK1P3 | TMEM185B | p.Pro341fs | frameshift_variant | HIGH | 0.0666667 | 0 | 105 | 232 | . | . | . | . |
| PM1548-SK1P3 | CLASP1 | p.Ala1225Val | missense_variant | MODERATE | 0.189189 | 0 | 37 | 85 | . | . | . | . |
| PM1548-SK1P3 | CYP27C1 | p.Ala103Thr | missense_variant | MODERATE | 0.222222 | 0 | 9 | 179 | . | . | . | . |
| PM1548-SK1P3 | ACVR2A | p.Lys437fs | frameshift_variant | HIGH | 0.988889 | 0 | 90 | 199 | . | true | . | . |
| PM1548-SK1P3 | NEB | p.Phe2843Tyr | missense_variant | MODERATE | 0.60396 | 0 | 202 | 178 | . | . | . | . |
| PM1548-SK1P3 | TBR1 | p.Gly599Ser | missense_variant | MODERATE | 1 | 0 | 35 | 40 | . | . | . | . |
| PM1548-SK1P3 | SCN3A | p.Ala1768Val | missense_variant | MODERATE | 0.75 | 0 | 4 | 51 | . | . | . | . |
| PM1548-SK1P3 | TLK1 | p.Gln18Arg | missense_variant | MODERATE | 0.3 | 0 | 80 | 550 | . | . | . | . |
| PM1548-SK1P3 | ITGA6 | p.Val266Leu | missense_variant | MODERATE | 0.30303 | 0 | 33 | 156 | . | . | . | . |
| PM1548-SK1P3 | SP3 | p.Val461Ile | missense_variant | MODERATE | 0.16875 | 0 | 160 | 262 | . | . | . | . |
| PM1548-SK1P3 | EVX2 | p.Gly438Ser | missense_variant | MODERATE | 1 | 0 | 135 | 165 | . | . | . | . |
| PM1548-SK1P3 | EVX2 | p.Asp320Glu | missense_variant | MODERATE | 0 | 0 | 11 | 48 | . | . | . | . |
| PM1548-SK1P3 | CCDC141 | p.Ala1030Val | missense_variant | MODERATE | 0 | 0 | 61 | 69 | . | . | . | . |
| PM1548-SK1P3 | PDE1A | p.Ala371Val | missense_variant | MODERATE | 0.933333 | 0 | 15 | 75 | . | . | . | . |
| PM1548-SK1P3 | FSIP2 | p.Arg829Cys | missense_variant | MODERATE | 0.2 | 0 | 15 | 51 | . | . | . | . |
| PM1548-SK1P3 | DNAH7 | p.Ala3252Glu | missense_variant | MODERATE | 0.0645161 | 0 | 124 | 64 | . | . | . | . |
| PM1548-SK1P3 | GTF3C3 | p.Pro248Leu | missense_variant | MODERATE | 0.285714 | 0 | 35 | 201 | . | . | . | . |
| PM1548-SK1P3 | AOX1 | p.Phe1162Ser | missense_variant | MODERATE | 0.150442 | 0 | 113 | 36 | . | . | . | . |
| PM1548-SK1P3 | PIKFYVE | p.Leu419Leu | synonymous_variant | LOW | 0.342857 | 0 | 35 | 192 | . | . | . | . |
| PM1548-SK1P3 | MAP2 | p.Gly1784Ser | missense_variant | MODERATE | 0.705882 | 0 | 34 | 92 | . | . | . | . |
| PM1548-SK1P3 | SMARCAL1 | p.Arg499Gln | missense_variant | MODERATE | 0.68 | 0 | 175 | 137 | . | . | . | . |
| PM1548-SK1P3 | AAMP | p.Glu63Asp | missense_variant | MODERATE | 0.268293 | 0 | 82 | 163 | . | . | . | . |
| PM1548-SK1P3 | ZNF142 | p.Ala335Pro | missense_variant | MODERATE | 0 | 0 | 45 | 317 | . | . | . | . |
| PM1548-SK1P3 | CYP27A1 | p.Ser203Leu | missense_variant | MODERATE | 0.333333 | 0 | 15 | 39 | . | . | . | . |
| PM1548-SK1P3 | WNT6 | p.Asp283Val | missense_variant | MODERATE | 0.408669 | 0 | 323 | 120 | . | . | . | . |
| PM1548-SK1P3 | FAM134A | p.Asp391Val | missense_variant | MODERATE | 0.378788 | 0 | 66 | 285 | . | . | . | . |
| PM1548-SK1P3 | DNPEP | p.Arg191Gln | missense_variant | MODERATE | 0 | 0 | 29 | 296 | . | . | . | . |
| PM1548-SK1P3 | CHPF | p.Arg721Trp | missense_variant | MODERATE | 0.1375 | 0 | 560 | 74 | . | . | . | . |
| PM1548-SK1P3 | WDFY1 | p.Ala75Asp | missense_variant | MODERATE | 0 | 0 | 22 | 181 | . | . | . | . |
| PM1548-SK1P3 | SERPINE2 | p.Met296Val | missense_variant | MODERATE | 1 | 0 | 6 | 111 | . | . | . | . |
| PM1548-SK1P3 | B3GNT7 | p.Gly302Asp | missense_variant | MODERATE | 0.666667 | 0 | 3 | 60 | . | . | . | . |
| PM1548-SK1P3 | DGKD | p.Arg935Trp | missense_variant | MODERATE | 0 | 0 | 11 | 67 | . | . | . | . |
| PM1548-SK1P3 | ACKR3 | p.Val42Ala | missense_variant | MODERATE | 0.608696 | 0 | 46 | 137 | true | . | . | . |
| PM1548-SK1P3 | COL6A3 | p.Gly1714Glu | missense_variant | MODERATE | 0.714286 | 0 | 14 | 50 | . | . | . | . |
| PM1548-SK1P3 | COL6A3 | p.Ile1609Asn | missense_variant | MODERATE | 0.307692 | 0 | 26 | 113 | . | . | . | . |
| PM1548-SK1P3 | ING5 | p.Arg106His | missense_variant | MODERATE | 0 | 0 | 76 | 105 | . | . | . | . |
| PM1548-SK1P3 | TBC1D20 | p.Pro71Ser | missense_variant | MODERATE | 0 | 0 | 25 | 49 | . | . | . | . |
| PM1548-SK1P3 | VPS16 | p.Thr818Met | missense_variant | MODERATE | 0.439353 | 0 | 371 | 238 | . | . | . | . |
| PM1548-SK1P3 | ATRN | p.Ala122Thr | missense_variant | MODERATE | 0.333333 | 0 | 108 | 154 | . | . | . | . |
| PM1548-SK1P3 | SLC23A2 | p.Ile412fs | frameshift_variant | HIGH | 0.37931 | 0 | 29 | 150 | . | . | . | . |
| PM1548-SK1P3 | BMP2 | p.Ala10Val | missense_variant | MODERATE | 0.276923 | 0 | 65 | 162 | . | . | . | . |
| PM1548-SK1P3 | PLCB1 | p.Arg768Gln | missense_variant | MODERATE | 0 | 0 | 13 | 91 | . | . | . | . |
| PM1548-SK1P3 | JAG1 | p.Gly537Ser | missense_variant | MODERATE | 0.0169492 | 0 | 59 | 184 | . | . | . | . |
| PM1548-SK1P3 | CSRP2BP | p.Trp173* | stop_gained | HIGH | 0.545455 | 0 | 11 | 168 | . | . | . | . |
| PM1548-SK1P3 | CRNKL1 | p.Phe352Leu | missense_variant | MODERATE | 0.684211 | 0 | 19 | 71 | . | . | . | . |
| PM1548-SK1P3 | PLAGL2 | p.Arg180His | missense_variant | MODERATE | 0.614286 | 0 | 140 | 145 | . | . | . | . |
| PM1548-SK1P3 | BPIFA2 | p.Arg221His | missense_variant | MODERATE | 0.53125 | 0 | 32 | 61 | . | . | . | . |
| PM1548-SK1P3 | NECAB3 | p.Arg278fs | frameshift_variant | HIGH | 1 | 0 | 1 | 39 | . | . | . | . |
| PM1548-SK1P3 | CEP250 | p.Lys787Glu | missense_variant | MODERATE | 0 | 0 | 65 | 64 | . | . | . | . |
| PM1548-SK1P3 | SOGA1 | p.Pro974Ser | missense_variant&splice_region_variant | MODERATE | 0.205882 | 0 | 34 | 259 | . | . | . | . |
| PM1548-SK1P3 | PLCG1 | p.Ile942Met | missense_variant | MODERATE | 0.0619469 | 0 | 226 | 140 | true | . | . | . |
| PM1548-SK1P3 | ZSWIM3 | p.Arg225Gln | missense_variant | MODERATE | 0.482759 | 0 | 29 | 55 | . | . | . | . |
| PM1548-SK1P3 | NFATC2 | p.Phe158Tyr | missense_variant | MODERATE | 0.0434783 | 0 | 92 | 59 | true | . | . | . |
| PM1548-SK1P3 | NFATC2 | p.Gly22fs | frameshift_variant | HIGH | 0.119266 | 0 | 109 | 146 | true | . | . | . |
| PM1548-SK1P3 | SALL4 | p.Phe391Leu | missense_variant | MODERATE | 0.144 | 0 | 125 | 33 | . | . | . | . |
| PM1548-SK1P3 | ZFP64 | n.50701484A>G | intragenic_variant | MODIFIER | 0.142857 | 0 | 28 | 142 | . | . | . | . |
| PM1548-SK1P3 | DOK5 | p.Val169Ala | missense_variant | MODERATE | 0.267442 | 0 | 258 | 338 | . | . | . | . |
| PM1548-SK1P3 | DOK5 | p.Ala267Asp | missense_variant | MODERATE | 0.32687 | 0 | 361 | 145 | . | . | . | . |
| PM1548-SK1P3 | TFAP2C | p.Ala354Val | missense_variant | MODERATE | 0.512195 | 0 | 41 | 269 | . | . | . | . |
| PM1548-SK1P3 | PMEPA1 | p.Ser209fs | frameshift_variant | HIGH | 0 | 0 | 580 | 249 | . | . | . | . |
| PM1548-SK1P3 | NELFCD | p.Val321Ile | missense_variant | MODERATE | 0.513514 | 0 | 37 | 73 | . | . | . | . |
| PM1548-SK1P3 | ZNF831 | p.Leu434Met | missense_variant | MODERATE | 0.405405 | 0 | 185 | 129 | . | . | . | . |
| PM1548-SK1P3 | HELZ2 | p.Ala639Thr | missense_variant | MODERATE | 0.0808081 | 0 | 396 | 139 | . | . | . | . |
| PM1548-SK1P3 | RTEL1 | p.Arg729Cys | missense_variant | MODERATE | 0.517647 | 0 | 170 | 260 | . | . | . | . |
| PM1548-SK1P3 | TNFRSF6B | p.Leu297Phe | missense_variant | MODERATE | 0 | 0 | 261 | 277 | . | . | . | . |
| PM1548-SK1P3 | SLC2A4RG | p.Arg82fs | frameshift_variant | HIGH | 0.204082 | 0 | 49 | 206 | . | . | . | . |
| PM1548-SK1P3 | PRPF6 | p.Arg317His | missense_variant | MODERATE | 0.235294 | 0 | 34 | 86 | . | . | . | . |
| PM1548-SK1P3 | LTN1 | p.Gly1705Val | missense_variant | MODERATE | 0.580645 | 0 | 31 | 36 | . | . | . | . |
| PM1548-SK1P3 | TIAM1 | p.His1010Tyr | missense_variant | MODERATE | 0.192771 | 0 | 83 | 418 | . | . | . | . |
| PM1548-SK1P3 | URB1 | p.Pro2036Leu | missense_variant | MODERATE | 0.666667 | 0 | 6 | 14 | . | . | . | . |
| PM1548-SK1P3 | PAXBP1 | p.Pro161Thr | missense_variant | MODERATE | 0 | 0 | 14 | 40 | . | . | . | . |
| PM1548-SK1P3 | PAXBP1 | p.Ser105Arg | missense_variant | MODERATE | 0 | 0 | 180 | 158 | . | . | . | . |
| PM1548-SK1P3 | GART | p.Glu297Gln | missense_variant | MODERATE | 0.432432 | 0 | 74 | 160 | . | . | . | . |
| PM1548-SK1P3 | SON | p.Thr790Ser | missense_variant | MODERATE | 0.382353 | 0 | 34 | 177 | . | . | . | . |
| PM1548-SK1P3 | SON | c.6657+163T>C | intron_variant | MODIFIER | 0.255319 | 0 | 94 | 33 | . | . | . | . |
| PM1548-SK1P3 | ITSN1 | p.Met768Thr | missense_variant | MODERATE | 0.5 | 0 | 8 | 46 | . | . | . | true |
| PM1548-SK1P3 | SLC5A3 | p.Arg252Gln | missense_variant | MODERATE | 0.433962 | 0 | 53 | 44 | . | . | . | . |
| PM1548-SK1P3 | UMODL1 | p.Gly1319Ser | missense_variant | MODERATE | 0.518519 | 0 | 108 | 131 | . | . | . | . |
| PM1548-SK1P3 | CRYAA | p.Ile61Val | missense_variant | MODERATE | 0.441758 | 0 | 455 | 344 | . | . | . | . |
| PM1548-SK1P3 | SLC19A1 | p.Asp488Gly | missense_variant | MODERATE | 0.0666667 | 0 | 165 | 131 | . | . | . | . |
| PM1548-SK1P3 | FTCD | p.Ile187Thr | missense_variant | MODERATE | 0.146341 | 0 | 41 | 175 | . | . | . | . |
| PM1548-SK1P3 | LSS | p.Thr457Met | missense_variant | MODERATE | 0.591398 | 0 | 372 | 295 | . | . | . | . |
| PM1548-SK1P3 | CECR2 | p.Ser425Ser | synonymous_variant | LOW | 0.428571 | 0 | 112 | 131 | . | . | . | . |
| PM1548-SK1P3 | CECR2 | p.Pro428Pro | synonymous_variant | LOW | 0.461538 | 0 | 104 | 130 | . | . | . | . |
| PM1548-SK1P3 | CECR2 | p.Tyr431Tyr | synonymous_variant | LOW | 0.466019 | 0 | 103 | 130 | . | . | . | . |
| PM1548-SK1P3 | ATP6V1E1 | p.Leu128Met | missense_variant | MODERATE | 0.423729 | 0 | 59 | 148 | . | . | . | . |
| PM1548-SK1P3 | CLTCL1 | p.Ala1567Thr | missense_variant | MODERATE | 0.495074 | 0 | 406 | 279 | true | . | . | . |
| PM1548-SK1P3 | SCARF2 | p.Arg335Cys | missense_variant | MODERATE | 0 | 0 | 9 | 100 | . | . | . | . |
| PM1548-SK1P3 | SLC7A4 | p.Ala256Thr | missense_variant | MODERATE | 0.653061 | 0 | 49 | 223 | . | . | . | . |
| PM1548-SK1P3 | IGLL1 | p.Arg60Gln | missense_variant | MODERATE | 0 | 0 | 30 | 81 | . | . | . | . |
| PM1548-SK1P3 | C22orf43 | p.Ser153Gly | missense_variant | MODERATE | 0 | 0 | 13 | 97 | . | . | . | . |
| PM1548-SK1P3 | ASPHD2 | p.Arg358Trp | missense_variant | MODERATE | 0.54878 | 0 | 82 | 411 | . | . | . | . |
| PM1548-SK1P3 | TFIP11 | p.Arg39* | stop_gained | HIGH | 0.460976 | 0 | 410 | 103 | . | . | . | . |
| PM1548-SK1P3 | RASL10A | p.Arg108Trp | missense_variant | MODERATE | 0 | 0 | 22 | 108 | . | . | . | . |
| PM1548-SK1P3 | AP1B1 | p.Ala655Asp | missense_variant | MODERATE | 0 | 0 | 20 | 31 | . | . | . | . |
| PM1548-SK1P3 | LIMK2 | p.Asn477Ser | missense_variant | MODERATE | 0.266833 | 0 | 401 | 257 | . | . | . | . |
| PM1548-SK1P3 | SFI1 | p.Thr710Ser | missense_variant | MODERATE | 0.285714 | 0 | 28 | 104 | . | . | . | . |
| PM1548-SK1P3 | CSF2RB | p.Gln642fs | frameshift_variant | HIGH | 0.584041 | 0 | 589 | 133 | . | . | . | . |
| PM1548-SK1P3 | C1QTNF6 | p.Met229Thr | missense_variant | MODERATE | 0.464789 | 0 | 71 | 41 | . | . | . | . |
| PM1548-SK1P3 | GRAP2 | p.Val129Ile | missense_variant | MODERATE | 0.491525 | 0 | 236 | 162 | . | . | . | . |
| PM1548-SK1P3 | MKL1 | p.Pro138Leu | missense_variant | MODERATE | 0.440252 | 0 | 159 | 100 | true | . | . | . |
| PM1548-SK1P3 | EP300 | p.Pro2107Thr | missense_variant | MODERATE | 0.151515 | 0 | 33 | 94 | true | true | true | . |
| PM1548-SK1P3 | ZC3H7B | p.Arg908Cys | missense_variant | MODERATE | 0.608108 | 0 | 74 | 73 | . | . | . | . |
| PM1548-SK1P3 | PARVB | c.211+24725C>A | intron_variant | MODIFIER | 0.428571 | 0 | 77 | 165 | . | . | . | . |
| PM1548-SK1P3 | CELSR1 | p.Val1411Gly | missense_variant | MODERATE | 0.117284 | 0 | 162 | 333 | . | . | . | . |
| PM1548-SK1P3 | CELSR1 | p.Glu1159Lys | missense_variant | MODERATE | 0.5 | 0 | 24 | 186 | . | . | . | . |
| PM1548-SK1P3 | TUBGCP6 | p.Cys1239Trp | missense_variant | MODERATE | 0.5 | 0.00947867 | 232 | 211 | . | . | . | . |
| PM1548-SK1P3 | TUBGCP6 | p.Ser1237Pro | missense_variant | MODERATE | 0.284768 | 0.00420168 | 151 | 238 | . | . | . | . |
| PM1548-SK1P3 | MIOX | p.Asp78Gly | missense_variant | MODERATE | 0 | 0 | 101 | 102 | . | . | . | . |
| PM1548-SK1P3 | MAPK8IP2 | p.Thr518Thr | synonymous_variant | LOW | 0.565041 | 0 | 246 | 166 | . | . | . | . |
| PM1548-SK1P3 | ARSA | c.684+1G>A | splice_donor_variant&intron_variant | HIGH | 0.485915 | 0 | 142 | 160 | . | . | . | . |

| | | | | | | | | | | | | |
|---|---|---|---|---|---|---|---|---|---|---|---|---|
| PM1548-SK1P3 | SETD5 | p.Arg1193* | stop_gained | HIGH | 0.444444 | 0 | 135 | 243 | . | . | . | . |
| PM1548-SK1P3 | CPNE9 | p.Ser284Ser | synonymous_variant | LOW | 0.274286 | 0 | 175 | 196 | . | . | . | . |
| PM1548-SK1P3 | CPNE9 | p.Asp473Asp | synonymous_variant | LOW | 0.469333 | 0 | 375 | 85 | . | . | . | . |
| PM1548-SK1P3 | BRPF1 | p.Gln938fs | frameshift_variant | HIGH | 0.222222 | 0 | 9 | 52 | . | . | . | true |
| PM1548-SK1P3 | OGG1 | p.Ala85Thr | missense_variant | MODERATE | 0.366667 | 0 | 60 | 334 | . | . | . | . |
| PM1548-SK1P3 | CRELD1 | p.Arg220* | stop_gained | HIGH | 0.376642 | 0 | 685 | 387 | . | . | . | . |
| PM1548-SK1P3 | PRRT3 | p.His679Arg | missense_variant | MODERATE | 0.62 | 0 | 100 | 166 | . | . | . | . |
| PM1548-SK1P3 | CAND2 | p.Arg489His | missense_variant | MODERATE | 0.608108 | 0 | 74 | 331 | . | . | . | . |
| PM1548-SK1P3 | SLC6A6 | p.Phe392fs | frameshift_variant | HIGH | 0.524887 | 0 | 663 | 115 | . | . | . | . |
| PM1548-SK1P3 | NR2C2 | p.Phe582fs | frameshift_variant | HIGH | 0.357759 | 0 | 464 | 75 | . | . | . | . |
| PM1548-SK1P3 | SH3BP5 | p.Thr119Ala | missense_variant | MODERATE | 0.416092 | 0 | 435 | 345 | . | . | . | . |
| PM1548-SK1P3 | PLCL2 | p.Gln628Arg | missense_variant | MODERATE | 0.30303 | 0 | 33 | 118 | . | . | . | . |
| PM1548-SK1P3 | SLC4A7 | p.Ile1097Val | missense_variant | MODERATE | 0.355932 | 0.00657895 | 59 | 152 | . | . | . | . |
| PM1548-SK1P3 | CLASP2 | p.Cys10Cys | synonymous_variant | LOW | 0.451299 | 0.00520833 | 308 | 192 | . | . | . | . |
| PM1548-SK1P3 | SCN10A | p.Val378Leu | missense_variant | MODERATE | 0.0625 | 0.00809717 | 112 | 247 | . | . | . | . |
| PM1548-SK1P3 | TTC21A | p.Ser332Thr | missense_variant | MODERATE | 0.185185 | 0 | 54 | 40 | . | . | . | . |
| PM1548-SK1P3 | SNRK | p.Pro598Thr | missense_variant | MODERATE | 0 | 0 | 185 | 93 | . | . | . | . |
| PM1548-SK1P3 | SLC6A20 | p.Asn103Lys | missense_variant | MODERATE | 0 | 0 | 295 | 172 | . | . | . | . |
| PM1548-SK1P3 | SLC6A20 | p.Gly66Trp | missense_variant | MODERATE | 0.0392157 | 0.00847458 | 51 | 118 | . | . | . | . |
| PM1548-SK1P3 | FYCO1 | p.Lys501del | inframe_deletion | MODERATE | 0.434783 | 0 | 253 | 171 | . | . | . | . |
| PM1548-SK1P3 | ALS2CL | p.Asn384Lys | missense_variant | MODERATE | 0.2 | 0 | 30 | 77 | . | . | . | . |
| PM1548-SK1P3 | DHX30 | n.47859525delC | intragenic_variant | MODIFIER | 0.424603 | 0 | 252 | 119 | . | . | . | . |
| PM1548-SK1P3 | UCN2 | p.Thr27Pro | missense_variant | MODERATE | 0.114713 | 0 | 401 | 135 | . | . | . | . |
| PM1548-SK1P3 | CELSR3 | p.Pro1425Ser | missense_variant | MODERATE | 0 | 0 | 117 | 95 | . | . | . | . |
| PM1548-SK1P3 | SLC25A20 | p.Ala227Val | missense_variant | MODERATE | 0.444444 | 0 | 18 | 126 | . | . | . | . |
| PM1548-SK1P3 | KLHDC8B | p.Gly38fs | frameshift_variant | HIGH | 0.229167 | 0 | 96 | 199 | . | . | . | . |
| PM1548-SK1P3 | RRP9 | p.Ala342Thr | missense_variant | MODERATE | 0.37619 | 0 | 210 | 161 | . | . | . | . |
| PM1548-SK1P3 | DNAH1 | p.Leu2072Ile | missense_variant | MODERATE | 0.47541 | 0 | 61 | 99 | . | . | . | . |
| PM1548-SK1P3 | FAM208A | p.Ala227Val | missense_variant | MODERATE | 0.588235 | 0 | 34 | 80 | . | . | . | . |
| PM1548-SK1P3 | DNAH12 | p.Pro1035Ala | missense_variant | MODERATE | 0.16 | 0 | 75 | 189 | . | . | . | . |
| PM1548-SK1P3 | PDE12 | p.Ala293Ser | missense_variant | MODERATE | 0.289474 | 0 | 38 | 150 | . | . | . | . |
| PM1548-SK1P3 | PDE12 | p.Leu294Phe | missense_variant | MODERATE | 0.289474 | 0 | 38 | 150 | . | . | . | . |
| PM1548-SK1P3 | ADAMTS9 | p.Pro596His | missense_variant | MODERATE | 0.7 | 0 | 40 | 126 | . | . | . | . |
| PM1548-SK1P3 | ADAMTS9 | p.Ser593Gly | missense_variant | MODERATE | 0.682927 | 0 | 41 | 130 | . | . | . | . |
| PM1548-SK1P3 | ROBO1 | p.Tyr1009* | stop_gained | HIGH | 0 | 0 | 43 | 54 | . | . | . | . |
| PM1548-SK1P3 | OR5K4 | p.Val51Ile | missense_variant | MODERATE | 0.333333 | 0 | 84 | 441 | . | . | . | . |
| PM1548-SK1P3 | OR5K4 | p.Arg53Pro | missense_variant | MODERATE | 0.333333 | 0 | 84 | 437 | . | . | . | . |
| PM1548-SK1P3 | OR5K4 | p.Leu56His | missense_variant | MODERATE | 0.27451 | 0 | 102 | 510 | . | . | . | . |
| PM1548-SK1P3 | DCBLD2 | p.Thr718Met | missense_variant | MODERATE | 0.514286 | 0 | 315 | 81 | . | . | . | . |
| PM1548-SK1P3 | TFG | p.Ser213Pro | missense_variant | MODERATE | 0.460938 | 0 | 128 | 76 | true | . | . | . |
| PM1548-SK1P3 | CBLB | p.Pro823Ser | missense_variant | MODERATE | 0.117647 | 0 | 17 | 275 | true | . | . | . |
| PM1548-SK1P3 | BOC | p.Pro593Leu | missense_variant | MODERATE | 0.485876 | 0.00199203 | 177 | 502 | . | . | . | . |
| PM1548-SK1P3 | BOC | p.His601Gln | missense_variant | MODERATE | 0.359649 | 0 | 114 | 258 | . | . | . | . |
| PM1548-SK1P3 | KIAA2018 | p.Asn1016fs | frameshift_variant | HIGH | 0.0313901 | 0 | 223 | 36 | . | . | . | . |
| PM1548-SK1P3 | KIAA1407 | p.Glu683Gly | missense_variant | MODERATE | 0.0625 | 0 | 16 | 85 | . | . | . | . |
| PM1548-SK1P3 | LSAMP | p.Val277Ile | missense_variant | MODERATE | 0.514793 | 0 | 169 | 91 | . | . | . | . |
| PM1548-SK1P3 | ARHGAP31 | p.Lys857Glu | missense_variant | MODERATE | 0 | 0 | 3 | 33 | . | . | . | . |
| PM1548-SK1P3 | FSTL1 | p.Val13Ala | missense_variant | MODERATE | 0.401639 | 0 | 122 | 223 | . | . | . | . |
| PM1548-SK1P3 | EAF2 | p.Thr112fs | frameshift_variant | HIGH | 0.6 | 0 | 20 | 119 | . | . | . | . |
| PM1548-SK1P3 | ILDR1 | p.Ala76Val | missense_variant&splice_region_variant | MODERATE | 0.464286 | 0 | 28 | 159 | . | . | . | . |
| PM1548-SK1P3 | MYLK | p.Ala1684Thr | missense_variant | MODERATE | 0.244275 | 0 | 262 | 98 | . | . | . | . |
| PM1548-SK1P3 | ALDH1L1 | p.Glu439fs | frameshift_variant | HIGH | 0.489691 | 0 | 194 | 55 | . | . | . | . |
| PM1548-SK1P3 | CHCHD6 | p.Ser220Leu | missense_variant | MODERATE | 0 | 0 | 220 | 164 | . | . | . | . |
| PM1548-SK1P3 | C3orf27 | p.Asp85fs | frameshift_variant | HIGH | 0.420455 | 0 | 176 | 229 | . | . | . | . |
| PM1548-SK1P3 | TMCC1 | p.Arg620Ser | missense_variant | MODERATE | 0 | 0 | 132 | 62 | . | . | . | . |
| PM1548-SK1P3 | CDV3 | p.Gly226Asp | missense_variant | MODERATE | 0.108108 | 0 | 37 | 44 | . | . | . | . |
| PM1548-SK1P3 | CDV3 | p.Asp228Glu | missense_variant | MODERATE | 0.153846 | 0 | 39 | 55 | . | . | . | . |
| PM1548-SK1P3 | AMOTL2 | p.Arg292Cys | missense_variant | MODERATE | 0 | 0 | 66 | 43 | . | . | . | . |
| PM1548-SK1P3 | EPHB1 | p.Ala82fs | frameshift_variant | HIGH | 0.0831234 | 0 | 397 | 300 | . | . | . | . |
| PM1548-SK1P3 | SLC35G2 | p.Asn97fs | frameshift_variant | HIGH | 0.25 | 0 | 4 | 31 | . | . | . | . |
| PM1548-SK1P3 | ESYT3 | p.Val238Met | missense_variant | MODERATE | 0.468504 | 0 | 254 | 78 | . | . | . | . |
| PM1548-SK1P3 | MED12L | p.Ser970Leu | missense_variant | MODERATE | 0.375 | 0 | 16 | 123 | . | . | . | . |
| PM1548-SK1P3 | DHX36 | p.Met154fs | frameshift_variant | HIGH | 0 | 0 | 32 | 44 | . | . | . | . |
| PM1548-SK1P3 | CCNL1 | p.Ile354Val | missense_variant | MODERATE | 0.25 | 0 | 56 | 120 | . | . | . | . |
| PM1548-SK1P3 | PPM1L | p.Glu106Glu | synonymous_variant | LOW | 0.363636 | 0 | 55 | 274 | . | . | . | . |
| PM1548-SK1P3 | SLITRK3 | p.Arg759Cys | missense_variant | MODERATE | 0.54878 | 0 | 164 | 99 | . | . | . | . |
| PM1548-SK1P3 | ZBBX | p.Ile496fs | frameshift_variant | HIGH | 0.35 | 0 | 20 | 76 | . | . | . | . |
| PM1548-SK1P3 | FNDC3B | p.Glu687Gly | missense_variant | MODERATE | 0.533333 | 0 | 30 | 96 | . | . | . | . |
| PM1548-SK1P3 | NCEH1 | p.Ser105fs | frameshift_variant | HIGH | 0.30137 | 0 | 73 | 227 | . | . | . | . |
| PM1548-SK1P3 | KLHL24 | p.Asn555fs | frameshift_variant | HIGH | 0 | 0 | 7 | 55 | . | . | . | . |
| PM1548-SK1P3 | CHRD | p.Arg76Cys | missense_variant | MODERATE | 0.695652 | 0 | 23 | 91 | . | . | . | . |
| PM1548-SK1P3 | TP63 | p.Thr368Ala | missense_variant | MODERATE | 0.52381 | 0 | 147 | 121 | . | . | . | . |
| PM1548-SK1P3 | LEPREL1 | p.Asp204Glu | missense_variant | MODERATE | 0 | 0 | 60 | 177 | . | . | . | . |
| PM1548-SK1P3 | CCDC50 | p.Ala343Ser | missense_variant | MODERATE | 0 | 0 | 18 | 55 | . | . | . | . |
| PM1548-SK1P3 | MUC20 | p.Gly514Arg | missense_variant | MODERATE | 0.282051 | 0 | 39 | 228 | . | . | . | . |
| PM1548-SK1P3 | LRCH3 | p.Leu49Val | missense_variant | MODERATE | 0.181818 | 0.00369004 | 44 | 271 | . | . | . | . |
| PM1548-SK1P3 | GAK | p.Ala690Val | missense_variant | MODERATE | 0 | 0 | 91 | 111 | . | . | . | . |
| PM1548-SK1P3 | DGKQ | p.Thr299Met | missense_variant | MODERATE | 0.5209 | 0 | 311 | 253 | . | . | . | . |
| PM1548-SK1P3 | SLC26A1 | p.Ala624Thr | missense_variant | MODERATE | 0 | 0 | 92 | 466 | . | . | . | . |
| PM1548-SK1P3 | POLN | p.Tyr686Cys | missense_variant | MODERATE | 0 | 0 | 99 | 37 | . | . | . | . |
| PM1548-SK1P3 | HTT | p.Ala2259Thr | missense_variant&splice_region_variant | MODERATE | 0.142857 | 0 | 7 | 52 | . | . | . | . |
| PM1548-SK1P3 | PPP2R2C | p.Gly397fs | frameshift_variant | HIGH | 0 | 0 | 44 | 188 | . | . | . | . |
| PM1548-SK1P3 | SH3TC1 | p.Glu203Gly | missense_variant | MODERATE | 0.133333 | 0.0021322 | 30 | 469 | . | . | . | . |
| PM1548-SK1P3 | GPR78 | p.Val209Ile | missense_variant | MODERATE | 0.607692 | 0 | 130 | 34 | . | . | . | . |
| PM1548-SK1P3 | KCTD8 | p.Leu62Leu | synonymous_variant | LOW | 0.268116 | 0 | 138 | 212 | . | . | . | . |
| PM1548-SK1P3 | YTHDC1 | p.Ser684Asn | missense_variant | MODERATE | 0.125 | 0 | 8 | 59 | . | . | . | . |
| PM1548-SK1P3 | ANKRD17 | p.Ile2241Val | missense_variant | MODERATE | 0.3 | 0 | 30 | 146 | . | . | . | . |
| PM1548-SK1P3 | PF4 | p.Ala4Thr | missense_variant | MODERATE | 0.121212 | 0 | 99 | 40 | . | . | . | . |
| PM1548-SK1P3 | PARM1 | p.Ser82Pro | missense_variant | MODERATE | 0 | 0 | 32 | 388 | . | . | . | . |
| PM1548-SK1P3 | CDKL2 | p.Pro452Ser | missense_variant | MODERATE | 0 | 0 | 9 | 51 | . | . | . | . |
| PM1548-SK1P3 | SHROOM3 | p.Lys1017del | inframe_deletion | MODERATE | 0.462617 | 0 | 214 | 97 | . | . | . | . |
| PM1548-SK1P3 | BMP3 | p.Lys358fs | frameshift_variant | HIGH | 0.5 | 0 | 30 | 124 | . | . | . | . |
| PM1548-SK1P3 | PRKG2 | p.Val461Phe | missense_variant | MODERATE | 0 | 0 | 8 | 44 | . | . | . | . |
| PM1548-SK1P3 | SEC31A | p.Ile462fs | frameshift_variant | HIGH | 0.967741935 | 0 | 31 | 87 | . | . | . | . |
| PM1548-SK1P3 | SEC31A | p.Lys461fs | frameshift_variant | HIGH | 0 | 0 | 31 | 87 | . | . | . | . |
| PM1548-SK1P3 | SEC31A | p.Gln422Pro | missense_variant | MODERATE | 0.134328 | 0 | 67 | 143 | . | . | . | . |
| PM1548-SK1P3 | AGPAT9 | p.Met372fs | frameshift_variant | HIGH | 0.61165 | 0 | 103 | 70 | . | . | . | . |
| PM1548-SK1P3 | NAP1L5 | p.Lys108fs | frameshift_variant | HIGH | 0.502538 | 0 | 197 | 237 | . | . | . | . |
| PM1548-SK1P3 | CASP6 | p.Pro33Leu | missense_variant | MODERATE | 0.4 | 0.00722022 | 30 | 277 | . | . | . | . |
| PM1548-SK1P3 | PITX2 | c.411+284C>T | intron_variant | MODIFIER | 0.509632 | 0 | 571 | 59 | . | . | . | . |
| PM1548-SK1P3 | ALPK1 | p.Tyr1216fs | frameshift_variant | HIGH | 0.5 | 0 | 4 | 44 | . | . | . | . |
| PM1548-SK1P3 | TNIP3 | p.Ala59Thr | missense_variant | MODERATE | 0.25 | 0 | 4 | 33 | . | . | . | . |
| PM1548-SK1P3 | PCDH10 | p.Tyr324Cys | missense_variant | MODERATE | 0.214815 | 0 | 135 | 50 | . | . | . | . |
| PM1548-SK1P3 | SETD7 | p.Leu331Ile | missense_variant | MODERATE | 0.502075 | 0 | 241 | 328 | . | . | . | . |
| PM1548-SK1P3 | FREM3 | p.Arg502* | stop_gained | HIGH | 0 | 0 | 10 | 34 | . | . | . | . |
| PM1548-SK1P3 | NEK1 | p.Thr137Asn | missense_variant | MODERATE | 0.0135135 | 0 | 74 | 84 | . | . | . | . |
| PM1548-SK1P3 | AHRR | p.Arg72His | missense_variant | MODERATE | 0.425532 | 0 | 376 | 75 | . | . | . | . |
| PM1548-SK1P3 | EXOC3 | p.Val576Met | missense_variant | MODERATE | 0.567568 | 0 | 37 | 107 | . | . | . | . |
| PM1548-SK1P3 | LPCAT1 | p.Val411Ile | missense_variant | MODERATE | 0 | 0 | 152 | 136 | . | . | . | . |
| PM1548-SK1P3 | IRX2 | p.Tyr110Cys | missense_variant | MODERATE | 0.514056 | 0 | 249 | 189 | . | . | . | . |
| PM1548-SK1P3 | NSUN2 | p.Ala653Thr | missense_variant&splice_region_variant | MODERATE | 0.128713 | 0 | 101 | 83 | . | . | . | . |
| PM1548-SK1P3 | SRD5A1 | p.Met162Ile | missense_variant | MODERATE | 0.445652 | 0 | 92 | 54 | . | . | . | . |
| PM1548-SK1P3 | SEMA5A | p.Ile679Thr | missense_variant | MODERATE | 0.0208333 | 0 | 96 | 101 | . | . | . | . |
| PM1548-SK1P3 | NIPBL | p.Glu1060Glu | synonymous_variant | LOW | 0.3 | 0 | 40 | 120 | . | . | . | . |
| PM1548-SK1P3 | CARD6 | p.Gln393Glu | missense_variant | MODERATE | 0.461538 | 0 | 52 | 370 | . | . | . | . |
| PM1548-SK1P3 | CARD6 | p.Gln400His | missense_variant | MODERATE | 0.45283 | 0 | 53 | 367 | . | . | . | . |
| PM1548-SK1P3 | CARD6 | p.Gln402* | stop_gained | HIGH | 0.45283 | 0 | 53 | 367 | . | . | . | . |
| PM1548-SK1P3 | CARD6 | p.Gln402Arg | missense_variant | MODERATE | 0.45283 | 0 | 53 | 337 | . | . | . | . |
| PM1548-SK1P3 | CARD6 | p.Gln402His | missense_variant | MODERATE | 0.461538 | 0 | 52 | 337 | . | . | . | . |
| PM1548-SK1P3 | C6 | p.Ala2Val | missense_variant | MODERATE | 0.325 | 0 | 80 | 129 | . | . | . | . |
| PM1548-SK1P3 | SNX18 | n.53839150G>A | intragenic_variant | MODIFIER | 0.586207 | 0 | 29 | 117 | . | . | . | . |
| PM1548-SK1P3 | GPX8 | p.Phe125fs | frameshift_variant | HIGH | 0.466667 | 0 | 75 | 228 | . | . | . | . |
| PM1548-SK1P3 | UTP15 | c.169-826A>G | intron_variant | MODIFIER | 0 | 0 | 60 | 55 | . | . | . | . |
| PM1548-SK1P3 | TTC37 | p.Ala1407Thr | missense_variant | MODERATE | 0.714286 | 0 | 7 | 33 | . | . | . | . |
| PM1548-SK1P3 | CHD1 | p.Asn310fs | frameshift_variant | HIGH | 0.940299 | 0 | 67 | 201 | . | . | . | true |
| PM1548-SK1P3 | PAM | p.Arg260Gln | missense_variant | MODERATE | 0 | 0 | 45 | 107 | . | . | . | . |
| PM1548-SK1P3 | GIN1 | p.Leu345Pro | missense_variant | MODERATE | 0.4 | 0 | 15 | 72 | . | . | . | . |

| | | | | | | | | | | | | |
|---|---|---|---|---|---|---|---|---|---|---|---|---|
| PM1548-SK1P3 | APC | p.Asp802fs | frameshift_variant&stop_gained | HIGH | 0.498956 | 0 | 479 | 125 | true | true | true | true |
| PM1548-SK1P3 | MCC | p.Arg800Lys | missense_variant | MODERATE | 0.8 | 0 | 5 | 65 | . | . | . | . |
| PM1548-SK1P3 | FTMT | p.Ala90Val | missense_variant | MODERATE | 0.578947 | 0 | 285 | 105 | . | . | . | . |
| PM1548-SK1P3 | FBN2 | p.Leu2853Arg | missense_variant | MODERATE | 0.494737 | 0 | 95 | 91 | . | . | . | . |
| PM1548-SK1P3 | FNIP1 | p.Leu90Met | missense_variant | MODERATE | 0 | 0 | 8 | 76 | . | . | . | . |
| PM1548-SK1P3 | JADE2 | p.Phe360Leu | missense_variant | MODERATE | 0.202899 | 0 | 207 | 343 | . | . | . | true |
| PM1548-SK1P3 | FBXL21 | n.956C>T | non_coding_exon_variant | MODIFIER | 0.5 | 0 | 10 | 13 | . | . | . | . |
| PM1548-SK1P3 | KDM3B | p.Arg921Trp | missense_variant | MODERATE | 0 | 0 | 45 | 178 | . | . | . | . |
| PM1548-SK1P3 | PSD2 | p.Arg574His | missense_variant | MODERATE | 0.412088 | 0.0147059 | 182 | 136 | . | . | . | . |
| PM1548-SK1P3 | PCDHA3 | p.Glu558Val | missense_variant | MODERATE | 0.291262 | 0 | 103 | 71 | . | . | . | . |
| PM1548-SK1P3 | PCDHA5 | p.Ala56Thr | missense_variant | MODERATE | 0.283422 | 0 | 187 | 32 | . | . | . | . |
| PM1548-SK1P3 | PCDHA5 | p.Arg64Arg | synonymous_variant | LOW | 0.291262 | 0 | 206 | 38 | . | . | . | . |
| PM1548-SK1P3 | PCDHA5 | p.Gly69Asp | missense_variant | MODERATE | 0.2723 | 0 | 213 | 47 | . | . | . | . |
| PM1548-SK1P3 | PCDHA8 | p.Ala57Thr | missense_variant | MODERATE | 0.315068 | 0 | 146 | 46 | . | . | . | . |
| PM1548-SK1P3 | PCDHA13 | p.Glu558Val | missense_variant | MODERATE | 0.511628 | 0 | 43 | 105 | . | . | . | . |
| PM1548-SK1P3 | PCDHAC2 | p.Arg543Trp | missense_variant | MODERATE | 0.448276 | 0 | 29 | 153 | . | . | . | . |
| PM1548-SK1P3 | PCDHAC2 | c.2565+37G>A | intron_variant | MODIFIER | 0.386139 | 0 | 101 | 61 | . | . | . | . |
| PM1548-SK1P3 | PCDHGA1 | p.Ser370Gly | missense_variant | MODERATE | 0.555556 | 0 | 45 | 385 | . | . | . | . |
| PM1548-SK1P3 | PCDHGA9 | p.Val422Ala | missense_variant | MODERATE | 0 | 0 | 5 | 94 | . | . | . | . |
| PM1548-SK1P3 | PCDH1 | p.Ala203Thr | missense_variant | MODERATE | 0.45 | 0 | 320 | 253 | . | . | . | . |
| PM1548-SK1P3 | PCDH12 | p.Gly10Glu | missense_variant | MODERATE | 0.168831 | 0 | 154 | 44 | . | . | . | . |
| PM1548-SK1P3 | CSF1R | p.Glu955Lys | missense_variant | MODERATE | 0.607143 | 0 | 28 | 65 | . | . | . | . |
| PM1548-SK1P3 | TIMD4 | p.Ala333Val | missense_variant | MODERATE | 0.435897 | 0 | 39 | 229 | . | . | . | . |
| PM1548-SK1P3 | ATP10B | p.Arg115Lys | missense_variant | MODERATE | 0.375963 | 0 | 649 | 155 | . | . | . | . |
| PM1548-SK1P3 | UBTD2 | p.Thr160Ile | missense_variant | MODERATE | 0.285714 | 0 | 7 | 33 | . | . | . | . |
| PM1548-SK1P3 | GPRIN1 | p.Pro806fs | frameshift_variant | HIGH | 0.534483 | 0 | 58 | 165 | . | . | . | . |
| PM1548-SK1P3 | GPRIN1 | p.Pro805Thr | missense_variant | MODERATE | 0.525424 | 0 | 59 | 165 | . | . | . | . |
| PM1548-SK1P3 | GPRIN1 | p.Ser803fs | frameshift_variant | HIGH | 0.525424 | 0 | 59 | 164 | . | . | . | . |
| PM1548-SK1P3 | GPRIN1 | p.Glu801Asp | missense_variant | MODERATE | 0.526316 | 0 | 57 | 165 | . | . | . | . |
| PM1548-SK1P3 | UIMC1 | p.Phe165Cys | missense_variant | MODERATE | 0.337662 | 0 | 77 | 220 | . | . | . | . |
| PM1548-SK1P3 | UIMC1 | p.Ile156Val | missense_variant&splice_region_variant | MODERATE | 0.285714 | 0 | 91 | 259 | . | . | . | . |
| PM1548-SK1P3 | FGFR4 | p.Pro568fs | frameshift_variant | HIGH | 0.517241 | 0 | 29 | 174 | true | . | . | . |
| PM1548-SK1P3 | FAM193B | p.Pro761Ser | missense_variant | MODERATE | 0.103448 | 0 | 58 | 84 | . | . | . | . |
| PM1548-SK1P3 | GRM6 | p.Ala93Thr | missense_variant | MODERATE | 0.454545 | 0 | 110 | 91 | . | . | . | . |
| PM1548-SK1P3 | RREB1 | p.Val754Met | missense_variant | MODERATE | 0.333333 | 0 | 135 | 159 | . | . | . | . |
| PM1548-SK1P3 | HIST1H4C | p.Val82Ala | missense_variant | MODERATE | 0.428571 | 0 | 7 | 187 | . | . | . | . |
| PM1548-SK1P3 | HIST1H2AM | p.Leu52Met | missense_variant | MODERATE | 0.319149 | 0 | 47 | 43 | . | . | . | . |
| PM1548-SK1P3 | TRIM27 | p.Ala24Val | missense_variant | MODERATE | 0.213592 | 0 | 103 | 147 | true | . | . | . |
| PM1548-SK1P3 | OR2H1 | p.Ile295Leu | missense_variant | MODERATE | 0.705882 | 0 | 17 | 181 | . | . | . | . |
| PM1548-SK1P3 | HLA-F | p.Leu396fs | frameshift_variant | HIGH | 0.625 | 0 | 8 | 38 | . | . | . | . |
| PM1548-SK1P3 | HLA-G | p.Arg285Ser | missense_variant | MODERATE | 0.366667 | 0 | 30 | 42 | . | . | . | . |
| PM1548-SK1P3 | TRIM26 | p.Glu241Lys | missense_variant | MODERATE | 0.580645 | 0 | 31 | 200 | . | . | . | . |
| PM1548-SK1P3 | MDC1 | p.Gln1431Arg | missense_variant | MODERATE | 0 | 0 | 103 | 163 | . | . | . | . |
| PM1548-SK1P3 | MUC22 | p.Ile1250Thr | missense_variant | MODERATE | 0.15625 | 0 | 96 | 195 | . | . | . | . |
| PM1548-SK1P3 | HLA-C | p.Arg243Trp | missense_variant | MODERATE | 0.541667 | 0 | 24 | 40 | . | . | . | . |
| PM1548-SK1P3 | HLA-B | p.Arg243Trp | missense_variant | MODERATE | 0.0635452 | 0.00879765 | 299 | 341 | . | . | true | . |
| PM1548-SK1P3 | VWA7 | p.Thr26fs | frameshift_variant | HIGH | 0.110236 | 0 | 254 | 239 | . | . | . | . |
| PM1548-SK1P3 | CFB | p.Asp1161fs | frameshift_variant | HIGH | 0.414286 | 0 | 140 | 133 | . | . | . | . |
| PM1548-SK1P3 | RGL2 | p.Glu372fs | frameshift_variant | HIGH | 0.0815217 | 0 | 368 | 192 | . | . | . | . |
| PM1548-SK1P3 | DNAH8 | p.Tyr1705Cys | missense_variant | MODERATE | 0 | 0 | 64 | 68 | . | . | . | . |
| PM1548-SK1P3 | KLHDC3 | p.Arg67Cys | missense_variant | MODERATE | 0.307692 | 0 | 13 | 263 | . | . | . | . |
| PM1548-SK1P3 | PKHD1 | p.Arg880His | missense_variant | MODERATE | 0.517241 | 0 | 58 | 253 | . | . | . | . |
| PM1548-SK1P3 | DST | p.Thr1051fs | frameshift_variant | HIGH | 0.3 | 0 | 10 | 228 | . | . | . | . |
| PM1548-SK1P3 | KIAA1586 | p.Arg617Leu | missense_variant | MODERATE | 0 | 0 | 91 | 71 | . | . | . | . |
| PM1548-SK1P3 | ZNF451 | p.Phe193Leu | missense_variant&splice_region_variant | MODERATE | 0.5 | 0 | 6 | 32 | . | . | . | . |
| PM1548-SK1P3 | COL12A1 | p.Pro2509Thr | missense_variant | MODERATE | 0 | 0 | 105 | 157 | . | . | . | . |
| PM1548-SK1P3 | SENP6 | p.Ser627Gly | missense_variant | MODERATE | 0 | 0 | 139 | 66 | . | . | . | . |
| PM1548-SK1P3 | ME1 | c.78+1G>A | splice_donor_variant&intron_variant | HIGH | 0.480769 | 0 | 312 | 306 | . | . | . | . |
| PM1548-SK1P3 | PRSS35 | p.Arg265* | stop_gained | HIGH | 0.666667 | 0 | 3 | 41 | . | . | . | . |
| PM1548-SK1P3 | ANKRD6 | p.Ala390Val | missense_variant | MODERATE | 0.857143 | 0 | 7 | 31 | . | . | . | . |
| PM1548-SK1P3 | MDN1 | p.Thr2167Lys | missense_variant | MODERATE | 0 | 0 | 9 | 72 | . | . | . | . |
| PM1548-SK1P3 | MMS22L | p.Gly37Glu | missense_variant | MODERATE | 0.55 | 0 | 20 | 240 | . | . | . | . |
| PM1548-SK1P3 | ATG5 | p.Asp149fs | frameshift_variant | HIGH | 0 | 0 | 277 | 95 | . | . | . | . |
| PM1548-SK1P3 | RFPL4B | p.Arg203Cys | missense_variant | MODERATE | 0.647059 | 0 | 17 | 78 | . | . | . | . |
| PM1548-SK1P3 | KPNA5 | p.Phe383Ser | missense_variant | MODERATE | 0.133333 | 0 | 15 | 67 | . | . | . | . |
| PM1548-SK1P3 | FAM184A | p.Glu79Asp | missense_variant | MODERATE | 0.364486 | 0 | 107 | 52 | . | . | . | . |
| PM1548-SK1P3 | TBC1D32 | p.Ile27Asn | missense_variant | MODERATE | 0.52381 | 0 | 21 | 42 | . | . | . | . |
| PM1548-SK1P3 | HSF2 | p.Arg196His | missense_variant | MODERATE | 0.111111 | 0 | 27 | 76 | . | . | . | . |
| PM1548-SK1P3 | KIAA0408 | p.Phe313Leu | missense_variant | MODERATE | 0.0227273 | 0 | 44 | 109 | . | . | . | . |
| PM1548-SK1P3 | SOGA3 | p.Ala214fs | frameshift_variant | HIGH | 0.5 | 0 | 42 | 39 | . | . | . | . |
| PM1548-SK1P3 | ENPP1 | c.240+2T>C | splice_donor_variant&intron_variant | HIGH | 0.72 | 0 | 50 | 63 | . | . | . | . |
| PM1548-SK1P3 | CTGF | p.Ser227Phe | missense_variant | MODERATE | 0.339623 | 0 | 53 | 36 | . | . | . | . |
| PM1548-SK1P3 | CTGF | p.Ala226Thr | missense_variant | MODERATE | 0.333333 | 0 | 54 | 46 | . | . | . | . |
| PM1548-SK1P3 | CTGF | p.Ile195Met | missense_variant | MODERATE | 0.5625 | 0 | 32 | 60 | . | . | . | . |
| PM1548-SK1P3 | ADGB | p.Asp890Val | missense_variant | MODERATE | 0 | 0 | 17 | 47 | . | . | . | . |
| PM1548-SK1P3 | SYNE1 | p.Pro1307Ser | missense_variant | MODERATE | 0.9 | 0 | 50 | 79 | . | . | . | . |
| PM1548-SK1P3 | ARID1B | p.Pro1578Ser | missense_variant | MODERATE | 0.191781 | 0 | 73 | 46 | true | . | . | . |
| PM1548-SK1P3 | ARID1B | p.Ser1617Thr | missense_variant | MODERATE | 0.116279 | 0 | 129 | 59 | true | . | . | . |
| PM1548-SK1P3 | TULP4 | p.Pro1275Thr | missense_variant | MODERATE | 0.35 | 0 | 40 | 171 | . | . | . | . |
| PM1548-SK1P3 | TULP4 | p.Asp1298Asn | missense_variant | MODERATE | 0.28125 | 0.00787402 | 32 | 254 | . | . | . | . |
| PM1548-SK1P3 | TULP4 | p.Asp1298Glu | missense_variant | MODERATE | 0.40625 | 0 | 32 | 255 | . | . | . | . |
| PM1548-SK1P3 | SMOC2 | p.Ala54Thr | missense_variant | MODERATE | 0.795181 | 0 | 83 | 36 | . | . | . | . |
| PM1548-SK1P3 | GPR146 | p.Pro248Gln | missense_variant | MODERATE | 0 | 0 | 35 | 46 | . | . | . | . |
| PM1548-SK1P3 | ELFN1 | p.Arg389Cys | missense_variant | MODERATE | 0.484444 | 0 | 225 | 43 | . | . | . | . |
| PM1548-SK1P3 | SDK1 | p.Pro1727fs | frameshift_variant | HIGH | 0.493976 | 0 | 249 | 269 | . | . | . | . |
| PM1548-SK1P3 | PMS2 | p.Thr277Ala | missense_variant | MODERATE | 0.142857 | 0 | 7 | 38 | true | . | . | . |
| PM1548-SK1P3 | MIOS | p.Leu213His | missense_variant | MODERATE | 0.666667 | 0 | 12 | 44 | . | . | . | . |
| PM1548-SK1P3 | ETV1 | p.Asn37Thr | missense_variant | MODERATE | 0 | 0 | 164 | 72 | true | . | . | . |
| PM1548-SK1P3 | MPP6 | p.Lys306fs | frameshift_variant | HIGH | 0 | 0 | 471 | 33 | . | . | . | . |
| PM1548-SK1P3 | HERPUD2 | p.Val266Ala | missense_variant | MODERATE | 0.333333 | 0 | 165 | 59 | . | . | . | . |
| PM1548-SK1P3 | POU6F2 | p.Glu562Ala | missense_variant | MODERATE | 0.317152 | 0 | 309 | 225 | . | . | . | . |
| PM1548-SK1P3 | MYO1G | p.Ala779Val | missense_variant | MODERATE | 0.129707 | 0 | 1195 | 358 | . | . | . | . |
| PM1548-SK1P3 | VWC2 | p.Ser3Arg | missense_variant | MODERATE | 0.103448 | 0 | 29 | 64 | . | . | . | . |
| PM1548-SK1P3 | GRB10 | p.Ala349Thr | missense_variant | MODERATE | 0.594595 | 0 | 37 | 372 | . | . | . | . |
| PM1548-SK1P3 | WBSCR22 | p.Pro8Gln | missense_variant | MODERATE | 0 | 0 | 6 | 35 | . | . | . | . |
| PM1548-SK1P3 | CLIP2 | p.Arg771Gln | missense_variant | MODERATE | 0.258065 | 0 | 248 | 150 | . | . | . | . |
| PM1548-SK1P3 | TMEM60 | p.Ala78fs | frameshift_variant | HIGH | 0.642857 | 0 | 14 | 31 | . | . | . | . |
| PM1548-SK1P3 | GNAI1 | p.Cys214Phe | missense_variant | MODERATE | 0 | 0 | 57 | 196 | . | . | . | . |
| PM1548-SK1P3 | HGF | p.Val529Ala | missense_variant | MODERATE | 0 | 0 | 341 | 125 | . | true | . | . |
| PM1548-SK1P3 | ABCB1 | p.Tyr1165* | stop_gained | HIGH | 0.4375 | 0 | 32 | 32 | . | . | . | . |
| PM1548-SK1P3 | ABCB1 | c.3490-1G>C | splice_acceptor_variant&intron_variant | HIGH | 0.4375 | 0 | 32 | 32 | . | . | . | . |
| PM1548-SK1P3 | ABCB1 | p.Ser196Ala | missense_variant | MODERATE | 0.229167 | 0 | 48 | 56 | . | . | . | . |
| PM1548-SK1P3 | MTERF | p.Ala221Thr | missense_variant | MODERATE | 0.5 | 0 | 12 | 212 | . | . | . | . |
| PM1548-SK1P3 | AKAP9 | p.Ile1377Thr | missense_variant | MODERATE | 1 | 0 | 1 | 76 | true | . | . | . |
| PM1548-SK1P3 | LMTK2 | p.Asp236fs | frameshift_variant | HIGH | 0.428571 | 0 | 21 | 300 | . | . | . | . |
| PM1548-SK1P3 | LMTK2 | p.Leu1130Ser | missense_variant | MODERATE | 0.46875 | 0 | 64 | 72 | . | . | . | . |
| PM1548-SK1P3 | TRRAP | p.Asn1675Thr | missense_variant | MODERATE | 0.25 | 0 | 120 | 117 | true | . | . | . |
| PM1548-SK1P3 | TRRAP | p.Arg1978Trp | missense_variant | MODERATE | 0.53125 | 0 | 32 | 221 | true | . | . | . |
| PM1548-SK1P3 | TRRAP | p.Pro2508Ala | missense_variant | MODERATE | 0.477612 | 0 | 67 | 384 | true | . | . | . |
| PM1548-SK1P3 | GIGYF1 | p.Leu580Pro | missense_variant | MODERATE | 0.105263 | 0 | 456 | 69 | . | . | . | . |
| PM1548-SK1P3 | SLC12A9 | p.Leu559fs | frameshift_variant | HIGH | 0.2 | 0 | 10 | 56 | . | . | . | . |
| PM1548-SK1P3 | MOGAT3 | p.Asn82Lys | missense_variant | MODERATE | 0 | 0 | 89 | 150 | . | . | . | . |
| PM1548-SK1P3 | SH2B2 | p.Ala74Ala | synonymous_variant | LOW | 0.481481 | 0 | 27 | 70 | . | . | . | . |
| PM1548-SK1P3 | LRWD1 | p.Leu87Phe | missense_variant | MODERATE | 0 | 0 | 20 | 78 | . | . | . | . |
| PM1548-SK1P3 | FBXL13 | p.Cys375Ser | missense_variant | MODERATE | 0 | 0 | 17 | 94 | . | . | . | . |
| PM1548-SK1P3 | RELN | p.Arg2363His | missense_variant | MODERATE | 0.458498 | 0 | 253 | 39 | . | . | . | . |
| PM1548-SK1P3 | ATXN7L1 | p.Val629Phe | missense_variant | MODERATE | 0 | 0 | 6 | 193 | . | . | . | . |
| PM1548-SK1P3 | PIK3CG | p.Thr607fs | frameshift_variant | HIGH | 0 | 0 | 52 | 44 | . | true | . | . |
| PM1548-SK1P3 | CBLL1 | p.Val277Ala | missense_variant | MODERATE | 0 | 0 | 101 | 200 | . | . | . | . |
| PM1548-SK1P3 | SLC26A3 | p.Thr479Ala | missense_variant | MODERATE | 0.341463 | 0 | 82 | 194 | . | . | true | . |
| PM1548-SK1P3 | FOXP2 | p.Gly98Val | missense_variant | MODERATE | 0 | 0 | 49 | 125 | . | . | . | . |
| PM1548-SK1P3 | PTPRZ1 | p.Pro751Leu | missense_variant | MODERATE | 0.529412 | 0 | 17 | 64 | . | . | . | . |
| PM1548-SK1P3 | SND1 | p.Ala331Pro | missense_variant | MODERATE | 0.107623 | 0 | 446 | 236 | true | . | . | . |
| PM1548-SK1P3 | CEP41 | p.Asp152Gly | missense_variant | MODERATE | 0.0535714 | 0 | 112 | 235 | . | . | . | . |
| PM1548-SK1P3 | TSGA13 | p.Lys151fs | frameshift_variant | HIGH | 0 | 0 | 5 | 195 | . | . | . | . |
| PM1548-SK1P3 | TTC26 | p.Ala221Thr | missense_variant | MODERATE | 0.4375 | 0 | 32 | 82 | . | . | . | . |
| PM1548-SK1P3 | MKRN1 | c.186-2435G>C | intron_variant | MODIFIER | 0.48 | 0 | 25 | 136 | . | . | . | . |

| | | | | | | | | | | | | |
|---|---|---|---|---|---|---|---|---|---|---|---|---|
| PM1548-SK1P3 | MGAM | p.Asn1603Asp | missense_variant | MODERATE | 0.483221 | 0 | 149 | 85 | . | . | . | . |
| PM1548-SK1P3 | EPHB6 | p.Ala253fs | frameshift_variant | HIGH | 0.272727 | 0 | 22 | 107 | . | true | . | . |
| PM1548-SK1P3 | ZNF775 | p.Gly65fs | frameshift_variant | HIGH | 0 | 0 | 14 | 50 | . | . | . | . |
| PM1548-SK1P3 | FASTK | p.Ile345Asn | missense_variant | MODERATE | 0.308824 | 0 | 68 | 255 | . | . | . | . |
| PM1548-SK1P3 | GALNT11 | p.Leu188Pro | missense_variant | MODERATE | 0.333333 | 0 | 6 | 157 | . | . | . | . |
| PM1548-SK1P3 | TNKS | p.His859Asn | missense_variant | MODERATE | 0 | 0 | 15 | 75 | . | . | . | . |
| PM1548-SK1P3 | RP1L1 | p.Gln1044fs | frameshift_variant | HIGH | 1 | 0 | 160 | 101 | . | . | . | . |
| PM1548-SK1P3 | XKR6 | p.Arg80Lys | missense_variant | MODERATE | 0.416667 | 0 | 24 | 127 | . | . | . | . |
| PM1548-SK1P3 | PCM1 | p.Met1321Val | missense_variant | MODERATE | 0.25 | 0 | 8 | 138 | true | . | . | . |
| PM1548-SK1P3 | SORBS3 | p.Thr100Ala | missense_variant | MODERATE | 0.258621 | 0 | 58 | 143 | . | . | . | . |
| PM1548-SK1P3 | LOXL2 | p.Pro40Leu | missense_variant | MODERATE | 0.974359 | 0 | 39 | 224 | . | . | . | . |
| PM1548-SK1P3 | NEFM | p.Leu5Leu | synonymous_variant | LOW | 0.290323 | 0 | 31 | 41 | . | . | . | . |
| PM1548-SK1P3 | NRG1 | p.Gly39Glu | missense_variant | MODERATE | 0.181818 | 0 | 11 | 55 | true | . | . | . |
| PM1548-SK1P3 | NRG1 | p.Pro44Ser | missense_variant | MODERATE | 0.166667 | 0 | 12 | 58 | true | . | . | . |
| PM1548-SK1P3 | NRG1 | p.Ala74Pro | missense_variant | MODERATE | 0.75 | 0 | 8 | 35 | true | . | . | . |
| PM1548-SK1P3 | NRG1 | c.1488-469C>G | intron_variant | MODIFIER | 0.136364 | 0 | 44 | 106 | true | . | . | . |
| PM1548-SK1P3 | ADAM2 | p.Arg3His | missense_variant | MODERATE | 0 | 0 | 92 | 64 | . | . | . | . |
| PM1548-SK1P3 | DKK4 | p.Asp195Asn | missense_variant | MODERATE | 0.532258 | 0 | 124 | 81 | . | . | . | . |
| PM1548-SK1P3 | MCM4 | p.Thr629fs | frameshift_variant | HIGH | 0.321429 | 0 | 56 | 85 | . | . | . | . |
| PM1548-SK1P3 | PXDNL | p.Ala353Thr | missense_variant | MODERATE | 0.392857 | 0 | 28 | 175 | . | . | . | . |
| PM1548-SK1P3 | RP1 | p.Tyr834fs | frameshift_variant | HIGH | 0.333333 | 0 | 21 | 39 | . | . | . | . |
| PM1548-SK1P3 | YTHDF3 | p.Gln323Gln | synonymous_variant | LOW | 0.302752 | 0 | 109 | 588 | . | . | . | . |
| PM1548-SK1P3 | C8orf46 | p.Thr78Ile | missense_variant | MODERATE | 0 | 0 | 24 | 109 | . | . | . | . |
| PM1548-SK1P3 | ZFHX4 | p.Ala2905Val | missense_variant | MODERATE | 0.285714 | 0 | 21 | 42 | . | . | . | . |
| PM1548-SK1P3 | HEY1 | p.Val187Ala | missense_variant | MODERATE | 0.120455 | 0 | 440 | 281 | true | . | . | . |
| PM1548-SK1P3 | HEY1 | p.Thr186Ser | missense_variant | MODERATE | 0.120729 | 0 | 439 | 272 | true | . | . | . |
| PM1548-SK1P3 | VPS13B | p.Asn2064Thr | missense_variant | MODERATE | 0 | 0 | 16 | 75 | . | . | . | . |
| PM1548-SK1P3 | TRPS1 | p.Arg297Cys | missense_variant | MODERATE | 0.427536 | 0 | 138 | 259 | . | . | . | . |
| PM1548-SK1P3 | TRPS1 | p.Asn265Asn | synonymous_variant | LOW | 0.192308 | 0 | 26 | 334 | . | . | . | . |
| PM1548-SK1P3 | EXT1 | p.Val66Phe | missense_variant | MODERATE | 0.142857 | 0 | 70 | 87 | true | . | . | . |
| PM1548-SK1P3 | TBC1D31 | p.Ala208Thr | missense_variant | MODERATE | 0.447368 | 0 | 228 | 184 | . | . | . | . |
| PM1548-SK1P3 | ATAD2 | p.Glu119fs | frameshift_variant | HIGH | 0.818182 | 0 | 11 | 106 | . | . | . | . |
| PM1548-SK1P3 | ZNF572 | p.Ser55Pro | missense_variant | MODERATE | 0.51049 | 0 | 143 | 39 | . | . | . | . |
| PM1548-SK1P3 | MYC | p.Gly175Ser | missense_variant | MODERATE | 0.103448 | 0 | 58 | 269 | true | . | . | . |
| PM1548-SK1P3 | ASAP1 | p.Glu919Lys | missense_variant | MODERATE | 0.242105 | 0 | 95 | 187 | . | . | . | . |
| PM1548-SK1P3 | WISP1 | p.Ile175Met | missense_variant | MODERATE | 0.362069 | 0 | 58 | 119 | . | . | . | . |
| PM1548-SK1P3 | KHDRBS3 | p.Tyr316Tyr | splice_region_variant&synonymous_variant | LOW | 0.4 | 0 | 80 | 229 | . | . | . | . |
| PM1548-SK1P3 | COL22A1 | p.Thr125Met | missense_variant | MODERATE | 0 | 0 | 3 | 59 | . | . | . | . |
| PM1548-SK1P3 | LY6D | p.Ala99Thr | missense_variant | MODERATE | 0.0766773 | 0.00956938 | 626 | 209 | . | . | . | . |
| PM1548-SK1P3 | TOP1MT | p.Pro193Leu | missense_variant | MODERATE | 0 | 0.00595238 | 24 | 168 | . | . | . | . |
| PM1548-SK1P3 | ZC3H3 | p.Gly101fs | frameshift_variant | HIGH | 0.08 | 0 | 25 | 100 | . | . | . | . |
| PM1548-SK1P3 | GSDMD | p.Glu451Ala | missense_variant | MODERATE | 0.401198 | 0 | 167 | 215 | . | . | . | . |
| PM1548-SK1P3 | MAPK15 | p.Pro493Leu | missense_variant | MODERATE | 0.443636 | 0.00271739 | 275 | 368 | . | . | . | . |
| PM1548-SK1P3 | SCRIB | p.Cys22Trp | missense_variant | MODERATE | 0.1 | 0 | 330 | 81 | . | . | . | . |
| PM1548-SK1P3 | PLEC | p.Arg3540Gln | missense_variant | MODERATE | 0 | 0 | 73 | 130 | . | . | . | . |
| PM1548-SK1P3 | PLEC | p.Val648Met | missense_variant | MODERATE | 0 | 0 | 49 | 166 | . | . | . | . |
| PM1548-SK1P3 | CYHR1 | n.145689659dup | intragenic_variant | MODIFIER | 0.1 | 0 | 50 | 56 | . | . | . | . |
| PM1548-SK1P3 | ZNF7 | p.Val136Ile | missense_variant | MODERATE | 0 | 0 | 12 | 154 | . | . | . | . |
| PM1548-SK1P3 | KANK1 | p.Thr192Ser | missense_variant | MODERATE | 0.285714 | 0 | 49 | 65 | . | . | . | . |
| PM1548-SK1P3 | DMRT3 | p.Leu385Leu | synonymous_variant | LOW | 0.212389 | 0 | 113 | 221 | . | . | . | . |
| PM1548-SK1P3 | RLN2 | p.His8fs | frameshift_variant | HIGH | 0.284916 | 0 | 179 | 225 | . | . | . | . |
| PM1548-SK1P3 | IFNA4 | p.Ala74Thr | missense_variant | MODERATE | 0.391304 | 0 | 23 | 51 | . | . | . | . |
| PM1548-SK1P3 | BAG1 | p.Glu321Lys | missense_variant | MODERATE | 0.518519 | 0 | 189 | 192 | . | . | . | . |
| PM1548-SK1P3 | PRSS3 | p.Ser102Tyr | missense_variant | MODERATE | 0.0625 | 0 | 32 | 102 | . | . | . | . |
| PM1548-SK1P3 | CD72 | p.Ala111Thr | missense_variant | MODERATE | 0.53271 | 0 | 107 | 227 | . | . | . | . |
| PM1548-SK1P3 | RGP1 | p.Arg328Trp | missense_variant | MODERATE | 0.425 | 0 | 40 | 168 | . | . | . | . |
| PM1548-SK1P3 | ANKRD18A | p.Gly900Ala | missense_variant | MODERATE | 0.469697 | 0 | 66 | 119 | . | . | . | . |
| PM1548-SK1P3 | ANKRD18A | p.Ala897Val | missense_variant | MODERATE | 0.533981 | 0 | 103 | 119 | . | . | . | . |
| PM1548-SK1P3 | CNTNAP3 | p.Arg841Ser | missense_variant | MODERATE | 0 | 0 | 4 | 45 | . | . | . | . |
| PM1548-SK1P3 | TMC1 | p.Ala755Pro | missense_variant&splice_region_variant | MODERATE | 0.294118 | 0 | 17 | 45 | . | . | . | . |
| PM1548-SK1P3 | RMI1 | p.Thr325Ala | missense_variant | MODERATE | 0.435443 | 0 | 395 | 482 | . | . | . | . |
| PM1548-SK1P3 | NAA35 | p.*726* | stop_retained_variant | LOW | 0 | 0 | 27 | 360 | . | . | . | . |
| PM1548-SK1P3 | SPATA31E1 | p.Arg436His | missense_variant | MODERATE | 0.611111 | 0 | 18 | 212 | . | . | . | . |
| PM1548-SK1P3 | ROR2 | p.Ala307Thr | missense_variant | MODERATE | 0 | 0 | 60 | 69 | . | . | . | . |
| PM1548-SK1P3 | NOL8 | p.Arg430Lys | missense_variant | MODERATE | 0.072 | 0 | 125 | 112 | . | . | . | . |
| PM1548-SK1P3 | NOL8 | p.Leu427Gln | missense_variant | MODERATE | 0.0789474 | 0 | 114 | 113 | . | . | . | . |
| PM1548-SK1P3 | IPPK | p.Phe34Ser | missense_variant | MODERATE | 0.416667 | 0 | 12 | 158 | . | . | . | . |
| PM1548-SK1P3 | SUSD3 | c.89-6671G>T | intron_variant | MODIFIER | 0 | 0 | 3 | 26 | . | . | . | . |
| PM1548-SK1P3 | WNK2 | p.His758fs | frameshift_variant | HIGH | 0.478992 | 0 | 119 | 277 | . | . | . | . |
| PM1548-SK1P3 | C9orf3 | p.Ala207Thr | missense_variant | MODERATE | 0.389908 | 0 | 218 | 105 | . | . | . | . |
| PM1548-SK1P3 | PTCH1 | p.Thr1064Met | missense_variant | MODERATE | 0.195652 | 0 | 46 | 167 | true | . | . | . |
| PM1548-SK1P3 | ZNF510 | p.Arg592Gly | missense_variant | MODERATE | 0 | 0 | 58 | 140 | . | . | . | . |
| PM1548-SK1P3 | CCDC180 | p.Met439Ile | missense_variant | MODERATE | 0.401786 | 0 | 112 | 95 | . | . | . | . |
| PM1548-SK1P3 | GABBR2 | p.Arg622* | stop_gained | HIGH | 0.3 | 0 | 10 | 80 | . | . | . | . |
| PM1548-SK1P3 | ANKS6 | p.Ala872Val | missense_variant | MODERATE | 0.368421 | 0 | 95 | 240 | . | . | . | . |
| PM1548-SK1P3 | SLC44A1 | p.Arg437His | missense_variant | MODERATE | 0.407407 | 0 | 27 | 72 | . | . | . | . |
| PM1548-SK1P3 | SVEP1 | p.Ile1787Thr | missense_variant | MODERATE | 0.538462 | 0 | 13 | 79 | . | . | . | . |
| PM1548-SK1P3 | TNC | p.Lys1068Asn | missense_variant | MODERATE | 0.308511 | 0 | 94 | 37 | . | . | . | . |
| PM1548-SK1P3 | BRINP1 | p.Arg106His | missense_variant | MODERATE | 0.32 | 0 | 25 | 219 | . | . | . | . |
| PM1548-SK1P3 | FBXW2 | p.Arg385His | missense_variant | MODERATE | 0.266667 | 0 | 60 | 162 | . | . | . | . |
| PM1548-SK1P3 | FBXW2 | p.Ile377Val | missense_variant | MODERATE | 0.266667 | 0 | 60 | 161 | . | . | . | . |
| PM1548-SK1P3 | DAB2IP | p.His1025fs | frameshift_variant | HIGH | 0.571429 | 0 | 84 | 32 | . | . | . | . |
| PM1548-SK1P3 | LRSAM1 | p.Val385Ile | missense_variant | MODERATE | 0.368421 | 0 | 19 | 115 | . | . | . | . |
| PM1548-SK1P3 | SH2D3C | p.Arg821His | missense_variant | MODERATE | 0.553488 | 0 | 430 | 143 | . | . | . | . |
| PM1548-SK1P3 | C9orf114 | p.Val77Ile | missense_variant | MODERATE | 0.469751 | 0 | 281 | 277 | . | . | . | . |
| PM1548-SK1P3 | C9orf50 | p.Ala252Val | missense_variant | MODERATE | 0.494737 | 0 | 380 | 411 | . | . | . | . |
| PM1548-SK1P3 | NUP214 | p.Arg887His | missense_variant | MODERATE | 0.352381 | 0 | 105 | 67 | true | . | . | . |
| PM1548-SK1P3 | ADAMTS13 | p.Arg7fs | frameshift_variant | HIGH | 0.708475 | 0 | 295 | 247 | . | . | . | . |
| PM1548-SK1P3 | SARDH | p.Ala819Val | missense_variant | MODERATE | 0.5 | 0 | 6 | 41 | . | . | . | . |
| PM1548-SK1P3 | SARDH | p.Arg280Trp | missense_variant | MODERATE | 0 | 0 | 9 | 43 | . | . | . | . |
| PM1548-SK1P3 | LHX3 | p.Ala180Thr | missense_variant | MODERATE | 0.395349 | 0 | 129 | 204 | . | . | . | . |
| PM1548-SK1P3 | MAMDC4 | p.Gln963His | missense_variant | MODERATE | 0.454545 | 0 | 110 | 91 | . | . | . | . |
| PM1548-SK1P3 | AL807752.1 | p.Trp56fs | frameshift_variant | HIGH | 0.451613 | 0 | 31 | 54 | . | . | . | . |
| PM1548-SK1P3 | TPRN | p.Asn499Asp | missense_variant | MODERATE | 0.13577 | 0 | 383 | 335 | . | . | . | . |
| PM1548-SK1P3 | NSMF | p.Arg366Cys | missense_variant | MODERATE | 0.352941 | 0 | 119 | 90 | . | . | . | . |
| PM1548-SK1P3 | PNPLA7 | p.Leu178Met | missense_variant | MODERATE | 0 | 0 | 42 | 199 | . | . | . | . |
| PM1548-SK1P3 | HDHD1 | p.Arg149Cys | missense_variant | MODERATE | 1 | 0 | 6 | 104 | . | . | . | . |
| PM1548-SK1P3 | EGFL6 | p.Trp5* | stop_gained | HIGH | 0 | 0 | 37 | 81 | . | . | . | . |
| PM1548-SK1P3 | YY2 | p.Glu5Val | missense_variant | MODERATE | 0.804444 | 0 | 225 | 51 | . | . | . | . |
| PM1548-SK1P3 | TMEM47 | p.Cys104Cys | synonymous_variant | LOW | 0.487805 | 0 | 41 | 132 | . | . | . | . |
| PM1548-SK1P3 | USP27X | p.Arg155His | missense_variant | MODERATE | 0 | 0 | 38 | 112 | . | . | . | . |
| PM1548-SK1P3 | KDM5C | p.Glu1003Asp | missense_variant | MODERATE | 0.138889 | 0 | 36 | 54 | true | true | true | . |
| PM1548-SK1P3 | IQSEC2 | p.Arg612Cys | missense_variant | MODERATE | 0 | 0 | 155 | 42 | . | . | . | . |
| PM1548-SK1P3 | FAM120C | p.Gln36_Gln37d | disruptive_inframe_deletion | MODERATE | 0.95 | 0 | 20 | 126 | . | . | . | . |
| PM1548-SK1P3 | MAGED2 | p.Glu53Asp | missense_variant | MODERATE | 0 | 0 | 47 | 102 | . | . | . | . |
| PM1548-SK1P3 | TRO | p.Ala1132Thr | missense_variant | MODERATE | 0.0350877 | 0 | 57 | 69 | . | . | . | . |
| PM1548-SK1P3 | DLG3 | p.Leu333Ile | missense_variant | MODERATE | 1 | 0 | 7 | 34 | . | . | . | . |
| PM1548-SK1P3 | ARMCX4 | p.Tyr2123fs | frameshift_variant | HIGH | 0 | 0 | 120 | 78 | . | . | . | . |
| PM1548-SK1P3 | PAK3 | p.Ala466Val | missense_variant | MODERATE | 1 | 0 | 73 | 37 | . | . | . | . |
| PM1548-SK1P3 | ZCCHC16 | p.Arg297Cys | missense_variant | MODERATE | 0.285714 | 0 | 49 | 98 | . | . | . | . |
| PM1548-SK1P3 | AVPR2 | p.Ala60Val | missense_variant | MODERATE | 0 | 0 | 7 | 59 | . | . | . | . |
| PM1548-SK1P4 | B3GALT6 | p.Glu303Lys | missense_variant | MODERATE | 0.480769 | 0 | 104 | 86 | . | . | . | . |
| PM1548-SK1P4 | SCNN1D | p.Arg344His | missense_variant | MODERATE | 0.405405 | 0 | 37 | 274 | . | . | . | . |
| PM1548-SK1P4 | ATAD3B | p.Arg290Cys | missense_variant | MODERATE | 0.504274 | 0.00687285 | 585 | 291 | . | . | . | . |
| PM1548-SK1P4 | C1orf86 | p.Pro72His | missense_variant | MODERATE | 0.454887 | 0 | 266 | 43 | . | . | . | . |
| PM1548-SK1P4 | PRDM16 | p.Arg759Gln | missense_variant | MODERATE | 0 | 0 | 23 | 40 | true | . | . | . |
| PM1548-SK1P4 | DFFB | p.Tyr290Phe | missense_variant | MODERATE | 0.333333 | 0 | 54 | 203 | . | . | . | . |
| PM1548-SK1P4 | NPHP4 | p.Arg488Gln | missense_variant | MODERATE | 0.477778 | 0 | 450 | 194 | . | . | . | . |
| PM1548-SK1P4 | PLEKHG5 | p.Thr301fs | frameshift_variant | HIGH | 0.507607 | 0 | 723 | 105 | . | . | . | . |
| PM1548-SK1P4 | ZBTB48 | p.Val626Ala | missense_variant | MODERATE | 0.25 | 0 | 488 | 142 | . | . | . | . |
| PM1548-SK1P4 | KLHL21 | p.Cys250fs | frameshift_variant | HIGH | 0.375691 | 0 | 181 | 136 | . | . | . | . |
| PM1548-SK1P4 | CAMTA1 | p.Ser510Thr | missense_variant | MODERATE | 0.00793651 | 0 | 126 | 185 | true | . | . | . |
| PM1548-SK1P4 | PER3 | p.Ala803Ser | missense_variant | MODERATE | 0 | 0 | 61 | 110 | . | . | . | . |
| PM1548-SK1P4 | SLC45A1 | p.Thr364Met | missense_variant | MODERATE | 0.4 | 0 | 10 | 56 | . | . | . | . |
| PM1548-SK1P4 | SLC45A1 | p.Tyr689* | stop_gained | HIGH | 0.45 | 0 | 60 | 120 | . | . | . | . |
| PM1548-SK1P4 | RERE | p.Lys83fs | frameshift_variant | HIGH | 0 | 0 | 4 | 49 | . | . | . | . |
| PM1548-SK1P4 | CORT | p.Gln133fs | frameshift_variant | HIGH | 0 | 0 | 45 | 48 | . | . | . | . |

| | | | | | | | | | | | | |
|---|---|---|---|---|---|---|---|---|---|---|---|---|
| PM1548-SK1P4 | EXOSC10 | p.Met1fs | frameshift_variant&start_lost | HIGH | 0.450644 | 0 | 233 | 61 | . | . | . | . |
| PM1548-SK1P4 | FHAD1 | p.Ile953fs | frameshift_variant | HIGH | 0.492308 | 0 | 260 | 108 | . | . | . | . |
| PM1548-SK1P4 | PLEKHM2 | p.Arg848Trp | missense_variant | MODERATE | 0 | 0 | 33 | 42 | . | . | . | . |
| PM1548-SK1P4 | SPEN | p.Asn2002fs | frameshift_variant | HIGH | 0 | 0 | 512 | 173 | true | . | true | true |
| PM1548-SK1P4 | EPHA2 | p.Asn106Ser | missense_variant | MODERATE | 0.24 | 0 | 50 | 84 | . | . | true | . |
| PM1548-SK1P4 | CROCC | p.Arg1590Cys | missense_variant | MODERATE | 0.237975 | 0 | 395 | 548 | . | . | . | . |
| PM1548-SK1P4 | RCC2 | p.Gly465Arg | missense_variant | MODERATE | 0.603175 | 0 | 63 | 148 | . | . | . | . |
| PM1548-SK1P4 | ARHGEF10L | p.Val965Met | missense_variant | MODERATE | 0 | 0 | 1001 | 230 | . | . | . | . |
| PM1548-SK1P4 | KLHDC7A | p.Ala652Val | missense_variant | MODERATE | 0.64 | 0 | 25 | 60 | . | . | . | . |
| PM1548-SK1P4 | RNF186 | p.Ile185Thr | missense_variant | MODERATE | 0.00358423 | 0 | 279 | 472 | . | . | . | . |
| PM1548-SK1P4 | KIF17 | p.Arg196His | missense_variant | MODERATE | 0.301887 | 0 | 53 | 71 | . | . | . | . |
| PM1548-SK1P4 | EIF4G3 | c.30+5101C>G | intron_variant | MODIFIER | 0.222222 | 0 | 27 | 302 | . | . | . | . |
| PM1548-SK1P4 | HSPG2 | p.Gly2259Asp | missense_variant | MODERATE | 0 | 0 | 23 | 55 | . | . | . | . |
| PM1548-SK1P4 | EPHA8 | p.Ser87Asn | missense_variant | MODERATE | 0.021978 | 0 | 91 | 190 | . | . | . | . |
| PM1548-SK1P4 | AIM1L | n.26671971delA | intragenic_variant | MODIFIER | 0.047619 | 0 | 21 | 55 | . | . | . | . |
| PM1548-SK1P4 | ARID1A | p.Thr1863Ile | missense_variant | MODERATE | 0.0411523 | 0 | 486 | 201 | true | true | true | true |
| PM1548-SK1P4 | PTPRU | p.Arg639Trp | missense_variant | MODERATE | 0 | 0 | 169 | 60 | . | . | . | . |
| PM1548-SK1P4 | FABP3 | p.Thr54fs | frameshift_variant | HIGH | 0.652174 | 0 | 23 | 207 | . | . | . | . |
| PM1548-SK1P4 | LCK | p.Arg222Cys | missense_variant | MODERATE | 0.0120482 | 0 | 83 | 146 | true | . | . | . |
| PM1548-SK1P4 | KIAA1522 | p.Ala943Thr | missense_variant | MODERATE | 0.498113 | 0 | 265 | 198 | . | . | . | . |
| PM1548-SK1P4 | ZNF362 | p.Ala105Val | missense_variant | MODERATE | 0 | 0 | 26 | 295 | . | . | . | . |
| PM1548-SK1P4 | LSM10 | p.Ala2Val | missense_variant | MODERATE | 0.431193 | 0 | 109 | 400 | . | . | . | . |
| PM1548-SK1P4 | CSF3R | p.Tyr508Cys | missense_variant | MODERATE | 0.13253 | 0 | 249 | 166 | true | . | . | . |
| PM1548-SK1P4 | EPHA10 | p.Pro330Ser | missense_variant | MODERATE | 0.382576 | 0 | 264 | 71 | . | . | . | . |
| PM1548-SK1P4 | MACF1 | p.Gln6033Pro | missense_variant&splice_region_variant | MODERATE | 0 | 0 | 14 | 48 | . | . | . | . |
| PM1548-SK1P4 | PABPC4 | p.Ser206Asn | missense_variant | MODERATE | 0 | 0 | 45 | 90 | . | . | . | . |
| PM1548-SK1P4 | PABPC4 | p.Glu205Gly | missense_variant | MODERATE | 0 | 0 | 40 | 73 | . | . | . | . |
| PM1548-SK1P4 | ZMPSTE24 | p.Leu362fs | frameshift_variant | HIGH | 0.818182 | 0 | 11 | 53 | . | . | . | . |
| PM1548-SK1P4 | HIVEP3 | p.Pro1037Leu | missense_variant | MODERATE | 0.55814 | 0 | 43 | 35 | . | . | . | . |
| PM1548-SK1P4 | MED8 | p.Asn295fs | frameshift_variant | HIGH | 0.645621 | 0 | 491 | 97 | . | . | . | . |
| PM1548-SK1P4 | HYI | p.Ala55Ser | missense_variant | MODERATE | 0 | 0 | 446 | 158 | . | . | . | . |
| PM1548-SK1P4 | PTPRF | p.Arg1669Trp | missense_variant | MODERATE | 0.557951 | 0 | 371 | 112 | . | . | . | . |
| PM1548-SK1P4 | SLC6A9 | p.Arg662Gln | missense_variant | MODERATE | 0.449123 | 0 | 285 | 99 | . | . | . | . |
| PM1548-SK1P4 | PTCH2 | p.Ala967Val | missense_variant | MODERATE | 0 | 0 | 232 | 194 | . | . | . | . |
| PM1548-SK1P4 | PRDX1 | p.His10Tyr | missense_variant | MODERATE | 0.0377358 | 0 | 53 | 151 | . | . | . | . |
| PM1548-SK1P4 | MAST2 | p.Arg1086Arg | synonymous_variant | LOW | 0.0322581 | 0 | 93 | 187 | . | . | . | . |
| PM1548-SK1P4 | TAL1 | p.His86Leu | missense_variant | MODERATE | 0 | 0 | 144 | 170 | true | . | . | . |
| PM1548-SK1P4 | LRP8 | p.Asp50Gly | missense_variant | MODERATE | 0.39819 | 0 | 442 | 175 | . | . | . | . |
| PM1548-SK1P4 | CDCP2 | p.Pro209fs | frameshift_variant | HIGH | 0.48 | 0 | 25 | 107 | . | . | . | . |
| PM1548-SK1P4 | MRPL37 | p.Asp142Glu | missense_variant | MODERATE | 0.0546875 | 0 | 128 | 91 | . | . | . | . |
| PM1548-SK1P4 | CACHD1 | p.Glu211Lys | missense_variant | MODERATE | 0 | 0 | 75 | 147 | . | . | . | . |
| PM1548-SK1P4 | JAK1 | p.Val433Ile | missense_variant | MODERATE | 0.195378 | 0 | 476 | 203 | true | . | . | true |
| PM1548-SK1P4 | FPGT | p.Asp273Asn | missense_variant | MODERATE | 0 | 0 | 5 | 168 | . | . | . | . |
| PM1548-SK1P4 | LPHN2 | p.Asn759Ile | missense_variant | MODERATE | 0 | 0 | 1 | 85 | . | . | . | . |
| PM1548-SK1P4 | SYDE2 | p.Gly14fs | frameshift_variant | HIGH | 0 | 0 | 116 | 141 | . | . | . | . |
| PM1548-SK1P4 | CLCA1 | p.Arg676Trp | missense_variant | MODERATE | 1 | 0 | 5 | 91 | . | . | . | . |
| PM1548-SK1P4 | GFI1 | p.Arg348Met | missense_variant | MODERATE | 0.489642 | 0 | 531 | 270 | . | . | . | . |
| PM1548-SK1P4 | ABCA4 | p.His423Gln | missense_variant | MODERATE | 0.130435 | 0 | 46 | 257 | . | . | . | . |
| PM1548-SK1P4 | ABCA4 | p.Glu422Asp | missense_variant | MODERATE | 0.130435 | 0.00389105 | 46 | 257 | . | . | . | . |
| PM1548-SK1P4 | LPPR4 | p.Ala343Thr | missense_variant | MODERATE | 0.394737 | 0 | 38 | 96 | . | . | . | . |
| PM1548-SK1P4 | HIAT1 | p.Ala235Val | missense_variant&splice_region_variant | MODERATE | 0.472973 | 0 | 74 | 80 | . | . | . | . |
| PM1548-SK1P4 | TRMT13 | p.Glu307fs | frameshift_variant | HIGH | 0 | 0 | 47 | 85 | . | . | . | . |
| PM1548-SK1P4 | AKNAD1 | p.Leu577Met | missense_variant | MODERATE | 0.4 | 0 | 30 | 457 | . | . | . | . |
| PM1548-SK1P4 | CELSR2 | p.Asp1795Gly | missense_variant | MODERATE | 0 | 0 | 63 | 362 | . | . | . | . |
| PM1548-SK1P4 | LAMTOR5 | p.Thr39Ala | missense_variant | MODERATE | 0 | 0 | 170 | 96 | . | . | . | . |
| PM1548-SK1P4 | HIPK1 | p.Ala314Thr | missense_variant | MODERATE | 0.1 | 0 | 10 | 273 | . | . | . | . |
| PM1548-SK1P4 | TRIM33 | p.Val961Met | missense_variant | MODERATE | 0.666667 | 0 | 3 | 37 | true | . | . | . |
| PM1548-SK1P4 | PDE4DIP | p.Gln2125Lys | missense_variant | MODERATE | 0.2 | 0 | 55 | 86 | true | . | . | . |
| PM1548-SK1P4 | PDE4DIP | p.Arg759* | stop_gained | HIGH | 0.312195 | 0 | 205 | 104 | true | . | . | . |
| PM1548-SK1P4 | BCL9 | p.Pro517fs | frameshift_variant | HIGH | 0.42487 | 0 | 193 | 142 | true | . | . | . |
| PM1548-SK1P4 | TARS2 | p.Tyr371* | stop_gained | HIGH | 0.265625 | 0 | 64 | 48 | . | . | . | . |
| PM1548-SK1P4 | ADAMTSL4 | p.Gly780fs | frameshift_variant | HIGH | 0.44702 | 0 | 302 | 108 | . | . | . | . |
| PM1548-SK1P4 | ZNF687 | p.Arg885His | missense_variant | MODERATE | 0.44186 | 0 | 645 | 322 | . | . | . | . |
| PM1548-SK1P4 | PI4KB | p.Val202Met | missense_variant | MODERATE | 0 | 0 | 15 | 157 | . | . | . | . |
| PM1548-SK1P4 | CELF3 | p.Val279Ile | missense_variant | MODERATE | 0.6 | 0 | 20 | 54 | . | . | . | . |
| PM1548-SK1P4 | DENND4B | p.Ala154fs | frameshift_variant | HIGH | 0.208333 | 0 | 384 | 107 | . | . | . | . |
| PM1548-SK1P4 | CREB3L4 | p.Val263fs | frameshift_variant | HIGH | 0.398734 | 0 | 158 | 42 | . | . | . | . |
| PM1548-SK1P4 | EFNA4 | c.470-101delG | intron_variant | MODIFIER | 0 | 0 | 32 | 82 | . | . | . | . |
| PM1548-SK1P4 | ASH1L | p.Asn1907Lys | missense_variant | MODERATE | 0 | 0 | 46 | 55 | . | . | . | . |
| PM1548-SK1P4 | SMG5 | p.Gln846Lys | missense_variant | MODERATE | 0 | 0 | 40 | 128 | . | . | . | . |
| PM1548-SK1P4 | CD1D | p.Asp140Asn | missense_variant | MODERATE | 0.405405 | 0 | 37 | 198 | . | . | true | . |
| PM1548-SK1P4 | AIM2 | p.Lys342fs | frameshift_variant | HIGH | 0.444444 | 0 | 27 | 33 | . | . | . | . |
| PM1548-SK1P4 | NIT1 | p.Arg261His | missense_variant | MODERATE | 0.334302 | 0 | 344 | 402 | . | . | . | . |
| PM1548-SK1P4 | ILDR2 | p.Gly453Ala | missense_variant | MODERATE | 0 | 0 | 96 | 328 | . | . | . | . |
| PM1548-SK1P4 | ILDR2 | p.Arg415Trp | missense_variant | MODERATE | 0.486014 | 0 | 572 | 213 | . | . | . | . |
| PM1548-SK1P4 | MYOC | p.Arg91Gln | missense_variant | MODERATE | 0 | 0 | 470 | 242 | . | . | . | . |
| PM1548-SK1P4 | DNM3 | p.Glu7Asp | missense_variant | MODERATE | 0 | 0 | 28 | 77 | . | . | . | . |
| PM1548-SK1P4 | DNM3 | p.Phe20Phe | synonymous_variant | LOW | 0.169811 | 0.0060241 | 53 | 166 | . | . | . | . |
| PM1548-SK1P4 | DNM3 | p.His85Gln | missense_variant | MODERATE | 0.306667 | 0 | 75 | 56 | . | . | . | . |
| PM1548-SK1P4 | ANKRD45 | p.Leu39fs | frameshift_variant | HIGH | 0.512821 | 0 | 78 | 229 | . | . | . | . |
| PM1548-SK1P4 | ZBTB37 | p.Val284Ile | missense_variant | MODERATE | 0 | 0 | 33 | 152 | . | . | . | . |
| PM1548-SK1P4 | ZBTB37 | p.Asn288Ser | missense_variant | MODERATE | 0 | 0 | 28 | 160 | . | . | . | . |
| PM1548-SK1P4 | PAPPA2 | p.Thr1174fs | frameshift_variant | HIGH | 0.857143 | 0 | 7 | 36 | . | . | . | . |
| PM1548-SK1P4 | CACNA1E | p.Asn2301Ser | missense_variant | MODERATE | 0.05 | 0 | 80 | 120 | . | . | . | . |
| PM1548-SK1P4 | CACNA1E | p.Asn2302Asp | missense_variant | MODERATE | 0.05 | 0 | 80 | 119 | . | . | . | . |
| PM1548-SK1P4 | CACNA1E | p.Asp2310Asp | synonymous_variant | LOW | 0.0506329 | 0 | 79 | 90 | . | . | . | . |
| PM1548-SK1P4 | ZNF648 | p.Ser436Tyr | missense_variant | MODERATE | 0.109091 | 0 | 55 | 389 | . | . | . | . |
| PM1548-SK1P4 | ZNF648 | p.Ser436Pro | missense_variant | MODERATE | 0.105263 | 0 | 57 | 439 | . | . | . | . |
| PM1548-SK1P4 | ZNF648 | p.Pro423Ser | missense_variant | MODERATE | 0.101695 | 0.00228311 | 59 | 438 | . | . | . | . |
| PM1548-SK1P4 | C1orf27 | p.Thr381Ser | missense_variant | MODERATE | 0.5 | 0 | 2 | 10 | . | . | . | . |
| PM1548-SK1P4 | PLA2G4A | c.1033+2T>C | splice_donor_variant&intron_variant | HIGH | 0.307692 | 0 | 13 | 80 | . | . | . | . |
| PM1548-SK1P4 | ASPM | p.Arg2700Gln | missense_variant | MODERATE | 0 | 0.00653595 | 47 | 153 | . | . | . | . |
| PM1548-SK1P4 | DENND1B | c.83-20669G>A | intron_variant | MODIFIER | 0.403846 | 0 | 52 | 39 | . | . | . | . |
| PM1548-SK1P4 | NR5A2 | p.Gly169Val | missense_variant | MODERATE | 0.636364 | 0 | 11 | 71 | . | . | . | . |
| PM1548-SK1P4 | NAV1 | p.Thr719Ser | missense_variant | MODERATE | 0.0283019 | 0 | 212 | 176 | . | . | . | . |
| PM1548-SK1P4 | NAV1 | p.Thr726Ser | missense_variant | MODERATE | 0.0273973 | 0.00930233 | 219 | 215 | . | . | . | . |
| PM1548-SK1P4 | ATP2B4 | p.Asp867Asn | missense_variant&splice_region_variant | MODERATE | 0 | 0 | 24 | 121 | . | . | . | . |
| PM1548-SK1P4 | NUAK2 | p.Arg378His | missense_variant | MODERATE | 0.403727 | 0 | 161 | 162 | . | . | . | . |
| PM1548-SK1P4 | PFKFB2 | p.Glu261Ala | missense_variant | MODERATE | 0 | 0 | 103 | 81 | . | . | . | . |
| PM1548-SK1P4 | CR2 | p.Thr209fs | frameshift_variant | HIGH | 0.366667 | 0 | 30 | 118 | . | . | . | . |
| PM1548-SK1P4 | SMYD2 | p.Asn99fs | frameshift_variant | HIGH | 0.518519 | 0 | 27 | 181 | . | . | . | . |
| PM1548-SK1P4 | HHIPL2 | p.Gly156Ser | missense_variant | MODERATE | 0.255814 | 0 | 43 | 209 | . | . | . | . |
| PM1548-SK1P4 | MIA3 | p.Arg1545Gln | missense_variant | MODERATE | 0.470588 | 0 | 34 | 38 | . | . | . | . |
| PM1548-SK1P4 | LIN9 | p.Arg245Trp | missense_variant | MODERATE | 0.25 | 0 | 12 | 86 | . | . | . | . |
| PM1548-SK1P4 | ITPKB | p.Arg331His | missense_variant | MODERATE | 0.405467 | 0 | 439 | 395 | . | . | true | . |
| PM1548-SK1P4 | ITPKB | p.Leu42Val | missense_variant | MODERATE | 0.193277 | 0 | 119 | 35 | . | . | true | . |
| PM1548-SK1P4 | SNAP47 | p.Phe343Leu | missense_variant | MODERATE | 0 | 0 | 149 | 78 | . | . | . | . |
| PM1548-SK1P4 | IBA57 | p.Ala161Val | missense_variant | MODERATE | 0.428571 | 0 | 77 | 268 | . | . | . | . |
| PM1548-SK1P4 | OBSCN | p.Tyr2035Phe | missense_variant | MODERATE | 0.0427807 | 0 | 374 | 508 | . | . | . | . |
| PM1548-SK1P4 | OBSCN | p.Gln2065Lys | missense_variant | MODERATE | 0.0575916 | 0 | 191 | 171 | . | . | . | . |
| PM1548-SK1P4 | OBSCN | p.Cys2073Trp | missense_variant | MODERATE | 0 | 0 | 60 | 54 | . | . | . | . |
| PM1548-SK1P4 | ACTA1 | p.Thr231Met | missense_variant | MODERATE | 0 | 0 | 24 | 311 | . | . | . | . |
| PM1548-SK1P4 | GNPAT | p.Val551Met | missense_variant | MODERATE | 0.423077 | 0 | 52 | 79 | . | . | . | . |
| PM1548-SK1P4 | DISC1 | p.Asp580Gly | missense_variant | MODERATE | 0.694444 | 0 | 36 | 167 | . | . | . | . |
| PM1548-SK1P4 | GGPS1 | p.Val112Met | missense_variant | MODERATE | 0.0327869 | 0 | 61 | 405 | . | . | . | . |
| PM1548-SK1P4 | NID1 | p.Pro918fs | frameshift_variant&splice_region_variant | HIGH | 0.292683 | 0 | 41 | 46 | . | . | . | . |
| PM1548-SK1P4 | CEP170 | p.Glu1460Lys | missense_variant | MODERATE | 0.75 | 0 | 8 | 32 | . | . | . | . |
| PM1548-SK1P4 | ZNF496 | p.Arg606His | missense_variant | MODERATE | 0.411765 | 0 | 204 | 143 | . | . | . | . |
| PM1548-SK1P4 | OR2T4 | p.Cys60Ser | missense_variant | MODERATE | 1 | 0 | 1 | 107 | . | . | . | . |
| PM1548-SK1P4 | OR2T6 | p.Tyr60His | missense_variant | MODERATE | 0.454545 | 0 | 77 | 68 | . | . | . | . |
| PM1548-SK1P4 | DIP2C | p.Val1150Ile | missense_variant | MODERATE | 0 | 0 | 56 | 183 | . | . | . | . |
| PM1548-SK1P4 | WDR37 | p.Pro169Ser | missense_variant | MODERATE | 0.541284 | 0 | 109 | 220 | . | . | . | . |
| PM1548-SK1P4 | CALML5 | p.Glu102Lys | missense_variant | MODERATE | 0.514563 | 0 | 103 | 79 | . | . | . | . |
| PM1548-SK1P4 | FRMD4A | p.Pro1005fs | frameshift_variant | HIGH | 0 | 0 | 20 | 118 | . | . | . | . |
| PM1548-SK1P4 | CUBN | p.Trp3323Arg | missense_variant | MODERATE | 0 | 0 | 35 | 170 | . | . | . | . |
| PM1548-SK1P4 | GPR158 | p.Arg448Gln | missense_variant | MODERATE | 0.428571 | 0 | 21 | 52 | . | . | . | . |
| PM1548-SK1P4 | SVIL | p.Arg2075Cys | missense_variant | MODERATE | 0.6 | 0 | 20 | 40 | . | . | . | . |

| | | | | | | | | | | | | |
|---|---|---|---|---|---|---|---|---|---|---|---|---|
| PM1548-SK1P4 | PARD3 | p.Arg1278Gly | missense_variant | MODERATE | 0 | 0 | 70 | 91 | . | . | . | . |
| PM1548-SK1P4 | FRMPD2 | p.Pro226Leu | missense_variant | MODERATE | 0.413793 | 0 | 87 | 178 | . | . | . | . |
| PM1548-SK1P4 | PRKG1 | p.Asn601fs | frameshift_variant | HIGH | 0.62963 | 0 | 27 | 56 | . | . | . | . |
| PM1548-SK1P4 | PBLD | p.Asp167Tyr | missense_variant | MODERATE | 0 | 0 | 73 | 186 | . | . | . | . |
| PM1548-SK1P4 | DDX50 | p.Tyr39Cys | missense_variant | MODERATE | 0.5 | 0 | 10 | 39 | . | . | . | . |
| PM1548-SK1P4 | ADAMTS14 | p.Gly710Arg | missense_variant | MODERATE | 0.468085 | 0 | 47 | 78 | . | . | . | . |
| PM1548-SK1P4 | DDIT4 | p.Ser46Cys | missense_variant | MODERATE | 0.0952381 | 0 | 84 | 263 | . | . | . | . |
| PM1548-SK1P4 | DDIT4 | p.Thr47Ala | missense_variant | MODERATE | 0.0952381 | 0 | 84 | 263 | . | . | . | . |
| PM1548-SK1P4 | KAT6B | p.Arg463His | missense_variant | MODERATE | 0.42446 | 0 | 139 | 104 | true | . | . | . |
| PM1548-SK1P4 | KCNMA1 | p.Ala1039Val | missense_variant | MODERATE | 0.636364 | 0 | 22 | 74 | . | . | . | . |
| PM1548-SK1P4 | ZMIZ1 | p.Asp1051fs | frameshift_variant | HIGH | 0.416667 | 0 | 72 | 67 | . | . | . | . |
| PM1548-SK1P4 | MAT1A | p.Arg81Gln | missense_variant | MODERATE | 0.297297 | 0 | 37 | 225 | . | . | . | . |
| PM1548-SK1P4 | TSPAN14 | p.Asp190Gly | missense_variant | MODERATE | 0 | 0 | 29 | 134 | . | . | . | . |
| PM1548-SK1P4 | SH2D4B | p.Ala186Val | missense_variant | MODERATE | 0.48913 | 0 | 92 | 42 | . | . | . | . |
| PM1548-SK1P4 | LRIT2 | p.Thr520fs | frameshift_variant | HIGH | 0.548387 | 0 | 93 | 224 | . | . | . | . |
| PM1548-SK1P4 | CCSER2 | c.2326-13521C> | intron_variant | MODIFIER | 0 | 0 | 2 | 179 | . | . | . | . |
| PM1548-SK1P4 | CYP26C1 | p.Ser70Leu | missense_variant | MODERATE | 0 | 0 | 119 | 149 | . | . | . | . |
| PM1548-SK1P4 | MYOF | p.Arg726Gln | missense_variant | MODERATE | 0.4375 | 0 | 16 | 43 | . | . | . | . |
| PM1548-SK1P4 | SORBS1 | p.Arg585Cys | missense_variant | MODERATE | 0.267857 | 0 | 56 | 42 | . | . | . | . |
| PM1548-SK1P4 | ALDH18A1 | p.Arg622Trp | missense_variant | MODERATE | 0.551724 | 0 | 29 | 86 | . | . | . | . |
| PM1548-SK1P4 | ENTPD1 | p.Arg252Leu | missense_variant | MODERATE | 0 | 0 | 67 | 84 | . | . | . | . |
| PM1548-SK1P4 | MORN4 | p.Leu202Ile | missense_variant | MODERATE | 0 | 0 | 22 | 87 | . | . | . | . |
| PM1548-SK1P4 | PKD2L1 | p.Arg206Cys | missense_variant | MODERATE | 0.497908 | 0 | 239 | 130 | . | . | . | . |
| PM1548-SK1P4 | MRPL43 | p.Asn66Ser | missense_variant | MODERATE | 0.610922 | 0 | 293 | 81 | . | . | . | . |
| PM1548-SK1P4 | POLL | p.Arg419* | stop_gained | HIGH | 0 | 0 | 22 | 44 | . | . | . | . |
| PM1548-SK1P4 | TAF5 | p.Phe374fs | frameshift_variant | HIGH | 1 | 0 | 2 | 50 | . | . | . | . |
| PM1548-SK1P4 | WDR96 | p.Val70Ala | missense_variant | MODERATE | 0.5 | 0 | 14 | 59 | . | . | . | . |
| PM1548-SK1P4 | SORCS3 | p.Lys1023Gln | missense_variant | MODERATE | 0.0555556 | 0 | 36 | 58 | . | . | . | . |
| PM1548-SK1P4 | TCF7L2 | p.Ser587Leu | missense_variant | MODERATE | 0 | 0 | 149 | 171 | true | . | true | . |
| PM1548-SK1P4 | ABLIM1 | p.Leu633Met | missense_variant | MODERATE | 0 | 0 | 87 | 266 | . | . | . | . |
| PM1548-SK1P4 | GFRA1 | p.Ala173Val | missense_variant | MODERATE | 0 | 0 | 9 | 73 | . | . | . | . |
| PM1548-SK1P4 | KIAA1598 | p.Arg482His | missense_variant | MODERATE | 0.292683 | 0 | 41 | 181 | true | . | . | . |
| PM1548-SK1P4 | VAX1 | p.Cys200* | stop_gained | HIGH | 0 | 0 | 151 | 96 | . | . | . | . |
| PM1548-SK1P4 | EIF3A | p.His257His | synonymous_variant | LOW | 0 | 0 | 55 | 131 | . | . | . | . |
| PM1548-SK1P4 | ATE1 | p.Leu289Gln | missense_variant | MODERATE | 0.1875 | 0 | 16 | 77 | . | . | . | . |
| PM1548-SK1P4 | PLEKHA1 | p.Arg9His | missense_variant | MODERATE | 0.625 | 0 | 8 | 43 | . | . | . | . |
| PM1548-SK1P4 | PLEKHA1 | p.Val404Met | missense_variant | MODERATE | 0.275 | 0 | 40 | 76 | . | . | . | . |
| PM1548-SK1P4 | HMX3 | p.Pro23fs | frameshift_variant | HIGH | 0.6625 | 0 | 160 | 56 | . | . | . | . |
| PM1548-SK1P4 | CLRN3 | p.Pro71Thr | missense_variant | MODERATE | 0.0869565 | 0.00613497 | 69 | 163 | . | . | . | . |
| PM1548-SK1P4 | MKI67 | p.Arg2258* | stop_gained | HIGH | 0.37037 | 0 | 27 | 90 | . | . | . | . |
| PM1548-SK1P4 | TTC40 | p.Thr1864Met | missense_variant | MODERATE | 0.410714 | 0 | 56 | 72 | . | . | . | . |
| PM1548-SK1P4 | KNDC1 | p.Cys79fs | frameshift_variant | HIGH | 0.577778 | 0 | 45 | 162 | . | . | . | . |
| PM1548-SK1P4 | PIDD | p.Ala727Glu | missense_variant | MODERATE | 0 | 0 | 260 | 108 | . | . | . | . |
| PM1548-SK1P4 | MUC6 | p.Thr1861Ala | missense_variant | MODERATE | 0.652174 | 0 | 92 | 78 | . | . | . | . |
| PM1548-SK1P4 | MUC6 | p.Ile1857Thr | missense_variant | MODERATE | 0.655556 | 0 | 90 | 78 | . | . | . | . |
| PM1548-SK1P4 | MUC6 | p.His1752Gln | missense_variant | MODERATE | 0.0557621 | 0 | 269 | 31 | . | . | . | . |
| PM1548-SK1P4 | MUC6 | p.Thr1711Ile | missense_variant | MODERATE | 0.0810811 | 0 | 111 | 51 | . | . | . | . |
| PM1548-SK1P4 | MUC2 | p.Gly1331Arg | missense_variant | MODERATE | 0.581818 | 0 | 55 | 70 | . | . | . | . |
| PM1548-SK1P4 | MUC5B | p.Ala110Thr | missense_variant | MODERATE | 0.642612 | 0 | 291 | 216 | . | . | . | . |
| PM1548-SK1P4 | BRSK2 | p.Pro440fs | frameshift_variant | HIGH | 0.473149 | 0 | 689 | 593 | . | . | . | . |
| PM1548-SK1P4 | TSPAN32 | p.Ala178Val | missense_variant | MODERATE | 0.490196 | 0 | 153 | 69 | . | . | . | . |
| PM1548-SK1P4 | TSSC4 | p.Arg92Cys | missense_variant | MODERATE | 0.517094 | 0 | 1404 | 524 | . | . | . | . |
| PM1548-SK1P4 | OSBPL5 | p.Thr843Met | missense_variant | MODERATE | 0 | 0 | 20 | 106 | . | . | . | . |
| PM1548-SK1P4 | STIM1 | p.Lys371Arg | missense_variant | MODERATE | 0.0481928 | 0 | 83 | 322 | . | . | . | . |
| PM1548-SK1P4 | OR51E1 | p.Tyr63His | missense_variant | MODERATE | 0.227891 | 0 | 294 | 116 | . | . | . | . |
| PM1548-SK1P4 | OR51A2 | p.Lys289Asn | missense_variant | MODERATE | 0.272727 | 0 | 11 | 30 | . | . | . | . |
| PM1548-SK1P4 | SOX6 | p.Met803Ile | missense_variant | MODERATE | 0.0517241 | 0 | 58 | 33 | . | . | . | . |
| PM1548-SK1P4 | SOX6 | p.Met803Thr | missense_variant | MODERATE | 0.0517241 | 0 | 58 | 33 | . | . | . | . |
| PM1548-SK1P4 | SOX6 | p.Met803Val | missense_variant | MODERATE | 0.0517241 | 0 | 58 | 33 | . | . | . | . |
| PM1548-SK1P4 | ABCC8 | p.Asp761Asn | missense_variant | MODERATE | 0.313953 | 0 | 86 | 230 | . | . | . | . |
| PM1548-SK1P4 | LDHC | p.Lys5Glu | missense_variant | MODERATE | 0.375 | 0 | 8 | 119 | . | . | . | . |
| PM1548-SK1P4 | MRGPRX2 | p.Asn62Asn | synonymous_variant | LOW | 0.390625 | 0 | 64 | 83 | . | . | . | . |
| PM1548-SK1P4 | ZDHHC13 | p.Gly591Asp | missense_variant | MODERATE | 0.8 | 0 | 5 | 33 | . | . | . | . |
| PM1548-SK1P4 | ANO3 | p.Leu273Pro | missense_variant | MODERATE | 0.387097 | 0 | 62 | 432 | . | . | . | . |
| PM1548-SK1P4 | HIPK3 | p.Asp532Ala | missense_variant | MODERATE | 0.0141844 | 0 | 141 | 54 | . | . | . | . |
| PM1548-SK1P4 | ABTB2 | p.Asp793Asn | missense_variant | MODERATE | 0.429864 | 0.00668896 | 221 | 299 | . | . | . | . |
| PM1548-SK1P4 | RAG1 | p.Arg507Trp | missense_variant | MODERATE | 0.321767 | 0 | 317 | 86 | . | . | . | true |
| PM1548-SK1P4 | TP53I11 | p.Leu62Arg | missense_variant | MODERATE | 0.00359712 | 0 | 556 | 34 | . | . | . | . |
| PM1548-SK1P4 | MAPK8IP1 | p.Thr321Met | missense_variant | MODERATE | 0.541502 | 0 | 253 | 188 | . | true | . | . |
| PM1548-SK1P4 | DGKZ | n.46369267G>A | intragenic_variant | MODIFIER | 0.0204082 | 0 | 49 | 32 | . | . | . | . |
| PM1548-SK1P4 | DGKZ | p.Asp574Asn | missense_variant | MODERATE | 0.530973 | 0 | 113 | 74 | . | . | . | . |
| PM1548-SK1P4 | ACP2 | p.Arg231His | missense_variant | MODERATE | 0.515957 | 0 | 376 | 199 | . | . | . | . |
| PM1548-SK1P4 | MYBPC3 | p.Arg654Ser | missense_variant | MODERATE | 0 | 0 | 25 | 97 | . | . | . | . |
| PM1548-SK1P4 | OR5D14 | p.Tyr261Cys | missense_variant | MODERATE | 0.333333 | 0 | 6 | 71 | . | . | . | . |
| PM1548-SK1P4 | OR5W2 | p.Ala29Ser | missense_variant | MODERATE | 0.189189 | 0 | 37 | 99 | . | . | . | . |
| PM1548-SK1P4 | APLNR | p.Ala161Thr | missense_variant | MODERATE | 0.208333 | 0 | 24 | 160 | . | . | . | . |
| PM1548-SK1P4 | OR10Q1 | p.Thr287Ile | missense_variant | MODERATE | 0.571429 | 0 | 7 | 90 | . | . | . | . |
| PM1548-SK1P4 | DTX4 | p.Lys396fs | frameshift_variant | HIGH | 0.449541 | 0 | 109 | 34 | . | . | . | . |
| PM1548-SK1P4 | DTX4 | p.Gly510fs | frameshift_variant | HIGH | 0.458333 | 0.00719424 | 48 | 139 | . | . | . | . |
| PM1548-SK1P4 | DAK | p.Arg380Gln | missense_variant | MODERATE | 0.444816 | 0 | 598 | 316 | . | . | . | . |
| PM1548-SK1P4 | FEN1 | p.Arg332Cys | missense_variant | MODERATE | 0 | 0 | 59 | 80 | . | . | . | . |
| PM1548-SK1P4 | BEST1 | p.Ser413Asn | missense_variant | MODERATE | 0.555556 | 0 | 63 | 159 | . | . | . | . |
| PM1548-SK1P4 | AHNAK | p.Ser4908Leu | missense_variant | MODERATE | 0.464052 | 0 | 459 | 39 | . | . | . | . |
| PM1548-SK1P4 | AHNAK | p.His4772Asn | missense_variant | MODERATE | 0 | 0 | 156 | 54 | . | . | . | . |
| PM1548-SK1P4 | AHNAK | p.Gly4561Asp | missense_variant | MODERATE | 0.0664062 | 0 | 256 | 206 | . | . | . | . |
| PM1548-SK1P4 | AHNAK | p.Val4560Leu | missense_variant | MODERATE | 0.0625 | 0 | 256 | 203 | . | . | . | . |
| PM1548-SK1P4 | AHNAK | p.Val3640Ile | missense_variant | MODERATE | 0.361111 | 0 | 36 | 44 | . | . | . | . |
| PM1548-SK1P4 | AHNAK | p.Asn3637Lys | missense_variant | MODERATE | 0.322581 | 0 | 31 | 44 | . | . | . | . |
| PM1548-SK1P4 | AHNAK | p.Lys2706fs | frameshift_variant | HIGH | 0.540741 | 0 | 270 | 43 | . | . | . | . |
| PM1548-SK1P4 | AHNAK | p.Thr1986Ala | missense_variant | MODERATE | 0.130841 | 0 | 107 | 324 | . | . | . | . |
| PM1548-SK1P4 | CHRM1 | p.Pro319fs | frameshift_variant | HIGH | 0 | 0 | 11 | 31 | . | . | . | . |
| PM1548-SK1P4 | SLC22A6 | p.Val240Met | missense_variant | MODERATE | 0 | 0 | 33 | 227 | . | . | . | . |
| PM1548-SK1P4 | SLC22A25 | p.Asn62Asp | missense_variant | MODERATE | 0 | 0 | 94 | 110 | . | . | . | . |
| PM1548-SK1P4 | C11orf95 | n.1245C>T | non_coding_exon_variant | MODIFIER | 0.444444 | 0 | 18 | 34 | . | . | . | . |
| PM1548-SK1P4 | NRXN2 | p.Asp277Asn | missense_variant | MODERATE | 0.290698 | 0 | 86 | 222 | . | . | . | . |
| PM1548-SK1P4 | SNX15 | p.Leu10Met | missense_variant | MODERATE | 0 | 0 | 6 | 37 | . | . | . | . |
| PM1548-SK1P4 | RBM4 | p.Tyr303His | missense_variant | MODERATE | 0 | 0 | 18 | 269 | . | . | . | . |
| PM1548-SK1P4 | SPTBN2 | p.Cys1393Ser | missense_variant | MODERATE | 0.076087 | 0 | 184 | 300 | . | . | . | . |
| PM1548-SK1P4 | SPTBN2 | p.Arg465Gln | missense_variant | MODERATE | 0.28125 | 0 | 96 | 212 | . | . | . | . |
| PM1548-SK1P4 | LRP5 | p.Ala724Val | missense_variant | MODERATE | 0.3125 | 0 | 16 | 227 | . | . | . | . |
| PM1548-SK1P4 | IGHMBP2 | p.Pro809Ser | missense_variant | MODERATE | 0.48062 | 0 | 129 | 132 | . | . | . | . |
| PM1548-SK1P4 | TPCN2 | p.Gly492Val | missense_variant | MODERATE | 0 | 0 | 31 | 21 | . | . | . | . |
| PM1548-SK1P4 | CCND1 | p.Val94Leu | missense_variant | MODERATE | 0.041543 | 0 | 337 | 259 | true | true | true | . |
| PM1548-SK1P4 | INPPL1 | p.Leu957Phe | missense_variant | MODERATE | 0.230769 | 0 | 65 | 30 | . | . | . | . |
| PM1548-SK1P4 | PDE2A | p.His99fs | frameshift_variant | HIGH | 0.598276 | 0 | 580 | 160 | . | . | . | . |
| PM1548-SK1P4 | RNF169 | p.Glu131* | stop_gained | HIGH | 0 | 0 | 74 | 52 | . | . | . | . |
| PM1548-SK1P4 | TENM4 | p.Ala235Thr | missense_variant | MODERATE | 0.686275 | 0 | 102 | 201 | . | . | . | . |
| PM1548-SK1P4 | PCF11 | p.Leu745Leu | synonymous_variant | LOW | 0.0496894 | 0 | 161 | 203 | . | . | . | . |
| PM1548-SK1P4 | PCF11 | p.Ser756Gly | missense_variant | MODERATE | 0.0481928 | 0 | 166 | 215 | . | . | . | . |
| PM1548-SK1P4 | PCF11 | p.Arg833Trp | missense_variant | MODERATE | 0.423398 | 0 | 718 | 258 | . | . | . | . |
| PM1548-SK1P4 | TYR | p.Gly191fs | frameshift_variant | HIGH | 0.3 | 0 | 10 | 94 | . | . | . | . |
| PM1548-SK1P4 | FAT3 | p.Ala2495Ser | missense_variant | MODERATE | 0 | 0 | 95 | 229 | . | . | . | . |
| PM1548-SK1P4 | PANX1 | p.Ser250Thr | missense_variant | MODERATE | 0.625 | 0 | 24 | 236 | . | . | . | . |
| PM1548-SK1P4 | MTMR2 | p.Lys451Glu | missense_variant | MODERATE | 0 | 0 | 48 | 77 | . | . | . | . |
| PM1548-SK1P4 | MTMR2 | p.Asp436Asn | missense_variant | MODERATE | 0 | 0 | 47 | 75 | . | . | . | . |
| PM1548-SK1P4 | MTMR2 | p.Leu433Val | missense_variant | MODERATE | 0 | 0 | 53 | 282 | . | . | . | . |
| PM1548-SK1P4 | MMP3 | p.Asp200Gly | missense_variant | MODERATE | 0 | 0 | 34 | 91 | . | . | . | . |
| PM1548-SK1P4 | DYNC2H1 | p.Tyr3523His | missense_variant | MODERATE | 0.411765 | 0 | 17 | 62 | . | . | . | . |
| PM1548-SK1P4 | CASP1 | p.Phe262Tyr | missense_variant | MODERATE | 0.25 | 0 | 4 | 177 | . | . | . | . |
| PM1548-SK1P4 | KBTBD3 | p.Lys478fs | frameshift_variant | HIGH | 0.307692 | 0 | 52 | 50 | . | . | . | . |
| PM1548-SK1P4 | NXPE1 | p.Thr169Ile | missense_variant | MODERATE | 0 | 0 | 20 | 97 | . | . | . | . |
| PM1548-SK1P4 | NXPE1 | p.Met157Ile | missense_variant | MODERATE | 0.26087 | 0 | 23 | 97 | . | . | . | . |
| PM1548-SK1P4 | NXPE1 | p.Met157Thr | missense_variant | MODERATE | 0.227273 | 0 | 22 | 96 | . | . | . | . |
| PM1548-SK1P4 | NXPE1 | p.Thr149Lys | missense_variant | MODERATE | 0.157895 | 0 | 19 | 118 | . | . | . | . |
| PM1548-SK1P4 | DSCAML1 | p.Val1762Ile | missense_variant | MODERATE | 0.383117 | 0 | 154 | 226 | . | . | . | . |
| PM1548-SK1P4 | DSCAML1 | p.Asp949fs | frameshift_variant | HIGH | 0.561024 | 0 | 508 | 244 | . | . | . | . |
| PM1548-SK1P4 | ATP5L | p.Leu84Trp | missense_variant | MODERATE | 0.1125 | 0 | 80 | 125 | . | . | . | . |

| | | | | | | | | | | | | |
|---|---|---|---|---|---|---|---|---|---|---|---|---|
| PM1548-SK1P4 | ATP5L | p.Val89Ile | missense_variant | MODERATE | 0.0963855 | 0 | 83 | 140 | . | . | . | . |
| PM1548-SK1P4 | ATP5L | p.Ile99Val | missense_variant | MODERATE | 0.0963855 | 0 | 83 | 141 | . | . | . | . |
| PM1548-SK1P4 | PHLDB1 | p.Ala932Ser | missense_variant | MODERATE | 0.105263 | 0 | 76 | 223 | . | . | . | . |
| PM1548-SK1P4 | PHLDB1 | p.Pro938Leu | missense_variant | MODERATE | 0.106667 | 0 | 75 | 220 | . | . | . | . |
| PM1548-SK1P4 | CBL | p.Ser237Leu | missense_variant | MODERATE | 0.363636 | 0 | 11 | 128 | true | . | . | . |
| PM1548-SK1P4 | CBL | p.Pro504Gln | missense_variant | MODERATE | 0 | 0 | 89 | 111 | true | . | . | . |
| PM1548-SK1P4 | CRTAM | p.Leu60fs | frameshift_variant | HIGH | 0.485782 | 0 | 422 | 169 | . | . | . | . |
| PM1548-SK1P4 | OR10G8 | p.Lys265Glu | missense_variant | MODERATE | 0.140845 | 0 | 142 | 232 | . | . | . | . |
| PM1548-SK1P4 | OR10G8 | p.Lys265Asn | missense_variant | MODERATE | 0.155556 | 0 | 135 | 193 | . | . | . | . |
| PM1548-SK1P4 | OR8G2 | p.Val2Gly | missense_variant | MODERATE | 1 | 0 | 1 | 1 | . | . | . | . |
| PM1548-SK1P4 | ROBO4 | p.Arg776His | missense_variant | MODERATE | 0.495868 | 0 | 121 | 263 | . | . | . | . |
| PM1548-SK1P4 | ADAMTS8 | p.Arg9Gly | missense_variant | MODERATE | 0.5 | 0 | 86 | 48 | . | . | . | true |
| PM1548-SK1P4 | IQSEC3 | p.Gln1075Lys | missense_variant | MODERATE | 0 | 0 | 161 | 59 | . | . | . | . |
| PM1548-SK1P4 | FOXM1 | p.Leu334Ser | missense_variant | MODERATE | 0.145455 | 0 | 55 | 177 | . | . | . | . |
| PM1548-SK1P4 | FOXM1 | p.Asp328Glu | missense_variant | MODERATE | 0.163265 | 0 | 49 | 203 | . | . | . | . |
| PM1548-SK1P4 | VWF | p.Ala865Asp | missense_variant | MODERATE | 0.243728 | 0 | 279 | 226 | . | . | . | . |
| PM1548-SK1P4 | IFFO1 | p.Val126Met | missense_variant | MODERATE | 0.453901 | 0 | 141 | 191 | . | . | . | . |
| PM1548-SK1P4 | PHB2 | p.Arg270Arg | synonymous_variant | LOW | 0.0550459 | 0 | 218 | 405 | . | . | . | . |
| PM1548-SK1P4 | PHB2 | p.Asn269Asn | synonymous_variant | LOW | 0.0552995 | 0 | 217 | 404 | . | . | . | . |
| PM1548-SK1P4 | TAS2R10 | p.Ser41Thr | missense_variant | MODERATE | 0.645161 | 0 | 31 | 57 | . | . | . | . |
| PM1548-SK1P4 | TAS2R31 | p.Ala177Thr | missense_variant | MODERATE | 0.0666667 | 0 | 30 | 47 | . | . | . | . |
| PM1548-SK1P4 | PLCZ1 | p.Gly133Val | missense_variant | MODERATE | 0 | 0 | 6 | 42 | . | . | . | . |
| PM1548-SK1P4 | PDE3A | p.Glu171fs | frameshift_variant | HIGH | 0.186667 | 0 | 75 | 56 | . | . | . | . |
| PM1548-SK1P4 | TM7SF3 | c.869-699C>G | intron_variant | MODIFIER | 0.460177 | 0 | 113 | 125 | . | . | . | . |
| PM1548-SK1P4 | DDX11 | p.Asp935Asn | missense_variant | MODERATE | 0.625 | 0 | 8 | 35 | . | . | . | . |
| PM1548-SK1P4 | COL2A1 | p.Asp682Asn | missense_variant | MODERATE | 0.565032 | 0 | 469 | 53 | true | . | . | . |
| PM1548-SK1P4 | KMT2D | p.Ala4804Val | missense_variant | MODERATE | 0.453333 | 0.0060241 | 75 | 166 | true | true | true | true |
| PM1548-SK1P4 | KMT2D | p.Leu4214Leu | synonymous_variant | LOW | 0.0357143 | 0 | 140 | 376 | true | true | true | true |
| PM1548-SK1P4 | KMT2D | p.Thr3166Ile | missense_variant | MODERATE | 0.354962 | 0 | 262 | 193 | true | true | true | true |
| PM1548-SK1P4 | KMT2D | p.Thr2581Ala | missense_variant | MODERATE | 0.0191388 | 0 | 418 | 134 | true | true | true | true |
| PM1548-SK1P4 | KMT2D | p.Gly1628fs | frameshift_variant | HIGH | 0 | 0 | 99 | 136 | true | true | true | true |
| PM1548-SK1P4 | PRPH | p.Ala348Thr | missense_variant | MODERATE | 0.48 | 0 | 175 | 109 | . | . | . | . |
| PM1548-SK1P4 | FAM186B | p.Leu494Met | missense_variant | MODERATE | 0.516854 | 0 | 267 | 412 | . | . | . | . |
| PM1548-SK1P4 | ANKRD33 | p.Ala173Val | missense_variant | MODERATE | 0.319495 | 0 | 554 | 241 | . | . | . | . |
| PM1548-SK1P4 | KRT74 | p.Gln440His | missense_variant | MODERATE | 0 | 0 | 12 | 34 | . | . | . | . |
| PM1548-SK1P4 | PCBP2 | p.Pro175fs | frameshift_variant | HIGH | 0.75 | 0 | 12 | 35 | . | . | . | . |
| PM1548-SK1P4 | HNRNPA1 | p.Thr61Ala | missense_variant | MODERATE | 0.46875 | 0 | 32 | 168 | . | . | . | . |
| PM1548-SK1P4 | ITGA5 | p.Arg193His | missense_variant | MODERATE | 0.235955 | 0 | 89 | 104 | . | . | . | . |
| PM1548-SK1P4 | MYO1A | p.Arg559Cys | missense_variant | MODERATE | 0.47013 | 0 | 385 | 190 | . | . | . | . |
| PM1548-SK1P4 | NAB2 | p.Pro388Thr | missense_variant | MODERATE | 0.0113636 | 0 | 88 | 78 | true | . | . | . |
| PM1548-SK1P4 | NAB2 | p.Pro388His | missense_variant | MODERATE | 0.0113636 | 0 | 88 | 78 | true | . | . | . |
| PM1548-SK1P4 | R3HDM2 | p.Arg956His | missense_variant | MODERATE | 0 | 0 | 50 | 33 | . | . | . | . |
| PM1548-SK1P4 | CTDSP2 | p.Glu259Asp | missense_variant | MODERATE | 0.0930233 | 0 | 43 | 90 | . | . | . | . |
| PM1548-SK1P4 | CTDSP2 | p.Ala258Thr | missense_variant | MODERATE | 0.0930233 | 0 | 43 | 90 | . | . | . | . |
| PM1548-SK1P4 | CTDSP2 | p.Ile251Val | missense_variant | MODERATE | 0.105263 | 0 | 38 | 94 | . | . | . | . |
| PM1548-SK1P4 | PTPRB | p.Val1992Ala | missense_variant | MODERATE | 0.101266 | 0 | 79 | 50 | true | . | . | . |
| PM1548-SK1P4 | NAV3 | p.Met1275Val | missense_variant | MODERATE | 0.08 | 0 | 25 | 67 | . | true | . | . |
| PM1548-SK1P4 | NAV3 | p.Pro1276Leu | missense_variant | MODERATE | 0.111111 | 0 | 36 | 130 | . | true | . | . |
| PM1548-SK1P4 | NAV3 | p.Thr1281Ser | missense_variant | MODERATE | 0.108108 | 0 | 37 | 146 | . | true | . | . |
| PM1548-SK1P4 | TMTC2 | p.Gly143Arg | missense_variant | MODERATE | 0.607843 | 0 | 51 | 67 | . | . | . | . |
| PM1548-SK1P4 | ALX1 | p.Thr68Ser | missense_variant | MODERATE | 0.0625 | 0 | 16 | 191 | . | . | . | . |
| PM1548-SK1P4 | FGD6 | p.Met965fs | frameshift_variant | HIGH | 0.789474 | 0 | 19 | 56 | . | . | . | . |
| PM1548-SK1P4 | APAF1 | p.Gln578Arg | missense_variant | MODERATE | 0.0344828 | 0 | 29 | 126 | . | . | . | . |
| PM1548-SK1P4 | SLC41A2 | p.Pro266Ser | missense_variant | MODERATE | 0 | 0 | 5 | 65 | . | . | . | . |
| PM1548-SK1P4 | PRDM4 | p.Glu313Glu | synonymous_variant | LOW | 0.0377358 | 0 | 106 | 274 | . | . | . | . |
| PM1548-SK1P4 | TMEM119 | p.Thr81fs | frameshift_variant | HIGH | 0.187925 | 0 | 1325 | 297 | . | . | . | . |
| PM1548-SK1P4 | FOXN4 | p.Ala462Thr | missense_variant | MODERATE | 0.527778 | 0.00490196 | 72 | 204 | . | . | . | . |
| PM1548-SK1P4 | FAM222A | p.Pro392Thr | missense_variant | MODERATE | 0.0569106 | 0 | 123 | 422 | . | . | . | . |
| PM1548-SK1P4 | FAM222A | p.Ser400Ala | missense_variant | MODERATE | 0.0551181 | 0 | 127 | 423 | . | . | . | . |
| PM1548-SK1P4 | TCHP | p.Ala371Thr | missense_variant | MODERATE | 0.32 | 0 | 25 | 45 | . | . | . | . |
| PM1548-SK1P4 | SH2B3 | p.Glu523fs | frameshift_variant | HIGH | 0 | 0 | 466 | 158 | true | . | . | . |
| PM1548-SK1P4 | HECTD4 | p.Ala3707Val | missense_variant | MODERATE | 0.589286 | 0 | 56 | 62 | . | . | . | . |
| PM1548-SK1P4 | TPCN1 | p.Leu234Arg | missense_variant | MODERATE | 0.238267 | 0 | 277 | 134 | . | . | . | . |
| PM1548-SK1P4 | RBM19 | p.Asn533Asn | synonymous_variant | LOW | 0.414286 | 0 | 210 | 250 | . | . | . | . |
| PM1548-SK1P4 | RAB35 | n.120541738C>T | intragenic_variant | MODIFIER | 0.488827 | 0 | 358 | 243 | . | . | . | . |
| PM1548-SK1P4 | GCN1L1 | p.Ser2359Pro | missense_variant | MODERATE | 0.296296 | 0 | 27 | 85 | . | . | . | . |
| PM1548-SK1P4 | SETD1B | p.His8fs | frameshift_variant | HIGH | 0.0333333 | 0 | 30 | 141 | . | . | . | . |
| PM1548-SK1P4 | SETD1B | p.Pro1361fs | frameshift_variant | HIGH | 0.624365 | 0 | 197 | 190 | . | . | . | . |
| PM1548-SK1P4 | ZCCHC8 | p.Gly551Asp | missense_variant | MODERATE | 0.416667 | 0 | 24 | 34 | true | . | . | . |
| PM1548-SK1P4 | ABCB9 | p.His709Gln | missense_variant | MODERATE | 0.0763889 | 0 | 144 | 333 | . | . | . | . |
| PM1548-SK1P4 | ABCB9 | p.His709Arg | missense_variant | MODERATE | 0.0833333 | 0 | 144 | 333 | . | . | . | . |
| PM1548-SK1P4 | ABCB9 | p.Lys694Arg | missense_variant | MODERATE | 0.100719 | 0 | 139 | 301 | . | . | . | . |
| PM1548-SK1P4 | PITPNM2 | p.Arg749Trp | missense_variant | MODERATE | 0.457143 | 0 | 105 | 160 | . | . | . | . |
| PM1548-SK1P4 | SBNO1 | p.Arg1283Cys | missense_variant&splice_region_variant | MODERATE | 0.269006 | 0 | 171 | 244 | . | . | . | . |
| PM1548-SK1P4 | DNAH10 | p.Pro3133His | missense_variant | MODERATE | 0.428571 | 0.00452489 | 42 | 221 | . | . | . | . |
| PM1548-SK1P4 | TMEM132B | p.Ala297Thr | missense_variant | MODERATE | 0 | 0 | 5 | 79 | . | . | . | . |
| PM1548-SK1P4 | GPR133 | p.Gly731Arg | missense_variant | MODERATE | 0.733333 | 0 | 15 | 110 | . | . | . | . |
| PM1548-SK1P4 | FBRSL1 | p.Glu251Asp | missense_variant | MODERATE | 0.0023753 | 0 | 421 | 153 | . | . | . | . |
| PM1548-SK1P4 | MPHOSPH8 | p.Val686Ile | missense_variant | MODERATE | 0 | 0 | 69 | 75 | . | . | . | . |
| PM1548-SK1P4 | GJA3 | p.Ala213Thr | missense_variant | MODERATE | 0.18797 | 0 | 133 | 153 | . | . | . | . |
| PM1548-SK1P4 | LATS2 | p.Pro41Gln | missense_variant | MODERATE | 0.6 | 0.025641 | 25 | 39 | . | . | . | . |
| PM1548-SK1P4 | CDK8 | p.Asn372Thr | missense_variant | MODERATE | 0.0666667 | 0 | 30 | 249 | . | . | . | . |
| PM1548-SK1P4 | POLR1D | n.28240022G>A | intragenic_variant | MODIFIER | 0.591429 | 0 | 350 | 101 | . | . | . | . |
| PM1548-SK1P4 | FRY | p.Gln2970His | missense_variant | MODERATE | 0.111111 | 0 | 45 | 39 | . | . | . | . |
| PM1548-SK1P4 | BRCA2 | p.Asn863fs | frameshift_variant | HIGH | 0.333333 | 0 | 57 | 84 | true | true | . | true |
| PM1548-SK1P4 | STARD13 | p.Arg155His | missense_variant | MODERATE | 0.111111 | 0 | 18 | 32 | . | . | . | . |
| PM1548-SK1P4 | MAB21L1 | p.Ser103Ser | synonymous_variant | LOW | 0.666667 | 0 | 6 | 101 | . | . | . | . |
| PM1548-SK1P4 | FOXO1 | p.Ser152Thr | missense_variant | MODERATE | 0.0333333 | 0 | 60 | 140 | true | . | . | . |
| PM1548-SK1P4 | KBTBD7 | p.Tyr195Phe | missense_variant | MODERATE | 0.153846 | 0 | 78 | 644 | . | . | . | . |
| PM1548-SK1P4 | GPALPP1 | p.Pro37Arg | missense_variant | MODERATE | 1 | 0 | 9 | 103 | . | . | . | . |
| PM1548-SK1P4 | NEK3 | p.Asn343Lys | missense_variant | MODERATE | 0 | 0 | 27 | 44 | . | . | . | . |
| PM1548-SK1P4 | DACH1 | p.Arg196Arg | synonymous_variant | LOW | 0.226415 | 0 | 53 | 115 | . | . | . | . |
| PM1548-SK1P4 | ABCC4 | p.Ile1219fs | frameshift_variant | HIGH | 0 | 0 | 168 | 130 | . | . | . | . |
| PM1548-SK1P4 | FGF14 | p.Asp33Gly | missense_variant | MODERATE | 0.111111 | 0 | 36 | 97 | . | . | . | . |
| PM1548-SK1P4 | CCDC168 | p.Met4045fs | frameshift_variant | HIGH | 0.352941 | 0 | 85 | 95 | . | . | . | . |
| PM1548-SK1P4 | ARHGEF7 | p.Thr23fs | frameshift_variant | HIGH | 0.243328 | 0 | 637 | 189 | . | . | . | . |
| PM1548-SK1P4 | MCF2L | p.Arg590Gln | missense_variant | MODERATE | 0.454545 | 0 | 22 | 34 | . | . | . | . |
| PM1548-SK1P4 | CDC16 | p.Asn557fs | frameshift_variant | HIGH | 0 | 0 | 20 | 90 | . | . | . | . |
| PM1548-SK1P4 | UPF3A | p.Asp129Glu | missense_variant | MODERATE | 0.50774 | 0 | 323 | 60 | . | . | . | . |
| PM1548-SK1P4 | OR11G2 | p.Phe141Phe | synonymous_variant | LOW | 0.238095 | 0 | 42 | 117 | . | . | . | . |
| PM1548-SK1P4 | ACIN1 | p.Glu192Asp | missense_variant | MODERATE | 0.00704225 | 0 | 142 | 129 | . | . | . | . |
| PM1548-SK1P4 | ACIN1 | p.Glu192Lys | missense_variant | MODERATE | 0 | 0 | 142 | 129 | . | . | . | . |
| PM1548-SK1P4 | PABPN1 | c.*5dupA | frameshift_variant&stop_retained_variant | HIGH | 0.26087 | 0 | 23 | 92 | . | . | . | . |
| PM1548-SK1P4 | RNF31 | p.Ala9Gly | missense_variant | MODERATE | 0 | 0 | 70 | 104 | . | . | . | true |
| PM1548-SK1P4 | RNF31 | p.Val12Ala | missense_variant | MODERATE | 0 | 0 | 70 | 104 | . | . | . | true |
| PM1548-SK1P4 | LTB4R2 | p.Ala92Val | missense_variant | MODERATE | 0.0608696 | 0 | 115 | 173 | . | . | . | . |
| PM1548-SK1P4 | FOXG1 | p.Glu173Asp | missense_variant | MODERATE | 0.032967 | 0 | 91 | 31 | . | . | . | . |
| PM1548-SK1P4 | CLEC14A | p.Ser404Asn | missense_variant | MODERATE | 0 | 0 | 76 | 138 | . | . | . | . |
| PM1548-SK1P4 | KTN1 | p.Phe1182Val | missense_variant | MODERATE | 0.652174 | 0 | 23 | 114 | true | . | . | . |
| PM1548-SK1P4 | KCNH5 | p.Arg333His | missense_variant | MODERATE | 0.642857 | 0 | 42 | 68 | . | . | . | . |
| PM1548-SK1P4 | ZBTB25 | p.His371Asn | missense_variant | MODERATE | 0 | 0 | 60 | 47 | . | . | . | . |
| PM1548-SK1P4 | HSPA2 | p.Gly138fs | frameshift_variant | HIGH | 0.172043 | 0 | 279 | 148 | . | . | . | . |
| PM1548-SK1P4 | HSPA2 | p.Phe220Phe | synonymous_variant | LOW | 0.0588235 | 0 | 85 | 96 | . | . | . | . |
| PM1548-SK1P4 | FUT8 | p.Arg249Cys | missense_variant | MODERATE | 0.6 | 0 | 35 | 316 | . | . | . | . |
| PM1548-SK1P4 | TMEM229B | p.Leu37Phe | missense_variant | MODERATE | 0.0833333 | 0 | 216 | 483 | . | . | . | . |
| PM1548-SK1P4 | ZFP36L1 | p.Gly271fs | frameshift_variant | HIGH | 0.505464 | 0 | 366 | 215 | . | . | . | . |
| PM1548-SK1P4 | ELMSAN1 | p.Arg739Cys | missense_variant | MODERATE | 0.463277 | 0 | 177 | 46 | . | . | . | . |
| PM1548-SK1P4 | ELMSAN1 | p.Gln36fs | frameshift_variant | HIGH | 0.531792 | 0 | 173 | 43 | . | . | . | . |
| PM1548-SK1P4 | VRTN | p.Gly328fs | frameshift_variant | HIGH | 0.4375 | 0 | 32 | 58 | . | . | . | . |
| PM1548-SK1P4 | GALC | p.Arg685His | missense_variant | MODERATE | 0 | 0 | 23 | 31 | . | . | . | . |
| PM1548-SK1P4 | PTPN21 | p.Ile849fs | frameshift_variant | HIGH | 0.571429 | 0 | 7 | 156 | . | . | . | . |
| PM1548-SK1P4 | ZC3H14 | p.Asn577Asp | missense_variant | MODERATE | 0.0392157 | 0 | 51 | 222 | . | . | . | . |
| PM1548-SK1P4 | CCDC88C | p.Lys818Glu | missense_variant | MODERATE | 0 | 0 | 44 | 69 | . | . | . | . |
| PM1548-SK1P4 | FBLN5 | p.Asp357Asn | missense_variant | MODERATE | 0.352941 | 0 | 17 | 147 | . | . | . | . |
| PM1548-SK1P4 | UNC79 | p.Arg746Gln | missense_variant | MODERATE | 0.333333 | 0 | 9 | 31 | . | . | . | . |
| PM1548-SK1P4 | SYNE3 | p.Ile83Val | missense_variant | MODERATE | 0.421875 | 0 | 128 | 217 | . | . | . | . |
| PM1548-SK1P4 | SETD3 | p.Phe388Leu | missense_variant | MODERATE | 0.5 | 0 | 4 | 32 | . | . | . | . |

| | | | | | | | | | | | | |
|---|---|---|---|---|---|---|---|---|---|---|---|---|
| PM1548-SK1P4 | TECPR2 | p.Gln498fs | frameshift_variant | HIGH | 0.3 | 0 | 10 | 55 | . | . | . | . |
| PM1548-SK1P4 | BRF1 | p.Ala129Ser | missense_variant | MODERATE | 0 | 0 | 41 | 135 | . | . | . | . |
| PM1548-SK1P4 | CYFIP1 | p.Arg1085Leu | missense_variant | MODERATE | 0 | 0 | 37 | 334 | . | . | . | . |
| PM1548-SK1P4 | MAGEL2 | p.Ala11Val | missense_variant | MODERATE | 0.171975 | 0 | 785 | 323 | . | . | . | . |
| PM1548-SK1P4 | PAR5 | n.2021_2022del | intron_variant | MODIFIER | 0.422222 | 0 | 45 | 260 | . | . | . | . |
| PM1548-SK1P4 | UBE3A | p.Val133Ala | missense_variant | MODERATE | 0.384615 | 0 | 13 | 79 | . | . | . | . |
| PM1548-SK1P4 | ACTC1 | p.Ala137Thr | missense_variant | MODERATE | 0.666667 | 0 | 18 | 73 | . | . | . | . |
| PM1548-SK1P4 | SPRED1 | p.Arg332His | missense_variant | MODERATE | 0.555556 | 0.0070922 | 36 | 141 | . | . | . | . |
| PM1548-SK1P4 | CASC5 | p.Asp178Gly | missense_variant | MODERATE | 0.363636 | 0 | 33 | 58 | true | . | . | . |
| PM1548-SK1P4 | SPTBN5 | p.Arg1943His | missense_variant | MODERATE | 0.170732 | 0 | 123 | 148 | . | . | . | . |
| PM1548-SK1P4 | SPTBN5 | p.Val1842Gly | missense_variant | MODERATE | 0.0675676 | 0 | 74 | 87 | . | . | . | . |
| PM1548-SK1P4 | GANC | p.Ala394Val | missense_variant | MODERATE | 0 | 0 | 39 | 106 | . | . | . | . |
| PM1548-SK1P4 | GANC | p.Gly404Asp | missense_variant | MODERATE | 0 | 0 | 7 | 105 | . | . | . | . |
| PM1548-SK1P4 | TTBK2 | p.Ala197Thr | missense_variant | MODERATE | 0.333333 | 0 | 12 | 69 | . | . | . | . |
| PM1548-SK1P4 | CCNDBP1 | p.Asp237Asn | missense_variant | MODERATE | 0.461538 | 0 | 13 | 44 | . | . | . | . |
| PM1548-SK1P4 | TGM7 | p.Ser305Phe | missense_variant | MODERATE | 0.45679 | 0 | 81 | 54 | . | . | . | . |
| PM1548-SK1P4 | ZSCAN29 | p.Arg33Trp | missense_variant | MODERATE | 0 | 0 | 67 | 209 | . | . | . | . |
| PM1548-SK1P4 | SLC28A2 | p.Gly128Asp | missense_variant | MODERATE | 0.09375 | 0 | 64 | 169 | . | . | . | . |
| PM1548-SK1P4 | MYO5A | p.Arg831His | missense_variant | MODERATE | 0.516304 | 0 | 368 | 78 | true | . | . | . |
| PM1548-SK1P4 | TCF12 | p.Val183fs | frameshift_variant | HIGH | 0 | 0 | 10 | 200 | true | . | . | . |
| PM1548-SK1P4 | LIPC | p.Phe286fs | frameshift_variant | HIGH | 0.666667 | 0 | 9 | 89 | . | . | . | . |
| PM1548-SK1P4 | OAZ2 | p.Ser92Ser | synonymous_variant | LOW | 0 | 0 | 94 | 163 | . | . | . | . |
| PM1548-SK1P4 | OAZ2 | p.Glu87Gly | missense_variant | MODERATE | 0 | 0 | 93 | 155 | . | . | . | . |
| PM1548-SK1P4 | OAZ2 | p.Glu84Val | missense_variant | MODERATE | 0.00970874 | 0 | 103 | 248 | . | . | . | . |
| PM1548-SK1P4 | OAZ2 | p.Val79Gly | missense_variant | MODERATE | 0 | 0 | 92 | 233 | . | . | . | . |
| PM1548-SK1P4 | SPG21 | p.Ala282Val | missense_variant | MODERATE | 0.392857 | 0 | 28 | 32 | . | . | . | . |
| PM1548-SK1P4 | CILP | p.Asn930Asn | synonymous_variant | LOW | 0.276596 | 0.00877193 | 47 | 114 | . | . | . | . |
| PM1548-SK1P4 | IGDCC3 | p.Pro358Leu | missense_variant | MODERATE | 0 | 0 | 310 | 237 | . | . | . | . |
| PM1548-SK1P4 | NOX5 | p.His470fs | frameshift_variant | HIGH | 0.0875 | 0 | 80 | 164 | . | . | . | . |
| PM1548-SK1P4 | MYO9A | p.Cys1767Ser | missense_variant | MODERATE | 0.157895 | 0 | 38 | 186 | . | . | . | . |
| PM1548-SK1P4 | NPTN | p.Arg334Trp | missense_variant | MODERATE | 0.3125 | 0 | 64 | 86 | . | . | . | . |
| PM1548-SK1P4 | ISLR2 | p.Val74Ala | missense_variant | MODERATE | 0.469799 | 0 | 149 | 331 | . | . | . | . |
| PM1548-SK1P4 | C15orf39 | p.Ala475Val | missense_variant | MODERATE | 0.507028 | 0.00393701 | 996 | 254 | . | . | . | . |
| PM1548-SK1P4 | ETFA | p.Thr45Arg | missense_variant | MODERATE | 0.129032 | 0 | 31 | 214 | . | . | . | . |
| PM1548-SK1P4 | ETFA | p.Thr45Ala | missense_variant | MODERATE | 0.129032 | 0 | 31 | 214 | . | . | . | . |
| PM1548-SK1P4 | PSTPIP1 | p.Ala280fs | frameshift_variant&splice_region_variant | HIGH | 0.449807 | 0 | 518 | 438 | . | . | . | . |
| PM1548-SK1P4 | PEAK1 | p.Ala36Thr | missense_variant | MODERATE | 0 | 0 | 19 | 189 | . | . | . | . |
| PM1548-SK1P4 | CHRNB4 | p.Gly261Asp | missense_variant | MODERATE | 0.176471 | 0 | 17 | 56 | . | . | . | . |
| PM1548-SK1P4 | MESDC1 | p.Ser224Asn | missense_variant | MODERATE | 0.626506 | 0 | 249 | 161 | . | . | . | . |
| PM1548-SK1P4 | IL16 | p.Thr417Met | missense_variant | MODERATE | 0 | 0 | 58 | 34 | . | . | . | . |
| PM1548-SK1P4 | BTBD1 | p.Arg236Gln | missense_variant | MODERATE | 0 | 0 | 23 | 77 | . | . | . | . |
| PM1548-SK1P4 | ZNF592 | p.Ala1232Val | missense_variant | MODERATE | 0.454545 | 0 | 11 | 35 | . | . | . | . |
| PM1548-SK1P4 | AEN | p.Ala194Val | missense_variant | MODERATE | 0 | 0 | 28 | 122 | . | . | . | . |
| PM1548-SK1P4 | IQGAP1 | p.Met1231fs | frameshift_variant | HIGH | 0.548611 | 0 | 288 | 170 | . | . | . | . |
| PM1548-SK1P4 | FES | c.1531-47G>A | intron_variant | MODIFIER | 0.378882 | 0 | 483 | 35 | . | . | . | . |
| PM1548-SK1P4 | MAN2A2 | p.Pro139Leu | missense_variant | MODERATE | 0.470314 | 0 | 859 | 490 | . | . | . | . |
| PM1548-SK1P4 | RCCD1 | p.Arg87Ser | missense_variant | MODERATE | 0 | 0 | 16 | 63 | . | . | . | . |
| PM1548-SK1P4 | ADAMTS17 | p.Cys921Tyr | missense_variant | MODERATE | 0.368421 | 0 | 38 | 57 | . | . | . | . |
| PM1548-SK1P4 | WDR90 | p.Ala1738Thr | missense_variant | MODERATE | 0.522727 | 0 | 88 | 185 | . | . | . | . |
| PM1548-SK1P4 | RPUSD1 | p.Arg255Gln | missense_variant | MODERATE | 0.421687 | 0 | 83 | 198 | . | . | . | . |
| PM1548-SK1P4 | SOX8 | p.Pro11Thr | missense_variant | MODERATE | 0 | 0 | 82 | 62 | . | . | . | . |
| PM1548-SK1P4 | SOX8 | p.Asp27Asn | missense_variant | MODERATE | 0 | 0 | 165 | 120 | . | . | . | . |
| PM1548-SK1P4 | CACNA1H | p.Arg1892His | missense_variant | MODERATE | 0.223684 | 0 | 76 | 231 | . | . | . | . |
| PM1548-SK1P4 | BAIAP3 | p.Ala740Asp | missense_variant | MODERATE | 0 | 0 | 31 | 90 | . | . | . | . |
| PM1548-SK1P4 | IFT140 | p.Ala1330Val | missense_variant | MODERATE | 0 | 0 | 377 | 188 | . | . | . | . |
| PM1548-SK1P4 | NME3 | p.Ala20Ser | missense_variant | MODERATE | 0 | 0 | 183 | 162 | . | . | . | . |
| PM1548-SK1P4 | PRSS33 | p.Leu174fs | frameshift_variant | HIGH | 0 | 0 | 284 | 199 | . | . | . | . |
| PM1548-SK1P4 | CCDC64B | p.Arg144Gln | missense_variant | MODERATE | 0 | 0.00254453 | 793 | 393 | . | . | . | . |
| PM1548-SK1P4 | CREBBP | p.Gly1479Val | missense_variant | MODERATE | 0 | 0 | 80 | 193 | true | . | true | . |
| PM1548-SK1P4 | CORO7 | p.Thr230Ala | missense_variant | MODERATE | 0.263323 | 0 | 319 | 171 | . | . | . | . |
| PM1548-SK1P4 | GLYR1 | p.Ser148Thr | missense_variant | MODERATE | 0.280488 | 0 | 82 | 42 | . | . | . | . |
| PM1548-SK1P4 | TVP23A | p.Phe140fs | frameshift_variant | HIGH | 0.522152 | 0 | 316 | 182 | . | . | . | . |
| PM1548-SK1P4 | CIITA | p.Arg580His | missense_variant | MODERATE | 0.602273 | 0 | 528 | 335 | true | . | . | . |
| PM1548-SK1P4 | RMI2 | p.Ile130Val | missense_variant | MODERATE | 0 | 0 | 49 | 171 | true | . | . | . |
| PM1548-SK1P4 | TNFRSF17 | p.Pro145Gln | missense_variant | MODERATE | 0.204545 | 0 | 44 | 38 | true | . | . | . |
| PM1548-SK1P4 | CDR2 | p.Gln130Arg | missense_variant | MODERATE | 0.488038 | 0 | 209 | 85 | . | . | . | . |
| PM1548-SK1P4 | USP31 | p.Gly1267Arg | missense_variant | MODERATE | 0.363636 | 0 | 11 | 32 | . | . | . | . |
| PM1548-SK1P4 | COG7 | p.Pro67Ala | missense_variant | MODERATE | 0.413043 | 0 | 46 | 71 | . | . | . | . |
| PM1548-SK1P4 | GTF3C1 | p.Glu1761Gly | missense_variant | MODERATE | 0.147368 | 0 | 95 | 99 | . | . | . | . |
| PM1548-SK1P4 | ATXN2L | c.3140-64G>T | intron_variant | MODIFIER | 0.0677966 | 0 | 59 | 124 | . | . | . | . |
| PM1548-SK1P4 | ATP2A1 | p.Gly229Ser | missense_variant | MODERATE | 0.025 | 0 | 120 | 168 | . | . | . | . |
| PM1548-SK1P4 | KIF22 | p.Ile592fs | frameshift_variant | HIGH | 0.6 | 0 | 65 | 127 | . | . | . | . |
| PM1548-SK1P4 | PRR14 | p.Arg96Gln | missense_variant | MODERATE | 0.531532 | 0 | 111 | 271 | . | . | . | . |
| PM1548-SK1P4 | ZNF629 | p.Asp434Asn | missense_variant | MODERATE | 0.420732 | 0 | 164 | 34 | . | . | . | . |
| PM1548-SK1P4 | HSD3B7 | p.Val361Ile | missense_variant | MODERATE | 0.48 | 0 | 75 | 165 | . | . | . | . |
| PM1548-SK1P4 | KAT8 | p.Val438Ala | missense_variant | MODERATE | 0.321839 | 0 | 87 | 67 | . | . | . | . |
| PM1548-SK1P4 | FUS | c.1295+1G>A | splice_donor_variant&intron_variant | HIGH | 0.548673 | 0 | 113 | 234 | true | . | . | . |
| PM1548-SK1P4 | PHKB | p.Tyr52His | missense_variant | MODERATE | 0.0041841 | 0 | 239 | 155 | . | . | . | . |
| PM1548-SK1P4 | LONP2 | p.Gly681Cys | missense_variant | MODERATE | 0.428571 | 0 | 35 | 276 | . | . | . | . |
| PM1548-SK1P4 | N4BP1 | p.Arg640Cys | missense_variant | MODERATE | 0.235294 | 0 | 17 | 94 | . | . | . | . |
| PM1548-SK1P4 | SALL1 | p.Ser1318Asn | missense_variant | MODERATE | 0.44 | 0 | 25 | 197 | . | . | . | . |
| PM1548-SK1P4 | SALL1 | p.His756Asp | missense_variant | MODERATE | 0.259259 | 0 | 27 | 53 | . | . | . | . |
| PM1548-SK1P4 | LPCAT2 | c.1216-7923dup | intron_variant | MODIFIER | 0.333333 | 0 | 36 | 56 | . | . | . | . |
| PM1548-SK1P4 | SLC6A2 | p.Gly591Asp | missense_variant | MODERATE | 0.537234 | 0 | 188 | 342 | . | . | . | . |
| PM1548-SK1P4 | MMP15 | p.Ala603Asp | missense_variant | MODERATE | 0 | 0 | 91 | 388 | . | . | . | . |
| PM1548-SK1P4 | CDH11 | p.Ala166Thr | missense_variant | MODERATE | 0.0891089 | 0 | 101 | 177 | true | . | . | . |
| PM1548-SK1P4 | CDH11 | p.Thr163Ile | missense_variant | MODERATE | 0.118812 | 0 | 101 | 177 | true | . | . | . |
| PM1548-SK1P4 | CDH11 | p.His161His | synonymous_variant | LOW | 0.121212 | 0 | 99 | 175 | true | . | . | . |
| PM1548-SK1P4 | CDH11 | p.Val149Asp | missense_variant | MODERATE | 0.209302 | 0 | 129 | 62 | true | . | . | . |
| PM1548-SK1P4 | CDH11 | p.Val147Asp | missense_variant | MODERATE | 0.253846 | 0 | 130 | 55 | true | . | . | . |
| PM1548-SK1P4 | CDH11 | p.Glu140Gly | missense_variant | MODERATE | 0.242424 | 0 | 66 | 80 | true | . | . | . |
| PM1548-SK1P4 | CDH11 | p.Arg137Gln | missense_variant | MODERATE | 0.287879 | 0 | 66 | 79 | true | . | . | . |
| PM1548-SK1P4 | TK2 | p.Asn114Asn | synonymous_variant | LOW | 0.3 | 0 | 10 | 102 | . | . | . | . |
| PM1548-SK1P4 | B3GNT9 | p.Asp221Asn | missense_variant | MODERATE | 0.102564 | 0 | 39 | 208 | . | . | . | . |
| PM1548-SK1P4 | LRRC29 | p.Gln205His | missense_variant | MODERATE | 0.117188 | 0 | 128 | 214 | . | . | . | . |
| PM1548-SK1P4 | KCTD19 | p.Gly313Arg | missense_variant | MODERATE | 0.458182 | 0 | 275 | 109 | . | . | . | . |
| PM1548-SK1P4 | TPPP3 | p.Ser49Ala | missense_variant | MODERATE | 0 | 0 | 47 | 59 | . | . | . | . |
| PM1548-SK1P4 | CTCF | p.Arg283His | missense_variant | MODERATE | 0.173913 | 0 | 23 | 196 | true | true | true | . |
| PM1548-SK1P4 | CDH3 | p.Asp264Glu | missense_variant | MODERATE | 0 | 0 | 58 | 403 | . | . | . | . |
| PM1548-SK1P4 | WWP2 | p.Arg633Gln | missense_variant | MODERATE | 0.384615 | 0 | 26 | 88 | . | . | . | . |
| PM1548-SK1P4 | PMFBP1 | p.Glu205fs | frameshift_variant | HIGH | 0.243902 | 0 | 41 | 212 | . | . | . | . |
| PM1548-SK1P4 | ZFHX3 | p.Ala1099Ser | missense_variant | MODERATE | 0.0107527 | 0 | 93 | 40 | true | . | true | true |
| PM1548-SK1P4 | RFWD3 | p.Val155Ala | missense_variant | MODERATE | 0.294737 | 0 | 475 | 69 | . | . | . | . |
| PM1548-SK1P4 | PKD1L2 | n.81164209C>T | intragenic_variant | MODIFIER | 0.628205 | 0 | 78 | 77 | . | . | . | . |
| PM1548-SK1P4 | TAF1C | p.Pro707fs | frameshift_variant | HIGH | 0.536723 | 0 | 177 | 205 | . | . | . | . |
| PM1548-SK1P4 | PIEZO1 | p.Pro1061Leu | missense_variant | MODERATE | 0.477778 | 0 | 270 | 171 | . | . | . | . |
| PM1548-SK1P4 | PIEZO1 | p.Met1002Ile | missense_variant | MODERATE | 0.407407 | 0 | 27 | 33 | . | . | . | . |
| PM1548-SK1P4 | GALNS | p.Thr109Thr | synonymous_variant | LOW | 0.336283 | 0 | 113 | 187 | . | . | . | . |
| PM1548-SK1P4 | ANKRD11 | p.Asp2130Asn | missense_variant | MODERATE | 0 | 0 | 30 | 42 | . | . | . | . |
| PM1548-SK1P4 | GEMIN4 | p.Arg229Gln | missense_variant | MODERATE | 0.157895 | 0 | 19 | 149 | . | . | . | . |
| PM1548-SK1P4 | RNMTL1 | p.Met143fs | frameshift_variant | HIGH | 0.235772 | 0 | 123 | 36 | . | . | . | . |
| PM1548-SK1P4 | PRPF8 | p.Arg1057Gln | missense_variant | MODERATE | 0.540948 | 0 | 464 | 146 | . | . | . | . |
| PM1548-SK1P4 | HIC1 | p.Pro693Leu | missense_variant | MODERATE | 0 | 0.0126582 | 39 | 79 | . | . | . | . |
| PM1548-SK1P4 | SMG6 | p.Arg603His | missense_variant | MODERATE | 0.13253 | 0.00434783 | 166 | 230 | . | . | . | . |
| PM1548-SK1P4 | CTNS | p.Asp74Asn | missense_variant | MODERATE | 0.594203 | 0.00314465 | 69 | 318 | . | . | . | . |
| PM1548-SK1P4 | ITGAE | p.Arg113Gln | missense_variant | MODERATE | 0.40678 | 0 | 118 | 94 | . | . | . | . |
| PM1548-SK1P4 | ZZEF1 | p.Arg194Trp | missense_variant | MODERATE | 0.333333 | 0 | 12 | 95 | . | . | . | . |
| PM1548-SK1P4 | KIF1C | p.Ser511Phe | missense_variant | MODERATE | 0.487179 | 0 | 39 | 140 | . | . | . | . |
| PM1548-SK1P4 | USP6 | p.Pro181Leu | missense_variant&splice_region_variant | MODERATE | 0.68 | 0 | 25 | 73 | true | . | . | . |
| PM1548-SK1P4 | DVL2 | p.Arg442Cys | missense_variant | MODERATE | 0.483283 | 0 | 329 | 34 | . | . | . | . |
| PM1548-SK1P4 | EIF5A | p.Thr168Met | missense_variant | MODERATE | 0.375 | 0 | 160 | 86 | . | . | true | . |
| PM1548-SK1P4 | DNAH2 | p.Gly4241fs | frameshift_variant | HIGH | 0.545455 | 0 | 88 | 74 | . | . | . | . |
| PM1548-SK1P4 | ALOX12B | p.Ala355Thr | missense_variant | MODERATE | 0.440355 | 0 | 788 | 295 | . | . | . | . |
| PM1548-SK1P4 | GLP2R | p.Ala515Thr | missense_variant | MODERATE | 0.4 | 0.00983607 | 20 | 305 | . | . | . | . |
| PM1548-SK1P4 | MYH13 | p.Glu984Lys | missense_variant | MODERATE | 0.38961 | 0 | 154 | 72 | . | . | . | . |
| PM1548-SK1P4 | MYH8 | p.Gly186Arg | missense_variant | MODERATE | 0.328125 | 0 | 64 | 81 | . | . | . | . |
| PM1548-SK1P4 | MYH4 | p.Tyr424Cys | missense_variant | MODERATE | 0.510638 | 0 | 94 | 298 | . | . | . | . |

| | | | | | | | | | | | | |
|---|---|---|---|---|---|---|---|---|---|---|---|---|
| PM1548-SK1P4 | DNAH9 | p.Arg1542Met | missense_variant | MODERATE | 0 | 0 | 71 | 342 | . | . | . | . |
| PM1548-SK1P4 | DNAH9 | p.Ala2514Thr | missense_variant | MODERATE | 0.214815 | 0 | 135 | 101 | . | . | . | . |
| PM1548-SK1P4 | ARHGAP44 | p.Ala495Thr | missense_variant | MODERATE | 0.333333 | 0 | 6 | 53 | . | . | . | . |
| PM1548-SK1P4 | FLCN | p.His429fs | frameshift_variant | HIGH | 0.0178571 | 0 | 224 | 269 | true | . | . | . |
| PM1548-SK1P4 | FLCN | p.Thr174Ala | missense_variant | MODERATE | 0.0137931 | 0 | 145 | 227 | true | . | . | . |
| PM1548-SK1P4 | TOM1L2 | p.Met482fs | frameshift_variant | HIGH | 0.234848 | 0 | 132 | 63 | . | . | . | . |
| PM1548-SK1P4 | KIAA0100 | p.Arg689* | stop_gained | HIGH | 0.277778 | 0 | 180 | 219 | . | . | . | . |
| PM1548-SK1P4 | SUPT6H | p.Ile1356Leu | missense_variant | MODERATE | 0.416667 | 0.00465116 | 24 | 215 | . | . | . | . |
| PM1548-SK1P4 | PHF12 | p.Leu393fs | frameshift_variant | HIGH | 0.428571 | 0 | 196 | 60 | . | . | . | . |
| PM1548-SK1P4 | SEZ6 | p.Gly660Asp | missense_variant | MODERATE | 0.636364 | 0 | 22 | 83 | . | . | . | . |
| PM1548-SK1P4 | ANKRD13B | p.Arg233Gln | missense_variant | MODERATE | 0 | 0 | 122 | 98 | . | . | . | . |
| PM1548-SK1P4 | FNDC8 | p.Asn57Ser | missense_variant | MODERATE | 0.5 | 0 | 4 | 150 | . | . | . | . |
| PM1548-SK1P4 | NLE1 | p.Val10Ala | missense_variant | MODERATE | 0.319444 | 0 | 144 | 97 | . | . | . | . |
| PM1548-SK1P4 | SRCIN1 | p.Leu460Phe | missense_variant | MODERATE | 0.363636 | 0 | 11 | 53 | . | . | . | . |
| PM1548-SK1P4 | PCGF2 | p.Asp157Asn | missense_variant | MODERATE | 0.3125 | 0 | 32 | 53 | . | . | . | . |
| PM1548-SK1P4 | STAC2 | p.Cys141Cys | synonymous_variant | LOW | 0.285714 | 0 | 35 | 250 | . | . | . | . |
| PM1548-SK1P4 | GRB7 | p.Arg552Gln | missense_variant | MODERATE | 0 | 0 | 539 | 159 | . | . | . | . |
| PM1548-SK1P4 | NR1D1 | p.Pro593Pro | synonymous_variant | LOW | 0.216828 | 0.00961538 | 309 | 104 | . | . | . | . |
| PM1548-SK1P4 | MSL1 | p.Leu372Ile | missense_variant | MODERATE | 0 | 0 | 12 | 43 | . | . | . | . |
| PM1548-SK1P4 | WIPF2 | p.Ala39Thr | missense_variant | MODERATE | 0.406977 | 0 | 86 | 66 | . | . | . | . |
| PM1548-SK1P4 | KRT25 | p.Ala91Thr | missense_variant | MODERATE | 0.463415 | 0 | 41 | 148 | . | . | . | . |
| PM1548-SK1P4 | KRT15 | p.Arg398Trp | missense_variant | MODERATE | 0.577778 | 0 | 45 | 53 | . | . | . | . |
| PM1548-SK1P4 | KRT16 | p.Gly243Ser | missense_variant | MODERATE | 0.0789474 | 0 | 38 | 130 | . | . | . | . |
| PM1548-SK1P4 | DHX58 | p.Arg224Cys | missense_variant | MODERATE | 0.353659 | 0 | 82 | 87 | . | . | . | . |
| PM1548-SK1P4 | PLEKHH3 | p.Pro318Thr | missense_variant | MODERATE | 0.307692 | 0 | 26 | 121 | . | . | . | . |
| PM1548-SK1P4 | ETV4 | p.Gln13Arg | missense_variant | MODERATE | 0.0226415 | 0 | 265 | 215 | true | . | . | . |
| PM1548-SK1P4 | MPP3 | p.Arg265Gln | missense_variant | MODERATE | 0.354839 | 0 | 31 | 78 | . | . | . | . |
| PM1548-SK1P4 | ATXN7L3 | p.Ile246Phe | missense_variant | MODERATE | 0.473118 | 0 | 93 | 37 | . | . | . | . |
| PM1548-SK1P4 | ITGA2B | p.Asn330Ser | missense_variant | MODERATE | 0 | 0 | 11 | 60 | . | . | . | . |
| PM1548-SK1P4 | ARHGAP27 | p.Leu247fs | frameshift_variant | HIGH | 0.472222 | 0 | 144 | 82 | . | . | . | . |
| PM1548-SK1P4 | WNT3 | p.Ala223Ser | missense_variant | MODERATE | 0.0217391 | 0 | 46 | 125 | . | . | . | . |
| PM1548-SK1P4 | KPNB1 | p.Ile686Leu | missense_variant | MODERATE | 0.028169 | 0 | 71 | 143 | . | . | . | . |
| PM1548-SK1P4 | KPNB1 | p.Ile686Met | missense_variant | MODERATE | 0.028169 | 0 | 71 | 143 | . | . | . | . |
| PM1548-SK1P4 | SP6 | p.Ala138Thr | missense_variant | MODERATE | 0 | 0 | 372 | 221 | . | . | . | . |
| PM1548-SK1P4 | PPP1R9B | p.Glu24Glu | synonymous_variant | LOW | 0.0229885 | 0 | 174 | 79 | . | . | . | . |
| PM1548-SK1P4 | CUEDC1 | p.Arg71Cys | missense_variant | MODERATE | 0.587629 | 0 | 97 | 107 | . | . | . | . |
| PM1548-SK1P4 | LPO | p.Arg82Cys | missense_variant | MODERATE | 0.521739 | 0 | 92 | 404 | . | . | . | . |
| PM1548-SK1P4 | RNF43 | p.Gly659fs | frameshift_variant | HIGH | 0.535565 | 0 | 239 | 76 | true | . | . | true |
| PM1548-SK1P4 | TBX2 | p.Leu460fs | frameshift_variant | HIGH | 0.509091 | 0 | 110 | 331 | . | . | . | . |
| PM1548-SK1P4 | MRC2 | p.Arg490Cys | missense_variant | MODERATE | 0 | 0 | 17 | 51 | . | . | . | . |
| PM1548-SK1P4 | CCDC47 | p.Ser303Pro | missense_variant | MODERATE | 0 | 0 | 20 | 131 | . | . | . | . |
| PM1548-SK1P4 | FTSJ3 | p.Met493Ile | missense_variant | MODERATE | 0.333333 | 0 | 12 | 61 | . | . | . | . |
| PM1548-SK1P4 | ABCA10 | p.Tyr1399His | missense_variant | MODERATE | 0.52 | 0 | 25 | 336 | . | . | . | . |
| PM1548-SK1P4 | SLC16A5 | p.Gly93Cys | missense_variant | MODERATE | 0 | 0 | 11 | 58 | . | . | . | . |
| PM1548-SK1P4 | ITGB4 | p.Pro305Leu | missense_variant | MODERATE | 0.421053 | 0 | 19 | 78 | . | . | . | . |
| PM1548-SK1P4 | ITGB4 | p.Pro723Ser | missense_variant | MODERATE | 0.270471 | 0 | 403 | 167 | . | . | . | . |
| PM1548-SK1P4 | H3F3B | p.Ala8Asp | missense_variant | MODERATE | 0 | 0 | 12 | 88 | true | . | . | . |
| PM1548-SK1P4 | EVPL | p.Arg266Trp | missense_variant | MODERATE | 0.371795 | 0 | 78 | 157 | . | . | . | . |
| PM1548-SK1P4 | RHBDF2 | p.Ala354Thr | missense_variant | MODERATE | 0 | 0 | 76 | 138 | . | . | . | . |
| PM1548-SK1P4 | TNRC6C | p.Arg1160His | missense_variant | MODERATE | 0.277778 | 0 | 54 | 184 | . | . | . | . |
| PM1548-SK1P4 | TMC6 | p.His31His | synonymous_variant | LOW | 0.506494 | 0 | 154 | 487 | . | . | . | . |
| PM1548-SK1P4 | CCDC40 | p.Gly929Ser | missense_variant | MODERATE | 0 | 0 | 4 | 34 | . | . | . | . |
| PM1548-SK1P4 | GAA | p.Gly828Asp | missense_variant&splice_region_variant | MODERATE | 0.264706 | 0 | 68 | 76 | . | . | . | . |
| PM1548-SK1P4 | ENDOV | p.Pro259Leu | missense_variant | MODERATE | 0.507812 | 0 | 128 | 82 | . | . | . | . |
| PM1548-SK1P4 | RPTOR | p.Arg899* | stop_gained | HIGH | 0 | 0 | 10 | 72 | . | . | . | . |
| PM1548-SK1P4 | SIRT7 | p.Thr181Met | missense_variant | MODERATE | 0.146067 | 0 | 89 | 217 | . | . | . | . |
| PM1548-SK1P4 | OGFOD3 | p.Ser127Ser | splice_region_variant&synonymous_variant | LOW | 0.557692 | 0 | 52 | 110 | . | . | . | . |
| PM1548-SK1P4 | TXNDC2 | p.Ile284Leu | missense_variant | MODERATE | 0 | 0 | 52 | 275 | . | . | . | . |
| PM1548-SK1P4 | TXNDC2 | p.Ser285Pro | missense_variant | MODERATE | 0 | 0 | 52 | 275 | . | . | . | . |
| PM1548-SK1P4 | TXNDC2 | p.Pro288Leu | missense_variant | MODERATE | 0.0208333 | 0.00816327 | 48 | 245 | . | . | . | . |
| PM1548-SK1P4 | ANKRD62 | p.Gln745His | missense_variant | MODERATE | 1 | 0 | 6 | 165 | . | . | . | . |
| PM1548-SK1P4 | ZNF521 | p.Cys754Tyr | missense_variant | MODERATE | 0.2 | 0 | 35 | 57 | true | . | . | . |
| PM1548-SK1P4 | KCTD1 | p.Asn124His | missense_variant | MODERATE | 0.147368 | 0 | 95 | 69 | . | . | . | . |
| PM1548-SK1P4 | DSG4 | p.Thr1001Met | missense_variant | MODERATE | 0.975 | 0 | 80 | 104 | . | . | . | . |
| PM1548-SK1P4 | CCDC178 | p.Arg827Met | missense_variant | MODERATE | 0 | 0 | 49 | 192 | . | . | . | . |
| PM1548-SK1P4 | INO80C | p.Gly216Ser | missense_variant | MODERATE | 0 | 0 | 126 | 89 | . | . | . | . |
| PM1548-SK1P4 | SETBP1 | p.Glu244Asp | missense_variant | MODERATE | 0.0169492 | 0 | 59 | 133 | true | true | . | . |
| PM1548-SK1P4 | SETBP1 | p.Ile251Val | missense_variant | MODERATE | 0.010989 | 0 | 91 | 201 | true | true | . | . |
| PM1548-SK1P4 | SETBP1 | p.Ala733Thr | missense_variant | MODERATE | 0 | 0 | 36 | 315 | true | true | . | . |
| PM1548-SK1P4 | SIGLEC15 | p.Ala96Thr | missense_variant | MODERATE | 0 | 0 | 17 | 175 | . | . | . | . |
| PM1548-SK1P4 | SIGLEC15 | p.Arg99Pro | missense_variant | MODERATE | 0 | 0 | 17 | 176 | . | . | . | . |
| PM1548-SK1P4 | SKOR2 | p.Asp736Glu | missense_variant | MODERATE | 0.0536913 | 0 | 149 | 157 | . | . | . | . |
| PM1548-SK1P4 | SKOR2 | p.Gly720Glu | missense_variant | MODERATE | 0.0526316 | 0 | 133 | 134 | . | . | . | . |
| PM1548-SK1P4 | SMAD7 | p.Pro55Ala | missense_variant | MODERATE | 0.153846 | 0 | 13 | 32 | . | . | . | . |
| PM1548-SK1P4 | DSEL | p.Arg1009His | missense_variant | MODERATE | 1 | 0 | 16 | 67 | . | . | . | . |
| PM1548-SK1P4 | FAM69C | p.Gly48Cys | missense_variant | MODERATE | 0 | 0 | 37 | 170 | . | . | . | . |
| PM1548-SK1P4 | TSHZ1 | p.Ala570Val | missense_variant | MODERATE | 0 | 0 | 85 | 279 | . | . | . | . |
| PM1548-SK1P4 | TSHZ1 | p.Tyr775Cys | missense_variant | MODERATE | 0.9 | 0.00806452 | 20 | 124 | . | . | . | . |
| PM1548-SK1P4 | ZNF516 | p.Gly232del | disruptive_inframe_deletion | MODERATE | 0.0833333 | 0 | 24 | 279 | . | . | . | . |
| PM1548-SK1P4 | ZNF516 | p.Pro229_Gly23( | inframe_insertion | MODERATE | 0.0909091 | 0 | 22 | 280 | . | . | . | . |
| PM1548-SK1P4 | GALR1 | p.Leu46Met | missense_variant | MODERATE | 0.092233 | 0 | 206 | 102 | . | . | . | . |
| PM1548-SK1P4 | GALR1 | p.Lys62Lys | synonymous_variant | LOW | 0.099723 | 0 | 361 | 218 | . | . | . | . |
| PM1548-SK1P4 | SALL3 | p.Pro1105fs | frameshift_variant | HIGH | 1 | 0 | 9 | 200 | . | . | . | . |
| PM1548-SK1P4 | PPAP2C | n.291320T>C | intragenic_variant | MODIFIER | 0.627451 | 0 | 51 | 59 | . | . | . | . |
| PM1548-SK1P4 | LPPR3 | p.Gly646Ser | missense_variant | MODERATE | 0.111111 | 0 | 9 | 120 | . | . | . | . |
| PM1548-SK1P4 | CFD | p.Arg199Cys | missense_variant | MODERATE | 0.413793 | 0 | 145 | 193 | . | . | . | . |
| PM1548-SK1P4 | ARID3A | p.Arg503Cys | missense_variant | MODERATE | 0.285714 | 0 | 14 | 124 | . | . | . | . |
| PM1548-SK1P4 | APC2 | p.Asp501Asn | missense_variant | MODERATE | 0.518667 | 0 | 750 | 282 | . | . | . | . |
| PM1548-SK1P4 | PCSK4 | p.Gly57Asp | missense_variant | MODERATE | 0.267574 | 0 | 882 | 340 | . | . | . | . |
| PM1548-SK1P4 | ADAMTSL5 | p.Ala343fs | frameshift_variant | HIGH | 0.232731 | 0 | 941 | 366 | . | . | . | . |
| PM1548-SK1P4 | ADAMTSL5 | p.Pro341Ala | missense_variant | MODERATE | 0.311787 | 0 | 526 | 312 | . | . | . | . |
| PM1548-SK1P4 | MEX3D | p.Gly262Asp | missense_variant | MODERATE | 0.411765 | 0 | 51 | 217 | . | . | . | . |
| PM1548-SK1P4 | TCF3 | p.Gly247fs | frameshift_variant | HIGH | 0 | 0 | 113 | 71 | true | . | . | . |
| PM1548-SK1P4 | C19orf35 | p.Arg325Cys | missense_variant | MODERATE | 0 | 0 | 229 | 53 | . | . | . | . |
| PM1548-SK1P4 | NCLN | p.Thr70Met | missense_variant | MODERATE | 0.561772 | 0 | 429 | 140 | . | . | . | . |
| PM1548-SK1P4 | DOHH | p.Arg286Trp | missense_variant | MODERATE | 0.273381 | 0.0136054 | 139 | 147 | . | . | . | . |
| PM1548-SK1P4 | EEF2 | p.Pro738Ser | missense_variant | MODERATE | 0.482456 | 0 | 228 | 230 | . | . | . | . |
| PM1548-SK1P4 | PIAS4 | p.Tyr265Tyr | synonymous_variant | LOW | 0.563202 | 0 | 712 | 240 | . | . | . | . |
| PM1548-SK1P4 | MAP2K2 | p.Lys51Arg | missense_variant | MODERATE | 0 | 0 | 37 | 75 | true | . | . | . |
| PM1548-SK1P4 | MAP2K2 | p.Lys51Gln | missense_variant | MODERATE | 0 | 0 | 37 | 75 | true | . | . | . |
| PM1548-SK1P4 | CREB3L3 | p.Val140Ala | missense_variant | MODERATE | 0 | 0 | 429 | 205 | . | . | . | . |
| PM1548-SK1P4 | UBXN6 | p.Thr337Met | missense_variant | MODERATE | 0 | 0 | 116 | 93 | . | . | . | . |
| PM1548-SK1P4 | PLIN4 | p.Val592Met | missense_variant | MODERATE | 0.122449 | 0 | 98 | 40 | . | . | . | . |
| PM1548-SK1P4 | PLIN5 | p.Thr456Ile | missense_variant | MODERATE | 0.389831 | 0 | 59 | 37 | . | . | . | . |
| PM1548-SK1P4 | AC005594.3 | n.381G>A | non_coding_exon_variant | MODIFIER | 0.515464 | 0.0037037 | 291 | 270 | . | . | . | . |
| PM1548-SK1P4 | DPP9 | p.Arg125Trp | missense_variant | MODERATE | 0.220708 | 0 | 367 | 40 | . | . | . | . |
| PM1548-SK1P4 | KDM4B | p.Arg618Cys | missense_variant | MODERATE | 0 | 0 | 167 | 167 | . | . | . | . |
| PM1548-SK1P4 | KDM4B | p.Glu1013Lys | missense_variant | MODERATE | 0.305501 | 0 | 1018 | 111 | . | . | . | . |
| PM1548-SK1P4 | DENND1C | p.Thr165fs | frameshift_variant | HIGH | 0.338028 | 0 | 71 | 34 | . | . | . | . |
| PM1548-SK1P4 | PEX11G | p.Arg124Trp | missense_variant | MODERATE | 0.506964 | 0 | 359 | 176 | . | . | . | . |
| PM1548-SK1P4 | CTD-3193O13.9 | p.Thr838Thr | synonymous_variant | LOW | 0.60241 | 0 | 83 | 174 | . | . | . | . |
| PM1548-SK1P4 | COL5A3 | p.Glu1677Lys | missense_variant | MODERATE | 0.217391 | 0 | 23 | 264 | . | . | true | true |
| PM1548-SK1P4 | ZNF653 | p.Asp247Gly | missense_variant | MODERATE | 0.21875 | 0 | 32 | 166 | . | . | . | . |
| PM1548-SK1P4 | ELOF1 | p.Arg41Cys | missense_variant | MODERATE | 0.505929 | 0 | 253 | 108 | . | . | . | . |
| PM1548-SK1P4 | MRI1 | p.His330fs | frameshift_variant | HIGH | 0.546875 | 0 | 64 | 82 | . | . | . | . |
| PM1548-SK1P4 | RFX1 | p.Val600Ile | missense_variant | MODERATE | 0.267442 | 0 | 430 | 345 | . | . | . | . |
| PM1548-SK1P4 | PRKACA | p.Ala219Val | missense_variant | MODERATE | 0.333333 | 0 | 21 | 150 | true | . | . | . |
| PM1548-SK1P4 | TECR | p.Val85Met | missense_variant | MODERATE | 0.4625 | 0.00641026 | 80 | 156 | . | . | . | . |
| PM1548-SK1P4 | EMR2 | p.Pro294Thr | missense_variant | MODERATE | 0.0469799 | 0 | 149 | 117 | . | . | . | . |
| PM1548-SK1P4 | EMR2 | p.Tyr292Ser | missense_variant | MODERATE | 0.0555556 | 0 | 126 | 117 | . | . | . | . |
| PM1548-SK1P4 | KLF2 | p.Ala238fs | frameshift_variant | HIGH | 0.363158 | 0 | 190 | 137 | . | . | . | . |
| PM1548-SK1P4 | NWD1 | p.Arg788Cys | missense_variant | MODERATE | 0.692308 | 0 | 117 | 334 | . | . | . | . |
| PM1548-SK1P4 | BST2 | p.Ile120fs | frameshift_variant | HIGH | 0.363636 | 0 | 11 | 64 | . | . | . | . |
| PM1548-SK1P4 | FCHO1 | p.Ala286Val | missense_variant&splice_region_variant | MODERATE | 0.436709 | 0 | 158 | 272 | . | . | . | . |
| PM1548-SK1P4 | DDX49 | p.Arg310Gln | missense_variant&splice_region_variant | MODERATE | 0.453271 | 0 | 642 | 300 | . | . | . | . |
| PM1548-SK1P4 | SUGP2 | p.Cys490Tyr | missense_variant | MODERATE | 0.434783 | 0 | 23 | 81 | . | . | . | . |
| PM1548-SK1P4 | NR2C2AP | p.Lys28fs | frameshift_variant | HIGH | 0.511111 | 0 | 315 | 183 | . | . | . | . |

| | | | | | | | | | | | | |
|---|---|---|---|---|---|---|---|---|---|---|---|---|
| PM1548-SK1P4 | NCAN | p.Arg256Gln | missense_variant | MODERATE | 0.544643 | 0 | 112 | 124 | . | . | . | . |
| PM1548-SK1P4 | NDUFA13 | p.Trp131Leu | missense_variant | MODERATE | 0 | 0 | 38 | 265 | . | . | . | . |
| PM1548-SK1P4 | ZNF728 | p.Pro380Ser | missense_variant | MODERATE | 0.333333 | 0 | 3 | 32 | . | . | . | . |
| PM1548-SK1P4 | ZNF728 | p.Arg323Pro | missense_variant | MODERATE | 0.0555556 | 0 | 18 | 48 | . | . | . | . |
| PM1548-SK1P4 | ZNF728 | p.Arg323Trp | missense_variant | MODERATE | 0.0555556 | 0 | 18 | 48 | . | . | . | . |
| PM1548-SK1P4 | ZNF507 | p.Asp475Gly | missense_variant | MODERATE | 0.5 | 0 | 4 | 42 | . | . | . | . |
| PM1548-SK1P4 | KIAA0355 | p.Val801Met | missense_variant | MODERATE | 0.0472103 | 0.0060241 | 233 | 166 | . | . | . | . |
| PM1548-SK1P4 | GRAMD1A | p.Gly605Ser | missense_variant | MODERATE | 0.0218978 | 0 | 137 | 68 | . | . | . | . |
| PM1548-SK1P4 | RBM42 | p.Pro117Arg | missense_variant | MODERATE | 0.25 | 0 | 80 | 129 | . | . | . | . |
| PM1548-SK1P4 | ARHGAP33 | p.Gln144fs | frameshift_variant | HIGH | 0.551266 | 0 | 1580 | 430 | . | . | . | . |
| PM1548-SK1P4 | ZNF850 | p.Ala575Thr | missense_variant | MODERATE | 0.109756 | 0 | 82 | 89 | . | . | . | . |
| PM1548-SK1P4 | ZNF850 | p.Ser572Ala | missense_variant | MODERATE | 0.103093 | 0 | 97 | 133 | . | . | . | . |
| PM1548-SK1P4 | ZNF850 | p.Val554Ile | missense_variant | MODERATE | 0.104478 | 0 | 67 | 86 | . | . | . | . |
| PM1548-SK1P4 | DPF1 | p.Cys58Cys | synonymous_variant | LOW | 0.0350877 | 0 | 114 | 394 | . | . | . | . |
| PM1548-SK1P4 | DPF1 | p.Cys50Cys | synonymous_variant | LOW | 0.036036 | 0 | 111 | 391 | . | . | . | . |
| PM1548-SK1P4 | YIF1B | p.Phe39Phe | synonymous_variant | LOW | 0.315068 | 0 | 146 | 480 | . | . | . | . |
| PM1548-SK1P4 | RYR1 | p.Val3218Met | missense_variant | MODERATE | 0.00490196 | 0 | 408 | 114 | . | . | . | . |
| PM1548-SK1P4 | FCGBP | p.Ala5241Val | missense_variant | MODERATE | 0.416667 | 0 | 168 | 284 | . | . | . | . |
| PM1548-SK1P4 | PRX | p.Glu495Gln | missense_variant | MODERATE | 0.288462 | 0 | 52 | 130 | . | true | . | . |
| PM1548-SK1P4 | B3GNT8 | p.Glu333Ala | missense_variant | MODERATE | 0.517442 | 0 | 516 | 219 | . | . | . | . |
| PM1548-SK1P4 | ARHGEF1 | p.Arg137Trp | missense_variant | MODERATE | 0.479452 | 0 | 219 | 130 | . | . | . | . |
| PM1548-SK1P4 | ZNF227 | p.Gly390Ser | missense_variant | MODERATE | 0.454545 | 0 | 11 | 73 | . | . | . | . |
| PM1548-SK1P4 | PVRL2 | p.Pro53Leu | missense_variant | MODERATE | 0.541284 | 0 | 109 | 119 | . | . | . | . |
| PM1548-SK1P4 | MARK4 | p.Thr651Thr | synonymous_variant | LOW | 0.307692 | 0 | 104 | 352 | . | . | . | . |
| PM1548-SK1P4 | KLC3 | p.Arg320Cys | missense_variant | MODERATE | 0.526316 | 0 | 171 | 102 | . | . | . | . |
| PM1548-SK1P4 | PPP1R13L | p.Leu397Pro | missense_variant | MODERATE | 0.0393701 | 0 | 127 | 478 | . | . | . | . |
| PM1548-SK1P4 | GIPR | p.Gly144Ser | missense_variant | MODERATE | 0.214286 | 0 | 14 | 35 | . | . | . | . |
| PM1548-SK1P4 | STRN4 | p.Ala484Val | missense_variant | MODERATE | 0.468657 | 0 | 335 | 141 | . | . | . | . |
| PM1548-SK1P4 | FKRP | p.Ala193Thr | missense_variant | MODERATE | 0 | 0 | 433 | 98 | . | . | . | . |
| PM1548-SK1P4 | SLC1A5 | p.Ala534Thr | missense_variant | MODERATE | 0 | 0 | 15 | 44 | . | . | . | . |
| PM1548-SK1P4 | ZC3H4 | p.Arg894His | missense_variant | MODERATE | 0 | 0 | 450 | 254 | . | . | . | . |
| PM1548-SK1P4 | ZC3H4 | p.Gly668Ser | missense_variant | MODERATE | 0.5 | 0 | 38 | 220 | . | . | . | . |
| PM1548-SK1P4 | ZC3H4 | p.Lys500Arg | missense_variant | MODERATE | 0 | 0 | 47 | 129 | . | . | . | . |
| PM1548-SK1P4 | ZNF541 | p.Arg214His | missense_variant | MODERATE | 0 | 0 | 37 | 53 | . | . | . | . |
| PM1548-SK1P4 | GLTSCR1 | p.Ser1126fs | frameshift_variant | HIGH | 0.463744 | 0 | 593 | 148 | . | . | . | . |
| PM1548-SK1P4 | CRX | p.Ala35Thr | missense_variant&splice_region_variant | MODERATE | 0.436533 | 0 | 323 | 73 | . | . | . | . |
| PM1548-SK1P4 | PLEKHA4 | p.Arg225Trp | missense_variant | MODERATE | 0.589041 | 0 | 73 | 36 | . | . | . | . |
| PM1548-SK1P4 | BAX | p.Glu41fs | frameshift_variant | HIGH | 0.00854701 | 0 | 234 | 76 | . | . | . | . |
| PM1548-SK1P4 | FLT3LG | p.Ser118fs | frameshift_variant | HIGH | 0.52 | 0 | 50 | 82 | . | . | . | . |
| PM1548-SK1P4 | SCAF1 | p.Gly651fs | frameshift_variant | HIGH | 0.473684 | 0 | 114 | 93 | . | . | . | . |
| PM1548-SK1P4 | ASPDH | p.Glu177Lys | missense_variant | MODERATE | 0.315789 | 0 | 76 | 225 | . | . | . | . |
| PM1548-SK1P4 | LRRC4B | p.Gly50fs | frameshift_variant | HIGH | 0.621951 | 0 | 82 | 58 | . | . | . | . |
| PM1548-SK1P4 | ZNF611 | p.Met226Thr | missense_variant | MODERATE | 0 | 0 | 12 | 80 | . | . | . | . |
| PM1548-SK1P4 | CNOT3 | p.Tyr637His | missense_variant | MODERATE | 0.258065 | 0 | 31 | 41 | true | . | . | true |
| PM1548-SK1P4 | LILRB3 | c.355+1720T>A | intron_variant | MODIFIER | 0.305556 | 0 | 72 | 186 | . | . | . | . |
| PM1548-SK1P4 | LILRB3 | c.355+1720T>A | intron_variant | MODIFIER | 0.305556 | 0 | 72 | 187 | . | . | . | . |
| PM1548-SK1P4 | LILRB3 | c.355+1720T>A | intron_variant | MODIFIER | 0.289855 | 0 | 69 | 184 | . | . | . | . |
| PM1548-SK1P4 | LILRB3 | c.355+1720T>A | intron_variant | MODIFIER | 0.289855 | 0 | 69 | 184 | . | . | . | . |
| PM1548-SK1P4 | EPS8L1 | p.Arg608Cys | missense_variant | MODERATE | 0.577778 | 0 | 90 | 167 | . | . | . | . |
| PM1548-SK1P4 | SBK2 | p.Ala115Thr | missense_variant | MODERATE | 0.509804 | 0 | 204 | 186 | . | . | . | . |
| PM1548-SK1P4 | ZNF579 | p.Arg101Cys | missense_variant | MODERATE | 0.475248 | 0 | 202 | 216 | . | . | . | . |
| PM1548-SK1P4 | ZNF865 | p.Leu610Phe | missense_variant | MODERATE | 0.0318627 | 0 | 408 | 212 | . | . | . | . |
| PM1548-SK1P4 | ZNF865 | p.Leu632Leu | synonymous_variant | LOW | 0.062069 | 0 | 145 | 209 | . | . | . | . |
| PM1548-SK1P4 | ZNF71 | p.Gln343Gln | synonymous_variant | LOW | 0.491882 | 0 | 1047 | 331 | . | . | . | . |
| PM1548-SK1P4 | TRIM28 | p.Arg492Cys | missense_variant | MODERATE | 0.442308 | 0 | 52 | 142 | . | . | . | . |
| PM1548-SK1P4 | ACP1 | p.His67Asn | missense_variant | MODERATE | 0 | 0 | 4 | 36 | . | . | . | . |
| PM1548-SK1P4 | PXDN | p.Val850Met | missense_variant | MODERATE | 0 | 0 | 923 | 308 | . | . | . | . |
| PM1548-SK1P4 | ROCK2 | p.Glu1060Gly | missense_variant | MODERATE | 0 | 0 | 20 | 93 | . | . | . | . |
| PM1548-SK1P4 | FAM228B | p.Asp82Gly | missense_variant | MODERATE | 0.419355 | 0 | 31 | 174 | . | . | . | . |
| PM1548-SK1P4 | NCOA1 | p.Asn1244Thr | missense_variant | MODERATE | 0.102041 | 0 | 49 | 206 | true | . | . | . |
| PM1548-SK1P4 | NCOA1 | p.Ile1260Val | missense_variant | MODERATE | 0.133333 | 0 | 75 | 203 | true | . | . | . |
| PM1548-SK1P4 | NCOA1 | p.Ala1274Thr | missense_variant | MODERATE | 0.121951 | 0 | 41 | 84 | true | . | . | . |
| PM1548-SK1P4 | ADCY3 | p.Thr182Met | missense_variant | MODERATE | 0.575893 | 0 | 224 | 84 | . | . | . | . |
| PM1548-SK1P4 | TRIM54 | p.Arg97Gln | missense_variant | MODERATE | 0.472222 | 0 | 36 | 313 | . | . | . | . |
| PM1548-SK1P4 | TRIM54 | p.Glu131Asp | missense_variant | MODERATE | 0.0847458 | 0 | 59 | 142 | . | . | . | . |
| PM1548-SK1P4 | HEATR5B | p.Lys48fs | frameshift_variant | HIGH | 0 | 0 | 113 | 85 | . | . | . | . |
| PM1548-SK1P4 | RMDN2 | p.Arg130His | missense_variant | MODERATE | 0.153846 | 0 | 65 | 121 | . | . | . | . |
| PM1548-SK1P4 | SOS1 | p.Asn15Lys | missense_variant | MODERATE | 0.486874 | 0 | 419 | 93 | . | . | . | . |
| PM1548-SK1P4 | FSHR | p.Arg569Lys | missense_variant | MODERATE | 0.100719 | 0 | 139 | 177 | . | . | . | . |
| PM1548-SK1P4 | FSHR | p.Ser566Arg | missense_variant | MODERATE | 0.0992908 | 0 | 141 | 133 | . | . | . | . |
| PM1548-SK1P4 | FSHR | p.Asn558Asn | synonymous_variant | LOW | 0.0992908 | 0 | 141 | 139 | . | . | . | . |
| PM1548-SK1P4 | FSHR | p.Ile550Thr | missense_variant | MODERATE | 0.0887097 | 0 | 124 | 160 | . | . | . | . |
| PM1548-SK1P4 | CCDC88A | p.Pro1470Ser | missense_variant | MODERATE | 0.538462 | 0 | 104 | 211 | . | . | . | . |
| PM1548-SK1P4 | CCDC85A | p.His295Tyr | missense_variant | MODERATE | 0 | 0 | 2 | 103 | . | . | . | . |
| PM1548-SK1P4 | CCDC85A | p.Pro359Leu | missense_variant | MODERATE | 0 | 0 | 1 | 45 | . | . | . | . |
| PM1548-SK1P4 | FANCL | c.555+2T>C | splice_donor_variant&intron_variant | HIGH | 0.294118 | 0 | 17 | 47 | . | . | . | . |
| PM1548-SK1P4 | OTX1 | p.Arg39Trp | missense_variant | MODERATE | 0.534884 | 0 | 43 | 144 | . | . | . | . |
| PM1548-SK1P4 | AFTPH | p.Asp198fs | frameshift_variant | HIGH | 0 | 0 | 48 | 159 | . | . | . | . |
| PM1548-SK1P4 | NAGK | p.Lys90Gln | missense_variant | MODERATE | 0.107143 | 0 | 112 | 344 | . | . | . | . |
| PM1548-SK1P4 | NAGK | p.Asn100Asp | missense_variant | MODERATE | 0.108108 | 0 | 111 | 271 | . | . | . | . |
| PM1548-SK1P4 | FBXO41 | p.Arg594Gln | missense_variant | MODERATE | 0 | 0 | 17 | 69 | . | . | . | . |
| PM1548-SK1P4 | TET3 | p.Pro1195Leu | missense_variant | MODERATE | 0.727273 | 0 | 22 | 50 | . | . | . | . |
| PM1548-SK1P4 | C2orf81 | p.Ala344Val | missense_variant | MODERATE | 0.269091 | 0 | 275 | 234 | . | . | . | . |
| PM1548-SK1P4 | TCF7L1 | p.His246Pro | missense_variant | MODERATE | 0.116883 | 0.00719424 | 77 | 139 | . | . | . | . |
| PM1548-SK1P4 | KDM3A | p.Glu1141* | stop_gained | HIGH | 0 | 0 | 52 | 166 | . | . | . | . |
| PM1548-SK1P4 | ZNF514 | p.Gly396Glu | missense_variant | MODERATE | 0.585185 | 0 | 135 | 187 | . | . | . | . |
| PM1548-SK1P4 | NCAPH | p.Gln453* | stop_gained&splice_region_variant | HIGH | 0.272727 | 0 | 11 | 53 | . | . | . | . |
| PM1548-SK1P4 | ZAP70 | p.Ala387Val | missense_variant | MODERATE | 0.521739 | 0 | 92 | 293 | . | . | . | . |
| PM1548-SK1P4 | LONRF2 | p.Asp406Gly | missense_variant | MODERATE | 0 | 0 | 18 | 174 | . | . | . | . |
| PM1548-SK1P4 | CHST10 | p.Asn138Asn | synonymous_variant | LOW | 0.527397 | 0 | 584 | 236 | . | . | . | . |
| PM1548-SK1P4 | IL18R1 | p.Pro527fs | frameshift_variant | HIGH | 0.453901 | 0 | 141 | 255 | . | . | . | . |
| PM1548-SK1P4 | FBLN7 | c.670+86C>T | intron_variant | MODIFIER | 0.542601 | 0 | 223 | 109 | . | . | . | . |
| PM1548-SK1P4 | FBLN7 | p.Arg381Cys | missense_variant | MODERATE | 0.268293 | 0 | 41 | 87 | . | . | . | . |
| PM1548-SK1P4 | ZC3H6 | p.Arg99fs | frameshift_variant | HIGH | 0.6 | 0 | 35 | 90 | . | . | . | . |
| PM1548-SK1P4 | POLR1B | p.Cys893Tyr | missense_variant | MODERATE | 0.301887 | 0 | 53 | 125 | . | . | . | . |
| PM1548-SK1P4 | IL36B | c.261+1014delT | intron_variant | MODIFIER | 0.454545 | 0 | 11 | 101 | . | . | . | . |
| PM1548-SK1P4 | TMEM185B | p.Pro341fs | frameshift_variant | HIGH | 0.188406 | 0 | 69 | 232 | . | . | . | . |
| PM1548-SK1P4 | CLASP1 | p.Ala1225Val | missense_variant | MODERATE | 0.225806 | 0 | 31 | 85 | . | . | . | . |
| PM1548-SK1P4 | CYP27C1 | p.Ala103Thr | missense_variant | MODERATE | 0.5625 | 0 | 16 | 179 | . | . | . | . |
| PM1548-SK1P4 | ACVR2A | p.Lys437fs | frameshift_variant | HIGH | 0.87037 | 0 | 54 | 199 | . | true | . | . |
| PM1548-SK1P4 | NEB | p.Phe2843Tyr | missense_variant | MODERATE | 0.428571 | 0 | 49 | 178 | . | . | . | . |
| PM1548-SK1P4 | TBR1 | p.Gly599Ser | missense_variant | MODERATE | 1 | 0 | 66 | 40 | . | . | . | . |
| PM1548-SK1P4 | SCN3A | p.Ala1768Val | missense_variant | MODERATE | 0.5 | 0 | 2 | 51 | . | . | . | . |
| PM1548-SK1P4 | TLK1 | p.Gln18Arg | missense_variant | MODERATE | 0.0243902 | 0 | 82 | 529 | . | . | . | . |
| PM1548-SK1P4 | ITGA6 | p.Val266Leu | missense_variant | MODERATE | 0.047619 | 0 | 42 | 156 | . | . | . | . |
| PM1548-SK1P4 | SP3 | p.Val461Ile | missense_variant | MODERATE | 0.034188 | 0 | 117 | 262 | . | . | . | . |
| PM1548-SK1P4 | EVX2 | p.Gly438Ser | missense_variant | MODERATE | 1 | 0 | 184 | 165 | . | . | . | . |
| PM1548-SK1P4 | EVX2 | p.Asp320Glu | missense_variant | MODERATE | 0 | 0 | 13 | 48 | . | . | . | . |
| PM1548-SK1P4 | CCDC141 | p.Ala1030Val | missense_variant | MODERATE | 0.047619 | 0 | 63 | 69 | . | . | . | . |
| PM1548-SK1P4 | PDE1A | p.Ala371Val | missense_variant | MODERATE | 0.857143 | 0 | 7 | 75 | . | . | . | . |
| PM1548-SK1P4 | FSIP2 | p.Arg829Cys | missense_variant | MODERATE | 0.5 | 0 | 8 | 51 | . | . | . | . |
| PM1548-SK1P4 | DNAH7 | p.Ala3252Glu | missense_variant | MODERATE | 0.107143 | 0 | 84 | 64 | . | . | . | . |
| PM1548-SK1P4 | GTF3C3 | p.Pro248Leu | missense_variant | MODERATE | 0 | 0 | 16 | 201 | . | . | . | . |
| PM1548-SK1P4 | AOX1 | p.Phe1162Ser | missense_variant | MODERATE | 0.2 | 0 | 100 | 36 | . | . | . | . |
| PM1548-SK1P4 | PIKFYVE | p.Leu419Leu | synonymous_variant | LOW | 0 | 0 | 32 | 195 | . | . | . | . |
| PM1548-SK1P4 | MAP2 | p.Gly1784Ser | missense_variant | MODERATE | 0 | 0 | 4 | 93 | . | . | . | . |
| PM1548-SK1P4 | SMARCAL1 | p.Arg499Gln | missense_variant | MODERATE | 0.558824 | 0 | 204 | 137 | . | . | . | . |
| PM1548-SK1P4 | AAMP | p.Glu63Asp | missense_variant | MODERATE | 0.0294118 | 0.00606061 | 68 | 165 | . | . | . | . |
| PM1548-SK1P4 | ZNF142 | p.Ala335Pro | missense_variant | MODERATE | 0 | 0 | 32 | 317 | . | . | . | . |
| PM1548-SK1P4 | CYP27A1 | p.Ser203Leu | missense_variant | MODERATE | 0.473684 | 0 | 19 | 39 | . | . | . | . |
| PM1548-SK1P4 | WNT6 | p.Asp283Val | missense_variant | MODERATE | 0.48 | 0 | 400 | 120 | . | . | . | . |
| PM1548-SK1P4 | FAM134A | p.Asp391Val | missense_variant | MODERATE | 0.611111 | 0 | 90 | 285 | . | . | . | . |
| PM1548-SK1P4 | DNPEP | p.Arg191Gln | missense_variant | MODERATE | 0 | 0 | 36 | 296 | . | . | . | . |
| PM1548-SK1P4 | CHPF | p.Arg721Trp | missense_variant | MODERATE | 0.196281 | 0 | 484 | 74 | . | . | . | . |
| PM1548-SK1P4 | WDFY1 | p.Ala75Asp | missense_variant | MODERATE | 0 | 0 | 36 | 181 | . | . | . | . |
| PM1548-SK1P4 | SERPINE2 | p.Met296Val | missense_variant | MODERATE | 0.583333 | 0 | 12 | 111 | . | . | . | . |
| PM1548-SK1P4 | B3GNT7 | p.Gly302Asp | missense_variant | MODERATE | 0.363636 | 0 | 11 | 60 | . | . | . | . |

| | | | | | | | | | | | | |
|---|---|---|---|---|---|---|---|---|---|---|---|---|
| PM1548-SK1P4 | DGKD | p.Arg935Trp | missense_variant | MODERATE | 0 | 0 | 30 | 67 | . | . | . | . |
| PM1548-SK1P4 | ACKR3 | p.Val42Ala | missense_variant | MODERATE | 0.634921 | 0 | 63 | 137 | true | . | . | . |
| PM1548-SK1P4 | COL6A3 | p.Gly1714Glu | missense_variant | MODERATE | 0.421053 | 0 | 19 | 50 | . | . | . | . |
| PM1548-SK1P4 | COL6A3 | p.Ile1609Asn | missense_variant | MODERATE | 0.62069 | 0 | 29 | 113 | . | . | . | . |
| PM1548-SK1P4 | ING5 | p.Arg106His | missense_variant | MODERATE | 0 | 0 | 76 | 105 | . | . | . | . |
| PM1548-SK1P4 | TBC1D20 | p.Pro71Ser | missense_variant | MODERATE | 0 | 0 | 27 | 49 | . | . | . | . |
| PM1548-SK1P4 | VPS16 | p.Thr818Met | missense_variant | MODERATE | 0.476378 | 0 | 508 | 238 | . | . | . | . |
| PM1548-SK1P4 | ATRN | p.Ala122Thr | missense_variant | MODERATE | 0.0691824 | 0.00645161 | 159 | 155 | . | . | . | . |
| PM1548-SK1P4 | SLC23A2 | p.Ile412fs | frameshift_variant | HIGH | 0.3 | 0 | 10 | 150 | . | . | . | . |
| PM1548-SK1P4 | BMP2 | p.Ala10Val | missense_variant | MODERATE | 0.0576923 | 0 | 104 | 162 | . | . | . | . |
| PM1548-SK1P4 | PLCB1 | p.Arg768Gln | missense_variant | MODERATE | 0 | 0 | 16 | 91 | . | . | . | . |
| PM1548-SK1P4 | JAG1 | p.Gly537Ser | missense_variant | MODERATE | 0 | 0 | 109 | 184 | . | . | . | . |
| PM1548-SK1P4 | CSRP2BP | p.Trp173* | stop_gained | HIGH | 0.333333 | 0 | 12 | 168 | . | . | . | . |
| PM1548-SK1P4 | CRNKL1 | p.Phe352Leu | missense_variant | MODERATE | 0.333333 | 0 | 24 | 71 | . | . | . | . |
| PM1548-SK1P4 | PLAGL2 | p.Arg180His | missense_variant | MODERATE | 0.516484 | 0 | 182 | 145 | . | . | . | . |
| PM1548-SK1P4 | BPIFA2 | p.Arg221His | missense_variant | MODERATE | 0.516129 | 0 | 31 | 61 | . | . | . | . |
| PM1548-SK1P4 | NECAB3 | p.Arg278fs | frameshift_variant | HIGH | 0.333333 | 0 | 15 | 39 | . | . | . | . |
| PM1548-SK1P4 | CEP250 | p.Lys787Glu | missense_variant | MODERATE | 0 | 0 | 39 | 64 | . | . | . | . |
| PM1548-SK1P4 | SOGA1 | p.Pro974Ser | missense_variant&splice_region_variant | MODERATE | 0.271186 | 0 | 59 | 259 | . | . | . | . |
| PM1548-SK1P4 | PLCG1 | p.Ile942Met | missense_variant | MODERATE | 0.0260417 | 0 | 192 | 140 | true | . | . | . |
| PM1548-SK1P4 | ZSWIM3 | p.Arg225Gln | missense_variant | MODERATE | 0.423077 | 0 | 52 | 55 | . | . | . | . |
| PM1548-SK1P4 | NFATC2 | p.Phe158Tyr | missense_variant | MODERATE | 0 | 0 | 107 | 59 | true | . | . | . |
| PM1548-SK1P4 | NFATC2 | p.Gly22fs | frameshift_variant | HIGH | 0.0410959 | 0 | 146 | 146 | true | . | . | . |
| PM1548-SK1P4 | SALL4 | p.Phe391Leu | missense_variant | MODERATE | 0.216981 | 0 | 106 | 33 | . | . | . | . |
| PM1548-SK1P4 | ZFP64 | n.50701484A>G | intragenic_variant | MODIFIER | 0.135135 | 0 | 111 | 142 | . | . | . | . |
| PM1548-SK1P4 | DOK5 | p.Val169Ala | missense_variant | MODERATE | 0.419689 | 0 | 193 | 338 | . | . | . | . |
| PM1548-SK1P4 | DOK5 | p.Ala267Asp | missense_variant | MODERATE | 0.25 | 0 | 296 | 146 | . | . | . | . |
| PM1548-SK1P4 | TFAP2C | p.Ala354Val | missense_variant | MODERATE | 0.487805 | 0 | 41 | 269 | . | . | . | . |
| PM1548-SK1P4 | PMEPA1 | p.Ser209fs | frameshift_variant | HIGH | 0 | 0 | 702 | 249 | . | . | . | . |
| PM1548-SK1P4 | NELFCD | p.Val321Ile | missense_variant | MODERATE | 0.648649 | 0 | 37 | 73 | . | . | . | . |
| PM1548-SK1P4 | ZNF831 | p.Leu434Met | missense_variant | MODERATE | 0.462733 | 0 | 322 | 129 | . | . | . | . |
| PM1548-SK1P4 | HELZ2 | p.Ala639Thr | missense_variant | MODERATE | 0.144131 | 0 | 673 | 139 | . | . | . | . |
| PM1548-SK1P4 | RTEL1 | p.Arg729Cys | missense_variant | MODERATE | 0.56338 | 0 | 142 | 260 | . | . | . | . |
| PM1548-SK1P4 | TNFRSF6B | p.Leu297Phe | missense_variant | MODERATE | 0 | 0 | 414 | 277 | . | . | . | . |
| PM1548-SK1P4 | SLC2A4RG | p.Arg82fs | frameshift_variant | HIGH | 0.3 | 0 | 30 | 201 | . | . | . | . |
| PM1548-SK1P4 | PRPF6 | p.Arg317His | missense_variant | MODERATE | 0.132353 | 0 | 68 | 86 | . | . | . | . |
| PM1548-SK1P4 | LTN1 | p.Gly1705Val | missense_variant | MODERATE | 0.425 | 0 | 40 | 36 | . | . | . | . |
| PM1548-SK1P4 | TIAM1 | p.His1010Tyr | missense_variant | MODERATE | 0.181818 | 0 | 88 | 418 | . | . | . | . |
| PM1548-SK1P4 | URB1 | p.Pro2036Leu | missense_variant | MODERATE | 0.666667 | 0 | 12 | 14 | . | . | . | . |
| PM1548-SK1P4 | PAXBP1 | p.Pro161Thr | missense_variant | MODERATE | 0 | 0 | 10 | 40 | . | . | . | . |
| PM1548-SK1P4 | PAXBP1 | p.Ser105Arg | missense_variant | MODERATE | 0 | 0 | 248 | 158 | . | . | . | . |
| PM1548-SK1P4 | GART | p.Glu297Gln | missense_variant | MODERATE | 0.171053 | 0 | 76 | 160 | . | . | . | . |
| PM1548-SK1P4 | SON | p.Thr790Ser | missense_variant | MODERATE | 0 | 0 | 23 | 177 | . | . | . | . |
| PM1548-SK1P4 | SON | c.6657+163T>C | intron_variant | MODIFIER | 0.416667 | 0 | 60 | 33 | . | . | . | . |
| PM1548-SK1P4 | ITSN1 | p.Met768Thr | missense_variant | MODERATE | 0.153846 | 0 | 13 | 46 | . | . | . | true |
| PM1548-SK1P4 | SLC5A3 | p.Arg252Gln | missense_variant | MODERATE | 0.695652 | 0 | 46 | 44 | . | . | . | . |
| PM1548-SK1P4 | UMODL1 | p.Gly1319Ser | missense_variant | MODERATE | 0.560976 | 0 | 123 | 131 | . | . | . | . |
| PM1548-SK1P4 | CRYAA | p.Ile61Val | missense_variant | MODERATE | 0.437628 | 0 | 489 | 344 | . | . | . | . |
| PM1548-SK1P4 | SLC19A1 | p.Asp488Gly | missense_variant | MODERATE | 0.0760234 | 0 | 171 | 131 | . | . | . | . |
| PM1548-SK1P4 | FTCD | p.Ile187Thr | missense_variant | MODERATE | 0.431373 | 0 | 51 | 175 | . | . | . | . |
| PM1548-SK1P4 | LSS | p.Thr457Met | missense_variant | MODERATE | 0.451791 | 0 | 363 | 295 | . | . | . | . |
| PM1548-SK1P4 | CECR2 | p.Ser425Ser | synonymous_variant | LOW | 0.15873 | 0 | 63 | 131 | . | . | . | . |
| PM1548-SK1P4 | CECR2 | p.Pro428Pro | synonymous_variant | LOW | 0.163934 | 0 | 61 | 130 | . | . | . | . |
| PM1548-SK1P4 | CECR2 | p.Tyr431Tyr | synonymous_variant | LOW | 0.163934 | 0 | 61 | 130 | . | . | . | . |
| PM1548-SK1P4 | ATP6V1E1 | p.Leu128Met | missense_variant | MODERATE | 0.592593 | 0 | 54 | 148 | . | . | . | . |
| PM1548-SK1P4 | CLTCL1 | p.Ala1567Thr | missense_variant | MODERATE | 0.418421 | 0 | 380 | 279 | true | . | . | . |
| PM1548-SK1P4 | SCARF2 | p.Arg335Cys | missense_variant | MODERATE | 0 | 0 | 37 | 100 | . | . | . | . |
| PM1548-SK1P4 | SLC7A4 | p.Ala256Thr | missense_variant | MODERATE | 0.571429 | 0 | 98 | 223 | . | . | . | . |
| PM1548-SK1P4 | IGLL1 | p.Arg60Gln | missense_variant | MODERATE | 0 | 0 | 56 | 81 | . | . | . | . |
| PM1548-SK1P4 | C22orf43 | p.Ser153Gly | missense_variant | MODERATE | 0 | 0 | 19 | 97 | . | . | . | . |
| PM1548-SK1P4 | CABIN1 | p.Pro585Gln | missense_variant | MODERATE | 0 | 0 | 10 | 59 | . | . | . | . |
| PM1548-SK1P4 | ASPHD2 | p.Arg358Trp | missense_variant | MODERATE | 0.371681 | 0 | 113 | 411 | . | . | . | . |
| PM1548-SK1P4 | TFIP11 | p.Arg39* | stop_gained | HIGH | 0.547541 | 0 | 305 | 103 | . | . | . | . |
| PM1548-SK1P4 | RASL10A | p.Arg108Trp | missense_variant | MODERATE | 0 | 0 | 36 | 108 | . | . | . | . |
| PM1548-SK1P4 | AP1B1 | p.Ala655Asp | missense_variant | MODERATE | 0 | 0 | 28 | 31 | . | . | . | . |
| PM1548-SK1P4 | LIMK2 | p.Asn477Ser | missense_variant | MODERATE | 0.0282258 | 0 | 496 | 257 | . | . | . | . |
| PM1548-SK1P4 | SFI1 | p.Thr710Ser | missense_variant | MODERATE | 0.25 | 0 | 40 | 103 | . | . | . | . |
| PM1548-SK1P4 | CSF2RB | p.Gln642fs | frameshift_variant | HIGH | 0.581864 | 0 | 794 | 133 | . | . | . | . |
| PM1548-SK1P4 | C1QTNF6 | p.Met229Thr | missense_variant | MODERATE | 0.457143 | 0 | 175 | 41 | . | . | . | . |
| PM1548-SK1P4 | GRAP2 | p.Val129Ile | missense_variant | MODERATE | 0.524444 | 0 | 225 | 162 | . | . | . | . |
| PM1548-SK1P4 | MKL1 | p.Pro138Leu | missense_variant | MODERATE | 0.391473 | 0 | 258 | 100 | true | . | . | . |
| PM1548-SK1P4 | EP300 | p.Pro2107Thr | missense_variant | MODERATE | 0.0833333 | 0 | 36 | 94 | true | true | true | . |
| PM1548-SK1P4 | ZC3H7B | p.Arg908Cys | missense_variant | MODERATE | 0.648649 | 0 | 74 | 73 | . | . | . | . |
| PM1548-SK1P4 | PARVB | c.211+24725C>A | intron_variant | MODIFIER | 0.474576 | 0 | 177 | 165 | . | . | . | . |
| PM1548-SK1P4 | CELSR1 | p.Val1411Gly | missense_variant | MODERATE | 0.209231 | 0 | 325 | 333 | . | . | . | . |
| PM1548-SK1P4 | CELSR1 | p.Glu1159Lys | missense_variant | MODERATE | 0.318182 | 0 | 44 | 186 | . | . | . | . |
| PM1548-SK1P4 | TUBGCP6 | p.Cys1239Trp | missense_variant | MODERATE | 0.462069 | 0.00947867 | 145 | 211 | . | . | . | . |
| PM1548-SK1P4 | TUBGCP6 | p.Ser1237Pro | missense_variant | MODERATE | 0.197183 | 0 | 142 | 238 | . | . | . | . |
| PM1548-SK1P4 | MIOX | p.Asp78Gly | missense_variant | MODERATE | 0 | 0 | 121 | 102 | . | . | . | . |
| PM1548-SK1P4 | MAPK8IP2 | p.Thr518Thr | synonymous_variant | LOW | 0.3 | 0 | 320 | 166 | . | . | . | . |
| PM1548-SK1P4 | ARSA | c.684+1G>A | splice_donor_variant&intron_variant | HIGH | 0.557143 | 0 | 210 | 132 | . | . | . | . |
| PM1548-SK1P4 | SETD5 | p.Arg1193* | stop_gained | HIGH | 0.553957 | 0 | 139 | 243 | . | . | . | . |
| PM1548-SK1P4 | CPNE9 | p.Ser284Ser | synonymous_variant | LOW | 0.117647 | 0 | 153 | 197 | . | . | . | . |
| PM1548-SK1P4 | CPNE9 | p.Asp473Asp | synonymous_variant | LOW | 0.481297 | 0 | 401 | 85 | . | . | . | . |
| PM1548-SK1P4 | BRPF1 | p.Gln938fs | frameshift_variant | HIGH | 0.3 | 0 | 20 | 52 | . | . | . | true |
| PM1548-SK1P4 | OGG1 | p.Ala85Thr | missense_variant | MODERATE | 0.568627 | 0 | 51 | 334 | . | . | . | . |
| PM1548-SK1P4 | CRELD1 | p.Arg220* | stop_gained | HIGH | 0.365217 | 0 | 690 | 387 | . | . | . | . |
| PM1548-SK1P4 | PRRT3 | p.His679Arg | missense_variant | MODERATE | 0.525714 | 0 | 175 | 166 | . | . | . | . |
| PM1548-SK1P4 | CAND2 | p.Arg489His | missense_variant | MODERATE | 0.468085 | 0 | 94 | 331 | . | . | . | . |
| PM1548-SK1P4 | SLC6A6 | p.Phe392fs | frameshift_variant | HIGH | 0.510917 | 0 | 687 | 115 | . | . | . | . |
| PM1548-SK1P4 | NR2C2 | p.Phe582fs | frameshift_variant | HIGH | 0.367609 | 0 | 389 | 75 | . | . | . | . |
| PM1548-SK1P4 | SH3BP5 | p.Thr119Ala | missense_variant | MODERATE | 0.451827 | 0 | 602 | 345 | . | . | . | . |
| PM1548-SK1P4 | PLCL2 | p.Gln628Arg | missense_variant | MODERATE | 0.130435 | 0 | 46 | 118 | . | . | . | . |
| PM1548-SK1P4 | SLC4A7 | p.Ile1097Val | missense_variant | MODERATE | 0.387755 | 0.00657895 | 49 | 152 | . | . | . | . |
| PM1548-SK1P4 | CLASP2 | p.Cys10Cys | synonymous_variant | LOW | 0.536017 | 0 | 472 | 190 | . | . | . | . |
| PM1548-SK1P4 | SCN10A | p.Val378Leu | missense_variant | MODERATE | 0.0630631 | 0.00809717 | 111 | 247 | . | . | . | . |
| PM1548-SK1P4 | TTC21A | p.Ser332Thr | missense_variant | MODERATE | 0.333333 | 0 | 39 | 40 | . | . | . | . |
| PM1548-SK1P4 | SNRK | p.Pro598Thr | missense_variant | MODERATE | 0 | 0 | 105 | 93 | . | . | . | . |
| PM1548-SK1P4 | SLC6A20 | p.Asn103Lys | missense_variant | MODERATE | 0 | 0 | 338 | 172 | . | . | . | . |
| PM1548-SK1P4 | SLC6A20 | p.Gly66Trp | missense_variant | MODERATE | 0 | 0.00847458 | 54 | 118 | . | . | . | . |
| PM1548-SK1P4 | FYCO1 | p.Lys501del | inframe_deletion | MODERATE | 0.383607 | 0 | 305 | 171 | . | . | . | . |
| PM1548-SK1P4 | ALS2CL | p.Asn384Lys | missense_variant | MODERATE | 0.466667 | 0 | 30 | 77 | . | . | . | . |
| PM1548-SK1P4 | DHX30 | n.47859525delC | intragenic_variant | MODIFIER | 0.41115 | 0 | 287 | 119 | . | . | . | . |
| PM1548-SK1P4 | UCN2 | p.Thr27Pro | missense_variant | MODERATE | 0.106267 | 0 | 367 | 135 | . | . | . | . |
| PM1548-SK1P4 | CELSR3 | p.Pro1425Ser | missense_variant | MODERATE | 0 | 0 | 137 | 95 | . | . | . | . |
| PM1548-SK1P4 | SLC25A20 | p.Ala227Val | missense_variant | MODERATE | 0.458333 | 0 | 24 | 126 | . | . | . | . |
| PM1548-SK1P4 | KLHDC8B | p.Gly38fs | frameshift_variant | HIGH | 0.337079 | 0 | 89 | 198 | . | . | . | . |
| PM1548-SK1P4 | RRP9 | p.Ala342Thr | missense_variant | MODERATE | 0.384328 | 0 | 268 | 161 | . | . | . | . |
| PM1548-SK1P4 | DNAH1 | p.Leu2072Ile | missense_variant | MODERATE | 0.522727 | 0 | 88 | 99 | . | . | . | . |
| PM1548-SK1P4 | FAM208A | p.Ala227Val | missense_variant | MODERATE | 0.222222 | 0 | 18 | 81 | . | . | . | . |
| PM1548-SK1P4 | DNAH12 | p.Pro1035Ala | missense_variant | MODERATE | 0 | 0 | 64 | 189 | . | . | . | . |
| PM1548-SK1P4 | PDE12 | p.Ala293Ser | missense_variant | MODERATE | 0.0909091 | 0 | 44 | 150 | . | . | . | . |
| PM1548-SK1P4 | PDE12 | p.Leu294Phe | missense_variant | MODERATE | 0.0909091 | 0 | 44 | 150 | . | . | . | . |
| PM1548-SK1P4 | ADAMTS9 | p.Pro596His | missense_variant | MODERATE | 0.0384615 | 0 | 26 | 126 | . | . | . | . |
| PM1548-SK1P4 | ADAMTS9 | p.Ser593Gly | missense_variant | MODERATE | 0.037037 | 0 | 27 | 130 | . | . | . | . |
| PM1548-SK1P4 | ROBO1 | p.Tyr1009* | stop_gained | HIGH | 0 | 0 | 35 | 54 | . | . | . | . |
| PM1548-SK1P4 | OR5K4 | p.Val51Ile | missense_variant | MODERATE | 0.2 | 0 | 20 | 442 | . | . | . | . |
| PM1548-SK1P4 | OR5K4 | p.Arg53Pro | missense_variant | MODERATE | 0.2 | 0 | 20 | 441 | . | . | . | . |
| PM1548-SK1P4 | OR5K4 | p.Leu56His | missense_variant | MODERATE | 0.117647 | 0 | 34 | 510 | . | . | . | . |
| PM1548-SK1P4 | DCBLD2 | p.Thr718Met | missense_variant | MODERATE | 0.516746 | 0 | 209 | 81 | . | . | . | . |
| PM1548-SK1P4 | TFG | p.Ser213Pro | missense_variant | MODERATE | 0.487179 | 0 | 78 | 76 | true | . | . | . |
| PM1548-SK1P4 | CBLB | p.Pro823Ser | missense_variant | MODERATE | 0.5 | 0 | 12 | 275 | true | . | . | . |
| PM1548-SK1P4 | BOC | p.Pro593Leu | missense_variant | MODERATE | 0.168067 | 0.00197628 | 119 | 506 | . | . | . | . |
| PM1548-SK1P4 | BOC | p.His601Gln | missense_variant | MODERATE | 0 | 0 | 103 | 255 | . | . | . | . |
| PM1548-SK1P4 | KIAA2018 | p.Asn1016fs | frameshift_variant | HIGH | 0.054878 | 0 | 164 | 36 | . | . | . | . |
| PM1548-SK1P4 | KIAA1407 | p.Glu683Gly | missense_variant | MODERATE | 0.375 | 0 | 16 | 85 | . | . | . | . |
| PM1548-SK1P4 | LSAMP | p.Val277Ile | missense_variant | MODERATE | 0.466667 | 0 | 120 | 91 | . | . | . | . |

| | | | | | | | | | | | | |
|---|---|---|---|---|---|---|---|---|---|---|---|---|
| PM1548-SK1P4 | ARHGAP31 | p.Lys857Glu | missense_variant | MODERATE | 0.5 | 0 | 2 | 33 | . | . | . | . |
| PM1548-SK1P4 | FSTL1 | p.Val13Ala | missense_variant | MODERATE | 0.284615 | 0 | 130 | 223 | . | . | . | . |
| PM1548-SK1P4 | EAF2 | p.Thr112fs | frameshift_variant | HIGH | 0.428571 | 0 | 21 | 119 | . | . | . | . |
| PM1548-SK1P4 | ILDR1 | p.Ala76Val | missense_variant&splice_region_variant | MODERATE | 0.25 | 0 | 36 | 158 | . | . | . | . |
| PM1548-SK1P4 | MYLK | p.Ala1684Thr | missense_variant | MODERATE | 0.306122 | 0 | 196 | 98 | . | . | . | . |
| PM1548-SK1P4 | ALDH1L1 | p.Glu439fs | frameshift_variant | HIGH | 0.595092 | 0 | 163 | 55 | . | . | . | . |
| PM1548-SK1P4 | CHCHD6 | p.Ser220Leu | missense_variant | MODERATE | 0 | 0 | 288 | 164 | . | . | . | . |
| PM1548-SK1P4 | C3orf27 | p.Asp85fs | frameshift_variant | HIGH | 0.374359 | 0 | 195 | 229 | . | . | . | . |
| PM1548-SK1P4 | TMCC1 | p.Arg620Ser | missense_variant | MODERATE | 0 | 0 | 46 | 62 | . | . | . | . |
| PM1548-SK1P4 | CDV3 | p.Gly226Asp | missense_variant | MODERATE | 0 | 0 | 46 | 44 | . | . | . | . |
| PM1548-SK1P4 | CDV3 | p.Asp228Glu | missense_variant | MODERATE | 0 | 0 | 48 | 55 | . | . | . | . |
| PM1548-SK1P4 | AMOTL2 | p.Arg292Cys | missense_variant | MODERATE | 0 | 0 | 120 | 43 | . | . | . | . |
| PM1548-SK1P4 | EPHB1 | p.Ala82fs | frameshift_variant | HIGH | 0.139344 | 0 | 366 | 300 | . | . | . | . |
| PM1548-SK1P4 | SLC35G2 | p.Asn97fs | frameshift_variant | HIGH | 0.166667 | 0 | 6 | 31 | . | . | . | . |
| PM1548-SK1P4 | ESYT3 | p.Val238Met | missense_variant | MODERATE | 0.544601 | 0 | 213 | 78 | . | . | . | . |
| PM1548-SK1P4 | MED12L | p.Ser970Leu | missense_variant | MODERATE | 0 | 0 | 7 | 123 | . | . | . | . |
| PM1548-SK1P4 | DHX36 | p.Met154fs | frameshift_variant | HIGH | 0 | 0 | 17 | 44 | . | . | . | . |
| PM1548-SK1P4 | CCNL1 | p.Ile354Val | missense_variant | MODERATE | 0 | 0 | 18 | 120 | . | . | . | . |
| PM1548-SK1P4 | PPM1L | p.Glu106Glu | synonymous_variant | LOW | 0.235294 | 0 | 17 | 275 | . | . | . | . |
| PM1548-SK1P4 | SLITRK3 | p.Arg759Cys | missense_variant | MODERATE | 0.52 | 0 | 150 | 99 | . | . | . | . |
| PM1548-SK1P4 | ZBBX | p.Ile496fs | frameshift_variant | HIGH | 0.454545 | 0 | 33 | 76 | . | . | . | . |
| PM1548-SK1P4 | FNDC3B | p.Glu687Gly | missense_variant | MODERATE | 0.0909091 | 0 | 22 | 96 | . | . | . | . |
| PM1548-SK1P4 | NCEH1 | p.Ser105fs | frameshift_variant | HIGH | 0.152542 | 0 | 59 | 228 | . | . | . | . |
| PM1548-SK1P4 | KLHL24 | p.Asn555fs | frameshift_variant | HIGH | 0.777778 | 0 | 18 | 55 | . | . | . | . |
| PM1548-SK1P4 | CHRD | p.Arg76Cys | missense_variant | MODERATE | 0.392857 | 0 | 56 | 91 | . | . | . | . |
| PM1548-SK1P4 | TP63 | p.Thr368Ala | missense_variant | MODERATE | 0.145161 | 0 | 62 | 121 | . | . | . | . |
| PM1548-SK1P4 | LEPREL1 | p.Asp204Glu | missense_variant | MODERATE | 0 | 0 | 54 | 177 | . | . | . | . |
| PM1548-SK1P4 | CCDC50 | p.Ala343Ser | missense_variant | MODERATE | 0 | 0 | 5 | 55 | . | . | . | . |
| PM1548-SK1P4 | MUC20 | p.Gly514Arg | missense_variant | MODERATE | 0.178571 | 0 | 112 | 228 | . | . | . | . |
| PM1548-SK1P4 | LRCH3 | p.Leu49Val | missense_variant | MODERATE | 0.0285714 | 0.00369004 | 70 | 271 | . | . | . | . |
| PM1548-SK1P4 | GAK | p.Ala690Val | missense_variant | MODERATE | 0 | 0 | 122 | 111 | . | . | . | . |
| PM1548-SK1P4 | DGKQ | p.Thr299Met | missense_variant | MODERATE | 0.468153 | 0 | 314 | 253 | . | . | . | . |
| PM1548-SK1P4 | SLC26A1 | p.Ala624Thr | missense_variant | MODERATE | 0 | 0 | 86 | 466 | . | . | . | . |
| PM1548-SK1P4 | POLN | p.Tyr686Cys | missense_variant | MODERATE | 0 | 0 | 124 | 37 | . | . | . | . |
| PM1548-SK1P4 | HTT | p.Ala2259Thr | missense_variant&splice_region_variant | MODERATE | 0.5 | 0 | 6 | 52 | . | . | . | . |
| PM1548-SK1P4 | PPP2R2C | p.Gly397fs | frameshift_variant | HIGH | 0 | 0 | 64 | 188 | . | . | . | . |
| PM1548-SK1P4 | SH3TC1 | p.Glu203Gly | missense_variant | MODERATE | 0.377778 | 0.00213675 | 45 | 468 | . | . | . | . |
| PM1548-SK1P4 | GPR78 | p.Val209Ile | missense_variant | MODERATE | 0.552764 | 0 | 199 | 34 | . | . | . | . |
| PM1548-SK1P4 | KCTD8 | p.Leu62Leu | synonymous_variant | LOW | 0.045977 | 0 | 174 | 213 | . | . | . | . |
| PM1548-SK1P4 | YTHDC1 | p.Ser684Asn | missense_variant | MODERATE | 0.5 | 0 | 2 | 59 | . | . | . | . |
| PM1548-SK1P4 | ANKRD17 | p.Ile2241Val | missense_variant | MODERATE | 0 | 0 | 21 | 147 | . | . | . | . |
| PM1548-SK1P4 | PF4 | p.Ala4Thr | missense_variant | MODERATE | 0.117647 | 0 | 68 | 40 | . | . | . | . |
| PM1548-SK1P4 | PARM1 | p.Ser82Pro | missense_variant | MODERATE | 0 | 0 | 46 | 388 | . | . | . | . |
| PM1548-SK1P4 | CDKL2 | p.Pro452Ser | missense_variant | MODERATE | 0 | 0 | 7 | 51 | . | . | . | . |
| PM1548-SK1P4 | SHROOM3 | p.Lys1017del | inframe_deletion | MODERATE | 0.490272 | 0 | 257 | 97 | . | . | . | . |
| PM1548-SK1P4 | BMP3 | p.Lys358fs | frameshift_variant | HIGH | 0.55 | 0 | 20 | 124 | . | . | . | . |
| PM1548-SK1P4 | PRKG2 | p.Val461Phe | missense_variant | MODERATE | 0 | 0 | 8 | 44 | . | . | . | . |
| PM1548-SK1P4 | SEC31A | p.Ile462fs | frameshift_variant | HIGH | 1 | 0 | 36 | 87 | . | . | . | . |
| PM1548-SK1P4 | SEC31A | p.Lys461fs | frameshift_variant | HIGH | 0 | 0 | 36 | 87 | . | . | . | . |
| PM1548-SK1P4 | SEC31A | p.Gln422Pro | missense_variant | MODERATE | 0.192982 | 0 | 57 | 143 | . | . | . | . |
| PM1548-SK1P4 | AGPAT9 | p.Met372fs | frameshift_variant | HIGH | 0.45977 | 0 | 87 | 70 | . | . | . | . |
| PM1548-SK1P4 | NAP1L5 | p.Lys108fs | frameshift_variant | HIGH | 0.503268 | 0 | 153 | 237 | . | . | . | . |
| PM1548-SK1P4 | CASP6 | p.Pro33Leu | missense_variant | MODERATE | 0.03125 | 0.00719424 | 32 | 278 | . | . | . | . |
| PM1548-SK1P4 | PITX2 | c.411+284C>T | intron_variant | MODIFIER | 0.54013 | 0 | 461 | 59 | . | . | . | . |
| PM1548-SK1P4 | ALPK1 | p.Tyr1216fs | frameshift_variant | HIGH | 0 | 0 | 6 | 44 | . | . | . | . |
| PM1548-SK1P4 | PCDH10 | p.Tyr324Cys | missense_variant | MODERATE | 0.411348 | 0 | 141 | 49 | . | . | . | . |
| PM1548-SK1P4 | SETD7 | p.Leu331Ile | missense_variant | MODERATE | 0.416938 | 0 | 307 | 328 | . | . | . | . |
| PM1548-SK1P4 | FREM3 | p.Arg502* | stop_gained | HIGH | 0 | 0 | 7 | 34 | . | . | . | . |
| PM1548-SK1P4 | NEK1 | p.Thr137Asn | missense_variant | MODERATE | 0.0208333 | 0 | 48 | 84 | . | . | . | . |
| PM1548-SK1P4 | AHRR | p.Arg72His | missense_variant | MODERATE | 0.422414 | 0 | 348 | 75 | . | . | . | . |
| PM1548-SK1P4 | EXOC3 | p.Val576Met | missense_variant | MODERATE | 0.727273 | 0 | 22 | 107 | . | . | . | . |
| PM1548-SK1P4 | LPCAT1 | p.Val411Ile | missense_variant | MODERATE | 0.00529101 | 0 | 189 | 136 | . | . | . | . |
| PM1548-SK1P4 | IRX2 | p.Tyr110Cys | missense_variant | MODERATE | 0.503067 | 0 | 326 | 189 | . | . | . | . |
| PM1548-SK1P4 | NSUN2 | p.Ala653Thr | missense_variant&splice_region_variant | MODERATE | 0.215054 | 0 | 93 | 83 | . | . | . | . |
| PM1548-SK1P4 | SRD5A1 | p.Met162Ile | missense_variant | MODERATE | 0.535714 | 0 | 56 | 54 | . | . | . | . |
| PM1548-SK1P4 | SEMA5A | p.Ile679Thr | missense_variant | MODERATE | 0.0273973 | 0 | 219 | 101 | . | . | . | . |
| PM1548-SK1P4 | NIPBL | p.Glu1060Glu | synonymous_variant | LOW | 1 | 0 | 2 | 121 | . | . | . | . |
| PM1548-SK1P4 | CARD6 | p.Gln393Glu | missense_variant | MODERATE | 0.142857 | 0 | 28 | 371 | . | . | . | . |
| PM1548-SK1P4 | CARD6 | p.Gln400His | missense_variant | MODERATE | 0.148148 | 0 | 27 | 367 | . | . | . | . |
| PM1548-SK1P4 | CARD6 | p.Gln402* | stop_gained | HIGH | 0.148148 | 0 | 27 | 367 | . | . | . | . |
| PM1548-SK1P4 | CARD6 | p.Gln402Arg | missense_variant | MODERATE | 0.153846 | 0 | 26 | 337 | . | . | . | . |
| PM1548-SK1P4 | CARD6 | p.Gln402His | missense_variant | MODERATE | 0.153846 | 0 | 26 | 337 | . | . | . | . |
| PM1548-SK1P4 | C6 | p.Ala2Val | missense_variant | MODERATE | 0.42029 | 0 | 69 | 129 | . | . | . | . |
| PM1548-SK1P4 | SNX18 | n.53839150G>A | intragenic_variant | MODIFIER | 0.580645 | 0 | 31 | 117 | . | . | . | . |
| PM1548-SK1P4 | GPX8 | p.Phe125fs | frameshift_variant | HIGH | 0.272727 | 0 | 77 | 228 | . | . | . | . |
| PM1548-SK1P4 | UTP15 | c.169-826A>G | intron_variant | MODIFIER | 0 | 0 | 52 | 55 | . | . | . | . |
| PM1548-SK1P4 | TTC37 | p.Ala1407Thr | missense_variant | MODERATE | 1 | 0 | 3 | 33 | . | . | . | . |
| PM1548-SK1P4 | CHD1 | p.Asn310fs | frameshift_variant | HIGH | 0.942857 | 0 | 35 | 201 | . | . | . | true |
| PM1548-SK1P4 | PAM | p.Arg260Gln | missense_variant | MODERATE | 0 | 0 | 15 | 107 | . | . | . | . |
| PM1548-SK1P4 | GIN1 | p.Leu345Pro | missense_variant | MODERATE | 0.8 | 0 | 5 | 72 | . | . | . | . |
| PM1548-SK1P4 | APC | p.Asp802fs | frameshift_variant&stop_gained | HIGH | 0.481605 | 0 | 299 | 125 | true | true | true | true |
| PM1548-SK1P4 | MCC | p.Arg800Lys | missense_variant | MODERATE | 0.3 | 0 | 10 | 65 | . | . | . | . |
| PM1548-SK1P4 | FTMT | p.Ala90Val | missense_variant | MODERATE | 0.593137 | 0 | 204 | 105 | . | . | . | . |
| PM1548-SK1P4 | FBN2 | p.Leu2853Arg | missense_variant | MODERATE | 0.530303 | 0 | 66 | 91 | . | . | . | . |
| PM1548-SK1P4 | FNIP1 | p.Leu90Met | missense_variant | MODERATE | 0 | 0 | 14 | 76 | . | . | . | . |
| PM1548-SK1P4 | JADE2 | p.Phe360Leu | missense_variant | MODERATE | 0.0621762 | 0 | 193 | 343 | . | . | . | true |
| PM1548-SK1P4 | FBXL21 | n.956C>T | non_coding_exon_variant | MODIFIER | 0 | 0 | 5 | 13 | . | . | . | . |
| PM1548-SK1P4 | KDM3B | p.Arg921Trp | missense_variant | MODERATE | 0 | 0 | 15 | 178 | . | . | . | . |
| PM1548-SK1P4 | PSD2 | p.Arg574His | missense_variant | MODERATE | 0.474074 | 0.0147059 | 270 | 136 | . | . | . | . |
| PM1548-SK1P4 | PCDHA3 | p.Glu558Val | missense_variant | MODERATE | 0.387755 | 0 | 49 | 71 | . | . | . | . |
| PM1548-SK1P4 | PCDHA5 | p.Ala56Thr | missense_variant | MODERATE | 0.215054 | 0 | 93 | 32 | . | . | . | . |
| PM1548-SK1P4 | PCDHA5 | p.Arg64Arg | synonymous_variant | LOW | 0.223404 | 0 | 94 | 38 | . | . | . | . |
| PM1548-SK1P4 | PCDHA5 | p.Gly69Asp | missense_variant | MODERATE | 0.212121 | 0 | 99 | 45 | . | . | . | . |
| PM1548-SK1P4 | PCDHA8 | p.Ala57Thr | missense_variant | MODERATE | 0.0886076 | 0 | 79 | 46 | . | . | . | . |
| PM1548-SK1P4 | PCDHA13 | p.Glu558Val | missense_variant | MODERATE | 0.196078 | 0 | 51 | 106 | . | . | . | . |
| PM1548-SK1P4 | PCDHAC2 | p.Arg543Trp | missense_variant | MODERATE | 0.548387 | 0 | 31 | 153 | . | . | . | . |
| PM1548-SK1P4 | PCDHAC2 | c.2565+37G>A | intron_variant | MODIFIER | 0.507812 | 0 | 128 | 61 | . | . | . | . |
| PM1548-SK1P4 | PCDHGA1 | p.Ser370Gly | missense_variant | MODERATE | 0.246154 | 0 | 65 | 386 | . | . | . | . |
| PM1548-SK1P4 | PCDH1 | p.Ala203Thr | missense_variant | MODERATE | 0.510324 | 0 | 339 | 253 | . | . | . | . |
| PM1548-SK1P4 | PCDH12 | p.Gly10Glu | missense_variant | MODERATE | 0.188034 | 0 | 117 | 44 | . | . | . | . |
| PM1548-SK1P4 | CSF1R | p.Glu955Lys | missense_variant | MODERATE | 0.518519 | 0 | 27 | 65 | . | . | . | . |
| PM1548-SK1P4 | TIMD4 | p.Ala333Val | missense_variant | MODERATE | 0.42 | 0 | 50 | 229 | . | . | . | . |
| PM1548-SK1P4 | ATP10B | p.Arg115Lys | missense_variant | MODERATE | 0.236726 | 0 | 452 | 157 | . | . | . | . |
| PM1548-SK1P4 | UBTD2 | p.Thr160Ile | missense_variant | MODERATE | 0.428571 | 0 | 7 | 33 | . | . | . | . |
| PM1548-SK1P4 | GPRIN1 | p.Pro806fs | frameshift_variant | HIGH | 0.161765 | 0 | 68 | 165 | . | . | . | . |
| PM1548-SK1P4 | GPRIN1 | p.Pro805Thr | missense_variant | MODERATE | 0.164179 | 0 | 67 | 165 | . | . | . | . |
| PM1548-SK1P4 | GPRIN1 | p.Ser803fs | frameshift_variant | HIGH | 0.164179 | 0 | 67 | 165 | . | . | . | . |
| PM1548-SK1P4 | GPRIN1 | p.Glu801Asp | missense_variant | MODERATE | 0.164179 | 0 | 67 | 165 | . | . | . | . |
| PM1548-SK1P4 | UIMC1 | p.Phe165Cys | missense_variant | MODERATE | 0.0689655 | 0 | 58 | 180 | . | . | . | . |
| PM1548-SK1P4 | UIMC1 | p.Ile156Val | missense_variant&splice_region_variant | MODERATE | 0.0352941 | 0 | 85 | 259 | . | . | . | . |
| PM1548-SK1P4 | FGFR4 | p.Pro568fs | frameshift_variant | HIGH | 0.470588 | 0 | 17 | 174 | true | . | . | . |
| PM1548-SK1P4 | FAM193B | p.Pro761Ser | missense_variant | MODERATE | 0.0410959 | 0 | 73 | 84 | . | . | . | . |
| PM1548-SK1P4 | GRM6 | p.Ala93Thr | missense_variant | MODERATE | 0.559809 | 0 | 209 | 91 | . | . | . | . |
| PM1548-SK1P4 | RREB1 | p.Val754Met | missense_variant | MODERATE | 0.280112 | 0 | 357 | 159 | . | . | . | . |
| PM1548-SK1P4 | HIST1H4C | p.Val82Ala | missense_variant | MODERATE | 0.5 | 0 | 8 | 187 | . | . | . | . |
| PM1548-SK1P4 | HIST1H2AM | p.Leu52Met | missense_variant | MODERATE | 0.0377358 | 0 | 53 | 43 | . | . | . | . |
| PM1548-SK1P4 | TRIM27 | p.Ala24Val | missense_variant | MODERATE | 0.013245 | 0 | 151 | 147 | true | . | . | . |
| PM1548-SK1P4 | OR2H1 | p.Ile295Leu | missense_variant | MODERATE | 0.580645 | 0 | 31 | 181 | . | . | . | . |
| PM1548-SK1P4 | HLA-F | p.Leu396fs | frameshift_variant | HIGH | 0.333333 | 0 | 12 | 38 | . | . | . | . |
| PM1548-SK1P4 | HLA-G | p.Arg285Ser | missense_variant | MODERATE | 0 | 0 | 29 | 42 | . | . | . | . |
| PM1548-SK1P4 | TRIM26 | p.Glu241Lys | missense_variant | MODERATE | 0.591837 | 0 | 49 | 200 | . | . | . | . |
| PM1548-SK1P4 | MDC1 | p.Gln1431Arg | missense_variant | MODERATE | 0 | 0 | 89 | 163 | . | . | . | . |
| PM1548-SK1P4 | MUC22 | p.Ile1250Thr | missense_variant | MODERATE | 0.0333333 | 0 | 60 | 195 | . | . | . | . |
| PM1548-SK1P4 | HLA-C | p.Arg243Trp | missense_variant | MODERATE | 0.282609 | 0 | 46 | 41 | . | . | . | . |
| PM1548-SK1P4 | HLA-B | p.Arg243Trp | missense_variant | MODERATE | 0.0701754 | 0.00879765 | 228 | 341 | . | . | true | . |
| PM1548-SK1P4 | VWA7 | p.Thr26fs | frameshift_variant | HIGH | 0.2 | 0 | 245 | 239 | . | . | . | . |
| PM1548-SK1P4 | CFB | p.Asp1161fs | frameshift_variant | HIGH | 0.443038 | 0 | 158 | 133 | . | . | . | . |

| | | | | | | | | | | | | |
|---|---|---|---|---|---|---|---|---|---|---|---|---|
| PM1548-SK1P4 | RGL2 | p.Glu372fs | frameshift_variant | HIGH | 0.148459 | 0 | 357 | 192 | . | . | . | . |
| PM1548-SK1P4 | DNAH8 | p.Tyr1705Cys | missense_variant | MODERATE | 0 | 0 | 23 | 68 | . | . | . | . |
| PM1548-SK1P4 | KLHDC3 | p.Arg67Cys | missense_variant | MODERATE | 0.238095 | 0 | 21 | 263 | . | . | . | . |
| PM1548-SK1P4 | PKHD1 | p.Arg880His | missense_variant | MODERATE | 0.461538 | 0 | 39 | 253 | . | . | . | . |
| PM1548-SK1P4 | DST | p.Thr1051fs | frameshift_variant | HIGH | 0.545455 | 0 | 11 | 228 | . | . | . | . |
| PM1548-SK1P4 | KIAA1586 | p.Arg617Leu | missense_variant | MODERATE | 0 | 0 | 55 | 71 | . | . | . | . |
| PM1548-SK1P4 | ZNF451 | p.Phe193Leu | missense_variant&splice_region_variant | MODERATE | 0.833333 | 0 | 6 | 32 | . | . | . | . |
| PM1548-SK1P4 | COL12A1 | p.Pro2509Thr | missense_variant | MODERATE | 0 | 0 | 72 | 157 | . | . | . | . |
| PM1548-SK1P4 | SENP6 | p.Ser627Gly | missense_variant | MODERATE | 0 | 0 | 52 | 66 | . | . | . | . |
| PM1548-SK1P4 | ME1 | c.78+1G>A | splice_donor_variant&intron_variant | HIGH | 0.435216 | 0 | 301 | 306 | . | . | . | . |
| PM1548-SK1P4 | PRSS35 | p.Arg265* | stop_gained | HIGH | 0 | 0 | 2 | 41 | . | . | . | . |
| PM1548-SK1P4 | ANKRD6 | p.Ala390Val | missense_variant | MODERATE | 0 | 0 | 15 | 31 | . | . | . | . |
| PM1548-SK1P4 | MDN1 | p.Thr2167Lys | missense_variant | MODERATE | 0.258065 | 0 | 31 | 72 | . | . | . | . |
| PM1548-SK1P4 | MMS22L | p.Gly37Glu | missense_variant | MODERATE | 0.475 | 0 | 40 | 237 | . | . | . | . |
| PM1548-SK1P4 | ATG5 | p.Asp149fs | frameshift_variant | HIGH | 0 | 0 | 172 | 95 | . | . | . | . |
| PM1548-SK1P4 | RFPL4B | p.Arg203Cys | missense_variant | MODERATE | 0.380952 | 0 | 21 | 78 | . | . | . | . |
| PM1548-SK1P4 | KPNA5 | p.Phe383Ser | missense_variant | MODERATE | 0.454545 | 0 | 11 | 67 | . | . | . | . |
| PM1548-SK1P4 | FAM184A | p.Glu79Asp | missense_variant | MODERATE | 0.230159 | 0 | 126 | 52 | . | . | . | . |
| PM1548-SK1P4 | TBC1D32 | p.Ile27Asn | missense_variant | MODERATE | 0.357143 | 0 | 14 | 42 | . | . | . | . |
| PM1548-SK1P4 | HSF2 | p.Arg196His | missense_variant | MODERATE | 0.3 | 0 | 10 | 76 | . | . | . | . |
| PM1548-SK1P4 | KIAA0408 | p.Phe313Leu | missense_variant | MODERATE | 0 | 0 | 37 | 109 | . | . | . | . |
| PM1548-SK1P4 | SOGA3 | p.Ala214fs | frameshift_variant | HIGH | 0.304348 | 0 | 46 | 39 | . | . | . | . |
| PM1548-SK1P4 | ENPP1 | c.240+2T>C | splice_donor_variant&intron_variant | HIGH | 0.733333 | 0 | 45 | 63 | . | . | . | . |
| PM1548-SK1P4 | CTGF | p.Ser227Phe | missense_variant | MODERATE | 0.0731707 | 0 | 41 | 36 | . | . | . | . |
| PM1548-SK1P4 | CTGF | p.Ala226Thr | missense_variant | MODERATE | 0.0697674 | 0 | 43 | 46 | . | . | . | . |
| PM1548-SK1P4 | CTGF | p.Ile195Met | missense_variant | MODERATE | 0.0967742 | 0 | 31 | 61 | . | . | . | . |
| PM1548-SK1P4 | ADGB | p.Asp890Val | missense_variant | MODERATE | 0 | 0 | 5 | 47 | . | . | . | . |
| PM1548-SK1P4 | SYNE1 | p.Pro1307Ser | missense_variant | MODERATE | 0.675676 | 0 | 37 | 79 | . | . | . | . |
| PM1548-SK1P4 | ARID1B | p.Pro1578Ser | missense_variant | MODERATE | 0.125 | 0 | 56 | 46 | true | . | . | . |
| PM1548-SK1P4 | ARID1B | p.Ser1617Thr | missense_variant | MODERATE | 0.0630631 | 0 | 111 | 59 | true | . | . | . |
| PM1548-SK1P4 | TULP4 | p.Pro1275Thr | missense_variant | MODERATE | 0.108108 | 0 | 37 | 171 | . | . | . | . |
| PM1548-SK1P4 | TULP4 | p.Asp1298Asn | missense_variant | MODERATE | 0.0571429 | 0.0078125 | 35 | 256 | . | . | . | . |
| PM1548-SK1P4 | TULP4 | p.Asp1298Glu | missense_variant | MODERATE | 0.114286 | 0 | 35 | 255 | . | . | . | . |
| PM1548-SK1P4 | SMOC2 | p.Ala54Thr | missense_variant | MODERATE | 0.702703 | 0 | 74 | 36 | . | . | . | . |
| PM1548-SK1P4 | GPR146 | p.Pro248Gln | missense_variant | MODERATE | 0 | 0 | 53 | 46 | . | . | . | . |
| PM1548-SK1P4 | ELFN1 | p.Arg389Cys | missense_variant | MODERATE | 0.521246 | 0 | 353 | 43 | . | . | . | . |
| PM1548-SK1P4 | SDK1 | p.Pro1727fs | frameshift_variant | HIGH | 0.518519 | 0 | 324 | 269 | . | . | . | . |
| PM1548-SK1P4 | PMS2 | p.Thr277Ala | missense_variant | MODERATE | 0.272727 | 0 | 11 | 38 | true | . | . | . |
| PM1548-SK1P4 | MIOS | p.Leu213His | missense_variant | MODERATE | 0.5 | 0 | 8 | 44 | . | . | . | . |
| PM1548-SK1P4 | ETV1 | p.Asn37Thr | missense_variant | MODERATE | 0.0336134 | 0 | 119 | 72 | true | . | . | . |
| PM1548-SK1P4 | MPP6 | p.Lys306fs | frameshift_variant | HIGH | 0.0169492 | 0 | 236 | 33 | . | . | . | . |
| PM1548-SK1P4 | HERPUD2 | p.Val266Ala | missense_variant | MODERATE | 0.276423 | 0 | 123 | 59 | . | . | . | . |
| PM1548-SK1P4 | POU6F2 | p.Glu562Ala | missense_variant | MODERATE | 0.344615 | 0 | 325 | 225 | . | . | . | . |
| PM1548-SK1P4 | MYO1G | p.Ala779Val | missense_variant | MODERATE | 0.190476 | 0 | 924 | 358 | . | . | . | . |
| PM1548-SK1P4 | VWC2 | p.Ser3Arg | missense_variant | MODERATE | 0.037037 | 0 | 27 | 64 | . | . | . | . |
| PM1548-SK1P4 | GRB10 | p.Ala349Thr | missense_variant | MODERATE | 0.659091 | 0 | 44 | 372 | . | . | . | . |
| PM1548-SK1P4 | WBSCR22 | p.Pro8Gln | missense_variant | MODERATE | 0 | 0 | 8 | 35 | . | . | . | . |
| PM1548-SK1P4 | CLIP2 | p.Arg771Gln | missense_variant | MODERATE | 0.242105 | 0 | 380 | 150 | . | . | . | . |
| PM1548-SK1P4 | TMEM60 | p.Ala78fs | frameshift_variant | HIGH | 0.416667 | 0 | 12 | 31 | . | . | . | . |
| PM1548-SK1P4 | GNAI1 | p.Cys214Phe | missense_variant | MODERATE | 0 | 0 | 42 | 196 | . | . | . | . |
| PM1548-SK1P4 | HGF | p.Val529Ala | missense_variant | MODERATE | 0 | 0 | 125 | 125 | . | true | . | . |
| PM1548-SK1P4 | ABCB1 | p.Tyr1165* | stop_gained | HIGH | 0.409091 | 0 | 22 | 32 | . | . | . | . |
| PM1548-SK1P4 | ABCB1 | c.3490-1G>C | splice_acceptor_variant&intron_variant | HIGH | 0.380952 | 0 | 21 | 32 | . | . | . | . |
| PM1548-SK1P4 | ABCB1 | p.Ser196Ala | missense_variant | MODERATE | 0.189189 | 0 | 37 | 56 | . | . | . | . |
| PM1548-SK1P4 | MTERF | p.Ala221Thr | missense_variant | MODERATE | 0.666667 | 0 | 6 | 212 | . | . | . | . |
| PM1548-SK1P4 | AKAP9 | p.Ile1377Thr | missense_variant | MODERATE | 0.285714 | 0 | 7 | 76 | true | . | . | . |
| PM1548-SK1P4 | ASB4 | p.Thr229Met | missense_variant | MODERATE | 0.5 | 0 | 12 | 168 | . | . | . | . |
| PM1548-SK1P4 | LMTK2 | p.Asp236fs | frameshift_variant | HIGH | 0.547619 | 0 | 42 | 299 | . | . | . | . |
| PM1548-SK1P4 | LMTK2 | p.Leu1130Ser | missense_variant | MODERATE | 0.512821 | 0 | 39 | 72 | . | . | . | . |
| PM1548-SK1P4 | TRRAP | p.Asn1675Thr | missense_variant | MODERATE | 0.0347826 | 0 | 115 | 117 | true | . | . | . |
| PM1548-SK1P4 | TRRAP | p.Arg1978Trp | missense_variant | MODERATE | 0.25641 | 0 | 39 | 221 | true | . | . | . |
| PM1548-SK1P4 | TRRAP | p.Pro2508Ala | missense_variant | MODERATE | 0.136986 | 0 | 73 | 384 | true | . | . | . |
| PM1548-SK1P4 | GIGYF1 | p.Leu580Pro | missense_variant | MODERATE | 0.0918033 | 0 | 305 | 69 | . | . | . | . |
| PM1548-SK1P4 | SLC12A9 | p.Leu559fs | frameshift_variant | HIGH | 0.5 | 0 | 10 | 56 | . | . | . | . |
| PM1548-SK1P4 | MOGAT3 | p.Asn82Lys | missense_variant | MODERATE | 0 | 0 | 115 | 150 | . | . | . | . |
| PM1548-SK1P4 | SH2B2 | p.Ala74Ala | synonymous_variant | LOW | 0.467532 | 0 | 77 | 70 | . | . | . | . |
| PM1548-SK1P4 | LRWD1 | p.Leu87Phe | missense_variant | MODERATE | 0 | 0 | 44 | 78 | . | . | . | . |
| PM1548-SK1P4 | FBXL13 | p.Cys375Ser | missense_variant | MODERATE | 0 | 0 | 14 | 94 | . | . | . | . |
| PM1548-SK1P4 | RELN | p.Arg2363His | missense_variant | MODERATE | 0.446667 | 0 | 150 | 39 | . | . | . | . |
| PM1548-SK1P4 | ATXN7L1 | p.Val629Phe | missense_variant | MODERATE | 0 | 0 | 2 | 193 | . | . | . | . |
| PM1548-SK1P4 | PIK3CG | p.Thr607fs | frameshift_variant | HIGH | 0 | 0 | 38 | 44 | . | true | . | . |
| PM1548-SK1P4 | CBLL1 | p.Val277Ala | missense_variant | MODERATE | 0 | 0 | 55 | 200 | . | . | . | . |
| PM1548-SK1P4 | SLC26A3 | p.Thr479Ala | missense_variant | MODERATE | 0.135593 | 0 | 59 | 195 | . | . | true | . |
| PM1548-SK1P4 | FOXP2 | p.Gly98Val | missense_variant | MODERATE | 0 | 0 | 15 | 125 | . | . | . | . |
| PM1548-SK1P4 | SND1 | p.Ala331Pro | missense_variant | MODERATE | 0.0213523 | 0 | 562 | 236 | true | . | . | . |
| PM1548-SK1P4 | CEP41 | p.Asp152Gly | missense_variant | MODERATE | 0.215827 | 0 | 139 | 235 | . | . | . | . |
| PM1548-SK1P4 | TSGA13 | p.Lys151fs | frameshift_variant | HIGH | 0 | 0 | 13 | 195 | . | . | . | . |
| PM1548-SK1P4 | TTC26 | p.Ala221Thr | missense_variant | MODERATE | 0.257143 | 0 | 35 | 82 | . | . | . | . |
| PM1548-SK1P4 | MKRN1 | c.186-2435G>C | intron_variant | MODIFIER | 0.642857 | 0 | 28 | 136 | . | . | . | . |
| PM1548-SK1P4 | MGAM | p.Asn1603Asp | missense_variant | MODERATE | 0.415385 | 0 | 130 | 85 | . | . | . | . |
| PM1548-SK1P4 | EPHB6 | p.Ala253fs | frameshift_variant | HIGH | 0.548387 | 0 | 31 | 105 | . | true | . | . |
| PM1548-SK1P4 | ZNF775 | p.Gly65fs | frameshift_variant | HIGH | 0 | 0 | 8 | 50 | . | . | . | . |
| PM1548-SK1P4 | FASTK | p.Ile345Asn | missense_variant | MODERATE | 0.168539 | 0 | 89 | 260 | . | . | . | . |
| PM1548-SK1P4 | GALNT11 | p.Leu188Pro | missense_variant | MODERATE | 0.470588 | 0 | 17 | 157 | . | . | . | . |
| PM1548-SK1P4 | TNKS | p.His859Asn | missense_variant | MODERATE | 0 | 0 | 22 | 75 | . | . | . | . |
| PM1548-SK1P4 | RP1L1 | p.Gln1044fs | frameshift_variant | HIGH | 1 | 0 | 328 | 101 | . | . | . | . |
| PM1548-SK1P4 | XKR6 | p.Arg80Lys | missense_variant | MODERATE | 0.0952381 | 0 | 21 | 127 | . | . | . | . |
| PM1548-SK1P4 | PCM1 | p.Met1321Val | missense_variant | MODERATE | 0 | 0 | 6 | 138 | true | . | . | . |
| PM1548-SK1P4 | SORBS3 | p.Thr100Ala | missense_variant | MODERATE | 0.019802 | 0 | 101 | 140 | . | . | . | . |
| PM1548-SK1P4 | LOXL2 | p.Pro40Leu | missense_variant | MODERATE | 0.958333 | 0 | 48 | 224 | . | . | . | . |
| PM1548-SK1P4 | NEFM | p.Leu5Leu | synonymous_variant | LOW | 0.047619 | 0 | 21 | 41 | . | . | . | . |
| PM1548-SK1P4 | NRG1 | p.Gly39Glu | missense_variant | MODERATE | 0.0571429 | 0 | 35 | 55 | true | . | . | . |
| PM1548-SK1P4 | NRG1 | p.Pro44Ser | missense_variant | MODERATE | 0.105263 | 0 | 19 | 58 | true | . | . | . |
| PM1548-SK1P4 | NRG1 | p.Ala74Pro | missense_variant | MODERATE | 0.285714 | 0 | 7 | 35 | true | . | . | . |
| PM1548-SK1P4 | NRG1 | c.1488-469C>G | intron_variant | MODIFIER | 0.117647 | 0 | 34 | 107 | true | . | . | . |
| PM1548-SK1P4 | ADAM2 | p.Arg3His | missense_variant | MODERATE | 0 | 0 | 64 | 64 | . | . | . | . |
| PM1548-SK1P4 | DKK4 | p.Asp195Asn | missense_variant | MODERATE | 0.6 | 0 | 145 | 81 | . | . | . | . |
| PM1548-SK1P4 | MCM4 | p.Thr629fs | frameshift_variant | HIGH | 0.436364 | 0 | 55 | 85 | . | . | . | . |
| PM1548-SK1P4 | PXDNL | p.Ala353Thr | missense_variant | MODERATE | 0.5 | 0 | 18 | 175 | . | . | . | . |
| PM1548-SK1P4 | RP1 | p.Tyr834fs | frameshift_variant | HIGH | 0.153846 | 0 | 13 | 39 | . | . | . | . |
| PM1548-SK1P4 | YTHDF3 | p.Gln323Gln | synonymous_variant | LOW | 0.0952381 | 0 | 105 | 589 | . | . | . | . |
| PM1548-SK1P4 | C8orf46 | p.Thr78Ile | missense_variant | MODERATE | 0 | 0 | 31 | 109 | . | . | . | . |
| PM1548-SK1P4 | ZFHX4 | p.Ala2905Val | missense_variant | MODERATE | 0.392857 | 0 | 28 | 42 | . | . | . | . |
| PM1548-SK1P4 | HEY1 | p.Val187Ala | missense_variant | MODERATE | 0.0184758 | 0 | 433 | 281 | true | . | . | . |
| PM1548-SK1P4 | HEY1 | p.Thr186Ser | missense_variant | MODERATE | 0.0186047 | 0 | 430 | 272 | true | . | . | . |
| PM1548-SK1P4 | VPS13B | p.Asn2064Thr | missense_variant | MODERATE | 0.30303 | 0 | 33 | 75 | . | . | . | . |
| PM1548-SK1P4 | TRPS1 | p.Arg297Cys | missense_variant | MODERATE | 0.251701 | 0 | 147 | 261 | . | . | . | . |
| PM1548-SK1P4 | TRPS1 | p.Asn265Asn | synonymous_variant | LOW | 0 | 0 | 37 | 334 | . | . | . | . |
| PM1548-SK1P4 | EXT1 | p.Val66Phe | missense_variant | MODERATE | 0.05 | 0 | 60 | 88 | true | . | . | . |
| PM1548-SK1P4 | TBC1D31 | p.Ala208Thr | missense_variant | MODERATE | 0.538462 | 0 | 156 | 184 | . | . | . | . |
| PM1548-SK1P4 | ATAD2 | p.Glu119fs | frameshift_variant | HIGH | 0.555556 | 0 | 27 | 106 | . | . | . | . |
| PM1548-SK1P4 | ZNF572 | p.Ser55Pro | missense_variant | MODERATE | 0.482759 | 0 | 116 | 39 | . | . | . | . |
| PM1548-SK1P4 | MYC | p.Gly175Ser | missense_variant | MODERATE | 0 | 0 | 112 | 270 | true | . | . | . |
| PM1548-SK1P4 | ASAP1 | p.Glu919Lys | missense_variant | MODERATE | 0.208333 | 0 | 120 | 187 | . | . | . | . |
| PM1548-SK1P4 | WISP1 | p.Ile175Met | missense_variant | MODERATE | 0.386792 | 0 | 106 | 119 | . | . | . | . |
| PM1548-SK1P4 | KHDRBS3 | p.Tyr316Tyr | splice_region_variant&synonymous_variant | LOW | 0.439024 | 0 | 82 | 229 | . | . | . | . |
| PM1548-SK1P4 | COL22A1 | p.Thr125Met | missense_variant | MODERATE | 0.333333 | 0 | 6 | 59 | . | . | . | . |
| PM1548-SK1P4 | LY6D | p.Ala99Thr | missense_variant | MODERATE | 0.152805 | 0.00956938 | 517 | 209 | . | . | . | . |
| PM1548-SK1P4 | TOP1MT | p.Pro193Leu | missense_variant | MODERATE | 0 | 0.00595238 | 21 | 168 | . | . | . | . |
| PM1548-SK1P4 | ZC3H3 | p.Gly101fs | frameshift_variant | HIGH | 0.26087 | 0 | 46 | 178 | . | . | . | . |
| PM1548-SK1P4 | GSDMD | p.Glu451Ala | missense_variant | MODERATE | 0.501567 | 0 | 319 | 215 | . | . | . | . |
| PM1548-SK1P4 | MAPK15 | p.Pro493Leu | missense_variant | MODERATE | 0.420063 | 0 | 319 | 366 | . | . | . | . |
| PM1548-SK1P4 | SCRIB | p.Cys22Trp | missense_variant | MODERATE | 0.149171 | 0 | 362 | 81 | . | . | . | . |
| PM1548-SK1P4 | PLEC | p.Arg3540Gln | missense_variant | MODERATE | 0 | 0 | 122 | 130 | . | . | . | . |
| PM1548-SK1P4 | PLEC | p.Val648Met | missense_variant | MODERATE | 0 | 0 | 29 | 166 | . | . | . | . |
| PM1548-SK1P4 | CYHR1 | n.145689659dup | intragenic_variant | MODIFIER | 0.0454545 | 0 | 66 | 56 | . | . | . | . |
| PM1548-SK1P4 | ZNF7 | p.Val136Ile | missense_variant | MODERATE | 0 | 0 | 6 | 154 | . | . | . | . |

| | | | | | | | | | | | | |
|---|---|---|---|---|---|---|---|---|---|---|---|---|
| PM1548-SK1P4 | KANK1 | p.Thr192Ser | missense_variant | MODERATE | 0.0810811 | 0 | 37 | 65 | . | . | . | . |
| PM1548-SK1P4 | DMRT3 | p.Leu385Leu | synonymous_variant | LOW | 0 | 0 | 102 | 221 | . | . | . | . |
| PM1548-SK1P4 | RLN2 | p.His8fs | frameshift_variant | HIGH | 0.377551 | 0 | 98 | 225 | . | . | . | . |
| PM1548-SK1P4 | IFNA4 | p.Ala74Thr | missense_variant | MODERATE | 0.357143 | 0 | 14 | 51 | . | . | . | . |
| PM1548-SK1P4 | TAF1L | p.Gly4Ser | missense_variant | MODERATE | 0.333333 | 0 | 6 | 329 | . | . | . | true |
| PM1548-SK1P4 | BAG1 | p.Glu321Lys | missense_variant | MODERATE | 0.459184 | 0 | 196 | 192 | . | . | . | . |
| PM1548-SK1P4 | PRSS3 | p.Ser102Tyr | missense_variant | MODERATE | 0 | 0 | 44 | 102 | . | . | . | . |
| PM1548-SK1P4 | CD72 | p.Ala111Thr | missense_variant | MODERATE | 0.445652 | 0 | 92 | 227 | . | . | . | . |
| PM1548-SK1P4 | RGP1 | p.Arg328Trp | missense_variant | MODERATE | 0.416667 | 0 | 36 | 168 | . | . | . | . |
| PM1548-SK1P4 | ANKRD18A | p.Gly900Ala | missense_variant | MODERATE | 0.3 | 0 | 40 | 119 | . | . | . | . |
| PM1548-SK1P4 | ANKRD18A | p.Ala897Val | missense_variant | MODERATE | 0.387755 | 0 | 49 | 119 | . | . | . | . |
| PM1548-SK1P4 | CNTNAP3 | p.Arg841Ser | missense_variant | MODERATE | 0 | 0 | 6 | 45 | . | . | . | . |
| PM1548-SK1P4 | TMC1 | p.Ala755Pro | missense_variant&splice_region_variant | MODERATE | 0.111111 | 0 | 9 | 45 | . | . | . | . |
| PM1548-SK1P4 | RMI1 | p.Thr325Ala | missense_variant | MODERATE | 0.485714 | 0 | 280 | 482 | . | . | . | . |
| PM1548-SK1P4 | NAA35 | p.*726* | stop_retained_variant | LOW | 0 | 0 | 43 | 360 | . | . | . | . |
| PM1548-SK1P4 | SPATA31E1 | p.Arg436His | missense_variant | MODERATE | 0.6 | 0 | 20 | 212 | . | . | . | . |
| PM1548-SK1P4 | ROR2 | p.Ala307Thr | missense_variant | MODERATE | 0 | 0 | 53 | 69 | . | . | . | . |
| PM1548-SK1P4 | NOL8 | p.Arg430Lys | missense_variant | MODERATE | 0.0967742 | 0 | 93 | 112 | . | . | . | . |
| PM1548-SK1P4 | NOL8 | p.Leu427Gln | missense_variant | MODERATE | 0.130435 | 0 | 92 | 113 | . | . | . | . |
| PM1548-SK1P4 | IPPK | p.Phe34Ser | missense_variant | MODERATE | 0.48 | 0 | 25 | 158 | . | . | . | . |
| PM1548-SK1P4 | SUSD3 | c.89-6671G>T | intron_variant | MODIFIER | 0 | 0 | 7 | 26 | . | . | . | . |
| PM1548-SK1P4 | WNK2 | p.His758fs | frameshift_variant | HIGH | 0.457286 | 0 | 199 | 277 | . | . | . | . |
| PM1548-SK1P4 | C9orf3 | p.Ala207Thr | missense_variant | MODERATE | 0.157895 | 0 | 57 | 105 | . | . | . | . |
| PM1548-SK1P4 | PTCH1 | p.Thr1064Met | missense_variant | MODERATE | 0.211538 | 0 | 52 | 167 | true | . | . | . |
| PM1548-SK1P4 | ZNF510 | p.Arg592Gly | missense_variant | MODERATE | 0 | 0 | 46 | 140 | . | . | . | . |
| PM1548-SK1P4 | CCDC180 | p.Met439Ile | missense_variant | MODERATE | 0.453782 | 0 | 119 | 95 | . | . | . | . |
| PM1548-SK1P4 | GABBR2 | p.Arg622* | stop_gained | HIGH | 0.444444 | 0 | 9 | 80 | . | . | . | . |
| PM1548-SK1P4 | ANKS6 | p.Ala872Val | missense_variant | MODERATE | 0.243902 | 0.00413223 | 82 | 242 | . | . | . | . |
| PM1548-SK1P4 | SLC44A1 | p.Arg437His | missense_variant | MODERATE | 0.545455 | 0 | 11 | 72 | . | . | . | . |
| PM1548-SK1P4 | SVEP1 | p.Ile1787Thr | missense_variant | MODERATE | 0.5 | 0 | 14 | 79 | . | . | . | . |
| PM1548-SK1P4 | TNC | p.Lys1068Asn | missense_variant | MODERATE | 0.4375 | 0 | 80 | 37 | . | . | . | . |
| PM1548-SK1P4 | BRINP1 | p.Arg106His | missense_variant | MODERATE | 0.21875 | 0 | 32 | 219 | . | . | . | . |
| PM1548-SK1P4 | FBXW2 | p.Arg385His | missense_variant | MODERATE | 0 | 0 | 56 | 162 | . | . | . | . |
| PM1548-SK1P4 | FBXW2 | p.Ile377Val | missense_variant | MODERATE | 0 | 0 | 54 | 161 | . | . | . | . |
| PM1548-SK1P4 | DAB2IP | p.His1025fs | frameshift_variant | HIGH | 0.525773 | 0 | 97 | 32 | . | . | . | . |
| PM1548-SK1P4 | LRSAM1 | p.Val385Ile | missense_variant | MODERATE | 0.25 | 0 | 40 | 115 | . | . | . | . |
| PM1548-SK1P4 | SH2D3C | p.Arg821His | missense_variant | MODERATE | 0.52 | 0 | 400 | 143 | . | . | . | . |
| PM1548-SK1P4 | C9orf114 | p.Val77Ile | missense_variant | MODERATE | 0.488764 | 0 | 356 | 277 | . | . | . | . |
| PM1548-SK1P4 | C9orf50 | p.Ala252Val | missense_variant | MODERATE | 0.547826 | 0 | 460 | 411 | . | . | . | . |
| PM1548-SK1P4 | NUP214 | p.Arg887His | missense_variant | MODERATE | 0.547619 | 0 | 42 | 67 | true | . | . | . |
| PM1548-SK1P4 | ADAMTS13 | p.Arg7fs | frameshift_variant | HIGH | 0.650685 | 0 | 292 | 247 | . | . | . | . |
| PM1548-SK1P4 | SARDH | p.Ala819Val | missense_variant | MODERATE | 0.433333 | 0 | 30 | 41 | . | . | . | . |
| PM1548-SK1P4 | SARDH | p.Arg280Trp | missense_variant | MODERATE | 0 | 0 | 22 | 43 | . | . | . | . |
| PM1548-SK1P4 | LHX3 | p.Ala180Thr | missense_variant | MODERATE | 0.525253 | 0 | 297 | 204 | . | . | . | . |
| PM1548-SK1P4 | MAMDC4 | p.Gln963His | missense_variant | MODERATE | 0.45082 | 0 | 122 | 91 | . | . | . | . |
| PM1548-SK1P4 | AL807752.1 | p.Trp56fs | frameshift_variant | HIGH | 0.526316 | 0 | 38 | 54 | . | . | . | . |
| PM1548-SK1P4 | TPRN | p.Asn499Asp | missense_variant | MODERATE | 0.275472 | 0 | 530 | 333 | . | . | . | . |
| PM1548-SK1P4 | NSMF | p.Arg366Cys | missense_variant | MODERATE | 0.527778 | 0 | 108 | 90 | . | . | . | . |
| PM1548-SK1P4 | PNPLA7 | p.Leu178Met | missense_variant | MODERATE | 0 | 0 | 39 | 199 | . | . | . | . |
| PM1548-SK1P4 | HDHD1 | p.Arg149Cys | missense_variant | MODERATE | 1 | 0 | 19 | 104 | . | . | . | . |
| PM1548-SK1P4 | EGFL6 | p.Trp5* | stop_gained | HIGH | 0 | 0 | 36 | 81 | . | . | . | . |
| PM1548-SK1P4 | YY2 | p.Glu5Val | missense_variant | MODERATE | 0.647059 | 0 | 153 | 51 | . | . | . | . |
| PM1548-SK1P4 | TMEM47 | p.Cys104Cys | synonymous_variant | LOW | 0 | 0 | 25 | 132 | . | . | . | . |
| PM1548-SK1P4 | USP27X | p.Arg155His | missense_variant | MODERATE | 0 | 0 | 82 | 112 | . | . | . | . |
| PM1548-SK1P4 | KDM5C | p.Glu1003Asp | missense_variant | MODERATE | 0 | 0 | 59 | 54 | true | true | true | . |
| PM1548-SK1P4 | IQSEC2 | p.Arg612Cys | missense_variant | MODERATE | 0 | 0 | 163 | 42 | . | . | . | . |
| PM1548-SK1P4 | FAM120C | p.Gln36_Gln37de | disruptive_inframe_deletion | MODERATE | 0.958333 | 0 | 48 | 126 | . | . | . | . |
| PM1548-SK1P4 | MAGED2 | p.Glu53Asp | missense_variant | MODERATE | 0 | 0 | 77 | 102 | . | . | . | . |
| PM1548-SK1P4 | TRO | p.Ala1132Thr | missense_variant | MODERATE | 0.0185185 | 0 | 54 | 69 | . | . | . | . |
| PM1548-SK1P4 | DLG3 | p.Leu333Ile | missense_variant | MODERATE | 1 | 0 | 3 | 34 | . | . | . | . |
| PM1548-SK1P4 | ARMCX4 | p.Tyr2123fs | frameshift_variant | HIGH | 0 | 0 | 141 | 78 | . | . | . | . |
| PM1548-SK1P4 | PAK3 | p.Ala466Val | missense_variant | MODERATE | 0.963636 | 0 | 55 | 37 | . | . | . | . |
| PM1548-SK1P4 | ZCCHC16 | p.Arg297Cys | missense_variant | MODERATE | 0.478261 | 0 | 46 | 98 | . | . | . | . |
| PM1548-SK1P4 | AVPR2 | p.Ala60Val | missense_variant | MODERATE | 0 | 0 | 11 | 59 | . | . | . | . |
| PM1548-SK1T1 | B3GALT6 | p.Glu303Lys | missense_variant | MODERATE | 0.32 | 0 | 25 | 86 | . | . | . | . |
| PM1548-SK1T1 | SCNN1D | p.Arg344His | missense_variant | MODERATE | 0.0275862 | 0 | 145 | 274 | . | . | . | . |
| PM1548-SK1T1 | ATAD3B | p.Arg290Cys | missense_variant | MODERATE | 0.0867052 | 0.00687285 | 173 | 291 | . | . | . | . |
| PM1548-SK1T1 | C1orf86 | p.Pro72His | missense_variant | MODERATE | 0.163934 | 0 | 61 | 43 | . | . | . | . |
| PM1548-SK1T1 | PRDM16 | p.Arg759Gln | missense_variant | MODERATE | 0 | 0 | 75 | 40 | true | . | . | . |
| PM1548-SK1T1 | DFFB | p.Tyr290Phe | missense_variant | MODERATE | 0 | 0 | 266 | 203 | . | . | . | . |
| PM1548-SK1T1 | NPHP4 | p.Arg488Gln | missense_variant | MODERATE | 0.102273 | 0 | 88 | 194 | . | . | . | . |
| PM1548-SK1T1 | PLEKHG5 | p.Thr301fs | frameshift_variant | HIGH | 0.0416667 | 0 | 48 | 106 | . | . | . | . |
| PM1548-SK1T1 | ZBTB48 | p.Val626Ala | missense_variant | MODERATE | 0.00636943 | 0 | 157 | 143 | . | . | . | . |
| PM1548-SK1T1 | KLHL21 | p.Cys250fs | frameshift_variant | HIGH | 0.0555556 | 0 | 54 | 136 | . | . | . | . |
| PM1548-SK1T1 | CAMTA1 | p.Ser510Thr | missense_variant | MODERATE | 0 | 0 | 141 | 185 | true | . | . | . |
| PM1548-SK1T1 | PER3 | p.Ala803Ser | missense_variant | MODERATE | 0 | 0 | 234 | 110 | . | . | . | . |
| PM1548-SK1T1 | SLC45A1 | p.Thr364Met | missense_variant | MODERATE | 0.01 | 0 | 100 | 56 | . | . | . | . |
| PM1548-SK1T1 | SLC45A1 | p.Tyr689* | stop_gained | HIGH | 0.151515 | 0 | 132 | 120 | . | . | . | . |
| PM1548-SK1T1 | RERE | p.Lys83fs | frameshift_variant | HIGH | 0.018018 | 0 | 111 | 49 | . | . | . | . |
| PM1548-SK1T1 | CORT | p.Gln133fs | frameshift_variant | HIGH | 0.10084 | 0 | 119 | 48 | . | . | . | . |
| PM1548-SK1T1 | EXOSC10 | p.Met1fs | frameshift_variant&start_lost | HIGH | 0.0641026 | 0 | 78 | 61 | . | . | . | . |
| PM1548-SK1T1 | FHAD1 | p.Ile953fs | frameshift_variant | HIGH | 0.0483871 | 0 | 124 | 108 | . | . | . | . |
| PM1548-SK1T1 | PLEKHM2 | p.Arg848Trp | missense_variant | MODERATE | 0.105263 | 0 | 38 | 42 | . | . | . | . |
| PM1548-SK1T1 | SPEN | p.Asn2002fs | frameshift_variant | HIGH | 0.142241 | 0 | 232 | 173 | true | . | true | true |
| PM1548-SK1T1 | EPHA2 | p.Asn106Ser | missense_variant | MODERATE | 0 | 0 | 152 | 84 | . | . | true | . |
| PM1548-SK1T1 | CROCC | p.Arg1590Cys | missense_variant | MODERATE | 0.027027 | 0 | 222 | 548 | . | . | . | . |
| PM1548-SK1T1 | RCC2 | p.Gly465Arg | missense_variant | MODERATE | 0.0491803 | 0 | 122 | 148 | . | . | . | . |
| PM1548-SK1T1 | ARHGEF10L | p.Val965Met | missense_variant | MODERATE | 0.104762 | 0 | 105 | 230 | . | . | . | . |
| PM1548-SK1T1 | KLHDC7A | p.Ala652Val | missense_variant | MODERATE | 0.037037 | 0 | 54 | 60 | . | . | . | . |
| PM1548-SK1T1 | RNF186 | p.Ile185Thr | missense_variant | MODERATE | 0.144444 | 0 | 270 | 467 | . | . | . | . |
| PM1548-SK1T1 | KIF17 | p.Arg196His | missense_variant | MODERATE | 0.164948 | 0 | 97 | 71 | . | . | . | . |
| PM1548-SK1T1 | EIF4G3 | c.30+5101C>G | intron_variant | MODIFIER | 0 | 0 | 204 | 302 | . | . | . | . |
| PM1548-SK1T1 | HSPG2 | p.Gly2259Asp | missense_variant | MODERATE | 0.175258 | 0 | 97 | 55 | . | . | . | . |
| PM1548-SK1T1 | EPHA8 | p.Ser87Asn | missense_variant | MODERATE | 0 | 0 | 203 | 190 | . | . | . | . |
| PM1548-SK1T1 | AIM1L | n.26671971delA | intragenic_variant | MODIFIER | 0.135922 | 0 | 103 | 54 | . | . | . | . |
| PM1548-SK1T1 | ARID1A | p.Thr1863Ile | missense_variant | MODERATE | 0 | 0 | 110 | 201 | true | true | true | true |
| PM1548-SK1T1 | PTPRU | p.Arg639Trp | missense_variant | MODERATE | 0.114286 | 0 | 35 | 59 | . | . | . | . |
| PM1548-SK1T1 | FABP3 | p.Thr54fs | frameshift_variant | HIGH | 0.11811 | 0 | 127 | 207 | . | . | . | . |
| PM1548-SK1T1 | LCK | p.Arg222Cys | missense_variant | MODERATE | 0.117978 | 0 | 178 | 144 | true | . | . | . |
| PM1548-SK1T1 | KIAA1522 | p.Ala943Thr | missense_variant | MODERATE | 0.0150754 | 0 | 199 | 199 | . | . | . | . |
| PM1548-SK1T1 | ZNF362 | p.Ala105Val | missense_variant | MODERATE | 0.204082 | 0 | 49 | 293 | . | . | . | . |
| PM1548-SK1T1 | LSM10 | p.Ala2Val | missense_variant | MODERATE | 0.117949 | 0 | 195 | 400 | . | . | . | . |
| PM1548-SK1T1 | CSF3R | p.Tyr508Cys | missense_variant | MODERATE | 0 | 0 | 112 | 167 | true | . | . | . |
| PM1548-SK1T1 | EPHA10 | p.Pro330Ser | missense_variant | MODERATE | 0.0375 | 0 | 80 | 71 | . | . | . | . |
| PM1548-SK1T1 | MACF1 | p.Gln6033Pro | missense_variant&splice_region_variant | MODERATE | 0.179775 | 0 | 89 | 48 | . | . | . | . |
| PM1548-SK1T1 | PABPC4 | p.Ser206Asn | missense_variant | MODERATE | 0 | 0 | 250 | 90 | . | . | . | . |
| PM1548-SK1T1 | PABPC4 | p.Glu205Gly | missense_variant | MODERATE | 0 | 0 | 214 | 73 | . | . | . | . |
| PM1548-SK1T1 | ZMPSTE24 | p.Leu362fs | frameshift_variant | HIGH | 0.134454 | 0 | 119 | 53 | . | . | . | . |
| PM1548-SK1T1 | HIVEP3 | p.Pro1037Leu | missense_variant | MODERATE | 0.0280374 | 0 | 107 | 35 | . | . | . | . |
| PM1548-SK1T1 | MED8 | p.Asn295fs | frameshift_variant | HIGH | 0.0638298 | 0 | 94 | 97 | . | . | . | . |
| PM1548-SK1T1 | HYI | p.Ala55Ser | missense_variant | MODERATE | 0.225 | 0 | 40 | 157 | . | . | . | . |
| PM1548-SK1T1 | PTPRF | p.Arg1669Trp | missense_variant | MODERATE | 0.0301205 | 0 | 166 | 115 | . | . | . | . |
| PM1548-SK1T1 | SLC6A9 | p.Arg662Gln | missense_variant | MODERATE | 0.198276 | 0 | 116 | 99 | . | . | . | . |
| PM1548-SK1T1 | PTCH2 | p.Ala967Val | missense_variant | MODERATE | 0.108883 | 0 | 349 | 341 | . | . | . | . |
| PM1548-SK1T1 | PRDX1 | p.His10Tyr | missense_variant | MODERATE | 0 | 0 | 168 | 151 | . | . | . | . |
| PM1548-SK1T1 | MAST2 | p.Arg1086Arg | synonymous_variant | LOW | 0.00502513 | 0 | 199 | 187 | . | . | . | . |
| PM1548-SK1T1 | TAL1 | p.His86Leu | missense_variant | MODERATE | 0 | 0 | 66 | 170 | true | . | . | . |
| PM1548-SK1T1 | LRP8 | p.Asp50Gly | missense_variant | MODERATE | 0.0206186 | 0.00564972 | 194 | 177 | . | . | . | . |
| PM1548-SK1T1 | CDCP2 | p.Pro209fs | frameshift_variant | HIGH | 0.134529 | 0 | 223 | 107 | . | . | . | . |
| PM1548-SK1T1 | MRPL37 | p.Asp142Glu | missense_variant | MODERATE | 0.00735294 | 0 | 136 | 91 | . | . | . | . |
| PM1548-SK1T1 | CACHD1 | p.Glu211Lys | missense_variant | MODERATE | 0 | 0 | 157 | 147 | . | . | . | . |
| PM1548-SK1T1 | JAK1 | p.Val433Ile | missense_variant | MODERATE | 0 | 0 | 271 | 203 | true | . | . | true |
| PM1548-SK1T1 | FPGT | p.Asp273Asn | missense_variant | MODERATE | 0.215054 | 0 | 93 | 168 | . | . | . | . |
| PM1548-SK1T1 | LPHN2 | p.Asn759Ile | missense_variant | MODERATE | 0.046875 | 0 | 128 | 85 | . | . | . | . |
| PM1548-SK1T1 | SYDE2 | p.Gly14fs | frameshift_variant | HIGH | 0.115942 | 0 | 69 | 140 | . | . | . | . |
| PM1548-SK1T1 | CLCA1 | p.Arg676Trp | missense_variant | MODERATE | 0.0288462 | 0 | 104 | 91 | . | . | . | . |
| PM1548-SK1T1 | GBP6 | p.Gly77Asp | missense_variant | MODERATE | 0 | 0 | 104 | 98 | . | . | . | . |

| | | | | | | | | | | | | |
|---|---|---|---|---|---|---|---|---|---|---|---|---|
| PM1548-SK1T1 | GFI1 | p.Arg348Met | missense_variant | MODERATE | 0.030303 | 0 | 264 | 271 | . | . | . | . |
| PM1548-SK1T1 | ABCA4 | p.His423Gln | missense_variant | MODERATE | 0 | 0 | 164 | 257 | . | . | . | . |
| PM1548-SK1T1 | ABCA4 | p.Glu422Asp | missense_variant | MODERATE | 0 | 0.00389105 | 164 | 257 | . | . | . | . |
| PM1548-SK1T1 | LPPR4 | p.Ala343Thr | missense_variant | MODERATE | 0.0923077 | 0 | 130 | 97 | . | . | . | . |
| PM1548-SK1T1 | HIAT1 | p.Ala235Val | missense_variant&splice_region_variant | MODERATE | 0.0108696 | 0 | 92 | 80 | . | . | . | . |
| PM1548-SK1T1 | TRMT13 | p.Glu307fs | frameshift_variant | HIGH | 0.117117 | 0 | 111 | 85 | . | . | . | . |
| PM1548-SK1T1 | AKNAD1 | p.Leu577Met | missense_variant | MODERATE | 0.122563 | 0 | 359 | 457 | . | . | . | . |
| PM1548-SK1T1 | CELSR2 | p.Asp1795Gly | missense_variant | MODERATE | 0 | 0 | 422 | 362 | . | . | . | . |
| PM1548-SK1T1 | LAMTOR5 | p.Thr39Ala | missense_variant | MODERATE | 0.108696 | 0 | 138 | 96 | . | . | . | . |
| PM1548-SK1T1 | HIPK1 | p.Ala314Thr | missense_variant | MODERATE | 0.0764706 | 0 | 170 | 273 | . | . | . | . |
| PM1548-SK1T1 | TRIM33 | p.Val961Met | missense_variant | MODERATE | 0.0833333 | 0 | 48 | 37 | true | . | . | . |
| PM1548-SK1T1 | PDE4DIP | p.Gln2125Lys | missense_variant | MODERATE | 0.0189394 | 0 | 264 | 86 | true | . | . | . |
| PM1548-SK1T1 | PDE4DIP | p.Arg759* | stop_gained | HIGH | 0.0814815 | 0 | 135 | 105 | true | . | . | . |
| PM1548-SK1T1 | BCL9 | p.Pro517fs | frameshift_variant | HIGH | 0.0188679 | 0 | 53 | 144 | true | . | . | . |
| PM1548-SK1T1 | TARS2 | p.Tyr371* | stop_gained | HIGH | 0 | 0 | 79 | 48 | . | . | . | . |
| PM1548-SK1T1 | ADAMTSL4 | p.Gly780fs | frameshift_variant | HIGH | 0.0989011 | 0 | 91 | 112 | . | . | . | . |
| PM1548-SK1T1 | ZNF687 | p.Arg885His | missense_variant | MODERATE | 0.0828402 | 0 | 169 | 314 | . | . | . | . |
| PM1548-SK1T1 | PI4KB | p.Val202Met | missense_variant | MODERATE | 0.136882 | 0 | 263 | 156 | . | . | . | . |
| PM1548-SK1T1 | CELF3 | p.Val279Ile | missense_variant | MODERATE | 0.191489 | 0 | 47 | 53 | . | . | . | . |
| PM1548-SK1T1 | DENND4B | p.Ala154fs | frameshift_variant | HIGH | 0.00806452 | 0 | 124 | 107 | . | . | . | . |
| PM1548-SK1T1 | CREB3L4 | p.Val263fs | frameshift_variant | HIGH | 0.152542 | 0 | 118 | 42 | . | . | . | . |
| PM1548-SK1T1 | EFNA4 | c.470-101delG | intron_variant | MODIFIER | 0.147727 | 0 | 88 | 80 | . | . | . | . |
| PM1548-SK1T1 | ASH1L | p.Asn1907Lys | missense_variant | MODERATE | 0.124088 | 0 | 137 | 55 | . | . | . | . |
| PM1548-SK1T1 | SMG5 | p.Gln846Lys | missense_variant | MODERATE | 0.102564 | 0 | 39 | 128 | . | . | . | . |
| PM1548-SK1T1 | CD1D | p.Asp140Asn | missense_variant | MODERATE | 0.0282258 | 0 | 248 | 198 | . | . | true | . |
| PM1548-SK1T1 | AIM2 | p.Lys342fs | frameshift_variant | HIGH | 0.0714286 | 0 | 84 | 33 | . | . | . | . |
| PM1548-SK1T1 | NIT1 | p.Arg261His | missense_variant | MODERATE | 0.192157 | 0 | 255 | 402 | . | . | . | . |
| PM1548-SK1T1 | ILDR2 | p.Gly453Ala | missense_variant | MODERATE | 0 | 0 | 131 | 328 | . | . | . | . |
| PM1548-SK1T1 | ILDR2 | p.Arg415Trp | missense_variant | MODERATE | 0.161074 | 0 | 149 | 213 | . | . | . | . |
| PM1548-SK1T1 | MYOC | p.Arg91Gln | missense_variant | MODERATE | 0.124031 | 0 | 129 | 240 | . | . | . | . |
| PM1548-SK1T1 | DNM3 | p.Glu7Asp | missense_variant | MODERATE | 0 | 0 | 59 | 77 | . | . | . | . |
| PM1548-SK1T1 | DNM3 | p.Phe20Phe | synonymous_variant | LOW | 0 | 0.0060241 | 134 | 166 | . | . | . | . |
| PM1548-SK1T1 | DNM3 | p.His85Gln | missense_variant | MODERATE | 0 | 0 | 73 | 56 | . | . | . | . |
| PM1548-SK1T1 | ANKRD45 | p.Leu39fs | frameshift_variant | HIGH | 0.0412371 | 0 | 194 | 229 | . | . | . | . |
| PM1548-SK1T1 | ZBTB37 | p.Val284Ile | missense_variant | MODERATE | 0 | 0 | 89 | 152 | . | . | . | . |
| PM1548-SK1T1 | ZBTB37 | p.Asn288Ser | missense_variant | MODERATE | 0 | 0 | 93 | 160 | . | . | . | . |
| PM1548-SK1T1 | PAPPA2 | p.Thr1174fs | frameshift_variant | HIGH | 0.0444444 | 0 | 90 | 36 | . | . | . | . |
| PM1548-SK1T1 | CACNA1E | p.Asn2301Ser | missense_variant | MODERATE | 0 | 0 | 183 | 120 | . | . | . | . |
| PM1548-SK1T1 | CACNA1E | p.Asn2302Asp | missense_variant | MODERATE | 0 | 0 | 183 | 119 | . | . | . | . |
| PM1548-SK1T1 | CACNA1E | p.Asp2310Asp | synonymous_variant | LOW | 0 | 0 | 154 | 90 | . | . | . | . |
| PM1548-SK1T1 | ZNF648 | p.Ser436Tyr | missense_variant | MODERATE | 0 | 0 | 160 | 389 | . | . | . | . |
| PM1548-SK1T1 | ZNF648 | p.Ser436Pro | missense_variant | MODERATE | 0 | 0 | 202 | 439 | . | . | . | . |
| PM1548-SK1T1 | ZNF648 | p.Pro423Ser | missense_variant | MODERATE | 0 | 0.00228311 | 202 | 438 | . | . | . | . |
| PM1548-SK1T1 | C1orf27 | p.Thr381Ser | missense_variant | MODERATE | 0.0379747 | 0 | 79 | 10 | . | . | . | . |
| PM1548-SK1T1 | PLA2G4A | c.1033+2T>C | splice_donor_variant&intron_variant | HIGH | 0.106667 | 0 | 150 | 80 | . | . | . | . |
| PM1548-SK1T1 | ASPM | p.Arg2700Gln | missense_variant | MODERATE | 0.103093 | 0 | 97 | 152 | . | . | . | . |
| PM1548-SK1T1 | DENND1B | c.83-20669G>A | intron_variant | MODIFIER | 0.0123457 | 0 | 81 | 39 | . | . | . | . |
| PM1548-SK1T1 | NR5A2 | p.Gly169Val | missense_variant | MODERATE | 0.163121 | 0 | 141 | 71 | . | . | . | . |
| PM1548-SK1T1 | NAV1 | p.Thr719Ser | missense_variant | MODERATE | 0.0103093 | 0 | 97 | 176 | . | . | . | . |
| PM1548-SK1T1 | NAV1 | p.Thr726Ser | missense_variant | MODERATE | 0 | 0.00930233 | 162 | 215 | . | . | . | . |
| PM1548-SK1T1 | ATP2B4 | p.Asp867Asn | missense_variant&splice_region_variant | MODERATE | 0 | 0 | 82 | 121 | . | . | . | . |
| PM1548-SK1T1 | NUAK2 | p.Arg378His | missense_variant | MODERATE | 0 | 0 | 84 | 162 | . | . | . | . |
| PM1548-SK1T1 | PFKFB2 | p.Glu261Ala | missense_variant | MODERATE | 0.111111 | 0 | 108 | 81 | . | . | . | . |
| PM1548-SK1T1 | CR2 | p.Thr209fs | frameshift_variant | HIGH | 0 | 0 | 96 | 118 | . | . | . | . |
| PM1548-SK1T1 | SMYD2 | p.Asn99fs | frameshift_variant | HIGH | 0.0364583 | 0 | 192 | 181 | . | . | . | . |
| PM1548-SK1T1 | HHIPL2 | p.Gly156Ser | missense_variant | MODERATE | 0.104712 | 0 | 191 | 209 | . | . | . | . |
| PM1548-SK1T1 | MIA3 | p.Arg1545Gln | missense_variant | MODERATE | 0.0933333 | 0 | 150 | 39 | . | . | . | . |
| PM1548-SK1T1 | LIN9 | p.Arg245Trp | missense_variant | MODERATE | 0 | 0 | 85 | 86 | . | . | . | . |
| PM1548-SK1T1 | ITPKB | p.Arg331His | missense_variant | MODERATE | 0.147059 | 0 | 238 | 395 | . | . | true | . |
| PM1548-SK1T1 | ITPKB | p.Leu42Val | missense_variant | MODERATE | 0 | 0 | 22 | 35 | . | . | true | . |
| PM1548-SK1T1 | SNAP47 | p.Phe343Leu | missense_variant | MODERATE | 0 | 0 | 141 | 78 | . | . | . | . |
| PM1548-SK1T1 | IBA57 | p.Ala161Val | missense_variant | MODERATE | 0 | 0 | 195 | 268 | . | . | . | . |
| PM1548-SK1T1 | OBSCN | p.Tyr2035Phe | missense_variant | MODERATE | 0 | 0 | 336 | 508 | . | . | . | . |
| PM1548-SK1T1 | OBSCN | p.Gln2065Lys | missense_variant | MODERATE | 0 | 0 | 196 | 171 | . | . | . | . |
| PM1548-SK1T1 | OBSCN | p.Cys2073Trp | missense_variant | MODERATE | 0 | 0 | 83 | 54 | . | . | . | . |
| PM1548-SK1T1 | ACTA1 | p.Thr231Met | missense_variant | MODERATE | 0.137931 | 0 | 87 | 310 | . | . | . | . |
| PM1548-SK1T1 | GNPAT | p.Val551Met | missense_variant | MODERATE | 0.00581395 | 0 | 172 | 71 | . | . | . | . |
| PM1548-SK1T1 | DISC1 | p.Asp580Gly | missense_variant | MODERATE | 0.0402299 | 0 | 174 | 167 | . | . | . | . |
| PM1548-SK1T1 | GGPS1 | p.Val112Met | missense_variant | MODERATE | 0 | 0 | 272 | 405 | . | . | . | . |
| PM1548-SK1T1 | NID1 | p.Pro918fs | frameshift_variant&splice_region_variant | HIGH | 0.25 | 0 | 64 | 46 | . | . | . | . |
| PM1548-SK1T1 | CEP170 | p.Glu1460Lys | missense_variant | MODERATE | 0.0507614 | 0 | 197 | 32 | . | . | . | . |
| PM1548-SK1T1 | ZNF496 | p.Arg606His | missense_variant | MODERATE | 0.0561798 | 0 | 89 | 143 | . | . | . | . |
| PM1548-SK1T1 | OR2T4 | p.Cys60Ser | missense_variant | MODERATE | 0.117647 | 0 | 51 | 107 | . | . | . | . |
| PM1548-SK1T1 | OR2T6 | p.Tyr60His | missense_variant | MODERATE | 0.109244 | 0 | 119 | 68 | . | . | . | . |
| PM1548-SK1T1 | DIP2C | p.Val1150Ile | missense_variant | MODERATE | 0 | 0 | 239 | 183 | . | . | . | . |
| PM1548-SK1T1 | WDR37 | p.Pro169Ser | missense_variant | MODERATE | 0.015625 | 0 | 128 | 220 | . | . | . | . |
| PM1548-SK1T1 | CALML5 | p.Glu102Lys | missense_variant | MODERATE | 0.174757 | 0 | 103 | 79 | . | . | . | . |
| PM1548-SK1T1 | FRMD4A | p.Pro1005fs | frameshift_variant | HIGH | 0.138122 | 0 | 181 | 118 | . | . | . | . |
| PM1548-SK1T1 | CUBN | p.Trp3323Arg | missense_variant | MODERATE | 0.152709 | 0 | 203 | 170 | . | . | . | . |
| PM1548-SK1T1 | GPR158 | p.Arg448Gln | missense_variant | MODERATE | 0.0416667 | 0 | 96 | 52 | . | . | . | . |
| PM1548-SK1T1 | SVIL | p.Arg2075Cys | missense_variant | MODERATE | 0.113475 | 0 | 141 | 40 | . | . | . | . |
| PM1548-SK1T1 | PARD3 | p.Arg1278Gly | missense_variant | MODERATE | 0.126437 | 0 | 174 | 90 | . | . | . | . |
| PM1548-SK1T1 | FRMPD2 | p.Pro226Leu | missense_variant | MODERATE | 0.0389105 | 0 | 257 | 178 | . | . | . | . |
| PM1548-SK1T1 | PRKG1 | p.Asn601fs | frameshift_variant | HIGH | 0.134146 | 0 | 82 | 55 | . | . | . | . |
| PM1548-SK1T1 | PBLD | p.Asp167Tyr | missense_variant | MODERATE | 0 | 0 | 162 | 186 | . | . | . | . |
| PM1548-SK1T1 | DDX50 | p.Tyr39Cys | missense_variant | MODERATE | 0.0526316 | 0 | 57 | 39 | . | . | . | . |
| PM1548-SK1T1 | ADAMTS14 | p.Gly710Arg | missense_variant | MODERATE | 0.0674157 | 0 | 89 | 80 | . | . | . | . |
| PM1548-SK1T1 | DDIT4 | p.Ser46Cys | missense_variant | MODERATE | 0 | 0 | 188 | 263 | . | . | . | . |
| PM1548-SK1T1 | DDIT4 | p.Thr47Ala | missense_variant | MODERATE | 0 | 0 | 188 | 263 | . | . | . | . |
| PM1548-SK1T1 | KAT6B | p.Arg463His | missense_variant | MODERATE | 0.0833333 | 0 | 144 | 106 | true | . | . | . |
| PM1548-SK1T1 | KCNMA1 | p.Ala1039Val | missense_variant | MODERATE | 0.136364 | 0 | 154 | 73 | . | . | . | . |
| PM1548-SK1T1 | ZMIZ1 | p.Asp1051fs | frameshift_variant | HIGH | 0 | 0 | 120 | 74 | . | . | . | . |
| PM1548-SK1T1 | MAT1A | p.Arg81Gln | missense_variant | MODERATE | 0.0503145 | 0 | 159 | 225 | . | . | . | . |
| PM1548-SK1T1 | TSPAN14 | p.Asp190Gly | missense_variant | MODERATE | 0 | 0 | 203 | 134 | . | . | . | . |
| PM1548-SK1T1 | SH2D4B | p.Ala186Val | missense_variant | MODERATE | 0.0869565 | 0 | 115 | 42 | . | . | . | . |
| PM1548-SK1T1 | LRIT2 | p.Thr520fs | frameshift_variant | HIGH | 0.202186 | 0.0580357 | 183 | 224 | . | . | . | . |
| PM1548-SK1T1 | CCSER2 | c.2326-13521C> | intron_variant | MODIFIER | 0 | 0 | 117 | 179 | . | . | . | . |
| PM1548-SK1T1 | CYP26C1 | p.Ser70Leu | missense_variant | MODERATE | 0.117647 | 0 | 34 | 149 | . | . | . | . |
| PM1548-SK1T1 | MYOF | p.Arg726Gln | missense_variant | MODERATE | 0.011976 | 0 | 167 | 43 | . | . | . | . |
| PM1548-SK1T1 | SORBS1 | p.Arg585Cys | missense_variant | MODERATE | 0.25 | 0 | 24 | 42 | . | . | . | . |
| PM1548-SK1T1 | ALDH18A1 | p.Arg622Trp | missense_variant | MODERATE | 0.0382166 | 0 | 157 | 86 | . | . | . | . |
| PM1548-SK1T1 | ENTPD1 | p.Arg252Leu | missense_variant | MODERATE | 0 | 0 | 276 | 84 | . | . | . | . |
| PM1548-SK1T1 | MORN4 | p.Leu202Ile | missense_variant | MODERATE | 0 | 0 | 144 | 87 | . | . | . | . |
| PM1548-SK1T1 | PKD2L1 | p.Arg206Cys | missense_variant | MODERATE | 0.141935 | 0 | 155 | 130 | . | . | . | . |
| PM1548-SK1T1 | MRPL43 | p.Asn66Ser | missense_variant | MODERATE | 0.00653595 | 0 | 153 | 81 | . | . | . | . |
| PM1548-SK1T1 | POLL | p.Arg419* | stop_gained | HIGH | 0.157895 | 0 | 76 | 44 | . | . | . | . |
| PM1548-SK1T1 | TAF5 | p.Phe374fs | frameshift_variant | HIGH | 0.0294118 | 0 | 68 | 50 | . | . | . | . |
| PM1548-SK1T1 | WDR96 | p.Val70Ala | missense_variant | MODERATE | 0.0357143 | 0 | 84 | 59 | . | . | . | . |
| PM1548-SK1T1 | SORCS3 | p.Lys1023Gln | missense_variant | MODERATE | 0.020979 | 0 | 143 | 58 | . | . | . | . |
| PM1548-SK1T1 | TCF7L2 | p.Ser587Leu | missense_variant | MODERATE | 0 | 0 | 149 | 171 | true | . | true | . |
| PM1548-SK1T1 | ABLIM1 | p.Leu633Met | missense_variant | MODERATE | 0 | 0 | 244 | 266 | . | . | . | . |
| PM1548-SK1T1 | GFRA1 | p.Ala173Val | missense_variant | MODERATE | 0.037037 | 0 | 108 | 73 | . | . | . | . |
| PM1548-SK1T1 | KIAA1598 | p.Arg482His | missense_variant | MODERATE | 0 | 0 | 145 | 182 | true | . | . | . |
| PM1548-SK1T1 | VAX1 | p.Cys200* | stop_gained | HIGH | 0.125 | 0 | 32 | 96 | . | . | . | . |
| PM1548-SK1T1 | EIF3A | p.His257His | synonymous_variant | LOW | 0 | 0 | 161 | 131 | . | . | . | . |
| PM1548-SK1T1 | ATE1 | p.Leu289Gln | missense_variant | MODERATE | 0 | 0 | 92 | 77 | . | . | . | . |
| PM1548-SK1T1 | PLEKHA1 | p.Arg9His | missense_variant | MODERATE | 0 | 0 | 89 | 43 | . | . | . | . |
| PM1548-SK1T1 | PLEKHA1 | p.Val404Met | missense_variant | MODERATE | 0 | 0 | 72 | 76 | . | . | . | . |
| PM1548-SK1T1 | HMX3 | p.Pro23fs | frameshift_variant | HIGH | 0.2 | 0 | 25 | 68 | . | . | . | . |
| PM1548-SK1T1 | CLRN3 | p.Pro71Thr | missense_variant | MODERATE | 0.0104167 | 0.00613497 | 96 | 163 | . | . | . | . |
| PM1548-SK1T1 | MKI67 | p.Arg2258* | stop_gained | HIGH | 0.0833333 | 0 | 168 | 90 | . | . | . | . |
| PM1548-SK1T1 | TTC40 | p.Thr1864Met | missense_variant | MODERATE | 0.183673 | 0 | 98 | 72 | . | . | . | . |
| PM1548-SK1T1 | KNDC1 | p.Cys79fs | frameshift_variant | HIGH | 0.0165746 | 0 | 181 | 162 | . | . | . | . |
| PM1548-SK1T1 | PIDD | p.Ala727Glu | missense_variant | MODERATE | 0.105263 | 0 | 38 | 108 | . | . | . | . |
| PM1548-SK1T1 | MUC6 | p.Thr1861Ala | missense_variant | MODERATE | 0.196203 | 0 | 158 | 78 | . | . | . | . |
| PM1548-SK1T1 | MUC6 | p.Ile1857Thr | missense_variant | MODERATE | 0.16 | 0 | 150 | 78 | . | . | . | . |
| PM1548-SK1T1 | MUC6 | p.His1752Gln | missense_variant | MODERATE | 0.0172414 | 0 | 58 | 31 | . | . | . | . |

| | | | | | | | | | | | | |
|---|---|---|---|---|---|---|---|---|---|---|---|---|
| PM1548-SK1T1 | MUC6 | p.Thr1711Ile | missense_variant | MODERATE | 0.0462185 | 0 | 238 | 51 | . | . | . | . |
| PM1548-SK1T1 | MUC2 | p.Gly1331Arg | missense_variant | MODERATE | 0 | 0 | 66 | 32 | . | . | . | . |
| PM1548-SK1T1 | MUC5B | p.Ala110Thr | missense_variant | MODERATE | 0 | 0 | 91 | 216 | . | . | . | . |
| PM1548-SK1T1 | BRSK2 | p.Pro440fs | frameshift_variant | HIGH | 0.165919 | 0 | 223 | 593 | . | . | . | . |
| PM1548-SK1T1 | TSPAN32 | p.Ala178Val | missense_variant | MODERATE | 0.136986 | 0 | 73 | 69 | . | . | . | . |
| PM1548-SK1T1 | TSSC4 | p.Arg92Cys | missense_variant | MODERATE | 0.0408805 | 0 | 318 | 525 | . | . | . | . |
| PM1548-SK1T1 | OSBPL5 | p.Thr843Met | missense_variant | MODERATE | 0.106557 | 0 | 122 | 106 | . | . | . | . |
| PM1548-SK1T1 | STIM1 | p.Lys371Arg | missense_variant | MODERATE | 0 | 0 | 322 | 322 | . | . | . | . |
| PM1548-SK1T1 | OR51E1 | p.Tyr63His | missense_variant | MODERATE | 0 | 0 | 242 | 116 | . | . | . | . |
| PM1548-SK1T1 | OR51A2 | p.Lys289Asn | missense_variant | MODERATE | 0.0941176 | 0 | 85 | 30 | . | . | . | . |
| PM1548-SK1T1 | SOX6 | p.Met803Ile | missense_variant | MODERATE | 0 | 0 | 66 | 33 | . | . | . | . |
| PM1548-SK1T1 | SOX6 | p.Met803Thr | missense_variant | MODERATE | 0 | 0 | 66 | 33 | . | . | . | . |
| PM1548-SK1T1 | SOX6 | p.Met803Val | missense_variant | MODERATE | 0 | 0 | 68 | 33 | . | . | . | . |
| PM1548-SK1T1 | ABCC8 | p.Asp761Asn | missense_variant | MODERATE | 0 | 0 | 344 | 231 | . | . | . | . |
| PM1548-SK1T1 | LDHC | p.Lys5Glu | missense_variant | MODERATE | 0.0148148 | 0 | 135 | 119 | . | . | . | . |
| PM1548-SK1T1 | MRGPRX2 | p.Asn62Asn | synonymous_variant | LOW | 0 | 0 | 131 | 83 | . | . | . | . |
| PM1548-SK1T1 | ZDHHC13 | p.Gly591Asp | missense_variant | MODERATE | 0 | 0 | 95 | 33 | . | . | . | . |
| PM1548-SK1T1 | ANO3 | p.Leu273Pro | missense_variant | MODERATE | 0 | 0 | 359 | 435 | . | . | . | . |
| PM1548-SK1T1 | HIPK3 | p.Asp532Ala | missense_variant | MODERATE | 0.133333 | 0 | 105 | 48 | . | . | . | . |
| PM1548-SK1T1 | ABTB2 | p.Asp793Asn | missense_variant | MODERATE | 0.0277778 | 0.00668896 | 180 | 299 | . | . | . | . |
| PM1548-SK1T1 | RAG1 | p.Arg507Trp | missense_variant | MODERATE | 0 | 0 | 87 | 86 | . | . | . | true |
| PM1548-SK1T1 | TP53I11 | p.Leu62Arg | missense_variant | MODERATE | 0.106383 | 0 | 47 | 33 | . | . | . | . |
| PM1548-SK1T1 | MAPK8IP1 | p.Thr321Met | missense_variant | MODERATE | 0.0872093 | 0 | 172 | 140 | . | true | . | . |
| PM1548-SK1T1 | DGKZ | n.46369267G>A | intragenic_variant | MODIFIER | 0.194444 | 0 | 36 | 32 | . | . | . | . |
| PM1548-SK1T1 | DGKZ | p.Asp574Asn | missense_variant | MODERATE | 0 | 0 | 64 | 74 | . | . | . | . |
| PM1548-SK1T1 | ACP2 | p.Arg231His | missense_variant | MODERATE | 0.0116959 | 0 | 171 | 203 | . | . | . | . |
| PM1548-SK1T1 | MYBPC3 | p.Arg654Ser | missense_variant | MODERATE | 0 | 0 | 131 | 97 | . | . | . | . |
| PM1548-SK1T1 | OR5D14 | p.Tyr261Cys | missense_variant | MODERATE | 0 | 0 | 200 | 71 | . | . | . | . |
| PM1548-SK1T1 | OR5W2 | p.Ala29Ser | missense_variant | MODERATE | 0.0198675 | 0 | 151 | 99 | . | . | . | . |
| PM1548-SK1T1 | APLNR | p.Ala161Thr | missense_variant | MODERATE | 0.013245 | 0 | 151 | 160 | . | . | . | . |
| PM1548-SK1T1 | OR10Q1 | p.Thr287Ile | missense_variant | MODERATE | 0.0116279 | 0 | 86 | 90 | . | . | . | . |
| PM1548-SK1T1 | DTX4 | p.Lys396fs | frameshift_variant | HIGH | 0.134615 | 0 | 104 | 34 | . | . | . | . |
| PM1548-SK1T1 | DTX4 | p.Gly510fs | frameshift_variant | HIGH | 0.0208333 | 0.00704225 | 192 | 142 | . | . | . | . |
| PM1548-SK1T1 | DAK | p.Arg380Gln | missense_variant | MODERATE | 0.0324324 | 0 | 185 | 319 | . | . | . | . |
| PM1548-SK1T1 | FEN1 | p.Arg332Cys | missense_variant | MODERATE | 0 | 0 | 88 | 80 | . | . | . | . |
| PM1548-SK1T1 | BEST1 | p.Ser413Asn | missense_variant | MODERATE | 0.0070922 | 0 | 141 | 159 | . | . | . | . |
| PM1548-SK1T1 | AHNAK | p.Ser4908Leu | missense_variant | MODERATE | 0.146341 | 0 | 41 | 39 | . | . | . | . |
| PM1548-SK1T1 | AHNAK | p.His4772Asn | missense_variant | MODERATE | 0.113208 | 0 | 53 | 54 | . | . | . | . |
| PM1548-SK1T1 | AHNAK | p.Gly4561Asp | missense_variant | MODERATE | 0 | 0 | 264 | 206 | . | . | . | . |
| PM1548-SK1T1 | AHNAK | p.Val4560Leu | missense_variant | MODERATE | 0 | 0 | 263 | 203 | . | . | . | . |
| PM1548-SK1T1 | AHNAK | p.Val3640Ile | missense_variant | MODERATE | 0 | 0 | 85 | 44 | . | . | . | . |
| PM1548-SK1T1 | AHNAK | p.Asn3637Lys | missense_variant | MODERATE | 0 | 0 | 84 | 44 | . | . | . | . |
| PM1548-SK1T1 | AHNAK | p.Lys2706fs | frameshift_variant | HIGH | 0 | 0 | 10 | 43 | . | . | . | . |
| PM1548-SK1T1 | AHNAK | p.Thr1986Ala | missense_variant | MODERATE | 0 | 0 | 300 | 324 | . | . | . | . |
| PM1548-SK1T1 | CHRM1 | p.Pro319fs | frameshift_variant | HIGH | 0.158537 | 0 | 82 | 88 | . | . | . | . |
| PM1548-SK1T1 | SLC22A6 | p.Val240Met | missense_variant | MODERATE | 0 | 0 | 190 | 227 | . | . | . | . |
| PM1548-SK1T1 | SLC22A25 | p.Asn62Asp | missense_variant | MODERATE | 0.108108 | 0 | 111 | 110 | . | . | . | . |
| PM1548-SK1T1 | C11orf95 | n.1245C>T | non_coding_exon_variant | MODIFIER | 0 | 0 | 16 | 34 | . | . | . | . |
| PM1548-SK1T1 | NRXN2 | p.Asp277Asn | missense_variant | MODERATE | 0 | 0 | 152 | 229 | . | . | . | . |
| PM1548-SK1T1 | SNX15 | p.Leu10Met | missense_variant | MODERATE | 0.1 | 0 | 60 | 37 | . | . | . | . |
| PM1548-SK1T1 | RBM4 | p.Tyr303His | missense_variant | MODERATE | 0 | 0 | 254 | 269 | . | . | . | . |
| PM1548-SK1T1 | SPTBN2 | p.Cys1393Ser | missense_variant | MODERATE | 0 | 0 | 180 | 300 | . | . | . | . |
| PM1548-SK1T1 | SPTBN2 | p.Arg465Gln | missense_variant | MODERATE | 0 | 0 | 250 | 212 | . | . | . | . |
| PM1548-SK1T1 | LRP5 | p.Ala724Val | missense_variant | MODERATE | 0.0425532 | 0 | 47 | 227 | . | . | . | . |
| PM1548-SK1T1 | IGHMBP2 | p.Pro809Ser | missense_variant | MODERATE | 0.0416667 | 0 | 48 | 134 | . | . | . | . |
| PM1548-SK1T1 | TPCN2 | p.Gly492Val | missense_variant | MODERATE | 0 | 0 | 56 | 21 | . | . | . | . |
| PM1548-SK1T1 | CCND1 | p.Val94Leu | missense_variant | MODERATE | 0 | 0 | 97 | 259 | true | true | true | . |
| PM1548-SK1T1 | INPPL1 | p.Leu957Phe | missense_variant | MODERATE | 0.011236 | 0 | 89 | 30 | . | . | . | . |
| PM1548-SK1T1 | PDE2A | p.His99fs | frameshift_variant | HIGH | 0.136364 | 0 | 176 | 161 | . | . | . | . |
| PM1548-SK1T1 | RNF169 | p.Glu131* | stop_gained | HIGH | 0.133333 | 0 | 30 | 52 | . | . | . | . |
| PM1548-SK1T1 | TENM4 | p.Ala235Thr | missense_variant | MODERATE | 0.0138889 | 0 | 144 | 205 | . | . | . | . |
| PM1548-SK1T1 | PCF11 | p.Leu745Leu | synonymous_variant | LOW | 0 | 0 | 111 | 203 | . | . | . | . |
| PM1548-SK1T1 | PCF11 | p.Ser756Gly | missense_variant | MODERATE | 0 | 0 | 129 | 215 | . | . | . | . |
| PM1548-SK1T1 | PCF11 | p.Arg833Trp | missense_variant | MODERATE | 0.277311 | 0 | 119 | 258 | . | . | . | . |
| PM1548-SK1T1 | TYR | p.Gly191fs | frameshift_variant | HIGH | 0.0491071 | 0 | 224 | 94 | . | . | . | . |
| PM1548-SK1T1 | FAT3 | p.Ala2495Ser | missense_variant | MODERATE | 0.00480769 | 0 | 208 | 229 | . | . | . | . |
| PM1548-SK1T1 | PANX1 | p.Ser250Thr | missense_variant | MODERATE | 0.0382353 | 0 | 340 | 236 | . | . | . | . |
| PM1548-SK1T1 | MTMR2 | p.Lys451Glu | missense_variant | MODERATE | 0 | 0 | 72 | 77 | . | . | . | . |
| PM1548-SK1T1 | MTMR2 | p.Asp436Asn | missense_variant | MODERATE | 0 | 0 | 54 | 75 | . | . | . | . |
| PM1548-SK1T1 | MTMR2 | p.Leu433Val | missense_variant | MODERATE | 0 | 0 | 100 | 282 | . | . | . | . |
| PM1548-SK1T1 | MMP3 | p.Asp200Gly | missense_variant | MODERATE | 0 | 0 | 95 | 91 | . | . | . | . |
| PM1548-SK1T1 | DYNC2H1 | p.Tyr3523His | missense_variant | MODERATE | 0.00925926 | 0 | 108 | 62 | . | . | . | . |
| PM1548-SK1T1 | CASP1 | p.Phe262Tyr | missense_variant | MODERATE | 0.0248963 | 0 | 241 | 177 | . | . | . | . |
| PM1548-SK1T1 | KBTBD3 | p.Lys478fs | frameshift_variant | HIGH | 0 | 0 | 24 | 12 | . | . | . | . |
| PM1548-SK1T1 | NXPE1 | p.Thr169Ile | missense_variant | MODERATE | 0 | 0 | 81 | 97 | . | . | . | . |
| PM1548-SK1T1 | NXPE1 | p.Met157Ile | missense_variant | MODERATE | 0 | 0 | 88 | 97 | . | . | . | . |
| PM1548-SK1T1 | NXPE1 | p.Met157Thr | missense_variant | MODERATE | 0 | 0 | 88 | 96 | . | . | . | . |
| PM1548-SK1T1 | NXPE1 | p.Thr149Lys | missense_variant | MODERATE | 0 | 0 | 114 | 118 | . | . | . | . |
| PM1548-SK1T1 | DSCAML1 | p.Val1762Ile | missense_variant | MODERATE | 0.13879 | 0 | 281 | 226 | . | . | . | . |
| PM1548-SK1T1 | DSCAML1 | p.Asp949fs | frameshift_variant | HIGH | 0.0991736 | 0 | 242 | 244 | . | . | . | . |
| PM1548-SK1T1 | ATP5L | p.Leu84Trp | missense_variant | MODERATE | 0 | 0 | 89 | 125 | . | . | . | . |
| PM1548-SK1T1 | ATP5L | p.Val89Ile | missense_variant | MODERATE | 0 | 0 | 96 | 140 | . | . | . | . |
| PM1548-SK1T1 | ATP5L | p.Ile99Val | missense_variant | MODERATE | 0 | 0 | 96 | 141 | . | . | . | . |
| PM1548-SK1T1 | PHLDB1 | p.Ala932Ser | missense_variant | MODERATE | 0 | 0 | 130 | 223 | . | . | . | . |
| PM1548-SK1T1 | PHLDB1 | p.Pro938Leu | missense_variant | MODERATE | 0 | 0 | 124 | 220 | . | . | . | . |
| PM1548-SK1T1 | CBL | p.Ser237Leu | missense_variant | MODERATE | 0 | 0 | 79 | 128 | true | . | . | . |
| PM1548-SK1T1 | CBL | p.Pro504Gln | missense_variant | MODERATE | 0 | 0 | 97 | 111 | true | . | . | . |
| PM1548-SK1T1 | CRTAM | p.Leu60fs | frameshift_variant | HIGH | 0.0377358 | 0 | 106 | 170 | . | . | . | . |
| PM1548-SK1T1 | OR10G8 | p.Lys265Glu | missense_variant | MODERATE | 0 | 0 | 276 | 232 | . | . | . | . |
| PM1548-SK1T1 | OR10G8 | p.Lys265Asn | missense_variant | MODERATE | 0 | 0 | 237 | 193 | . | . | . | . |
| PM1548-SK1T1 | OR8G2 | p.Val2Gly | missense_variant | MODERATE | 0 | 0 | 4 | 1 | . | . | . | . |
| PM1548-SK1T1 | ROBO4 | p.Arg776His | missense_variant | MODERATE | 0.0052356 | 0 | 191 | 269 | . | . | . | . |
| PM1548-SK1T1 | ADAMTS8 | p.Arg9Gly | missense_variant | MODERATE | 0 | 0 | 26 | 48 | . | . | . | true |
| PM1548-SK1T1 | IQSEC3 | p.Gln1075Lys | missense_variant | MODERATE | 0.1 | 0 | 40 | 59 | . | . | . | . |
| PM1548-SK1T1 | FOXM1 | p.Leu334Ser | missense_variant | MODERATE | 0 | 0 | 116 | 177 | . | . | . | . |
| PM1548-SK1T1 | FOXM1 | p.Asp328Glu | missense_variant | MODERATE | 0 | 0 | 128 | 203 | . | . | . | . |
| PM1548-SK1T1 | VWF | p.Ala865Asp | missense_variant | MODERATE | 0 | 0 | 184 | 226 | . | . | . | . |
| PM1548-SK1T1 | IFFO1 | p.Val126Met | missense_variant | MODERATE | 0.0506329 | 0 | 79 | 165 | . | . | . | . |
| PM1548-SK1T1 | PHB2 | p.Arg270Arg | synonymous_variant | LOW | 0 | 0 | 261 | 405 | . | . | . | . |
| PM1548-SK1T1 | PHB2 | p.Asn269Asn | synonymous_variant | LOW | 0 | 0 | 261 | 404 | . | . | . | . |
| PM1548-SK1T1 | TAS2R10 | p.Ser41Thr | missense_variant | MODERATE | 0.0636364 | 0 | 110 | 57 | . | . | . | . |
| PM1548-SK1T1 | TAS2R31 | p.Ala177Thr | missense_variant | MODERATE | 0.126761 | 0 | 71 | 47 | . | . | . | . |
| PM1548-SK1T1 | PLCZ1 | p.Gly133Val | missense_variant | MODERATE | 0.102041 | 0 | 49 | 42 | . | . | . | . |
| PM1548-SK1T1 | PDE3A | p.Glu171fs | frameshift_variant | HIGH | 0 | 0 | 73 | 56 | . | . | . | . |
| PM1548-SK1T1 | TM7SF3 | c.869-699C>G | intron_variant | MODIFIER | 0 | 0 | 85 | 125 | . | . | . | . |
| PM1548-SK1T1 | DDX11 | p.Asp935Asn | missense_variant | MODERATE | 0.140845 | 0 | 71 | 35 | . | . | . | . |
| PM1548-SK1T1 | COL2A1 | p.Asp682Asn | missense_variant | MODERATE | 0.0844156 | 0 | 154 | 53 | true | . | . | . |
| PM1548-SK1T1 | KMT2D | p.Ala4804Val | missense_variant | MODERATE | 0.168421 | 0.0060241 | 285 | 166 | true | true | true | true |
| PM1548-SK1T1 | KMT2D | p.Leu4214Leu | synonymous_variant | LOW | 0 | 0 | 257 | 376 | true | true | true | true |
| PM1548-SK1T1 | KMT2D | p.Thr3166Ile | missense_variant | MODERATE | 0.183673 | 0 | 98 | 193 | true | true | true | true |
| PM1548-SK1T1 | KMT2D | p.Thr2581Ala | missense_variant | MODERATE | 0.021978 | 0 | 91 | 134 | true | true | true | true |
| PM1548-SK1T1 | KMT2D | p.Gly1628fs | frameshift_variant | HIGH | 0 | 0 | 166 | 136 | true | true | true | true |
| PM1548-SK1T1 | PRPH | p.Ala348Thr | missense_variant | MODERATE | 0.0211268 | 0 | 142 | 109 | . | . | . | . |
| PM1548-SK1T1 | FAM186B | p.Leu494Met | missense_variant | MODERATE | 0.052 | 0 | 250 | 422 | . | . | . | . |
| PM1548-SK1T1 | ANKRD33 | p.Ala173Val | missense_variant | MODERATE | 0.00775194 | 0 | 129 | 242 | . | . | . | . |
| PM1548-SK1T1 | KRT74 | p.Gln440His | missense_variant | MODERATE | 0 | 0 | 44 | 34 | . | . | . | . |
| PM1548-SK1T1 | PCBP2 | p.Pro175fs | frameshift_variant | HIGH | 0.0810811 | 0 | 74 | 35 | . | . | . | . |
| PM1548-SK1T1 | HNRNPA1 | p.Thr61Ala | missense_variant | MODERATE | 0.0402685 | 0 | 149 | 168 | . | . | . | . |
| PM1548-SK1T1 | ITGA5 | p.Arg193His | missense_variant | MODERATE | 0 | 0 | 92 | 104 | . | . | . | . |
| PM1548-SK1T1 | MYO1A | p.Arg559Cys | missense_variant | MODERATE | 0.124352 | 0 | 193 | 190 | . | . | . | . |
| PM1548-SK1T1 | NAB2 | p.Pro388Thr | missense_variant | MODERATE | 0 | 0 | 135 | 78 | true | . | . | . |
| PM1548-SK1T1 | NAB2 | p.Pro388His | missense_variant | MODERATE | 0 | 0 | 135 | 78 | true | . | . | . |
| PM1548-SK1T1 | R3HDM2 | p.Arg956His | missense_variant | MODERATE | 0.105263 | 0 | 95 | 33 | . | . | . | . |
| PM1548-SK1T1 | CTDSP2 | p.Glu259Asp | missense_variant | MODERATE | 0 | 0 | 169 | 90 | . | . | . | . |
| PM1548-SK1T1 | CTDSP2 | p.Ala258Thr | missense_variant | MODERATE | 0.00591716 | 0 | 169 | 90 | . | . | . | . |
| PM1548-SK1T1 | CTDSP2 | p.Ile251Val | missense_variant | MODERATE | 0 | 0 | 150 | 94 | . | . | . | . |
| PM1548-SK1T1 | PTPRB | p.Val1992Ala | missense_variant | MODERATE | 0 | 0 | 100 | 51 | true | . | . | . |

| | | | | | | | | | | | | |
|---|---|---|---|---|---|---|---|---|---|---|---|---|
| PM1548-SK1T1 | NAV3 | p.Met1275Val | missense_variant | MODERATE | 0 | 0 | 71 | 67 | . | true | . | . |
| PM1548-SK1T1 | NAV3 | p.Pro1276Leu | missense_variant | MODERATE | 0 | 0 | 109 | 130 | . | true | . | . |
| PM1548-SK1T1 | NAV3 | p.Thr1281Ser | missense_variant | MODERATE | 0 | 0 | 145 | 146 | . | true | . | . |
| PM1548-SK1T1 | TMTC2 | p.Gly143Arg | missense_variant | MODERATE | 0.12931 | 0 | 116 | 67 | . | . | . | . |
| PM1548-SK1T1 | ALX1 | p.Thr68Ser | missense_variant | MODERATE | 0.0650407 | 0 | 123 | 191 | . | . | . | . |
| PM1548-SK1T1 | FGD6 | p.Met965fs | frameshift_variant | HIGH | 0.0131579 | 0 | 76 | 56 | . | . | . | . |
| PM1548-SK1T1 | APAF1 | p.Gln578Arg | missense_variant | MODERATE | 0 | 0 | 198 | 126 | . | . | . | . |
| PM1548-SK1T1 | SLC41A2 | p.Pro266Ser | missense_variant | MODERATE | 0.157025 | 0 | 121 | 65 | . | . | . | . |
| PM1548-SK1T1 | PRDM4 | p.Glu313Glu | synonymous_variant | LOW | 0 | 0 | 140 | 274 | . | . | . | . |
| PM1548-SK1T1 | TMEM119 | p.Thr81fs | frameshift_variant | HIGH | 0.00816327 | 0 | 245 | 297 | . | . | . | . |
| PM1548-SK1T1 | FOXN4 | p.Ala462Thr | missense_variant | MODERATE | 0.0160643 | 0.00492611 | 249 | 203 | . | . | . | . |
| PM1548-SK1T1 | FAM222A | p.Pro392Thr | missense_variant | MODERATE | 0 | 0 | 108 | 422 | . | . | . | . |
| PM1548-SK1T1 | FAM222A | p.Ser400Ala | missense_variant | MODERATE | 0 | 0 | 119 | 423 | . | . | . | . |
| PM1548-SK1T1 | TCHP | p.Ala371Thr | missense_variant | MODERATE | 0.0535714 | 0 | 112 | 45 | . | . | . | . |
| PM1548-SK1T1 | SH2B3 | p.Glu523fs | frameshift_variant | HIGH | 0.113924 | 0 | 158 | 156 | true | . | . | . |
| PM1548-SK1T1 | HECTD4 | p.Ala3707Val | missense_variant | MODERATE | 0.0531915 | 0 | 94 | 63 | . | . | . | . |
| PM1548-SK1T1 | TPCN1 | p.Leu234Arg | missense_variant | MODERATE | 0.104651 | 0 | 172 | 134 | . | . | . | . |
| PM1548-SK1T1 | RBM19 | p.Asn533Asn | synonymous_variant | LOW | 0.166667 | 0 | 102 | 250 | . | . | . | . |
| PM1548-SK1T1 | RAB35 | n.120541738C> | intragenic_variant | MODIFIER | 0.0221239 | 0 | 226 | 244 | . | . | . | . |
| PM1548-SK1T1 | GCN1L1 | p.Ser2359Pro | missense_variant | MODERATE | 0.237179 | 0 | 156 | 83 | . | . | . | . |
| PM1548-SK1T1 | SETD1B | p.His8fs | frameshift_variant | HIGH | 0.215686 | 0 | 51 | 140 | . | . | . | . |
| PM1548-SK1T1 | SETD1B | p.Pro1361fs | frameshift_variant | HIGH | 0 | 0 | 94 | 190 | . | . | . | . |
| PM1548-SK1T1 | ZCCHC8 | p.Gly551Asp | missense_variant | MODERATE | 0 | 0 | 83 | 34 | true | . | . | . |
| PM1548-SK1T1 | ABCB9 | p.His709Gln | missense_variant | MODERATE | 0.00440529 | 0 | 227 | 333 | . | . | . | . |
| PM1548-SK1T1 | ABCB9 | p.His709Arg | missense_variant | MODERATE | 0 | 0 | 227 | 333 | . | . | . | . |
| PM1548-SK1T1 | ABCB9 | p.Lys694Arg | missense_variant | MODERATE | 0 | 0 | 192 | 301 | . | . | . | . |
| PM1548-SK1T1 | PITPNM2 | p.Arg749Trp | missense_variant | MODERATE | 0.0592105 | 0 | 152 | 82 | . | . | . | . |
| PM1548-SK1T1 | SBNO1 | p.Arg1283Cys | missense_variant&splice_region_variant | MODERATE | 0.0961538 | 0 | 156 | 246 | . | . | . | . |
| PM1548-SK1T1 | DNAH10 | p.Pro3133His | missense_variant | MODERATE | 0.00873362 | 0.00452489 | 229 | 221 | . | . | . | . |
| PM1548-SK1T1 | TMEM132B | p.Ala297Thr | missense_variant | MODERATE | 0.130435 | 0 | 138 | 79 | . | . | . | . |
| PM1548-SK1T1 | GPR133 | p.Gly731Arg | missense_variant | MODERATE | 0.0458015 | 0 | 131 | 110 | . | . | . | . |
| PM1548-SK1T1 | FBRSL1 | p.Glu251Asp | missense_variant | MODERATE | 0.1875 | 0 | 32 | 153 | . | . | . | . |
| PM1548-SK1T1 | MPHOSPH8 | p.Val686Ile | missense_variant | MODERATE | 0.102941 | 0 | 204 | 75 | . | . | . | . |
| PM1548-SK1T1 | GJA3 | p.Ala213Thr | missense_variant | MODERATE | 0.0133333 | 0 | 75 | 153 | . | . | . | . |
| PM1548-SK1T1 | LATS2 | p.Pro41Gln | missense_variant | MODERATE | 0.0545455 | 0.025641 | 110 | 39 | . | . | . | . |
| PM1548-SK1T1 | CDK8 | p.Asn372Thr | missense_variant | MODERATE | 0 | 0 | 251 | 249 | . | . | . | . |
| PM1548-SK1T1 | POLR1D | n.28240022G>A | intragenic_variant | MODIFIER | 0.0662651 | 0 | 166 | 101 | . | . | . | . |
| PM1548-SK1T1 | FRY | p.Gln2970His | missense_variant | MODERATE | 0 | 0 | 59 | 39 | . | . | . | . |
| PM1548-SK1T1 | BRCA2 | p.Asn863fs | frameshift_variant | HIGH | 0.019802 | 0 | 101 | 84 | true | true | . | true |
| PM1548-SK1T1 | STARD13 | p.Arg155His | missense_variant | MODERATE | 0 | 0 | 151 | 32 | . | . | . | . |
| PM1548-SK1T1 | MAB21L1 | p.Ser103Ser | synonymous_variant | LOW | 0.0431034 | 0 | 116 | 101 | . | . | . | . |
| PM1548-SK1T1 | FOXO1 | p.Ser152Thr | missense_variant | MODERATE | 0 | 0 | 56 | 140 | true | . | . | . |
| PM1548-SK1T1 | KBTBD7 | p.Tyr195Phe | missense_variant | MODERATE | 0 | 0 | 404 | 644 | . | . | . | . |
| PM1548-SK1T1 | GPALPP1 | p.Pro37Arg | missense_variant | MODERATE | 0.115646 | 0 | 147 | 103 | . | . | . | . |
| PM1548-SK1T1 | FAM124A | p.Ser257Pro | missense_variant | MODERATE | 0.101449 | 0 | 138 | 101 | . | . | . | . |
| PM1548-SK1T1 | NEK3 | p.Asn343Lys | missense_variant | MODERATE | 0.109091 | 0 | 55 | 44 | . | . | . | . |
| PM1548-SK1T1 | DACH1 | p.Arg196Arg | synonymous_variant | LOW | 0 | 0 | 79 | 115 | . | . | . | . |
| PM1548-SK1T1 | ABCC4 | p.Ile1219fs | frameshift_variant | HIGH | 0.144068 | 0 | 118 | 130 | . | . | . | . |
| PM1548-SK1T1 | FGF14 | p.Asp33Gly | missense_variant | MODERATE | 0 | 0 | 191 | 97 | . | . | . | . |
| PM1548-SK1T1 | CCDC168 | p.Met4045fs | frameshift_variant | HIGH | 0.015625 | 0 | 128 | 95 | . | . | . | . |
| PM1548-SK1T1 | ARHGEF7 | p.Thr23fs | frameshift_variant | HIGH | 0.0155039 | 0 | 129 | 189 | . | . | . | . |
| PM1548-SK1T1 | MCF2L | p.Arg590Gln | missense_variant | MODERATE | 0.0862069 | 0 | 58 | 34 | . | . | . | . |
| PM1548-SK1T1 | CDC16 | p.Asn557fs | frameshift_variant | HIGH | 0.125 | 0 | 80 | 89 | . | . | . | . |
| PM1548-SK1T1 | UPF3A | p.Asp129Glu | missense_variant | MODERATE | 0.0246914 | 0.0163934 | 81 | 61 | . | . | . | . |
| PM1548-SK1T1 | OR11G2 | p.Phe141Phe | synonymous_variant | LOW | 0 | 0 | 189 | 117 | . | . | . | . |
| PM1548-SK1T1 | ACIN1 | p.Glu192Asp | missense_variant | MODERATE | 0 | 0 | 149 | 129 | . | . | . | . |
| PM1548-SK1T1 | ACIN1 | p.Glu192Lys | missense_variant | MODERATE | 0 | 0 | 149 | 129 | . | . | . | . |
| PM1548-SK1T1 | PABPN1 | c.*5dupA | frameshift_variant&stop_retained_variant | HIGH | 0 | 0 | 96 | 92 | . | . | . | . |
| PM1548-SK1T1 | RNF31 | p.Ala9Gly | missense_variant | MODERATE | 0 | 0 | 34 | 104 | . | . | . | true |
| PM1548-SK1T1 | RNF31 | p.Val12Ala | missense_variant | MODERATE | 0 | 0 | 34 | 104 | . | . | . | true |
| PM1548-SK1T1 | LTB4R2 | p.Ala92Val | missense_variant | MODERATE | 0 | 0 | 72 | 173 | . | . | . | . |
| PM1548-SK1T1 | FOXG1 | p.Glu173Asp | missense_variant | MODERATE | 0 | 0 | 33 | 31 | . | . | . | . |
| PM1548-SK1T1 | CLEC14A | p.Ser404Asn | missense_variant | MODERATE | 0.112069 | 0 | 116 | 138 | . | . | . | . |
| PM1548-SK1T1 | KTN1 | p.Phe1182Val | missense_variant | MODERATE | 0.0447761 | 0 | 67 | 114 | true | . | . | . |
| PM1548-SK1T1 | KCNH5 | p.Arg333His | missense_variant | MODERATE | 0.203704 | 0 | 54 | 68 | . | . | . | . |
| PM1548-SK1T1 | ZBTB25 | p.His371Asn | missense_variant | MODERATE | 0 | 0 | 115 | 47 | . | . | . | . |
| PM1548-SK1T1 | HSPA2 | p.Gly138fs | frameshift_variant | HIGH | 0 | 0 | 199 | 148 | . | . | . | . |
| PM1548-SK1T1 | HSPA2 | p.Phe220Phe | synonymous_variant | LOW | 0 | 0 | 66 | 96 | . | . | . | . |
| PM1548-SK1T1 | FUT8 | p.Arg249Cys | missense_variant | MODERATE | 0.168539 | 0 | 178 | 316 | . | . | . | . |
| PM1548-SK1T1 | TMEM229B | p.Leu37Phe | missense_variant | MODERATE | 0 | 0 | 172 | 483 | . | . | . | . |
| PM1548-SK1T1 | ZFP36L1 | p.Gly271fs | frameshift_variant | HIGH | 0.0810811 | 0 | 185 | 217 | . | . | . | . |
| PM1548-SK1T1 | ELMSAN1 | p.Arg739Cys | missense_variant | MODERATE | 0.0465116 | 0 | 86 | 46 | . | . | . | . |
| PM1548-SK1T1 | ELMSAN1 | p.Gln36fs | frameshift_variant | HIGH | 0.25 | 0 | 80 | 43 | . | . | . | . |
| PM1548-SK1T1 | VRTN | p.Gly328fs | frameshift_variant | HIGH | 0 | 0 | 139 | 58 | . | . | . | . |
| PM1548-SK1T1 | GALC | p.Arg685His | missense_variant | MODERATE | 0.104478 | 0 | 67 | 31 | . | . | . | . |
| PM1548-SK1T1 | PTPN21 | p.Ile849fs | frameshift_variant | HIGH | 0.0581395 | 0 | 86 | 156 | . | . | . | . |
| PM1548-SK1T1 | ZC3H14 | p.Asn577Asp | missense_variant | MODERATE | 0 | 0 | 179 | 222 | . | . | . | . |
| PM1548-SK1T1 | CCDC88C | p.Lys818Glu | missense_variant | MODERATE | 0 | 0 | 120 | 69 | . | . | . | . |
| PM1548-SK1T1 | FBLN5 | p.Asp357Asn | missense_variant | MODERATE | 0.0131579 | 0 | 228 | 147 | . | . | . | . |
| PM1548-SK1T1 | UNC79 | p.Arg746Gln | missense_variant | MODERATE | 0.135593 | 0 | 118 | 30 | . | . | . | . |
| PM1548-SK1T1 | SYNE3 | p.Ile83Val | missense_variant | MODERATE | 0.0518519 | 0 | 135 | 217 | . | . | . | . |
| PM1548-SK1T1 | SETD3 | p.Phe388Leu | missense_variant | MODERATE | 0.0508475 | 0 | 59 | 32 | . | . | . | . |
| PM1548-SK1T1 | TECPR2 | p.Gln498fs | frameshift_variant | HIGH | 0.0294118 | 0 | 102 | 55 | . | . | . | . |
| PM1548-SK1T1 | BRF1 | p.Ala129Ser | missense_variant | MODERATE | 0 | 0 | 137 | 135 | . | . | . | . |
| PM1548-SK1T1 | CYFIP1 | p.Arg1085Leu | missense_variant | MODERATE | 0 | 0 | 162 | 334 | . | . | . | . |
| PM1548-SK1T1 | MAGEL2 | p.Ala11Val | missense_variant | MODERATE | 0 | 0 | 187 | 323 | . | . | . | . |
| PM1548-SK1T1 | PAR5 | n.2021_2022del | intron_variant | MODIFIER | 0.0178571 | 0 | 168 | 165 | . | . | . | . |
| PM1548-SK1T1 | UBE3A | p.Val133Ala | missense_variant | MODERATE | 0.0857143 | 0 | 70 | 79 | . | . | . | . |
| PM1548-SK1T1 | ACTC1 | p.Ala137Thr | missense_variant | MODERATE | 0.0103093 | 0 | 97 | 73 | . | . | . | . |
| PM1548-SK1T1 | SPRED1 | p.Arg332His | missense_variant | MODERATE | 0.0204082 | 0.0070922 | 98 | 141 | . | . | . | . |
| PM1548-SK1T1 | CASC5 | p.Asp178Gly | missense_variant | MODERATE | 0 | 0 | 95 | 58 | true | . | . | . |
| PM1548-SK1T1 | SPTBN5 | p.Arg1943His | missense_variant | MODERATE | 0 | 0 | 121 | 148 | . | . | . | . |
| PM1548-SK1T1 | SPTBN5 | p.Val1842Gly | missense_variant | MODERATE | 0.0697674 | 0 | 43 | 87 | . | . | . | . |
| PM1548-SK1T1 | GANC | p.Ala394Val | missense_variant | MODERATE | 0 | 0 | 170 | 106 | . | . | . | . |
| PM1548-SK1T1 | GANC | p.Gly404Asp | missense_variant | MODERATE | 0 | 0 | 161 | 105 | . | . | . | . |
| PM1548-SK1T1 | TTBK2 | p.Ala197Thr | missense_variant | MODERATE | 0 | 0 | 131 | 69 | . | . | . | . |
| PM1548-SK1T1 | CCNDBP1 | p.Asp237Asn | missense_variant | MODERATE | 0 | 0 | 96 | 44 | . | . | . | . |
| PM1548-SK1T1 | TGM7 | p.Ser305Phe | missense_variant | MODERATE | 0.181818 | 0 | 55 | 54 | . | . | . | . |
| PM1548-SK1T1 | ZSCAN29 | p.Arg33Trp | missense_variant | MODERATE | 0.132743 | 0 | 113 | 208 | . | . | . | . |
| PM1548-SK1T1 | SLC28A2 | p.Gly128Asp | missense_variant | MODERATE | 0.00543478 | 0 | 184 | 169 | . | . | . | . |
| PM1548-SK1T1 | MYO5A | p.Arg831His | missense_variant | MODERATE | 0.0612245 | 0 | 98 | 78 | true | . | . | . |
| PM1548-SK1T1 | TCF12 | p.Val183fs | frameshift_variant | HIGH | 0 | 0 | 168 | 200 | true | . | . | . |
| PM1548-SK1T1 | LIPC | p.Phe286fs | frameshift_variant | HIGH | 0.0434783 | 0 | 46 | 89 | . | . | . | . |
| PM1548-SK1T1 | OAZ2 | p.Ser92Ser | synonymous_variant | LOW | 0 | 0 | 127 | 163 | . | . | . | . |
| PM1548-SK1T1 | OAZ2 | p.Glu87Gly | missense_variant | MODERATE | 0 | 0 | 94 | 155 | . | . | . | . |
| PM1548-SK1T1 | OAZ2 | p.Glu84Val | missense_variant | MODERATE | 0 | 0 | 209 | 248 | . | . | . | . |
| PM1548-SK1T1 | OAZ2 | p.Val79Gly | missense_variant | MODERATE | 0 | 0 | 202 | 233 | . | . | . | . |
| PM1548-SK1T1 | SPG21 | p.Ala282Val | missense_variant | MODERATE | 0.0384615 | 0 | 52 | 32 | . | . | . | . |
| PM1548-SK1T1 | CILP | p.Asn930Asn | synonymous_variant | LOW | 0 | 0.00877193 | 173 | 114 | . | . | . | . |
| PM1548-SK1T1 | IGDCC3 | p.Pro358Leu | missense_variant | MODERATE | 0.104712 | 0 | 191 | 237 | . | . | . | . |
| PM1548-SK1T1 | NOX5 | p.His470fs | frameshift_variant | HIGH | 0 | 0 | 140 | 164 | . | . | . | . |
| PM1548-SK1T1 | MYO9A | p.Cys1767Ser | missense_variant | MODERATE | 0 | 0 | 178 | 186 | . | . | . | . |
| PM1548-SK1T1 | NPTN | p.Arg334Trp | missense_variant | MODERATE | 0.115702 | 0 | 121 | 86 | . | . | . | . |
| PM1548-SK1T1 | ISLR2 | p.Val74Ala | missense_variant | MODERATE | 0.0634921 | 0 | 126 | 333 | . | . | . | . |
| PM1548-SK1T1 | C15orf39 | p.Ala475Val | missense_variant | MODERATE | 0.208835 | 0.00393701 | 249 | 254 | . | . | . | . |
| PM1548-SK1T1 | ETFA | p.Thr45Arg | missense_variant | MODERATE | 0 | 0 | 137 | 214 | . | . | . | . |
| PM1548-SK1T1 | ETFA | p.Thr45Ala | missense_variant | MODERATE | 0.00729927 | 0 | 137 | 214 | . | . | . | . |
| PM1548-SK1T1 | PSTPIP1 | p.Ala280fs | frameshift_variant&splice_region_variant | HIGH | 0.0711462 | 0 | 253 | 442 | . | . | . | . |
| PM1548-SK1T1 | PEAK1 | p.Ala36Thr | missense_variant | MODERATE | 0 | 0 | 215 | 189 | . | . | . | . |
| PM1548-SK1T1 | CHRNB4 | p.Gly261Asp | missense_variant | MODERATE | 0.0576923 | 0 | 156 | 56 | . | . | . | . |
| PM1548-SK1T1 | MESDC1 | p.Ser224Asn | missense_variant | MODERATE | 0.219178 | 0 | 73 | 161 | . | . | . | . |
| PM1548-SK1T1 | IL16 | p.Thr417Met | missense_variant | MODERATE | 0.104651 | 0 | 86 | 34 | . | . | . | . |
| PM1548-SK1T1 | BTBD1 | p.Arg236Gln | missense_variant | MODERATE | 0.14881 | 0 | 168 | 74 | . | . | . | . |
| PM1548-SK1T1 | ZNF592 | p.Ala1232Val | missense_variant | MODERATE | 0.114286 | 0 | 35 | 35 | . | . | . | . |
| PM1548-SK1T1 | AEN | p.Ala194Val | missense_variant | MODERATE | 0.119718 | 0 | 142 | 121 | . | . | . | . |
| PM1548-SK1T1 | IQGAP1 | p.Met1231fs | frameshift_variant | HIGH | 0.0366972 | 0 | 109 | 170 | . | . | . | . |
| PM1548-SK1T1 | FES | c.1531-47G>A | intron_variant | MODIFIER | 0.0405405 | 0 | 74 | 35 | . | . | . | . |

| | | | | | | | | | | | | |
|---|---|---|---|---|---|---|---|---|---|---|---|---|
| PM1548-SK1T1 | MAN2A2 | p.Pro139Leu | missense_variant | MODERATE | 0.0357143 | 0 | 364 | 493 | . | . | . | . |
| PM1548-SK1T1 | RCCD1 | p.Arg87Ser | missense_variant | MODERATE | 0.103448 | 0 | 58 | 63 | . | . | . | . |
| PM1548-SK1T1 | ADAMTS17 | p.Cys921Tyr | missense_variant | MODERATE | 0 | 0 | 157 | 58 | . | . | . | . |
| PM1548-SK1T1 | WDR90 | p.Ala1738Thr | missense_variant | MODERATE | 0.121053 | 0 | 190 | 185 | . | . | . | . |
| PM1548-SK1T1 | RPUSD1 | p.Arg255Gln | missense_variant | MODERATE | 0.0938776 | 0 | 245 | 202 | . | . | . | . |
| PM1548-SK1T1 | SOX8 | p.Pro11Thr | missense_variant | MODERATE | 0.111111 | 0 | 36 | 62 | . | . | . | . |
| PM1548-SK1T1 | SOX8 | p.Asp27Asn | missense_variant | MODERATE | 0.114286 | 0 | 35 | 120 | . | . | . | . |
| PM1548-SK1T1 | CACNA1H | p.Arg1892His | missense_variant | MODERATE | 0 | 0 | 218 | 231 | . | . | . | . |
| PM1548-SK1T1 | BAIAP3 | p.Ala740Asp | missense_variant | MODERATE | 0.133333 | 0 | 30 | 90 | . | . | . | . |
| PM1548-SK1T1 | IFT140 | p.Ala1330Val | missense_variant | MODERATE | 0.111111 | 0 | 72 | 187 | . | . | . | . |
| PM1548-SK1T1 | NME3 | p.Ala20Ser | missense_variant | MODERATE | 0.121212 | 0 | 33 | 162 | . | . | . | . |
| PM1548-SK1T1 | PRSS33 | p.Leu174fs | frameshift_variant | HIGH | 0.128834 | 0 | 163 | 197 | . | . | . | . |
| PM1548-SK1T1 | CCDC64B | p.Arg144Gln | missense_variant | MODERATE | 0.111675 | 0 | 197 | 390 | . | . | . | . |
| PM1548-SK1T1 | CREBBP | p.Gly1479Val | missense_variant | MODERATE | 0 | 0 | 212 | 193 | true | . | true | . |
| PM1548-SK1T1 | CORO7 | p.Thr230Ala | missense_variant | MODERATE | 0 | 0 | 116 | 172 | . | . | . | . |
| PM1548-SK1T1 | GLYR1 | p.Ser148Thr | missense_variant | MODERATE | 0 | 0 | 58 | 42 | . | . | . | . |
| PM1548-SK1T1 | TVP23A | p.Phe140fs | frameshift_variant | HIGH | 0.0186335 | 0 | 161 | 183 | . | . | . | . |
| PM1548-SK1T1 | CIITA | p.Arg580His | missense_variant | MODERATE | 0.0788177 | 0 | 203 | 337 | true | . | . | . |
| PM1548-SK1T1 | RMI2 | p.Ile130Val | missense_variant | MODERATE | 0 | 0 | 147 | 171 | true | . | . | . |
| PM1548-SK1T1 | TNFRSF17 | p.Pro145Gln | missense_variant | MODERATE | 0 | 0 | 56 | 38 | true | . | . | . |
| PM1548-SK1T1 | CDR2 | p.Gln130Arg | missense_variant | MODERATE | 0.0387597 | 0 | 129 | 85 | . | . | . | . |
| PM1548-SK1T1 | USP31 | p.Gly1267Arg | missense_variant | MODERATE | 0.172414 | 0 | 58 | 32 | . | . | . | . |
| PM1548-SK1T1 | COG7 | p.Pro67Ala | missense_variant | MODERATE | 0.0493827 | 0 | 81 | 71 | . | . | . | . |
| PM1548-SK1T1 | GTF3C1 | p.Glu1761Gly | missense_variant | MODERATE | 0 | 0 | 182 | 99 | . | . | . | . |
| PM1548-SK1T1 | ATXN2L | c.3140-64G>T | intron_variant | MODIFIER | 0 | 0 | 139 | 124 | . | . | . | . |
| PM1548-SK1T1 | ATP2A1 | p.Gly229Ser | missense_variant | MODERATE | 0 | 0 | 182 | 168 | . | . | . | . |
| PM1548-SK1T1 | KIF22 | p.Ile592fs | frameshift_variant | HIGH | 0.11236 | 0 | 89 | 127 | . | . | . | . |
| PM1548-SK1T1 | PRR14 | p.Arg96Gln | missense_variant | MODERATE | 0.0220588 | 0 | 136 | 273 | . | . | . | . |
| PM1548-SK1T1 | ZNF629 | p.Asp434Asn | missense_variant | MODERATE | 0.0212766 | 0 | 94 | 34 | . | . | . | . |
| PM1548-SK1T1 | HSD3B7 | p.Val361Ile | missense_variant | MODERATE | 0.0738636 | 0 | 176 | 165 | . | . | . | . |
| PM1548-SK1T1 | KAT8 | p.Val438Ala | missense_variant | MODERATE | 0.102273 | 0 | 88 | 67 | . | . | . | . |
| PM1548-SK1T1 | FUS | c.1295+1G>A | splice_donor_variant&intron_variant | HIGH | 0.136364 | 0 | 242 | 234 | true | . | . | . |
| PM1548-SK1T1 | PHKB | p.Tyr52His | missense_variant | MODERATE | 0.109195 | 0 | 174 | 155 | . | . | . | . |
| PM1548-SK1T1 | LONP2 | p.Gly681Cys | missense_variant | MODERATE | 0.0308642 | 0 | 162 | 276 | . | . | . | . |
| PM1548-SK1T1 | N4BP1 | p.Arg640Cys | missense_variant | MODERATE | 0 | 0 | 113 | 94 | . | . | . | . |
| PM1548-SK1T1 | SALL1 | p.Ser1318Asn | missense_variant | MODERATE | 0.0222222 | 0 | 180 | 197 | . | . | . | . |
| PM1548-SK1T1 | SALL1 | p.His756Asp | missense_variant | MODERATE | 0.0638298 | 0 | 47 | 53 | . | . | . | . |
| PM1548-SK1T1 | LPCAT2 | c.1216-7923dup | intron_variant | MODIFIER | 0.030303 | 0 | 99 | 56 | . | . | . | . |
| PM1548-SK1T1 | SLC6A2 | p.Gly591Asp | missense_variant | MODERATE | 0.171521 | 0 | 309 | 342 | . | . | . | . |
| PM1548-SK1T1 | MMP15 | p.Ala603Asp | missense_variant | MODERATE | 0 | 0 | 325 | 388 | . | . | . | . |
| PM1548-SK1T1 | CDH11 | p.Ala166Thr | missense_variant | MODERATE | 0 | 0 | 146 | 177 | true | . | . | . |
| PM1548-SK1T1 | CDH11 | p.Thr163Ile | missense_variant | MODERATE | 0 | 0 | 146 | 177 | true | . | . | . |
| PM1548-SK1T1 | CDH11 | p.His161His | synonymous_variant | LOW | 0 | 0 | 137 | 175 | true | . | . | . |
| PM1548-SK1T1 | CDH11 | p.Val149Asp | missense_variant | MODERATE | 0 | 0 | 168 | 64 | true | . | . | . |
| PM1548-SK1T1 | CDH11 | p.Val147Asp | missense_variant | MODERATE | 0 | 0 | 129 | 55 | true | . | . | . |
| PM1548-SK1T1 | CDH11 | p.Glu140Gly | missense_variant | MODERATE | 0 | 0 | 135 | 80 | true | . | . | . |
| PM1548-SK1T1 | CDH11 | p.Arg137Gln | missense_variant | MODERATE | 0.0149254 | 0 | 134 | 79 | true | . | . | . |
| PM1548-SK1T1 | TK2 | p.Asn114Asn | synonymous_variant | LOW | 0.0289855 | 0 | 138 | 102 | . | . | . | . |
| PM1548-SK1T1 | B3GNT9 | p.Asp221Asn | missense_variant | MODERATE | 0 | 0 | 117 | 208 | . | . | . | . |
| PM1548-SK1T1 | LRRC29 | p.Gln205His | missense_variant | MODERATE | 0 | 0 | 169 | 214 | . | . | . | . |
| PM1548-SK1T1 | KCTD19 | p.Gly313Arg | missense_variant | MODERATE | 0.0396825 | 0 | 126 | 109 | . | . | . | . |
| PM1548-SK1T1 | TPPP3 | p.Ser49Ala | missense_variant | MODERATE | 0 | 0 | 139 | 59 | . | . | . | . |
| PM1548-SK1T1 | CTCF | p.Arg283His | missense_variant | MODERATE | 0.135135 | 0 | 185 | 194 | true | true | true | . |
| PM1548-SK1T1 | CDH3 | p.Asp264Glu | missense_variant | MODERATE | 0 | 0 | 277 | 403 | . | . | . | . |
| PM1548-SK1T1 | WWP2 | p.Arg633Gln | missense_variant | MODERATE | 0.183486 | 0 | 109 | 88 | . | . | . | . |
| PM1548-SK1T1 | PMFBP1 | p.Glu205fs | frameshift_variant | HIGH | 0.0664062 | 0 | 256 | 212 | . | . | . | . |
| PM1548-SK1T1 | ZFHX3 | p.Ala1099Ser | missense_variant | MODERATE | 0 | 0 | 114 | 40 | true | . | true | true |
| PM1548-SK1T1 | RFWD3 | p.Val155Ala | missense_variant | MODERATE | 0 | 0 | 95 | 69 | . | . | . | . |
| PM1548-SK1T1 | PKD1L2 | n.81164209C>T | intragenic_variant | MODIFIER | 0.039604 | 0 | 101 | 78 | . | . | . | . |
| PM1548-SK1T1 | TAF1C | p.Pro707fs | frameshift_variant | HIGH | 0.0277778 | 0 | 144 | 205 | . | . | . | . |
| PM1548-SK1T1 | PIEZO1 | p.Pro1061Leu | missense_variant | MODERATE | 0 | 0 | 30 | 23 | . | . | . | . |
| PM1548-SK1T1 | PIEZO1 | p.Met1002Ile | missense_variant | MODERATE | 0.136364 | 0 | 44 | 33 | . | . | . | . |
| PM1548-SK1T1 | GALNS | p.Thr109Thr | synonymous_variant | LOW | 0 | 0 | 73 | 187 | . | . | . | . |
| PM1548-SK1T1 | ANKRD11 | p.Asp2130Asn | missense_variant | MODERATE | 0.121951 | 0 | 41 | 42 | . | . | . | . |
| PM1548-SK1T1 | GEMIN4 | p.Arg229Gln | missense_variant | MODERATE | 0.105263 | 0 | 57 | 146 | . | . | . | . |
| PM1548-SK1T1 | RNMTL1 | p.Met143fs | frameshift_variant | HIGH | 0.0363636 | 0 | 55 | 36 | . | . | . | . |
| PM1548-SK1T1 | PRPF8 | p.Arg1057Gln | missense_variant | MODERATE | 0.0216216 | 0 | 185 | 146 | . | . | . | . |
| PM1548-SK1T1 | HIC1 | p.Pro693Leu | missense_variant | MODERATE | 0.129032 | 0 | 31 | 77 | . | . | . | . |
| PM1548-SK1T1 | SMG6 | p.Arg603His | missense_variant | MODERATE | 0 | 0.00434783 | 235 | 230 | . | . | . | . |
| PM1548-SK1T1 | CTNS | p.Asp74Asn | missense_variant | MODERATE | 0.171875 | 0.00314465 | 320 | 318 | . | . | . | . |
| PM1548-SK1T1 | ITGAE | p.Arg113Gln | missense_variant | MODERATE | 0.131034 | 0 | 145 | 94 | . | . | . | . |
| PM1548-SK1T1 | ZZEF1 | p.Arg194Trp | missense_variant | MODERATE | 0.103448 | 0 | 58 | 95 | . | . | . | . |
| PM1548-SK1T1 | KIF1C | p.Ser511Phe | missense_variant | MODERATE | 0.117647 | 0 | 17 | 140 | . | . | . | . |
| PM1548-SK1T1 | USP6 | p.Pro181Leu | missense_variant&splice_region_variant | MODERATE | 0.0727273 | 0 | 55 | 73 | true | . | . | . |
| PM1548-SK1T1 | DVL2 | p.Arg442Cys | missense_variant | MODERATE | 0.134831 | 0 | 89 | 34 | . | . | . | . |
| PM1548-SK1T1 | EIF5A | p.Thr168Met | missense_variant | MODERATE | 0.0316456 | 0 | 158 | 85 | . | . | true | . |
| PM1548-SK1T1 | DNAH2 | p.Gly4241fs | frameshift_variant | HIGH | 0.15 | 0 | 120 | 74 | . | . | . | . |
| PM1548-SK1T1 | ALOX12B | p.Ala355Thr | missense_variant | MODERATE | 0.065 | 0 | 200 | 296 | . | . | . | . |
| PM1548-SK1T1 | GLP2R | p.Ala515Thr | missense_variant | MODERATE | 0.231481 | 0.00334448 | 216 | 299 | . | . | . | . |
| PM1548-SK1T1 | MYH13 | p.Glu984Lys | missense_variant | MODERATE | 0.146667 | 0 | 75 | 72 | . | . | . | . |
| PM1548-SK1T1 | MYH8 | p.Gly186Arg | missense_variant | MODERATE | 0.139896 | 0 | 193 | 81 | . | . | . | . |
| PM1548-SK1T1 | MYH4 | p.Tyr424Cys | missense_variant | MODERATE | 0.00840336 | 0 | 119 | 298 | . | . | . | . |
| PM1548-SK1T1 | DNAH9 | p.Arg1542Met | missense_variant | MODERATE | 0.00303951 | 0 | 329 | 342 | . | . | . | . |
| PM1548-SK1T1 | DNAH9 | p.Ala2514Thr | missense_variant | MODERATE | 0.00666667 | 0 | 150 | 101 | . | . | . | . |
| PM1548-SK1T1 | ARHGAP44 | p.Ala495Thr | missense_variant | MODERATE | 0.0142857 | 0 | 70 | 53 | . | . | . | . |
| PM1548-SK1T1 | FLCN | p.His429fs | frameshift_variant | HIGH | 0 | 0 | 161 | 269 | true | . | . | . |
| PM1548-SK1T1 | FLCN | p.Thr174Ala | missense_variant | MODERATE | 0 | 0 | 93 | 227 | true | . | . | . |
| PM1548-SK1T1 | TOM1L2 | p.Met482fs | frameshift_variant | HIGH | 0 | 0 | 82 | 63 | . | . | . | . |
| PM1548-SK1T1 | KIAA0100 | p.Arg689* | stop_gained | HIGH | 0 | 0 | 209 | 219 | . | . | . | . |
| PM1548-SK1T1 | SUPT6H | p.Ile1356Leu | missense_variant | MODERATE | 0 | 0.00465116 | 115 | 215 | . | . | . | . |
| PM1548-SK1T1 | PHF12 | p.Leu393fs | frameshift_variant | HIGH | 0.00684932 | 0 | 146 | 61 | . | . | . | . |
| PM1548-SK1T1 | SEZ6 | p.Gly660Asp | missense_variant | MODERATE | 0.0757576 | 0 | 132 | 83 | . | . | . | . |
| PM1548-SK1T1 | ANKRD13B | p.Arg233Gln | missense_variant | MODERATE | 0.131034 | 0 | 145 | 97 | . | . | . | . |
| PM1548-SK1T1 | FNDC8 | p.Asn57Ser | missense_variant | MODERATE | 0.0208333 | 0 | 96 | 150 | . | . | . | . |
| PM1548-SK1T1 | NLE1 | p.Val10Ala | missense_variant | MODERATE | 0 | 0 | 93 | 97 | . | . | . | . |
| PM1548-SK1T1 | SRCIN1 | p.Leu460Phe | missense_variant | MODERATE | 0 | 0 | 52 | 53 | . | . | . | . |
| PM1548-SK1T1 | PCGF2 | p.Asp157Asn | missense_variant | MODERATE | 0.0350877 | 0 | 114 | 53 | . | . | . | . |
| PM1548-SK1T1 | STAC2 | p.Cys141Cys | synonymous_variant | LOW | 0 | 0 | 204 | 250 | . | . | . | . |
| PM1548-SK1T1 | GRB7 | p.Arg552Gln | missense_variant | MODERATE | 0.115385 | 0 | 78 | 159 | . | . | . | . |
| PM1548-SK1T1 | NR1D1 | p.Pro593Pro | synonymous_variant | LOW | 0 | 0.00961538 | 77 | 104 | . | . | . | . |
| PM1548-SK1T1 | MSL1 | p.Leu372Ile | missense_variant | MODERATE | 0 | 0 | 63 | 43 | . | . | . | . |
| PM1548-SK1T1 | WIPF2 | p.Ala39Thr | missense_variant | MODERATE | 0.178082 | 0 | 73 | 66 | . | . | . | . |
| PM1548-SK1T1 | KRT25 | p.Ala91Thr | missense_variant | MODERATE | 0.08 | 0 | 150 | 148 | . | . | . | . |
| PM1548-SK1T1 | KRT15 | p.Arg398Trp | missense_variant | MODERATE | 0.140625 | 0 | 128 | 53 | . | . | . | . |
| PM1548-SK1T1 | KRT16 | p.Gly243Ser | missense_variant | MODERATE | 0 | 0 | 113 | 130 | . | . | . | . |
| PM1548-SK1T1 | DHX58 | p.Arg224Cys | missense_variant | MODERATE | 0.0233918 | 0 | 171 | 91 | . | . | . | . |
| PM1548-SK1T1 | PLEKHH3 | p.Pro318Thr | missense_variant | MODERATE | 0.015625 | 0 | 128 | 121 | . | . | . | . |
| PM1548-SK1T1 | ETV4 | p.Gln13Arg | missense_variant | MODERATE | 0 | 0 | 133 | 215 | true | . | . | . |
| PM1548-SK1T1 | MPP3 | p.Arg265Gln | missense_variant | MODERATE | 0.0333333 | 0 | 30 | 79 | . | . | . | . |
| PM1548-SK1T1 | ATXN7L3 | p.Ile246Phe | missense_variant | MODERATE | 0.0227273 | 0 | 44 | 37 | . | . | . | . |
| PM1548-SK1T1 | ITGA2B | p.Asn330Ser | missense_variant | MODERATE | 0 | 0 | 52 | 60 | . | . | . | . |
| PM1548-SK1T1 | ARHGAP27 | p.Leu247fs | frameshift_variant | HIGH | 0 | 0 | 76 | 82 | . | . | . | . |
| PM1548-SK1T1 | WNT3 | p.Ala223Ser | missense_variant | MODERATE | 0 | 0 | 183 | 125 | . | . | . | . |
| PM1548-SK1T1 | KPNB1 | p.Ile686Leu | missense_variant | MODERATE | 0 | 0 | 210 | 143 | . | . | . | . |
| PM1548-SK1T1 | KPNB1 | p.Ile686Met | missense_variant | MODERATE | 0.0047619 | 0 | 210 | 143 | . | . | . | . |
| PM1548-SK1T1 | SP6 | p.Ala138Thr | missense_variant | MODERATE | 0.11 | 0 | 100 | 220 | . | . | . | . |
| PM1548-SK1T1 | PPP1R9B | p.Glu24Glu | synonymous_variant | LOW | 0 | 0 | 6 | 79 | . | . | . | . |
| PM1548-SK1T1 | CUEDC1 | p.Arg71Cys | missense_variant | MODERATE | 0.0204082 | 0 | 196 | 109 | . | . | . | . |
| PM1548-SK1T1 | LPO | p.Arg82Cys | missense_variant | MODERATE | 0.0150376 | 0 | 133 | 404 | . | . | . | . |
| PM1548-SK1T1 | RNF43 | p.Gly659fs | frameshift_variant | HIGH | 0.15625 | 0 | 64 | 76 | true | . | . | true |
| PM1548-SK1T1 | TBX2 | p.Leu460fs | frameshift_variant | HIGH | 0.103226 | 0 | 155 | 331 | . | . | . | . |
| PM1548-SK1T1 | MRC2 | p.Arg490Cys | missense_variant | MODERATE | 0.138889 | 0 | 108 | 131 | . | . | . | . |
| PM1548-SK1T1 | CCDC47 | p.Ser303Pro | missense_variant | MODERATE | 0 | 0 | 97 | 131 | . | . | . | . |
| PM1548-SK1T1 | FTSJ3 | p.Met493Ile | missense_variant | MODERATE | 0.0181818 | 0 | 55 | 61 | . | . | . | . |
| PM1548-SK1T1 | ABCA10 | p.Tyr1399His | missense_variant | MODERATE | 0.0327869 | 0 | 244 | 336 | . | . | . | . |
| PM1548-SK1T1 | SLC16A5 | p.Gly93Cys | missense_variant | MODERATE | 0.116279 | 0 | 86 | 59 | . | . | . | . |
| PM1548-SK1T1 | ITGB4 | p.Pro305Leu | missense_variant | MODERATE | 0.0534351 | 0 | 131 | 78 | . | . | . | . |

| | | | | | | | | | | | | |
|---|---|---|---|---|---|---|---|---|---|---|---|---|
| PM1548-SK1T1 | ITGB4 | p.Pro723Ser | missense_variant | MODERATE | 0 | 0 | 172 | 167 | . | . | . | . |
| PM1548-SK1T1 | H3F3B | p.Ala8Asp | missense_variant | MODERATE | 0 | 0 | 43 | 88 | true | . | . | . |
| PM1548-SK1T1 | EVPL | p.Arg266Trp | missense_variant | MODERATE | 0 | 0 | 13 | 157 | . | . | . | . |
| PM1548-SK1T1 | RHBDF2 | p.Ala354Thr | missense_variant | MODERATE | 0.137931 | 0 | 87 | 138 | . | . | . | . |
| PM1548-SK1T1 | TNRC6C | p.Arg1160His | missense_variant | MODERATE | 0.0707071 | 0 | 99 | 185 | . | . | . | . |
| PM1548-SK1T1 | TMC6 | p.His31His | synonymous_variant | LOW | 0.106762 | 0 | 281 | 491 | . | . | . | . |
| PM1548-SK1T1 | CCDC40 | p.Gly929Ser | missense_variant | MODERATE | 0.1 | 0 | 40 | 34 | . | . | . | . |
| PM1548-SK1T1 | GAA | p.Gly828Asp | missense_variant&splice_region_variant | MODERATE | 0 | 0 | 172 | 76 | . | . | . | . |
| PM1548-SK1T1 | ENDOV | p.Pro259Leu | missense_variant | MODERATE | 0.0533333 | 0 | 75 | 83 | . | . | . | . |
| PM1548-SK1T1 | RPTOR | p.Arg899* | stop_gained | HIGH | 0.140187 | 0 | 107 | 71 | . | . | . | . |
| PM1548-SK1T1 | SIRT7 | p.Thr181Met | missense_variant | MODERATE | 0 | 0 | 201 | 217 | . | . | . | . |
| PM1548-SK1T1 | OGFOD3 | p.Ser127Ser | splice_region_variant&synonymous_variant | LOW | 0.0944444 | 0 | 180 | 110 | . | . | . | . |
| PM1548-SK1T1 | TXNDC2 | p.Ile284Leu | missense_variant | MODERATE | 0 | 0 | 191 | 275 | . | . | . | . |
| PM1548-SK1T1 | TXNDC2 | p.Ser285Pro | missense_variant | MODERATE | 0.0052356 | 0 | 191 | 275 | . | . | . | . |
| PM1548-SK1T1 | TXNDC2 | p.Pro288Leu | missense_variant | MODERATE | 0 | 0.00816327 | 175 | 245 | . | . | . | . |
| PM1548-SK1T1 | ANKRD62 | p.Gln745His | missense_variant | MODERATE | 0.0294118 | 0 | 136 | 165 | . | . | . | . |
| PM1548-SK1T1 | ZNF521 | p.Cys754Tyr | missense_variant | MODERATE | 0 | 0 | 141 | 57 | true | . | . | . |
| PM1548-SK1T1 | KCTD1 | p.Asn124His | missense_variant | MODERATE | 0 | 0 | 25 | 69 | . | . | . | . |
| PM1548-SK1T1 | DSG4 | p.Thr1001Met | missense_variant | MODERATE | 0.0235294 | 0 | 85 | 104 | . | . | . | . |
| PM1548-SK1T1 | CCDC178 | p.Arg827Met | missense_variant | MODERATE | 0.121053 | 0 | 190 | 192 | . | . | . | . |
| PM1548-SK1T1 | INO80C | p.Gly216Ser | missense_variant | MODERATE | 0.113924 | 0 | 79 | 89 | . | . | . | . |
| PM1548-SK1T1 | SETBP1 | p.Glu244Asp | missense_variant | MODERATE | 0 | 0 | 63 | 133 | true | true | . | . |
| PM1548-SK1T1 | SETBP1 | p.Ile251Val | missense_variant | MODERATE | 0 | 0 | 82 | 201 | true | true | . | . |
| PM1548-SK1T1 | SETBP1 | p.Ala733Thr | missense_variant | MODERATE | 0 | 0 | 120 | 315 | true | true | . | . |
| PM1548-SK1T1 | SIGLEC15 | p.Ala96Thr | missense_variant | MODERATE | 0.0714286 | 0 | 28 | 175 | . | . | . | . |
| PM1548-SK1T1 | SIGLEC15 | p.Arg99Pro | missense_variant | MODERATE | 0.0714286 | 0 | 28 | 176 | . | . | . | . |
| PM1548-SK1T1 | SKOR2 | p.Asp736Glu | missense_variant | MODERATE | 0 | 0 | 72 | 157 | . | . | . | . |
| PM1548-SK1T1 | SKOR2 | p.Gly720Glu | missense_variant | MODERATE | 0 | 0 | 60 | 134 | . | . | . | . |
| PM1548-SK1T1 | SMAD7 | p.Pro55Ala | missense_variant | MODERATE | 0 | 0 | 64 | 32 | . | . | . | . |
| PM1548-SK1T1 | DSEL | p.Arg1009His | missense_variant | MODERATE | 0.134146 | 0 | 82 | 67 | . | . | . | . |
| PM1548-SK1T1 | FAM69C | p.Gly48Cys | missense_variant | MODERATE | 0 | 0 | 74 | 170 | . | . | . | . |
| PM1548-SK1T1 | TSHZ1 | p.Ala570Val | missense_variant | MODERATE | 0.147541 | 0 | 183 | 278 | . | . | . | . |
| PM1548-SK1T1 | TSHZ1 | p.Tyr775Cys | missense_variant | MODERATE | 0.109375 | 0 | 64 | 122 | . | . | . | . |
| PM1548-SK1T1 | ZNF516 | p.Gly232del | disruptive_inframe_deletion | MODERATE | 0 | 0 | 138 | 279 | . | . | . | . |
| PM1548-SK1T1 | ZNF516 | p.Pro229_Gly23( | inframe_insertion | MODERATE | 0 | 0 | 130 | 280 | . | . | . | . |
| PM1548-SK1T1 | GALR1 | p.Leu46Met | missense_variant | MODERATE | 0 | 0 | 64 | 102 | . | . | . | . |
| PM1548-SK1T1 | GALR1 | p.Lys62Lys | synonymous_variant | LOW | 0 | 0 | 143 | 218 | . | . | . | . |
| PM1548-SK1T1 | SALL3 | p.Pro1105fs | frameshift_variant | HIGH | 0.148148 | 0 | 108 | 197 | . | . | . | . |
| PM1548-SK1T1 | PPAP2C | n.291320T>C | intragenic_variant | MODIFIER | 0.019802 | 0 | 101 | 60 | . | . | . | . |
| PM1548-SK1T1 | LPPR3 | p.Gly646Ser | missense_variant | MODERATE | 0 | 0 | 12 | 120 | . | . | . | . |
| PM1548-SK1T1 | CFD | p.Arg199Cys | missense_variant | MODERATE | 0.02 | 0 | 200 | 193 | . | . | . | . |
| PM1548-SK1T1 | ARID3A | p.Arg503Cys | missense_variant | MODERATE | 0 | 0 | 64 | 124 | . | . | . | . |
| PM1548-SK1T1 | APC2 | p.Asp501Asn | missense_variant | MODERATE | 0.22695 | 0 | 141 | 282 | . | . | . | . |
| PM1548-SK1T1 | PCSK4 | p.Gly57Asp | missense_variant | MODERATE | 0.00505051 | 0 | 198 | 340 | . | . | . | . |
| PM1548-SK1T1 | ADAMTSL5 | p.Ala343fs | frameshift_variant | HIGH | 0.044335 | 0 | 203 | 366 | . | . | . | . |
| PM1548-SK1T1 | ADAMTSL5 | p.Pro341Ala | missense_variant | MODERATE | 0.0231214 | 0 | 173 | 313 | . | . | . | . |
| PM1548-SK1T1 | MEX3D | p.Gly262Asp | missense_variant | MODERATE | 0.0625 | 0 | 32 | 217 | . | . | . | . |
| PM1548-SK1T1 | TCF3 | p.Gly247fs | frameshift_variant | HIGH | 0.104651 | 0 | 86 | 79 | true | . | . | . |
| PM1548-SK1T1 | C19orf35 | p.Arg325Cys | missense_variant | MODERATE | 0.119403 | 0 | 67 | 53 | . | . | . | . |
| PM1548-SK1T1 | NCLN | p.Thr70Met | missense_variant | MODERATE | 0.0869565 | 0 | 92 | 140 | . | . | . | . |
| PM1548-SK1T1 | DOHH | p.Arg286Trp | missense_variant | MODERATE | 0.0136054 | 0.0136054 | 147 | 147 | . | . | . | . |
| PM1548-SK1T1 | EEF2 | p.Pro738Ser | missense_variant | MODERATE | 0.122807 | 0 | 57 | 230 | . | . | . | . |
| PM1548-SK1T1 | PIAS4 | p.Tyr265Tyr | synonymous_variant | LOW | 0.0196078 | 0 | 102 | 240 | . | . | . | . |
| PM1548-SK1T1 | MAP2K2 | p.Lys51Arg | missense_variant | MODERATE | 0 | 0 | 134 | 75 | true | . | . | . |
| PM1548-SK1T1 | MAP2K2 | p.Lys51Gln | missense_variant | MODERATE | 0 | 0 | 134 | 75 | true | . | . | . |
| PM1548-SK1T1 | CREB3L3 | p.Val140Ala | missense_variant | MODERATE | 0.113636 | 0 | 132 | 204 | . | . | . | . |
| PM1548-SK1T1 | UBXN6 | p.Thr337Met | missense_variant | MODERATE | 0.125 | 0 | 96 | 93 | . | . | . | . |
| PM1548-SK1T1 | PLIN4 | p.Val592Met | missense_variant | MODERATE | 0 | 0 | 116 | 40 | . | . | . | . |
| PM1548-SK1T1 | PLIN5 | p.Thr456Ile | missense_variant | MODERATE | 0.114286 | 0 | 35 | 37 | . | . | . | . |
| PM1548-SK1T1 | AC005594.3 | n.381G>A | non_coding_exon_variant | MODIFIER | 0.0855615 | 0.0037037 | 187 | 270 | . | . | . | . |
| PM1548-SK1T1 | DPP9 | p.Arg125Trp | missense_variant | MODERATE | 0 | 0 | 55 | 40 | . | . | . | . |
| PM1548-SK1T1 | KDM4B | p.Arg618Cys | missense_variant | MODERATE | 0.103448 | 0 | 58 | 167 | . | . | . | . |
| PM1548-SK1T1 | KDM4B | p.Glu1013Lys | missense_variant | MODERATE | 0.0190476 | 0 | 105 | 113 | . | . | . | . |
| PM1548-SK1T1 | DENND1C | p.Thr165fs | frameshift_variant | HIGH | 0.123711 | 0 | 97 | 34 | . | . | . | . |
| PM1548-SK1T1 | PEX11G | p.Arg124Trp | missense_variant | MODERATE | 0.142857 | 0 | 77 | 176 | . | . | . | . |
| PM1548-SK1T1 | CTD-3193O13.9 | p.Thr838Thr | synonymous_variant | LOW | 0.134615 | 0 | 156 | 174 | . | . | . | . |
| PM1548-SK1T1 | COL5A3 | p.Glu1677Lys | missense_variant | MODERATE | 0.0447761 | 0 | 134 | 264 | . | . | true | true |
| PM1548-SK1T1 | ZNF653 | p.Asp247Gly | missense_variant | MODERATE | 0 | 0 | 104 | 166 | . | . | . | . |
| PM1548-SK1T1 | ELOF1 | p.Arg41Cys | missense_variant | MODERATE | 0.182482 | 0 | 137 | 108 | . | . | . | . |
| PM1548-SK1T1 | MRI1 | p.His330fs | frameshift_variant | HIGH | 0.155039 | 0 | 129 | 82 | . | . | . | . |
| PM1548-SK1T1 | RFX1 | p.Val600Ile | missense_variant | MODERATE | 0 | 0 | 199 | 347 | . | . | . | . |
| PM1548-SK1T1 | PRKACA | p.Ala219Val | missense_variant | MODERATE | 0.111111 | 0 | 45 | 150 | true | . | . | . |
| PM1548-SK1T1 | TECR | p.Val85Met | missense_variant | MODERATE | 0 | 0.00641026 | 85 | 156 | . | . | . | . |
| PM1548-SK1T1 | EMR2 | p.Pro294Thr | missense_variant | MODERATE | 0.0108108 | 0 | 185 | 117 | . | . | . | . |
| PM1548-SK1T1 | EMR2 | p.Tyr292Ser | missense_variant | MODERATE | 0.0108696 | 0 | 184 | 117 | . | . | . | . |
| PM1548-SK1T1 | KLF2 | p.Ala238fs | frameshift_variant | HIGH | 0.133333 | 0 | 30 | 137 | . | . | . | . |
| PM1548-SK1T1 | NWD1 | p.Arg788Cys | missense_variant | MODERATE | 0.04 | 0 | 175 | 334 | . | . | . | . |
| PM1548-SK1T1 | BST2 | p.Ile120fs | frameshift_variant | HIGH | 0.0701754 | 0 | 57 | 64 | . | . | . | . |
| PM1548-SK1T1 | FCHO1 | p.Ala286Val | missense_variant&splice_region_variant | MODERATE | 0.03003 | 0 | 333 | 272 | . | . | . | . |
| PM1548-SK1T1 | DDX49 | p.Arg310Gln | missense_variant&splice_region_variant | MODERATE | 0.137615 | 0 | 218 | 300 | . | . | . | . |
| PM1548-SK1T1 | SUGP2 | p.Cys490Tyr | missense_variant | MODERATE | 0 | 0 | 141 | 81 | . | . | . | . |
| PM1548-SK1T1 | NR2C2AP | p.Lys28fs | frameshift_variant | HIGH | 0.0326087 | 0 | 184 | 183 | . | . | . | . |
| PM1548-SK1T1 | NCAN | p.Arg256Gln | missense_variant | MODERATE | 0.0479452 | 0 | 146 | 125 | . | . | . | . |
| PM1548-SK1T1 | NDUFA13 | p.Trp131Leu | missense_variant | MODERATE | 0 | 0 | 150 | 265 | . | . | . | . |
| PM1548-SK1T1 | ZNF728 | p.Pro380Ser | missense_variant | MODERATE | 0.265823 | 0 | 79 | 32 | . | . | . | . |
| PM1548-SK1T1 | ZNF728 | p.Arg323Pro | missense_variant | MODERATE | 0.207921 | 0 | 101 | 48 | . | . | . | . |
| PM1548-SK1T1 | ZNF728 | p.Arg323Trp | missense_variant | MODERATE | 0.207921 | 0 | 101 | 48 | . | . | . | . |
| PM1548-SK1T1 | ZNF507 | p.Asp475Gly | missense_variant | MODERATE | 0.00787402 | 0 | 127 | 42 | . | . | . | . |
| PM1548-SK1T1 | KIAA0355 | p.Val801Met | missense_variant | MODERATE | 0.00854701 | 0.0060241 | 117 | 166 | . | . | . | . |
| PM1548-SK1T1 | GRAMD1A | p.Gly605Ser | missense_variant | MODERATE | 0 | 0 | 67 | 68 | . | . | . | . |
| PM1548-SK1T1 | RBM42 | p.Pro117Arg | missense_variant | MODERATE | 0 | 0 | 197 | 129 | . | . | . | . |
| PM1548-SK1T1 | ARHGAP33 | p.Gln144fs | frameshift_variant | HIGH | 0.0600522 | 0 | 383 | 432 | . | . | . | . |
| PM1548-SK1T1 | ZNF850 | p.Ala575Thr | missense_variant | MODERATE | 0 | 0 | 117 | 89 | . | . | . | . |
| PM1548-SK1T1 | ZNF850 | p.Ser572Ala | missense_variant | MODERATE | 0 | 0 | 137 | 133 | . | . | . | . |
| PM1548-SK1T1 | ZNF850 | p.Val554Ile | missense_variant | MODERATE | 0 | 0 | 61 | 86 | . | . | . | . |
| PM1548-SK1T1 | DPF1 | p.Cys58Cys | synonymous_variant | LOW | 0 | 0 | 118 | 394 | . | . | . | . |
| PM1548-SK1T1 | DPF1 | p.Cys50Cys | synonymous_variant | LOW | 0 | 0 | 117 | 391 | . | . | . | . |
| PM1548-SK1T1 | YIF1B | p.Phe39Phe | synonymous_variant | LOW | 0.13 | 0 | 300 | 480 | . | . | . | . |
| PM1548-SK1T1 | RYR1 | p.Val3218Met | missense_variant | MODERATE | 0.142857 | 0 | 35 | 114 | . | . | . | . |
| PM1548-SK1T1 | FCGBP | p.Ala5241Val | missense_variant | MODERATE | 0.057971 | 0 | 207 | 284 | . | . | . | . |
| PM1548-SK1T1 | PRX | p.Glu495Gln | missense_variant | MODERATE | 0.0294118 | 0 | 34 | 97 | . | true | . | . |
| PM1548-SK1T1 | B3GNT8 | p.Glu333Ala | missense_variant | MODERATE | 0.0898876 | 0 | 178 | 219 | . | . | . | . |
| PM1548-SK1T1 | ARHGEF1 | p.Arg137Trp | missense_variant | MODERATE | 0.11 | 0 | 100 | 130 | . | . | . | . |
| PM1548-SK1T1 | ZNF227 | p.Gly390Ser | missense_variant | MODERATE | 0.104348 | 0 | 115 | 73 | . | . | . | . |
| PM1548-SK1T1 | PVRL2 | p.Pro53Leu | missense_variant | MODERATE | 0 | 0 | 66 | 119 | . | . | . | . |
| PM1548-SK1T1 | MARK4 | p.Thr651Thr | synonymous_variant | LOW | 0.00492611 | 0 | 203 | 352 | . | . | . | . |
| PM1548-SK1T1 | KLC3 | p.Arg320Cys | missense_variant | MODERATE | 0.0144928 | 0 | 69 | 102 | . | . | . | . |
| PM1548-SK1T1 | PPP1R13L | p.Leu397Pro | missense_variant | MODERATE | 0 | 0 | 283 | 478 | . | . | . | . |
| PM1548-SK1T1 | GIPR | p.Gly144Ser | missense_variant | MODERATE | 0.12 | 0 | 25 | 35 | . | . | . | . |
| PM1548-SK1T1 | STRN4 | p.Ala484Val | missense_variant | MODERATE | 0.147059 | 0 | 136 | 141 | . | . | . | . |
| PM1548-SK1T1 | FKRP | p.Ala193Thr | missense_variant | MODERATE | 0.1 | 0 | 40 | 97 | . | . | . | . |
| PM1548-SK1T1 | SLC1A5 | p.Ala534Thr | missense_variant | MODERATE | 0.0461538 | 0 | 65 | 44 | . | . | . | . |
| PM1548-SK1T1 | ZC3H4 | p.Arg894His | missense_variant | MODERATE | 0.162162 | 0 | 148 | 253 | . | . | . | . |
| PM1548-SK1T1 | ZC3H4 | p.Gly668Ser | missense_variant | MODERATE | 0.126126 | 0 | 111 | 220 | . | . | . | . |
| PM1548-SK1T1 | ZC3H4 | p.Lys500Arg | missense_variant | MODERATE | 0 | 0 | 101 | 129 | . | . | . | . |
| PM1548-SK1T1 | ZNF541 | p.Arg214His | missense_variant | MODERATE | 0.139535 | 0 | 43 | 53 | . | . | . | . |
| PM1548-SK1T1 | GLTSCR1 | p.Ser1126fs | frameshift_variant | HIGH | 0.186275 | 0 | 102 | 148 | . | . | . | . |
| PM1548-SK1T1 | CRX | p.Ala35Thr | missense_variant&splice_region_variant | MODERATE | 0.129252 | 0 | 147 | 73 | . | . | . | . |
| PM1548-SK1T1 | PLEKHA4 | p.Arg225Trp | missense_variant | MODERATE | 0.0857143 | 0 | 35 | 37 | . | . | . | . |
| PM1548-SK1T1 | BAX | p.Glu41fs | frameshift_variant | HIGH | 0.100719 | 0 | 139 | 75 | . | . | . | . |
| PM1548-SK1T1 | FLT3LG | p.Ser118fs | frameshift_variant | HIGH | 0 | 0 | 95 | 82 | . | . | . | . |
| PM1548-SK1T1 | SCAF1 | p.Gly651fs | frameshift_variant | HIGH | 0.0555556 | 0 | 36 | 93 | . | . | . | . |
| PM1548-SK1T1 | ASPDH | p.Glu177Lys | missense_variant | MODERATE | 0.00584795 | 0 | 171 | 226 | . | . | . | . |
| PM1548-SK1T1 | LRRC4B | p.Gly50fs | frameshift_variant | HIGH | 0.075 | 0 | 40 | 59 | . | . | . | . |
| PM1548-SK1T1 | ZNF611 | p.Met226Thr | missense_variant | MODERATE | 0.171053 | 0 | 76 | 80 | . | . | . | . |
| PM1548-SK1T1 | CNOT3 | p.Tyr637His | missense_variant | MODERATE | 0 | 0 | 117 | 41 | true | . | . | true |
| PM1548-SK1T1 | LILRB3 | c.355+1720T>A | intron_variant | MODIFIER | 0 | 0 | 239 | 187 | . | . | . | . |

| | | | | | | | | | | | | |
|---|---|---|---|---|---|---|---|---|---|---|---|---|
| PM1548-SK1T1 | LILRB3 | c.355+1720T>A | intron_variant | MODIFIER | 0 | 0 | 238 | 187 | . | . | . | . |
| PM1548-SK1T1 | LILRB3 | c.355+1720T>A | intron_variant | MODIFIER | 0 | 0 | 237 | 184 | . | . | . | . |
| PM1548-SK1T1 | LILRB3 | c.355+1720T>A | intron_variant | MODIFIER | 0 | 0 | 237 | 184 | . | . | . | . |
| PM1548-SK1T1 | EPS8L1 | p.Arg608Cys | missense_variant | MODERATE | 0 | 0 | 94 | 168 | . | . | . | . |
| PM1548-SK1T1 | SBK2 | p.Ala115Thr | missense_variant | MODERATE | 0.105991 | 0 | 217 | 186 | . | . | . | . |
| PM1548-SK1T1 | ZNF579 | p.Arg101Cys | missense_variant | MODERATE | 0.107843 | 0 | 102 | 216 | . | . | . | . |
| PM1548-SK1T1 | ZNF865 | p.Leu610Phe | missense_variant | MODERATE | 0 | 0 | 92 | 212 | . | . | . | . |
| PM1548-SK1T1 | ZNF865 | p.Leu632Leu | synonymous_variant | LOW | 0 | 0 | 88 | 209 | . | . | . | . |
| PM1548-SK1T1 | ZNF71 | p.Gln343Gln | synonymous_variant | LOW | 0.0544747 | 0 | 257 | 303 | . | . | . | . |
| PM1548-SK1T1 | TRIM28 | p.Arg492Cys | missense_variant | MODERATE | 0.17757 | 0 | 107 | 142 | . | . | . | . |
| PM1548-SK1T1 | ACP1 | p.His67Asn | missense_variant | MODERATE | 0.105263 | 0 | 38 | 36 | . | . | . | . |
| PM1548-SK1T1 | PXDN | p.Val850Met | missense_variant | MODERATE | 0.130233 | 0 | 215 | 303 | . | . | . | . |
| PM1548-SK1T1 | ROCK2 | p.Glu1060Gly | missense_variant | MODERATE | 0 | 0 | 71 | 93 | . | . | . | . |
| PM1548-SK1T1 | FAM228B | p.Asp82Gly | missense_variant | MODERATE | 0.128571 | 0 | 210 | 174 | . | . | . | . |
| PM1548-SK1T1 | NCOA1 | p.Asn1244Thr | missense_variant | MODERATE | 0 | 0 | 310 | 206 | true | . | . | . |
| PM1548-SK1T1 | NCOA1 | p.Ile1260Val | missense_variant | MODERATE | 0 | 0 | 303 | 204 | true | . | . | . |
| PM1548-SK1T1 | NCOA1 | p.Ala1274Thr | missense_variant | MODERATE | 0 | 0 | 144 | 87 | true | . | . | . |
| PM1548-SK1T1 | ADCY3 | p.Thr182Met | missense_variant | MODERATE | 0.0437956 | 0 | 137 | 85 | . | . | . | . |
| PM1548-SK1T1 | TRIM54 | p.Arg97Gln | missense_variant | MODERATE | 0.149533 | 0 | 214 | 313 | . | . | . | . |
| PM1548-SK1T1 | TRIM54 | p.Glu131Asp | missense_variant | MODERATE | 0 | 0 | 149 | 142 | . | . | . | . |
| PM1548-SK1T1 | HEATR5B | p.Lys48fs | frameshift_variant | HIGH | 0.116279 | 0 | 86 | 85 | . | . | . | . |
| PM1548-SK1T1 | RMDN2 | p.Arg130His | missense_variant | MODERATE | 0 | 0 | 195 | 121 | . | . | . | . |
| PM1548-SK1T1 | SOS1 | p.Asn15Lys | missense_variant | MODERATE | 0.173333 | 0 | 75 | 93 | . | . | . | . |
| PM1548-SK1T1 | FSHR | p.Arg569Lys | missense_variant | MODERATE | 0 | 0 | 139 | 177 | . | . | . | . |
| PM1548-SK1T1 | FSHR | p.Ser566Arg | missense_variant | MODERATE | 0 | 0 | 113 | 133 | . | . | . | . |
| PM1548-SK1T1 | FSHR | p.Asn558Asn | synonymous_variant | LOW | 0 | 0 | 137 | 139 | . | . | . | . |
| PM1548-SK1T1 | FSHR | p.Ile550Thr | missense_variant | MODERATE | 0 | 0 | 171 | 160 | . | . | . | . |
| PM1548-SK1T1 | CCDC88A | p.Pro1470Ser | missense_variant | MODERATE | 0.0150754 | 0 | 199 | 212 | . | . | . | . |
| PM1548-SK1T1 | CCDC85A | p.His295Tyr | missense_variant | MODERATE | 0.0105263 | 0 | 95 | 103 | . | . | . | . |
| PM1548-SK1T1 | CCDC85A | p.Pro359Leu | missense_variant | MODERATE | 0.193548 | 0 | 93 | 45 | . | . | . | . |
| PM1548-SK1T1 | FANCL | c.555+2T>C | splice_donor_variant&intron_variant | HIGH | 0.0241935 | 0 | 124 | 47 | . | . | . | . |
| PM1548-SK1T1 | OTX1 | p.Arg39Trp | missense_variant | MODERATE | 0.02 | 0 | 100 | 146 | . | . | . | . |
| PM1548-SK1T1 | AFTPH | p.Asp198fs | frameshift_variant | HIGH | 0 | 0 | 225 | 159 | . | . | . | . |
| PM1548-SK1T1 | NAGK | p.Lys90Gln | missense_variant | MODERATE | 0 | 0 | 286 | 344 | . | . | . | . |
| PM1548-SK1T1 | NAGK | p.Asn100Asp | missense_variant | MODERATE | 0 | 0 | 245 | 271 | . | . | . | . |
| PM1548-SK1T1 | FBXO41 | p.Arg594Gln | missense_variant | MODERATE | 0.12963 | 0 | 108 | 67 | . | . | . | . |
| PM1548-SK1T1 | TET3 | p.Pro1195Leu | missense_variant | MODERATE | 0.133333 | 0 | 105 | 50 | . | . | . | . |
| PM1548-SK1T1 | C2orf81 | p.Ala344Val | missense_variant | MODERATE | 0 | 0 | 140 | 154 | . | . | . | . |
| PM1548-SK1T1 | TCF7L1 | p.His246Pro | missense_variant | MODERATE | 0.0205128 | 0.00719424 | 195 | 139 | . | . | . | . |
| PM1548-SK1T1 | KDM3A | p.Glu1141* | stop_gained | HIGH | 0 | 0 | 205 | 166 | . | . | . | . |
| PM1548-SK1T1 | ZNF514 | p.Gly396Glu | missense_variant | MODERATE | 0.158824 | 0 | 170 | 187 | . | . | . | . |
| PM1548-SK1T1 | NCAPH | p.Gln453* | stop_gained&splice_region_variant | HIGH | 0.232323 | 0 | 99 | 53 | . | . | . | . |
| PM1548-SK1T1 | ZAP70 | p.Ala387Val | missense_variant | MODERATE | 0.183908 | 0 | 174 | 293 | . | . | . | . |
| PM1548-SK1T1 | LONRF2 | p.Asp406Gly | missense_variant | MODERATE | 0.104478 | 0 | 134 | 174 | . | . | . | . |
| PM1548-SK1T1 | CHST10 | p.Asn138Asn | synonymous_variant | LOW | 0.00621118 | 0 | 161 | 236 | . | . | . | . |
| PM1548-SK1T1 | IL18R1 | p.Pro527fs | frameshift_variant | HIGH | 0 | 0 | 52 | 190 | . | . | . | . |
| PM1548-SK1T1 | FBLN7 | c.670+86C>T | intron_variant | MODIFIER | 0.0192308 | 0 | 104 | 109 | . | . | . | . |
| PM1548-SK1T1 | FBLN7 | p.Arg381Cys | missense_variant | MODERATE | 0.220472 | 0 | 127 | 87 | . | . | . | . |
| PM1548-SK1T1 | ZC3H6 | p.Arg99fs | frameshift_variant | HIGH | 0.0747664 | 0 | 107 | 90 | . | . | . | . |
| PM1548-SK1T1 | POLR1B | p.Cys893Tyr | missense_variant | MODERATE | 0 | 0 | 115 | 127 | . | . | . | . |
| PM1548-SK1T1 | IL36B | c.261+1014delT | intron_variant | MODIFIER | 0.198113 | 0 | 106 | 101 | . | . | . | . |
| PM1548-SK1T1 | TMEM185B | p.Pro341fs | frameshift_variant | HIGH | 0.0425532 | 0 | 188 | 232 | . | . | . | . |
| PM1548-SK1T1 | CLASP1 | p.Ala1225Val | missense_variant | MODERATE | 0 | 0 | 95 | 85 | . | . | . | . |
| PM1548-SK1T1 | CYP27C1 | p.Ala103Thr | missense_variant | MODERATE | 0.025 | 0 | 160 | 179 | . | . | . | . |
| PM1548-SK1T1 | ACVR2A | p.Lys437fs | frameshift_variant | HIGH | 0.0202703 | 0 | 148 | 201 | . | true | . | . |
| PM1548-SK1T1 | NEB | p.Phe2843Tyr | missense_variant | MODERATE | 0 | 0 | 153 | 178 | . | . | . | . |
| PM1548-SK1T1 | TBR1 | p.Gly599Ser | missense_variant | MODERATE | 0.272727 | 0 | 11 | 40 | . | . | . | . |
| PM1548-SK1T1 | SCN3A | p.Ala1768Val | missense_variant | MODERATE | 0.131579 | 0 | 76 | 50 | . | . | . | . |
| PM1548-SK1T1 | TLK1 | p.Gln18Arg | missense_variant | MODERATE | 0 | 0 | 146 | 529 | . | . | . | . |
| PM1548-SK1T1 | ITGA6 | p.Val266Leu | missense_variant | MODERATE | 0 | 0 | 112 | 156 | . | . | . | . |
| PM1548-SK1T1 | SP3 | p.Val461Ile | missense_variant | MODERATE | 0 | 0 | 280 | 262 | . | . | . | . |
| PM1548-SK1T1 | EVX2 | p.Gly438Ser | missense_variant | MODERATE | 0.12069 | 0 | 58 | 165 | . | . | . | . |
| PM1548-SK1T1 | EVX2 | p.Asp320Glu | missense_variant | MODERATE | 0.190476 | 0 | 42 | 48 | . | . | . | . |
| PM1548-SK1T1 | CCDC141 | p.Ala1030Val | missense_variant | MODERATE | 0 | 0 | 101 | 69 | . | . | . | . |
| PM1548-SK1T1 | PDE1A | p.Ala371Val | missense_variant | MODERATE | 0.121495 | 0 | 107 | 75 | . | . | . | . |
| PM1548-SK1T1 | FSIP2 | p.Arg829Cys | missense_variant | MODERATE | 0.0108108 | 0 | 185 | 51 | . | . | . | . |
| PM1548-SK1T1 | DNAH7 | p.Ala3252Glu | missense_variant | MODERATE | 0.0483871 | 0 | 62 | 64 | . | . | . | . |
| PM1548-SK1T1 | GTF3C3 | p.Pro248Leu | missense_variant | MODERATE | 0 | 0 | 205 | 201 | . | . | . | . |
| PM1548-SK1T1 | AOX1 | p.Phe1162Ser | missense_variant | MODERATE | 0 | 0 | 103 | 36 | . | . | . | . |
| PM1548-SK1T1 | PIKFYVE | p.Leu419Leu | synonymous_variant | LOW | 0 | 0 | 273 | 195 | . | . | . | . |
| PM1548-SK1T1 | MAP2 | p.Gly1784Ser | missense_variant | MODERATE | 0 | 0 | 116 | 93 | . | . | . | . |
| PM1548-SK1T1 | SMARCAL1 | p.Arg499Gln | missense_variant | MODERATE | 0.104651 | 0 | 172 | 137 | . | . | . | . |
| PM1548-SK1T1 | AAMP | p.Glu63Asp | missense_variant | MODERATE | 0 | 0.00606061 | 194 | 165 | . | . | . | . |
| PM1548-SK1T1 | ZNF142 | p.Ala335Pro | missense_variant | MODERATE | 0.00793651 | 0 | 252 | 317 | . | . | . | . |
| PM1548-SK1T1 | CYP27A1 | p.Ser203Leu | missense_variant | MODERATE | 0.0571429 | 0 | 105 | 39 | . | . | . | . |
| PM1548-SK1T1 | WNT6 | p.Asp283Val | missense_variant | MODERATE | 0.121212 | 0 | 99 | 120 | . | . | . | . |
| PM1548-SK1T1 | FAM134A | p.Asp391Val | missense_variant | MODERATE | 0.0481928 | 0 | 249 | 285 | . | . | . | . |
| PM1548-SK1T1 | DNPEP | p.Arg191Gln | missense_variant | MODERATE | 0.102302 | 0 | 391 | 294 | . | . | . | . |
| PM1548-SK1T1 | CHPF | p.Arg721Trp | missense_variant | MODERATE | 0 | 0 | 79 | 74 | . | . | . | . |
| PM1548-SK1T1 | WDFY1 | p.Ala75Asp | missense_variant | MODERATE | 0 | 0 | 209 | 181 | . | . | . | . |
| PM1548-SK1T1 | SERPINE2 | p.Met296Val | missense_variant | MODERATE | 0.0844156 | 0 | 154 | 111 | . | . | . | . |
| PM1548-SK1T1 | B3GNT7 | p.Gly302Asp | missense_variant | MODERATE | 0.020202 | 0 | 99 | 60 | . | . | . | . |
| PM1548-SK1T1 | DGKD | p.Arg935Trp | missense_variant | MODERATE | 0.138462 | 0 | 65 | 67 | . | . | . | . |
| PM1548-SK1T1 | ACKR3 | p.Val42Ala | missense_variant | MODERATE | 0.0122699 | 0 | 163 | 137 | true | . | . | . |
| PM1548-SK1T1 | COL6A3 | p.Gly1714Glu | missense_variant | MODERATE | 0.114583 | 0 | 96 | 48 | . | . | . | . |
| PM1548-SK1T1 | COL6A3 | p.Ile1609Asn | missense_variant | MODERATE | 0.0558659 | 0 | 179 | 113 | . | . | . | . |
| PM1548-SK1T1 | ING5 | p.Arg106His | missense_variant | MODERATE | 0.100719 | 0 | 139 | 105 | . | . | . | . |
| PM1548-SK1T1 | TBC1D20 | p.Pro71Ser | missense_variant | MODERATE | 0.012987 | 0 | 77 | 49 | . | . | . | . |
| PM1548-SK1T1 | VPS16 | p.Thr818Met | missense_variant | MODERATE | 0.013986 | 0 | 143 | 238 | . | . | . | . |
| PM1548-SK1T1 | ATRN | p.Ala122Thr | missense_variant | MODERATE | 0 | 0.00645161 | 146 | 155 | . | . | . | . |
| PM1548-SK1T1 | SLC23A2 | p.Ile412fs | frameshift_variant | HIGH | 0.0833333 | 0 | 96 | 150 | . | . | . | . |
| PM1548-SK1T1 | BMP2 | p.Ala10Val | missense_variant | MODERATE | 0 | 0 | 60 | 162 | . | . | . | . |
| PM1548-SK1T1 | PLCB1 | p.Arg768Gln | missense_variant | MODERATE | 0.12963 | 0 | 54 | 91 | . | . | . | . |
| PM1548-SK1T1 | JAG1 | p.Gly537Ser | missense_variant | MODERATE | 0.0126582 | 0 | 158 | 184 | . | . | . | . |
| PM1548-SK1T1 | CSRP2BP | p.Trp173* | stop_gained | HIGH | 0 | 0 | 113 | 168 | . | . | . | . |
| PM1548-SK1T1 | CRNKL1 | p.Phe352Leu | missense_variant | MODERATE | 0 | 0 | 161 | 71 | . | . | . | . |
| PM1548-SK1T1 | PLAGL2 | p.Arg180His | missense_variant | MODERATE | 0.0416667 | 0 | 120 | 145 | . | . | . | . |
| PM1548-SK1T1 | BPIFA2 | p.Arg221His | missense_variant | MODERATE | 0.163934 | 0 | 122 | 61 | . | . | . | . |
| PM1548-SK1T1 | NECAB3 | p.Arg278fs | frameshift_variant | HIGH | 0.0588235 | 0 | 34 | 39 | . | . | . | . |
| PM1548-SK1T1 | CEP250 | p.Lys787Glu | missense_variant | MODERATE | 0 | 0 | 122 | 64 | . | . | . | . |
| PM1548-SK1T1 | SOGA1 | p.Pro974Ser | missense_variant&splice_region_variant | MODERATE | 0 | 0 | 252 | 259 | . | . | . | . |
| PM1548-SK1T1 | PLCG1 | p.Ile942Met | missense_variant | MODERATE | 0 | 0 | 85 | 140 | true | . | . | . |
| PM1548-SK1T1 | ZSWIM3 | p.Arg225Gln | missense_variant | MODERATE | 0.142857 | 0 | 189 | 55 | . | . | . | . |
| PM1548-SK1T1 | NFATC2 | p.Phe158Tyr | missense_variant | MODERATE | 0 | 0 | 54 | 59 | true | . | . | . |
| PM1548-SK1T1 | NFATC2 | p.Gly22fs | frameshift_variant | HIGH | 0.104762 | 0 | 105 | 145 | true | . | . | . |
| PM1548-SK1T1 | SALL4 | p.Phe391Leu | missense_variant | MODERATE | 0 | 0 | 89 | 33 | . | . | . | . |
| PM1548-SK1T1 | ZFP64 | n.50701484A>G | intragenic_variant | MODIFIER | 0.0454545 | 0 | 44 | 142 | . | . | . | . |
| PM1548-SK1T1 | DOK5 | p.Val169Ala | missense_variant | MODERATE | 0.0552764 | 0 | 199 | 340 | . | . | . | . |
| PM1548-SK1T1 | DOK5 | p.Ala267Asp | missense_variant | MODERATE | 0 | 0 | 197 | 146 | . | . | . | . |
| PM1548-SK1T1 | TFAP2C | p.Ala354Val | missense_variant | MODERATE | 0.0430108 | 0 | 186 | 269 | . | . | . | . |
| PM1548-SK1T1 | PMEPA1 | p.Ser209fs | frameshift_variant | HIGH | 0.105882 | 0 | 85 | 248 | . | . | . | . |
| PM1548-SK1T1 | NELFCD | p.Val321Ile | missense_variant | MODERATE | 0.0268817 | 0 | 186 | 74 | . | . | . | . |
| PM1548-SK1T1 | ZNF831 | p.Leu434Met | missense_variant | MODERATE | 0.140351 | 0 | 57 | 129 | . | . | . | . |
| PM1548-SK1T1 | HELZ2 | p.Ala639Thr | missense_variant | MODERATE | 0 | 0 | 106 | 139 | . | . | . | . |
| PM1548-SK1T1 | RTEL1 | p.Arg729Cys | missense_variant | MODERATE | 0.0735294 | 0 | 136 | 264 | . | . | . | . |
| PM1548-SK1T1 | TNFRSF6B | p.Leu297Phe | missense_variant | MODERATE | 0.118644 | 0 | 118 | 277 | . | . | . | . |
| PM1548-SK1T1 | SLC2A4RG | p.Arg82fs | frameshift_variant | HIGH | 0.0512821 | 0 | 39 | 206 | . | . | . | . |
| PM1548-SK1T1 | PRPF6 | p.Arg317His | missense_variant | MODERATE | 0 | 0 | 189 | 86 | . | . | . | . |
| PM1548-SK1T1 | LTN1 | p.Gly1705Val | missense_variant | MODERATE | 0.1 | 0 | 120 | 36 | . | . | . | . |
| PM1548-SK1T1 | TIAM1 | p.His1010Tyr | missense_variant | MODERATE | 0 | 0 | 486 | 418 | . | . | . | . |
| PM1548-SK1T1 | URB1 | p.Pro2036Leu | missense_variant | MODERATE | 0.172414 | 0 | 58 | 14 | . | . | . | . |
| PM1548-SK1T1 | PAXBP1 | p.Pro161Thr | missense_variant | MODERATE | 0.142857 | 0 | 56 | 40 | . | . | . | . |
| PM1548-SK1T1 | PAXBP1 | p.Ser105Arg | missense_variant | MODERATE | 0.117647 | 0 | 34 | 156 | . | . | . | . |
| PM1548-SK1T1 | GART | p.Glu297Gln | missense_variant | MODERATE | 0 | 0 | 112 | 160 | . | . | . | . |
| PM1548-SK1T1 | SON | p.Thr790Ser | missense_variant | MODERATE | 0 | 0 | 330 | 177 | . | . | . | . |
| PM1548-SK1T1 | SON | c.6657+163T>C | intron_variant | MODIFIER | 0.0795455 | 0 | 88 | 34 | . | . | . | true |
| PM1548-SK1T1 | ITSN1 | p.Met768Thr | missense_variant | MODERATE | 0.119403 | 0 | 134 | 46 | . | . | . | . |

| | | | | | | | | | | | | |
|---|---|---|---|---|---|---|---|---|---|---|---|---|
| PM1548-SK1T1 | SLC5A3 | p.Arg252Gln | missense_variant | MODERATE | 0 | 0 | 77 | 44 | . | . | . | . |
| PM1548-SK1T1 | UMODL1 | p.Gly1319Ser | missense_variant | MODERATE | 0.192118 | 0 | 203 | 131 | . | . | . | . |
| PM1548-SK1T1 | CRYAA | p.Ile61Val | missense_variant | MODERATE | 0.0377358 | 0 | 159 | 344 | . | . | . | . |
| PM1548-SK1T1 | SLC19A1 | p.Asp488Gly | missense_variant | MODERATE | 0 | 0 | 40 | 131 | . | . | . | . |
| PM1548-SK1T1 | FTCD | p.Ile187Thr | missense_variant | MODERATE | 0.0681818 | 0 | 44 | 175 | . | . | . | . |
| PM1548-SK1T1 | LSS | p.Thr457Met | missense_variant | MODERATE | 0.0820513 | 0 | 195 | 297 | . | . | . | . |
| PM1548-SK1T1 | CECR2 | p.Ser425Ser | synonymous_variant | LOW | 0 | 0 | 115 | 131 | . | . | . | . |
| PM1548-SK1T1 | CECR2 | p.Pro428Pro | synonymous_variant | LOW | 0 | 0 | 115 | 130 | . | . | . | . |
| PM1548-SK1T1 | CECR2 | p.Tyr431Tyr | synonymous_variant | LOW | 0 | 0 | 115 | 130 | . | . | . | . |
| PM1548-SK1T1 | ATP6V1E1 | p.Leu128Met | missense_variant | MODERATE | 0.0304183 | 0 | 263 | 148 | . | . | . | . |
| PM1548-SK1T1 | CLTCL1 | p.Ala1567Thr | missense_variant | MODERATE | 0.0180723 | 0 | 332 | 281 | true | . | . | . |
| PM1548-SK1T1 | SCARF2 | p.Arg335Cys | missense_variant | MODERATE | 0.109091 | 0 | 110 | 100 | . | . | . | . |
| PM1548-SK1T1 | SLC7A4 | p.Ala256Thr | missense_variant | MODERATE | 0.166667 | 0 | 120 | 223 | . | . | . | . |
| PM1548-SK1T1 | IGLL1 | p.Arg60Gln | missense_variant | MODERATE | 0.113402 | 0 | 97 | 81 | . | . | . | . |
| PM1548-SK1T1 | C22orf43 | p.Ser153Gly | missense_variant | MODERATE | 0.100719 | 0 | 139 | 97 | . | . | . | . |
| PM1548-SK1T1 | CABIN1 | p.Pro585Gln | missense_variant | MODERATE | 0.133333 | 0 | 30 | 58 | . | . | . | . |
| PM1548-SK1T1 | ASPHD2 | p.Arg358Trp | missense_variant | MODERATE | 0.0731707 | 0 | 205 | 411 | . | . | . | . |
| PM1548-SK1T1 | TFIP11 | p.Arg39* | stop_gained | HIGH | 0.0288462 | 0 | 104 | 103 | . | . | . | . |
| PM1548-SK1T1 | RASL10A | p.Arg108Trp | missense_variant | MODERATE | 0 | 0 | 171 | 108 | . | . | . | . |
| PM1548-SK1T1 | AP1B1 | p.Ala655Asp | missense_variant | MODERATE | 0.1 | 0 | 30 | 31 | . | . | . | . |
| PM1548-SK1T1 | LIMK2 | p.Asn477Ser | missense_variant | MODERATE | 0 | 0 | 229 | 257 | . | . | . | . |
| PM1548-SK1T1 | SFI1 | p.Thr710Ser | missense_variant | MODERATE | 0 | 0 | 187 | 104 | . | . | . | . |
| PM1548-SK1T1 | CSF2RB | p.Gln642fs | frameshift_variant | HIGH | 0.0230769 | 0 | 130 | 133 | . | . | . | . |
| PM1548-SK1T1 | C1QTNF6 | p.Met229Thr | missense_variant | MODERATE | 0.25 | 0 | 4 | 20 | . | . | . | . |
| PM1548-SK1T1 | GRAP2 | p.Val129Ile | missense_variant | MODERATE | 0.0491803 | 0 | 183 | 162 | . | . | . | . |
| PM1548-SK1T1 | MKL1 | p.Pro138Leu | missense_variant | MODERATE | 0 | 0 | 102 | 100 | true | . | . | . |
| PM1548-SK1T1 | EP300 | p.Pro2107Thr | missense_variant | MODERATE | 0 | 0 | 144 | 94 | true | true | true | . |
| PM1548-SK1T1 | ZC3H7B | p.Arg908Cys | missense_variant | MODERATE | 0.158273 | 0 | 139 | 73 | . | . | . | . |
| PM1548-SK1T1 | PARVB | c.211+24725C>A | intron_variant | MODIFIER | 0.0408163 | 0 | 98 | 168 | . | . | . | . |
| PM1548-SK1T1 | CELSR1 | p.Val1411Gly | missense_variant | MODERATE | 0 | 0 | 228 | 333 | . | . | . | . |
| PM1548-SK1T1 | CELSR1 | p.Glu1159Lys | missense_variant | MODERATE | 0.193717 | 0 | 191 | 186 | . | . | . | . |
| PM1548-SK1T1 | TUBGCP6 | p.Cys1239Trp | missense_variant | MODERATE | 0.00719424 | 0.00947867 | 139 | 211 | . | . | . | . |
| PM1548-SK1T1 | TUBGCP6 | p.Ser1237Pro | missense_variant | MODERATE | 0.00460829 | 0 | 217 | 238 | . | . | . | . |
| PM1548-SK1T1 | MIOX | p.Asp78Gly | missense_variant | MODERATE | 0.111111 | 0 | 126 | 102 | . | . | . | . |
| PM1548-SK1T1 | MAPK8IP2 | p.Thr518Thr | synonymous_variant | LOW | 0 | 0 | 110 | 166 | . | . | . | . |
| PM1548-SK1T1 | ARSA | c.684+1G>A | splice_donor_variant&intron_variant | HIGH | 0.0666667 | 0 | 30 | 132 | . | . | . | . |
| PM1548-SK1T1 | SETD5 | p.Arg1193* | stop_gained | HIGH | 0.165179 | 0 | 224 | 243 | . | . | . | . |
| PM1548-SK1T1 | CPNE9 | p.Ser284Ser | synonymous_variant | LOW | 0 | 0 | 120 | 197 | . | . | . | . |
| PM1548-SK1T1 | CPNE9 | p.Asp473Asp | synonymous_variant | LOW | 0.0634146 | 0 | 205 | 87 | . | . | . | . |
| PM1548-SK1T1 | BRPF1 | p.Gln938fs | frameshift_variant | HIGH | 0.225806 | 0 | 31 | 52 | . | . | . | true |
| PM1548-SK1T1 | OGG1 | p.Ala85Thr | missense_variant | MODERATE | 0.115523 | 0 | 277 | 334 | . | . | . | . |
| PM1548-SK1T1 | CRELD1 | p.Arg220* | stop_gained | HIGH | 0.126354 | 0 | 277 | 387 | . | . | . | . |
| PM1548-SK1T1 | PRRT3 | p.His679Arg | missense_variant | MODERATE | 0 | 0 | 91 | 166 | . | . | . | . |
| PM1548-SK1T1 | CAND2 | p.Arg489His | missense_variant | MODERATE | 0.0285714 | 0 | 315 | 333 | . | . | . | . |
| PM1548-SK1T1 | SLC6A6 | p.Phe392fs | frameshift_variant | HIGH | 0.0277778 | 0 | 72 | 115 | . | . | . | . |
| PM1548-SK1T1 | NR2C2 | p.Phe582fs | frameshift_variant | HIGH | 0.037037 | 0 | 135 | 75 | . | . | . | . |
| PM1548-SK1T1 | SH3BP5 | p.Thr119Ala | missense_variant | MODERATE | 0.0898438 | 0 | 256 | 346 | . | . | . | . |
| PM1548-SK1T1 | PLCL2 | p.Gln628Arg | missense_variant | MODERATE | 0 | 0 | 50 | 118 | . | . | . | . |
| PM1548-SK1T1 | SLC4A7 | p.Ile1097Val | missense_variant | MODERATE | 0 | 0.00657895 | 144 | 152 | . | . | . | . |
| PM1548-SK1T1 | CLASP2 | p.Cys10Cys | synonymous_variant | LOW | 0.0615385 | 0.00520833 | 65 | 192 | . | . | . | . |
| PM1548-SK1T1 | SCN10A | p.Val378Leu | missense_variant | MODERATE | 0.0172414 | 0.00809717 | 232 | 247 | . | . | . | . |
| PM1548-SK1T1 | TTC21A | p.Ser332Thr | missense_variant | MODERATE | 0 | 0 | 109 | 40 | . | . | . | . |
| PM1548-SK1T1 | SNRK | p.Pro598Thr | missense_variant | MODERATE | 0.102564 | 0 | 39 | 93 | . | . | . | . |
| PM1548-SK1T1 | SLC6A20 | p.Asn103Lys | missense_variant | MODERATE | 0.105263 | 0 | 76 | 172 | . | . | . | . |
| PM1548-SK1T1 | SLC6A20 | p.Gly66Trp | missense_variant | MODERATE | 0 | 0.00847458 | 173 | 118 | . | . | . | . |
| PM1548-SK1T1 | FYCO1 | p.Lys501del | inframe_deletion | MODERATE | 0.104803 | 0 | 229 | 171 | . | . | . | . |
| PM1548-SK1T1 | ALS2CL | p.Asn384Lys | missense_variant | MODERATE | 0.0207254 | 0 | 193 | 77 | . | . | . | . |
| PM1548-SK1T1 | DHX30 | n.47859525delC | intragenic_variant | MODIFIER | 0.157895 | 0 | 209 | 119 | . | . | . | . |
| PM1548-SK1T1 | UCN2 | p.Thr27Pro | missense_variant | MODERATE | 0.00917431 | 0 | 218 | 135 | . | . | . | . |
| PM1548-SK1T1 | CELSR3 | p.Pro1425Ser | missense_variant | MODERATE | 0.135338 | 0 | 133 | 95 | . | . | . | . |
| PM1548-SK1T1 | SLC25A20 | p.Ala227Val | missense_variant | MODERATE | 0.262712 | 0 | 118 | 125 | . | . | . | . |
| PM1548-SK1T1 | KLHDC8B | p.Gly38fs | frameshift_variant | HIGH | 0.0136986 | 0 | 146 | 199 | . | . | . | . |
| PM1548-SK1T1 | RRP9 | p.Ala342Thr | missense_variant | MODERATE | 0.139535 | 0 | 172 | 161 | . | . | . | . |
| PM1548-SK1T1 | DNAH1 | p.Leu2072Ile | missense_variant | MODERATE | 0.155172 | 0 | 58 | 99 | . | . | . | . |
| PM1548-SK1T1 | FAM208A | p.Ala227Val | missense_variant | MODERATE | 0.0885417 | 0 | 192 | 81 | . | . | . | . |
| PM1548-SK1T1 | DNAH12 | p.Pro1035Ala | missense_variant | MODERATE | 0 | 0 | 211 | 189 | . | . | . | . |
| PM1548-SK1T1 | PDE12 | p.Ala293Ser | missense_variant | MODERATE | 0 | 0 | 146 | 150 | . | . | . | . |
| PM1548-SK1T1 | PDE12 | p.Leu294Phe | missense_variant | MODERATE | 0 | 0 | 146 | 150 | . | . | . | . |
| PM1548-SK1T1 | ADAMTS9 | p.Pro596His | missense_variant | MODERATE | 0 | 0 | 202 | 126 | . | . | . | . |
| PM1548-SK1T1 | ADAMTS9 | p.Ser593Gly | missense_variant | MODERATE | 0 | 0 | 233 | 130 | . | . | . | . |
| PM1548-SK1T1 | ROBO1 | p.Tyr1009* | stop_gained | HIGH | 0.1 | 0 | 40 | 53 | . | . | . | . |
| PM1548-SK1T1 | OR5K4 | p.Val51Ile | missense_variant | MODERATE | 0 | 0 | 445 | 442 | . | . | . | . |
| PM1548-SK1T1 | OR5K4 | p.Arg53Pro | missense_variant | MODERATE | 0 | 0 | 445 | 441 | . | . | . | . |
| PM1548-SK1T1 | OR5K4 | p.Leu56His | missense_variant | MODERATE | 0.00187617 | 0 | 533 | 510 | . | . | . | . |
| PM1548-SK1T1 | DCBLD2 | p.Thr718Met | missense_variant | MODERATE | 0.104839 | 0 | 248 | 81 | . | . | . | . |
| PM1548-SK1T1 | TFG | p.Ser213Pro | missense_variant | MODERATE | 0.017341 | 0 | 173 | 76 | true | . | . | . |
| PM1548-SK1T1 | CBLB | p.Pro823Ser | missense_variant | MODERATE | 0 | 0 | 197 | 275 | true | . | . | . |
| PM1548-SK1T1 | BOC | p.Pro593Leu | missense_variant | MODERATE | 0 | 0.00197628 | 253 | 506 | . | . | . | . |
| PM1548-SK1T1 | BOC | p.His601Gln | missense_variant | MODERATE | 0 | 0 | 136 | 255 | . | . | . | . |
| PM1548-SK1T1 | KIAA2018 | p.Asn1016fs | frameshift_variant | HIGH | 0.12963 | 0 | 54 | 36 | . | . | . | . |
| PM1548-SK1T1 | KIAA1407 | p.Glu683Gly | missense_variant | MODERATE | 0.186047 | 0 | 86 | 85 | . | . | . | . |
| PM1548-SK1T1 | LSAMP | p.Val277Ile | missense_variant | MODERATE | 0.206349 | 0 | 63 | 91 | . | . | . | . |
| PM1548-SK1T1 | ARHGAP31 | p.Lys857Glu | missense_variant | MODERATE | 0.0724638 | 0 | 69 | 33 | . | . | . | . |
| PM1548-SK1T1 | FSTL1 | p.Val13Ala | missense_variant | MODERATE | 0.0133333 | 0.00438596 | 75 | 228 | . | . | . | . |
| PM1548-SK1T1 | EAF2 | p.Thr112fs | frameshift_variant | HIGH | 0.0574713 | 0 | 87 | 119 | . | . | . | . |
| PM1548-SK1T1 | ILDR1 | p.Ala76Val | missense_variant&splice_region_variant | MODERATE | 0 | 0 | 301 | 159 | . | . | . | . |
| PM1548-SK1T1 | MYLK | p.Ala1684Thr | missense_variant | MODERATE | 0.013245 | 0 | 151 | 98 | . | . | . | . |
| PM1548-SK1T1 | ALDH1L1 | p.Glu439fs | frameshift_variant | HIGH | 0.269231 | 0 | 52 | 55 | . | . | . | . |
| PM1548-SK1T1 | CHCHD6 | p.Ser220Leu | missense_variant | MODERATE | 0.144231 | 0 | 104 | 164 | . | . | . | . |
| PM1548-SK1T1 | C3orf27 | p.Asp85fs | frameshift_variant | HIGH | 0.0375587 | 0 | 213 | 231 | . | . | . | . |
| PM1548-SK1T1 | TMCC1 | p.Arg620Ser | missense_variant | MODERATE | 0 | 0 | 151 | 62 | . | . | . | . |
| PM1548-SK1T1 | CDV3 | p.Gly226Asp | missense_variant | MODERATE | 0 | 0 | 78 | 44 | . | . | . | . |
| PM1548-SK1T1 | CDV3 | p.Asp228Glu | missense_variant | MODERATE | 0 | 0 | 111 | 55 | . | . | . | . |
| PM1548-SK1T1 | AMOTL2 | p.Arg292Cys | missense_variant | MODERATE | 0.1625 | 0 | 80 | 43 | . | . | . | . |
| PM1548-SK1T1 | EPHB1 | p.Ala82fs | frameshift_variant | HIGH | 0.0177778 | 0 | 225 | 300 | . | . | . | . |
| PM1548-SK1T1 | SLC35G2 | p.Asn97fs | frameshift_variant | HIGH | 0 | 0 | 77 | 31 | . | . | . | . |
| PM1548-SK1T1 | ESYT3 | p.Val238Met | missense_variant | MODERATE | 0.0193548 | 0 | 155 | 79 | . | . | . | . |
| PM1548-SK1T1 | MED12L | p.Ser970Leu | missense_variant | MODERATE | 0 | 0 | 198 | 123 | . | . | . | . |
| PM1548-SK1T1 | DHX36 | p.Met154fs | frameshift_variant | HIGH | 0.133333 | 0 | 30 | 30 | . | . | . | . |
| PM1548-SK1T1 | CCNL1 | p.Ile354Val | missense_variant | MODERATE | 0 | 0 | 86 | 120 | . | . | . | . |
| PM1548-SK1T1 | PPM1L | p.Glu106Glu | synonymous_variant | LOW | 0 | 0 | 195 | 275 | . | . | . | . |
| PM1548-SK1T1 | SLITRK3 | p.Arg759Cys | missense_variant | MODERATE | 0 | 0 | 106 | 99 | . | . | . | . |
| PM1548-SK1T1 | ZBBX | p.Ile496fs | frameshift_variant | HIGH | 0.0111111 | 0 | 90 | 76 | . | . | . | . |
| PM1548-SK1T1 | FNDC3B | p.Glu687Gly | missense_variant | MODERATE | 0 | 0 | 84 | 96 | . | . | . | . |
| PM1548-SK1T1 | NCEH1 | p.Ser105fs | frameshift_variant | HIGH | 0 | 0 | 112 | 228 | . | . | . | . |
| PM1548-SK1T1 | KLHL24 | p.Asn555fs | frameshift_variant | HIGH | 0.16092 | 0 | 87 | 55 | . | . | . | . |
| PM1548-SK1T1 | CHRD | p.Arg76Cys | missense_variant | MODERATE | 0.173913 | 0 | 115 | 91 | . | . | . | . |
| PM1548-SK1T1 | TP63 | p.Thr368Ala | missense_variant | MODERATE | 0 | 0 | 128 | 121 | . | . | . | . |
| PM1548-SK1T1 | LEPREL1 | p.Asp204Glu | missense_variant | MODERATE | 0 | 0 | 244 | 177 | . | . | . | . |
| PM1548-SK1T1 | CCDC50 | p.Ala343Ser | missense_variant | MODERATE | 0 | 0 | 29 | 55 | . | . | . | . |
| PM1548-SK1T1 | MUC20 | p.Gly514Arg | missense_variant | MODERATE | 0 | 0 | 173 | 228 | . | . | . | . |
| PM1548-SK1T1 | LRCH3 | p.Leu49Val | missense_variant | MODERATE | 0.00724638 | 0.00369004 | 138 | 271 | . | . | . | . |
| PM1548-SK1T1 | GAK | p.Ala690Val | missense_variant | MODERATE | 0.144 | 0 | 125 | 111 | . | . | . | . |
| PM1548-SK1T1 | DGKQ | p.Thr299Met | missense_variant | MODERATE | 0.0141844 | 0 | 141 | 253 | . | . | . | . |
| PM1548-SK1T1 | SLC26A1 | p.Ala624Thr | missense_variant | MODERATE | 0.134021 | 0 | 194 | 464 | . | . | . | . |
| PM1548-SK1T1 | POLN | p.Tyr686Cys | missense_variant | MODERATE | 0.101266 | 0 | 79 | 37 | . | . | . | . |
| PM1548-SK1T1 | HTT | p.Ala2259Thr | missense_variant&splice_region_variant | MODERATE | 0.0384615 | 0 | 52 | 52 | . | . | . | . |
| PM1548-SK1T1 | PPP2R2C | p.Gly397fs | frameshift_variant | HIGH | 0.116505 | 0 | 103 | 188 | . | . | . | . |
| PM1548-SK1T1 | SH3TC1 | p.Glu203Gly | missense_variant | MODERATE | 0.00847458 | 0.0021322 | 236 | 469 | . | . | . | . |
| PM1548-SK1T1 | GPR78 | p.Val209Ile | missense_variant | MODERATE | 0.0307692 | 0 | 65 | 34 | . | . | . | . |
| PM1548-SK1T1 | KCTD8 | p.Leu62Leu | synonymous_variant | LOW | 0 | 0 | 118 | 213 | . | . | . | . |
| PM1548-SK1T1 | YTHDC1 | p.Ser684Asn | missense_variant | MODERATE | 0.225225 | 0 | 111 | 59 | . | . | . | . |
| PM1548-SK1T1 | ANKRD17 | p.Ile2241Val | missense_variant | MODERATE | 0 | 0 | 145 | 147 | . | . | . | . |
| PM1548-SK1T1 | PF4 | p.Ala4Thr | missense_variant | MODERATE | 0 | 0 | 78 | 40 | . | . | . | . |
| PM1548-SK1T1 | PARM1 | p.Ser82Pro | missense_variant | MODERATE | 0 | 0 | 303 | 388 | . | . | . | . |
| PM1548-SK1T1 | CDKL2 | p.Pro452Ser | missense_variant | MODERATE | 0.102564 | 0 | 39 | 51 | . | . | . | . |
| PM1548-SK1T1 | SHROOM3 | p.Lys1017del | inframe_deletion | MODERATE | 0.106061 | 0 | 66 | 97 | . | . | . | . |

| | | | | | | | | | | | | |
|---|---|---|---|---|---|---|---|---|---|---|---|---|
| PM1548-SK1T1 | BMP3 | p.Lys358fs | frameshift_variant | HIGH | 0.115044 | 0 | 113 | 124 | . | . | . | . |
| PM1548-SK1T1 | PRKG2 | p.Val461Phe | missense_variant | MODERATE | 0.12 | 0 | 50 | 41 | . | . | . | . |
| PM1548-SK1T1 | SEC31A | p.Ile462fs | frameshift_variant | HIGH | 0.344632768 | 0 | 177 | 87 | . | . | . | . |
| PM1548-SK1T1 | SEC31A | p.Lys461fs | frameshift_variant | HIGH | 0.13559322 | 0 | 177 | 87 | . | . | . | . |
| PM1548-SK1T1 | SEC31A | p.Gln422Pro | missense_variant | MODERATE | 0 | 0 | 191 | 143 | . | . | . | . |
| PM1548-SK1T1 | AGPAT9 | p.Met372fs | frameshift_variant | HIGH | 0.0848485 | 0 | 165 | 70 | . | . | . | . |
| PM1548-SK1T1 | NAP1L5 | p.Lys108fs | frameshift_variant | HIGH | 0.0304878 | 0 | 328 | 237 | . | . | . | . |
| PM1548-SK1T1 | CASP6 | p.Pro33Leu | missense_variant | MODERATE | 0 | 0.00719424 | 289 | 278 | . | . | . | . |
| PM1548-SK1T1 | PITX2 | c.411+284C>T | intron_variant | MODIFIER | 0.176471 | 0 | 51 | 59 | . | . | . | . |
| PM1548-SK1T1 | ALPK1 | p.Tyr1216fs | frameshift_variant | HIGH | 0 | 0 | 17 | 44 | . | . | . | . |
| PM1548-SK1T1 | TNIP3 | p.Ala59Thr | missense_variant | MODERATE | 0.00970874 | 0 | 103 | 33 | . | . | . | . |
| PM1548-SK1T1 | PCDH10 | p.Tyr324Cys | missense_variant | MODERATE | 0 | 0 | 42 | 50 | . | . | . | . |
| PM1548-SK1T1 | SETD7 | p.Leu331Ile | missense_variant | MODERATE | 0.0346821 | 0 | 173 | 311 | . | . | . | . |
| PM1548-SK1T1 | FREM3 | p.Arg502* | stop_gained | HIGH | 0.219512 | 0 | 82 | 32 | . | . | . | . |
| PM1548-SK1T1 | NEK1 | p.Thr137Asn | missense_variant | MODERATE | 0.101695 | 0 | 59 | 71 | . | . | . | . |
| PM1548-SK1T1 | AHRR | p.Arg72His | missense_variant | MODERATE | 0.0735294 | 0 | 136 | 76 | . | . | . | . |
| PM1548-SK1T1 | EXOC3 | p.Val576Met | missense_variant | MODERATE | 0.0614035 | 0 | 114 | 107 | . | . | . | . |
| PM1548-SK1T1 | LPCAT1 | p.Val411Ile | missense_variant | MODERATE | 0.108434 | 0 | 83 | 136 | . | . | . | . |
| PM1548-SK1T1 | IRX2 | p.Tyr110Cys | missense_variant | MODERATE | 0 | 0 | 48 | 189 | . | . | . | . |
| PM1548-SK1T1 | NSUN2 | p.Ala653Thr | missense_variant&splice_region_variant | MODERATE | 0 | 0 | 109 | 83 | . | . | . | . |
| PM1548-SK1T1 | SRD5A1 | p.Met162Ile | missense_variant | MODERATE | 0.136364 | 0 | 88 | 54 | . | . | . | . |
| PM1548-SK1T1 | SEMA5A | p.Ile679Thr | missense_variant | MODERATE | 0 | 0 | 135 | 101 | . | . | . | . |
| PM1548-SK1T1 | NIPBL | p.Glu1060Glu | synonymous_variant | LOW | 0 | 0 | 80 | 121 | . | . | . | . |
| PM1548-SK1T1 | CARD6 | p.Gln393Glu | missense_variant | MODERATE | 0 | 0 | 321 | 371 | . | . | . | . |
| PM1548-SK1T1 | CARD6 | p.Gln400His | missense_variant | MODERATE | 0 | 0 | 308 | 367 | . | . | . | . |
| PM1548-SK1T1 | CARD6 | p.Gln402* | stop_gained | HIGH | 0 | 0 | 308 | 367 | . | . | . | . |
| PM1548-SK1T1 | CARD6 | p.Gln402Arg | missense_variant | MODERATE | 0 | 0 | 282 | 337 | . | . | . | . |
| PM1548-SK1T1 | CARD6 | p.Gln402His | missense_variant | MODERATE | 0 | 0 | 282 | 337 | . | . | . | . |
| PM1548-SK1T1 | C6 | p.Ala2Val | missense_variant | MODERATE | 0.016129 | 0 | 62 | 33 | . | . | . | . |
| PM1548-SK1T1 | SNX18 | n.53839150G>A | intragenic_variant | MODIFIER | 0.118421 | 0 | 152 | 117 | . | . | . | . |
| PM1548-SK1T1 | GPX8 | p.Phe125fs | frameshift_variant | HIGH | 0.015625 | 0 | 192 | 229 | . | . | . | . |
| PM1548-SK1T1 | UTP15 | c.169-826A>G | intron_variant | MODIFIER | 0 | 0 | 139 | 55 | . | . | . | . |
| PM1548-SK1T1 | TTC37 | p.Ala1407Thr | missense_variant | MODERATE | 0.138462 | 0 | 65 | 32 | . | . | . | . |
| PM1548-SK1T1 | CHD1 | p.Asn310fs | frameshift_variant | HIGH | 0.296296 | 0 | 135 | 201 | . | . | . | true |
| PM1548-SK1T1 | PAM | p.Arg260Gln | missense_variant | MODERATE | 0.1 | 0 | 120 | 104 | . | . | . | . |
| PM1548-SK1T1 | GIN1 | p.Leu345Pro | missense_variant | MODERATE | 0.00952381 | 0 | 105 | 72 | . | . | . | . |
| PM1548-SK1T1 | APC | p.Asp802fs | frameshift_variant&stop_gained | HIGH | 0.138365 | 0 | 159 | 125 | true | true | true | true |
| PM1548-SK1T1 | MCC | p.Arg800Lys | missense_variant | MODERATE | 0 | 0 | 85 | 65 | . | . | . | . |
| PM1548-SK1T1 | FTMT | p.Ala90Val | missense_variant | MODERATE | 0.176471 | 0 | 85 | 105 | . | . | . | . |
| PM1548-SK1T1 | FBN2 | p.Leu2853Arg | missense_variant | MODERATE | 0.0254777 | 0 | 157 | 91 | . | . | . | . |
| PM1548-SK1T1 | FNIP1 | p.Leu90Met | missense_variant | MODERATE | 0.117647 | 0 | 51 | 76 | . | . | . | . |
| PM1548-SK1T1 | JADE2 | p.Phe360Leu | missense_variant | MODERATE | 0 | 0 | 192 | 343 | . | . | . | true |
| PM1548-SK1T1 | FBXL21 | n.956C>T | non_coding_exon_variant | MODIFIER | 0 | 0 | 34 | 13 | . | . | . | . |
| PM1548-SK1T1 | KDM3B | p.Arg921Trp | missense_variant | MODERATE | 0.105991 | 0 | 217 | 177 | . | . | . | . |
| PM1548-SK1T1 | PSD2 | p.Arg574His | missense_variant | MODERATE | 0 | 0.0147059 | 107 | 136 | . | . | . | . |
| PM1548-SK1T1 | PCDHA3 | p.Glu558Val | missense_variant | MODERATE | 0 | 0 | 142 | 71 | . | . | . | . |
| PM1548-SK1T1 | PCDHA5 | p.Ala56Thr | missense_variant | MODERATE | 0 | 0 | 52 | 32 | . | . | . | . |
| PM1548-SK1T1 | PCDHA5 | p.Arg64Arg | synonymous_variant | LOW | 0 | 0 | 73 | 38 | . | . | . | . |
| PM1548-SK1T1 | PCDHA5 | p.Gly69Asp | missense_variant | MODERATE | 0 | 0 | 90 | 45 | . | . | . | . |
| PM1548-SK1T1 | PCDHA8 | p.Ala57Thr | missense_variant | MODERATE | 0 | 0 | 113 | 46 | . | . | . | . |
| PM1548-SK1T1 | PCDHA13 | p.Glu558Val | missense_variant | MODERATE | 0 | 0 | 148 | 106 | . | . | . | . |
| PM1548-SK1T1 | PCDHAC2 | p.Arg543Trp | missense_variant | MODERATE | 0.052381 | 0 | 210 | 153 | . | . | . | . |
| PM1548-SK1T1 | PCDHAC2 | c.2565+37G>A | intron_variant | MODIFIER | 0.0493827 | 0 | 81 | 61 | . | . | . | . |
| PM1548-SK1T1 | PCDHGA1 | p.Ser370Gly | missense_variant | MODERATE | 0.0179856 | 0 | 278 | 386 | . | . | . | . |
| PM1548-SK1T1 | PCDHGA9 | p.Val422Ala | missense_variant | MODERATE | 0.139241 | 0 | 79 | 94 | . | . | . | . |
| PM1548-SK1T1 | PCDH1 | p.Ala203Thr | missense_variant | MODERATE | 0.0280899 | 0 | 178 | 253 | . | . | . | . |
| PM1548-SK1T1 | PCDH12 | p.Gly10Glu | missense_variant | MODERATE | 0 | 0 | 111 | 44 | . | . | . | . |
| PM1548-SK1T1 | CSF1R | p.Glu955Lys | missense_variant | MODERATE | 0.105263 | 0 | 95 | 65 | . | . | . | . |
| PM1548-SK1T1 | TIMD4 | p.Ala333Val | missense_variant | MODERATE | 0.0382775 | 0 | 209 | 229 | . | . | . | . |
| PM1548-SK1T1 | ATP10B | p.Arg115Lys | missense_variant | MODERATE | 0.00952381 | 0 | 210 | 157 | . | . | . | . |
| PM1548-SK1T1 | UBTD2 | p.Thr160Ile | missense_variant | MODERATE | 0.00775194 | 0 | 129 | 33 | . | . | . | . |
| PM1548-SK1T1 | GPRIN1 | p.Pro806fs | frameshift_variant | HIGH | 0 | 0 | 44 | 165 | . | . | . | . |
| PM1548-SK1T1 | GPRIN1 | p.Pro805Thr | missense_variant | MODERATE | 0 | 0 | 44 | 165 | . | . | . | . |
| PM1548-SK1T1 | GPRIN1 | p.Ser803fs | frameshift_variant | HIGH | 0 | 0 | 44 | 165 | . | . | . | . |
| PM1548-SK1T1 | GPRIN1 | p.Glu801Asp | missense_variant | MODERATE | 0 | 0 | 44 | 165 | . | . | . | . |
| PM1548-SK1T1 | UIMC1 | p.Phe165Cys | missense_variant | MODERATE | 0 | 0 | 240 | 180 | . | . | . | . |
| PM1548-SK1T1 | UIMC1 | p.Ile156Val | missense_variant&splice_region_variant | MODERATE | 0 | 0 | 311 | 259 | . | . | . | . |
| PM1548-SK1T1 | FGFR4 | p.Pro568fs | frameshift_variant | HIGH | 0.040201 | 0 | 199 | 174 | true | . | . | . |
| PM1548-SK1T1 | FAM193B | p.Pro761Ser | missense_variant | MODERATE | 0 | 0 | 121 | 84 | . | . | . | . |
| PM1548-SK1T1 | GRM6 | p.Ala93Thr | missense_variant | MODERATE | 0 | 0 | 80 | 91 | . | . | . | . |
| PM1548-SK1T1 | RREB1 | p.Val754Met | missense_variant | MODERATE | 0 | 0 | 38 | 159 | . | . | . | . |
| PM1548-SK1T1 | HIST1H4C | p.Val82Ala | missense_variant | MODERATE | 0.0212766 | 0 | 141 | 187 | . | . | . | . |
| PM1548-SK1T1 | HIST1H2AM | p.Leu52Met | missense_variant | MODERATE | 0.0111111 | 0 | 90 | 43 | . | . | . | . |
| PM1548-SK1T1 | TRIM27 | p.Ala24Val | missense_variant | MODERATE | 0 | 0 | 118 | 147 | true | . | . | . |
| PM1548-SK1T1 | OR2H1 | p.Ile295Leu | missense_variant | MODERATE | 0.104895 | 0 | 143 | 181 | . | . | . | . |
| PM1548-SK1T1 | HLA-F | p.Leu396fs | frameshift_variant | HIGH | 0.0300752 | 0 | 133 | 38 | . | . | . | . |
| PM1548-SK1T1 | HLA-G | p.Arg285Ser | missense_variant | MODERATE | 0 | 0 | 112 | 42 | . | . | . | . |
| PM1548-SK1T1 | TRIM26 | p.Glu241Lys | missense_variant | MODERATE | 0.09 | 0 | 100 | 201 | . | . | . | . |
| PM1548-SK1T1 | MDC1 | p.Gln1431Arg | missense_variant | MODERATE | 0 | 0 | 287 | 163 | . | . | . | . |
| PM1548-SK1T1 | MUC22 | p.Ile1250Thr | missense_variant | MODERATE | 0.00763359 | 0 | 131 | 195 | . | . | . | . |
| PM1548-SK1T1 | HLA-C | p.Arg243Trp | missense_variant | MODERATE | 0.0714286 | 0 | 14 | 40 | . | . | . | . |
| PM1548-SK1T1 | HLA-B | p.Arg243Trp | missense_variant | MODERATE | 0.0183486 | 0.00879765 | 218 | 341 | . | . | true | . |
| PM1548-SK1T1 | VWA7 | p.Thr26fs | frameshift_variant | HIGH | 0 | 0 | 266 | 239 | . | . | . | . |
| PM1548-SK1T1 | CFB | p.Asp1161fs | frameshift_variant | HIGH | 0.039801 | 0 | 201 | 133 | . | . | . | . |
| PM1548-SK1T1 | RGL2 | p.Glu372fs | frameshift_variant | HIGH | 0.0148148 | 0 | 135 | 192 | . | . | . | . |
| PM1548-SK1T1 | DNAH8 | p.Tyr1705Cys | missense_variant | MODERATE | 0 | 0 | 116 | 68 | . | . | . | . |
| PM1548-SK1T1 | KLHDC3 | p.Arg67Cys | missense_variant | MODERATE | 0.00625 | 0 | 160 | 263 | . | . | . | . |
| PM1548-SK1T1 | PKHD1 | p.Arg880His | missense_variant | MODERATE | 0.0275229 | 0 | 109 | 255 | . | . | . | . |
| PM1548-SK1T1 | DST | p.Thr1051fs | frameshift_variant | HIGH | 0.0792453 | 0 | 265 | 228 | . | . | . | . |
| PM1548-SK1T1 | KIAA1586 | p.Arg617Leu | missense_variant | MODERATE | 0 | 0 | 99 | 71 | . | . | . | . |
| PM1548-SK1T1 | ZNF451 | p.Phe193Leu | missense_variant&splice_region_variant | MODERATE | 0 | 0 | 85 | 32 | . | . | . | . |
| PM1548-SK1T1 | COL12A1 | p.Pro2509Thr | missense_variant | MODERATE | 0 | 0 | 211 | 157 | . | . | . | . |
| PM1548-SK1T1 | SENP6 | p.Ser627Gly | missense_variant | MODERATE | 0.117647 | 0 | 34 | 66 | . | . | . | . |
| PM1548-SK1T1 | ME1 | c.78+1G>A | splice_donor_variant&intron_variant | HIGH | 0.289308 | 0 | 159 | 306 | . | . | . | . |
| PM1548-SK1T1 | PRSS35 | p.Arg265* | stop_gained | HIGH | 0.0517241 | 0 | 58 | 41 | . | . | . | . |
| PM1548-SK1T1 | ANKRD6 | p.Ala390Val | missense_variant | MODERATE | 0 | 0 | 54 | 31 | . | . | . | . |
| PM1548-SK1T1 | MDN1 | p.Thr2167Lys | missense_variant | MODERATE | 0 | 0 | 83 | 72 | . | . | . | . |
| PM1548-SK1T1 | MMS22L | p.Gly37Glu | missense_variant | MODERATE | 0.115578 | 0 | 199 | 237 | . | . | . | . |
| PM1548-SK1T1 | ATG5 | p.Asp149fs | frameshift_variant | HIGH | 0.115044 | 0 | 113 | 95 | . | . | . | . |
| PM1548-SK1T1 | RFPL4B | p.Arg203Cys | missense_variant | MODERATE | 0.103448 | 0 | 58 | 78 | . | . | . | . |
| PM1548-SK1T1 | KPNA5 | p.Phe383Ser | missense_variant | MODERATE | 0 | 0 | 72 | 67 | . | . | . | . |
| PM1548-SK1T1 | FAM184A | p.Glu79Asp | missense_variant | MODERATE | 0 | 0 | 130 | 52 | . | . | . | . |
| PM1548-SK1T1 | TBC1D32 | p.Ile27Asn | missense_variant | MODERATE | 0.194444 | 0 | 144 | 41 | . | . | . | . |
| PM1548-SK1T1 | HSF2 | p.Arg196His | missense_variant | MODERATE | 0.166667 | 0 | 24 | 76 | . | . | . | . |
| PM1548-SK1T1 | KIAA0408 | p.Phe313Leu | missense_variant | MODERATE | 0 | 0 | 120 | 109 | . | . | . | . |
| PM1548-SK1T1 | SOGA3 | p.Ala214fs | frameshift_variant | HIGH | 0 | 0 | 48 | 41 | . | . | . | . |
| PM1548-SK1T1 | ENPP1 | c.240+2T>C | splice_donor_variant&intron_variant | HIGH | 0.181818 | 0 | 66 | 63 | . | . | . | . |
| PM1548-SK1T1 | CTGF | p.Ser227Phe | missense_variant | MODERATE | 0 | 0 | 110 | 36 | . | . | . | . |
| PM1548-SK1T1 | CTGF | p.Ala226Thr | missense_variant | MODERATE | 0 | 0 | 144 | 46 | . | . | . | . |
| PM1548-SK1T1 | CTGF | p.Ile195Met | missense_variant | MODERATE | 0 | 0 | 58 | 61 | . | . | . | . |
| PM1548-SK1T1 | ADGB | p.Asp890Val | missense_variant | MODERATE | 0.15 | 0 | 40 | 47 | . | . | . | . |
| PM1548-SK1T1 | SYNE1 | p.Pro1307Ser | missense_variant | MODERATE | 0.0176991 | 0 | 113 | 79 | . | . | . | . |
| PM1548-SK1T1 | ARID1B | p.Pro1578Ser | missense_variant | MODERATE | 0 | 0 | 83 | 48 | true | . | . | . |
| PM1548-SK1T1 | ARID1B | p.Ser1617Thr | missense_variant | MODERATE | 0 | 0 | 84 | 59 | true | . | . | . |
| PM1548-SK1T1 | TULP4 | p.Pro1275Thr | missense_variant | MODERATE | 0 | 0 | 194 | 171 | . | . | . | . |
| PM1548-SK1T1 | TULP4 | p.Asp1298Asn | missense_variant | MODERATE | 0.00699301 | 0.0078125 | 286 | 256 | . | . | . | . |
| PM1548-SK1T1 | TULP4 | p.Asp1298Glu | missense_variant | MODERATE | 0 | 0 | 285 | 255 | . | . | . | . |
| PM1548-SK1T1 | SMOC2 | p.Ala54Thr | missense_variant | MODERATE | 0.123077 | 0 | 65 | 36 | . | . | . | . |
| PM1548-SK1T1 | GPR146 | p.Pro248Gln | missense_variant | MODERATE | 0.121212 | 0 | 33 | 45 | . | . | . | . |
| PM1548-SK1T1 | ELFN1 | p.Arg389Cys | missense_variant | MODERATE | 0.0408163 | 0 | 49 | 43 | . | . | . | . |
| PM1548-SK1T1 | SDK1 | p.Pro1727fs | frameshift_variant | HIGH | 0.0778443 | 0 | 167 | 269 | . | . | . | . |
| PM1548-SK1T1 | PMS2 | p.Thr277Ala | missense_variant | MODERATE | 0 | 0 | 40 | 38 | true | . | . | . |
| PM1548-SK1T1 | MIOS | p.Leu213His | missense_variant | MODERATE | 0.185567 | 0 | 97 | 44 | . | . | . | . |
| PM1548-SK1T1 | ETV1 | p.Asn37Thr | missense_variant | MODERATE | 0 | 0 | 108 | 72 | true | . | . | . |
| PM1548-SK1T1 | MPP6 | p.Lys306fs | frameshift_variant | HIGH | 0.109375 | 0 | 64 | 33 | . | . | . | . |
| PM1548-SK1T1 | HERPUD2 | p.Val266Ala | missense_variant | MODERATE | 0 | 0 | 155 | 59 | . | . | . | . |
| PM1548-SK1T1 | POU6F2 | p.Glu562Ala | missense_variant | MODERATE | 0 | 0 | 273 | 226 | . | . | . | . |

| | | | | | | | | | | | | |
|---|---|---|---|---|---|---|---|---|---|---|---|---|
| PM1548-SK1T1 | MYO1G | p.Ala779Val | missense_variant | MODERATE | 0 | 0 | 162 | 358 | . | . | . | . |
| PM1548-SK1T1 | VWC2 | p.Ser3Arg | missense_variant | MODERATE | 0 | 0 | 12 | 64 | . | . | . | . |
| PM1548-SK1T1 | GRB10 | p.Ala349Thr | missense_variant | MODERATE | 0.0387097 | 0 | 310 | 375 | . | . | . | . |
| PM1548-SK1T1 | WBSCR22 | p.Pro8Gln | missense_variant | MODERATE | 0.102564 | 0 | 39 | 35 | . | . | . | . |
| PM1548-SK1T1 | CLIP2 | p.Arg771Gln | missense_variant | MODERATE | 0 | 0 | 157 | 150 | . | . | . | . |
| PM1548-SK1T1 | TMEM60 | p.Ala78fs | frameshift_variant | HIGH | 0.0777778 | 0 | 90 | 31 | . | . | . | . |
| PM1548-SK1T1 | GNAI1 | p.Cys214Phe | missense_variant | MODERATE | 0 | 0 | 205 | 196 | . | . | . | . |
| PM1548-SK1T1 | HGF | p.Val529Ala | missense_variant | MODERATE | 0.132 | 0 | 250 | 125 | . | true | . | . |
| PM1548-SK1T1 | ABCB1 | p.Tyr1165* | stop_gained | HIGH | 0.0138889 | 0 | 72 | 32 | . | . | . | . |
| PM1548-SK1T1 | ABCB1 | c.3490-1G>C | splice_acceptor_variant&intron_variant | HIGH | 0.0153846 | 0 | 65 | 32 | . | . | . | . |
| PM1548-SK1T1 | ABCB1 | p.Ser196Ala | missense_variant | MODERATE | 0 | 0 | 47 | 56 | . | . | . | . |
| PM1548-SK1T1 | MTERF | p.Ala221Thr | missense_variant | MODERATE | 0.00966184 | 0 | 207 | 212 | . | . | . | . |
| PM1548-SK1T1 | AKAP9 | p.Ile1377Thr | missense_variant | MODERATE | 0.0621118 | 0 | 161 | 76 | true | . | . | . |
| PM1548-SK1T1 | ASB4 | p.Thr229Met | missense_variant | MODERATE | 0.0454545 | 0 | 176 | 168 | . | . | . | . |
| PM1548-SK1T1 | LMTK2 | p.Asp236fs | frameshift_variant | HIGH | 0.0383142 | 0 | 261 | 300 | . | . | . | . |
| PM1548-SK1T1 | LMTK2 | p.Leu1130Ser | missense_variant | MODERATE | 0.0631579 | 0 | 190 | 72 | . | . | . | . |
| PM1548-SK1T1 | TRRAP | p.Asn1675Thr | missense_variant | MODERATE | 0 | 0 | 139 | 117 | true | . | . | . |
| PM1548-SK1T1 | TRRAP | p.Arg1978Trp | missense_variant | MODERATE | 0.0105263 | 0 | 190 | 221 | true | . | . | . |
| PM1548-SK1T1 | TRRAP | p.Pro2508Ala | missense_variant | MODERATE | 0 | 0 | 227 | 385 | true | . | . | . |
| PM1548-SK1T1 | GIGYF1 | p.Leu580Pro | missense_variant | MODERATE | 0.00877193 | 0 | 114 | 69 | . | . | . | . |
| PM1548-SK1T1 | SLC12A9 | p.Leu559fs | frameshift_variant | HIGH | 0.0740741 | 0 | 81 | 56 | . | . | . | . |
| PM1548-SK1T1 | MOGAT3 | p.Asn82Lys | missense_variant | MODERATE | 0 | 0 | 163 | 150 | . | . | . | . |
| PM1548-SK1T1 | SH2B2 | p.Ala74Ala | synonymous_variant | LOW | 0 | 0 | 38 | 70 | . | . | . | . |
| PM1548-SK1T1 | LRWD1 | p.Leu87Phe | missense_variant | MODERATE | 0 | 0 | 64 | 78 | . | . | . | . |
| PM1548-SK1T1 | FBXL13 | p.Cys375Ser | missense_variant | MODERATE | 0.105691 | 0 | 123 | 94 | . | . | . | . |
| PM1548-SK1T1 | RELN | p.Arg2363His | missense_variant | MODERATE | 0.02 | 0 | 150 | 39 | . | . | . | . |
| PM1548-SK1T1 | ATXN7L1 | p.Val629Phe | missense_variant | MODERATE | 0.101695 | 0 | 59 | 192 | . | . | . | . |
| PM1548-SK1T1 | PIK3CG | p.Thr607fs | frameshift_variant | HIGH | 0.215686 | 0 | 102 | 43 | . | true | . | . |
| PM1548-SK1T1 | CBLL1 | p.Val277Ala | missense_variant | MODERATE | 0 | 0 | 208 | 200 | . | . | . | . |
| PM1548-SK1T1 | SLC26A3 | p.Thr479Ala | missense_variant | MODERATE | 0 | 0 | 180 | 195 | . | . | true | . |
| PM1548-SK1T1 | FOXP2 | p.Gly98Val | missense_variant | MODERATE | 0 | 0 | 190 | 125 | . | . | . | . |
| PM1548-SK1T1 | PTPRZ1 | p.Pro751Leu | missense_variant | MODERATE | 0.0133333 | 0 | 75 | 64 | . | . | . | . |
| PM1548-SK1T1 | SND1 | p.Ala331Pro | missense_variant | MODERATE | 0 | 0 | 292 | 236 | true | . | . | . |
| PM1548-SK1T1 | CEP41 | p.Asp152Gly | missense_variant | MODERATE | 0 | 0 | 205 | 235 | . | . | . | . |
| PM1548-SK1T1 | TSGA13 | p.Lys151fs | frameshift_variant | HIGH | 0.137931 | 0 | 174 | 190 | . | . | . | . |
| PM1548-SK1T1 | TTC26 | p.Ala221Thr | missense_variant | MODERATE | 0.10303 | 0 | 165 | 82 | . | . | . | . |
| PM1548-SK1T1 | MKRN1 | c.186-2435G>C | intron_variant | MODIFIER | 0.0322581 | 0 | 62 | 136 | . | . | . | . |
| PM1548-SK1T1 | MGAM | p.Asn1603Asp | missense_variant | MODERATE | 0.0140845 | 0 | 71 | 85 | . | . | . | . |
| PM1548-SK1T1 | EPHB6 | p.Ala253fs | frameshift_variant | HIGH | 0 | 0 | 162 | 107 | . | true | . | . |
| PM1548-SK1T1 | ZNF775 | p.Gly65fs | frameshift_variant | HIGH | 0.1 | 0 | 50 | 50 | . | . | . | . |
| PM1548-SK1T1 | FASTK | p.Ile345Asn | missense_variant | MODERATE | 0 | 0 | 278 | 260 | . | . | . | . |
| PM1548-SK1T1 | GALNT11 | p.Leu188Pro | missense_variant | MODERATE | 0.0117647 | 0 | 170 | 157 | . | . | . | . |
| PM1548-SK1T1 | TNKS | p.His859Asn | missense_variant | MODERATE | 0.103448 | 0 | 58 | 75 | . | . | . | . |
| PM1548-SK1T1 | RP1L1 | p.Gln1044fs | frameshift_variant | HIGH | 0.176471 | 0 | 51 | 101 | . | . | . | . |
| PM1548-SK1T1 | XKR6 | p.Arg80Lys | missense_variant | MODERATE | 0 | 0 | 67 | 127 | . | . | . | . |
| PM1548-SK1T1 | PCM1 | p.Met1321Val | missense_variant | MODERATE | 0 | 0 | 55 | 138 | true | . | . | . |
| PM1548-SK1T1 | SORBS3 | p.Thr100Ala | missense_variant | MODERATE | 0 | 0 | 138 | 140 | . | . | . | . |
| PM1548-SK1T1 | LOXL2 | p.Pro40Leu | missense_variant | MODERATE | 0.123711 | 0 | 97 | 224 | . | . | . | . |
| PM1548-SK1T1 | NEFM | p.Leu5Leu | synonymous_variant | LOW | 0 | 0 | 25 | 41 | . | . | . | . |
| PM1548-SK1T1 | NRG1 | p.Gly39Glu | missense_variant | MODERATE | 0 | 0 | 55 | 55 | true | . | . | . |
| PM1548-SK1T1 | NRG1 | p.Pro44Ser | missense_variant | MODERATE | 0 | 0 | 43 | 58 | true | . | . | . |
| PM1548-SK1T1 | NRG1 | p.Ala74Pro | missense_variant | MODERATE | 0 | 0 | 68 | 35 | true | . | . | . |
| PM1548-SK1T1 | NRG1 | c.1488-469C>G | intron_variant | MODIFIER | 0 | 0 | 81 | 107 | true | . | . | . |
| PM1548-SK1T1 | ADAM2 | p.Arg3His | missense_variant | MODERATE | 0.136364 | 0 | 44 | 59 | . | . | . | . |
| PM1548-SK1T1 | DKK4 | p.Asp195Asn | missense_variant | MODERATE | 0.00961538 | 0 | 104 | 81 | . | . | . | . |
| PM1548-SK1T1 | MCM4 | p.Thr629fs | frameshift_variant | HIGH | 0.0347826 | 0 | 230 | 87 | . | . | . | . |
| PM1548-SK1T1 | PXDNL | p.Ala353Thr | missense_variant | MODERATE | 0.0384615 | 0 | 234 | 175 | . | . | . | . |
| PM1548-SK1T1 | RP1 | p.Tyr834fs | frameshift_variant | HIGH | 0.0140845 | 0 | 71 | 39 | . | . | . | . |
| PM1548-SK1T1 | YTHDF3 | p.Gln323Gln | synonymous_variant | LOW | 0 | 0 | 398 | 589 | . | . | . | . |
| PM1548-SK1T1 | C8orf46 | p.Thr78Ile | missense_variant | MODERATE | 0.111111 | 0 | 36 | 109 | . | . | . | . |
| PM1548-SK1T1 | ZFHX4 | p.Ala2905Val | missense_variant | MODERATE | 0.0462963 | 0 | 108 | 42 | . | . | . | . |
| PM1548-SK1T1 | HEY1 | p.Val187Ala | missense_variant | MODERATE | 0 | 0 | 185 | 281 | true | . | . | . |
| PM1548-SK1T1 | HEY1 | p.Thr186Ser | missense_variant | MODERATE | 0 | 0 | 156 | 272 | true | . | . | . |
| PM1548-SK1T1 | VPS13B | p.Asn2064Thr | missense_variant | MODERATE | 0 | 0 | 140 | 75 | . | . | . | . |
| PM1548-SK1T1 | TRPS1 | p.Arg297Cys | missense_variant | MODERATE | 0.0430108 | 0 | 186 | 261 | . | . | . | . |
| PM1548-SK1T1 | TRPS1 | p.Asn265Asn | synonymous_variant | LOW | 0 | 0 | 201 | 334 | . | . | . | . |
| PM1548-SK1T1 | EXT1 | p.Val66Phe | missense_variant | MODERATE | 0 | 0 | 114 | 88 | true | . | . | . |
| PM1548-SK1T1 | TBC1D31 | p.Ala208Thr | missense_variant | MODERATE | 0.0158103 | 0 | 253 | 174 | . | . | . | . |
| PM1548-SK1T1 | ATAD2 | p.Glu119fs | frameshift_variant | HIGH | 0.0614035 | 0 | 114 | 106 | . | . | . | . |
| PM1548-SK1T1 | ZNF572 | p.Ser55Pro | missense_variant | MODERATE | 0.0666667 | 0 | 90 | 41 | . | . | . | . |
| PM1548-SK1T1 | MYC | p.Gly175Ser | missense_variant | MODERATE | 0 | 0 | 176 | 270 | true | . | . | . |
| PM1548-SK1T1 | ASAP1 | p.Glu919Lys | missense_variant | MODERATE | 0 | 0 | 189 | 187 | . | . | . | . |
| PM1548-SK1T1 | WISP1 | p.Ile175Met | missense_variant | MODERATE | 0 | 0 | 159 | 119 | . | . | . | . |
| PM1548-SK1T1 | KHDRBS3 | p.Tyr316Tyr | splice_region_variant&synonymous_variant | LOW | 0.149425 | 0 | 87 | 229 | . | . | . | . |
| PM1548-SK1T1 | COL22A1 | p.Thr125Met | missense_variant | MODERATE | 0.0178571 | 0 | 56 | 59 | . | . | . | . |
| PM1548-SK1T1 | LY6D | p.Ala99Thr | missense_variant | MODERATE | 0 | 0.00956938 | 146 | 209 | . | . | . | . |
| PM1548-SK1T1 | TOP1MT | p.Pro193Leu | missense_variant | MODERATE | 0.137931 | 0.00595238 | 87 | 168 | . | . | . | . |
| PM1548-SK1T1 | ZC3H3 | p.Gly101fs | frameshift_variant | HIGH | 0 | 0 | 40 | 100 | . | . | . | . |
| PM1548-SK1T1 | GSDMD | p.Glu451Ala | missense_variant | MODERATE | 0.0472973 | 0 | 148 | 219 | . | . | . | . |
| PM1548-SK1T1 | MAPK15 | p.Pro493Leu | missense_variant | MODERATE | 0.100775 | 0 | 258 | 366 | . | . | . | . |
| PM1548-SK1T1 | SCRIB | p.Cys22Trp | missense_variant | MODERATE | 0 | 0 | 25 | 81 | . | . | . | . |
| PM1548-SK1T1 | PLEC | p.Arg3540Gln | missense_variant | MODERATE | 0.108108 | 0 | 74 | 130 | . | . | . | . |
| PM1548-SK1T1 | PLEC | p.Val648Met | missense_variant | MODERATE | 0.114286 | 0 | 35 | 164 | . | . | . | . |
| PM1548-SK1T1 | CYHR1 | n.145689659dup | intragenic_variant | MODIFIER | 0.0352941 | 0 | 85 | 56 | . | . | . | . |
| PM1548-SK1T1 | ZNF7 | p.Val136Ile | missense_variant | MODERATE | 0.127273 | 0 | 110 | 154 | . | . | . | . |
| PM1548-SK1T1 | KANK1 | p.Thr192Ser | missense_variant | MODERATE | 0 | 0 | 111 | 65 | . | . | . | . |
| PM1548-SK1T1 | DMRT3 | p.Leu385Leu | synonymous_variant | LOW | 0.00431034 | 0 | 232 | 221 | . | . | . | . |
| PM1548-SK1T1 | RLN2 | p.His8fs | frameshift_variant | HIGH | 0.0355731 | 0 | 253 | 225 | . | . | . | . |
| PM1548-SK1T1 | IFNA4 | p.Ala74Thr | missense_variant | MODERATE | 0.0185185 | 0 | 54 | 51 | . | . | . | . |
| PM1548-SK1T1 | TAF1L | p.Gly4Ser | missense_variant | MODERATE | 0.0282486 | 0 | 177 | 329 | . | . | . | true |
| PM1548-SK1T1 | BAG1 | p.Glu321Lys | missense_variant | MODERATE | 0.010101 | 0 | 198 | 192 | . | . | . | . |
| PM1548-SK1T1 | PRSS3 | p.Ser102Tyr | missense_variant | MODERATE | 0.102941 | 0 | 136 | 102 | . | . | . | . |
| PM1548-SK1T1 | CD72 | p.Ala111Thr | missense_variant | MODERATE | 0.17801 | 0 | 191 | 227 | . | . | . | . |
| PM1548-SK1T1 | RGP1 | p.Arg328Trp | missense_variant | MODERATE | 0.108108 | 0 | 185 | 168 | . | . | . | . |
| PM1548-SK1T1 | ANKRD18A | p.Gly900Ala | missense_variant | MODERATE | 0.194444 | 0 | 72 | 119 | . | . | . | . |
| PM1548-SK1T1 | ANKRD18A | p.Ala897Val | missense_variant | MODERATE | 0.194444 | 0 | 72 | 119 | . | . | . | . |
| PM1548-SK1T1 | CNTNAP3 | p.Arg841Ser | missense_variant | MODERATE | 0.108108 | 0 | 37 | 45 | . | . | . | . |
| PM1548-SK1T1 | TMC1 | p.Ala755Pro | missense_variant&splice_region_variant | MODERATE | 0 | 0 | 88 | 45 | . | . | . | . |
| PM1548-SK1T1 | RMI1 | p.Thr325Ala | missense_variant | MODERATE | 0.0381944 | 0 | 288 | 483 | . | . | . | . |
| PM1548-SK1T1 | NAA35 | p.*726* | stop_retained_variant | LOW | 0.113879 | 0 | 281 | 360 | . | . | . | . |
| PM1548-SK1T1 | SPATA31E1 | p.Arg436His | missense_variant | MODERATE | 0.130631 | 0 | 222 | 212 | . | . | . | . |
| PM1548-SK1T1 | ROR2 | p.Ala307Thr | missense_variant | MODERATE | 0.153846 | 0 | 130 | 69 | . | . | . | . |
| PM1548-SK1T1 | NOL8 | p.Arg430Lys | missense_variant | MODERATE | 0.00970874 | 0 | 103 | 112 | . | . | . | . |
| PM1548-SK1T1 | NOL8 | p.Leu427Gln | missense_variant | MODERATE | 0.00943396 | 0 | 106 | 113 | . | . | . | . |
| PM1548-SK1T1 | IPPK | p.Phe34Ser | missense_variant | MODERATE | 0.0772947 | 0 | 207 | 158 | . | . | . | . |
| PM1548-SK1T1 | SUSD3 | c.89-6671G>T | intron_variant | MODIFIER | 0 | 0 | 79 | 26 | . | . | . | . |
| PM1548-SK1T1 | WNK2 | p.His758fs | frameshift_variant | HIGH | 0.103774 | 0 | 106 | 277 | . | . | . | . |
| PM1548-SK1T1 | C9orf3 | p.Ala207Thr | missense_variant | MODERATE | 0 | 0 | 205 | 105 | . | . | . | . |
| PM1548-SK1T1 | PTCH1 | p.Thr1064Met | missense_variant | MODERATE | 0 | 0.00588235 | 198 | 170 | true | . | . | . |
| PM1548-SK1T1 | ZNF510 | p.Arg592Gly | missense_variant | MODERATE | 0 | 0 | 213 | 140 | . | . | . | . |
| PM1548-SK1T1 | CCDC180 | p.Met439Ile | missense_variant | MODERATE | 0.101266 | 0 | 79 | 95 | . | . | . | . |
| PM1548-SK1T1 | GABBR2 | p.Arg622* | stop_gained | HIGH | 0.0545455 | 0 | 110 | 80 | . | . | . | . |
| PM1548-SK1T1 | ANKS6 | p.Ala872Val | missense_variant | MODERATE | 0 | 0.00413223 | 161 | 242 | . | . | . | . |
| PM1548-SK1T1 | SLC44A1 | p.Arg437His | missense_variant | MODERATE | 0.104478 | 0 | 134 | 72 | . | . | . | . |
| PM1548-SK1T1 | SVEP1 | p.Ile1787Thr | missense_variant | MODERATE | 0.0204082 | 0 | 98 | 79 | . | . | . | . |
| PM1548-SK1T1 | TNC | p.Lys1068Asn | missense_variant | MODERATE | 0 | 0 | 87 | 37 | . | . | . | . |
| PM1548-SK1T1 | BRINP1 | p.Arg106His | missense_variant | MODERATE | 0.0778443 | 0 | 167 | 219 | . | . | . | . |
| PM1548-SK1T1 | FBXW2 | p.Arg385His | missense_variant | MODERATE | 0 | 0 | 103 | 162 | . | . | . | . |
| PM1548-SK1T1 | FBXW2 | p.Ile377Val | missense_variant | MODERATE | 0 | 0 | 107 | 161 | . | . | . | . |
| PM1548-SK1T1 | DAB2IP | p.His1025fs | frameshift_variant | HIGH | 0.0526316 | 0 | 57 | 33 | . | . | . | . |
| PM1548-SK1T1 | LRSAM1 | p.Val385Ile | missense_variant | MODERATE | 0.17757 | 0 | 107 | 115 | . | . | . | . |
| PM1548-SK1T1 | SH2D3C | p.Arg821His | missense_variant | MODERATE | 0.0973451 | 0 | 113 | 144 | . | . | . | . |
| PM1548-SK1T1 | C9orf114 | p.Val77Ile | missense_variant | MODERATE | 0.0448718 | 0 | 156 | 278 | . | . | . | . |
| PM1548-SK1T1 | C9orf50 | p.Ala252Val | missense_variant | MODERATE | 0.169903 | 0 | 206 | 411 | . | . | . | . |
| PM1548-SK1T1 | NUP214 | p.Arg887His | missense_variant | MODERATE | 0.221239 | 0 | 113 | 67 | true | . | . | . |
| PM1548-SK1T1 | ADAMTS13 | p.Arg7fs | frameshift_variant | HIGH | 0.0925926 | 0 | 216 | 249 | . | . | . | . |
| PM1548-SK1T1 | SARDH | p.Ala819Val | missense_variant | MODERATE | 0.125 | 0 | 104 | 41 | . | . | . | . |

| | | | | | | | | | | | | |
|---|---|---|---|---|---|---|---|---|---|---|---|---|
| PM1548-SK1T1 | SARDH | p.Arg280Trp | missense_variant | MODERATE | 0.109244 | 0 | 119 | 43 | . | . | . | . |
| PM1548-SK1T1 | LHX3 | p.Ala180Thr | missense_variant | MODERATE | 0.0188679 | 0 | 106 | 204 | . | . | . | . |
| PM1548-SK1T1 | MAMDC4 | p.Gln963His | missense_variant | MODERATE | 0.0888889 | 0 | 45 | 91 | . | . | . | . |
| PM1548-SK1T1 | AL807752.1 | p.Trp56fs | frameshift_variant | HIGH | 0.115385 | 0 | 26 | 54 | . | . | . | . |
| PM1548-SK1T1 | TPRN | p.Asn499Asp | missense_variant | MODERATE | 0.00411523 | 0 | 243 | 335 | . | . | . | . |
| PM1548-SK1T1 | NSMF | p.Arg366Cys | missense_variant | MODERATE | 0.260274 | 0 | 73 | 90 | . | . | . | . |
| PM1548-SK1T1 | PNPLA7 | p.Leu178Met | missense_variant | MODERATE | 0 | 0 | 192 | 199 | . | . | . | . |
| PM1548-SK1T1 | HDHD1 | p.Arg149Cys | missense_variant | MODERATE | 0 | 0 | 111 | 104 | . | . | . | . |
| PM1548-SK1T1 | EGFL6 | p.Trp5* | stop_gained | HIGH | 0.15625 | 0 | 32 | 80 | . | . | . | . |
| PM1548-SK1T1 | YY2 | p.Glu5Val | missense_variant | MODERATE | 0 | 0 | 56 | 38 | . | . | . | . |
| PM1548-SK1T1 | TMEM47 | p.Cys104Cys | synonymous_variant | LOW | 0 | 0 | 101 | 132 | . | . | . | . |
| PM1548-SK1T1 | USP27X | p.Arg155His | missense_variant | MODERATE | 0.238532 | 0 | 109 | 112 | . | . | . | . |
| PM1548-SK1T1 | KDM5C | p.Glu1003Asp | missense_variant | MODERATE | 0 | 0 | 43 | 54 | true | true | true | . |
| PM1548-SK1T1 | IQSEC2 | p.Arg612Cys | missense_variant | MODERATE | 0.163934 | 0 | 61 | 40 | . | . | . | . |
| PM1548-SK1T1 | FAM120C | p.Gln36_Gln37de | disruptive_inframe_deletion | MODERATE | 0.25 | 0 | 92 | 126 | . | . | . | . |
| PM1548-SK1T1 | MAGED2 | p.Glu53Asp | missense_variant | MODERATE | 0.193548 | 0 | 31 | 102 | . | . | . | . |
| PM1548-SK1T1 | TRO | p.Ala1132Thr | missense_variant | MODERATE | 0 | 0 | 72 | 69 | . | . | . | . |
| PM1548-SK1T1 | DLG3 | p.Leu333Ile | missense_variant | MODERATE | 0.0144928 | 0 | 69 | 34 | . | . | . | . |
| PM1548-SK1T1 | ARMCX4 | p.Tyr2123fs | frameshift_variant | HIGH | 0.149254 | 0 | 134 | 78 | . | . | . | . |
| PM1548-SK1T1 | PAK3 | p.Ala466Val | missense_variant | MODERATE | 0 | 0 | 25 | 37 | . | . | . | . |
| PM1548-SK1T1 | ZCCHC16 | p.Arg297Cys | missense_variant | MODERATE | 0.0102564 | 0 | 195 | 98 | . | . | . | . |
| PM1548-SK1T1 | AVPR2 | p.Ala60Val | missense_variant | MODERATE | 0.16129 | 0 | 31 | 59 | . | . | . | . |
