## Supplementary Information Tables for "Patient-derived xenografts and organoids model therapy response in prostate cancer": S.Table 3. List of Nexus FDA approved library.pdf

| Well | Catalog Numbe | Item Name | MW | CAS Number | Indication | Target | Brief Descriptio | Smiles | Copy 1 | Copy 2 |
|---|---|---|---|---|---|---|---|---|---|---|
| A02 | S1044 | Temsirolimus (Torisel) | 1030.29 | 162635-04-3 | Cancer | mTOR | mTOR inhibitor. | [C@H]1(C[C@ | kh_1A_10mM | kh_1B_10mM |
| A03 | S1178 | Regorafenib (BAY 73-4506) | 482.82 | 755037-03-7 | Cancer | VEGFR | Regorafenib (BA | C1=NC(=CC(=C | kh_1A_10mM | kh_1B_10mM |
| A04 | S1224 | Oxaliplatin (Eloxatin) | 397.29 | 61825-94-3 | Cancer | DNA/RNA Synth | Oxaliplatin (Elox | [Pt]1(OC(C(O1) | kh_1A_10mM | kh_1B_10mM |
| A05 | S1304 | Megestrol Acetate | 384.51 | 595-33-5 | Infection | Androgen Rece | Megestrol Aceta | C1C(C=C2[C@ | kh_1A_10mM | kh_1B_10mM |
| A06 | S1680 | Disulfiram (Antabuse) | 296.54 | 97-77-8 | Neurological Disease | | Disulfiram (Anta | CCN(C(SSC(N( | kh_1A_10mM | kh_1B_10mM |
| A07 | S2205 | OSI-420 (Desmethyl Erlotinib) | 415.87 | 183320-51-6 | Cancer | EGFR | OSI-420 (Desm | C1(=C(C=CC=C | kh_1A_10mM | kh_1B_10mM |
| A08 | S3031 | Linagliptin (BI-1356) | 472.54 | 668270-12-0 | Cancer | DPP-4 | Linagliptin is a h | C1=CC=C2C(= | kh_1A_10mM | kh_1B_10mM |
| A09 | S4244 | Serotonin HCl | 212.68 | 153-98-0 | Neurological Disease | 5-HT | Serotonin HCl is | C1(=CC=C2C(= | kh_1A_10mM | kh_1B_10mM |
| A10 | S1005 | Axitinib | 386.47 | 319460-85-0 | Cancer | VEGFR, PDGFI | Axitinib blocked | C1(=CC=CC=C | kh_1A_10mM | kh_1B_10mM |
| A11 | S1149 | Gemcitabine HCl (Gemzar) | 299.66 | 122111-03-9 | Cancer | DNA/RNA Synth | Gemcitabine HC | C1(N=C(C=CN1 | kh_1A_10mM | kh_1B_10mM |
| B02 | S1047 | Vorinostat (SAHA) | 264.3 | 149647-78-9 | Cancer | HDAC | Vorinostat also I | C1=CC=CC(=C | kh_1A_10mM | kh_1B_10mM |
| B03 | S1190 | Bicalutamide (Casodex) | 430.37 | 90357-06-5 | Endocrinology | Androgen Rece | An oral non-ster | C(C(CS(=O)(=C | kh_1A_10mM | kh_1B_10mM |
| B04 | S1225 | Etoposide (VP-16)□ | 588.56 | 33419-42-0 | Cancer | Topoisomerase | Etoposide (Etop | [C@@]12([C@ | kh_1A_10mM | kh_1B_10mM |
| B05 | S1322 | Dexamethasone | 392.46 | 50-02-2 | Inflammation | IL Receptor | Dexamethasone | C1C(C=C2[C@ | kh_1A_10mM | kh_1B_10mM |
| B06 | S1732 | Mitotane (Lysodren) | 320.04 | 53-19-0 | Cancer | | Mitotane(Lysodr | C1=C(C=CC(=C | kh_1A_10mM | kh_1B_10mM |
| B07 | S2221 | Apatinib (YN968D1) | 493.58 | 811803-05-1 | Cancer | VEGFR | Apatinib (YN968 | S(=O)(O)(C)=O | kh_1A_10mM | kh_1B_10mM |
| B08 | S3032 | Bindarit | 324.37 | 130641-38-2 | Cancer | | Bindarit exhibits | C(C(C)(OCC1=I | kh_1A_10mM | kh_1B_10mM |
| B09 | S7781 | Sunitinib | 532.56 | 341031-54-7 | Cancer | VEGFR, PDGFI | Sunitinib Malate | C1(=CC=C2C(= | kh_1A_10mM | kh_1B_10mM |
| B10 | S1014 | Bosutinib (SKI-606) | 530.45 | 380843-75-4 | Cancer | Src,Abl□ | Bosutinib (SKI-6 | N1=CC(=C(C2= | kh_1A_10mM | kh_1B_10mM |
| B11 | S1215 | Carboplatin | 371.25 | 41575-94-4 | Cancer | DNA/RNA Synth | This drug perfor | [Pt]1(OC(C2(C( | kh_1A_10mM | kh_1B_10mM |
| C02 | S1068 | Crizotinib (PF-02341066) | 450.34 | 877399-52-5 | Cancer | c-Met | PF-2341066 (Cr | C1(=CC(=C(N= | kh_1A_10mM | kh_1B_10mM |
| C03 | S1197 | Finasteride | 372.54 | 98319-26-7 | Endocrinology | 5-alpha Reducta | Inhibitor of stero | C1C(N[C@]2([C | kh_1A_10mM | kh_1B_10mM |
| C04 | S1229 | Fludarabine Phosphate | 285.23 | 21679-14-1 | Cancer | STAT, DNA/RN/ | Fludarabine (Flu | O1[C@@H]([C@ | kh_1A_10mM | kh_1B_10mM |
| C05 | S1342 | Genistein | 270.24 | 446-72-0 | Cancer | Topoisomerase | Genistein is a sc | C1(=CC2=C(C( | kh_1A_10mM | kh_1B_10mM |
| C06 | S1735 | Mesna (Uromitexan, Mesnex) | 164.18 | 19767-45-4 | Cancer | | Mesna (Uromite | SCCS(O[Na])(= | kh_1A_10mM | kh_1B_10mM |
| C07 | S2229 | Eltrombopag (SB-497115-GR) | 442.47 | 496775-61-2 | Cancer | | Eltrombopag (S | C1=C(C(=C(C= | kh_1A_10mM | kh_1B_10mM |
| C08 | S3033 | Vildagliptin (LAF-237) | 303.4 | 274901-16-5 | Metabolic Disease | DPP-4 | Vildagliptin (LAF | C1N([C@@H]( | kh_1A_10mM | kh_1B_10mM |
| C09 | S7786 | Erlotinib | 429.9 | 183319-69-9 | Cancer | EGFR | Erlotinib also kn | C1(=CC=CC(=C | kh_1A_10mM | kh_1B_10mM |
| C10 | S1025 | Gefitinib (Iressa) | 446.9 | 184475-35-2 | Cancer | EGFR | Gefitinib also kn | C1(=C(C=C2C( | kh_1A_10mM | kh_1B_10mM |
| C11 | S1311 | Pamidronate Disodium | 279.03 | 57248-88-1 | Metabolic Disease | | Pamidronate dis | C(O)(P([O-])(O) | kh_1A_10mM | kh_1B_10mM |
| D02 | S1082 | Vismodegib (GDC-0449) | 421.3 | 879085-55-9 | Cancer | Hedgehog | GDC-0449 (Visr | C1(=C(C=C(C= | kh_1A_10mM | kh_1B_10mM |
| D03 | S1202 | Dutasteride | 528.53 | 164656-23-9 | Endocrinology | 5-alpha Reducta | 5-alpha-reducta | C1C(N[C@]2([C | kh_1A_10mM | kh_1B_10mM |
| D04 | S1231 | Topotecan HCl | 457.91 | 119413-54-6 | Cancer | Topoisomerase | Topotecan Hydr | C1(C=CC2C(C= | kh_1A_10mM | kh_1B_10mM |
| D05 | S1353 | Ketoconazole | 531.43 | 65277-42-1 | Infection | P450 | Ketoconazole is | C1(=CC=C(C=C | kh_1A_10mM | kh_1B_10mM |
| D06 | S1859 | Diethylstilbestrol (Stilbestrol) | 268.35 | 56-53-1 | Cancer | | Diethylstilbestrol | C1(=CC=C(C=C | kh_1A_10mM | kh_1B_10mM |
| D07 | S2264 | Artemether (SM-224) | 298.37 | 71963-77-4 | Cancer | | Artemether is ar | [C@]12([C@]34 | kh_1A_10mM | kh_1B_10mM |
| D08 | S3035 | Daunorubicin HCl (Daunomycin HCl) | 563.98 | 23541-50-6 | Cancer | Telomerase | Daunorubicin H( | C1(=CC=C2C(= | kh_1A_10mM | kh_1B_10mM |
| D09 | S7787 | Docetaxel Trihydrate | 807.88 | 114977-28-5 | Cancer | Microtubule Ass | An microtubule ( | C1=CC=CC=C1 | kh_1A_10mM | kh_1B_10mM |
| D10 | S1028 | Lapatinib Ditosylate (Tykerb) | 925.46 | 388082-77-7 | Cancer | EGFR, HER2 | Lapatinib Ditosy | C1(=C(C=C2C( | kh_1A_10mM | kh_1B_10mM |
| D11 | S1995 | Procarbazine HCl (Matulane) | 257.76 | 366-70-1 | Cancer | DNA/RNA Synth | Procarbazine hy | C1(=CC=C(C=C | kh_1A_10mM | kh_1B_10mM |
| E02 | S1120 | Everolimus (RAD001) | 958.22 | 159351-69-6 | Cancer | mTOR | Everolimus (RAI | C(O)CO[C@H]1 | kh_1A_10mM | kh_1B_10mM |
| E03 | S1208 | Doxorubicin (Adriamycin) | 579.98 | 25316-40-9 | Cancer | Topoisomerase | Doxorubicin is a | C1(=CC=C2C(= | kh_1A_10mM | kh_1B_10mM |
| E04 | S1241 | Vincristine | 923.04 | 2068-78-2 | Cancer | Microtubule Ass | Vincristine Sulfa | C1(=CC2=C(C= | kh_1A_10mM | kh_1B_10mM |
| E05 | S1396 | Resveratrol | 228.24 | 501-36-0 | Infection | Sirtuin | Resveratrol is a | C1(=CC=C(C=C | kh_1A_10mM | kh_1B_10mM |
| E06 | S1908 | Flutamide (Eulexin) | 276.21 | 13311-84-7 | Cancer | P450 (e.g. CYP | Flutamide (Eule: | C1(=C(C=C(C= | kh_1A_10mM | kh_1B_10mM |
| E07 | S2408 | Cephalomannine | 831.9 | 71610-00-9 | Cancer | | Cephalomannine | C1=CC=CC(=C | kh_1A_10mM | kh_1B_10mM |
| E08 | S4001 | Cabozantinib malate (XL184) | 501.51 | 849217-68-1 | Cancer | c-Met | XL184 (Cabozai | C12=C(C(=CC= | kh_1A_10mM | kh_1B_10mM |
| E09 | S7810 | Afatinib (BIBW2992) Dimaleate | 485.94 | 439081-18-2 | Cancer | EGFR, HER2 | Afatinib (BIBW2 | C1(=C(C=C(C= | kh_1A_10mM | kh_1B_10mM |
| E10 | S1029 | Lenalidomide | 259.26 | 191732-72-6 | Cardiovascular Disea | TNF-alpha | Lenalidomide al: | C1=CC(=C2C(= | kh_1A_10mM | kh_1B_10mM |
| E11 | S4684 | Sildenafil | 666.7 | 171599-83-0 | Cardiovascular Disea | PDE | Sildenafil citrate | C1(=CC=C(C(= | kh_1A_10mM | kh_1B_10mM |
| F02 | S1150 | Paclitaxel (Taxol) | 853.91 | 33069-62-4 | Cancer | Microtubule Ass | Paclitaxel also k | C1=CC=CC(=C | kh_1A_10mM | kh_1B_10mM |
| F03 | S1209 | Adrucil (Fluorouracil) | 130.08 | 51-21-8 | Cancer | DNA/RNA Synth | Adrucil (Fluorou | C1(NC(C(=CN1 | kh_1A_10mM | kh_1B_10mM |
| F04 | S1250 | MDV3100 (Enzalutamide) | 464.44 | 915087-33-1 | Cancer | Androgen Rece | MDV3100 is an | C1(=C(C=CC(= | kh_1A_10mM | kh_1B_10mM |
| F05 | S1490 | Ponatinib (AP24534) | 532.56 | 943319-70-8 | Cancer | Abl | AP24534 is a nc | C1C=NN2C(C= | kh_1A_10mM | kh_1B_10mM |
| F06 | S2046 | Pioglitazone HCl | 356.44 | 111025-46-8 | Cancer | | Pioglitazone (Ac | C1(=CC=C(N=C | kh_1A_10mM | kh_1B_10mM |
| F07 | S2410 | Paeoniflorin | 480.46 | 23180-57-6 | Cancer | | Paeoniflorin is a | [C@H]1([C@H] | kh_1A_10mM | kh_1B_10mM |
| F08 | S4063 | Vitamin D3 (Cholecalciferol) | 384.64 | 67-97-0 | Cardiovascular Disease | | Vitamin D3 (Chc | C1[C@H](CCC( | kh_1A_10mM | kh_1B_10mM |
| F10 | S1035 | Pazopanib HCl | 473.98 | 635702-64-6 | Cancer | VEGFR, PDGFI | Pazopanib (GW | C1(=C(C(=CC(= | kh_1A_10mM | kh_1B_10mM |
| G02 | S1156 | Capecitabine (Xeloda) | 359.35 | 154361-50-9 | Cancer | DNA/RNA Synth | Capecitabine (X | N1=C(C(=CN(C | kh_1A_10mM | kh_1B_10mM |
| G03 | S1221 | Dacarbazine (DTIC-Dome) | 182.18 | 4342-03-4 | Cancer | DNA/RNA Synth | Dacarbazine (D | C1=NC(=C(N1)/ | kh_1A_10mM | kh_1B_10mM |
| G04 | S1289 | Carmofur | 257.26 | 61422-45-5 | Cancer | DNA/RNA Synth | Carmofur (INN) | C1(C(NC(N(C= | kh_1A_10mM | kh_1B_10mM |
| G05 | S1508 | Alprostadil (Caverject) | 354.48 | 745-65-3 | Endocrinology | | Alprostadil(Cave | [C@@H]1([C@ | kh_1A_10mM | kh_1B_10mM |
| G06 | S2057 | Cyclophosphamide monohydrate | 279.1 | 6055-19-2 | Cancer | | Cyclophospham | C1(CNP(OC1)( | kh_1A_10mM | kh_1B_10mM |
| G07 | S2521 | Epinephrine bitartrate (Adrenalinium) | 333.29 | 51-42-3 | Cancer | Adrenergic Rece | Epinephrine bita | C1(=C(C=CC(= | kh_1A_10mM | kh_1B_10mM |
| G08 | S4065 | Guanabenz acetate | 291.13 | 23256-50-0 | Endocrinology | | Guanabenz Ace | C1(=CC=C(C(= | kh_1A_10mM | kh_1B_10mM |
| G10 | S1039 | Rapamycin (Sirolimus) | 914.18 | 53123-88-9 | Immunology | mTOR | Rapamycin also | [C@@H]1([C@ | kh_1A_10mM | kh_1B_10mM |
| H02 | S1168 | Valproic acid sodium salt (Sodium valproate) | 166.19 | 1069-66-5 | Cardiovascular Disea | GABA Receptor | Valproic acid so | CCCC(C(O[Na] | kh_1A_10mM | kh_1B_10mM |
| H03 | S1223 | Epirubicin Hydrochloride | 579.98 | 56390-09-1 | Cancer | Topoisomerase | Epirubicin Hydrc | C1(=CC=C2C(= | kh_1A_10mM | kh_1B_10mM |
| H04 | S1302 | Ifosfamide | 261.09 | 3778-73-2 | Cancer | DNA/RNA Synth | Ifosfamide is a r | C1COP(N(C1)C | kh_1A_10mM | kh_1B_10mM |
| H05 | S1567 | Pomalidomide | 273.24 | 19171-19-8 | Cancer | TNF-α | Pomalidomide ir | C1=CC=C2C(= | kh_1A_10mM | kh_1B_10mM |
| H06 | S2075 | Rosiglitazone HCl | 473.5 | 155141-29-0 | Infection | PPAR | Rosiglitazone, a | C1(=CC=CC(=N | kh_1A_10mM | kh_1B_10mM |
| H07 | S3022 | Cabazitaxel (Jevtana) | 835.93 | 183133-96-2 | Neurological Disease | | Cabazitaxel (Je | [C@@]12([C@ | kh_1A_10mM | kh_1B_10mM |
| H08 | S4080 | Triamterene | 253.26 | 396-01-0 | Inflammation | Sodium Channe | Triamterene blo | C1(=NC2=C(C( | kh_1A_10mM | kh_1B_10mM |
| H10 | S1040 | Sorafenib (Nexavar) | 637.03 | 475207-59-1 | Cancer | VEGFR, PDGFI | Sorafenib Tosyl | C1(=CC=C(C=C | kh_1A_10mM | kh_1B_10mM |
