## Supplementary Information Tables for "Patient-derived xenografts and organoids model therapy response in prostate cancer": S.table 4.WES statistics.pdf

| SAMPLE | MEAN TARGET COVERAGE |
|---|---|
| BM18-1-Organoids | 88.323708 |
| BM18-1-Tumor | 117.680089 |
| BM18-2-Organoids | 109.371015 |
| BM18-2-Tumor | 117.048572 |
| BM18-3-Organoids | 107.088719 |
| BM18-3-Tumor | 116.873862 |
| LAPC9-1-Organoids | 114.966307 |
| LAPC9-2-Organoids | 76.999323 |
| LAPC9-3-Organoids | 116.123366 |
| LAPC9-m5L (Tumor) | 114.18574 |
| LAPC9-m5R (Tumor) | 108.821301 |
| PNPCa N1 | 97.89533 |
| PNPCa Org1 | 84.889893 |
| PNPCa Org2 | 118.989748 |
| PNPCa P2 | 109.014545 |
| PNPCa P3 | 101.59251 |
| PNPCa P4 | 99.498631 |
| PNPCa T1 | 106.700317 |
