## Supplementary Information Tables for "Patient-derived xenografts and organoids model therapy response in prostate cancer": Supplementary Information Tables.pdf

**SI Table 1. List of all somatic mutations identified in the PNPCa PDX and organoids, related to Fig.1**

List of somatic mutations, single nucleotide variants and insertion-deletions identified by WES.

**SI Table 2. Statistical analysis *in vivo* tumor growth in subcutaneous PDX PNPCa model, related to Fig.1**

Two-way ANOVA statistical test was performed on the tumor growth measurements among treatment groups; castrated versus castrated-testosterone in the different time points.

**SI Table 3. List of Nexus FDA approved compound library used for the organoid screens, related to Fig.4**

The following information are included; drug name, catalog number (Sellchekchem), Molecular Weight (MW), Chemical Abstracts Service identifier (CAS Number), indication of area of clinical use, molecular target description, chemical structure information (SMILES) and stock concentration.

**SI Table 4. Mean coverage WES**

Mean target coverage per sample obtained by whole exome sequencing.
